# Cross-species Study of Canine and Human Peripheral Nerve Sheath Tumors: Clinical and Molecular Perspectives

**DOI:** 10.64898/2026.07.28.740600

**Authors:** Jace P. Landry, Angela D. Bhalla, Sharon M. Landers, Rossana Lazcano, Lindsay A. Parker, Tasha M. Miller, Noelle Niemi, Heather Lyu, Heather Lillemoe, Emily Z. Keung, Christopher P. Scally, Christina L. Roland, Kelly K. Hunt, John M. Slopis, Ian E. McCutcheon, Beth Boudreau, Heather Wilson-Robles, Alexander J. Lazar, Kunal Rai, Dominique J. Wiener, Brian W. Davis, Brandan Wustefeld-Janssens, Keila E. Torres

## Abstract

Malignant peripheral nerve sheath tumors (MPNSTs) are aggressive sarcomas. An obstacle to treating MPNSTs is a lack of effective systemic therapies. Although over 70% of human MPNSTs have lost or inactivated the epigenome regulator polycomb repressive complex 2 (*PRC2*), its activity and contribution to canine PNST progression remain unclear. This study compared canine peripheral nerve sheath tumors (PNSTs) and human MPNSTs across biological and clinical features, including PRC2 activity. Immunohistochemical analysis was performed for a human tissue microarray of 54 neurofibromas and 139 MPNSTs, and 63 canine PNSTs for H3K27me3, a repressive histone mark deposited by intact PRC2, and H3K27ac, which increases globally upon H3K27me3 loss. To understand the genomic alterations present in canine PNSTs, we analyzed tumor mutation burden, copy number alteration, and transcriptomes of eight canine PNST/normal pairs. The results suggested that H3K27me3 loss and associated gain of H3K27ac epigenetically drive human and canine tumors. These findings warrant further studies to evaluate whether these epigenetic deregulations alter similar gene signatures across species.

## Introduction

Malignant peripheral nerve sheath tumors (MPNSTs) are rare sarcomas originating from peripheral nerves with a significant propensity for local recurrence and metastatic spread. In humans, one-third of MPNSTs locally recur despite aggressive resections, and about 44% progress to metastatic disease, with an overall 5-year survival rate of 52% (*1*). A major obstacle in treating these highly chemo- and radio-resistant tumors is a lack of effective systemic therapies. Due to their rarity and heterogeneous nature, the mechanisms driving MPNSTs are poorly understood (*2*). Although uncommon among human cancers, canine PNSTs account for approximately 27% of all neural tumors diagnosed in dogs (*3*). These complex tumors have an estimated annual incidence of 1309 – 11,872 per 100,000 dogs in the United States alone (*3-5*); however, humans are much less likely than canines to develop MPNSTs, with approximately 0.5 new diagnoses per 100,000 people per year (*6*). Emerging evidence shows that soft-tissue sarcomas (STS) and PNSTs in dogs closely resemble the same disease in humans. Thus, dogs present an opportunity for a preclinical model of these rare human malignancies (*7*).

While no apparent breed predilection exists for PNSTs, they most commonly affect medium to large breeds (*8*). In canines, PNSTs often present as solid tumors arising from a nerve in the subcutaneous tissues or a larger spinal nerve (*8, 9*). The mainstay of treatment for STS in canines is surgical resection with a negative margin (or R0) (*10-13*). Adjuvant radiation therapy is the most commonly utilized strategy in veterinary medicine, with an overall local cure rate of 82% (*14*). No evidence exists in veterinary medicine for the use of neoadjuvant radiation, although some institutions may use this strategy. Doxorubicin is the standard-of-care chemotherapeutic agent for canine soft-tissue sarcoma (STS); however, no significant improvement in median survival has been observed (*12*). Outcomes for canines with PNST are poor, with reported median survival between 14.7 and 43.3 months, depending on the treatment modality and anatomic location (*15, 16*). Disease-specific mortality in the first year after diagnosis in canines with PNST is reported at 31% (*15*). Low survival rates and poor responses to treatment in humans and canines alike stress the need for improved treatment strategies for this aggressive malignancy.

Sarcomas and PNSTs are complex and poorly understood malignancies known for their inherent heterogeneity. Even sensitive techniques, such as reverse transcriptase polymerase chain reaction (RT-PCR), are unable to unequivocally differentiate between fibrosarcoma and PNST in dogs (*17*). These findings suggest that the molecular aberrations of individual PNSTs vary greatly and underscore the need to thoroughly identify both the genetic mutations and epigenetic drivers of these tumors (18). Despite emerging research into epigenetic mechanisms in canine cancer, most studies have focused on lymphoid cancers (*18, 19*). However, some studies have shown promising results in epigenetics for solid tumors, such as breast cancer, bladder cancer, melanoma, and bone cancer (*20-23*). One study of mammary carcinoma in dogs demonstrated overexpression of enhancer of zeste homolog 2 (EZH2) protein (*20*). EZH2 is the catalytic subunit of polycomb repressive complex 2 (PRC2) which plays a critical role in epigenetics and chromatin structure. Similar findings were shown in human breast cancer (*20*). One study recently found that 25% of the canine nerve sheath tumors in their cohort had loss of the H3K27me3 mark with another 26% showing mosaic loss of H3K27me3 (*24*). The multi-subunit gene silencing complex, PRC2, regulates decisions involving cell fate, maintenance of stemness, and cell differentiation (*2*). PRC2 methylates lysine 27 on the tail of histone 3 (H3K27), repressing transcription of target genes. Mutations in PRC2 components and subsequent loss of function may promote MPNST development in humans and present new avenues for diagnosis and treatment. Recent studies have demonstrated that loss or inactivation of genes that regulate the epigenome (PRC2 loss) drives over 70% of human MPNSTs (*2*). Loss of PRC2 globally increases H3K27 acetylation (H3K27ac). This histone modification recruits bromodomain and extra-terminal domain (BET) proteins, which then recruit transcription activating proteins (*25*). Expression of the H3K27ac mark in canine STS remains unknown but is an exciting potential avenue to improve our molecular understanding of the disease and to open opportunities for targeted treatments such as inhibition of bromodomain-containing proteins.

Emerging evidence suggests STS and PNST in dogs closely resemble disease in humans; thus, existing techniques and concepts in human sarcoma research could translate to dogs (*7*). This study represents a multi-institutional collaboration to expedite better treatments for patients affected by these aggressive tumors. Our hypothesis was that canine PNSTs and human MPNSTs are clinically similar diseases driven by abnormal epigenetic reprogramming that supports tumor growth.

## Results

In total, 62 canine patients and 289 human patients were diagnosed with PNST or MPNST, respectively, during the study period. Table 1 summarizes the demographic and pathological characteristics of our canine cohort in comparison to our human cohort. Primary MPNSTs in our human cohort were most frequently located in the trunk (54%), followed by the extremities (31%). Similarly, most canine PNSTs were found in the trunk (44%) and then the extremities (40%). Almost all human primary MPNSTs (94%) were below the deep fascia, in contrast to 24% of canine primary PNSTs.

**Table 1.**
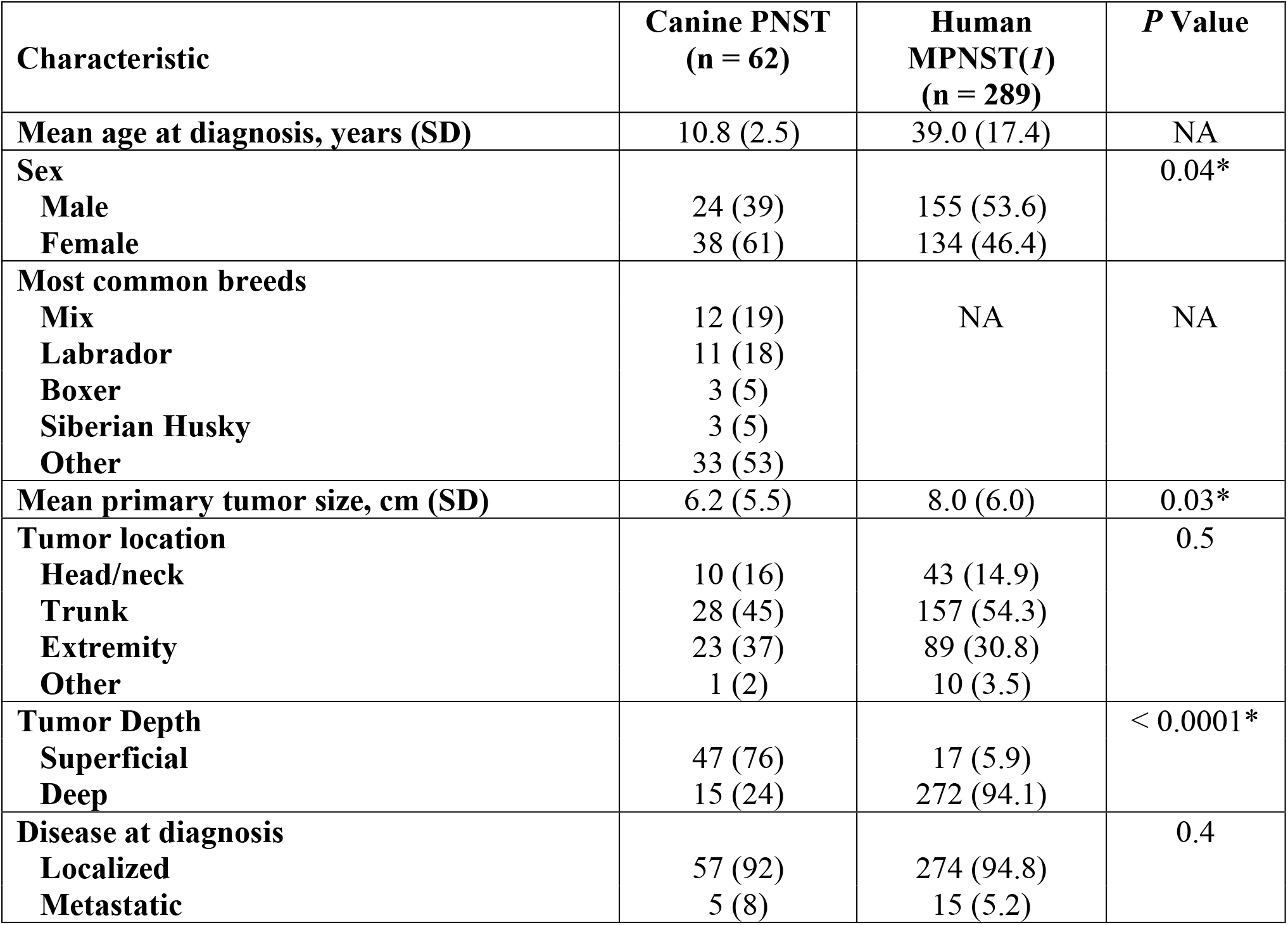
Demographics and histopathological characteristics of canines diagnosed with PNST compared to humans diagnosed with MPNST. Abbreviations: PNST, peripheral nerve sheath tumor; MPNST, malignant peripheral nerve sheath tumor; SD, standard deviation; NA, not applicable. ^*^Statistical significance defined as p < 0.05

Treatment characteristics of canine PNSTs were summarized and compared to human MPNSTs in Table 2. In patients with defined treatment characteristics, 259 (95%) human patients and 57 canines (100%) had surgical resection of their primary disease. Similar to our human cohort, surgical resection was the primary therapeutic approach for our canine patients, yet a significantly smaller percentage of canine patients received chemotherapy (18%) and radiation (12%). Local recurrence and progression to metastatic disease rates after primary tumor resection were significantly lower in our canine cohort compared to our human patients (12% vs 35.7% and 7% vs 43.8%, respectively).

**Table 2.**
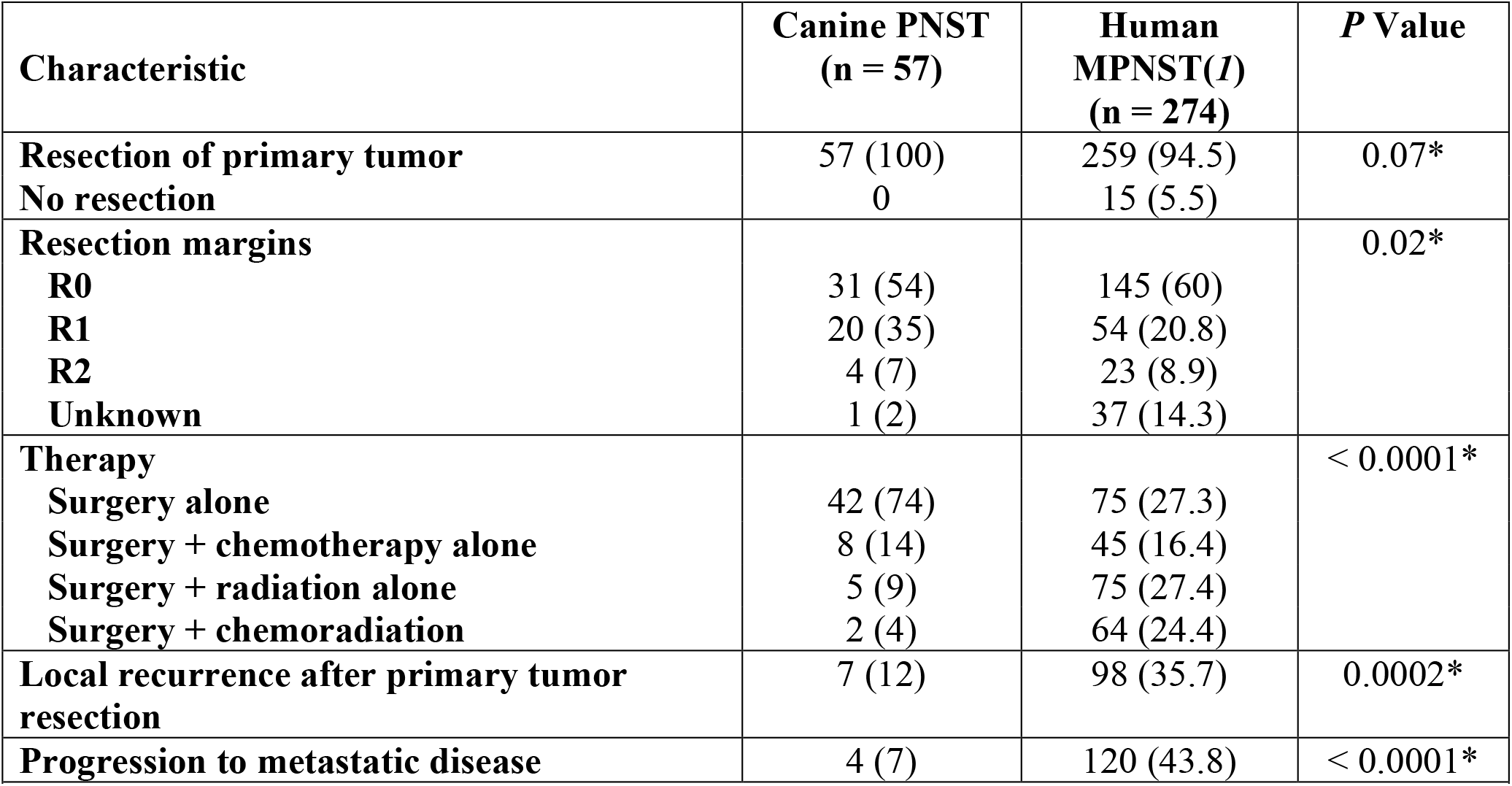
Treatment characteristics of canines diagnosed with localized MPNST with comparison to human MPNSTs with localized disease at diagnosis. Abbreviations: PNST, peripheral nerve sheath tumor; MPNST, malignant peripheral nerve sheath tumor; NA, no applicable. ^*^Statistical significance defined as p < 0.05

### Survival Analysis

Overall survival of our canine cohort is shown in Figure 1. The median survival in dogs diagnosed with PNST was 20.8 months, with a range between 14 and 44 months, depending on the treatment modality and anatomic location. The 2-year survival for canines with PNST was 40%.

**Figure 1.**
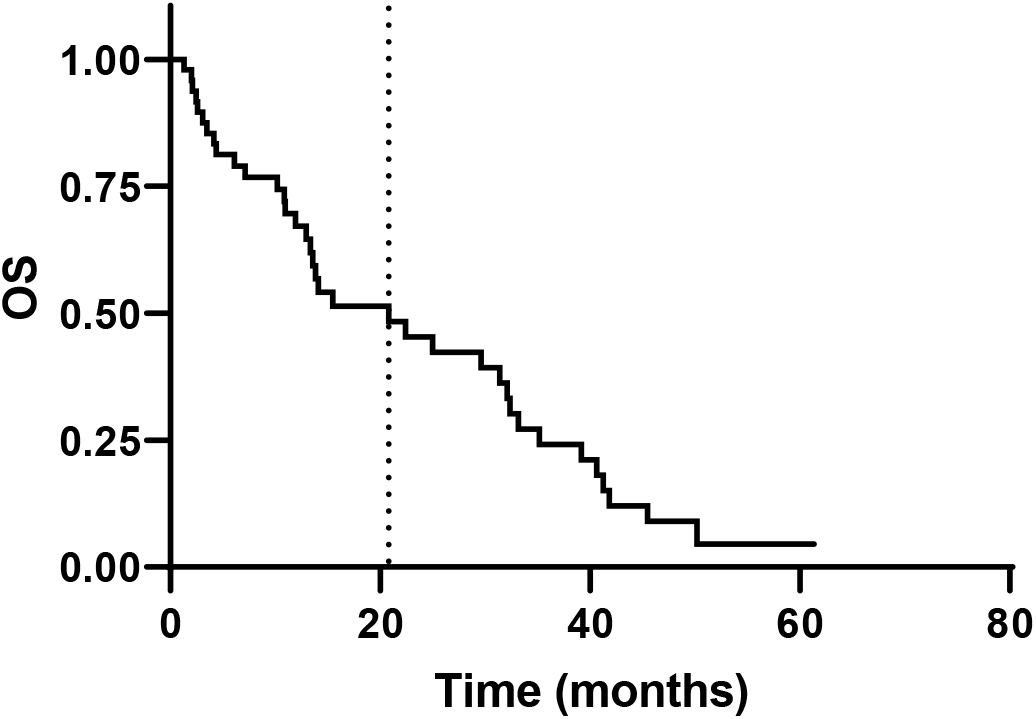
Survival analysis for canine PNST cohort. Overall survival (OS) of canines with peripheral nerve sheath tumors (PNSTs) after primary tumor resection (n = 62). Median survival (dotted line) was 20.8 months.

### Histopathologic Evaluation

Representative hematoxylin and eosin (H&E) and immunohistochemistry (IHC) images are shown in Figure 2A. All canine PNSTs had similar histological appearance on H&E compared to our human MPNST samples. We obtained MPNST FFPE and tissue microarray (TMA) samples for IHC analysis on 56 of our 62 canine patients and 139 of our 289 human patients, respectively. Our IHC analysis of calculated H scores demonstrated overall lower expression of H3K27me3 (H3K27me3 H score mean = 150.5) of canine PNSTs when compared to normal canine nerve (n = 5, Figure 2B). Conversely, high expression of H3K27ac based on calculated H scores was observed in canine PNSTs (H3K27ac mean H score = 217.5, Figure 2C). H3K27ac expression was also retained in all normal canine nerves (5/5). Human TMA analysis of neurofibroma (NF) and MPNST samples showed loss of H3K27me3 in 31% (38/121) of MPNSTs, while the mark was retained in NF samples (n = 54). In contrast, the mean percentage of H3K27ac positivity was found to be significantly higher in MPNST (45%, n = 139) compared to NF (16%, n = 54). The H scores for H3K27me2 and H3K27ac were higher for canine PNSTs than in humans (Figure 2D).

**Figure 2.**
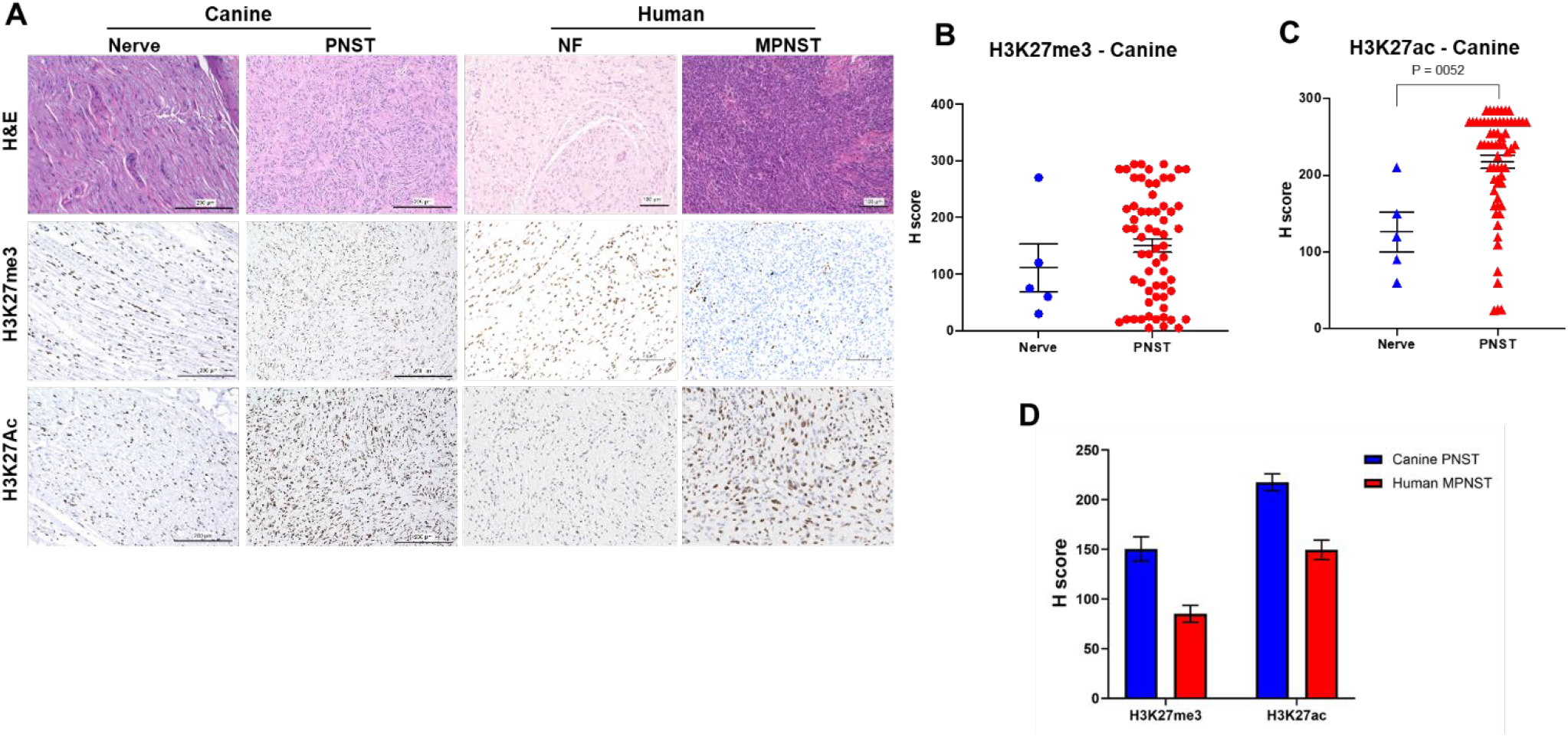
Histopathologic evaluation of canine and human PNSTs. (**A**) Representative H&E and IHC images of canine PNST (left) and human MPNST (right) samples. Quantification of canine H3K27me3 H scores (**B**) and H3K27ac (**C**) showing a wide range of H3K27me3 H score levels and increased H3K27ac H scores for canine PNSTs compared to canine nerves. (**D**) Comparison of canine and human H scores.

### Somatic Mutations and Transcriptomic Changes

The mutational burden of stop-gained, missense, and frameshift mutations was determined for genes functionally lost in human MPNST: *SUZ12, EED, CDKN2A, NF1*, and *TP53*. Only one *SUZ12* stop-gained mutation was found in sample six, the only tumor that did not express *SUZ12* in the transcriptomic data. One missense mutation in *SUZ12* was seen in each of samples five, six, and seven, but expression was not significantly altered (Figure S1A-S1B). *NF1* showed two stop gained mutations, both in sample five. Interestingly, *NF1* is expressed in sample five. Missense mutations in *NF1* are as follows: 1 in sample two, 2 in sample five, 3 in sample six, 2 in sample one, and 1 in sample four. A stop gained mutation was seen in samples five, six, and seven, but only sample six lacks *TP53* expression. Missense mutations in *TP53* were found in sample one, 2 in sample four, 3 in sample five, 8 in sample six, and 8 in sample seven. Sample eight was excluded from mutational counts due to comparative hypermutation (Table S1). Mutational signatures across all genes were compared to human using The Cancer Genome Atlas (TCGA) (*26, 27*). Comparison of mutated genes to documented human mutational signatures showed the highest similarity to adenoma, adenocarcinoma, or squamous cell carcinoma with primary site of bronchus, breast, brain, or kidney (Figure S1C-S1D). Somatic copy number was very heterogeneous as well, with all but sample eight showing aneuploidy. Most suffered a loss of chromosomes or chromosome segments, with only samples one and two having six or more chromosomes with whole-chromosome duplication. There was no consistent pattern of copy number changes between samples (Figure S2A-B).

Differential expression between all eight tumors and matched normal nerve tissue showed 5-fold more genes significantly downregulated than upregulated at FDR < 0.001 in normal tissue (Figure S3). Gene network enrichment utilizing this gene set and their associated FDR revealed three primary cell signaling pathways that encompassed the majority of dysregulated genes: the S100 family signaling pathway, the CREB neuron signaling pathway, and the FAK signaling pathway (Figure S4). Apart from gene regulatory networks, numerous individual gene misexpressions show similarities to human, as is seen in the Log_2_ fold-change (L_2_FC) compared to normal tissue (Table S2). CCN2 is extremely upregulated in tumors (L_2_FC=14.8, FDR 9.56 × 10^-17^) and encodes the connective tissue growth factor (Ctgf). *CAVIN2* (L_2_FC=3.58, FDR 8.4 × 10^-5^) produces the serum deprivation-response protein (SDPR). Upregulation of both is linked to embryonic neural crest cell mesenchymal behavior in human MPNST and CDH2 (neuronal cadherin, L_2_FC=10.8, FDR= 1.45 × 10^-7^) showing neural-crest-like expression as shown in single cell transcriptomic experiments(*28*). Interestingly, the most upregulated genes *IGFBP5* (L_2_FC=19.2 1.5 × 10-25) and *SFRP2* (L_2_FC=18, FDR=5.1×10^-10^) are secreted by Schwann cells, the cell type thought to give rise to MPNST (Table S2). *IGFBP5* is strongly associated with neuronal growth and glioblastoma progression via CREB signaling(*29*) stimulating keloid formation(*30*). SFRP2 is a SOX2-antagonist that can induce mesenchymal transition in glioma cells(*31*).

## Discussion

Comparative oncology is an evolving science with the potential to provide a better understanding of human disease mechanisms and the development of novel therapeutics (*32, 33*). Because of the increased occurrence of STS in canines, genetic and epigenetic analyses, as well as *in vivo* studies, may be simpler and more practical in canines (*34*). In humans and canines, PNSTs are aggressive and poorly understood, with little to no response to systemic therapies (*1, 15, 16*). Our multi-institutional collaboration sought to evaluate canines as a preclinical STS model for humans by comparing the clinical and histopathologic characteristics of canine and human PNSTs.

Initially, we compared canine and human PNSTs clinically on analogous aspects of diagnosis and care. PNSTs appear to follow similar etiologies in canines and humans; both arise from peripheral nerves, most frequently located on the trunk and extremities. Interestingly, the majority of canine MPNSTs developed from peripheral nerves above the deep fascia and were considered superficial in depth. In contrast, human MPNSTs in our cohort were almost all below or invaded through the deep fascia. The reason for this discrepancy in MPNST depth between our canine and human cohorts was not elucidated in this study.

Surgical resection of the primary PNST was the therapeutic approach used in all canine and nearly all (95%) human patients. The majority of human patients (68%) received adjuvant therapy in addition to surgical resection; however, only 26% of canines received additional adjuvant therapy. In addition to the standard cancer treatment options in humans (surgery, chemotherapy, radiotherapy, and palliative care), canine owners may also elect for euthanasia when their pets are diagnosed with malignant or terminal disease. The status of pets in society varies from owners who treat their pets as family members and spare no expense for care to others who abandon pets who they perceive as an inconvenience (*35, 36*). Despite this, surgery and/or adjuvant therapy is often chosen for pets, likely from positive and/or effective applications in human cancer treatments (*37, 38*). The decreased use of adjuvant therapy in canines who all received resection of their primary PNST may be multifactorial, factoring in cost of surgery compared to adjuvant therapy, discussion and guidance of their veterinarian, and perceived benefit of adjuvant therapy in the setting of PNST, which is known to be chemoradio-resistant.

Our canine cohort was significantly less likely than our human cohort to have a reported local recurrence or progression to metastatic disease by the end of the study. This is an interesting finding, considering 92% of our canine PNSTs were local at diagnosis and the poor OS observed in the canine cohort, with a median OS of 20.8 months after resection. The rate of local recurrence and metastatic progression may be higher but under-reported, depending on postoperative surveillance and care after resection. However, the poor OS may be explained by the mean age at diagnosis of 10.8 years in relatively large breed canines, making most of our canine cohort “senior citizens” (*39*). This is in contrast to the relatively young age of MPNST diagnosis in our human cohort (mean age, 39 years). The reason for the difference in relative age at diagnosis of MPNST between our canine and human cohorts is not elucidated.

Comparing canine and human PNSTs histopathologically yielded many interesting similarities. Although all PNST diagnoses were verified by respective veterinary or human pathologists with sarcoma expertise, canine PNST specimens were also reviewed by human pathologists with sarcoma expertise and were indistinguishable on H&E from human MPNSTs. We further explored PRC2 loss and increased H3K27ac in our canine PNST cohort as these are known prominent epigenetic deregulations in human MPNST. IHC analysis and H scores for H3K27me3 expression in our human cohort (TMA) showed loss in 31% of tested samples; this mark was inherently retained in all human NF samples tested (n = 55 for H3K27ac, n = 71 for H3K27me3). Our canine PNST cohort showed no complete loss of the H3K27me3 mark, but the majority (84%) had lower expression of this repressive mark when compared to normal canine nerve (n = 5). Inversely, IHC analysis for H3K27ac expression in our human cohort (TMA) resulted in significantly higher expression in MPNSTs than in NFs (45% vs 16%, respectively). Similarly, our canine cohort showed high expression of this enhancer mark when compared to normal canine nerve. These results suggest that canine PNSTs, like human MPNSTs, are epigenetically driven by decreased expression of H3K27me3 and associated gain in H3K27ac. In contrast with human MPNSTs, we found that canine PNSTs with high H3K27me3 staining also had high levels of H3K27ac staining. We attribute this discrepancy to the size differences in human core biopsies (3 mm diameter) compared to canine tissue blocks which are often 100 times larger, giving a larger area in which to observe heterogeneity. A more detailed comparison of the same areas of high H3K27me3 stained for H3K27ac could clarify this issue.

The landscape of somatic mutations was highly heterogeneous across the eight sequenced tumor-normal pairs, as well as within tumors. *SUZ12* was lost in one sample, with missense mutations in three others with small but insignificant decrease in expression. Multiple nonsense mutations were seen in *NF1* and *TP53*, though these genes were still expressed in the tumor; this could be only in the clonal compartment of these heterogeneous neoplasms. Increasing the sample size of canine PNST would contribute greatly to understanding the potential subtypes of this cancer and how it compares not only to human MPNST, but other human cancers. Due to tissue heterogeneity, a single-cell transcriptome approach would be advisable to identify which cells undergo both mutation and gene loss of expression. Though not identical in etiology to their human counterparts, canine PNST remains a viable model for human cancer accompanied by epigenetic dysregulation. Comparative transcriptomics showed primarily gene downregulation in tumors that upregulated the FAK, but downregulated the S100, and CREB signaling pathways, all of which have well-documented roles in cancer progression (Figure S3, S4) (*40-42*). Upregulation of FAK signaling, as seen in this work, is associated with an increase in H3K27me3 in human hepatocellular carcinoma, and downregulation decreases histone H3 levels at the *NOTCH2* gene (*43*). CREB-binding protein (CBP), a coactivator of CREB, operates as a histone acetyltransferase with multiple known epigenetic roles in neuronal plasticity and cancer (*42, 44*). As seen in these canine PNST, CBP is associated with an increase in H3K27ac across tissues, including human neurons (*42*). Our results show that the change in FAK and CREB signaling pathways may influence the patterns of H3K27me3 and H3K27ac observed by histopathology, similar to that seen in humans.

Given the advantageous characteristics of a spontaneous tumor-forming preclinical model like intact immunity and tumor microenvironment, we sought to evaluate canines as a preclinical sarcoma model. Our study focused on a particular sarcoma that is aggressive and poorly understood in humans and canines alike, PNST. Our findings demonstrate that canine PNSTs are similar clinically and histopathologically to human MPNSTs, and may also be similarly driven by epigenetic mechanisms. These similarities create an exciting opportunity to better understand the comparative oncology of canine and human sarcomas, especially PNSTs. Comparative oncology may open opportunities for rapid discovery of disease mechanisms, more applicable preclinical experiments, and more precise novel therapeutic trials in this rare, heterogeneous, and aggressive sarcoma subtype.

This study is the largest comparative oncology analysis of canine and human PNSTs to date. Some of the limitations in our study are inherent to its retrospective nature. First, some data are unknown or missing due to the rarity of PNSTs and the long study period required for this analysis. Second, some human and canine patients were eventually lost to follow-up, which may have affected our recurrence, progression, and survival outcomes. Despite these limitations, our study has shown important similarities between canine PNSTs and human MPNSTs, and thus such a canine model presents an opportunity to better understand disease processes and care for patients with this rare and aggressive malignancy.

Our results suggest that human and canine PNSTs are clinicopathologically similar and are epigenetically driven by H3K27me3 loss and associated gain in H3K27ac. Further studies are warranted to evaluate whether these epigenetic deregulations alter similar gene signatures in humans and canine patients. The knowledge gained from this work advances our understanding of the molecular drivers of MPNST and informs potential therapeutics to evaluate in future clinical studies.

## Materials and Methods

### Study Population

The University of Texas MD Anderson Cancer Center Institutional Review Board approved our retrospective review of patients diagnosed with MPNST evaluated between 1990 and 2015 (*1*). The retrospective canine cohort was collected under IACUC approval at Texas A&M University Veterinary Teaching Hospital between 2003 and 2018. Diagnoses were derived from veterinary pathologists and human pathologists with sarcoma expertise at the time of tumor resection or biopsy. Demographic information, histopathological characteristics, MPNST treatments, and outcomes were recorded in a clinical database. Resection margins obtained from operative notes and pathological reports were considered microscopically negative (R0), microscopically positive (R1), or macroscopically positive (R2).

### Human Tissue Microarray

An established tissue microarray (TMA) containing 162 MPNST and 97 neurofibroma tissues was used for the present study (*2, 45, 46*). The TMA was stained for H3K27ac (*47*) and H3K27me3 (*2*). Along with the TMA, a clinical database was annotated with patient, tumor, treatment and follow-up data.

### Immunohistochemical Studies

Hematoxylin and eosin (H&E) staining of canine and human PNST was performed for all samples. The human TMA was stained with anti-H3K27ac (Abcam, ab4729) (*47*) and scored by a sarcoma pathologists (RL) for staining intensity and percent positivity. Intensity was scored as negative (= 0), very low (= 1), low (= 2), high (= 3), and very high (= 4) and the percentage of positive cells for H3K27ac per field was estimated. The TMA staining with anti-H3K27me3 was performed by Cleven and colleagues (*2*). H3K27me3 stained sample intensity was scored as negative (= 0), weak (= 1), moderate (= 2), and strong (= 3) (*2*). The canine samples were stained against H3K27me3 (Cell Signaling Technology, 9733) and H3K27ac (Abcam, ab4729). Immunohistochemistry was evaluated by a sarcoma pathologist and veterinary pathologist (RL and DJW) for 61 canine PNST samples. Nuclear staining for H3K27me3 and H3K27ac was evaluated for staining intensity as negative (= 0), weak (= 1), moderate (= 2), and strong (= 3). In addition, the percentage of positive tumor cell nuclei was scored (0-100%). H-scores were calculated by multiplying the intensity by the percent positivity. For samples with mixed intensity, the percent positive for each intensity multiplied by percent positive were added together. Groups were defined as low (low intensity or % positive), high (Strong or Moderate staining), or mixed (Mixtures or areas with low staining and high staining). H-score calculated as product of H3K27me3 staining intensity and percent positivity. The samples were classified into distinct subgroups based on H-score as follows: 1–100, very low; 101–200, low; 201–300, high; 301–400, very high.

### Statistical Analysis

The Student’s *t*-test was used to evaluate differences between experimental groups. *P* values < 0.05 were considered statistically significant. Data are presented as the mean ± the standard deviation unless otherwise indicated. Log-rank methods were employed to estimate overall OS outcomes. Experiments were repeated in triplicate unless otherwise indicated. Data were analyzed using Prism 8 (GraphPad).

### Sequencing and Bioinformatics

DNA was extracted from eight tumor and matched blood samples using the phenol:choroform:isoamyl alcohol method. RNA was extracted from tumor and matched normal peripheral nerve tissue using the RNAeasy kit (Qiagen). Libraries were created using the NextFlex DNA and RNA kits respectively and sequencing was performed on an Illumina NovaSeq 6000 using a 150bp paired-end strategy at the Texas A&M Genomics and Bioinformatics core.

Genomic data was aligned to the UU_Cfam_GSD1 reference genome (*48*) using Broad Best Practices via the WAGS pipeline (*49*). Tumor-specific variants were called against the matched normal sample using Mutect2 and annotated using SNPeff to determine the impact of the somatic mutations (*50, 51*). Somatic copy number changes were estimated using iChorCNA (*52*). RNAseq data was aligned to the same reference using the HiSat algorithm and reads counted using featureCounts (*53, 54*). Differential expression of the eight tumor-normal pairs was estimated using a Fisher’s exact test after normalization using the trimmed means of m-values approach and Benjamini-Hochberg multiple testing correction using the edgeR algorithm (*55*). Expression network and pathway analyses were performed using Ingenuity Pathway Analysis (Qiagen) (*56*).

## Data availability

All sequencing data is deposited in the Sequence Read Archive at the National Library of Medicine under BioProject PRJNA1247493.

## Acknowledgments

The authors would like to acknowledge the Surgical Oncology Histology Facility at MD Anderson. The authors thank Department of Scientific Publications for editing this manuscript. The authors also thank all human and canine patients and their families for participating in this study.

## Funding

Funding for this research was provided in part by

National Institutes of Health grant T32 CA009599 (JPL)

Texas Neurofibromatosis Foundation (KET)

Sally M. Kingsbury Sarcoma Research Foundation (KET)

A Shelter for Cancer Families (KET)

Ferrin Randall Zeitlin Foundation for Sarcoma Research (KET)

Ann Jackson Family Foundation (KET)

National Institutes of Health/National Cancer Institute Cancer Center Support grant

P30CA016672 (UTMDACC Department of Veterinary Medicine Services Core Facility and Research Histology Core Laboratory)

## Author contributions

JPL, ADB, EZK, CS, CLR, KKH, JS, IM, BB, HWR, AJL, BWJ, BWD, and KET conceptualized the study. JPL, ADB, SML, LAP, and TMM designed and performed experiments, survival analysis, and clinical data analysis. RL, AJL, and DJW performed histopathologic analyses. BWD and NN performed canine genomics and transcriptomics study design and data analysis. JPL, ADB, BWD, and KET prepared the manuscript. BWD, BWJ, and KET oversaw the study and manuscript preparation.

## Data and materials availability

All data are available in the main text or the supplementary materials. All canine sequencing data is deposited in the Sequence Read Archive at the National Library of Medicine under BioProject PRJNA1247493.

## Supplementary Materials

The Supplementary Materials are included as a separate pdf.

Figs. S1 to S4

Table S1

Table S2

## References

1. K. L. Watson et al., Patterns of recurrence and survival in sporadic, neurofibromatosis Type 1-associated, and radiation-associated malignant peripheral nerve sheath tumors. J Neurosurg 126, 319–329 (2017).

2. A. H. Cleven et al., Loss of H3K27 tri-methylation is a diagnostic marker for malignant peripheral nerve sheath tumors and an indicator for an inferior survival. Mod Pathol 29, 582–590 (2016).

3. D. L. Gustafson, D. L. Duval, D. P. Regan, D. H. Thamm, Canine sarcomas as a surrogate for the human disease. Pharmacol Therapeut 188, 80–96 (2018).

4. K. Gruntzig et al., The Swiss Canine Cancer Registry: a retrospective study on the occurrence of tumours in dogs in Switzerland from 1955 to 2008. J Comp Pathol 152, 161–171 (2015).

5. ASPCA. (2017).

6. B. Seguin, Canine Soft Tissue Sarcomas: Can Being a Dog’s Best Friend Help a Child? Front Oncol 7, 285 (2017).

7. M. Milovancev et al., Comparative pathology of canine soft tissue sarcomas: possible models of human non-rhabdomyosarcoma soft tissue sarcomas. J Comp Pathol 152, 22–27 (2015).

8. J. P. Bray, Soft tissue sarcoma in the dog-part 1: a current review. J Small Anim Pract 57, 510–519 (2016).

9. L. Gaitero, S. Anor, D. Fondevila, M. Pumarola, Canine cutaneous spindle cell tumours with features of peripheral nerve sheath tumours: a histopathological and immunohistochemical study. J Comp Pathol 139, 16–23 (2008).

10. J. P. Bray, G. A. Polton, K. D. McSporran, J. Bridges, T. M. Whitbread, Canine soft tissue sarcoma managed in first opinion practice: outcome in 350 cases. Vet Surg 43, 774–782 (2014).

11. K. D. McSporran, Histologic grade predicts recurrence for marginally excised canine subcutaneous soft tissue sarcomas. Vet Pathol 46, 928–933 (2009).

12. K. A. Selting et al., Outcome of dogs with high-grade soft tissue sarcomas treated with and without adjuvant doxorubicin chemotherapy: 39 cases (1996-2004). J Am Vet Med Assoc 227, 1442–1448 (2005).

13. N. J. Bacon, W. S. Dernell, N. Ehrhart, B. E. Powers, S. J. Withrow, Evaluation of primary re-excision after recent inadequate resection of soft tissue sarcomas in dogs: 41 cases (1999-2004). J Am Vet Med Assoc 230, 548–554 (2007).

14. J. L. Demetriou, M. J. Brearley, F. Constantino-Casas, C. Addington, J. Dobson, Intentional marginal excision of canine limb soft tissue sarcomas followed by radiotherapy. J Small Anim Pract 53, 174–181 (2012).

15. L. van Stee, S. Boston, E. Teske, B. Meij, Compartmental resection of peripheral nerve tumours with limb preservation in 16 dogs (1995-2011). Vet J 226, 40–45 (2017).

16. K. E. Swift et al., Clinical and imaging findings, treatments, and outcomes in 27 dogs with imaging diagnosed trigeminal nerve sheath tumors: A multi-center study. Vet Radiol Ultrasound 58, 679–689 (2017).

17. S. Teixeira, I. Amorim, A. Rema, F. Faria, F. Gartner, Molecular Heterogeneity of Canine Cutaneous Peripheral Nerve Sheath Tumors: A Drawback in the Diagnosis Refinement. In Vivo 30, 819–827 (2016).

18. S. Da Ros et al., Validation of epigenetic mechanisms regulating gene expression in canine B-cell lymphoma: An in vitro and in vivo approach. PLoS One 13, e0208709 (2018).

19. S. Ferraresso et al., DNA methylation profiling reveals common signatures of tumorigenesis and defines epigenetic prognostic subtypes of canine Diffuse Large B-cell Lymphoma. Sci Rep 7, 11591 (2017).

20. H. J. Choi et al., Significance of EZH2 expression in canine mammary tumors. BMC Vet Res 12, 164 (2016).

21. S. Noguchi et al., DNA methylation contributes toward silencing of antioncogenic microRNA-203 in human and canine melanoma cells. Melanoma Res 25, 390–398 (2015).

22. C. M. Fulkerson, D. Dhawan, T. L. Ratliff, N. M. Hahn, D. W. Knapp, Naturally Occurring Canine Invasive Urinary Bladder Cancer: A Complementary Animal Model to Improve the Success Rate in Human Clinical Trials of New Cancer Drugs. Int J Genomics 2017, 6589529 (2017).

23. M. C. Scott et al., Aberrant Retinoblastoma (RB)-E2F Transcriptional Regulation Defines Molecular Phenotypes of Osteosarcoma. J Biol Chem 290, 28070–28083 (2015).

24. K. Tekavec et al., Loss of H3K27me3 expression in canine nerve sheath tumors. Front Vet Sci 9, 921720 (2022).

25. T. De Raedt et al., PRC2 loss amplifies Ras-driven transcription and confers sensitivity to BRD4-based therapies. Nature 514, 247–251 (2014).

26. R. L. Grossman et al., Toward a Shared Vision for Cancer Genomic Data. N Engl J Med 375, 1109–1112 (2016).

27. in Genomic Data Commons Data Portal. (National Cancer Institute, https://portal.gdc.cancer.gov/).

28. L. M. N. Wu et al., Single-cell multiomics identifies clinically relevant mesenchymal stem-like cells and key regulators for MPNST malignancy. Science Advances 8, eabo5442 (2022).

29. W. Lin et al., IGFBP5 is an ROR1 ligand promoting glioblastoma invasion via ROR1/HER2-CREB signaling axis. Nature Communications 14, 1578 (2023).

30. K. Wei et al., Schwann cells secrete IGFBP5 to facilitate the growth of keloids. Life Sciences 369, 123534 (2025).

31. M. Guo et al., SFRP2 induces a mesenchymal subtype transition by suppression of SOX2 in glioblastoma. Oncogene 40, 5066–5080 (2021).

32. I. Gordon, M. Paoloni, C. Mazcko, C. Khanna, The Comparative Oncology Trials Consortium: using spontaneously occurring cancers in dogs to inform the cancer drug development pathway. PLoS medicine 6, e1000161 (2009).

33. I. K. Gordon, C. Khanna, Modeling opportunities in comparative oncology for drug development. ILAR journal 51, 214–220 (2010).

34. A. Y. Angstadt, V. Thayanithy, S. Subramanian, J. F. Modiano, M. Breen, A genome-wide approach to comparative oncology: high-resolution oligonucleotide aCGH of canine and human osteosarcoma pinpoints shared microaberrations. Cancer Genet 205, 572–587 (2012).

35. F. Pirrone, L. Pierantoni, S. M. Mazzola, D. Vigo, M. Albertini, Owner and animal factors predict the incidence of, and owner reaction toward, problematic behaviors in companion dogs. J Vet Behav 10, 295–301 (2015).

36. P. Arkow, Application of ethics to animal welfare. Appl Anim Behav Sci 59, 193–200 (1998).

37. S. J. Withrow, D. M. Vail, Withrow & MacEwen’s small animal clinical oncology. (Saunders Elsevier, St. Louis, Mo., ed. 4th, 2007), pp. xvii, 846 p.

38. J. M. Dobson, B. D. X. Lascelles, British Small Animal Veterinary Association., BSAVA manual of canine and feline oncology. BSAVA manual series (British Small Animal Veterinary Association, Quedgeley, Gloucester, ed. 3rd, 2011), pp. viii, 364 p.

39. G. J. Patronek, D. J. Waters, L. T. Glickman, Comparative longevity of pet dogs and humans: implications for gerontology research. J Gerontol A Biol Sci Med Sci 52, B171–178 (1997).

40. C. Allgöwer et al., Friend or Foe: S100 Proteins in Cancer. Cancers 12, 2037 (2020).

41. S. Pomella et al., New Insights on the Nuclear Functions and Targeting of FAK in Cancer. International Journal of Molecular Sciences 23, 1998 (2022).

42. L. Sapio et al., Targeting CREB in Cancer Therapy: A Key Candidate or One of Many? An Update. Cancers 12, 3166 (2020).

43. D. Gnani et al., Focal adhesion kinase depletion reduces human hepatocellular carcinoma growth by repressing enhancer of zeste homolog 2. Cell Death & Differentiation 24, 889–902 (2017).

44. C. G. Vecsey et al., Histone Deacetylase Inhibitors Enhance Memory and Synaptic Plasticity via CREB: CBP-Dependent Transcriptional Activation. The Journal of Neuroscience 27, 6128–6140 (2007).

45. K. E. Torres et al., Activated MET is a molecular prognosticator and potential therapeutic target for malignant peripheral nerve sheath tumors. Clinical cancer research : an official journal of the American Association for Cancer Research 17, 3943–3955 (2011).

46. C. Zou et al., Clinical, pathological, and molecular variables predictive of malignant peripheral nerve sheath tumor outcome. Annals of surgery 249, 1014–1022 (2009).

47. V. Kochat et al., Enhancer reprogramming in PRC2-deficient malignant peripheral nerve sheath tumors induces a targetable de-differentiated state. Acta Neuropathol 142, 565–590 (2021).

48. C. Wang et al., A novel canine reference genome resolves genomic architecture and uncovers transcript complexity. Communications Biology 4, 185 (2021).

49. J. N. Cullen, S. G. Friedenberg, Whole Animal Genome Sequencing: user-friendly, rapid, containerized pipelines for processing, variant discovery, and annotation of short-read whole genome sequencing data. G3 Genes|Genomes|Genetics 13, (2023).

50. Z. Chen et al., Systematic comparison of somatic variant calling performance among different sequencing depth and mutation frequency. Scientific Reports 10, 3501 (2020).

51. P. Cingolani et al., A program for annotating and predicting the effects of single nucleotide polymorphisms, SnpEff. Fly 6, 80–92 (2012).

52. V. A. Adalsteinsson et al., Scalable whole-exome sequencing of cell-free DNA reveals high concordance with metastatic tumors. Nature Communications 8, 1324 (2017).

53. D. Kim, J. M. Paggi, C. Park, C. Bennett, S. L. Salzberg, Graph-based genome alignment and genotyping with HISAT2 and HISAT-genotype. Nature Biotechnology 37, 907–915 (2019).

54. Y. Liao, G. K. Smyth, W. Shi, featureCounts: an efficient general purpose program for assigning sequence reads to genomic features. Bioinformatics 30, 923–930 (2013).

55. Y. Chen, A. T. L. Lun, G. K. Smyth, in Statistical Analysis of Next Generation Sequencing Data, S. Datta, D. Nettleton, Eds. (Springer International Publishing, Cham, 2014), pp. 51–74.

56. A. Krämer, J. Green, J. Pollard, Jr, S. Tugendreich, Causal analysis approaches in Ingenuity Pathway Analysis. Bioinformatics 30, 523–530 (2013).

