## Supplementary material for "Cross-species Study of Canine and Human Peripheral Nerve Sheath Tumors: Clinical and Molecular Perspectives": Figs S1-S4, Table S1

Jace P. Landry *et al.*

**This PDF file includes:**

Figs. S1 to S4
Table S1

**Fig. S1.**

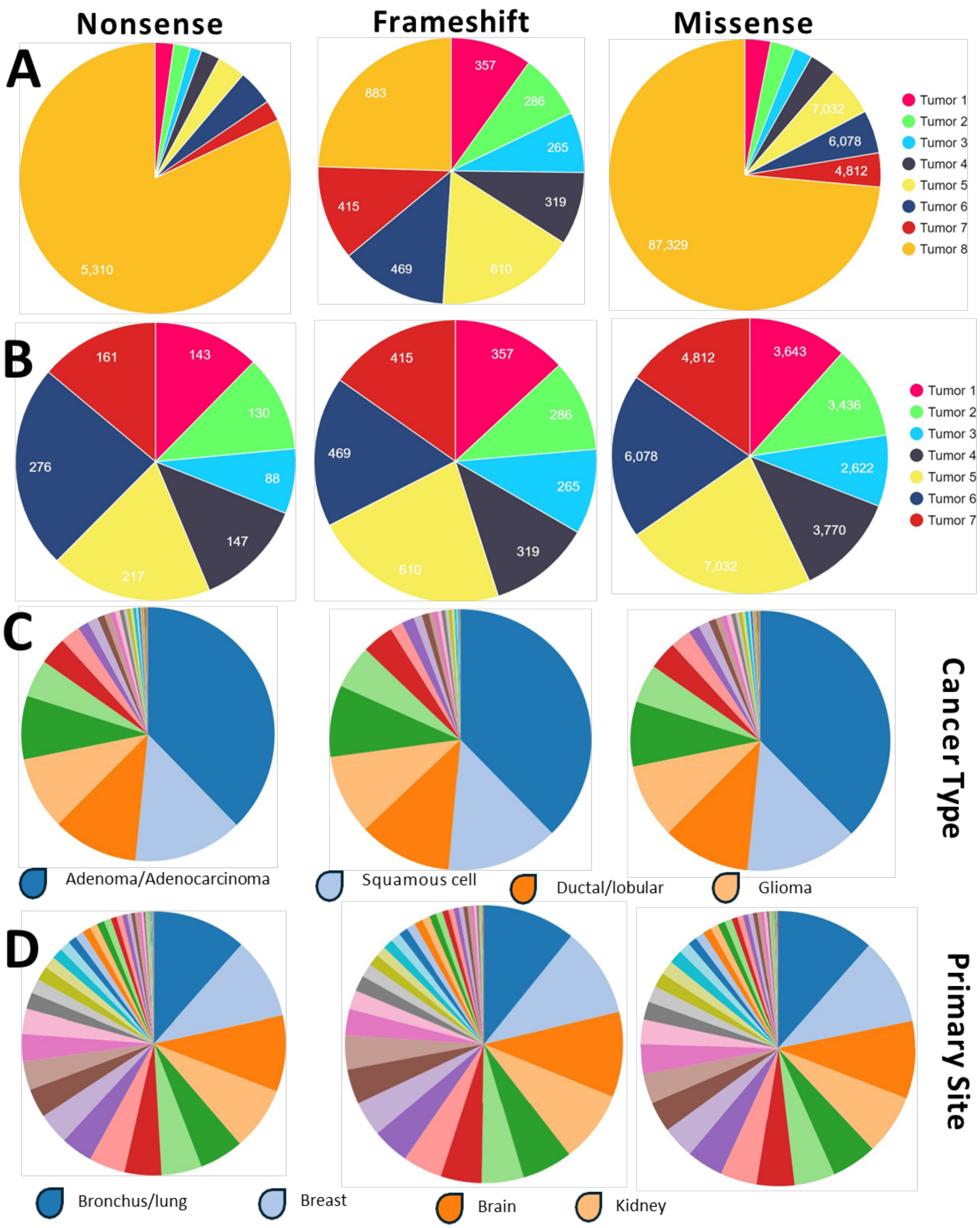


**Fig. S1. Total number of nonsense, frameshift, and missense somatic mutations** across

(A - B) all samples and with sample 8 removed due to relative hypermutation compared to other samples.

(C - D) Missense and higher severity mutations that affected genes in half of the tumors were compared to genes mutated in the Cancer Genome Atlas based on (C) disease type and (D) tissue of origin. Top four of (C) and (D) are shown.

**Fig. S2a.**

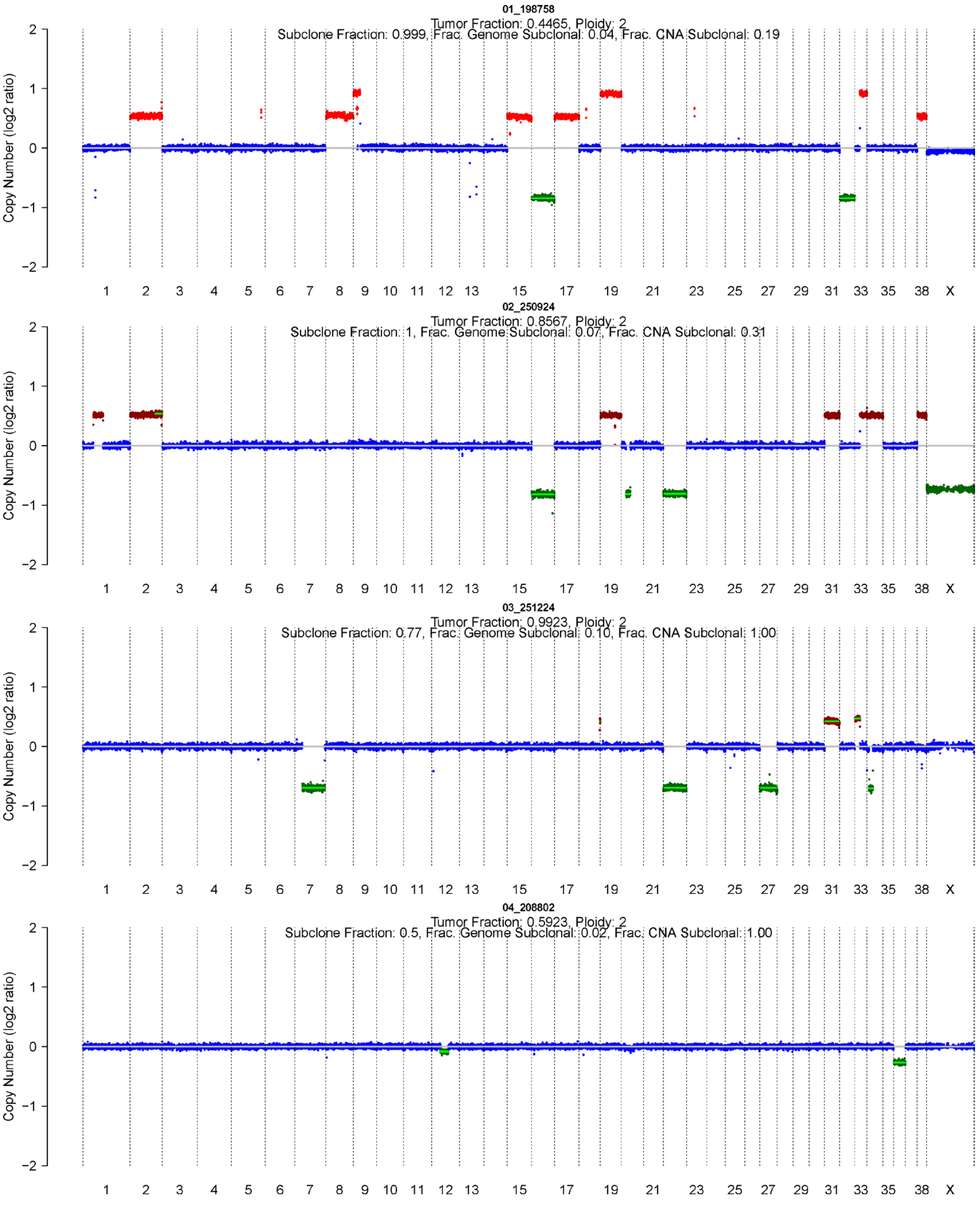


**Fig. S2a. Somatic copy number estimation for samples 1-4.** Departure from strict diploidy is indicated by an increase or decrease in $log_2$ copy number on the y-axis. Chromosome number on the x-axis. No consistent patterns of copy number change were observed across samples.

Sample 1: net gain of chromosome 2, 8, 15, 17, 19, and 38, with partial gain of the p arm of 9 and the q of 34. Net loss of 16 and 32.

Sample 2: net gain of 2, 19, 32, 34, and 38 with a partial gain on 1 and 33.

Sample 3: net gain of 31 and part of 33. Net loss of 22 and 27, with a partial loss of 7 and 34.

Sample 4: net loss of 36 and partial loss of 12.

**Fig. S2b.**

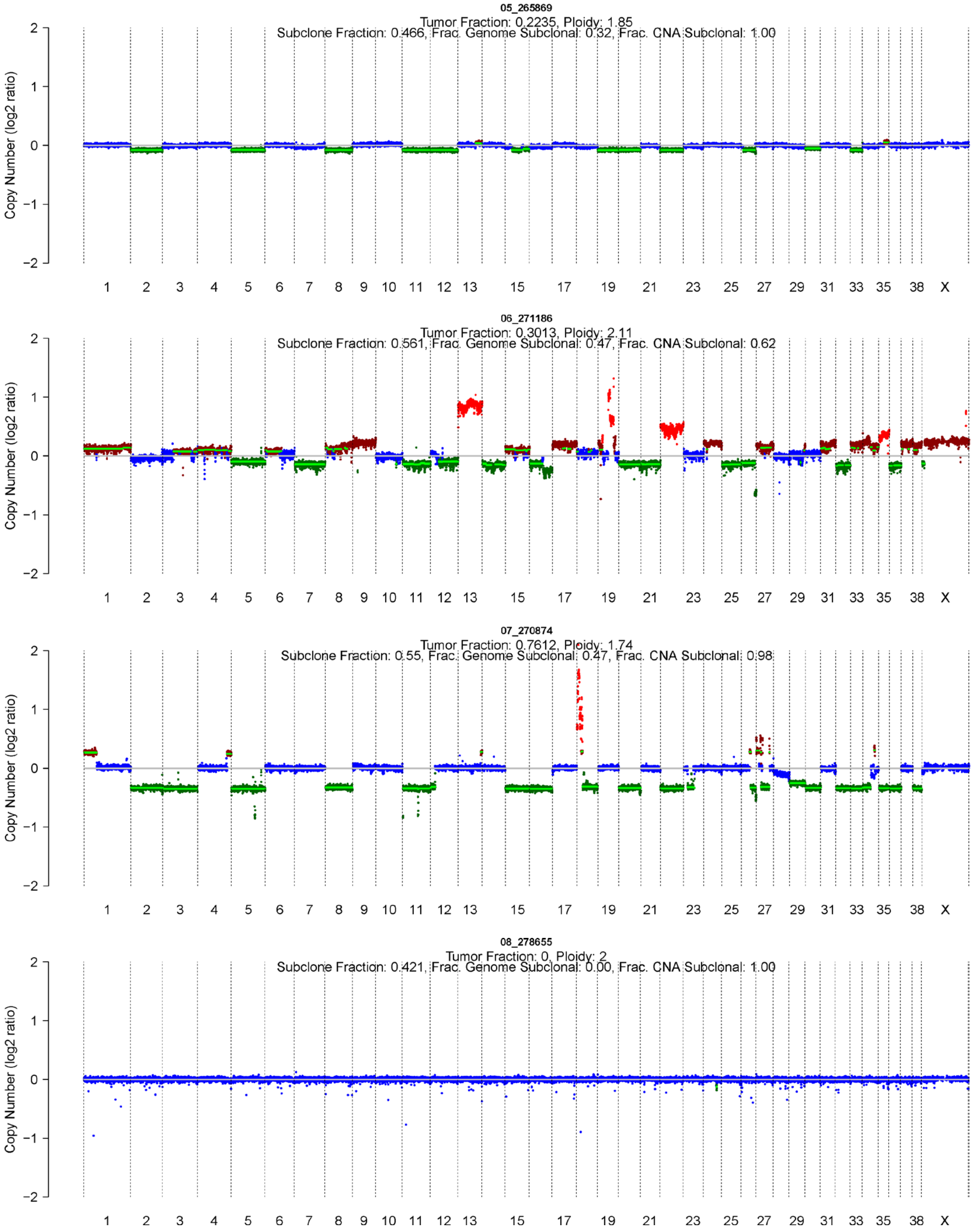


**Fig. S2b. Somatic copy number estimates for samples 5-8.**

Sample 5: net loss of chromosome 2, 5, 8, 11, 12, 15, 19, 20, 22, 27. 30, and 34. The magnitude is small, so this is likely a subclonal population.

Sample 6: Loss of 2, 6, 7, 11, 20, 21, 25, 26, 32, and 36, and the q arm of 12 and 16. Net gain of chromosome 13,22, and 34, with multiple duplication events on 19.

Sample 7: Gain of p arm segment on chromosome 1, and the q of 4. Loss of 2, 3, 5, 8, 11, 15, 16, 18, 20, 22, 29, 30, 32, 33, 35, and 36. Extreme duplication event on the p arm of 18 and complex copy loss and gain on 27.

Sample 8: no detectable copy number alterations, though this tumor has an order of magnitude more mutations (Figure S1)

**Fig. S3.**

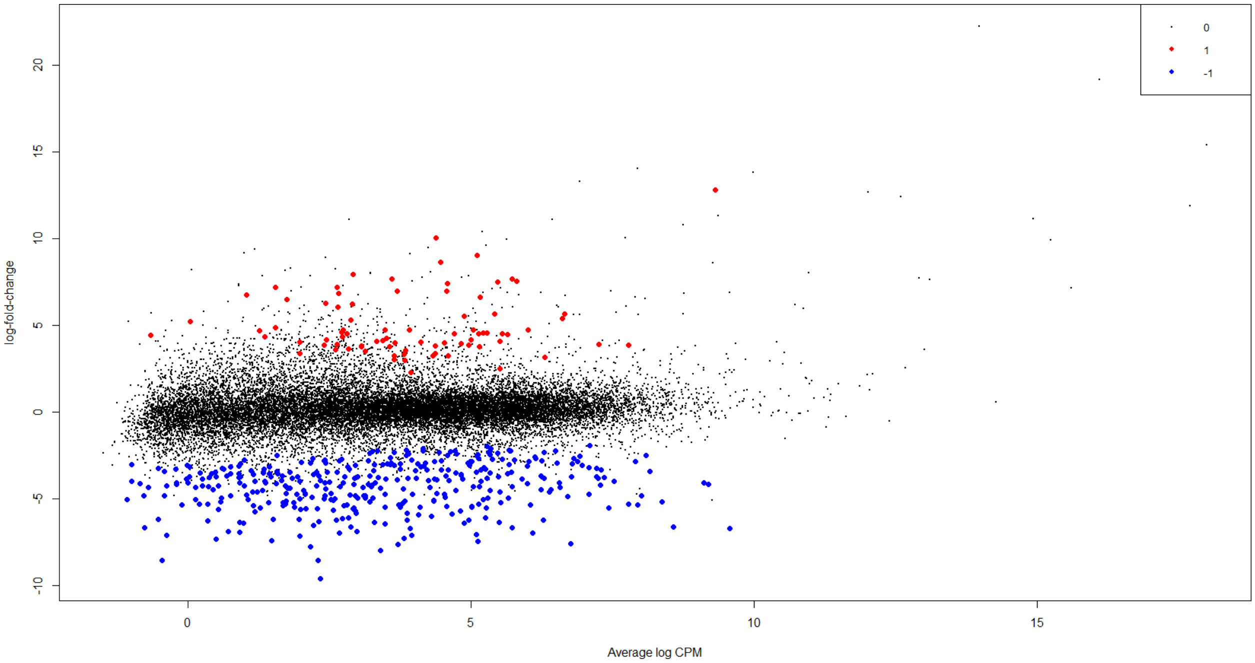


**Fig. S3. Smear plot of differentially expressed genes** (FDR < 0.001) between MPNST and normal nerve tissue determined by Fisher's exact test and Benjamini-Hochberg multiple testing correction. Red indicates an upregulated gene in tumor and blue a downregulated gene. X axis is average log of the counts per million reads and the Y axis is the $\log_2$ fold-change between tumor and normal.

**Fig. S4.**

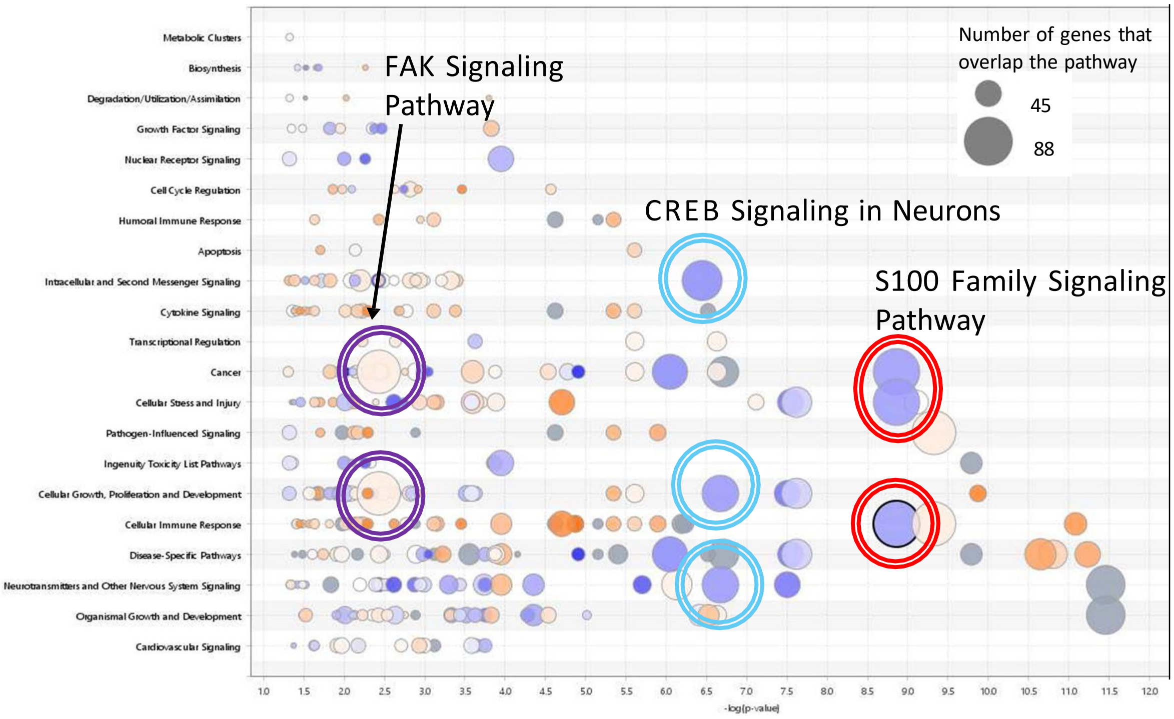


**Fig. S4. Pathway enrichment analysis.** Three cell signaling pathways were identified as the most enriched using a network-pathway analysis in Ingenuity (Qiagen). Size of the circle indicates the number of genes dysregulated. Purple indicates downregulation and orange indicates upregulation in the tumor. Three pathways contained the most dysregulation. FAK signaling circled in purple with many genes slightly upregulated at a moderate $\log_{10}p$, CREB neuronal signaling circled in light blue with many genes downregulated at $-\log_{10}p = 6.5\text{-}6.75$, and S100 signaling circled in red with the most genes downregulated at $-\log_{10}p = 8.8$.

### Table S1. Total counts of somatic mutations by impact in 8 sequenced tumors

| Mutation Type | Total Mutations | S1 | S2 | S3 | S4 | S5 | S6 | S7 | S8 | SNP | INS | DEL |
|---|---|---|---|---|---|---|---|---|---|---|---|---|
| **Stop gained** | 6103 | 143 | 130 | 88 | 147 | 217 | 276 | 161 | 5310 | 6016 | 105 | 16 |
| **Stop lost** | 1346 | 61 | 70 | 39 | 61 | 97 | 90 | 102 | 1024 | 1263 | 44 | 64 |
| **Start lost** | 513 | 24 | 18 | 12 | 16 | 39 | 18 | 22 | 411 | 500 | 4 | 11 |
| **Gene fusion** | 15 | 2 | 1 | 0 | 2 | 3 | 3 | 2 | 3 | 0 | 0 | 15 |
| **Inframe insertion** | 327 | 29 | 32 | 32 | 20 | 40 | 42 | 39 | 174 | 37 | 327 | 60 |
| **Inframe Deletion** | 883 | 138 | 120 | 115 | 118 | 230 | 155 | 148 | 198 | 29 | 43 | 883 |
| **Frameshift** | 2472 | 357 | 286 | 265 | 319 | 610 | 469 | 415 | 883 | 199 | 1380 | 1606 |
| **Missense** | 107,071 | 3643 | 3436 | 2622 | 3770 | 7032 | 6078 | 4812 | 87329 | 107071 | 115 | 98 |
