## Supplementary material for "Cross-species Study of Canine and Human Peripheral Nerve Sheath Tumors: Clinical and Molecular Perspectives": Table S2

Table S2. Differentially expressed genes between canine PNST and normal nerve tissue

| | logFC | logCPM | PValue | FDR |
|---|---|---|---|---|
| IGFBP5 | 19.19967788 | 10.71006354 | 1.88E-28 | 1.51E-25 |
| COL6A1 | 19.17333623 | 10.63615983 | 4.90E-34 | 1.06E-30 |
| LOC119878321 | 18.90227873 | 13.73367164 | 0.000985351 | 0.004043075 |
| LOC119879332 | 18.49699359 | 15.86758305 | 0.000125184 | 0.000691768 |
| LOC119879331 | 18.46015282 | 9.744148047 | 0.000208642 | 0.001068251 |
| SFRP2 | 18.0295553 | 9.319875633 | 1.56E-11 | 5.09E-10 |
| DCN | 17.81797486 | 9.509291344 | 1.76E-19 | 2.66E-17 |
| COL6A3 | 17.629137 | 10.55889169 | 4.25E-33 | 7.68E-30 |
| COL12A1 | 17.29859263 | 9.019824818 | 2.24E-26 | 1.13E-23 |
| LOC119879273 | 17.19027836 | 17.45291676 | 1.37E-07 | 1.79E-06 |
| LAMA2 | 17.18590616 | 8.630283561 | 1.67E-28 | 1.39E-25 |
| MMP2 | 17.07431502 | 9.979552291 | 1.49E-32 | 2.49E-29 |
| COL5A3 | 17.0443998 | 8.728353452 | 1.15E-21 | 2.60E-19 |
| COL3A1 | 16.73223912 | 11.7522638 | 1.27E-23 | 3.71E-21 |
| C1S | 16.55152267 | 8.113840522 | 4.24E-25 | 1.64E-22 |
| BGN | 16.54240531 | 8.031024378 | 1.30E-24 | 4.56E-22 |
| GLI2 | 16.42425116 | 7.770965551 | 9.91E-20 | 1.60E-17 |
| EFEMP1 | 16.42308154 | 7.890613973 | 3.76E-23 | 1.05E-20 |
| CLDN1 | 16.15930156 | 7.976687337 | 1.63E-14 | 9.66E-13 |
| HHIP | 16.08431771 | 7.38349108 | 8.29E-09 | 1.46E-07 |
| FNDC1 | 16.04042434 | 7.891376733 | 3.34E-17 | 3.18E-15 |
| LUM | 15.98549902 | 7.627848408 | 6.08E-10 | 1.40E-08 |
| CFH | 15.92390552 | 7.845087082 | 5.44E-18 | 6.11E-16 |
| HTRA3 | 15.90505167 | 7.240007311 | 3.11E-16 | 2.50E-14 |
| ADAMTS5 | 15.82595369 | 8.66054096 | 5.88E-18 | 6.57E-16 |
| TSHZ3 | 15.73666224 | 7.208105403 | 9.10E-21 | 1.84E-18 |
| PDGFRA | 15.72531639 | 7.293918272 | 1.22E-28 | 1.10E-25 |
| PRRX1 | 15.71880504 | 7.254534957 | 5.73E-27 | 3.27E-24 |
| MATN2 | 15.66567082 | 7.696585946 | 2.83E-34 | 6.81E-31 |
| BICC1 | 15.62642492 | 7.075466799 | 1.68E-35 | 5.19E-32 |
| RXRG | 15.57565624 | 7.098303699 | 6.52E-14 | 3.45E-12 |
| CDH11 | 15.56382746 | 7.16998515 | 5.18E-24 | 1.67E-21 |
| RBMS3 | 15.47550469 | 7.207626282 | 4.44E-44 | 4.82E-40 |
| COL16A1 | 15.46724019 | 7.37475934 | 8.28E-28 | 6.19E-25 |
| LOC119878328 | 15.40460288 | 6.670159308 | 0.000423087 | 0.001958052 |
| GJA1 | 15.33110872 | 6.9444827 | 1.32E-11 | 4.35E-10 |
| APOD | 15.33046106 | 8.047265251 | 8.59E-12 | 2.98E-10 |

| | | | | | |
|---|---|---|---|---|---|
| ZNF469 | 15.32047665 | 6.99878262 | 5.58E-17 | 5.16E-15 | |
| KIF1A | 15.31671963 | 7.219615677 | 1.27E-11 | 4.19E-10 | |
| COL1A1 | 15.22441029 | 12.75502992 | 4.13E-15 | 2.76E-13 | |
| ZNF423 | 15.19782746 | 6.711136517 | 1.26E-22 | 3.20E-20 | |
| KRT7 | 15.10261349 | 6.427332893 | 1.01E-09 | 2.21E-08 | |
| PRELP | 15.05297958 | 7.173414814 | 1.14E-15 | 8.43E-14 | |
| SCARA3 | 15.05210942 | 6.884251131 | 1.08E-23 | 3.22E-21 | |
| SFRP1 | 15.0502944 | 7.243322064 | 1.40E-07 | 1.83E-06 | |
| POSTN | 15.03075519 | 6.332750616 | 1.52E-23 | 4.39E-21 | |
| ROBO2 | 14.90343588 | 6.327492738 | 6.42E-13 | 2.85E-11 | |
| IGFBP3 | 14.87417791 | 6.431491239 | 2.54E-17 | 2.45E-15 | |
| CCN2 | 14.8115814 | 6.840881217 | 7.19E-19 | 9.56E-17 | encodes the connective tissue growth factor (CTGF). Upregulation is linked to epithelial to mesenchymal transition. |
| PAPPA | 14.78882374 | 6.231791213 | 5.29E-21 | 1.11E-18 | |
| LOC119865237 | 14.78712397 | 6.561828282 | 2.05E-26 | 1.06E-23 | |
| GALNT15 | 14.7438015 | 6.716265745 | 8.98E-15 | 5.59E-13 | |
| FLRT2 | 14.74151793 | 6.315968752 | 8.66E-38 | 3.13E-34 | |
| CRABP2 | 14.73990114 | 6.173123388 | 6.93E-21 | 1.43E-18 | |
| NGFR | 14.72088235 | 7.204804272 | 8.89E-08 | 1.22E-06 | |
| CGNL1 | 14.70615045 | 8.41816762 | 1.59E-17 | 1.60E-15 | |
| ZFHX4 | 14.63212143 | 6.367259148 | 8.13E-13 | 3.54E-11 | |
| NES | 14.58612989 | 6.847035077 | 5.53E-20 | 9.58E-18 | |
| CD248 | 14.5712806 | 8.061660972 | 7.45E-26 | 3.59E-23 | |
| VIT | 14.56096482 | 6.863528986 | 2.27E-10 | 5.74E-09 | |
| FZD1 | 14.54439463 | 6.670444426 | 2.36E-17 | 2.29E-15 | |
| PIEZO2 | 14.52966021 | 6.218250369 | 1.52E-09 | 3.18E-08 | |
| DACT1 | 14.50324275 | 6.139928139 | 3.30E-18 | 3.96E-16 | |
| C8H14orf132 | 14.49534232 | 5.96696616 | 6.27E-08 | 8.99E-07 | |
| ANXA8L1 | 14.4897632 | 6.05475453 | 1.25E-12 | 5.24E-11 | |
| RGS5 | 14.48415901 | 6.772580652 | 2.59E-15 | 1.81E-13 | |

| | | | | |
|---|---|---|---|---|
| GJB3 | 14.45587764 | 5.893182693 | 2.60E-13 | 1.24E-11 |
| CMSS1_1 | 14.43289486 | 5.980803143 | 1.05E-11 | 3.55E-10 |
| MEOX2 | 14.43038487 | 6.208456764 | 5.09E-13 | 2.31E-11 |
| HMCN1 | 14.41713573 | 6.101927286 | 8.59E-14 | 4.43E-12 |
| CHRDL1 | 14.40907893 | 6.878079556 | 1.42E-17 | 1.46E-15 |
| C7 | 14.3097046 | 6.219925837 | 4.42E-15 | 2.95E-13 |
| GAP43 | 14.25077235 | 5.547285528 | 3.41E-09 | 6.53E-08 |
| PREX2 | 14.23708975 | 6.222326388 | 1.68E-31 | 2.42E-28 |
| COL11A1 | 14.22953229 | 5.729198877 | 1.85E-06 | 1.76E-05 |
| HEYL | 14.22190644 | 6.007736882 | 1.81E-22 | 4.45E-20 |
| VGLL3 | 14.21368696 | 6.33946831 | 1.21E-19 | 1.91E-17 |
| SOGA3 | 14.16806478 | 5.581300004 | 7.24E-09 | 1.29E-07 |
| MXRA5 | 14.11894434 | 9.411253285 | 5.04E-18 | 5.69E-16 |
| SLC4A3 | 14.11634908 | 5.68262701 | 1.13E-18 | 1.46E-16 |
| VCAM1 | 14.05253769 | 5.739703474 | 3.60E-14 | 2.01E-12 |
| MEDAG | 14.04069444 | 5.869944215 | 5.14E-21 | 1.09E-18 |
| LOC119869296 | 14.01545138 | 5.314127023 | 1.13E-07 | 1.51E-06 |
| PAPSS2 | 14.00755261 | 5.673056652 | 1.73E-15 | 1.23E-13 |
| TRABD2B | 13.94546859 | 5.626842654 | 2.88E-24 | 9.46E-22 |
| LOC119879275 | 13.94119644 | 17.75686947 | 4.85E-05 | 0.000305887 |
| CCDC85A | 13.91059196 | 5.756846141 | 7.40E-14 | 3.86E-12 |
| PTGDS | 13.89086388 | 5.885449558 | 2.95E-15 | 2.03E-13 |
| LOC119864741 | 13.87164018 | 5.767150758 | 2.53E-27 | 1.61E-24 |
| EBF2 | 13.85731548 | 5.952552217 | 2.12E-10 | 5.41E-09 |
| LOC119873679 | 13.85084041 | 5.347530844 | 4.83E-19 | 6.62E-17 |
| LOC119877828 | 13.78564781 | 5.265350123 | 7.15E-13 | 3.15E-11 |
| PTPRG | 13.78463841 | 6.144773181 | 1.14E-17 | 1.21E-15 |
| LARP6 | 13.77015875 | 5.702433884 | 1.20E-22 | 3.13E-20 |
| EVC2 | 13.76285419 | 5.314406121 | 4.05E-23 | 1.11E-20 |
| TBX15 | 13.75671007 | 5.438366497 | 8.48E-20 | 1.39E-17 |
| LOC106558423 | 13.74103124 | 5.045417678 | 4.62E-06 | 3.89E-05 |
| LOC119878310 | 13.70116462 | 8.504472532 | 0.000159099 | 0.00084431 |
| ENPP1 | 13.66489259 | 5.222298029 | 4.13E-20 | 7.33E-18 |
| OSBPL5 | 13.65868924 | 5.517923086 | 1.44E-28 | 1.25E-25 |
| KIAA0408 | 13.65844443 | 5.412608996 | 3.92E-11 | 1.16E-09 |
| CPXM1 | 13.64986773 | 5.012312956 | 1.71E-14 | 1.01E-12 |
| GEM | 13.64835566 | 5.582412638 | 7.75E-30 | 8.84E-27 |
| CCN3 | 13.64268983 | 5.168408285 | 1.48E-12 | 6.06E-11 |
| CCDC8 | 13.61205289 | 5.24310623 | 6.83E-24 | 2.15E-21 |

| | | | | |
|---|---|---|---|---|
| GUCY1A2 | 13.60252832 | 6.360978599 | 1.19E-14 | 7.22E-13 |
| UNC13A | 13.60135125 | 4.900423118 | 1.04E-08 | 1.78E-07 |
| PCDH7 | 13.59653137 | 5.511819059 | 2.89E-15 | 1.99E-13 |
| SEMA3D | 13.58059381 | 5.551446621 | 3.18E-13 | 1.49E-11 |
| SMARCA1 | 13.55797846 | 5.180676957 | 5.42E-27 | 3.17E-24 |
| COL6A2 | 13.5513253 | 10.74682901 | 1.07E-20 | 2.12E-18 |
| TEAD4 | 13.53128444 | 5.974536917 | 1.79E-25 | 8.10E-23 |
| SCN7A | 13.51853623 | 9.00218279 | 1.07E-05 | 8.07E-05 |
| EDIL3 | 13.50879297 | 5.453661118 | 4.15E-10 | 9.97E-09 |
| CAV1 | 13.50464686 | 5.939884945 | 4.49E-32 | 6.96E-29 |
| NCAM1 | 13.49197969 | 6.404287053 | 5.09E-17 | 4.74E-15 |
| LOC111091587 | 13.47106137 | 4.779784959 | 1.03E-07 | 1.39E-06 |
| IGSF9B | 13.4629722 | 4.896675855 | 2.83E-08 | 4.41E-07 |
| IGF2 | 13.46059778 | 5.132189278 | 5.98E-10 | 1.38E-08 |
| ALDH1L1 | 13.44935255 | 5.06439033 | 9.88E-14 | 5.06E-12 |
| ALDH1A2 | 13.44658311 | 4.801131213 | 2.26E-10 | 5.73E-09 |
| TMEM100 | 13.44294702 | 4.845363455 | 4.50E-11 | 1.32E-09 |
| NTN4 | 13.44045025 | 5.670322774 | 1.73E-19 | 2.64E-17 |
| ITGA11 | 13.41085692 | 5.064373852 | 6.96E-11 | 1.96E-09 |
| EPHA3 | 13.38789731 | 4.812435628 | 1.16E-11 | 3.90E-10 |
| LIFR | 13.37680504 | 5.355601014 | 2.01E-20 | 3.89E-18 |
| DHRS3 | 13.3733051 | 6.107759619 | 1.88E-26 | 9.96E-24 |
| EEF1A2 | 13.36478465 | 4.673069069 | 7.10E-07 | 7.58E-06 |
| RGMA | 13.35827225 | 5.033652575 | 2.12E-19 | 3.10E-17 |
| WIF1 | 13.31930385 | 4.749472964 | 3.04E-05 | 0.000203658 |
| SPEG | 13.31003317 | 5.590054489 | 6.82E-17 | 6.21E-15 |
| SHANK1 | 13.29362383 | 4.597980021 | 6.13E-13 | 2.74E-11 |
| MOXD1 | 13.28728768 | 5.064869677 | 1.44E-13 | 7.15E-12 |
| RASL12 | 13.22663401 | 5.397181026 | 1.65E-16 | 1.43E-14 |
| LOC119879511 | 13.21853012 | 9.029273631 | 7.00E-06 | 5.57E-05 |
| CEMIP | 13.21199376 | 4.571570613 | 5.84E-09 | 1.06E-07 |
| EFS | 13.18212451 | 5.25967476 | 7.97E-29 | 7.85E-26 |
| NYNRIN | 13.17334091 | 5.693339214 | 3.02E-18 | 3.64E-16 |
| TBX3 | 13.16536056 | 4.766295879 | 1.54E-12 | 6.31E-11 |
| KERA | 13.14849007 | 4.775854388 | 2.11E-07 | 2.61E-06 |
| NOVA2 | 13.11687691 | 4.817920253 | 1.30E-19 | 2.03E-17 |
| DSEL | 13.11086327 | 4.932812685 | 2.82E-16 | 2.32E-14 |
| GULP1 | 13.07453656 | 6.205773202 | 5.89E-25 | 2.20E-22 |
| TBX18 | 13.07206846 | 4.954935686 | 6.84E-16 | 5.33E-14 |

| | | | | |
|---|---|---|---|---|
| EGFLAM | 13.05455271 | 4.635892136 | 1.22E-12 | 5.10E-11 |
| FJX1 | 13.05219893 | 4.577941652 | 3.00E-16 | 2.44E-14 |
| ETV5 | 13.04132792 | 4.770310165 | 1.08E-28 | 1.02E-25 |
| MMRN2 | 13.03511188 | 5.648837952 | 1.24E-17 | 1.29E-15 |
| FIBIN | 12.96225281 | 4.83288974 | 7.14E-15 | 4.55E-13 |
| PHLDB2 | 12.95259841 | 5.435569026 | 3.12E-20 | 5.73E-18 |
| S1PR3 | 12.90909864 | 5.351390931 | 9.99E-08 | 1.35E-06 |
| SHISAL1 | 12.88939622 | 4.806581946 | 4.75E-10 | 1.12E-08 |
| OLFML1 | 12.8720113 | 4.96057721 | 4.19E-25 | 1.64E-22 |
| MAGI1 | 12.87035328 | 4.962442061 | 1.81E-27 | 1.26E-24 |
| MMP16 | 12.86277249 | 4.258831381 | 7.55E-14 | 3.92E-12 |
| LGR6 | 12.85592838 | 4.177854828 | 2.11E-12 | 8.42E-11 |
| TRIL | 12.84857706 | 4.44761793 | 1.52E-15 | 1.08E-13 |
| CAV2 | 12.81693666 | 6.569660774 | 3.26E-25 | 1.41E-22 |
| LOC119872564 | 12.81668034 | 4.270167136 | 2.03E-16 | 1.73E-14 |
| PRRX2 | 12.80563023 | 4.257936983 | 7.00E-24 | 2.17E-21 |
| COL1A2 | 12.79848265 | 11.23287211 | 6.78E-17 | 6.20E-15 |
| SCRT1 | 12.79051413 | 4.085631119 | 4.33E-07 | 4.93E-06 |
| CLCNKA | 12.79014363 | 4.32237657 | 4.20E-14 | 2.32E-12 |
| ANGPTL4 | 12.7799846 | 4.647723011 | 6.02E-20 | 1.03E-17 |
| CYYR1 | 12.75748395 | 5.460021629 | 3.78E-13 | 1.76E-11 |
| PLCXD3 | 12.7523433 | 4.2982448 | 4.04E-12 | 1.52E-10 |
| APLNR | 12.73156286 | 4.814113598 | 1.18E-08 | 2.00E-07 |
| IGFBP2 | 12.71188912 | 4.23572884 | 8.99E-10 | 2.00E-08 |
| MEIS2 | 12.70458916 | 4.147381158 | 2.23E-16 | 1.86E-14 |
| ADCYAP1R1 | 12.70432706 | 4.096303407 | 3.18E-06 | 2.82E-05 |
| LOC119864392 | 12.69824946 | 4.001893364 | 1.50E-06 | 1.46E-05 |
| LOC106559497 | 12.6731187 | 4.838413974 | 2.09E-19 | 3.07E-17 |
| COL21A1 | 12.64616987 | 4.363752276 | 7.06E-17 | 6.35E-15 |
| TNC | 12.6261406 | 7.192386912 | 3.59E-12 | 1.38E-10 |
| DCX | 12.62545067 | 4.003912989 | 1.27E-09 | 2.72E-08 |
| MRGPRF | 12.59464364 | 4.727355096 | 3.19E-19 | 4.52E-17 |
| ACTG2 | 12.58657121 | 4.969615936 | 7.60E-10 | 1.72E-08 |
| LOC102152259 | 12.57812867 | 4.017470459 | 2.00E-14 | 1.17E-12 |
| DES | 12.57602323 | 4.285044797 | 8.96E-12 | 3.09E-10 |
| LOC119877844 | 12.57569605 | 3.879345115 | 1.23E-05 | 9.19E-05 |
| HSPG2 | 12.57143663 | 10.29206403 | 1.78E-27 | 1.26E-24 |
| LOC119867967 | 12.56178657 | 4.131071715 | 5.28E-15 | 3.47E-13 |
| BRINP1 | 12.56042196 | 4.206977211 | 4.14E-07 | 4.74E-06 |

| | | | | |
|---|---|---|---|---|
| LDOC1 | 12.55401999 | 4.250265756 | 4.53E-19 | 6.26E-17 |
| RFPL4B | 12.54823977 | 3.884024801 | 1.24E-06 | 1.24E-05 |
| MMP15 | 12.54576246 | 5.869901515 | 1.97E-20 | 3.84E-18 |
| SPRY4 | 12.54060927 | 4.356240254 | 1.86E-19 | 2.78E-17 |
| ROBO4 | 12.53957281 | 5.290481424 | 1.50E-17 | 1.53E-15 |
| CDH19 | 12.52721386 | 7.671395704 | 2.41E-05 | 0.000166627 |
| TWIST1 | 12.52624244 | 3.942604862 | 4.93E-15 | 3.26E-13 |
| COPZ2 | 12.49891992 | 4.374159788 | 1.02E-18 | 1.32E-16 |
| IGSF11 | 12.49817556 | 5.215672557 | 1.09E-08 | 1.85E-07 |
| SEMA6A | 12.49324067 | 5.875053922 | 1.50E-24 | 5.06E-22 |
| KIF26B | 12.46921106 | 6.104668033 | 3.01E-20 | 5.58E-18 |
| FABP4 | 12.42538735 | 6.068990739 | 2.95E-07 | 3.52E-06 |
| LOC106558182 | 12.42125113 | 5.264041938 | 6.59E-09 | 1.19E-07 |
| COL4A1 | 12.41569556 | 9.554980578 | 6.48E-20 | 1.09E-17 |
| KANK4 | 12.40428936 | 5.61919181 | 2.62E-10 | 6.50E-09 |
| FRMD3 | 12.35815848 | 4.778849301 | 1.42E-21 | 3.17E-19 |
| LOC119878330 | 12.33854257 | 14.99599871 | 4.03E-05 | 0.000260216 |
| CHL1 | 12.32792884 | 7.020470826 | 3.32E-10 | 8.08E-09 |
| CAVIN3 | 12.31329053 | 7.085159352 | 3.26E-30 | 3.93E-27 |
| KCNF1 | 12.2969004 | 3.89256218 | 1.58E-10 | 4.13E-09 |
| LOC119878273 | 12.28729686 | 7.699390105 | 0.000215511 | 0.001096888 |
| MPZ | 12.27483269 | 9.323378747 | 3.12E-05 | 0.0002081 |
| MGAT5B | 12.26981508 | 4.194537156 | 4.47E-16 | 3.55E-14 |
| PALMD | 12.25326592 | 5.058759543 | 2.24E-15 | 1.58E-13 |
| S100A16 | 12.2341982 | 4.357202295 | 3.44E-20 | 6.26E-18 |
| LOC111090688 | 12.22628639 | 3.833272048 | 2.56E-14 | 1.47E-12 |
| EMCN | 12.22593259 | 4.787314923 | 1.57E-18 | 1.98E-16 |
| EPHB2 | 12.22213935 | 4.132487391 | 6.88E-15 | 4.41E-13 |
| SGCA | 12.19950727 | 3.880283369 | 2.63E-15 | 1.83E-13 |
| PDZRN3 | 12.19656128 | 4.184390962 | 1.37E-17 | 1.41E-15 |
| SOX18 | 12.1816592 | 5.255825069 | 2.90E-19 | 4.14E-17 |
| RTL1 | 12.17094902 | 3.534828318 | 5.02E-09 | 9.24E-08 |
| LOC487177 | 12.15525376 | 3.681084018 | 1.59E-10 | 4.15E-09 |
| NOX4 | 12.15445504 | 4.028453378 | 6.07E-25 | 2.23E-22 |
| NR2F2 | 12.15131969 | 7.365818365 | 1.69E-22 | 4.21E-20 |
| STOX2 | 12.14856393 | 5.110823753 | 1.11E-14 | 6.79E-13 |
| COL28A1 | 12.13286924 | 5.683905229 | 1.78E-11 | 5.69E-10 |
| GTF2IRD1 | 12.12323343 | 3.941145528 | 2.74E-19 | 3.95E-17 |
| ALPL | 12.10659898 | 3.884274653 | 2.63E-05 | 0.000179192 |

| | | | | |
|---|---|---|---|---|
| SPHKAP | 12.10615328 | 3.524452198 | 2.44E-07 | 2.98E-06 |
| TFAP2C | 12.09402271 | 3.403617536 | 2.30E-09 | 4.64E-08 |
| ESM1 | 12.07477132 | 3.311197393 | 4.41E-10 | 1.05E-08 |
| GPX8 | 12.06806069 | 3.80028493 | 7.05E-15 | 4.50E-13 |
| JAKMIP2 | 12.06103957 | 3.720585145 | 9.34E-08 | 1.27E-06 |
| ZPLD1 | 12.05501595 | 3.536282762 | 2.49E-06 | 2.28E-05 |
| GFRA2 | 12.05372995 | 3.607608175 | 4.48E-09 | 8.33E-08 |
| AGMO | 12.05252115 | 3.958320085 | 2.10E-08 | 3.35E-07 |
| CX3CL1 | 12.04886208 | 3.678178006 | 1.49E-19 | 2.28E-17 |
| TC2N | 12.04870439 | 4.71602315 | 3.42E-11 | 1.03E-09 |
| CLDN11 | 12.04386847 | 4.044267938 | 3.55E-07 | 4.14E-06 |
| CARMIL3 | 12.03028183 | 7.773315059 | 2.72E-08 | 4.24E-07 |
| LOC119876084 | 12.02981328 | 3.613624623 | 1.66E-13 | 8.22E-12 |
| LOC102154504 | 12.02297727 | 3.359328341 | 3.37E-09 | 6.47E-08 |
| MECOM | 12.01982517 | 4.467735076 | 2.88E-16 | 2.35E-14 |
| B4GALNT3 | 12.0190251 | 4.403273675 | 1.99E-11 | 6.34E-10 |
| SERPINB11 | 12.01248404 | 3.31662447 | 7.24E-09 | 1.29E-07 |
| NDN | 12.00843338 | 3.672134405 | 1.85E-19 | 2.78E-17 |
| HID1 | 11.99615574 | 5.067227118 | 4.10E-25 | 1.64E-22 |
| SFRP4 | 11.99121694 | 8.119386903 | 2.81E-06 | 2.54E-05 |
| HOXD8 | 11.96131058 | 3.895003356 | 4.07E-21 | 8.82E-19 |
| DIO2 | 11.9558978 | 3.280785668 | 5.48E-07 | 6.07E-06 |
| C1QTNF3 | 11.94968539 | 3.193999964 | 7.65E-06 | 6.03E-05 |
| TPBG | 11.936942 | 3.362081484 | 6.68E-12 | 2.41E-10 |
| LOC111095505 | 11.93526491 | 3.717640573 | 3.00E-08 | 4.63E-07 |
| LOC102156205 | 11.93252996 | 3.393745003 | 4.28E-16 | 3.41E-14 |
| NPR1 | 11.93016221 | 5.218163456 | 6.66E-16 | 5.21E-14 |
| MID1 | 11.92506894 | 3.890675618 | 1.57E-25 | 7.24E-23 |
| PI15 | 11.90638767 | 3.850571099 | 5.07E-15 | 3.34E-13 |
| HSPB7 | 11.9051579 | 3.682197636 | 2.31E-13 | 1.11E-11 |
| MEOX1 | 11.88994747 | 3.804138403 | 1.13E-08 | 1.92E-07 |
| MARK1 | 11.88022938 | 4.092912298 | 9.48E-12 | 3.26E-10 |
| CYGB | 11.87548059 | 3.7114943 | 1.62E-12 | 6.59E-11 |
| CLSTN3 | 11.87277044 | 3.388682308 | 3.70E-13 | 1.72E-11 |
| ADAMTS17 | 11.86557498 | 3.753144176 | 5.81E-11 | 1.67E-09 |
| FAM13C | 11.84889661 | 4.453171218 | 6.14E-18 | 6.82E-16 |
| KRT8 | 11.83397254 | 3.145559915 | 5.21E-08 | 7.61E-07 |
| TSPAN7 | 11.83289885 | 4.357018954 | 4.60E-31 | 6.23E-28 |
| CRYAB | 11.81689793 | 4.748457511 | 3.86E-13 | 1.78E-11 |

| | | | | |
|---|---|---|---|---|
| SOX9 | 11.80633482 | 3.989021702 | 2.54E-09 | 5.04E-08 |
| ARHGEF5 | 11.80326835 | 3.553621375 | 2.20E-16 | 1.86E-14 |
| HSPB8 | 11.79859639 | 3.78070562 | 5.58E-15 | 3.63E-13 |
| TLE6 | 11.76936958 | 3.200675248 | 1.78E-09 | 3.66E-08 |
| SOSTDC1 | 11.76212537 | 3.900902777 | 2.28E-09 | 4.61E-08 |
| IGLON5 | 11.75794312 | 3.10733562 | 1.27E-09 | 2.71E-08 |
| THSD7A | 11.7374799 | 4.992155607 | 6.96E-08 | 9.84E-07 |
| RBP1 | 11.7373235 | 3.679218828 | 3.82E-15 | 2.56E-13 |
| LOC111093104 | 11.73619554 | 3.547672434 | 3.52E-10 | 8.54E-09 |
| LOC119866907 | 11.72279984 | 3.79673963 | 4.55E-25 | 1.73E-22 |
| DCLK1 | 11.72128722 | 6.218444428 | 8.64E-19 | 1.14E-16 |
| LOC111091714 | 11.69997133 | 3.980732145 | 2.79E-23 | 7.87E-21 |
| SRPX2 | 11.68690662 | 3.411439402 | 8.60E-13 | 3.72E-11 |
| COL24A1 | 11.68143009 | 3.509472319 | 7.38E-09 | 1.31E-07 |
| AEBP1 | 11.67940417 | 10.2327865 | 6.30E-14 | 3.36E-12 |
| GABRE | 11.67314304 | 3.349752128 | 2.57E-11 | 7.97E-10 |
| LMCD1 | 11.67251661 | 4.382125544 | 1.71E-18 | 2.14E-16 |
| TNNT2 | 11.66564038 | 2.977306909 | 3.65E-08 | 5.53E-07 |
| CRABP1 | 11.66563711 | 3.378170802 | 7.30E-09 | 1.30E-07 |
| EMX2 | 11.64748794 | 2.969019488 | 6.32E-07 | 6.85E-06 |
| KCTD16 | 11.63351633 | 3.48667817 | 4.10E-19 | 5.74E-17 |
| MYO10 | 11.62505805 | 6.546923267 | 1.18E-19 | 1.87E-17 |
| LOC610632 | 11.61986124 | 3.480744978 | 8.00E-11 | 2.23E-09 |
| LOC111091069 | 11.61290659 | 2.931187514 | 6.15E-06 | 4.97E-05 |
| SCD5 | 11.60588386 | 4.097536566 | 1.52E-15 | 1.08E-13 |
| ADGRL4 | 11.59223126 | 5.267986385 | 3.28E-12 | 1.27E-10 |
| P3H2 | 11.57812651 | 3.987354581 | 7.69E-13 | 3.36E-11 |
| APOC1 | 11.57554014 | 3.061604416 | 1.93E-11 | 6.16E-10 |
| LOC119874319 | 11.57521748 | 2.888663309 | 2.42E-06 | 2.23E-05 |
| ISM1 | 11.57348102 | 4.406443734 | 6.34E-12 | 2.31E-10 |
| LOC111097810 | 11.56880073 | 3.372292027 | 2.29E-13 | 1.10E-11 |
| LOC119879328 | 11.56533135 | 15.36162142 | 7.89E-06 | 6.20E-05 |
| TCIM | 11.55962802 | 4.18146574 | 7.74E-11 | 2.16E-09 |
| ADAMTS15 | 11.55089887 | 3.95777953 | 3.09E-16 | 2.50E-14 |
| LRRC17 | 11.52412794 | 2.863190761 | 8.22E-11 | 2.28E-09 |
| RORB | 11.51238471 | 3.518624989 | 2.12E-07 | 2.63E-06 |
| DOCK6 | 11.48919038 | 8.160869452 | 7.33E-16 | 5.68E-14 |
| CAP2 | 11.48099146 | 3.378682166 | 4.64E-13 | 2.13E-11 |
| SLC2A10 | 11.48004996 | 3.192295794 | 8.32E-19 | 1.10E-16 |

| | | | | |
|---|---|---|---|---|
| KCNMA1 | 11.47602799 | 3.679863162 | 1.69E-09 | 3.49E-08 |
| GFRA1 | 11.47037763 | 5.283936631 | 8.68E-07 | 9.02E-06 |
| MME | 11.46206059 | 4.76268786 | 3.84E-06 | 3.32E-05 |
| UNC5C | 11.43977013 | 3.064504864 | 2.23E-05 | 0.00015507 |
| LOC111093353 | 11.43614628 | 5.944812484 | 2.32E-29 | 2.40E-26 |
| TM4SF18 | 11.43229483 | 3.593695066 | 2.77E-09 | 5.44E-08 |
| OSR1 | 11.42425091 | 3.182186319 | 8.90E-09 | 1.55E-07 |
| PDLIM3 | 11.42296182 | 3.61189254 | 5.56E-09 | 1.02E-07 |
| LOC119864159 | 11.40593679 | 2.728620699 | 3.85E-06 | 3.32E-05 |
| MASP1 | 11.38117351 | 3.03482701 | 7.70E-07 | 8.15E-06 |
| LOC119877031 | 11.33805248 | 4.487395831 | 1.47E-15 | 1.06E-13 |
| ALDH1A3 | 11.33544672 | 4.598106088 | 1.21E-09 | 2.61E-08 |
| LOC102154980 | 11.32658258 | 3.561132485 | 1.32E-17 | 1.37E-15 |
| LOC119878322 | 11.31936695 | 14.69566193 | 3.27E-05 | 0.000216558 |
| LOC102152218 | 11.30730463 | 2.727525934 | 1.61E-05 | 0.00011641 |
| SLC17A7 | 11.30432038 | 2.616667672 | 1.47E-08 | 2.44E-07 |
| MALL | 11.30237586 | 3.941123307 | 1.23E-09 | 2.65E-08 |
| PMP22 | 11.29233114 | 8.620808931 | 1.71E-13 | 8.39E-12 |
| PDPN | 11.286973 | 6.647758028 | 2.47E-14 | 1.43E-12 |
| LOC102151524 | 11.2790967 | 2.598681466 | 2.10E-05 | 0.000146769 |
| HSF4_1 | 11.27393627 | 2.773183174 | 8.30E-13 | 3.60E-11 |
| PRKD1 | 11.26777637 | 3.715585777 | 1.19E-10 | 3.19E-09 |
| RHOJ | 11.26678625 | 4.013607109 | 6.13E-22 | 1.43E-19 |
| LOC102154920 | 11.26238477 | 2.616921063 | 3.10E-08 | 4.77E-07 |
| STAC | 11.25399292 | 3.261280381 | 1.40E-15 | 1.02E-13 |
| MAMSTR | 11.23130509 | 4.089798662 | 2.00E-12 | 8.02E-11 |
| LOC119866445 | 11.22608518 | 3.296729671 | 1.43E-09 | 3.01E-08 |
| C1QC | 11.22594005 | 6.399336754 | 1.90E-15 | 1.34E-13 |
| CXCL13 | 11.2209881 | 2.528879498 | 3.67E-05 | 0.000239602 |
| LOC119864153 | 11.21481344 | 3.935451002 | 4.79E-06 | 4.01E-05 |
| CA3 | 11.21365741 | 3.762635605 | 0.000149602 | 0.000799755 |
| LOC119869324 | 11.17202199 | 3.432740134 | 2.76E-11 | 8.47E-10 |
| ADAMTSL1 | 11.1690002 | 4.727055178 | 9.14E-18 | 9.96E-16 |
| TNS4 | 11.16702061 | 3.065862218 | 2.75E-09 | 5.41E-08 |
| C5H11orf87 | 11.15654823 | 2.450042549 | 3.37E-06 | 2.96E-05 |
| TENM4 | 11.15140435 | 5.271283545 | 3.25E-09 | 6.29E-08 |
| MAPK4 | 11.14924935 | 2.478895616 | 0.000317449 | 0.001529583 |
| LOC111092369 | 11.14755623 | 2.609599655 | 3.21E-15 | 2.17E-13 |
| BFSP1 | 11.14430185 | 2.985442105 | 7.03E-17 | 6.35E-15 |

| | | | | |
|---|---|---|---|---|
| LOC111095260 | 11.14217265 | 4.140440403 | 1.26E-09 | 2.70E-08 |
| FAT3 | 11.14132248 | 3.131720569 | 5.38E-08 | 7.83E-07 |
| LOC119872510 | 11.13900577 | 2.613160193 | 1.21E-05 | 9.04E-05 |
| LOC487172 | 11.12640094 | 3.132704052 | 5.43E-16 | 4.29E-14 |
| EVA1C | 11.12469766 | 3.491935816 | 3.29E-14 | 1.85E-12 |
| NPTX1 | 11.11997326 | 4.987712299 | 7.04E-17 | 6.35E-15 |
| TWIST2 | 11.11094576 | 2.623191437 | 2.92E-07 | 3.50E-06 |
| PLK5 | 11.11003587 | 2.643836384 | 7.04E-08 | 9.92E-07 |
| GFPT2 | 11.10527659 | 5.356387582 | 4.52E-18 | 5.15E-16 |
| SYT5 | 11.1039384 | 2.458909434 | 4.06E-07 | 4.66E-06 |
| LOC119864158 | 11.10149688 | 2.42767034 | 1.41E-06 | 1.39E-05 |
| ZBTB47 | 11.09896117 | 6.755068166 | 1.12E-26 | 6.22E-24 |
| IL33 | 11.09481843 | 7.085685151 | 1.35E-19 | 2.10E-17 |
| H19 | 11.08034399 | 2.521139521 | 5.58E-05 | 0.000345104 |
| RTN4RL2 | 11.07797109 | 2.429426293 | 2.52E-09 | 5.02E-08 |
| KITLG | 11.06290533 | 4.044293174 | 1.29E-12 | 5.38E-11 |
| LTBP4 | 11.06050328 | 9.207060682 | 2.12E-16 | 1.79E-14 |
| LAMA4 | 11.05955299 | 8.980349123 | 5.23E-24 | 1.67E-21 |
| PTGFR | 11.05684176 | 2.750295219 | 1.48E-08 | 2.44E-07 |
| GJB5 | 11.0471877 | 2.423380785 | 7.89E-08 | 1.09E-06 |
| LOC480552 | 11.04717699 | 2.525863563 | 4.01E-10 | 9.66E-09 |
| DKK2 | 11.03319992 | 4.262574177 | 1.57E-09 | 3.28E-08 |
| CACHD1 | 11.03252019 | 7.148159665 | 1.21E-08 | 2.03E-07 |
| SULT4A1 | 11.02322109 | 2.363131979 | 3.08E-09 | 6.00E-08 |
| SBSPON | 11.02277777 | 5.200334323 | 1.35E-07 | 1.76E-06 |
| CPE | 11.01995639 | 6.080782634 | 6.40E-20 | 1.08E-17 |
| ANTXR1 | 11.01854002 | 8.115955534 | 1.17E-13 | 5.91E-12 |
| LOC111095882 | 11.01501188 | 3.31523493 | 1.37E-08 | 2.28E-07 |
| WSCD1 | 11.00759663 | 3.169859854 | 1.80E-12 | 7.27E-11 |
| METTL11B | 11.00381744 | 2.328364457 | 5.24E-07 | 5.83E-06 |
| ZNF385D | 11.00065626 | 2.499794939 | 7.26E-07 | 7.73E-06 |
| SLC8A3 | 10.99790886 | 2.718709933 | 4.59E-07 | 5.21E-06 |
| SHISA9 | 10.9974435 | 2.508087526 | 2.02E-11 | 6.43E-10 |
| OR10J4 | 10.98781728 | 2.317032259 | 5.06E-06 | 4.20E-05 |
| C1QTNF2 | 10.98561167 | 2.802910731 | 5.46E-13 | 2.46E-11 |
| DRD1 | 10.97848029 | 2.663199757 | 2.10E-13 | 1.02E-11 |
| SNTA1 | 10.9666812 | 3.596492855 | 9.71E-13 | 4.17E-11 |
| WNT5A | 10.95794439 | 2.790312136 | 5.80E-12 | 2.13E-10 |
| LOC106558746 | 10.94245039 | 3.00941126 | 1.10E-13 | 5.56E-12 |

| | | | | |
|---|---|---|---|---|
| CASP14 | 10.9363529 | 2.258139915 | 5.14E-05 | 0.000321874 |
| LOC111096707 | 10.93443465 | 2.326078629 | 8.25E-11 | 2.29E-09 |
| LOC111090577 | 10.91718808 | 2.343799608 | 1.01E-09 | 2.21E-08 |
| EVA1A | 10.9166543 | 2.909587669 | 1.12E-12 | 4.69E-11 |
| SOX7 | 10.91482607 | 3.444068124 | 6.06E-11 | 1.72E-09 |
| LOC111095434 | 10.90805715 | 2.4461978 | 1.91E-10 | 4.92E-09 |
| LOC102152030 | 10.88680798 | 5.670654573 | 2.57E-26 | 1.27E-23 |
| ARHGEF25 | 10.87651903 | 5.623244227 | 4.07E-25 | 1.64E-22 |
| PNMA8A | 10.86487642 | 2.411852165 | 8.52E-15 | 5.32E-13 |
| LOC119864161 | 10.85277499 | 2.189886227 | 4.43E-07 | 5.04E-06 |
| MEGF10 | 10.85200609 | 2.881849571 | 1.49E-10 | 3.92E-09 |
| KCNJ8 | 10.85091665 | 2.967981804 | 3.61E-09 | 6.88E-08 |
| HRCT1 | 10.8494513 | 2.959705234 | 3.12E-14 | 1.76E-12 |
| FBN1 | 10.84325598 | 9.828047387 | 2.23E-16 | 1.86E-14 |
| APLN | 10.83755613 | 2.225292755 | 1.06E-11 | 3.59E-10 |
| LOC119870290 | 10.83264972 | 2.307527116 | 1.56E-09 | 3.27E-08 |
| KCNG3 | 10.82362191 | 2.511663459 | 1.78E-08 | 2.89E-07 |
| CXCL12 | 10.81915184 | 5.863387201 | 1.78E-30 | 2.27E-27 |
| LOC487174 | 10.81388157 | 2.71061161 | 2.34E-14 | 1.36E-12 |
| LOC119865381 | 10.81018161 | 2.845173809 | 1.70E-07 | 2.15E-06 |
| APOLD1 | 10.80896456 | 6.368641914 | 6.42E-13 | 2.85E-11 |
| HAPLN3 | 10.79762696 | 3.054062448 | 4.23E-12 | 1.59E-10 |
| SVEP1 | 10.79740105 | 7.763481647 | 2.38E-11 | 7.42E-10 |
| LOC119867049 | 10.7964311 | 2.148205297 | 8.28E-05 | 0.000486174 |
| CDH2 | 10.79512549 | 3.218099294 | 8.26E-09 | 1.45E-07 |
| PRR16 | 10.7874054 | 2.728281104 | 4.23E-10 | 1.01E-08 |
| IRX1 | 10.78258716 | 2.105754435 | 6.10E-10 | 1.40E-08 |
| GJC1 | 10.78143607 | 2.565965351 | 1.32E-08 | 2.19E-07 |
| TAFA5 | 10.78050392 | 4.001865554 | 9.42E-13 | 4.06E-11 |
| SEMA3F | 10.77544154 | 4.721786834 | 1.88E-22 | 4.59E-20 |
| LINGO1 | 10.77190363 | 2.513856883 | 3.30E-09 | 6.35E-08 |
| LOC119867040 | 10.76855395 | 2.208895703 | 9.40E-07 | 9.69E-06 |
| LOC100856382 | 10.76631725 | 7.186524798 | 3.88E-12 | 1.47E-10 |
| PRRG1 | 10.76626889 | 3.893427111 | 6.34E-17 | 5.83E-15 |
| LOC119870796 | 10.76625298 | 2.265695961 | 7.14E-09 | 1.28E-07 |
| SNCB | 10.76553107 | 2.061766124 | 2.59E-07 | 3.15E-06 |
| LOC119878274 | 10.76308069 | 12.85617867 | 3.02E-06 | 2.69E-05 |
| LOC119870941 | 10.76206854 | 2.264071226 | 2.25E-11 | 7.05E-10 |
| MAP6 | 10.75974117 | 2.925014047 | 1.35E-14 | 8.07E-13 |

| | | | | |
|---|---|---|---|---|
| SAMD5 | 10.74485015 | 2.862964391 | 9.84E-06 | 7.51E-05 |
| RHOU | 10.74463198 | 5.527759803 | 1.16E-19 | 1.87E-17 |
| LOC487176 | 10.74130281 | 2.875464867 | 2.51E-16 | 2.08E-14 |
| SORCS1 | 10.74063613 | 5.409485371 | 3.65E-07 | 4.24E-06 |
| LOC111090866 | 10.73584059 | 2.146687868 | 1.01E-07 | 1.36E-06 |
| DSC3 | 10.73458965 | 2.75759817 | 5.57E-07 | 6.15E-06 |
| LOC489707 | 10.70801234 | 2.50409101 | 2.03E-07 | 2.52E-06 |
| LOC119870146 | 10.69965968 | 2.203007422 | 2.35E-10 | 5.93E-09 |
| CLVS2 | 10.69078782 | 2.321817791 | 7.73E-09 | 1.37E-07 |
| EBF4 | 10.68555448 | 2.264531378 | 7.60E-10 | 1.72E-08 |
| IRX3 | 10.6811063 | 2.134969431 | 8.44E-08 | 1.17E-06 |
| ADIPOQ | 10.67591619 | 5.084153283 | 2.47E-05 | 0.000170202 |
| RERG | 10.67542664 | 2.596099559 | 4.47E-10 | 1.06E-08 |
| LOC484960 | 10.67372194 | 2.134560974 | 8.87E-07 | 9.18E-06 |
| KCNJ4 | 10.66271148 | 2.052779714 | 2.79E-08 | 4.35E-07 |
| EXOC3L1 | 10.65950992 | 2.778601576 | 2.55E-12 | 1.01E-10 |
| MDFI | 10.65320255 | 2.147216433 | 6.10E-06 | 4.93E-05 |
| HTR1B | 10.62765431 | 2.863548883 | 1.35E-11 | 4.44E-10 |
| GASK1A | 10.62085858 | 2.432308449 | 5.48E-11 | 1.58E-09 |
| PAPPA2 | 10.62026489 | 2.202379555 | 0.00018614 | 0.000964891 |
| CAPN6 | 10.60869867 | 1.931748437 | 2.42E-06 | 2.23E-05 |
| TMEM200A | 10.59207523 | 2.945520823 | 2.96E-14 | 1.68E-12 |
| CHMP4C | 10.59130583 | 2.246956191 | 1.28E-08 | 2.13E-07 |
| LAMB1 | 10.58296253 | 9.063892079 | 1.58E-22 | 3.98E-20 |
| SLC22A3 | 10.57785523 | 3.405082199 | 5.59E-06 | 4.58E-05 |
| SLIT2 | 10.57501398 | 8.664107242 | 1.87E-10 | 4.83E-09 |
| TFPI2 | 10.55296881 | 2.056309236 | 2.64E-05 | 0.000179439 |
| AMER3 | 10.54537537 | 2.586113926 | 2.78E-06 | 2.51E-05 |
| SIX4 | 10.52654583 | 2.065320461 | 4.84E-05 | 0.000305622 |
| IRX2 | 10.52583703 | 1.941932297 | 1.56E-08 | 2.56E-07 |
| COL26A1 | 10.52580655 | 2.148027729 | 0.00012909 | 0.000709553 |
| LOC111093976 | 10.50578704 | 2.059675442 | 3.47E-10 | 8.42E-09 |
| PRTG | 10.50313749 | 2.58026065 | 8.66E-09 | 1.52E-07 |
| LOC119875995 | 10.5003591 | 2.30542493 | 9.74E-12 | 3.34E-10 |
| CSMD2 | 10.49940982 | 4.806203396 | 1.37E-07 | 1.79E-06 |
| WWC3 | 10.49527948 | 6.001089169 | 6.17E-25 | 2.23E-22 |
| LOC119876505 | 10.48999822 | 2.46375219 | 4.29E-12 | 1.61E-10 |
| LOC102152011 | 10.47998749 | 2.110976669 | 8.22E-09 | 1.45E-07 |
| LOC119876269 | 10.47123349 | 1.840082039 | 3.02E-05 | 0.000202254 |

| | | | | |
|---|---|---|---|---|
| LOC111097168 | 10.46877929 | 1.854681372 | 7.24E-06 | 5.73E-05 |
| MROH5 | 10.44985113 | 2.99373809 | 5.09E-13 | 2.31E-11 |
| LOC119864948 | 10.43832244 | 1.89507154 | 3.72E-06 | 3.22E-05 |
| LOC119876393 | 10.43573155 | 1.870435583 | 7.72E-07 | 8.16E-06 |
| HUNK | 10.43185232 | 1.97624531 | 3.17E-09 | 6.14E-08 |
| OSR2 | 10.43174592 | 2.885339644 | 6.17E-06 | 4.98E-05 |
| LAPTM4B | 10.41316266 | 2.725917843 | 1.44E-12 | 5.93E-11 |
| ADAMTS1 | 10.40717846 | 7.215895678 | 2.80E-23 | 7.87E-21 |
| ABCB4 | 10.40455761 | 3.183952176 | 9.39E-11 | 2.56E-09 |
| CD34 | 10.40448012 | 8.106469101 | 5.48E-20 | 9.58E-18 |
| ADGRB1 | 10.40445192 | 3.038941594 | 2.67E-09 | 5.26E-08 |
| LOC119868377 | 10.40079998 | 2.222786006 | 1.40E-09 | 2.95E-08 |
| GJA5 | 10.40079159 | 2.721162974 | 1.37E-08 | 2.27E-07 |
| PLPP2 | 10.39875406 | 2.092247402 | 1.02E-14 | 6.22E-13 |
| CHSY3 | 10.39806797 | 2.337170965 | 7.12E-08 | 1.00E-06 |
| DSP | 10.39693185 | 3.144619997 | 2.54E-09 | 5.04E-08 |
| LOC119878305 | 10.39603455 | 7.224594621 | 2.95E-05 | 0.000198258 |
| LOC102152083 | 10.38854892 | 1.922306187 | 8.16E-11 | 2.27E-09 |
| EFCAB6 | 10.38586731 | 1.886038004 | 2.13E-09 | 4.34E-08 |
| LOC102156194 | 10.37925581 | 1.998064438 | 1.31E-12 | 5.43E-11 |
| THBS2 | 10.3748014 | 8.269298859 | 1.21E-07 | 1.60E-06 |
| LOC111095249 | 10.37214887 | 2.202763966 | 1.86E-08 | 3.01E-07 |
| HOXB6 | 10.36938878 | 2.21071781 | 3.62E-08 | 5.49E-07 |
| LOC111096582 | 10.36542509 | 3.39506284 | 2.50E-15 | 1.76E-13 |
| ID4 | 10.36376142 | 3.061394956 | 9.53E-09 | 1.65E-07 |
| LOC102153553 | 10.36178951 | 2.074573206 | 1.71E-11 | 5.53E-10 |
| LOC119873976 | 10.34584491 | 2.278191547 | 1.32E-07 | 1.73E-06 |
| SPARC | 10.34192757 | 10.3162628 | 2.76E-17 | 2.65E-15 |
| MLC1 | 10.34013339 | 1.741533375 | 0.000157924 | 0.000838688 |
| SH3GL2 | 10.33990765 | 2.408409303 | 5.88E-08 | 8.46E-07 |
| LOC102152258 | 10.3353368 | 2.141423865 | 5.18E-07 | 5.77E-06 |
| PDGFRB | 10.33521545 | 7.819219111 | 1.47E-16 | 1.29E-14 |
| TMEM132E | 10.32390246 | 2.03106693 | 1.93E-05 | 0.00013665 |
| SIX5 | 10.3122427 | 4.168694422 | 6.54E-16 | 5.13E-14 |
| LOC111094336 | 10.30560915 | 1.875759367 | 2.25E-09 | 4.55E-08 |
| LOC119867509 | 10.2992394 | 2.317232027 | 2.81E-15 | 1.95E-13 |
| LOC111093074 | 10.29873092 | 2.687733797 | 5.43E-14 | 2.95E-12 |
| PPP1R3C | 10.29407848 | 2.604184627 | 5.63E-08 | 8.14E-07 |
| GLIS1 | 10.28436893 | 2.285599208 | 2.80E-08 | 4.37E-07 |

| | | | | |
|---|---|---|---|---|
| RTN4R | 10.28122629 | 2.283874902 | 6.38E-12 | 2.31E-10 |
| ARHGEF16 | 10.28028842 | 2.848350293 | 5.55E-08 | 8.04E-07 |
| LOC119876427 | 10.28019733 | 2.672285777 | 2.49E-10 | 6.21E-09 |
| LOC119864784 | 10.27900483 | 2.320441711 | 5.31E-11 | 1.54E-09 |
| HOXD3 | 10.2646045 | 2.244500217 | 7.71E-15 | 4.86E-13 |
| APOE | 10.26422391 | 7.824263046 | 2.75E-20 | 5.14E-18 |
| LOC119866370 | 10.26377738 | 2.093344233 | 2.65E-07 | 3.19E-06 |
| LOC119866516 | 10.26246173 | 5.340274841 | 1.24E-22 | 3.20E-20 |
| LOC119867328 | 10.2601113 | 1.773130757 | 3.61E-09 | 6.87E-08 |
| LOC119877926 | 10.25500308 | 3.096624813 | 7.32E-16 | 5.68E-14 |
| NETO1 | 10.25317444 | 1.834448103 | 0.000443909 | 0.002036587 |
| PRSS23 | 10.25253501 | 7.105137701 | 6.34E-14 | 3.37E-12 |
| KIF6 | 10.25243848 | 1.85990253 | 4.83E-09 | 8.92E-08 |
| CDH6 | 10.24519269 | 2.831888164 | 7.25E-08 | 1.02E-06 |
| LOC102155734 | 10.23688577 | 1.812230168 | 6.51E-09 | 1.17E-07 |
| LOC111091339 | 10.220537 | 2.285883126 | 2.97E-15 | 2.03E-13 |
| SH3TC2 | 10.21054428 | 5.257230564 | 1.36E-11 | 4.45E-10 |
| LOC102155974 | 10.20542091 | 2.117778128 | 2.83E-11 | 8.64E-10 |
| LOC111097169 | 10.19698878 | 1.625971526 | 4.28E-06 | 3.64E-05 |
| FAH | 10.18711887 | 2.87719167 | 2.99E-14 | 1.69E-12 |
| LOC102153706 | 10.18196293 | 1.991746369 | 1.24E-11 | 4.12E-10 |
| SYT9 | 10.1695455 | 2.214797692 | 9.34E-10 | 2.07E-08 |
| ARSJ | 10.16104702 | 1.816260253 | 2.42E-08 | 3.82E-07 |
| ZNF662 | 10.15836074 | 2.712446005 | 7.19E-11 | 2.02E-09 |
| SOX17 | 10.13954693 | 3.524635499 | 4.55E-09 | 8.45E-08 |
| TMEM74 | 10.13920301 | 2.482138176 | 6.84E-08 | 9.71E-07 |
| GALNT18 | 10.139143 | 4.21280582 | 2.30E-22 | 5.53E-20 |
| C1QTNF1 | 10.13561278 | 7.229258076 | 7.14E-14 | 3.75E-12 |
| ANGPTL8 | 10.13517475 | 2.349523472 | 1.82E-13 | 8.89E-12 |
| LOC106559430 | 10.13441857 | 2.114048019 | 3.42E-07 | 4.01E-06 |
| MAP9 | 10.13407906 | 3.12235905 | 1.76E-16 | 1.52E-14 |
| SH3D19 | 10.12867397 | 6.973485987 | 6.76E-21 | 1.41E-18 |
| LOC106558485 | 10.12767381 | 2.333625549 | 6.39E-08 | 9.13E-07 |
| LOC111096062 | 10.12435088 | 2.151190979 | 4.70E-10 | 1.11E-08 |
| ADAMTSL2 | 10.1197525 | 2.57463659 | 4.50E-14 | 2.47E-12 |
| LOC119871500 | 10.10885848 | 1.464846036 | 3.28E-05 | 0.00021731 |
| LOC119868075 | 10.10638639 | 2.131200932 | 8.33E-10 | 1.87E-08 |
| TNNI3 | 10.10362431 | 4.739390491 | 2.26E-06 | 2.09E-05 |
| LOC119877537 | 10.103233 | 1.651016904 | 0.000274413 | 0.001345556 |

| | | | | |
|---|---|---|---|---|
| LOC119877202 | 10.09982468 | 7.646916635 | 0.000105076 | 0.000597875 |
| LOC111091594 | 10.09884377 | 3.16111792 | 2.87E-11 | 8.72E-10 |
| LOC111093430 | 10.09811106 | 2.00428287 | 1.54E-08 | 2.54E-07 |
| BMP7 | 10.09385618 | 2.224422543 | 1.18E-06 | 1.18E-05 |
| LAMA1 | 10.0865851 | 1.797032958 | 7.80E-07 | 8.22E-06 |
| GLRB | 10.07925753 | 3.451549712 | 3.65E-12 | 1.40E-10 |
| SYT14 | 10.0736381 | 1.442335373 | 5.33E-05 | 0.000332179 |
| LOC119877933 | 10.06755839 | 1.426713866 | 7.77E-05 | 0.000460521 |
| LOC102157275 | 10.06738335 | 1.670869313 | 1.64E-08 | 2.68E-07 |
| ARMCX1 | 10.06685623 | 4.104144911 | 9.44E-22 | 2.15E-19 |
| DSC2 | 10.06660279 | 1.78424467 | 0.000708707 | 0.003043352 |
| STK32B | 10.06451449 | 1.408455004 | 3.05E-08 | 4.69E-07 |
| LOC119867475 | 10.06449488 | 1.573030034 | 1.98E-06 | 1.86E-05 |
| EPB41L4B | 10.06054213 | 4.798424318 | 7.26E-15 | 4.61E-13 |
| LOC111090590 | 10.05709977 | 2.355256623 | 2.67E-11 | 8.23E-10 |
| LOC119873616 | 10.05166809 | 1.45672867 | 8.84E-07 | 9.16E-06 |
| XKR4 | 10.05094053 | 5.787915356 | 5.35E-06 | 4.40E-05 |
| BTNL9 | 10.04811647 | 3.966047926 | 8.28E-12 | 2.90E-10 |
| LOC111092166 | 10.04289057 | 1.844412294 | 4.54E-08 | 6.73E-07 |
| LOC111093754 | 10.0370722 | 1.868176207 | 4.04E-11 | 1.19E-09 |
| KLF15 | 10.03538844 | 3.119064053 | 4.17E-10 | 9.99E-09 |
| PPP1R1A | 10.03141467 | 1.623406751 | 2.95E-08 | 4.56E-07 |
| PARVA | 10.01988802 | 6.993882056 | 2.35E-25 | 1.04E-22 |
| LOC106557781 | 10.01520789 | 2.007390051 | 4.70E-11 | 1.37E-09 |
| ADAMTS19 | 10.01243643 | 2.598853171 | 3.29E-05 | 0.000217969 |
| EYA1 | 10.0120253 | 1.75518455 | 6.82E-06 | 5.44E-05 |
| LRCH2 | 10.01028229 | 2.192864816 | 4.70E-08 | 6.94E-07 |
| ABCC9 | 10.01011661 | 5.638237554 | 2.37E-09 | 4.75E-08 |
| HOXA7 | 10.00969823 | 2.277475384 | 1.75E-12 | 7.08E-11 |
| RASD2 | 10.00682855 | 1.584356468 | 1.60E-09 | 3.34E-08 |
| FBXO2 | 9.980132453 | 2.443271571 | 8.55E-09 | 1.50E-07 |
| CCL19 | 9.978142031 | 1.343455915 | 1.14E-06 | 1.16E-05 |
| ROBO3 | 9.977045866 | 2.288650391 | 9.02E-11 | 2.48E-09 |
| BNC2 | 9.975189067 | 7.279176197 | 2.46E-19 | 3.57E-17 |
| SLC39A8 | 9.974600435 | 2.286021364 | 9.26E-08 | 1.26E-06 |
| FOXL1 | 9.97065159 | 3.020078128 | 1.80E-10 | 4.66E-09 |
| FIGNL2 | 9.96886292 | 1.887515023 | 1.02E-12 | 4.34E-11 |
| LOC102151259 | 9.964115465 | 2.00737838 | 1.39E-07 | 1.81E-06 |
| MAGEL2 | 9.964057059 | 1.432266277 | 3.22E-05 | 0.00021396 |

| | | | | |
|---|---|---|---|---|
| SLC26A10 | 9.959893053 | 2.687708531 | 1.55E-06 | 1.50E-05 |
| LOC111095132 | 9.9591474 | 2.132290211 | 7.65E-12 | 2.70E-10 |
| TPD52L1 | 9.955709012 | 1.867142231 | 2.45E-09 | 4.90E-08 |
| DRAXIN | 9.948227026 | 1.323943663 | 2.59E-05 | 0.000177054 |
| HAP1 | 9.942568277 | 1.604936219 | 2.92E-09 | 5.72E-08 |
| LOC119866867 | 9.942170601 | 1.436923555 | 3.83E-09 | 7.25E-08 |
| LOC483954 | 9.934782131 | 1.264410753 | 5.52E-07 | 6.10E-06 |
| LOC111095621 | 9.9331564 | 1.712960599 | 8.24E-06 | 6.43E-05 |
| SOHLH2 | 9.931451294 | 1.312238493 | 8.03E-06 | 6.29E-05 |
| LOC119866909 | 9.925273472 | 1.894497703 | 1.38E-13 | 6.92E-12 |
| COBL | 9.924744945 | 2.611597145 | 8.49E-05 | 0.000497291 |
| KCNMB1 | 9.920058237 | 2.07356419 | 1.50E-06 | 1.47E-05 |
| LOC111097472 | 9.919638592 | 1.736957234 | 1.72E-11 | 5.54E-10 |
| VAT1L | 9.912453904 | 3.101160175 | 3.72E-08 | 5.62E-07 |
| LOC119872545 | 9.906934748 | 1.369327515 | 8.50E-09 | 1.49E-07 |
| LOC111091293 | 9.904886801 | 1.982109317 | 6.37E-13 | 2.83E-11 |
| LOC119878346 | 9.904579524 | 11.77314444 | 0.001186188 | 0.004742449 |
| CCDC80 | 9.899639147 | 7.524440186 | 3.81E-11 | 1.13E-09 |
| FAM110D | 9.899435729 | 2.922602827 | 3.46E-07 | 4.05E-06 |
| WDR86 | 9.894476764 | 1.826667419 | 1.60E-06 | 1.55E-05 |
| PDE1A | 9.893957055 | 2.628912726 | 3.05E-11 | 9.24E-10 |
| NAT8L | 9.887758883 | 3.088471351 | 1.75E-09 | 3.60E-08 |
| LOC491382 | 9.87949091 | 2.10698699 | 5.08E-14 | 2.76E-12 |
| DACT2 | 9.879193755 | 2.134865408 | 4.27E-09 | 8.01E-08 |
| TINAGL1 | 9.870462881 | 5.633109336 | 9.93E-12 | 3.39E-10 |
| LOC119877315 | 9.865035004 | 2.205562184 | 8.26E-14 | 4.28E-12 |
| NTRK3 | 9.864329197 | 5.899758102 | 1.63E-17 | 1.63E-15 |
| LOC119868664 | 9.863481567 | 1.743120528 | 1.89E-06 | 1.79E-05 |
| CLTCL1 | 9.855296834 | 1.642574222 | 3.18E-11 | 9.62E-10 |
| ADAMTSL3 | 9.850029926 | 5.464583213 | 2.32E-20 | 4.41E-18 |
| LOC106560206 | 9.847575924 | 1.211422207 | 7.42E-07 | 7.89E-06 |
| LOC102155082 | 9.842837342 | 2.08266333 | 2.37E-09 | 4.76E-08 |
| TMEM119 | 9.839291382 | 6.208518609 | 7.33E-24 | 2.24E-21 |
| TFAP2A | 9.832969432 | 1.750280055 | 8.96E-06 | 6.92E-05 |
| SIX1 | 9.831520287 | 1.875574481 | 4.14E-06 | 3.53E-05 |
| EPHB1 | 9.830609228 | 2.62089386 | 6.09E-11 | 1.72E-09 |
| NECAB1 | 9.829480256 | 2.103391404 | 4.07E-09 | 7.67E-08 |
| SAT2 | 9.821310394 | 2.137645573 | 8.42E-12 | 2.94E-10 |
| LOC119872629 | 9.814233323 | 1.799193622 | 1.73E-07 | 2.19E-06 |

| | | | | |
|---|---|---|---|---|
| LOC102153258 | 9.806433158 | 1.479884758 | 2.27E-09 | 4.59E-08 |
| AGTR1 | 9.803911398 | 2.470989911 | 1.50E-07 | 1.94E-06 |
| SOX8 | 9.803437005 | 2.80268555 | 2.01E-05 | 0.000141711 |
| LOC119871632 | 9.802843459 | 1.197873294 | 3.81E-07 | 4.41E-06 |
| GSTT2B | 9.800922767 | 1.804149814 | 1.11E-09 | 2.43E-08 |
| LRP1B | 9.798098524 | 3.844974481 | 6.50E-07 | 7.01E-06 |
| ZNF532 | 9.795781747 | 7.691441285 | 4.12E-21 | 8.83E-19 |
| LOC106560204 | 9.790516275 | 1.765517652 | 2.27E-10 | 5.75E-09 |
| SNCA | 9.789762332 | 4.757226326 | 0.000107625 | 0.00060966 |
| LOC119868046 | 9.779849386 | 2.860177984 | 1.59E-12 | 6.48E-11 |
| LOC119876140 | 9.779672238 | 1.955331267 | 1.17E-14 | 7.08E-13 |
| LOC119867861 | 9.768760488 | 1.761377614 | 1.03E-08 | 1.77E-07 |
| MAFA | 9.768402105 | 1.410512704 | 4.79E-06 | 4.01E-05 |
| CDC42EP1 | 9.768056621 | 5.624805505 | 3.72E-25 | 1.55E-22 |
| LOC119875969 | 9.767056054 | 1.199118111 | 2.33E-08 | 3.68E-07 |
| LOC119876216 | 9.766850012 | 1.862947592 | 1.81E-10 | 4.69E-09 |
| PODN | 9.766282862 | 7.547905957 | 4.03E-13 | 1.86E-11 |
| MIR8805 | 9.762917546 | 1.288121343 | 5.73E-06 | 4.68E-05 |
| LOC119869701 | 9.760321544 | 1.576660776 | 1.10E-09 | 2.41E-08 |
| RSPO3 | 9.757401482 | 1.888832661 | 2.10E-05 | 0.000146769 |
| LRRC6 | 9.756221484 | 1.172454908 | 4.90E-10 | 1.15E-08 |
| COX4I2 | 9.753157368 | 1.67294651 | 9.63E-08 | 1.31E-06 |
| THSD8 | 9.74879043 | 2.075694001 | 9.62E-12 | 3.30E-10 |
| KCNK2 | 9.745525304 | 2.295630575 | 5.13E-06 | 4.24E-05 |
| RELN | 9.7446922 | 7.199228626 | 7.04E-07 | 7.52E-06 |
| LOC119870797 | 9.742941264 | 1.255302282 | 1.22E-07 | 1.62E-06 |
| PLD5 | 9.741493034 | 2.070078433 | 2.86E-05 | 0.000193092 |
| KIAA1549L | 9.739222075 | 4.1459918 | 2.61E-07 | 3.16E-06 |
| RBFOX1 | 9.730806487 | 1.613895676 | 2.79E-07 | 3.34E-06 |
| LOC119876733 | 9.728847045 | 1.680949217 | 7.92E-11 | 2.21E-09 |
| LVRN | 9.724821193 | 3.331430566 | 2.70E-11 | 8.31E-10 |
| LOC119873286 | 9.724118256 | 1.692045349 | 1.49E-07 | 1.92E-06 |
| ENPP6 | 9.722705521 | 3.461962352 | 0.000147253 | 0.000790148 |
| TM4SF1 | 9.711136031 | 6.744977505 | 2.02E-13 | 9.82E-12 |
| TBC1D8B | 9.696660436 | 2.05989525 | 2.95E-10 | 7.23E-09 |
| FGFR2 | 9.694734703 | 5.489215798 | 1.14E-14 | 6.93E-13 |
| MYO3A | 9.684107482 | 1.259575785 | 3.13E-09 | 6.08E-08 |
| ADGRA1 | 9.683161788 | 1.045436977 | 0.000550371 | 0.002450326 |
| CDKL4 | 9.676826716 | 1.083059427 | 3.91E-07 | 4.52E-06 |

| | | | | |
|---|---|---|---|---|
| PTPRT | 9.671800295 | 6.167886468 | 1.17E-06 | 1.18E-05 |
| LOC111093449 | 9.670988653 | 1.265832563 | 1.40E-07 | 1.83E-06 |
| LOC119872479 | 9.667763371 | 1.221732941 | 5.86E-07 | 6.42E-06 |
| PDE10A | 9.661527569 | 4.250946154 | 2.74E-13 | 1.30E-11 |
| LOC119868368 | 9.66054643 | 1.740202386 | 1.27E-11 | 4.20E-10 |
| SMOC2 | 9.653414808 | 6.706994031 | 1.95E-10 | 5.01E-09 |
| LOC102153818 | 9.652192137 | 1.205616069 | 4.67E-07 | 5.28E-06 |
| LOC111095570 | 9.645423764 | 1.292427555 | 3.38E-09 | 6.48E-08 |
| GSC | 9.643833057 | 2.170642309 | 9.60E-05 | 0.000551793 |
| LOC102153763 | 9.638398509 | 2.35834912 | 1.15E-10 | 3.08E-09 |
| ERC2 | 9.633378124 | 1.169485123 | 3.98E-08 | 5.96E-07 |
| COL4A5 | 9.629427719 | 3.573259248 | 5.17E-06 | 4.27E-05 |
| LOC487175 | 9.627743872 | 1.717882711 | 1.52E-09 | 3.18E-08 |
| LOC111095204 | 9.625758451 | 2.752705539 | 1.18E-09 | 2.57E-08 |
| LOC106559789 | 9.625649205 | 1.01391344 | 0.000170027 | 0.000893549 |
| COLEC12 | 9.620711659 | 6.910589081 | 3.24E-14 | 1.82E-12 |
| LOC111092259 | 9.61626709 | 1.687421914 | 9.86E-05 | 0.000564945 |
| KCNK3 | 9.615734848 | 1.87152641 | 7.35E-05 | 0.000439243 |
| PTPRZ1 | 9.615638555 | 5.26772392 | 2.57E-05 | 0.0001757 |
| KRT24 | 9.615266556 | 1.050257612 | 6.13E-05 | 0.000374917 |
| LOC119873027 | 9.612697936 | 1.144815149 | 2.04E-06 | 1.91E-05 |
| ST6GALNAC3 | 9.604961098 | 1.475754178 | 9.48E-07 | 9.77E-06 |
| LOC119868032 | 9.602871477 | 1.625896019 | 3.32E-08 | 5.07E-07 |
| FBLN1 | 9.60127468 | 8.836607267 | 5.83E-10 | 1.35E-08 |
| TRPM4 | 9.583781662 | 5.471807187 | 7.45E-13 | 3.27E-11 |
| MPDZ | 9.581085697 | 6.640638989 | 1.59E-13 | 7.89E-12 |
| VSTM2L | 9.575855027 | 0.981197092 | 1.61E-07 | 2.05E-06 |
| PABPC4L | 9.575740653 | 1.502428776 | 3.66E-09 | 6.97E-08 |
| LOC106559945 | 9.563535553 | 1.260792637 | 1.68E-05 | 0.000120944 |
| PCSK1N | 9.56222193 | 2.344678849 | 2.12E-07 | 2.62E-06 |
| CPXM2 | 9.560375857 | 5.652967877 | 1.61E-09 | 3.34E-08 |
| SERPINH1 | 9.559031411 | 8.22376312 | 2.64E-28 | 2.05E-25 |
| LOC102155920 | 9.55264863 | 0.974913294 | 1.24E-07 | 1.64E-06 |
| LOC111096010 | 9.552595582 | 2.142764241 | 2.66E-10 | 6.59E-09 |
| LOC119878309 | 9.546881995 | 12.35022731 | 0.001289791 | 0.005084329 |
| LOC102156217 | 9.546683285 | 1.184951684 | 4.88E-06 | 4.06E-05 |
| LOC100688410 | 9.541971616 | 1.480059079 | 1.77E-09 | 3.64E-08 |
| SEC16B | 9.538623882 | 1.206820783 | 1.85E-08 | 2.99E-07 |
| LOC102154344 | 9.538372357 | 1.608484145 | 2.32E-08 | 3.67E-07 |

| | | | | |
|---|---|---|---|---|
| LOC119877048 | 9.536733845 | 1.395689281 | 5.84E-07 | 6.41E-06 |
| C1QB | 9.533753794 | 6.54354049 | 6.71E-10 | 1.54E-08 |
| LOC111095760 | 9.525775691 | 0.868932104 | 0.000166848 | 0.000878976 |
| LOC102152903 | 9.524064797 | 1.245038484 | 2.47E-11 | 7.65E-10 |
| LOC119873258 | 9.516417564 | 1.228902318 | 8.95E-08 | 1.23E-06 |
| PRCD | 9.513868464 | 1.200104624 | 1.36E-06 | 1.34E-05 |
| TNFSF15 | 9.513491105 | 1.231744748 | 1.56E-06 | 1.51E-05 |
| CLEC3B | 9.509300805 | 6.71390929 | 4.17E-10 | 9.99E-09 |
| ARSI | 9.505335864 | 1.400406118 | 2.30E-07 | 2.82E-06 |
| LOC102151805 | 9.50435268 | 3.921133022 | 6.47E-13 | 2.86E-11 |
| LGI2 | 9.501207212 | 4.973534171 | 3.63E-07 | 4.22E-06 |
| FNDC4 | 9.49836924 | 1.879923408 | 8.46E-12 | 2.95E-10 |
| KIAA1755 | 9.494496564 | 4.749839637 | 1.53E-16 | 1.33E-14 |
| PGBD5 | 9.491558315 | 1.206994629 | 7.45E-06 | 5.88E-05 |
| EVC | 9.488944795 | 5.553078013 | 2.53E-21 | 5.53E-19 |
| SNAP25 | 9.484122969 | 0.96910602 | 6.47E-06 | 5.19E-05 |
| LOC102153035 | 9.463438057 | 1.129010609 | 2.23E-08 | 3.54E-07 |
| KCNA6 | 9.459906361 | 3.439006765 | 1.39E-07 | 1.81E-06 |
| PAX1 | 9.458181346 | 1.473402403 | 0.000352381 | 0.001671879 |
| LDLRAD2 | 9.456697284 | 1.894521862 | 1.97E-10 | 5.05E-09 |
| CAVIN1 | 9.453517448 | 8.938950618 | 2.53E-24 | 8.43E-22 |
| FAP | 9.451236151 | 6.344322508 | 1.02E-08 | 1.75E-07 |
| EDNRA | 9.451230368 | 3.498905528 | 2.89E-14 | 1.65E-12 |
| RIMKLB | 9.450392995 | 2.597701874 | 3.68E-05 | 0.000240136 |
| GPC6 | 9.44906695 | 5.208763066 | 4.70E-06 | 3.95E-05 |
| SERPINB5 | 9.43611463 | 2.162230579 | 2.06E-08 | 3.29E-07 |
| PHF21B | 9.433988001 | 3.669704232 | 6.47E-09 | 1.17E-07 |
| LOC111092368 | 9.432355053 | 1.038229029 | 1.93E-08 | 3.10E-07 |
| LOC111095283 | 9.43062751 | 1.150704675 | 2.31E-10 | 5.83E-09 |
| LOC102154028 | 9.429062631 | 1.361006705 | 6.23E-06 | 5.02E-05 |
| ATP1A3 | 9.427404878 | 4.192698543 | 3.65E-06 | 3.17E-05 |
| LOC111091892 | 9.42684414 | 1.505117834 | 3.02E-08 | 4.66E-07 |
| LOC102153883 | 9.426071924 | 0.869956784 | 0.000545887 | 0.002433361 |
| LOC119870498 | 9.423894166 | 5.088193917 | 5.71E-20 | 9.83E-18 |
| SYBU | 9.419179309 | 5.36317804 | 4.72E-20 | 8.31E-18 |
| MYOC | 9.418555797 | 6.03375759 | 1.41E-07 | 1.84E-06 |
| LOC111096339 | 9.41692722 | 0.932052459 | 5.71E-07 | 6.29E-06 |
| LOC119865981 | 9.415753888 | 1.740964039 | 2.58E-10 | 6.41E-09 |
| LOC119876424 | 9.415374732 | 1.270424667 | 8.41E-05 | 0.000492541 |

| | | | | |
|---|---|---|---|---|
| BMPER | 9.415182497 | 3.670842763 | 6.18E-12 | 2.26E-10 |
| LOC119873282 | 9.407795345 | 1.046277419 | 6.75E-08 | 9.59E-07 |
| LOC119868367 | 9.405735863 | 1.308772348 | 8.01E-10 | 1.80E-08 |
| LMX1B | 9.39758028 | 0.797233279 | 0.000112895 | 0.000634868 |
| NCS1 | 9.397117524 | 3.919675171 | 1.88E-19 | 2.79E-17 |
| NEFH | 9.39178355 | 4.638252711 | 4.53E-05 | 0.00028809 |
| LOC102156352 | 9.389819441 | 1.681903966 | 1.18E-09 | 2.57E-08 |
| COL5A2 | 9.385603588 | 9.232281468 | 3.00E-18 | 3.63E-16 |
| DCHS2 | 9.383851441 | 0.723751487 | 1.51E-07 | 1.94E-06 |
| NAALADL2 | 9.383279176 | 4.419962096 | 7.06E-14 | 3.71E-12 |
| DNM3 | 9.380060719 | 4.836066722 | 1.03E-23 | 3.09E-21 |
| IGFBP4 | 9.364620545 | 8.257402433 | 2.14E-11 | 6.75E-10 |
| LOC111092286 | 9.362960002 | 1.382052734 | 1.11E-05 | 8.35E-05 |
| SOX2 | 9.361865638 | 3.431266124 | 4.66E-06 | 3.91E-05 |
| ADAMTS18 | 9.350849494 | 2.421252287 | 9.54E-10 | 2.11E-08 |
| APBA2 | 9.350591071 | 0.901272753 | 2.86E-06 | 2.57E-05 |
| ERICH2 | 9.349701265 | 0.68851489 | 3.25E-07 | 3.84E-06 |
| LOC119866889 | 9.348851855 | 1.37930271 | 5.02E-06 | 4.17E-05 |
| WWC1 | 9.343712733 | 2.096461368 | 3.11E-05 | 0.000207571 |
| LOC106558789 | 9.340297312 | 1.087047207 | 1.26E-08 | 2.11E-07 |
| NRN1 | 9.335658578 | 4.194815514 | 2.72E-05 | 0.000184301 |
| MSI1 | 9.335570322 | 0.954757794 | 2.39E-05 | 0.000165127 |
| MYBPH | 9.333372816 | 1.223258559 | 7.76E-08 | 1.08E-06 |
| NLGN1 | 9.329264198 | 2.811501125 | 1.66E-05 | 0.000119314 |
| LOC111097583 | 9.328430417 | 1.390218645 | 2.23E-09 | 4.54E-08 |
| CSPG4 | 9.328275611 | 7.653189908 | 2.89E-13 | 1.36E-11 |
| SOD3 | 9.322979549 | 6.934381831 | 4.81E-08 | 7.09E-07 |
| HOXB5 | 9.317891783 | 1.110002005 | 2.13E-06 | 1.99E-05 |
| DMPK | 9.317177152 | 4.970042783 | 4.18E-16 | 3.35E-14 |
| ONECUT2 | 9.313235639 | 0.699333234 | 1.17E-05 | 8.76E-05 |
| LOC111093250 | 9.312754757 | 0.792018447 | 4.77E-06 | 4.00E-05 |
| SPATA20 | 9.309956289 | 0.76625161 | 3.34E-07 | 3.93E-06 |
| WNT6 | 9.30906281 | 3.057337312 | 0.000148507 | 0.000795499 |
| ELFN1 | 9.304894633 | 1.646709293 | 2.55E-09 | 5.05E-08 |
| CACNA1G | 9.303307103 | 4.410125868 | 5.43E-12 | 2.01E-10 |
| LCNL1 | 9.301003448 | 0.924690775 | 1.25E-06 | 1.25E-05 |
| LOC111098870 | 9.297251018 | 1.049469388 | 1.83E-10 | 4.73E-09 |
| LOC111095627 | 9.295779086 | 2.083019544 | 2.34E-10 | 5.89E-09 |
| TMEM54 | 9.291684418 | 1.856469116 | 2.84E-11 | 8.64E-10 |

| | | | | |
|---|---|---|---|---|
| KCNG1 | 9.287627031 | 1.275109122 | 2.84E-06 | 2.56E-05 |
| GRIK4 | 9.287064803 | 1.369369902 | 4.58E-06 | 3.86E-05 |
| LOC119870339 | 9.286125671 | 0.821567355 | 4.02E-08 | 6.01E-07 |
| NDP | 9.282661984 | 1.496989487 | 0.000130922 | 0.000717078 |
| LOC119878294 | 9.280545244 | 10.72389025 | 0.000113334 | 0.000636672 |
| HIF3A | 9.276233527 | 5.693981554 | 2.25E-06 | 2.08E-05 |
| GPRIN2 | 9.273705116 | 0.726914823 | 1.23E-05 | 9.13E-05 |
| LOC111093370 | 9.2665797 | 0.656120306 | 0.000132282 | 0.000723429 |
| GRIA3 | 9.266366511 | 2.296196367 | 1.52E-07 | 1.96E-06 |
| LOC102151895 | 9.265939384 | 5.622801616 | 2.40E-10 | 6.03E-09 |
| SNCAIP | 9.260566492 | 3.771953614 | 1.33E-17 | 1.37E-15 |
| SERPINB13 | 9.259980641 | 0.653371159 | 8.48E-07 | 8.83E-06 |
| SLITRK6 | 9.256615052 | 1.246220205 | 3.02E-05 | 0.000202441 |
| ADAMDEC1 | 9.255564117 | 2.007094898 | 5.84E-06 | 4.76E-05 |
| LOC102152945 | 9.254714408 | 0.721879587 | 1.52E-06 | 1.48E-05 |
| CFAP300 | 9.253255663 | 1.749469263 | 6.88E-11 | 1.94E-09 |
| AKAP12 | 9.252486773 | 9.237439413 | 1.18E-11 | 3.95E-10 |
| BRME1 | 9.251653204 | 0.885346706 | 7.81E-09 | 1.38E-07 |
| PPP2R2B | 9.251051145 | 3.517301176 | 4.79E-07 | 5.40E-06 |
| LOC119865865 | 9.250677038 | 0.691414683 | 7.35E-08 | 1.03E-06 |
| LOC111094189 | 9.242702306 | 1.151224176 | 1.80E-09 | 3.69E-08 |
| MYOCD | 9.239027208 | 2.748024319 | 8.94E-08 | 1.23E-06 |
| WNT10A | 9.238031925 | 1.402689752 | 0.000595543 | 0.002621285 |
| PPFIA3 | 9.237745059 | 5.940046281 | 2.99E-07 | 3.57E-06 |
| LOC111092041 | 9.236812486 | 0.937941672 | 2.46E-09 | 4.91E-08 |
| A4GALT | 9.23468987 | 5.083028456 | 2.29E-27 | 1.51E-24 |
| LOC111096323 | 9.233918132 | 0.959856286 | 4.82E-07 | 5.42E-06 |
| RIBC2 | 9.231968591 | 0.579053986 | 1.17E-05 | 8.75E-05 |
| LOC119869534 | 9.231785879 | 1.744749196 | 1.30E-06 | 1.29E-05 |
| SLC1A7 | 9.230616537 | 3.100608586 | 1.37E-12 | 5.67E-11 |
| CYP2S1 | 9.219977129 | 0.799136454 | 4.13E-07 | 4.73E-06 |
| LOC487173 | 9.219531898 | 2.892170078 | 8.83E-20 | 1.44E-17 |
| GREM1 | 9.218073596 | 1.113247784 | 0.000395191 | 0.0018455 |
| LOC119865012 | 9.21695306 | 1.384959011 | 6.78E-11 | 1.91E-09 |
| LOC119865670 | 9.21544338 | 1.191980242 | 0.000245055 | 0.001221499 |
| HOXA6 | 9.214490404 | 1.611286016 | 7.44E-12 | 2.64E-10 |
| PLVAP | 9.211744257 | 6.844081072 | 1.05E-15 | 7.91E-14 |
| LOC106559420 | 9.206892901 | 0.831318718 | 1.06E-06 | 1.08E-05 |
| LOC119877529 | 9.205799865 | 1.393981717 | 9.50E-10 | 2.10E-08 |

| | | | | |
|---|---|---|---|---|
| ADGRB3 | 9.203764081 | 4.016068918 | 2.00E-07 | 2.49E-06 |
| OLFML2B | 9.197685153 | 8.244518873 | 1.21E-17 | 1.28E-15 |
| LOC111092370 | 9.192339738 | 0.765569799 | 1.52E-06 | 1.48E-05 |
| LOC119863925 | 9.191992654 | 2.155180712 | 6.24E-10 | 1.44E-08 |
| ADAMTS8 | 9.190137969 | 1.915735868 | 1.89E-08 | 3.04E-07 |
| SHBG | 9.18882682 | 1.405191381 | 1.39E-09 | 2.94E-08 |
| LOC119878422 | 9.187056144 | 12.67326731 | 8.80E-05 | 0.000512358 |
| LOC102152813 | 9.184820335 | 0.684892669 | 2.86E-05 | 0.000192756 |
| LOC111097118 | 9.183841937 | 0.858575015 | 4.73E-07 | 5.34E-06 |
| DPT | 9.182861265 | 5.494423656 | 1.56E-07 | 2.00E-06 |
| SEMA5B | 9.180664014 | 0.943035193 | 8.89E-08 | 1.22E-06 |
| C1QTNF7 | 9.174096138 | 1.811304297 | 5.23E-10 | 1.22E-08 |
| IFI6 | 9.170957723 | 1.31216397 | 1.11E-07 | 1.49E-06 |
| ATP6V1B1 | 9.161981254 | 0.570532163 | 1.91E-05 | 0.00013513 |
| SMIM9 | 9.1607151 | 2.44537441 | 4.19E-10 | 1.00E-08 |
| MPP3 | 9.151976934 | 1.670172066 | 4.67E-10 | 1.10E-08 |
| ACOT11 | 9.151337008 | 0.905975382 | 2.66E-07 | 3.20E-06 |
| LOC111090938_1 | 9.150922312 | 0.731506253 | 1.94E-05 | 0.000136991 |
| ASTN1 | 9.146432062 | 0.545523931 | 0.000327297 | 0.001567624 |
| LOC119877699 | 9.138738548 | 2.845259452 | 1.54E-13 | 7.65E-12 |
| MIRLET7B | 9.134950578 | 1.389778555 | 6.31E-08 | 9.03E-07 |
| WFIKKN2 | 9.134893591 | 1.875189437 | 4.49E-08 | 6.66E-07 |
| LOC119867461 | 9.13422269 | 0.763728282 | 2.90E-10 | 7.11E-09 |
| MPPED1 | 9.134130737 | 0.959790312 | 8.47E-08 | 1.17E-06 |
| FBXL7 | 9.132014972 | 6.419308761 | 8.29E-15 | 5.19E-13 |
| MFGE8 | 9.130990221 | 6.403690385 | 7.95E-16 | 6.13E-14 |
| LOC102154721 | 9.130637354 | 0.633745949 | 3.58E-08 | 5.43E-07 |
| CLDN34 | 9.129547098 | 1.666684807 | 2.20E-10 | 5.58E-09 |
| GSC2 | 9.129365073 | 0.625233507 | 3.15E-07 | 3.74E-06 |
| LOC111090975 | 9.122486664 | 1.11327821 | 1.20E-06 | 1.20E-05 |
| LOC106558158 | 9.119159468 | 1.289188567 | 3.83E-07 | 4.43E-06 |
| LOC111090872 | 9.1174637 | 1.0012425 | 3.14E-07 | 3.73E-06 |
| LHFPL6 | 9.112323244 | 5.726343947 | 3.02E-09 | 5.90E-08 |
| LOC102151543 | 9.112301042 | 0.536470323 | 4.82E-06 | 4.02E-05 |
| HOXA13 | 9.107416031 | 0.791012089 | 2.57E-05 | 0.0001757 |
| GAS6 | 9.107142631 | 5.553941781 | 9.97E-18 | 1.07E-15 |
| LY6H | 9.100071713 | 0.584060294 | 2.41E-05 | 0.00016629 |
| LOC102153450 | 9.097684958 | 0.914887604 | 1.17E-09 | 2.54E-08 |
| LOC119869718 | 9.096690943 | 1.410298824 | 5.48E-10 | 1.27E-08 |

| | | | | |
|---|---|---|---|---|
| STK32A | 9.092325817 | 1.280488037 | 4.19E-08 | 6.24E-07 |
| FAM43B | 9.088323674 | 1.288344018 | 4.74E-07 | 5.34E-06 |
| SLC6A7 | 9.063037506 | 0.902862901 | 1.93E-07 | 2.41E-06 |
| LOC119868498 | 9.054068812 | 2.664604167 | 8.65E-09 | 1.52E-07 |
| GRM7 | 9.053987822 | 0.957646021 | 4.82E-06 | 4.03E-05 |
| LOC119867066 | 9.051784122 | 0.573759145 | 3.99E-07 | 4.61E-06 |
| HS3ST5 | 9.049456431 | 1.17610131 | 8.35E-06 | 6.50E-05 |
| C6 | 9.046163805 | 2.189996264 | 9.75E-07 | 1.00E-05 |
| LOC111097347 | 9.045811424 | 0.741587982 | 8.27E-07 | 8.65E-06 |
| PTN | 9.044703176 | 2.178840714 | 3.00E-05 | 0.000201142 |
| FAM124A | 9.035412806 | 0.807446507 | 9.20E-06 | 7.08E-05 |
| KLK4 | 9.035334878 | 0.431013728 | 0.000144599 | 0.000778607 |
| OLFML2A | 9.034288124 | 6.902886595 | 2.27E-09 | 4.59E-08 |
| LOC111097167 | 9.031831682 | 0.489141197 | 3.11E-05 | 0.000207571 |
| ADAMTS9 | 9.026783199 | 6.08019245 | 2.45E-09 | 4.90E-08 |
| DOK7 | 9.018286669 | 0.823489178 | 1.42E-07 | 1.85E-06 |
| LOC111090578 | 9.012681165 | 0.315014898 | 0.000105553 | 0.000599962 |
| LOC111096667 | 9.011975287 | 0.419988757 | 0.000284255 | 0.00138785 |
| LOC119871850 | 9.0056592 | 1.705800138 | 1.95E-09 | 3.99E-08 |
| LOC102152485 | 9.004987142 | 0.786400003 | 1.21E-08 | 2.04E-07 |
| LOC119877449 | 8.998461772 | 1.993810088 | 5.84E-06 | 4.76E-05 |
| LOC119873138 | 8.998070538 | 0.61476517 | 4.19E-07 | 4.79E-06 |
| SYDE2 | 8.989405785 | 2.894388803 | 3.52E-06 | 3.08E-05 |
| CELF6 | 8.989332969 | 0.409955332 | 1.37E-06 | 1.35E-05 |
| LOC119873675 | 8.988651968 | 6.898448329 | 1.15E-15 | 8.43E-14 |
| SLITRK2 | 8.985620492 | 0.735179964 | 0.000422352 | 0.001955071 |
| LOC119865912 | 8.98328946 | 1.170876732 | 4.91E-09 | 9.06E-08 |
| SSUH2 | 8.968855552 | 1.233715133 | 0.000512894 | 0.002306694 |
| LGI3 | 8.96598191 | 4.279076653 | 5.34E-08 | 7.77E-07 |
| LOC102156498 | 8.962464118 | 1.429095533 | 3.36E-09 | 6.45E-08 |
| C32H4orf36 | 8.958705714 | 1.262960879 | 5.79E-11 | 1.66E-09 |
| LOC102155549 | 8.956796086 | 0.978484677 | 1.50E-11 | 4.88E-10 |
| THBD | 8.95414906 | 6.461925072 | 2.28E-10 | 5.75E-09 |
| RNF133 | 8.954111348 | 0.846633498 | 3.87E-08 | 5.81E-07 |
| LOC119868747 | 8.951844584 | 0.843217547 | 0.003750918 | 0.012580262 |
| SLIT3 | 8.950431638 | 9.356544436 | 4.50E-17 | 4.21E-15 |
| LOC106559446 | 8.948015677 | 0.986212154 | 3.80E-10 | 9.17E-09 |
| HSPB2 | 8.944095509 | 1.800795611 | 5.39E-11 | 1.56E-09 |
| CLCA2 | 8.941198378 | 5.414481936 | 4.00E-06 | 3.43E-05 |

| | | | | |
|---|---|---|---|---|
| LOC119871794 | 8.939663845 | 0.817451426 | 3.69E-07 | 4.29E-06 |
| LURAP1L | 8.938027775 | 0.754781747 | 7.35E-08 | 1.03E-06 |
| LOC119865668 | 8.930693793 | 1.528038518 | 0.000258119 | 0.001276341 |
| LOC119876425 | 8.928450784 | 0.666301974 | 6.23E-07 | 6.77E-06 |
| LOC111090635 | 8.928184351 | 1.395579631 | 8.08E-07 | 8.49E-06 |
| LOC106558208 | 8.927401015 | 2.285891036 | 4.52E-06 | 3.82E-05 |
| LOC119864411 | 8.924655549 | 0.672683592 | 2.04E-06 | 1.92E-05 |
| LOC102151142 | 8.914553725 | 0.828322808 | 8.63E-12 | 2.99E-10 |
| LOC111092445 | 8.912999569 | 0.551321517 | 0.000132534 | 0.000724441 |
| FBN3 | 8.911498996 | 4.599352944 | 8.44E-06 | 6.57E-05 |
| USH1C | 8.908930543 | 0.388398555 | 3.45E-07 | 4.03E-06 |
| PPP1R14C | 8.907647575 | 2.344057382 | 3.61E-11 | 1.08E-09 |
| LOC119864652 | 8.905438911 | 1.167799232 | 1.26E-06 | 1.26E-05 |
| LOC111093881 | 8.90341962 | 0.84027146 | 3.59E-07 | 4.18E-06 |
| TMEM200C | 8.897048779 | 0.691714803 | 1.15E-05 | 8.62E-05 |
| LOC102156399 | 8.894003802 | 0.712554251 | 2.09E-08 | 3.33E-07 |
| NDST3 | 8.891556111 | 2.116153649 | 3.02E-07 | 3.60E-06 |
| LOC100855450 | 8.88799401 | 0.597158774 | 0.001123259 | 0.004524139 |
| ITIH5 | 8.887904184 | 7.599688885 | 1.11E-12 | 4.69E-11 |
| SPARCL1 | 8.88761256 | 7.216741797 | 1.65E-09 | 3.43E-08 |
| LOC106559500 | 8.882746337 | 0.449675035 | 2.16E-08 | 3.44E-07 |
| LOC119868428 | 8.881900939 | 0.585877011 | 4.68E-09 | 8.68E-08 |
| PRPH | 8.88079631 | 7.273916647 | 2.24E-08 | 3.55E-07 |
| LOC111095193 | 8.875533572 | 2.187353939 | 2.03E-07 | 2.52E-06 |
| ERBB4 | 8.874034497 | 2.260359368 | 6.47E-06 | 5.19E-05 |
| ADGRL3 | 8.871011545 | 6.55387471 | 4.25E-07 | 4.85E-06 |
| LOC119867047 | 8.870779078 | 6.401860057 | 1.41E-19 | 2.19E-17 |
| LOC119867820 | 8.862764522 | 4.752625822 | 0.000161257 | 0.000854295 |
| CCN1 | 8.857649057 | 6.866807777 | 5.86E-14 | 3.14E-12 |
| PLPPR5 | 8.851577242 | 0.502069962 | 7.77E-07 | 8.20E-06 |
| DPYSL4 | 8.84151801 | 0.317704364 | 0.000106899 | 0.000606498 |
| KRT25 | 8.840582299 | 0.485645917 | 0.000441656 | 0.002028398 |
| LOC111098058 | 8.838523397 | 0.709008966 | 4.94E-05 | 0.000310844 |
| PRSS35 | 8.838173434 | 4.834214192 | 1.07E-10 | 2.88E-09 |
| TEX26 | 8.83741684 | 0.289347788 | 1.94E-07 | 2.43E-06 |
| ROBO1 | 8.835942407 | 5.619795146 | 4.56E-13 | 2.09E-11 |
| SCN1A | 8.83415392 | 0.627029734 | 1.18E-06 | 1.19E-05 |
| KANK2 | 8.827709513 | 8.127997733 | 3.83E-20 | 6.86E-18 |
| LOC119868057 | 8.822006093 | 0.560041947 | 2.80E-07 | 3.36E-06 |

| | | | | |
|---|---|---|---|---|
| PACRG | 8.816096197 | 0.781648989 | 4.17E-08 | 6.22E-07 |
| POU2F3 | 8.813747394 | 0.395624058 | 8.42E-07 | 8.77E-06 |
| SLC6A17 | 8.806456896 | 4.274190748 | 2.59E-08 | 4.06E-07 |
| LOC111090581 | 8.8058103 | 0.680646862 | 5.57E-11 | 1.60E-09 |
| AMOT | 8.799086117 | 4.933042567 | 5.69E-15 | 3.70E-13 |
| LRRC4B | 8.792264956 | 2.677573503 | 3.92E-09 | 7.39E-08 |
| LOC119867862 | 8.78813026 | 5.817298355 | 6.67E-12 | 2.41E-10 |
| F3 | 8.784042448 | 5.79056969 | 1.64E-08 | 2.68E-07 |
| BVES | 8.783548419 | 2.090380788 | 2.05E-05 | 0.000143704 |
| LOC119876141 | 8.778565195 | 0.421186217 | 6.83E-09 | 1.23E-07 |
| LOC119877530 | 8.777822749 | 0.611141563 | 0.000129581 | 0.000711529 |
| CENPV | 8.77693402 | 0.930907964 | 1.47E-07 | 1.90E-06 |
| ID1 | 8.773542261 | 3.543243504 | 3.54E-09 | 6.76E-08 |
| LOC119865454 | 8.771102879 | 1.671362597 | 1.56E-07 | 2.00E-06 |
| WWTR1 | 8.768372612 | 8.696542023 | 1.18E-22 | 3.11E-20 |
| LOC111093010 | 8.767768773 | 0.315372273 | 2.52E-05 | 0.000173154 |
| FBLL1 | 8.766322851 | 0.229853598 | 3.57E-07 | 4.16E-06 |
| MIR218-2 | 8.762544184 | 0.179607235 | 0.000103813 | 0.000591778 |
| TNXB | 8.760694347 | 10.32659983 | 1.71E-11 | 5.52E-10 |
| NTN1 | 8.760218012 | 4.515405923 | 1.12E-13 | 5.67E-12 |
| GRB14 | 8.754731055 | 1.411789512 | 3.70E-05 | 0.000241266 |
| C15H1orf216 | 8.753055433 | 3.560057904 | 2.22E-16 | 1.86E-14 |
| NT5C1A | 8.748854907 | 0.921031053 | 8.74E-05 | 0.000509543 |
| LOC102153504 | 8.742662494 | 0.332678273 | 0.000129884 | 0.00071283 |
| CHRDL2 | 8.742488687 | 0.096778064 | 1.95E-06 | 1.84E-05 |
| C1QA | 8.73762405 | 5.59743825 | 3.43E-09 | 6.57E-08 |
| KCNA5 | 8.731432082 | 1.473664648 | 2.04E-07 | 2.53E-06 |
| LOC119877043 | 8.728122414 | 0.514497013 | 5.87E-09 | 1.07E-07 |
| LOC111093728 | 8.724424304 | 1.691818998 | 1.33E-06 | 1.32E-05 |
| LUZP2 | 8.723782297 | 0.233131843 | 8.54E-05 | 0.000499588 |
| LOC119867821 | 8.72152172 | 3.920351587 | 0.000580674 | 0.002564692 |
| LOC119868381 | 8.719105769 | 0.419218957 | 1.16E-07 | 1.54E-06 |
| LIN7A | 8.716848206 | 2.953520286 | 8.15E-07 | 8.55E-06 |
| LOC119872502 | 8.712965835 | 1.473348507 | 1.19E-07 | 1.58E-06 |
| CACNG7 | 8.712186654 | 3.56630528 | 4.55E-05 | 0.000289235 |
| HOXB2 | 8.710227552 | 2.786106009 | 4.40E-11 | 1.29E-09 |
| LOC111092681 | 8.710111717 | 0.634421743 | 3.03E-07 | 3.61E-06 |
| PLS3 | 8.703457387 | 6.541508115 | 7.50E-18 | 8.25E-16 |
| LOC106560210 | 8.700246421 | 0.118895364 | 3.30E-05 | 0.00021852 |

| | | | | |
|---|---|---|---|---|
| NTN3 | 8.697386535 | 0.424814691 | 7.21E-08 | 1.01E-06 |
| DIO3 | 8.697301401 | 0.245422848 | 0.000143728 | 0.000775071 |
| LOC607729 | 8.69270648 | 0.548641729 | 8.99E-09 | 1.57E-07 |
| SLC6A11 | 8.692282448 | 0.135315055 | 0.000328093 | 0.001570743 |
| FOXG1 | 8.689307696 | 0.151313191 | 0.000241513 | 0.001206063 |
| PTGIS | 8.685114329 | 5.626292671 | 5.12E-08 | 7.51E-07 |
| LOC111091379 | 8.683564426 | 0.661291404 | 1.86E-08 | 3.01E-07 |
| LOC119865399 | 8.680969524 | 0.355617812 | 7.12E-10 | 1.62E-08 |
| COL18A1 | 8.680674984 | 8.642773917 | 1.18E-09 | 2.57E-08 |
| LOC102155499 | 8.679776229 | 0.19105562 | 6.57E-05 | 0.000398037 |
| LOC119870786 | 8.679391993 | 0.491421067 | 1.73E-07 | 2.19E-06 |
| NXPH4 | 8.67738519 | 2.504106341 | 4.25E-11 | 1.25E-09 |
| IQCD | 8.673676644 | 0.232128612 | 8.95E-08 | 1.23E-06 |
| ASPA | 8.669076786 | 4.818323228 | 5.39E-09 | 9.89E-08 |
| LOC111091529 | 8.666648139 | 1.6664616 | 5.90E-08 | 8.48E-07 |
| LOC111096977 | 8.661681754 | 0.782175851 | 1.39E-05 | 0.000102019 |
| LOC102151333 | 8.659586076 | 1.634345111 | 2.57E-08 | 4.02E-07 |
| LOC106558903 | 8.658395343 | 0.528622048 | 4.18E-07 | 4.78E-06 |
| MT-III | 8.654988257 | 3.421664877 | 2.29E-05 | 0.000158767 |
| LOC119866157 | 8.652529073 | 1.579017773 | 9.51E-05 | 0.00054729 |
| LOC111093578 | 8.651986061 | 0.728731005 | 2.12E-06 | 1.98E-05 |
| LOC119877349 | 8.646246117 | 0.715322312 | 1.34E-06 | 1.32E-05 |
| FOXE3 | 8.640082695 | 0.232555082 | 5.93E-06 | 4.81E-05 |
| GLI3 | 8.632975902 | 7.63471537 | 1.46E-12 | 6.02E-11 |
| CILP2 | 8.632241258 | 3.789892378 | 1.05E-07 | 1.42E-06 |
| CADPS | 8.6293465 | 0.084550855 | 0.000573157 | 0.00253666 |
| FOXF2 | 8.623620243 | 0.26811563 | 1.45E-05 | 0.000105874 |
| FAM241B | 8.619911185 | 0.288947996 | 9.17E-08 | 1.25E-06 |
| LOC111091298 | 8.619223697 | 0.830043779 | 1.26E-07 | 1.67E-06 |
| PRSS48 | 8.615729697 | 2.254781439 | 5.19E-08 | 7.58E-07 |
| LOC111093425 | 8.609597694 | 0.831980233 | 9.71E-07 | 9.98E-06 |
| LOC100856518 | 8.609264411 | 1.641877446 | 1.30E-06 | 1.29E-05 |
| PRG4 | 8.608261958 | 10.43645688 | 2.45E-07 | 2.99E-06 |
| C2CD4B | 8.606245821 | 0.426976916 | 2.59E-07 | 3.14E-06 |
| LOC119872615 | 8.605840997 | 0.871569424 | 1.06E-06 | 1.08E-05 |
| LOC102155021 | 8.604742533 | 0.622400869 | 1.89E-07 | 2.37E-06 |
| LOC106559547 | 8.602385326 | 0.162631615 | 1.58E-05 | 0.000114048 |
| GPRC5B | 8.600914778 | 5.42676334 | 7.04E-25 | 2.50E-22 |
| LOC119871572 | 8.599857604 | 0.690284419 | 1.15E-05 | 8.67E-05 |

| | | | | |
|---|---|---|---|---|
| CNN3 | 8.599028899 | 8.365420303 | 1.30E-18 | 1.66E-16 |
| MIR99B | 8.596234069 | 0.669041011 | 2.46E-07 | 2.99E-06 |
| LOC477556 | 8.593628922 | 1.646051246 | 1.78E-07 | 2.24E-06 |
| MADCAM1 | 8.593224236 | 0.17810674 | 1.62E-06 | 1.57E-05 |
| LOC119873706 | 8.591774945 | 0.27148594 | 1.19E-07 | 1.58E-06 |
| GREB1 | 8.588990737 | 2.817122854 | 2.44E-06 | 2.25E-05 |
| LAMC2 | 8.588323794 | 2.724319287 | 3.80E-11 | 1.13E-09 |
| LOC111090132 | 8.588232376 | 0.788541915 | 1.30E-06 | 1.29E-05 |
| LOC102152193 | 8.587066577 | 0.621878138 | 1.23E-06 | 1.23E-05 |
| LOC111091542 | 8.585915124 | 1.786716596 | 2.53E-08 | 3.98E-07 |
| DIP2C | 8.583649948 | 6.442845478 | 5.08E-14 | 2.76E-12 |
| LOC102151391 | 8.583075673 | 1.356953832 | 0.000110769 | 0.000625346 |
| SLC26A5 | 8.581208666 | 0.998728961 | 3.76E-08 | 5.67E-07 |
| HS6ST2 | 8.576840884 | 2.718489056 | 7.78E-05 | 0.000461191 |
| EMP2 | 8.572169184 | 5.814867946 | 5.49E-19 | 7.48E-17 |
| NGF | 8.570587391 | 0.894176924 | 3.86E-05 | 0.000250311 |
| AKAP6 | 8.568270983 | 7.034560673 | 5.44E-06 | 4.46E-05 |
| BTBD16 | 8.567939822 | 0.504726334 | 1.65E-07 | 2.09E-06 |
| LOC111092054 | 8.565619207 | 1.407180429 | 9.18E-08 | 1.25E-06 |
| PHYHIPL | 8.56388699 | 1.601162765 | 1.73E-07 | 2.19E-06 |
| FAM151A | 8.56288454 | 0.418402853 | 2.61E-05 | 0.000177906 |
| CALB2 | 8.561323766 | 0.736715405 | 0.000168649 | 0.000887411 |
| LOC119868074 | 8.560720984 | 2.051136339 | 2.99E-09 | 5.83E-08 |
| TPPP2 | 8.556489477 | 0.955414729 | 6.92E-08 | 9.79E-07 |
| AMOTL2 | 8.555417394 | 6.814629674 | 4.06E-19 | 5.71E-17 |
| LOC119871191 | 8.554732486 | 2.079305806 | 6.25E-13 | 2.79E-11 |
| CPEB1 | 8.552436679 | 4.390761726 | 3.98E-11 | 1.18E-09 |
| TMEM45A | 8.548852094 | 1.679953036 | 5.51E-09 | 1.01E-07 |
| DRP2 | 8.548779364 | 6.513989974 | 0.000121121 | 0.000672917 |
| UGT8 | 8.542229447 | 4.997436869 | 0.000126467 | 0.000697788 |
| COL14A1 | 8.539865555 | 7.293808155 | 4.45E-08 | 6.61E-07 |
| LOC111091517 | 8.532769175 | 0.311983888 | 0.000153116 | 0.000815952 |
| SERPING1 | 8.531414906 | 7.371906201 | 2.77E-19 | 3.98E-17 |
| SLC13A5 | 8.530964556 | 0.605718734 | 0.000747041 | 0.003184623 |
| LOC482646 | 8.52222037 | 1.937769617 | 2.24E-09 | 4.54E-08 |
| TBX5 | 8.520894859 | 0.963827637 | 0.000440169 | 0.002022857 |
| LOC111092582 | 8.51912099 | 1.105209234 | 8.39E-07 | 8.76E-06 |
| LOC119867863 | 8.512725364 | 0.364721958 | 1.37E-06 | 1.35E-05 |
| LOC111095583 | 8.512109821 | 0.018243351 | 0.000395031 | 0.00184515 |

| | | | | |
|---|---|---|---|---|
| LOC111098644 | 8.510321857 | 1.45584036 | 5.48E-05 | 0.00034003 |
| DDIT4L | 8.506640664 | 3.034172215 | 5.40E-06 | 4.44E-05 |
| FAM110C | 8.504783466 | -0.056379636 | 0.000555927 | 0.002470496 |
| KCNK17 | 8.503195769 | 1.689341239 | 9.14E-06 | 7.04E-05 |
| LOC119876665 | 8.502713347 | 2.348347174 | 8.60E-05 | 0.000502292 |
| LOC119868517 | 8.500849623 | 0.648730681 | 4.86E-06 | 4.05E-05 |
| PITX1 | 8.499824351 | 0.887745679 | 0.001225713 | 0.004868928 |
| PID1 | 8.497352346 | 5.25142092 | 7.40E-17 | 6.63E-15 |
| THBS3 | 8.495034352 | 6.797206678 | 7.22E-13 | 3.17E-11 |
| LOC111092728 | 8.49430261 | 0.931381133 | 0.000827372 | 0.003477821 |
| PLP1 | 8.49125435 | 5.241151863 | 0.000412272 | 0.001915774 |
| RRAD | 8.49069009 | 2.403886115 | 9.09E-07 | 9.39E-06 |
| LOC119871513 | 8.488846286 | 0.064097653 | 2.42E-06 | 2.22E-05 |
| DDI1 | 8.488054352 | 0.981900597 | 5.19E-08 | 7.58E-07 |
| C29H8orf34 | 8.48054348 | 0.168914102 | 7.12E-06 | 5.65E-05 |
| LOC119876312 | 8.479161448 | 4.332623749 | 3.88E-18 | 4.54E-16 |
| CACNA1C | 8.476993441 | 5.854843847 | 1.40E-11 | 4.60E-10 |
| ADORA1 | 8.474583858 | 1.273969599 | 5.74E-06 | 4.68E-05 |
| DLX5 | 8.473736684 | 0.034364615 | 3.39E-05 | 0.000223463 |
| TAGLN3 | 8.471575672 | 1.389044948 | 3.71E-07 | 4.30E-06 |
| LOC119864738 | 8.471308132 | 1.12094184 | 1.49E-08 | 2.46E-07 |
| LOC106558791 | 8.468873666 | 0.83557342 | 3.02E-08 | 4.66E-07 |
| AQP4 | 8.468011903 | 4.310628541 | 1.55E-05 | 0.000112588 |
| LOC111098748 | 8.467414464 | 0.510563889 | 2.60E-07 | 3.15E-06 |
| DKK3 | 8.465978029 | 5.489645306 | 4.78E-13 | 2.18E-11 |
| LOC102154791 | 8.455747662 | 0.058069746 | 3.07E-06 | 2.74E-05 |
| LOC111093230 | 8.45471712 | 1.045536028 | 0.000143509 | 0.000774472 |
| LOC111090278 | 8.454427177 | 0.379829424 | 0.000561303 | 0.002492341 |
| GPR148 | 8.452602652 | 0.239947877 | 0.000608725 | 0.002671909 |
| PRIMA1 | 8.451961781 | 4.56006583 | 1.97E-08 | 3.17E-07 |
| DLGAP1 | 8.448798004 | 0.099736747 | 5.05E-07 | 5.63E-06 |
| LOC111091683 | 8.446079914 | -0.020570958 | 0.000780935 | 0.003306589 |
| LOC111093330 | 8.44494169 | 1.626947848 | 1.21E-07 | 1.60E-06 |
| ADGRD1 | 8.440422078 | 7.063236082 | 7.16E-12 | 2.56E-10 |
| LOC612254 | 8.435452079 | 5.476758272 | 1.14E-06 | 1.15E-05 |
| ATP1A4 | 8.432897438 | 5.429752419 | 4.10E-14 | 2.27E-12 |
| LOC119870434 | 8.423405001 | 3.428107322 | 8.09E-09 | 1.43E-07 |
| PITPNM3 | 8.419100826 | 0.622602309 | 1.16E-07 | 1.54E-06 |
| MYH14 | 8.418896305 | 6.361759217 | 4.50E-10 | 1.07E-08 |

| | | | | |
|---|---|---|---|---|
| INHA | 8.411722667 | 0.29275723 | 5.36E-09 | 9.84E-08 |
| LOC111093428 | 8.411246258 | 0.807856049 | 5.10E-06 | 4.23E-05 |
| GRID2 | 8.406427884 | 1.385979312 | 4.50E-05 | 0.000286256 |
| RERGL | 8.402413227 | 2.935053798 | 0.000294665 | 0.001433928 |
| LOC119870730 | 8.40223402 | 0.107269586 | 1.82E-05 | 0.000129605 |
| LOC119871928 | 8.394121974 | 0.834683546 | 1.17E-07 | 1.56E-06 |
| CDH18 | 8.392668679 | -0.090671308 | 2.45E-05 | 0.000168447 |
| HOXC5 | 8.392235139 | 3.51741174 | 2.98E-08 | 4.60E-07 |
| MAB21L4 | 8.391211615 | -0.057493182 | 4.60E-06 | 3.88E-05 |
| LOC111093583 | 8.388437765 | 1.185535812 | 7.14E-08 | 1.00E-06 |
| ST6GAL2 | 8.388302758 | 0.991441812 | 5.99E-06 | 4.86E-05 |
| EDNRB | 8.388093084 | 3.944827381 | 5.44E-11 | 1.57E-09 |
| CDH5 | 8.385546248 | 6.237542777 | 7.66E-18 | 8.38E-16 |
| LOC111098613 | 8.383809529 | 0.098659644 | 3.25E-06 | 2.87E-05 |
| LOC100683344 | 8.382428158 | 0.558273458 | 3.66E-06 | 3.17E-05 |
| C9H17orf50 | 8.38110163 | 0.939476943 | 6.12E-07 | 6.67E-06 |
| FAM131C | 8.379208329 | 0.633466664 | 1.84E-06 | 1.75E-05 |
| LOC111098746 | 8.377826646 | 0.174777239 | 7.16E-08 | 1.00E-06 |
| WNT11 | 8.374956688 | 1.809688775 | 5.14E-08 | 7.53E-07 |
| LOC111094485 | 8.373170338 | 3.453108544 | 2.98E-17 | 2.85E-15 |
| LOC119871239 | 8.371214356 | 1.009061527 | 6.40E-08 | 9.14E-07 |
| CGREF1 | 8.371004292 | 1.021568805 | 9.69E-07 | 9.96E-06 |
| PNPLA5 | 8.368830577 | -0.22719924 | 0.000551272 | 0.002452848 |
| GALNT8 | 8.368791737 | 1.188374156 | 6.02E-06 | 4.88E-05 |
| LOC102152082 | 8.360196529 | 0.005038685 | 6.48E-07 | 7.00E-06 |
| ITGBL1 | 8.359347181 | 5.240664331 | 1.12E-06 | 1.14E-05 |
| LOC111096412 | 8.35635241 | 0.07494599 | 6.47E-07 | 6.99E-06 |
| LOC119873792 | 8.353464178 | -0.032621295 | 4.46E-05 | 0.00028385 |
| OR6W1 | 8.353268932 | 0.038940291 | 1.98E-06 | 1.86E-05 |
| KCNT2 | 8.352374361 | 3.827298788 | 2.07E-11 | 6.55E-10 |
| TBX1 | 8.349727032 | 1.317787953 | 2.57E-06 | 2.35E-05 |
| SPAAR | 8.346737845 | 0.652191755 | 6.49E-10 | 1.49E-08 |
| LDB3 | 8.342726391 | 1.583951536 | 8.02E-06 | 6.29E-05 |
| BMP1 | 8.335419092 | 6.748441864 | 5.01E-18 | 5.68E-16 |
| CLCA1 | 8.334922653 | 3.915203425 | 3.23E-07 | 3.82E-06 |
| SULF1 | 8.334677176 | 6.04657043 | 1.41E-07 | 1.84E-06 |
| LOC111096729 | 8.330412597 | 0.512919829 | 3.16E-07 | 3.75E-06 |
| LOC106559857 | 8.329850352 | 2.683285404 | 3.18E-06 | 2.82E-05 |
| HDX | 8.329161301 | 1.729021756 | 8.26E-11 | 2.29E-09 |

| | | | | |
|---|---|---|---|---|
| GREM2 | 8.327320708 | 0.283867052 | 1.03E-05 | 7.80E-05 |
| C1H9orf135 | 8.323467878 | -0.027540664 | 9.21E-08 | 1.26E-06 |
| LOC119869724 | 8.320658813 | 1.629833063 | 7.47E-09 | 1.33E-07 |
| FRRS1L | 8.314422787 | 1.293269827 | 4.55E-06 | 3.84E-05 |
| LOC102157230 | 8.314296218 | 0.659194117 | 1.18E-07 | 1.56E-06 |
| LOC111098518 | 8.314010352 | 0.289873316 | 6.68E-07 | 7.19E-06 |
| LOC102157027 | 8.313557356 | 0.907878081 | 3.82E-10 | 9.22E-09 |
| LOC111098342 | 8.311664995 | -0.144551088 | 8.29E-06 | 6.46E-05 |
| LOC119871225 | 8.310215785 | 0.251995116 | 1.03E-08 | 1.77E-07 |
| EPHA7 | 8.307588851 | 0.263937546 | 0.000443276 | 0.002034115 |
| LOC111096605 | 8.30720758 | 0.828499345 | 8.43E-07 | 8.78E-06 |
| EML1 | 8.306588621 | 5.208225819 | 2.14E-14 | 1.24E-12 |
| ISLR | 8.303224167 | 5.769549086 | 5.55E-10 | 1.29E-08 |
| LOC119873953 | 8.298958196 | 0.921941736 | 4.18E-08 | 6.23E-07 |
| ACKR1 | 8.298200421 | 4.85870951 | 5.19E-07 | 5.78E-06 |
| LOC102151536 | 8.295814061 | 0.784864055 | 4.80E-06 | 4.02E-05 |
| LOC111093527 | 8.290836212 | 0.895760304 | 1.02E-05 | 7.75E-05 |
| CLEC14A | 8.287355568 | 5.077632036 | 4.19E-14 | 2.32E-12 |
| ASTN2 | 8.286155335 | 0.610050648 | 4.54E-06 | 3.84E-05 |
| PLA2R1 | 8.274776191 | 6.277328273 | 3.31E-08 | 5.07E-07 |
| LOC119876663 | 8.273059228 | 0.282123268 | 6.74E-08 | 9.58E-07 |
| LOC111098841 | 8.27275025 | 0.97344906 | 0.000144878 | 0.000779916 |
| LOC119871885 | 8.267437335 | 0.80014932 | 1.87E-06 | 1.78E-05 |
| LOC111094852 | 8.265565772 | 2.670311704 | 4.56E-12 | 1.70E-10 |
| NDUFA4L2 | 8.265467886 | 4.90859406 | 1.63E-10 | 4.24E-09 |
| EDN1 | 8.257170435 | 3.252616345 | 2.13E-07 | 2.64E-06 |
| LOC119877025 | 8.255648896 | 0.715131828 | 5.00E-08 | 7.35E-07 |
| SHOX2 | 8.25153747 | 3.311766425 | 1.11E-09 | 2.43E-08 |
| PTCH1 | 8.250487247 | 8.898354576 | 2.00E-11 | 6.38E-10 |
| CNKSR2 | 8.249308858 | 3.632406491 | 9.79E-08 | 1.33E-06 |
| TLN2 | 8.249098699 | 6.796057541 | 7.88E-10 | 1.78E-08 |
| LOC111096033 | 8.248627101 | 1.14848918 | 7.09E-08 | 9.97E-07 |
| LOC119877698 | 8.247823606 | 1.142200867 | 1.65E-05 | 0.000118938 |
| CLDN5 | 8.243926088 | 3.766689887 | 3.16E-09 | 6.14E-08 |
| NXPH2 | 8.243850512 | 1.34341766 | 0.000448203 | 0.002053247 |
| LOC119864794 | 8.24298285 | -0.229696666 | 3.31E-08 | 5.06E-07 |
| LOC106560202 | 8.242868754 | 1.252007611 | 1.44E-07 | 1.87E-06 |
| LSAMP | 8.242189496 | 2.477581407 | 4.56E-05 | 0.000290033 |
| FMO3 | 8.241189976 | 2.987728736 | 2.34E-06 | 2.16E-05 |

| | | | | |
|---|---|---|---|---|
| LOC111093431 | 8.239570388 | 0.451968022 | 2.66E-06 | 2.42E-05 |
| LOC119864958 | 8.239229538 | -0.247241443 | 1.87E-05 | 0.000132802 |
| LOC119877953 | 8.237818494 | -0.070912091 | 7.60E-07 | 8.06E-06 |
| UBE2QL1 | 8.234099219 | 2.981251198 | 1.03E-05 | 7.80E-05 |
| LOC119870153 | 8.23353712 | 1.401462958 | 1.71E-07 | 2.17E-06 |
| GALNT16 | 8.232000434 | 3.860645394 | 1.14E-15 | 8.43E-14 |
| C11H9orf152 | 8.230735478 | 0.624048895 | 2.96E-06 | 2.65E-05 |
| GRM8 | 8.229705336 | -0.136344498 | 0.000251245 | 0.001247764 |
| KCNA4 | 8.2267689 | 0.079490964 | 2.63E-05 | 0.000179162 |
| SYTL5 | 8.223976156 | -0.243714821 | 1.10E-06 | 1.12E-05 |
| LOC111097151 | 8.222641595 | 0.184318899 | 0.000653346 | 0.002842796 |
| ODF3L1 | 8.215486342 | 0.04395481 | 2.78E-07 | 3.34E-06 |
| SLC7A2 | 8.211436592 | 6.186825115 | 3.52E-10 | 8.54E-09 |
| CHST1 | 8.210213418 | 4.638912435 | 7.67E-10 | 1.74E-08 |
| LOC119876391 | 8.207350813 | 0.414904768 | 4.68E-07 | 5.28E-06 |
| WNT9A | 8.205572508 | 3.120357257 | 1.05E-15 | 7.91E-14 |
| NTNG1 | 8.205478773 | 2.174423195 | 2.83E-05 | 0.000191012 |
| LOC111095455 | 8.203709392 | 0.274346861 | 1.11E-06 | 1.13E-05 |
| ATP8B1 | 8.202925753 | 6.384780976 | 1.34E-13 | 6.71E-12 |
| KIF26A | 8.202400915 | 4.925492616 | 8.19E-12 | 2.87E-10 |
| LOC119874569 | 8.201856061 | 1.549826454 | 7.32E-09 | 1.30E-07 |
| ARNT2 | 8.195906313 | 5.279311146 | 2.96E-18 | 3.60E-16 |
| LOC119864578 | 8.193499035 | 0.081552317 | 3.69E-06 | 3.20E-05 |
| NHSL1 | 8.186408756 | 5.651275362 | 9.92E-11 | 2.69E-09 |
| DCDC1 | 8.184865631 | -0.129303684 | 2.19E-06 | 2.04E-05 |
| SORBS2 | 8.179942974 | 6.520214865 | 2.66E-11 | 8.21E-10 |
| ZNF280B | 8.177973646 | 2.134385599 | 1.31E-14 | 7.86E-13 |
| LRRC10B | 8.173141318 | 0.457150724 | 2.44E-07 | 2.98E-06 |
| RAMP3 | 8.171156879 | 1.122259635 | 1.59E-06 | 1.54E-05 |
| PCDH17 | 8.165071529 | 5.349367252 | 1.64E-10 | 4.27E-09 |
| C1R | 8.162623905 | 7.240962243 | 9.53E-19 | 1.24E-16 |
| SHISA7 | 8.162318765 | 1.354962866 | 1.90E-06 | 1.80E-05 |
| LOC119870223 | 8.161865418 | 0.255143063 | 9.12E-06 | 7.03E-05 |
| GPR39 | 8.160328026 | -0.234467209 | 0.000424574 | 0.001964099 |
| CCL2 | 8.156380645 | 4.912737489 | 3.41E-13 | 1.59E-11 |
| EN1 | 8.155068515 | -0.381493734 | 7.59E-05 | 0.000451479 |
| LOC102156351 | 8.154503861 | 0.106790978 | 9.20E-09 | 1.60E-07 |
| INSYN1 | 8.147282054 | 3.414503434 | 1.71E-11 | 5.52E-10 |
| TLCD5 | 8.14646515 | 0.895232963 | 1.48E-10 | 3.92E-09 |

| | | | | |
|---|---|---|---|---|
| LMX1A | 8.127890974 | -0.078088595 | 8.10E-06 | 6.34E-05 |
| SLC51B | 8.126996221 | 3.365038132 | 5.97E-11 | 1.70E-09 |
| LOC119865404 | 8.126687895 | -0.018239303 | 8.39E-07 | 8.76E-06 |
| LOC102156875 | 8.124907757 | 0.924587693 | 7.50E-08 | 1.05E-06 |
| CCKBR | 8.123653247 | 0.061458884 | 0.000495521 | 0.002241584 |
| TACC2 | 8.1226139 | 6.413660388 | 6.31E-14 | 3.36E-12 |
| SELE | 8.11744737 | 4.095008426 | 6.59E-05 | 0.000398592 |
| LOC119878201 | 8.114027465 | 2.865343887 | 0.000156979 | 0.00083469 |
| LOC111097303 | 8.1118383 | 0.703229617 | 5.82E-07 | 6.39E-06 |
| ASPN | 8.100273564 | 5.702877188 | 7.88E-11 | 2.20E-09 |
| UNC13C | 8.098823566 | 1.048385292 | 3.99E-06 | 3.42E-05 |
| PLEKHH2 | 8.097975117 | 5.472810784 | 1.61E-14 | 9.57E-13 |
| LOC102154703 | 8.096072024 | 0.947325969 | 8.09E-07 | 8.49E-06 |
| SYNM | 8.092417285 | 6.029923787 | 5.72E-23 | 1.55E-20 |
| TMEM252 | 8.089757303 | 2.02327625 | 0.000169466 | 0.000890817 |
| LRRC32 | 8.086477936 | 5.448255675 | 3.33E-12 | 1.29E-10 |
| NMS | 8.085470257 | 0.435196295 | 2.76E-06 | 2.50E-05 |
| CHST8 | 8.084874583 | -0.192094506 | 2.87E-05 | 0.000193489 |
| SELENOM | 8.083948252 | 5.023964683 | 3.36E-18 | 4.00E-16 |
| LOC111092131 | 8.08088737 | 0.092654348 | 9.79E-07 | 1.00E-05 |
| SIX2 | 8.076913834 | -0.202156248 | 2.66E-06 | 2.42E-05 |
| LOC111096345 | 8.074579575 | -0.114288211 | 1.90E-06 | 1.80E-05 |
| ACKR3 | 8.063800236 | 6.239068478 | 1.48E-09 | 3.12E-08 |
| LOC111098054 | 8.058204172 | 0.090409058 | 1.30E-06 | 1.29E-05 |
| GPNMB | 8.055742435 | 6.084692171 | 4.01E-07 | 4.62E-06 |
| ITPRID1 | 8.052507658 | 0.788030196 | 0.000162177 | 0.000858329 |
| LOC102155159 | 8.050952931 | -0.085040809 | 2.93E-05 | 0.000196904 |
| LOC119870957 | 8.048095115 | 5.90135704 | 7.20E-20 | 1.19E-17 |
| DDAH1 | 8.046883266 | 2.549518708 | 3.77E-08 | 5.68E-07 |
| LOC111096417 | 8.041798859 | -0.077669339 | 0.001231287 | 0.004886592 |
| PRRG4 | 8.039593519 | 3.216365228 | 7.34E-05 | 0.000438415 |
| HMGCLL1 | 8.039496385 | 1.036154698 | 0.000118523 | 0.000661538 |
| LOC111094043 | 8.036762498 | -0.355804306 | 2.54E-05 | 0.000173857 |
| MLPH | 8.036135588 | 0.498440608 | 1.53E-06 | 1.49E-05 |
| TNFRSF12A | 8.033271338 | 4.670066757 | 2.41E-17 | 2.33E-15 |
| CNTNAP4 | 8.032959322 | -0.677192097 | 0.001467993 | 0.005657149 |
| LOC111095930 | 8.031405428 | -0.10425842 | 2.13E-07 | 2.63E-06 |
| GFRA3 | 8.030059975 | 3.58084879 | 4.25E-06 | 3.62E-05 |
| GRIK5 | 8.028583386 | 3.85709772 | 2.20E-11 | 6.91E-10 |

| | | | | |
|---|---|---|---|---|
| MIR214 | 8.028360636 | 0.069882933 | 5.87E-05 | 0.000361268 |
| LOC119869784 | 8.019261595 | -0.130373864 | 9.23E-09 | 1.60E-07 |
| CLEC3A | 8.016090997 | 0.020593842 | 0.000253679 | 0.001258695 |
| NT5M | 8.01567196 | 0.280588756 | 1.59E-06 | 1.54E-05 |
| FUT2 | 8.015669994 | -0.200090638 | 3.23E-07 | 3.82E-06 |
| PCDH9 | 8.011656393 | 2.825319202 | 4.99E-06 | 4.14E-05 |
| MTUS1 | 8.003177894 | 6.898352225 | 1.15E-20 | 2.26E-18 |
| TACR1 | 7.999675013 | 0.268078837 | 1.01E-06 | 1.04E-05 |
| LRRC3B | 7.998310208 | 0.482717992 | 8.85E-07 | 9.17E-06 |
| LOC111095802 | 7.993833554 | 0.038830682 | 1.24E-05 | 9.21E-05 |
| TRIM29 | 7.992532009 | -0.438388626 | 2.74E-05 | 0.000185679 |
| LRATD1 | 7.992525567 | 2.400937268 | 8.22E-05 | 0.000482933 |
| LOC119872501 | 7.984403617 | 0.986269112 | 2.48E-06 | 2.28E-05 |
| EFNA5 | 7.983348921 | 3.412502567 | 1.90E-10 | 4.89E-09 |
| LOC102153473 | 7.98158238 | 0.231720338 | 0.000665648 | 0.002887627 |
| LOC119873620 | 7.979003501 | 0.419086246 | 7.33E-09 | 1.30E-07 |
| GFAP | 7.978922988 | 8.323552466 | 0.000114861 | 0.000643915 |
| FAM240A | 7.97728381 | 0.31578524 | 2.48E-08 | 3.90E-07 |
| PXDC1 | 7.975052295 | 5.088029398 | 1.25E-15 | 9.15E-14 |
| LOC119873350 | 7.972822071 | 1.450504414 | 1.17E-11 | 3.94E-10 |
| FERMT2 | 7.969076607 | 6.586154362 | 2.54E-15 | 1.78E-13 |
| EFCAB1 | 7.968902178 | -0.086573686 | 1.47E-06 | 1.44E-05 |
| LOC106557821 | 7.968570544 | 4.489470996 | 1.86E-09 | 3.81E-08 |
| LOC119874664 | 7.96658596 | 1.489536654 | 8.50E-06 | 6.60E-05 |
| LOC102152820 | 7.964910264 | 0.614591371 | 6.10E-07 | 6.65E-06 |
| USP43 | 7.958705583 | -0.399767548 | 5.55E-06 | 4.55E-05 |
| LOC111090916 | 7.958448297 | -0.323225747 | 0.001077089 | 0.004364948 |
| LOC119867211 | 7.957668858 | -0.082183634 | 1.53E-05 | 0.000110908 |
| PWWP3B | 7.956312259 | 2.201651948 | 2.70E-09 | 5.31E-08 |
| HOXD11 | 7.95264779 | 0.143094706 | 0.000502791 | 0.002269734 |
| LOC111090583 | 7.949316071 | 0.460294435 | 1.54E-08 | 2.54E-07 |
| LOC111091285 | 7.949110608 | 0.581209882 | 8.62E-05 | 0.00050354 |
| LOC119867201 | 7.948873709 | -0.173094374 | 0.000328804 | 0.001573452 |
| LOC119878220 | 7.947507087 | 0.944495736 | 6.73E-05 | 0.000405563 |
| COL4A2 | 7.944695056 | 9.439885974 | 1.14E-22 | 3.05E-20 |
| LOC119876375 | 7.944263277 | 0.846868525 | 0.000266095 | 0.001310402 |
| LOC111091977 | 7.943370068 | -0.021594054 | 1.96E-06 | 1.85E-05 |
| LOC111091268 | 7.941713607 | -0.051929178 | 2.58E-06 | 2.35E-05 |
| LOC102155235 | 7.940962011 | 0.839461077 | 7.15E-05 | 0.000427863 |

| | | | | |
|---|---|---|---|---|
| COL5A1 | 7.93547559 | 10.21281252 | 5.79E-14 | 3.12E-12 |
| LOC119871199 | 7.924199422 | 0.389207336 | 4.61E-06 | 3.88E-05 |
| LOC102156409 | 7.918992744 | 0.490990009 | 0.000426361 | 0.001971101 |
| LOC119869349 | 7.91289968 | -0.579841762 | 0.000108366 | 0.00061354 |
| CFAP299 | 7.912085365 | 0.421115843 | 0.000222625 | 0.00112601 |
| LOC106558272 | 7.910543984 | -0.07570645 | 1.38E-06 | 1.36E-05 |
| PDZRN4 | 7.904991734 | 0.854280973 | 3.36E-06 | 2.96E-05 |
| AHNAK2 | 7.901855954 | 9.567712941 | 5.79E-09 | 1.05E-07 |
| DAB2 | 7.899419869 | 7.471066334 | 8.89E-16 | 6.78E-14 |
| HOXC8 | 7.898005174 | 1.832259835 | 1.36E-08 | 2.27E-07 |
| LOC119864774 | 7.897028877 | 0.409425164 | 7.79E-08 | 1.08E-06 |
| LOC111096418 | 7.892345915 | -0.193028474 | 0.000152408 | 0.000812782 |
| SCN3A | 7.88592625 | 0.900725356 | 7.53E-05 | 0.000448688 |
| SNX31 | 7.88338276 | 0.539377828 | 8.32E-07 | 8.70E-06 |
| LOC119869913 | 7.868539045 | 3.33131558 | 1.35E-05 | 9.97E-05 |
| LOC119873383 | 7.864889084 | -0.438183501 | 2.25E-06 | 2.08E-05 |
| ADAM33 | 7.863797381 | 6.120389292 | 9.63E-10 | 2.12E-08 |
| LOC119871186 | 7.862140049 | -0.667978632 | 0.000171996 | 0.000902149 |
| LOC487015 | 7.860062625 | 4.646678713 | 0.002107196 | 0.007706593 |
| MAP1B | 7.855545547 | 10.22954797 | 6.96E-10 | 1.59E-08 |
| LOC102154971_2 | 7.843693023 | 0.061200045 | 5.47E-06 | 4.49E-05 |
| LOC102156930 | 7.839904711 | -0.298260308 | 1.93E-06 | 1.82E-05 |
| CADM2 | 7.838851098 | 3.525607641 | 3.33E-05 | 0.000220072 |
| LOC111093427 | 7.836392039 | -0.111863546 | 1.03E-06 | 1.05E-05 |
| L1CAM | 7.836158587 | 6.950006434 | 4.32E-09 | 8.08E-08 |
| RAB36 | 7.834702375 | 0.964922949 | 4.12E-09 | 7.76E-08 |
| SEZ6L | 7.83411537 | 0.001617831 | 0.001893543 | 0.00701757 |
| LOC102154149 | 7.833382015 | -0.248983566 | 5.39E-07 | 5.97E-06 |
| LOC119864248 | 7.830285621 | 0.279164924 | 4.66E-05 | 0.000295569 |
| LOC111097807 | 7.829037931 | 5.116628693 | 8.02E-09 | 1.42E-07 |
| GNAT1 | 7.824246048 | 0.686506635 | 1.18E-06 | 1.19E-05 |
| LOC102152120 | 7.820924837 | -0.26497673 | 2.30E-06 | 2.13E-05 |
| FLYWCH2 | 7.812441527 | 2.429595431 | 5.62E-14 | 3.05E-12 |
| TEAD3 | 7.808840957 | 5.612726181 | 7.63E-12 | 2.70E-10 |
| LOC119865795 | 7.807476587 | 1.53070995 | 1.43E-05 | 0.000104936 |
| LOC102153932 | 7.806185915 | 0.581213365 | 1.86E-07 | 2.33E-06 |
| RSPO1 | 7.804964921 | -0.47843352 | 0.001455614 | 0.005617435 |
| LOC102154499 | 7.804410573 | -0.452140934 | 3.36E-07 | 3.95E-06 |
| LOC119865163 | 7.801796626 | 0.146076265 | 6.52E-07 | 7.03E-06 |

| | | | | |
|---|---|---|---|---|
| LOC119876532 | 7.800793374 | 2.407768917 | 1.79E-06 | 1.72E-05 |
| LOC111094968 | 7.797755806 | 0.88423287 | 4.77E-07 | 5.38E-06 |
| COL9A1 | 7.796438055 | 1.272412734 | 0.000780991 | 0.003306589 |
| ETV1 | 7.793250918 | 4.455745573 | 2.29E-09 | 4.61E-08 |
| LOC102157367 | 7.791868314 | 0.86420396 | 0.000431428 | 0.001989434 |
| SDSL | 7.790716504 | -0.394755539 | 0.000252955 | 0.001255389 |
| LOC111097369 | 7.789256654 | -0.21747711 | 2.65E-06 | 2.41E-05 |
| TMEM47 | 7.78917088 | 5.69129756 | 1.54E-10 | 4.05E-09 |
| LOC119878320 | 7.787375467 | 2.109769885 | 5.78E-05 | 0.000356278 |
| LOC111095162 | 7.786884059 | 5.368983658 | 1.05E-05 | 7.96E-05 |
| LOC119876766 | 7.786560456 | -0.336386863 | 0.000326822 | 0.001566388 |
| LOC119873458 | 7.778855988 | -0.347094129 | 1.88E-05 | 0.000133475 |
| TMEM158 | 7.775605966 | 0.874562769 | 7.87E-07 | 8.28E-06 |
| THBS4 | 7.774522156 | 7.058398981 | 2.52E-05 | 0.000173064 |
| LOC111095237 | 7.772711247 | -0.307969455 | 2.19E-05 | 0.000152931 |
| LOC106560184 | 7.771902561 | 0.911639433 | 2.33E-07 | 2.86E-06 |
| CLDN6 | 7.769700411 | -0.576329541 | 3.78E-06 | 3.27E-05 |
| LOC102153633 | 7.766045435 | -0.230526743 | 0.001280341 | 0.005057194 |
| LOC102152771 | 7.763891831 | -0.422467309 | 0.000910753 | 0.003786448 |
| LOC111090645 | 7.763626111 | 0.064015406 | 2.31E-05 | 0.000159988 |
| LOC102154628 | 7.760650222 | -0.104726398 | 2.97E-07 | 3.55E-06 |
| LOC119866939 | 7.758015416 | -0.266776625 | 8.64E-05 | 0.000504453 |
| FBLIM1 | 7.757275091 | 5.019309979 | 5.75E-14 | 3.10E-12 |
| LOC111091916 | 7.757261263 | -0.264759377 | 8.60E-06 | 6.68E-05 |
| CA10 | 7.756899133 | -0.058270829 | 1.80E-06 | 1.72E-05 |
| LOC111091291 | 7.748800336 | 0.222221932 | 8.70E-05 | 0.000507306 |
| LOC111091516 | 7.747603302 | -0.175649721 | 0.000847229 | 0.003549584 |
| LOC119867905 | 7.745506509 | -0.756113767 | 0.000226787 | 0.001143591 |
| EYA4 | 7.745455396 | 0.905366523 | 0.000413472 | 0.001920116 |
| FREM2 | 7.745212974 | 3.630195093 | 5.06E-06 | 4.20E-05 |
| LOC119865393 | 7.743132529 | 1.80027024 | 4.07E-06 | 3.48E-05 |
| RGL3 | 7.743118356 | -0.413544932 | 0.00011708 | 0.00065483 |
| LOC119866863 | 7.741442547 | -0.378331812 | 4.65E-06 | 3.91E-05 |
| SYT12 | 7.736413951 | -0.441151835 | 1.89E-06 | 1.79E-05 |
| LOC111098780 | 7.730422738 | -0.017333781 | 0.000439714 | 0.002021779 |
| FSD1 | 7.727826771 | 1.306787069 | 1.01E-07 | 1.36E-06 |
| C20H3orf14 | 7.726483122 | -0.503985008 | 2.30E-06 | 2.13E-05 |
| LOC102155408 | 7.726461375 | 0.643122495 | 5.98E-05 | 0.000367009 |
| CYP8B1 | 7.719988307 | 2.28352446 | 0.00018723 | 0.000969153 |

| | | | | |
|---|---|---|---|---|
| MYO3B | 7.716004479 | 2.420442316 | 8.55E-05 | 0.00049978 |
| ACOT4 | 7.710296335 | 0.551645276 | 2.67E-06 | 2.43E-05 |
| ABCD2 | 7.706886924 | 1.10475221 | 8.06E-05 | 0.000475119 |
| LOC102152272 | 7.70567438 | 0.151384298 | 2.86E-06 | 2.57E-05 |
| WDR31 | 7.704483906 | -0.343323395 | 3.15E-06 | 2.80E-05 |
| DLL4 | 7.704232543 | 4.705818238 | 3.89E-12 | 1.47E-10 |
| FOXD3 | 7.694083527 | 3.462294615 | 0.000349514 | 0.001659369 |
| LOC106558329 | 7.693691296 | -0.545609137 | 6.46E-05 | 0.00039204 |
| SHC3 | 7.692620491 | 3.427935448 | 1.49E-05 | 0.000108493 |
| PPP1R1B | 7.685937743 | 4.065389045 | 0.000148704 | 0.000796161 |
| LOC119867672 | 7.684018752 | 0.22797803 | 2.24E-05 | 0.000155587 |
| LOC111090331 | 7.681980438 | -0.178745039 | 0.000223744 | 0.001130351 |
| ZNF521 | 7.681025543 | 6.030669947 | 9.10E-17 | 8.09E-15 |
| BMP5 | 7.678221596 | 0.673005441 | 0.000142522 | 0.000769525 |
| LOC119866919 | 7.674279882 | 2.506792283 | 1.56E-07 | 2.00E-06 |
| EPHA2 | 7.672455263 | 5.265749176 | 1.41E-18 | 1.79E-16 |
| LOC106559534 | 7.672001096 | -0.607780696 | 4.42E-06 | 3.75E-05 |
| LOC102152695 | 7.668642543 | -0.166452179 | 0.000266795 | 0.001313254 |
| TFCP2L1 | 7.665744523 | 2.424482681 | 7.77E-06 | 6.11E-05 |
| FBN2 | 7.664561815 | 4.803824071 | 0.000208753 | 0.001068564 |
| LOC102156880 | 7.664555401 | 0.113763885 | 5.79E-08 | 8.35E-07 |
| LOC119867348 | 7.662723195 | -0.47340517 | 1.04E-06 | 1.07E-05 |
| MSX2 | 7.660241325 | -0.795131771 | 1.00E-05 | 7.62E-05 |
| TEAD1 | 7.659091116 | 7.697410813 | 4.10E-18 | 4.71E-16 |
| LOC102156172 | 7.658746498 | -0.094375104 | 1.50E-05 | 0.000109146 |
| ADAMTS4 | 7.654283125 | 6.045134067 | 3.24E-09 | 6.26E-08 |
| RAMP2 | 7.651860943 | 5.12846453 | 9.57E-12 | 3.29E-10 |
| LOC111097108 | 7.642563682 | 0.886922415 | 1.38E-10 | 3.66E-09 |
| LOC111096962 | 7.641792737 | 0.828532559 | 6.09E-06 | 4.93E-05 |
| LOC119866439 | 7.641595181 | 0.205876857 | 6.40E-07 | 6.93E-06 |
| LOC102151382 | 7.636308302 | 0.980349695 | 6.69E-12 | 2.41E-10 |
| ALDH3A1 | 7.633578634 | 0.168881586 | 0.000129775 | 0.000712415 |
| MIPOL1 | 7.629879596 | 4.340482732 | 3.23E-15 | 2.18E-13 |
| FGF7 | 7.627024705 | 4.121009487 | 2.01E-05 | 0.000141773 |
| LOC102152802 | 7.621707548 | 2.088194701 | 4.67E-05 | 0.000296071 |
| PHLDB1 | 7.6214962 | 8.864990368 | 3.03E-16 | 2.46E-14 |
| LOC102154871 | 7.619396117 | 0.219323113 | 2.91E-06 | 2.61E-05 |
| LOC111092938 | 7.619303496 | -0.853974097 | 3.95E-05 | 0.000255489 |
| LOC100684412 | 7.618534264 | -0.309015003 | 5.16E-08 | 7.55E-07 |

| | | | | |
|---|---|---|---|---|
| LOC102153278 | 7.616596503 | -0.656249087 | 9.57E-06 | 7.33E-05 |
| HOXD10 | 7.614932678 | -0.485998157 | 2.22E-05 | 0.000154595 |
| CCDC33 | 7.613997909 | -0.786961818 | 4.71E-06 | 3.95E-05 |
| LOC111094721 | 7.609208621 | -0.48166308 | 6.01E-06 | 4.87E-05 |
| LOC106557720 | 7.606694717 | -0.001909604 | 3.16E-05 | 0.000210472 |
| SLC35F1 | 7.60465348 | 2.870379435 | 0.000253836 | 0.001259188 |
| LOC119871841 | 7.603645303 | -0.468610214 | 4.03E-06 | 3.45E-05 |
| NR2F1 | 7.602855114 | 6.380131087 | 1.05E-06 | 1.07E-05 |
| LOC119876644 | 7.601497866 | -0.037897313 | 2.70E-06 | 2.45E-05 |
| NME5 | 7.597720005 | -0.5420633 | 2.57E-05 | 0.0001757 |
| LOC119873915 | 7.594299811 | -0.240826954 | 6.15E-05 | 0.000376015 |
| C1H18orf63 | 7.590759856 | 0.913549879 | 2.82E-05 | 0.000190454 |
| LOC119877654 | 7.588989113 | 3.481563773 | 2.62E-07 | 3.18E-06 |
| KRT124 | 7.586482166 | 2.405437208 | 0.000165301 | 0.000872518 |
| LOC111095598 | 7.584195709 | 0.545381373 | 1.79E-06 | 1.72E-05 |
| LOC475575 | 7.581564509 | 3.0194944 | 3.93E-18 | 4.55E-16 |
| ALOXE3 | 7.580503479 | 1.508373707 | 8.91E-06 | 6.89E-05 |
| LOC119877032 | 7.579868293 | 1.018860503 | 1.82E-06 | 1.74E-05 |
| CXCL14 | 7.579706741 | 8.24693719 | 6.07E-09 | 1.10E-07 |
| ZNF285 | 7.577749559 | -0.240743366 | 6.02E-09 | 1.09E-07 |
| FAM83B | 7.575369213 | -0.732400792 | 0.000157183 | 0.000835574 |
| BCAR1 | 7.573017232 | 5.776672391 | 8.52E-21 | 1.74E-18 |
| APOA1 | 7.572611228 | 4.808369067 | 1.63E-09 | 3.39E-08 |
| C35H6orf52 | 7.571049752 | -0.618348227 | 6.51E-06 | 5.22E-05 |
| LOC119866382 | 7.570342179 | -0.194606593 | 5.41E-06 | 4.45E-05 |
| LOC111097493 | 7.567920865 | -0.513046628 | 7.44E-06 | 5.88E-05 |
| LOC102156428 | 7.565616705 | 1.109227249 | 2.43E-11 | 7.55E-10 |
| LOC102152416 | 7.564848093 | -0.300580716 | 0.000211677 | 0.001079961 |
| LOC111096953 | 7.564302041 | -0.111901299 | 5.74E-05 | 0.000354611 |
| HOXA2 | 7.559896858 | -0.301164173 | 2.61E-07 | 3.16E-06 |
| LOC102155309 | 7.558141041 | 0.416347118 | 4.61E-05 | 0.000292613 |
| LOC119872146 | 7.55764195 | 5.527858627 | 1.69E-13 | 8.34E-12 |
| LOC102157065 | 7.557118643 | 0.41116458 | 8.02E-05 | 0.000473177 |
| LOC119870166 | 7.556034211 | -0.382607987 | 1.05E-05 | 7.98E-05 |
| LOC106559589 | 7.555044793 | -0.677158374 | 3.44E-06 | 3.02E-05 |
| LOC119866379 | 7.547478653 | 0.350176269 | 1.27E-06 | 1.26E-05 |
| LOC119877884 | 7.539164684 | -0.531578115 | 1.81E-08 | 2.93E-07 |
| LOC102156220 | 7.537937996 | -0.397308761 | 8.74E-05 | 0.0005094 |
| LOC102154924 | 7.534379311 | 0.521288573 | 8.13E-05 | 0.000478529 |

| | | | | |
|---|---|---|---|---|
| DOK5 | 7.52972209 | -0.222486689 | 0.001046191 | 0.004253255 |
| LOC119873382 | 7.523051049 | 0.177225694 | 2.12E-06 | 1.98E-05 |
| LOC119868357 | 7.522764034 | -0.315814247 | 1.50E-05 | 0.000108928 |
| CRISPLD1 | 7.520792729 | 0.042039546 | 0.000309419 | 0.001496847 |
| LOC111095541 | 7.52075582 | 4.684405249 | 3.82E-12 | 1.45E-10 |
| ARL4D | 7.520718331 | 2.165225974 | 5.27E-08 | 7.68E-07 |
| BOK | 7.520368264 | 4.411733942 | 5.68E-12 | 2.09E-10 |
| LOC111096191 | 7.520273722 | -0.559533458 | 6.06E-06 | 4.91E-05 |
| NPHS2 | 7.520136117 | -0.305737819 | 4.07E-05 | 0.000262159 |
| SDC1 | 7.516787459 | 5.614454307 | 1.07E-06 | 1.09E-05 |
| LOC102156472 | 7.51664319 | -0.551967697 | 0.000202372 | 0.001038844 |
| LOC106559851 | 7.516447943 | 1.040432819 | 1.55E-05 | 0.000112579 |
| LOC111091731 | 7.510353259 | 1.25229855 | 1.60E-09 | 3.34E-08 |
| PEG10 | 7.508025947 | 4.773527036 | 1.07E-06 | 1.09E-05 |
| CHRNA4 | 7.505897263 | -0.58619325 | 0.001093832 | 0.004420403 |
| LOC106559379 | 7.503689627 | 0.368240209 | 3.09E-06 | 2.75E-05 |
| TLE2 | 7.503278171 | 5.425777773 | 2.01E-10 | 5.14E-09 |
| LOC111095981 | 7.502613109 | -0.766050751 | 3.11E-05 | 0.000207571 |
| COL27A1 | 7.502551462 | 6.937883102 | 4.44E-10 | 1.06E-08 |
| LRFN2 | 7.49813437 | -0.259794775 | 9.84E-05 | 0.00056447 |
| FBXL16 | 7.497624124 | 5.429958795 | 0.000113325 | 0.000636672 |
| FGF14 | 7.49664812 | -0.428650966 | 2.95E-06 | 2.64E-05 |
| LOC102154532 | 7.495826755 | -0.720330055 | 3.76E-06 | 3.26E-05 |
| LOC119878421 | 7.491568771 | 6.193365637 | 0.004672014 | 0.01510841 |
| PXDNL | 7.49116282 | 1.432613149 | 8.85E-11 | 2.43E-09 |
| AFAP1L2 | 7.490129184 | 7.463426438 | 1.01E-11 | 3.45E-10 |
| LOC111092805 | 7.489114542 | 3.810797803 | 5.52E-07 | 6.10E-06 |
| LOC119877187 | 7.488328354 | 7.713906466 | 0.000122871 | 0.00068108 |
| LOC119868391 | 7.487878978 | 1.209447224 | 1.58E-05 | 0.00011436 |
| IGFBP7 | 7.486892621 | 8.047916243 | 3.35E-13 | 1.57E-11 |
| CERCAM | 7.480388681 | 5.433036903 | 1.03E-09 | 2.26E-08 |
| TMEM145 | 7.478712763 | 2.942181033 | 1.10E-06 | 1.12E-05 |
| LOC119867338 | 7.476838132 | 0.310754122 | 0.000177584 | 0.00092697 |
| DDR2 | 7.472422884 | 7.56422327 | 1.08E-17 | 1.15E-15 |
| LOC111096271 | 7.472044081 | 0.329862136 | 8.56E-05 | 0.000500185 |
| TMEM138 | 7.471872949 | 0.650117435 | 3.13E-13 | 1.47E-11 |
| LOC119870213 | 7.465963408 | -0.277127111 | 0.000464934 | 0.002118287 |
| ITGB8 | 7.463537638 | 7.732548015 | 2.28E-05 | 0.000158274 |
| KCNMB2 | 7.462232528 | 1.011722581 | 2.82E-05 | 0.000190659 |

| | | | | |
|---|---|---|---|---|
| LOC106558635 | 7.461457633 | 0.593967608 | 0.000138442 | 0.000750866 |
| MYO1B | 7.460373466 | 7.443425177 | 7.19E-19 | 9.56E-17 |
| AADAC | 7.458361225 | 0.577267479 | 0.000119442 | 0.000665807 |
| LOC102157149 | 7.458010395 | -0.518668711 | 3.12E-06 | 2.78E-05 |
| DYSF | 7.454211476 | 6.21989557 | 5.60E-09 | 1.03E-07 |
| PLPP3 | 7.453190391 | 6.312788876 | 8.29E-11 | 2.29E-09 |
| LOC478381 | 7.453029764 | 2.67333021 | 1.21E-05 | 9.04E-05 |
| CDH7 | 7.45109477 | 2.696168003 | 3.24E-05 | 0.000215248 |
| TMTC1 | 7.449057653 | 6.018896559 | 1.41E-10 | 3.72E-09 |
| SYT16 | 7.448701788 | 3.329275474 | 2.74E-06 | 2.48E-05 |
| KRT40 | 7.447746475 | -0.191125831 | 3.01E-06 | 2.69E-05 |
| MMP11 | 7.446994698 | 4.087524932 | 3.90E-18 | 4.54E-16 |
| RAC3 | 7.443983406 | 2.624501998 | 2.87E-11 | 8.71E-10 |
| TCP11 | 7.443333397 | -0.217553845 | 7.61E-06 | 5.99E-05 |
| PRSS22 | 7.440024455 | 1.01264006 | 1.99E-08 | 3.18E-07 |
| LOC111092297 | 7.439433275 | -0.31787125 | 0.000160362 | 0.000850177 |
| MYL7 | 7.436412396 | -0.532634116 | 2.09E-05 | 0.0001461 |
| LOC119870969 | 7.434120225 | -0.096009847 | 0.000176835 | 0.0009235 |
| SPP2 | 7.432149984 | 0.455205283 | 0.000116521 | 0.000652043 |
| LOC119865317 | 7.429620422 | -0.625618482 | 0.00012602 | 0.000695679 |
| MPPED2 | 7.428551428 | 0.585447425 | 9.04E-06 | 6.97E-05 |
| LOC119867320 | 7.427432402 | -0.375978116 | 1.30E-06 | 1.29E-05 |
| LOC119871683 | 7.424159679 | 0.216855273 | 9.39E-05 | 0.000541346 |
| PKP1 | 7.42265351 | 2.353986696 | 3.19E-07 | 3.79E-06 |
| LOC111091730 | 7.417116987 | 4.33764301 | 3.02E-15 | 2.07E-13 |
| GFY | 7.416924139 | 1.707925848 | 0.001074296 | 0.004355261 |
| THSD7B | 7.413874357 | -0.102561415 | 5.55E-05 | 0.000343622 |
| LOC119865693 | 7.410405731 | -0.070476726 | 0.000127763 | 0.000704222 |
| CTHRC1 | 7.406636281 | 3.425453574 | 6.92E-06 | 5.51E-05 |
| LOC480926 | 7.40533473 | 2.783272512 | 6.63E-08 | 9.43E-07 |
| PCK1 | 7.404787764 | 1.338136339 | 0.002937089 | 0.010232362 |
| GPC3 | 7.40435552 | 4.613102538 | 5.42E-06 | 4.45E-05 |
| BOC | 7.401758659 | 7.634144296 | 2.12E-08 | 3.38E-07 |
| SSPN | 7.39940072 | 5.916040825 | 1.12E-12 | 4.71E-11 |
| LOC102152920 | 7.398382174 | -0.545130603 | 3.22E-06 | 2.85E-05 |
| LOC119872696 | 7.397688787 | -0.676932281 | 7.16E-06 | 5.67E-05 |
| USP46 | 7.392329276 | 4.817999009 | 1.50E-10 | 3.94E-09 |
| FSCN1 | 7.390124664 | 7.460825835 | 9.85E-13 | 4.20E-11 |
| SERTAD4 | 7.380523658 | 4.647731584 | 3.13E-07 | 3.72E-06 |

| | | | | |
|---|---|---|---|---|
| KCNC4 | 7.379809395 | 3.730898468 | 2.34E-09 | 4.70E-08 |
| LOC119864187 | 7.378373398 | 0.228729742 | 0.000501945 | 0.002266857 |
| DAB2IP | 7.378218793 | 6.271724674 | 5.55E-13 | 2.50E-11 |
| LOC119869556 | 7.377090291 | 1.017352348 | 4.65E-07 | 5.27E-06 |
| PLXDC1 | 7.37635884 | 6.53911214 | 8.39E-14 | 4.34E-12 |
| NUDT14 | 7.373451137 | 2.838417924 | 3.15E-09 | 6.12E-08 |
| EXTL1 | 7.36966825 | 2.695902074 | 4.94E-08 | 7.27E-07 |
| LOC119873636 | 7.369120698 | -0.423502925 | 6.25E-05 | 0.000380874 |
| LOC111090858 | 7.368535823 | -0.104359089 | 0.002759782 | 0.009705599 |
| NBL1 | 7.367317776 | 7.046337537 | 7.18E-14 | 3.76E-12 |
| SCUBE3 | 7.365925612 | 4.704121534 | 1.54E-07 | 1.98E-06 |
| SEMA3A | 7.36344657 | 2.294430399 | 0.000512376 | 0.002305321 |
| SEMA6D | 7.360372445 | 5.241162045 | 1.71E-10 | 4.45E-09 |
| LOC119878227 | 7.357734862 | -0.424125271 | 2.81E-05 | 0.000190368 |
| LOC100686288 | 7.357232963 | -0.657791146 | 9.49E-05 | 0.000546317 |
| LOC102156830 | 7.357149137 | 0.484259419 | 1.61E-06 | 1.56E-05 |
| LOC119871542 | 7.356388592 | -0.098747307 | 1.22E-06 | 1.22E-05 |
| LOC119877324 | 7.354313359 | 0.154727041 | 1.25E-05 | 9.27E-05 |
| TMEM176B | 7.351047392 | 4.641208443 | 5.83E-12 | 2.14E-10 |
| LOC100683920 | 7.350774324 | -0.877524115 | 4.10E-05 | 0.000263927 |
| ADRA1B | 7.349868022 | -0.064190581 | 7.13E-05 | 0.000426595 |
| NOTO | 7.349712991 | -0.640874757 | 1.07E-05 | 8.12E-05 |
| LOC119873914 | 7.348798141 | -0.555471613 | 6.33E-07 | 6.86E-06 |
| LOC119866951 | 7.348361865 | -0.462731791 | 3.08E-06 | 2.75E-05 |
| ACADL | 7.346121061 | 4.51685471 | 3.55E-11 | 1.06E-09 |
| GLDN | 7.345872959 | 4.861895338 | 1.40E-09 | 2.95E-08 |
| LOC111093471 | 7.343099337 | 0.096065814 | 0.000228725 | 0.001151222 |
| LOC119870068 | 7.342657748 | -0.790774929 | 1.46E-05 | 0.000106725 |
| CTXN1 | 7.336314474 | 3.96904215 | 1.51E-05 | 0.00010975 |
| LOC106558464 | 7.33441906 | -0.548108431 | 7.78E-06 | 6.12E-05 |
| TMEM132B | 7.334382127 | 5.731161267 | 7.71E-17 | 6.87E-15 |
| LOC119876306 | 7.332070931 | 3.218826742 | 0.00025719 | 0.001272622 |
| LOC119867033 | 7.329644816 | -0.295704284 | 4.52E-06 | 3.82E-05 |
| PTP4A3 | 7.329057104 | 4.48399394 | 3.16E-13 | 1.48E-11 |
| LOC119872503 | 7.328593677 | -0.628882136 | 2.70E-05 | 0.000183532 |
| LOC119877770 | 7.327256588 | -0.479293809 | 0.00030707 | 0.001486397 |
| LOC119876731 | 7.325873135 | -0.854014953 | 0.000128055 | 0.000705297 |
| PRKG2 | 7.321687048 | 1.773239559 | 0.000510809 | 0.002299702 |
| LOC102151160 | 7.315202338 | 0.258929614 | 2.83E-05 | 0.000191256 |

| | | | | |
|---|---|---|---|---|
| MDGA2 | 7.310355833 | 1.701823852 | 0.002823835 | 0.009896665 |
| LOC486353 | 7.307508299 | -0.580424912 | 0.001269978 | 0.005020153 |
| LGALS2 | 7.301877355 | 1.460974896 | 3.84E-08 | 5.77E-07 |
| GASK1B | 7.299473165 | 5.73536055 | 3.80E-10 | 9.17E-09 |
| NUMBL | 7.299130622 | 6.317712268 | 1.03E-12 | 4.36E-11 |
| FLRT3 | 7.299025844 | 4.472512703 | 1.50E-08 | 2.47E-07 |
| LOC119867507 | 7.297180825 | 1.087421786 | 4.08E-11 | 1.20E-09 |
| GAL | 7.293100193 | -0.785572411 | 0.001602004 | 0.006094476 |
| MFAP4 | 7.292475119 | 6.427778212 | 1.82E-07 | 2.29E-06 |
| SPON2 | 7.287002206 | 5.318210655 | 1.51E-10 | 3.97E-09 |
| LRRC8E | 7.285423515 | 1.960853561 | 1.89E-06 | 1.79E-05 |
| CD36 | 7.283408878 | 5.989953483 | 5.17E-07 | 5.76E-06 |
| LOC111097989 | 7.27822646 | -0.749864429 | 0.000128815 | 0.000708583 |
| RAMP1 | 7.276316082 | 1.398033623 | 0.000191243 | 0.00098686 |
| LOC111097629 | 7.275748558 | -1.136015361 | 0.000698572 | 0.003006981 |
| LOC111096422 | 7.275274182 | -0.528507509 | 9.39E-06 | 7.21E-05 |
| RAB34 | 7.273827242 | 4.742033236 | 1.41E-16 | 1.24E-14 |
| OTOGL | 7.272707272 | 0.637577195 | 2.15E-06 | 2.00E-05 |
| CEP131 | 7.267644304 | 6.489169316 | 6.03E-09 | 1.09E-07 |
| SLC44A3 | 7.252328902 | 2.89552715 | 2.22E-14 | 1.29E-12 |
| LOC119863920 | 7.245739311 | 3.511831788 | 1.52E-06 | 1.48E-05 |
| LOC119865442 | 7.245323019 | 0.777604331 | 0.000704706 | 0.003027972 |
| KLHDC7B | 7.245173115 | 2.377589049 | 1.38E-08 | 2.29E-07 |
| LOC482919 | 7.238149033 | -0.001870562 | 2.56E-06 | 2.34E-05 |
| LOC119864566 | 7.236091878 | -0.785018876 | 0.000136344 | 0.000742435 |
| LOC119869351 | 7.234337969 | 0.425973294 | 1.00E-06 | 1.03E-05 |
| LOC106558031 | 7.234039984 | -0.712424046 | 3.27E-05 | 0.000216499 |
| LOC119867175 | 7.230939547 | -0.727423832 | 0.000175192 | 0.000916911 |
| LOC106558787 | 7.230109734 | -0.309135307 | 5.54E-05 | 0.000343375 |
| GABRA4 | 7.228834308 | 0.520111365 | 0.001982107 | 0.007314524 |
| LOC119871195 | 7.219076598 | -0.595466641 | 0.000427184 | 0.001974066 |
| DKK1 | 7.213825514 | 1.410351207 | 0.000512858 | 0.002306694 |
| GAS1 | 7.212760828 | 5.39672169 | 4.99E-10 | 1.17E-08 |
| LOC119867027 | 7.209476981 | -0.764138313 | 6.80E-05 | 0.000408809 |
| TSLP | 7.205705626 | -0.143812519 | 8.18E-07 | 8.57E-06 |
| PLXNA2 | 7.20409823 | 5.750491013 | 6.63E-10 | 1.52E-08 |
| LOC106558935 | 7.203793672 | -0.917590745 | 8.38E-05 | 0.000491322 |
| TRNP1 | 7.20175697 | 3.08965201 | 4.87E-11 | 1.41E-09 |
| LOC119865360 | 7.199866086 | -0.574582902 | 4.06E-07 | 4.66E-06 |

| | | | | |
|---|---|---|---|---|
| LOC106557970 | 7.19795727 | -0.775778877 | 2.00E-05 | 0.000141135 |
| LOC106558822 | 7.194572514 | 0.021568105 | 0.000218974 | 0.001111411 |
| LOC119870432 | 7.194076181 | -0.571835018 | 5.51E-05 | 0.000341743 |
| LOC106559361 | 7.192674056 | 0.94065133 | 3.96E-06 | 3.40E-05 |
| LOC111096181 | 7.190719135 | -0.440476489 | 0.000173537 | 0.00090913 |
| LOC111097304 | 7.186414708 | -0.564745109 | 0.004042387 | 0.013400222 |
| NHS | 7.185229971 | 4.984222543 | 2.32E-09 | 4.68E-08 |
| FOXO6 | 7.18403518 | 2.66790923 | 1.80E-06 | 1.72E-05 |
| PPP2R2C | 7.182166599 | 2.143117163 | 0.000174884 | 0.000915745 |
| LOC111095919 | 7.179254034 | -0.372184759 | 0.001230461 | 0.004885997 |
| ARHGEF15 | 7.174523196 | 5.167770677 | 1.65E-12 | 6.68E-11 |
| AVPR1A | 7.173688767 | 0.566801435 | 0.000250183 | 0.001243059 |
| PXDN | 7.171978404 | 7.784817266 | 3.99E-08 | 5.97E-07 |
| LOC111092500 | 7.170551653 | -0.258357271 | 0.000154533 | 0.000822897 |
| LOC119874670 | 7.170487642 | 5.624299747 | 1.49E-07 | 1.93E-06 |
| LOC119872628 | 7.169960978 | 3.960652123 | 1.08E-05 | 8.17E-05 |
| PCBP3 | 7.166748218 | 2.619644403 | 4.95E-07 | 5.54E-06 |
| LOC111097653 | 7.163985365 | 1.711078485 | 9.47E-06 | 7.27E-05 |
| LOC119873290 | 7.159706292 | 0.203266309 | 3.19E-06 | 2.83E-05 |
| LOC111095994 | 7.159425415 | 0.359495574 | 0.000670582 | 0.002906703 |
| WASF3 | 7.158994613 | 5.694531215 | 3.71E-17 | 3.51E-15 |
| SORCS2 | 7.153131906 | 6.148852936 | 3.02E-08 | 4.66E-07 |
| LOC484897 | 7.152860996 | 4.934366438 | 5.19E-13 | 2.35E-11 |
| LOC111093501 | 7.14838998 | -0.753183463 | 6.78E-05 | 0.0004081 |
| TMEM271 | 7.141511225 | -0.166350289 | 0.000839519 | 0.00352341 |
| LOC119866384 | 7.140562711 | -0.433090118 | 6.93E-06 | 5.52E-05 |
| NFATC4 | 7.139539754 | 4.778764407 | 2.19E-11 | 6.90E-10 |
| WNT7B | 7.135827462 | 1.23075448 | 0.000317903 | 0.001530412 |
| GPR37 | 7.135331593 | 0.757374173 | 0.002806235 | 0.009844534 |
| MIR126 | 7.134920775 | 0.940752032 | 9.13E-05 | 0.000528457 |
| BSND | 7.130092844 | 1.469889683 | 4.93E-06 | 4.10E-05 |
| SHROOM2 | 7.121319398 | 5.587981665 | 5.76E-12 | 2.12E-10 |
| FZD2 | 7.117476953 | 4.96203425 | 7.42E-11 | 2.08E-09 |
| LOC111090330 | 7.114820032 | -0.527104624 | 5.42E-06 | 4.45E-05 |
| LOC111092491 | 7.114648511 | -0.422665879 | 3.65E-05 | 0.00023893 |
| LOC102153011 | 7.114271956 | 1.100872298 | 3.40E-06 | 2.99E-05 |
| LOC102155852 | 7.112061718 | 2.94506593 | 1.14E-08 | 1.93E-07 |
| LMTK3 | 7.111532328 | 4.965754155 | 2.52E-05 | 0.000173154 |
| LOC119874665 | 7.111026005 | 0.778614999 | 5.28E-05 | 0.000329655 |

| | | | | |
|---|---|---|---|---|
| CEP126 | 7.109615792 | 2.110374873 | 9.33E-09 | 1.62E-07 |
| SORCS3 | 7.105241198 | 1.638209223 | 0.000668636 | 0.002898848 |
| LOC111090277 | 7.100817106 | -0.606311057 | 0.006504048 | 0.020025237 |
| HTRA1 | 7.098697874 | 7.028802388 | 1.88E-17 | 1.86E-15 |
| ELAVL3 | 7.097217583 | 5.227685415 | 4.98E-05 | 0.000313077 |
| ADAM18 | 7.087931955 | -0.904752663 | 9.19E-06 | 7.08E-05 |
| LYVE1 | 7.087405498 | 3.954120732 | 6.63E-07 | 7.14E-06 |
| LOC119866743 | 7.087097259 | -0.793798573 | 8.19E-06 | 6.40E-05 |
| TUBB3 | 7.086250702 | 4.820977765 | 1.98E-06 | 1.86E-05 |
| RIMKLA | 7.086145229 | 1.383366223 | 5.01E-09 | 9.23E-08 |
| TNNI1 | 7.086035638 | 5.439194872 | 6.28E-07 | 6.81E-06 |
| LOC119867339 | 7.085285947 | -0.144428939 | 0.000423583 | 0.00195993 |
| LHX8 | 7.082217965 | 1.414790997 | 0.005835115 | 0.018277851 |
| LOC102152925 | 7.077787996 | -0.902501639 | 4.26E-05 | 0.000272723 |
| TPM2 | 7.077408012 | 5.820110639 | 1.20E-08 | 2.02E-07 |
| FLNC | 7.077025778 | 7.386493188 | 3.93E-17 | 3.70E-15 |
| LOC111097507 | 7.063991444 | -0.681991238 | 0.000464942 | 0.002118287 |
| HOXA9 | 7.062727121 | 2.535195578 | 0.00014113 | 0.000762959 |
| LOC111093285 | 7.05970341 | -0.383733771 | 5.52E-05 | 0.000342222 |
| LOC119865348 | 7.050254838 | 1.256975774 | 0.000222538 | 0.00112601 |
| PER2 | 7.049760625 | 6.189458935 | 8.23E-13 | 3.58E-11 |
| LOC119879231 | 7.048770945 | -1.007282743 | 0.000332442 | 0.001588057 |
| SDK2 | 7.045563885 | 4.284501678 | 0.000216803 | 0.001101963 |
| LOC119877836 | 7.043688115 | 3.166526421 | 7.12E-10 | 1.62E-08 |
| LOC102155027 | 7.040419915 | 5.248353226 | 1.54E-07 | 1.98E-06 |
| LOC119869228 | 7.04017788 | -0.219777479 | 0.000386317 | 0.001808737 |
| LOC111090458 | 7.037783591 | -0.967440687 | 3.20E-05 | 0.000212811 |
| LOC111098563 | 7.029138851 | 4.859671774 | 0.003847757 | 0.012859282 |
| LOC102154971_1 | 7.026681761 | 0.458961189 | 1.16E-09 | 2.52E-08 |
| PLPP4 | 7.019217006 | -0.614511447 | 0.000588514 | 0.002592979 |
| SLITRK5 | 7.018466771 | 3.61752946 | 8.95E-05 | 0.000519267 |
| LOC102157245 | 7.013474016 | -0.610361127 | 0.000743853 | 0.003172905 |
| ATOH8 | 7.011015141 | 4.213815309 | 3.52E-14 | 1.97E-12 |
| TCHHL1 | 7.005859989 | -0.321453757 | 7.67E-05 | 0.000455187 |
| EPHB4 | 7.004331686 | 5.851962316 | 2.31E-16 | 1.92E-14 |
| LOC119872535 | 7.004233216 | -0.084670766 | 0.00136164 | 0.005320141 |
| SHISA3 | 6.997549654 | 5.939427003 | 6.17E-05 | 0.000376856 |
| LRRIQ1 | 6.996396664 | 0.813393816 | 4.10E-05 | 0.000263944 |
| LOC111096455 | 6.996205999 | -0.905970361 | 0.000225878 | 0.001140067 |

| | | | | |
|---|---|---|---|---|
| C1QTNF5 | 6.993685867 | 3.109805543 | 5.37E-12 | 1.99E-10 |
| ABI3BP | 6.990848702 | 9.194810897 | 3.45E-09 | 6.60E-08 |
| SCARA5 | 6.986597876 | 6.080409667 | 2.82E-08 | 4.39E-07 |
| LOC111093656 | 6.98636303 | -0.69621775 | 1.43E-05 | 0.000104637 |
| KL | 6.985682557 | -0.574948208 | 3.34E-06 | 2.94E-05 |
| LOC119876904 | 6.982383419 | 1.054476728 | 1.51E-05 | 0.000109873 |
| LOC111090994 | 6.981461237 | -0.849750361 | 1.24E-05 | 9.22E-05 |
| SHROOM4 | 6.980580083 | 5.343578602 | 5.40E-08 | 7.85E-07 |
| LOC119878272 | 6.977387996 | 7.04681938 | 4.70E-07 | 5.30E-06 |
| AATK | 6.977213586 | 7.176346449 | 2.20E-12 | 8.78E-11 |
| MIR8888 | 6.971757193 | -0.661541607 | 0.002104096 | 0.007696552 |
| LOC111096950 | 6.970173252 | -0.103970226 | 0.000264952 | 0.001305307 |
| EPB41L1 | 6.9685166 | 5.91682212 | 5.46E-15 | 3.57E-13 |
| LEP | 6.967202574 | 1.371028886 | 0.000267881 | 0.001317702 |
| LOC111090956 | 6.963624283 | -0.811616 | 2.63E-05 | 0.000179146 |
| LOC119872605 | 6.955419487 | 5.086395112 | 5.03E-09 | 9.25E-08 |
| LOC106559890 | 6.954685863 | 4.518489845 | 5.65E-16 | 4.45E-14 |
| MEX3A | 6.949618546 | 2.495373086 | 3.88E-09 | 7.32E-08 |
| ADM | 6.946967385 | 4.091720644 | 2.67E-11 | 8.23E-10 |
| PNCK | 6.94660384 | 0.421394221 | 0.000732885 | 0.00313105 |
| EGFR | 6.942709574 | 8.191356449 | 5.04E-05 | 0.000316764 |
| SYNPR | 6.934967425 | 0.26328628 | 0.00055573 | 0.002470127 |
| LOC111097750 | 6.933672494 | 0.152828062 | 0.002064106 | 0.007568156 |
| LOC119869700 | 6.933031226 | -0.648292943 | 6.06E-05 | 0.000370868 |
| ERICH3 | 6.932664459 | 1.076303631 | 0.000198379 | 0.001019797 |
| LOC119867204 | 6.93190212 | -0.889131706 | 0.000278313 | 0.001361598 |
| MATN4 | 6.931889958 | 3.843644676 | 2.45E-09 | 4.90E-08 |
| MAPK8IP2 | 6.929532879 | 6.523963395 | 8.89E-05 | 0.00051623 |
| LOC111093170 | 6.92929917 | -0.789082517 | 0.000171751 | 0.000901509 |
| LOC106560172 | 6.925802349 | -0.939512932 | 3.74E-05 | 0.00024358 |
| LOC119867505 | 6.919994627 | 0.838375143 | 1.97E-08 | 3.16E-07 |
| DPYSL3 | 6.919164711 | 8.621704862 | 1.02E-07 | 1.38E-06 |
| LOC111093600 | 6.911892011 | 0.105675946 | 1.32E-05 | 9.75E-05 |
| RAI14 | 6.90224079 | 6.801619193 | 3.19E-12 | 1.25E-10 |
| LOC102153282 | 6.901842114 | -0.636506418 | 0.005222511 | 0.016593973 |
| HOXC6 | 6.900118629 | 5.420321648 | 2.64E-07 | 3.19E-06 |
| LOC119872736 | 6.897814279 | 0.125445797 | 1.58E-07 | 2.02E-06 |
| ADAMTS12 | 6.896784216 | 5.281735333 | 6.20E-07 | 6.75E-06 |
| OSMR | 6.893733282 | 6.793667589 | 1.11E-15 | 8.28E-14 |

| | | | | |
|---|---|---|---|---|
| LOC111092354 | 6.893280295 | 0.229931881 | 0.000633845 | 0.002767388 |
| ADCY1 | 6.887057415 | 1.391834198 | 0.000271896 | 0.001335027 |
| GNG12 | 6.886202127 | 5.993255824 | 7.81E-22 | 1.80E-19 |
| ISLR2 | 6.882906555 | 4.091140759 | 1.50E-06 | 1.47E-05 |
| LOC119871913 | 6.873160654 | -0.60353439 | 1.11E-05 | 8.38E-05 |
| TLL1 | 6.87037517 | 4.838586464 | 7.00E-07 | 7.49E-06 |
| LOC111092581 | 6.86688981 | -0.557504513 | 0.000104211 | 0.000593578 |
| LOC100855970 | 6.866601111 | 4.199088293 | 5.78E-09 | 1.05E-07 |
| TMEM98 | 6.860533401 | 5.400413323 | 2.32E-09 | 4.68E-08 |
| MTARC1 | 6.856156731 | 4.737126905 | 7.04E-06 | 5.59E-05 |
| CDR2L | 6.855024479 | 5.779085522 | 6.43E-15 | 4.15E-13 |
| PCSK2 | 6.854050265 | 0.026833314 | 2.67E-05 | 0.000181407 |
| LOC119878909 | 6.853738816 | -0.608707488 | 0.001589358 | 0.006054869 |
| LOC111093519 | 6.850420178 | -0.7646738 | 5.43E-05 | 0.000337583 |
| ALX4 | 6.848911615 | 5.173182365 | 1.01E-06 | 1.03E-05 |
| LOC119873384 | 6.848338837 | 2.77679561 | 2.19E-08 | 3.49E-07 |
| CHRD | 6.846251691 | 3.035351502 | 3.42E-12 | 1.32E-10 |
| PI16 | 6.843218367 | 8.247775947 | 6.06E-06 | 4.91E-05 |
| UNC5D | 6.843176202 | 2.340243025 | 5.48E-06 | 4.49E-05 |
| LDB2 | 6.842187134 | 4.859166553 | 4.07E-08 | 6.08E-07 |
| LOC111095255 | 6.840742474 | -0.6954496 | 0.000180426 | 0.00093992 |
| AFDN | 6.835274886 | 8.206569597 | 1.59E-12 | 6.50E-11 |
| CD93 | 6.835148644 | 7.897984259 | 9.51E-16 | 7.18E-14 |
| LOC119879272 | 6.834621 | 10.62049532 | 1.20E-05 | 8.97E-05 |
| SLCO2A1 | 6.834349609 | 4.595465318 | 5.82E-09 | 1.06E-07 |
| CDON | 6.832822972 | 6.185150558 | 1.12E-11 | 3.78E-10 |
| CYP1B1 | 6.831179714 | 6.362208718 | 2.44E-10 | 6.12E-09 |
| SMTN | 6.830375512 | 7.044946436 | 1.01E-13 | 5.17E-12 |
| LOC119878319 | 6.815283562 | 8.484144977 | 0.000137284 | 0.00074608 |
| POMGNT2 | 6.814248056 | 3.565114143 | 2.45E-12 | 9.70E-11 |
| LOC102156927 | 6.811355931 | -0.647693157 | 0.000969788 | 0.003987161 |
| LOC111091270 | 6.808954363 | -0.756228026 | 0.000428692 | 0.001979342 |
| HOXD9 | 6.808050245 | 1.766408055 | 1.59E-09 | 3.32E-08 |
| LOC102152004 | 6.807552016 | -0.376151723 | 0.000375125 | 0.001764341 |
| ADRA2C | 6.80659304 | 0.857432569 | 6.04E-05 | 0.000370203 |
| LOC119864501 | 6.800864105 | 0.103267543 | 0.00072452 | 0.003100202 |
| LOC111094359 | 6.796937057 | 0.403755028 | 0.012986469 | 0.035899968 |
| DCC | 6.796207002 | 0.45129031 | 0.003293993 | 0.011260365 |
| LOC111098256 | 6.795903938 | -0.258672876 | 1.09E-07 | 1.47E-06 |

| | | | | |
|---|---|---|---|---|
| NEGR1 | 6.795749702 | 0.545801481 | 1.57E-05 | 0.000113887 |
| C2H10orf67 | 6.794840029 | -0.597276171 | 4.35E-05 | 0.000277806 |
| KBTBD11 | 6.79337084 | -0.62831228 | 0.000112391 | 0.000632525 |
| MYO1D | 6.79328641 | 6.851834285 | 1.88E-16 | 1.62E-14 |
| PTCH2 | 6.789702022 | 6.451342208 | 2.59E-08 | 4.06E-07 |
| PGR | 6.787199453 | -0.906431965 | 0.000211988 | 0.001081292 |
| ENPEP | 6.784564982 | 3.193508844 | 1.45E-06 | 1.43E-05 |
| DNAAF2 | 6.784012566 | 8.479973364 | 0.000292734 | 0.001426035 |
| LOC102151917 | 6.778377045 | 1.543965845 | 1.10E-12 | 4.67E-11 |
| PCSK5 | 6.774887204 | 5.545219444 | 4.77E-13 | 2.18E-11 |
| LOC609770 | 6.770901461 | 0.909882918 | 0.000145855 | 0.000784185 |
| ADRA2A | 6.770369095 | 5.536286982 | 1.93E-08 | 3.11E-07 |
| SCN8A | 6.770111002 | 3.325031188 | 8.66E-08 | 1.19E-06 |
| DBH | 6.76603791 | -1.135849839 | 0.001612421 | 0.006127651 |
| FGFRL1 | 6.765415724 | 3.438687364 | 1.51E-08 | 2.49E-07 |
| YPEL4 | 6.764085984 | 4.474767133 | 2.51E-09 | 5.00E-08 |
| SSC5D | 6.755524616 | 6.363949917 | 5.78E-09 | 1.05E-07 |
| LOC102156389 | 6.75501661 | 1.280246136 | 0.000599081 | 0.002634717 |
| HMCN2 | 6.751497756 | 6.545412395 | 3.87E-06 | 3.34E-05 |
| LOC119870691 | 6.75137692 | -0.999063163 | 0.000451675 | 0.002066965 |
| CTNNAL1 | 6.748845169 | 6.201509014 | 1.30E-08 | 2.17E-07 |
| LOC106559843 | 6.747372369 | -0.766847835 | 0.006710378 | 0.020576404 |
| LOC119872843 | 6.746634231 | -0.497623391 | 3.97E-06 | 3.41E-05 |
| KCNJ9 | 6.742490298 | 2.767627836 | 0.000114191 | 0.000640823 |
| LOC111096802 | 6.741089951 | 0.890258276 | 1.59E-06 | 1.54E-05 |
| LGI1 | 6.7372526 | 2.5515138 | 0.000508042 | 0.002290575 |
| LOC102152454 | 6.736973082 | -0.265879115 | 8.21E-05 | 0.000482922 |
| STK32C | 6.732287835 | 0.467618191 | 3.21E-05 | 0.000213221 |
| CLBA1 | 6.729924624 | 1.762230813 | 3.22E-08 | 4.94E-07 |
| WFDC1 | 6.728067732 | 4.085026161 | 1.06E-09 | 2.31E-08 |
| GAREM1 | 6.727720313 | 5.06940461 | 1.17E-11 | 3.92E-10 |
| C1GALT1C1L | 6.726283939 | -1.015621582 | 3.72E-05 | 0.000242873 |
| LOC111094605 | 6.723469151 | 5.238810954 | 7.35E-12 | 2.62E-10 |
| LOC119866403 | 6.719116676 | -0.971554936 | 0.000664108 | 0.0028821 |
| PRSS38 | 6.71909175 | 0.704602879 | 3.64E-08 | 5.52E-07 |
| DSPP | 6.717364676 | 2.666278807 | 0.000362282 | 0.001712112 |
| LOC119865115 | 6.714926351 | -0.564383049 | 6.18E-05 | 0.000377183 |
| MDGA1 | 6.714683399 | 4.813365531 | 2.87E-10 | 7.07E-09 |
| BEND6 | 6.714551524 | 2.41129249 | 1.33E-09 | 2.85E-08 |

| | | | | |
|---|---|---|---|---|
| HOXA3 | 6.709861439 | 2.457046044 | 2.43E-12 | 9.64E-11 |
| CCL21 | 6.70863872 | 2.018964644 | 0.000209069 | 0.001069677 |
| B3GAT1 | 6.707316485 | 2.156508971 | 0.001043596 | 0.004244295 |
| STMN2 | 6.701625092 | 5.758304083 | 0.001864483 | 0.006934675 |
| UACA | 6.688034833 | 6.072132645 | 5.31E-15 | 3.48E-13 |
| LOC102153673 | 6.685114153 | 0.165056324 | 0.000969885 | 0.003987161 |
| SOWAHA | 6.684372699 | 1.460621382 | 2.36E-05 | 0.000163414 |
| PAM | 6.6843612 | 7.239406091 | 4.48E-12 | 1.67E-10 |
| LOC487178 | 6.682179685 | 3.26653889 | 1.19E-06 | 1.19E-05 |
| DMP1 | 6.681717461 | 2.499075086 | 0.000420621 | 0.00194747 |
| ARHGEF10 | 6.677753507 | 7.623463858 | 1.19E-10 | 3.19E-09 |
| KRT23 | 6.676540873 | 0.505859105 | 0.000115539 | 0.000647548 |
| LRIG3 | 6.675918688 | 7.288777966 | 1.65E-11 | 5.34E-10 |
| LOC102157132 | 6.673610882 | -0.43490765 | 0.00022023 | 0.001115981 |
| WNT8B | 6.67007745 | -0.193694023 | 2.28E-05 | 0.000158274 |
| LOC102156514 | 6.666521135 | -0.778361006 | 0.000480078 | 0.002177633 |
| PARD3B | 6.663002895 | 6.506408233 | 1.10E-12 | 4.67E-11 |
| ARHGAP44 | 6.658222613 | 3.826534282 | 2.27E-11 | 7.11E-10 |
| LOC119867494 | 6.652769661 | 2.657018735 | 3.64E-05 | 0.0002379 |
| TSKU | 6.650946312 | 4.948175565 | 6.63E-08 | 9.43E-07 |
| FAM167A | 6.649782126 | 4.058198131 | 2.00E-05 | 0.000141135 |
| GCM2 | 6.6496174 | -1.024728287 | 0.000233979 | 0.001174664 |
| SARDH | 6.648453889 | 4.684045398 | 6.36E-08 | 9.09E-07 |
| LOC119864752 | 6.644463532 | 4.437460828 | 1.14E-08 | 1.93E-07 |
| RASSF10 | 6.642934657 | 1.297085248 | 0.000110049 | 0.000621767 |
| LOC477072 | 6.637942542 | 4.768475539 | 3.84E-05 | 0.000249203 |
| FMOD | 6.636836225 | 5.503722702 | 0.000241131 | 0.001204433 |
| ADD2 | 6.63467169 | 5.454217144 | 9.61E-05 | 0.00055228 |
| MSANTD3 | 6.634530524 | -0.598903935 | 0.001167877 | 0.00467863 |
| TRIM47 | 6.630733347 | 5.247674376 | 2.90E-12 | 1.14E-10 |
| TSPAN11 | 6.62938187 | 6.602390524 | 4.18E-13 | 1.92E-11 |
| LOC111096985 | 6.629255205 | 0.192427799 | 0.000323222 | 0.001552223 |
| PLEKHG4B | 6.625300358 | 1.813166058 | 0.00044973 | 0.002058935 |
| GDF10 | 6.624829803 | 1.835055943 | 0.002276059 | 0.008220135 |
| ALS2CL | 6.622289152 | 4.908144686 | 1.11E-10 | 2.99E-09 |
| REEP6 | 6.620033284 | 1.229463129 | 8.58E-05 | 0.000501322 |
| LOC611075 | 6.619405188 | 2.978385437 | 2.10E-12 | 8.41E-11 |
| KIRREL1 | 6.619313336 | 6.644107304 | 1.94E-17 | 1.91E-15 |
| LOC119876666 | 6.615247678 | 1.124968775 | 0.00099167 | 0.004065152 |

| | | | | |
|---|---|---|---|---|
| NDST4 | 6.612952482 | -0.583813499 | 0.00600858 | 0.018736588 |
| LOC119878702 | 6.612377385 | -0.978436851 | 0.000165605 | 0.000873913 |
| LOC119868427 | 6.611338844 | 1.542523321 | 5.40E-05 | 0.000335964 |
| LOC119876975 | 6.609317073 | 1.436864202 | 1.27E-10 | 3.38E-09 |
| LOC111093355 | 6.605834571 | -0.917481246 | 0.000104114 | 0.000593184 |
| STC1 | 6.605231675 | 3.853256406 | 1.69E-05 | 0.000121345 |
| C29H8orf88 | 6.603664002 | 1.882193754 | 5.86E-06 | 4.77E-05 |
| BCAN | 6.600824864 | -0.212757284 | 4.53E-05 | 0.000288091 |
| BICD1 | 6.600726639 | 5.93279941 | 3.77E-06 | 3.26E-05 |
| OAF | 6.597150235 | 7.020744263 | 1.56E-17 | 1.58E-15 |
| LOC119867489 | 6.590709428 | -0.489044591 | 0.000257595 | 0.001274045 |
| KCNA1 | 6.58913728 | 3.670470131 | 0.00055772 | 0.002477447 |
| LOC119865343 | 6.583350265 | -1.062171721 | 9.39E-05 | 0.000541346 |
| LOC106558668 | 6.582670162 | -0.98162831 | 0.001836451 | 0.0068505 |
| SBSN | 6.578048998 | 0.789305174 | 1.99E-05 | 0.00014046 |
| SNX33 | 6.57477699 | 6.004235921 | 2.43E-11 | 7.55E-10 |
| LRRC4C | 6.574502356 | 2.699063814 | 0.000591968 | 0.002606606 |
| LOC119877123 | 6.570969562 | 3.79846553 | 1.97E-09 | 4.04E-08 |
| RNASE4 | 6.569551205 | 3.735546401 | 9.53E-09 | 1.65E-07 |
| LOC111097003 | 6.568430109 | -0.639050982 | 4.64E-05 | 0.000294333 |
| PCDH18 | 6.565877661 | 5.109753588 | 6.13E-09 | 1.11E-07 |
| NEURL1B | 6.564502655 | 5.774017538 | 1.34E-08 | 2.23E-07 |
| LOC119870499 | 6.560972164 | 3.536990648 | 2.54E-10 | 6.32E-09 |
| LOC106557901 | 6.557807733 | 0.672122846 | 0.000518516 | 0.002330044 |
| PEX5L | 6.557113484 | 4.999605569 | 0.00358714 | 0.0121059 |
| ARHGAP32 | 6.556258618 | 6.430011703 | 6.18E-11 | 1.75E-09 |
| LOC106557652 | 6.554981381 | 2.115541629 | 5.78E-08 | 8.33E-07 |
| LOC102151347 | 6.554667344 | 2.227913867 | 8.30E-07 | 8.68E-06 |
| LOC119867508 | 6.549587104 | -0.770534738 | 0.000133362 | 0.000728233 |
| C18H11orf95 | 6.548977447 | 4.204054936 | 3.18E-10 | 7.80E-09 |
| LOC119870353 | 6.543546798 | -0.024353504 | 2.28E-06 | 2.10E-05 |
| CRTAC1 | 6.541084103 | 2.714068167 | 2.88E-10 | 7.09E-09 |
| GPIHBP1 | 6.537381435 | 2.024692695 | 1.73E-08 | 2.81E-07 |
| SLC22A17 | 6.536452514 | 6.16391165 | 1.24E-16 | 1.10E-14 |
| LOC119869366 | 6.533941479 | -1.050603358 | 0.00197223 | 0.007284276 |
| LOC111096489 | 6.531580129 | -1.247744029 | 0.002003091 | 0.007380651 |
| LOC119865691 | 6.528758883 | -1.314697633 | 0.001829784 | 0.006830333 |
| LOC119876392 | 6.526192881 | 3.662319151 | 1.46E-06 | 1.43E-05 |
| LOC119871824 | 6.525557325 | -0.989677851 | 3.77E-05 | 0.00024546 |

| | | | | |
|---|---|---|---|---|
| LOC487079 | 6.521835942 | -0.21332266 | 0.001007146 | 0.004120801 |
| LOC119871715 | 6.519245386 | -0.90918097 | 0.000405116 | 0.001886972 |
| FAM181B | 6.519071752 | 2.118208897 | 2.50E-07 | 3.04E-06 |
| LOC111095153 | 6.518743903 | -1.114573282 | 0.003076238 | 0.010632082 |
| LOC102157387 | 6.518575728 | -0.267257018 | 2.93E-05 | 0.000196818 |
| LOC119875968 | 6.515022486 | -0.185791382 | 0.000379449 | 0.001781202 |
| LOC102155832 | 6.51339368 | 1.090895091 | 3.03E-05 | 0.000202828 |
| LOC119878257 | 6.511157552 | 8.308502097 | 0.000146184 | 0.00078556 |
| MSC | 6.510525477 | 2.036494275 | 8.28E-07 | 8.66E-06 |
| LOC119864654 | 6.509606115 | 0.65543656 | 7.23E-06 | 5.72E-05 |
| LOC111097433 | 6.508940426 | -0.625171019 | 0.000201999 | 0.001037176 |
| PROKR1 | 6.506139166 | -0.367971597 | 0.001028811 | 0.004188881 |
| LOC102154778 | 6.504478026 | 3.688316585 | 0.003276581 | 0.011216766 |
| LOC119864492 | 6.504462899 | -1.136545168 | 0.000283474 | 0.001384657 |
| LOC111097630 | 6.502705353 | -0.542401434 | 0.000239235 | 0.001198277 |
| LOC111091635 | 6.499962995 | -0.573903047 | 0.000707557 | 0.003039017 |
| KRT27 | 6.499849522 | -0.63584008 | 0.008790199 | 0.025859914 |
| GHR | 6.497002828 | 6.042018649 | 4.16E-10 | 9.98E-09 |
| LOC119871912 | 6.496158616 | 1.420821144 | 8.56E-06 | 6.64E-05 |
| LOC119867310 | 6.491189406 | -0.629768585 | 0.000112595 | 0.000633509 |
| SHANK3 | 6.491039586 | 7.555992222 | 2.99E-09 | 5.83E-08 |
| KAZN | 6.488449382 | 4.776834169 | 1.62E-07 | 2.06E-06 |
| ROR2 | 6.487733958 | 5.334445583 | 7.82E-12 | 2.75E-10 |
| LOC111097308 | 6.486854739 | -0.727977032 | 8.02E-05 | 0.000473177 |
| LRP4 | 6.483848529 | 4.583107285 | 5.09E-09 | 9.35E-08 |
| FAM187A | 6.482809966 | 5.832838383 | 4.16E-05 | 0.000266973 |
| LOC119868394 | 6.476279537 | -0.925481246 | 0.002796673 | 0.009819172 |
| LOC119871988 | 6.475913789 | -0.299135389 | 0.000120464 | 0.000669957 |
| HRC | 6.474954558 | 1.573405776 | 1.28E-06 | 1.28E-05 |
| PIMREG | 6.473567375 | 0.707048267 | 4.20E-05 | 0.000269112 |
| HSPA12A | 6.470619998 | 4.735825743 | 3.99E-11 | 1.18E-09 |
| LAMC3 | 6.466740442 | 5.289137128 | 6.16E-05 | 0.000376483 |
| RET | 6.462835589 | 3.709116402 | 5.22E-13 | 2.36E-11 |
| CA9 | 6.462722323 | 1.944754393 | 3.65E-06 | 3.17E-05 |
| MXRA8 | 6.461775815 | 7.728456291 | 2.58E-12 | 1.02E-10 |
| EBF3 | 6.460945219 | 5.542240559 | 8.16E-10 | 1.83E-08 |
| GPC1 | 6.459548715 | 6.148136764 | 1.72E-09 | 3.55E-08 |
| FEZ1 | 6.45907668 | 3.625253886 | 5.00E-07 | 5.59E-06 |
| FXYD1 | 6.456316 | 5.752784161 | 2.29E-09 | 4.61E-08 |

| | | | | |
|---|---|---|---|---|
| RNF128 | 6.456264346 | 2.706676121 | 0.000210294 | 0.001074678 |
| GRIA1 | 6.450322019 | 4.463381935 | 1.54E-05 | 0.000112101 |
| CYP7B1 | 6.448289848 | 4.673211147 | 3.88E-08 | 5.83E-07 |
| ABCA9 | 6.448131116 | 7.055549397 | 3.76E-11 | 1.12E-09 |
| SNAI2 | 6.440571129 | 3.733707656 | 2.85E-08 | 4.43E-07 |
| HHATL | 6.438379374 | 0.629996214 | 0.000258272 | 0.001276557 |
| C1QTNF6 | 6.437589477 | 5.491513484 | 1.24E-11 | 4.12E-10 |
| NOS1 | 6.436049586 | 1.111081242 | 5.93E-05 | 0.000364325 |
| LOC119869621 | 6.432183076 | -0.306165339 | 0.000109231 | 0.000617785 |
| LOC119871827 | 6.431237755 | 1.977695668 | 1.22E-06 | 1.22E-05 |
| LOC119866734 | 6.430584762 | 5.233426822 | 5.98E-07 | 6.54E-06 |
| NLGN4X | 6.429551555 | 5.521383559 | 0.001363225 | 0.005324052 |
| THSD4 | 6.42933236 | 6.001465118 | 1.12E-05 | 8.42E-05 |
| LOC119865669 | 6.42714874 | -0.75527347 | 0.004865283 | 0.015630786 |
| LOC119864155 | 6.426719063 | 3.504402164 | 0.000197457 | 0.00101602 |
| LOC100686721 | 6.425131051 | -0.151149035 | 4.84E-05 | 0.000305622 |
| LOC106557985 | 6.422436957 | -0.913185839 | 0.001808549 | 0.006760381 |
| LOC119864667 | 6.42180536 | 3.799475348 | 2.08E-05 | 0.000145574 |
| ENAH | 6.415122401 | 6.830288752 | 1.95E-16 | 1.67E-14 |
| PTPN3 | 6.414278602 | 4.250574678 | 8.52E-10 | 1.90E-08 |
| SH3PXD2A | 6.411991486 | 8.806580235 | 3.03E-15 | 2.07E-13 |
| ADAMTS6 | 6.409848667 | 4.363785175 | 5.07E-08 | 7.45E-07 |
| LOC111096595 | 6.40702084 | -0.146968495 | 0.001776778 | 0.006653093 |
| LOC102151168 | 6.404787855 | 0.348015914 | 4.81E-06 | 4.02E-05 |
| LOC480601 | 6.404523531 | 3.663406658 | 0.001221628 | 0.004856264 |
| LOC111091878 | 6.402703539 | 0.221564487 | 0.001217464 | 0.004845261 |
| LOC111096335 | 6.401054863 | -0.898499849 | 0.006854251 | 0.020949397 |
| SH3RF2 | 6.398419261 | 2.749771972 | 3.99E-06 | 3.42E-05 |
| NFASC | 6.394583007 | 5.956922078 | 0.000117752 | 0.000657911 |
| CNTN5 | 6.39233857 | 0.715013182 | 0.008558567 | 0.025284873 |
| RGS11 | 6.390828245 | 6.449716739 | 9.92E-12 | 3.39E-10 |
| PRR18 | 6.387003377 | 0.902916592 | 2.15E-05 | 0.000149867 |
| RHOBTB3 | 6.383329824 | 5.003261112 | 1.95E-07 | 2.44E-06 |
| ELN | 6.374313803 | 6.208294681 | 3.59E-07 | 4.18E-06 |
| PARP3 | 6.373372554 | 4.42432317 | 1.75E-07 | 2.21E-06 |
| CCN5 | 6.371414773 | 4.708167811 | 9.90E-08 | 1.34E-06 |
| LOC119869725 | 6.371117922 | 0.352193689 | 4.64E-05 | 0.000294561 |
| NYAP1 | 6.36861314 | 5.738942846 | 8.46E-10 | 1.90E-08 |
| PDGFC | 6.368256711 | 4.689948579 | 5.94E-11 | 1.70E-09 |

| | | | | |
|---|---|---|---|---|
| LOC106557975 | 6.362154094 | 1.785333086 | 0.002408235 | 0.008617084 |
| LOC111094842 | 6.361002095 | -0.466082591 | 0.006568972 | 0.020197098 |
| LOC102157151 | 6.358951424 | -0.343734315 | 0.001524434 | 0.005847622 |
| GDA | 6.354485032 | 4.399284597 | 1.43E-11 | 4.68E-10 |
| LOC102153281 | 6.347103172 | -0.852021752 | 0.00041186 | 0.001914272 |
| LOC111091016 | 6.346586976 | -0.645958221 | 0.0002842 | 0.00138785 |
| TMEM116 | 6.34627173 | -1.236407613 | 0.000793984 | 0.003354389 |
| LOC479649 | 6.344327677 | 0.36749709 | 3.31E-08 | 5.06E-07 |
| PGM5 | 6.341183039 | 4.918797605 | 2.22E-06 | 2.06E-05 |
| LOC119866704 | 6.329063202 | -0.725700555 | 0.000294685 | 0.001433928 |
| LOC111090741 | 6.326218359 | 0.669176692 | 5.75E-05 | 0.000354906 |
| TTC12 | 6.322269822 | 1.297174965 | 5.89E-06 | 4.79E-05 |
| JAG2 | 6.321556952 | 4.873352275 | 4.50E-11 | 1.32E-09 |
| CLMP | 6.32114052 | 4.952647746 | 7.78E-06 | 6.12E-05 |
| TMEM9 | 6.317235575 | 4.592002802 | 7.24E-18 | 8.00E-16 |
| LOC106559289 | 6.31354072 | 7.044460262 | 5.53E-11 | 1.59E-09 |
| LOC119863960 | 6.311577198 | -0.894638313 | 3.68E-05 | 0.000240238 |
| LOC119872492 | 6.309080771 | 0.141910661 | 7.86E-07 | 8.28E-06 |
| GPR20 | 6.30619161 | -0.687926215 | 0.000502144 | 0.002267283 |
| LOC102154870 | 6.305634904 | 0.343001817 | 8.68E-06 | 6.72E-05 |
| LOC111092466 | 6.304148408 | 1.491575588 | 1.90E-07 | 2.38E-06 |
| LOC111093897 | 6.302763874 | -0.757799274 | 0.00057923 | 0.00255988 |
| LIPG | 6.301318359 | 0.090116614 | 7.96E-06 | 6.24E-05 |
| CPT1C | 6.298245802 | 4.359851564 | 1.23E-09 | 2.65E-08 |
| ART4 | 6.297869258 | 2.684897233 | 4.60E-06 | 3.88E-05 |
| LOC119868392 | 6.29571596 | 0.644898229 | 0.000960569 | 0.003956589 |
| LOC491316 | 6.293711841 | -0.133603217 | 0.000175932 | 0.000919875 |
| PRICKLE2 | 6.289972847 | 6.221973938 | 2.64E-11 | 8.16E-10 |
| HOXA11 | 6.289754234 | -0.038827217 | 0.002984226 | 0.010368248 |
| LOC111095647 | 6.283846093 | 2.761502966 | 1.64E-05 | 0.000118072 |
| CPNE8 | 6.282972784 | 3.258048974 | 4.47E-07 | 5.08E-06 |
| LOC119869241 | 6.28270448 | -0.570542932 | 0.002275495 | 0.008219469 |
| CTNNA3 | 6.282689728 | 5.004629639 | 3.46E-06 | 3.03E-05 |
| ADAMTS2 | 6.282287669 | 6.767427437 | 2.38E-09 | 4.76E-08 |
| LOC111094550 | 6.281369655 | -0.945777243 | 0.009508687 | 0.027636018 |
| LZTS1 | 6.278881297 | 3.526318377 | 4.40E-09 | 8.21E-08 |
| LOC119867829 | 6.27582806 | -0.217570221 | 0.000999608 | 0.004094595 |
| LOC102151393 | 6.27481938 | 3.395103291 | 9.29E-11 | 2.54E-09 |
| MIR99A-2 | 6.273543319 | -1.2375324 | 0.000242418 | 0.001209467 |

| | | | | |
|---|---|---|---|---|
| LOC119868670 | 6.272023579 | -0.890382787 | 0.000672877 | 0.002914294 |
| HECW1 | 6.270053999 | 2.130782681 | 0.002302649 | 0.008292674 |
| LOC102155178 | 6.262945692 | 1.471199352 | 0.000396678 | 0.001851252 |
| PTPRS | 6.260310184 | 8.34970993 | 1.47E-12 | 6.06E-11 |
| LOC489461 | 6.257466115 | -0.225767757 | 7.32E-06 | 5.79E-05 |
| RNASE9 | 6.256322499 | -0.753067102 | 0.000251122 | 0.001247439 |
| LOC102154320 | 6.253756551 | 0.832118841 | 1.52E-07 | 1.96E-06 |
| LRTM2 | 6.252384853 | -0.5451537 | 1.64E-07 | 2.09E-06 |
| EDN3 | 6.250815535 | 0.075544033 | 0.000104278 | 0.000593803 |
| ABCA8 | 6.250559781 | 7.822227557 | 8.63E-10 | 1.93E-08 |
| LOC119866715 | 6.249029938 | -1.382180788 | 0.002565083 | 0.009097209 |
| LOC111090243 | 6.248324844 | -0.420532362 | 0.003298837 | 0.011275144 |
| LOC111091089 | 6.245781182 | 0.050759809 | 0.000115686 | 0.000648207 |
| LOC111095831 | 6.245654538 | -0.441195187 | 0.002472795 | 0.008817521 |
| LOC111095197 | 6.245416487 | 0.840843936 | 1.34E-09 | 2.85E-08 |
| SLC9A3R2 | 6.241842919 | 6.289045171 | 1.89E-08 | 3.04E-07 |
| LOC102153923 | 6.241370646 | 1.134585602 | 1.61E-07 | 2.05E-06 |
| LOC119865207 | 6.237133677 | 0.054900316 | 0.000628054 | 0.00274598 |
| CTTN | 6.236797815 | 7.128184912 | 1.41E-15 | 1.02E-13 |
| FAM180A | 6.235370151 | 3.171202506 | 2.99E-06 | 2.68E-05 |
| LOC111095420 | 6.233099594 | -1.187367727 | 0.004046171 | 0.013408664 |
| NPY1R | 6.232589973 | 1.709722631 | 0.002134304 | 0.00778996 |
| VNN1 | 6.230446821 | 2.515618145 | 1.06E-07 | 1.43E-06 |
| LOC106558209 | 6.225693639 | -0.562469525 | 0.001011522 | 0.004134803 |
| FOXP2 | 6.224517412 | 2.539658706 | 8.27E-05 | 0.000485772 |
| ADIRF | 6.222558725 | 6.038223404 | 3.56E-08 | 5.41E-07 |
| LOC111095806 | 6.222444137 | -0.700209693 | 0.008235885 | 0.024468759 |
| LRP3 | 6.221668433 | 2.925397141 | 3.31E-07 | 3.90E-06 |
| LOC111094337 | 6.221150406 | -0.981086738 | 0.001383638 | 0.005388603 |
| FOXC2 | 6.220633343 | 0.244220935 | 0.002364634 | 0.008479272 |
| AXL | 6.220055081 | 6.569409598 | 8.65E-13 | 3.73E-11 |
| LRRN2 | 6.219258924 | 2.125733437 | 8.16E-06 | 6.38E-05 |
| MIR199-2 | 6.213408584 | -0.976614328 | 0.000442833 | 0.002032943 |
| LOC102151465 | 6.213328567 | -0.617789157 | 0.00034863 | 0.001655536 |
| CCDC24 | 6.210355735 | 0.787732712 | 3.22E-07 | 3.82E-06 |
| LOC111098840 | 6.209627861 | -0.062675936 | 0.002030093 | 0.007466182 |
| OVCH1 | 6.203940458 | -0.688340597 | 0.001159607 | 0.004651519 |
| LOC119871486 | 6.200402163 | -0.758848631 | 0.00241617 | 0.008638347 |
| CEP170B | 6.196865322 | 6.551995151 | 6.42E-08 | 9.16E-07 |

| | | | | |
|---|---|---|---|---|
| NXPH1 | 6.196010649 | -0.222571288 | 0.001780841 | 0.006666005 |
| LOC111096787 | 6.19278178 | -0.405288113 | 0.000960913 | 0.003957034 |
| SMPX | 6.190701227 | -1.41920382 | 0.004162894 | 0.013717865 |
| PPDPFL | 6.189844605 | 1.25445761 | 0.011685662 | 0.032831649 |
| PPP1R13L | 6.189464167 | 6.74368685 | 4.03E-07 | 4.64E-06 |
| PTX3 | 6.186646406 | 3.095360789 | 8.09E-05 | 0.000476302 |
| CAMK2A | 6.183059567 | 3.981828313 | 1.57E-06 | 1.52E-05 |
| MFAP5 | 6.180168429 | 6.345691052 | 0.000534474 | 0.00238937 |
| SP5 | 6.179603459 | 4.302693635 | 0.000150064 | 0.00080186 |
| LOC111095674 | 6.178183206 | 3.174101277 | 8.50E-12 | 2.96E-10 |
| NUPR1 | 6.173786189 | 6.650720266 | 1.98E-08 | 3.18E-07 |
| LOC111097888 | 6.173037378 | -0.834256337 | 0.000227903 | 0.001147881 |
| LAMA5 | 6.169090692 | 8.025357919 | 6.10E-13 | 2.73E-11 |
| ZCCHC24 | 6.16886022 | 8.119293648 | 1.75E-13 | 8.59E-12 |
| LOC111090938_2 | 6.168241988 | 1.088701087 | 5.23E-05 | 0.000326657 |
| C18H11orf96 | 6.163370423 | 1.697836549 | 1.55E-08 | 2.54E-07 |
| LOC119873278 | 6.1588677 | 3.889084473 | 1.95E-06 | 1.84E-05 |
| PLPPR2 | 6.156515604 | 3.839692672 | 1.63E-05 | 0.000117769 |
| LOC610160 | 6.153525649 | -0.631332799 | 0.000470162 | 0.002138471 |
| PCOLCE | 6.153478074 | 7.25930822 | 2.26E-13 | 1.09E-11 |
| SRGAP1 | 6.15239564 | 5.495403372 | 6.17E-15 | 3.99E-13 |
| FAM81A | 6.142307917 | 3.56700903 | 2.37E-07 | 2.90E-06 |
| ATP6AP1L | 6.141375041 | 0.655712489 | 9.77E-09 | 1.69E-07 |
| LOC111095673 | 6.134226381 | 2.188682448 | 5.55E-08 | 8.04E-07 |
| NFIX | 6.134073342 | 8.025895861 | 3.20E-12 | 1.25E-10 |
| SNPH | 6.131309663 | 2.614369058 | 3.88E-08 | 5.83E-07 |
| LOC119869870 | 6.130741267 | 0.210281085 | 0.00010677 | 0.000605947 |
| MINAR2 | 6.130524565 | -0.863626525 | 0.003254377 | 0.011154851 |
| LOC102151182 | 6.127633397 | 2.997482358 | 1.65E-05 | 0.00011898 |
| ANO1 | 6.127040758 | 3.340772337 | 4.82E-06 | 4.02E-05 |
| SRY | 6.125385788 | -0.753672395 | 0.011287382 | 0.031886265 |
| LOC111093375 | 6.125049547 | 0.873461001 | 1.40E-07 | 1.82E-06 |
| ADCY2 | 6.122842225 | 3.66154987 | 3.61E-05 | 0.000236162 |
| LOC119864041 | 6.122463165 | -0.433819894 | 0.001720317 | 0.006466254 |
| TMEM198 | 6.121771798 | 1.051375396 | 7.13E-06 | 5.66E-05 |
| LOC111094843 | 6.119863379 | -0.825020263 | 0.004596693 | 0.014909327 |
| LOC119867471 | 6.118302419 | 0.304114782 | 1.74E-05 | 0.000124726 |
| LOC102156272 | 6.118064293 | -0.114924286 | 0.001690145 | 0.006378973 |
| WNT2 | 6.114909267 | 3.822402232 | 5.81E-06 | 4.73E-05 |

| | | | | |
|---|---|---|---|---|
| LOC111098000 | 6.114647762 | -1.127649446 | 0.000412888 | 0.001918012 |
| TNS2 | 6.107357643 | 9.168436669 | 1.77E-12 | 7.15E-11 |
| LOC119867726 | 6.107040535 | -0.531673231 | 3.23E-07 | 3.82E-06 |
| LOC119870645 | 6.106891661 | -0.974135497 | 0.00106603 | 0.004324986 |
| LOC111091868 | 6.106782062 | 0.432010519 | 3.57E-05 | 0.000234116 |
| LOC111091947 | 6.106039031 | -0.855299896 | 0.001592238 | 0.006062706 |
| TREH | 6.104234714 | 6.620555106 | 3.77E-12 | 1.44E-10 |
| MTCL1 | 6.10313647 | 4.268952719 | 3.57E-06 | 3.11E-05 |
| CCDC180 | 6.102167786 | 2.978306525 | 0.000221411 | 0.001120653 |
| MAP1A | 6.101639936 | 7.978502816 | 3.30E-14 | 1.85E-12 |
| LOC106559274 | 6.100120867 | 0.009851664 | 0.000274804 | 0.001346909 |
| LOC102151699 | 6.098830839 | 3.147060642 | 5.25E-13 | 2.37E-11 |
| LOC102153897 | 6.090070363 | 0.416884496 | 0.000539401 | 0.002408414 |
| LOC102154646 | 6.086516438 | 1.782254544 | 4.44E-06 | 3.76E-05 |
| LEKR1 | 6.0821582 | 0.623205803 | 3.75E-08 | 5.66E-07 |
| ARL14EPL | 6.081963858 | -1.182981612 | 0.004514476 | 0.0146712 |
| ROR1 | 6.07993146 | 4.796122656 | 1.85E-14 | 1.08E-12 |
| LOC119871849 | 6.076640625 | 2.713930043 | 2.27E-07 | 2.79E-06 |
| LOC111095506 | 6.075939116 | -0.852238625 | 0.000235886 | 0.001182869 |
| FMN2 | 6.075001979 | 5.512327004 | 5.87E-05 | 0.00036131 |
| SOX11 | 6.06943406 | -1.087172885 | 0.007148699 | 0.021717656 |
| FSTL1 | 6.069252974 | 9.547172854 | 1.30E-14 | 7.81E-13 |
| TDRD9 | 6.061824227 | -0.954574368 | 0.000354452 | 0.00168039 |
| LOC119865128 | 6.060772559 | 0.994597332 | 5.19E-07 | 5.78E-06 |
| LOC111095809 | 6.057499161 | -1.286365312 | 0.000394196 | 0.001842842 |
| DUOX1 | 6.053312714 | 3.653538341 | 3.57E-06 | 3.12E-05 |
| LOC111091557 | 6.052615685 | 1.576229265 | 3.19E-07 | 3.79E-06 |
| LOC481674 | 6.050497195 | 1.545993625 | 4.86E-07 | 5.45E-06 |
| TPC3 | 6.050136474 | 4.112225148 | 1.13E-05 | 8.52E-05 |
| CADM3 | 6.049043294 | 5.508812285 | 1.33E-09 | 2.83E-08 |
| CLCA4 | 6.046378501 | 1.962821557 | 6.48E-07 | 7.00E-06 |
| TEX15 | 6.046261321 | 4.249050529 | 0.000369252 | 0.001740495 |
| LOC102155886 | 6.04218807 | 3.073237421 | 0.00248751 | 0.008865614 |
| LOC111098305 | 6.03973119 | 1.13835856 | 0.003686901 | 0.012402018 |
| LOC111092151 | 6.035074441 | -0.6914375 | 0.003177442 | 0.010932642 |
| BMP3 | 6.034433829 | 2.570293885 | 0.008236035 | 0.024468759 |
| LOC111094590 | 6.033849966 | 2.122979222 | 0.000102057 | 0.000582686 |
| LOC111098044 | 6.033586931 | -0.097482755 | 0.00025487 | 0.00126345 |
| LOC119864431 | 6.025999217 | -0.966223525 | 0.000655964 | 0.002850754 |

| | | | | |
|---|---|---|---|---|
| KY | 6.02158324 | -1.066830357 | 0.0009968 | 0.004084638 |
| LOC111091945 | 6.02082648 | -1.124967861 | 0.001450564 | 0.005599942 |
| ARMCX2 | 6.019865584 | 3.146397876 | 8.67E-06 | 6.72E-05 |
| LOC119868048 | 6.018089949 | -1.312286552 | 0.002308002 | 0.008309191 |
| LOC119872654 | 6.017098096 | -1.147641512 | 0.00765647 | 0.023037438 |
| MFAP2 | 6.012572448 | 4.868829219 | 8.97E-09 | 1.56E-07 |
| KCNK4 | 6.002661364 | 0.510839801 | 0.000134285 | 0.000732348 |
| SCRG1 | 6.000695283 | -0.361506755 | 0.002164699 | 0.007883636 |
| PALM | 6.000070181 | 6.896686969 | 4.57E-08 | 6.77E-07 |
| LOC111092014 | 5.998374156 | -0.192933256 | 0.002072416 | 0.007592205 |
| CYP2J2 | 5.993580934 | 4.09526309 | 9.05E-08 | 1.24E-06 |
| PTPRM | 5.991588243 | 6.961231743 | 7.43E-10 | 1.69E-08 |
| PPP1R1C | 5.99003941 | 2.020873667 | 0.000242152 | 0.001208417 |
| STAP2 | 5.987865803 | 1.496081848 | 6.44E-06 | 5.17E-05 |
| ATP1B2 | 5.986653711 | 2.712231008 | 6.25E-07 | 6.79E-06 |
| IRS1 | 5.984154132 | 7.269797944 | 1.80E-07 | 2.27E-06 |
| LOC111095006 | 5.983900824 | -0.351605409 | 0.000638969 | 0.002786952 |
| HOXC10 | 5.979777621 | 0.473106752 | 0.000382253 | 0.001792422 |
| LOC111092056 | 5.979024481 | -0.827367013 | 0.002058427 | 0.007549887 |
| PYCR1 | 5.976951147 | 5.40932033 | 5.52E-08 | 8.01E-07 |
| LOC102157297 | 5.975746334 | -0.680422934 | 0.000921193 | 0.003824716 |
| BCAM | 5.975705491 | 7.132547136 | 8.28E-11 | 2.29E-09 |
| KCNB1 | 5.97292262 | 3.336669655 | 4.73E-07 | 5.33E-06 |
| CA15 | 5.972109833 | 4.495624949 | 0.000162039 | 0.000858017 |
| ECM2 | 5.97196956 | 6.546893341 | 7.53E-15 | 4.77E-13 |
| SLC22A16 | 5.971745988 | 0.651517199 | 0.002769781 | 0.009735559 |
| LOC119877499 | 5.970967863 | 2.740745182 | 5.98E-06 | 4.85E-05 |
| FBLN7 | 5.965687514 | 3.805260109 | 9.20E-07 | 9.50E-06 |
| TMEM121 | 5.962402569 | 1.732601731 | 2.08E-08 | 3.33E-07 |
| LOC111092459 | 5.961825676 | 0.827566071 | 6.40E-09 | 1.15E-07 |
| CPZ | 5.959783411 | 3.020296002 | 0.000531216 | 0.002377744 |
| PCOLCE2 | 5.953349485 | 5.021578527 | 2.54E-09 | 5.04E-08 |
| LOC119873020 | 5.953110667 | -0.922686336 | 0.003354141 | 0.011438916 |
| LOC111092132 | 5.951837738 | 2.745194317 | 7.64E-11 | 2.14E-09 |
| LYPD6 | 5.949570804 | -0.021448509 | 0.002359287 | 0.008461498 |
| PTPRF | 5.946402107 | 6.100209661 | 2.02E-07 | 2.52E-06 |
| NOBOX | 5.946000461 | -0.522459615 | 1.68E-06 | 1.62E-05 |
| LOC119867479 | 5.942083208 | 0.1656263 | 0.000787543 | 0.003330422 |
| LOC119873896 | 5.935816563 | -0.647200354 | 0.001892596 | 0.007015263 |

| | | | | |
|---|---|---|---|---|
| VIPR2 | 5.935670837 | -0.486119255 | 0.006428553 | 0.019849839 |
| PRRG3 | 5.933088319 | 3.852082899 | 2.52E-07 | 3.06E-06 |
| LOC111098337 | 5.93119544 | -0.323513768 | 6.76E-07 | 7.26E-06 |
| LOC480441 | 5.930616882 | 4.562291445 | 8.48E-16 | 6.51E-14 |
| PCDH10 | 5.92878028 | 3.567478899 | 0.000246804 | 0.001229085 |
| KCNIP1 | 5.925472013 | -0.657649372 | 0.00078295 | 0.003313586 |
| LOC111092096 | 5.921827071 | -0.248070629 | 0.002667511 | 0.009422891 |
| HOXC9 | 5.921133916 | 2.438161803 | 0.000165966 | 0.000875178 |
| LOX | 5.920889606 | 6.008191508 | 6.20E-07 | 6.75E-06 |
| LOC111095438 | 5.91615669 | -0.785999593 | 0.001448877 | 0.005595423 |
| SLC2A4 | 5.913379381 | 2.095869762 | 6.00E-09 | 1.09E-07 |
| CNIH3 | 5.913053764 | 0.322081275 | 3.32E-07 | 3.91E-06 |
| ENTPD2 | 5.912135387 | 4.776391173 | 1.49E-06 | 1.46E-05 |
| GALNT13 | 5.911614128 | -0.222585722 | 0.004051447 | 0.013417942 |
| LOC111096232 | 5.911056388 | -0.362800919 | 0.000681734 | 0.002946792 |
| LOC102153175 | 5.910461736 | 2.063374442 | 1.47E-05 | 0.000107497 |
| NOSTRIN | 5.908965637 | 4.113281636 | 1.20E-06 | 1.21E-05 |
| ICAM5 | 5.908754392 | 1.725444272 | 3.60E-06 | 3.13E-05 |
| RHPN1 | 5.906118791 | 3.70954323 | 3.11E-08 | 4.78E-07 |
| LOC102151934 | 5.9059932 | 0.020218274 | 0.003935516 | 0.013108103 |
| EBF1 | 5.905190403 | 7.653807977 | 1.05E-12 | 4.44E-11 |
| SCN9A | 5.904031199 | 5.370994443 | 8.94E-08 | 1.23E-06 |
| FA2H | 5.903616662 | 2.491315295 | 0.001066689 | 0.00432685 |
| CH25H | 5.90315198 | 3.315206769 | 1.35E-06 | 1.34E-05 |
| LOC102156763 | 5.902217388 | 0.441327324 | 0.002599267 | 0.009206393 |
| CDK18 | 5.897427654 | 3.029752393 | 1.91E-07 | 2.39E-06 |
| LOC119868092 | 5.896403515 | -0.863901576 | 5.11E-05 | 0.000320392 |
| EMP1 | 5.888841388 | 8.187568793 | 1.09E-10 | 2.94E-09 |
| IL11 | 5.887993168 | 1.971302493 | 1.77E-07 | 2.23E-06 |
| MCIDAS | 5.886868872 | -1.177353484 | 0.003242397 | 0.011117303 |
| LOC119865692 | 5.885164816 | -1.397790451 | 0.004044595 | 0.01340549 |
| UCN2 | 5.884709475 | 1.152959373 | 0.000147839 | 0.0007929 |
| TSPAN18 | 5.884236119 | 5.442300154 | 5.33E-08 | 7.77E-07 |
| LOC102155496 | 5.883526885 | 2.930258149 | 9.39E-06 | 7.21E-05 |
| PHACTR3 | 5.877099745 | -0.632131405 | 0.004798947 | 0.015447436 |
| STPG2 | 5.876037722 | -0.379753824 | 6.80E-05 | 0.000408809 |
| LOC119865623 | 5.871170146 | 1.492745604 | 0.012440962 | 0.034626039 |
| CPNE5 | 5.869505314 | 1.331181129 | 1.73E-05 | 0.000123786 |
| CTXND1 | 5.86918697 | -0.278421319 | 0.000570005 | 0.002525805 |

| | | | | |
|---|---|---|---|---|
| LOC102156339 | 5.865741649 | -0.2752226 | 0.003095908 | 0.010691143 |
| MEX3B | 5.85579629 | 3.821703809 | 5.85E-07 | 6.42E-06 |
| LOC102151686 | 5.854203033 | 1.893302876 | 4.81E-05 | 0.000304139 |
| LOC119864981 | 5.853718673 | 4.734821606 | 2.48E-13 | 1.19E-11 |
| LOC119879487 | 5.846803888 | 5.988671959 | 0.000578216 | 0.00255592 |
| KRT80 | 5.845596735 | 3.086574194 | 0.000673296 | 0.00291439 |
| VWF | 5.845110053 | 8.88160684 | 6.06E-07 | 6.61E-06 |
| EPHX2 | 5.840693131 | 3.321782295 | 8.88E-10 | 1.98E-08 |
| SLC6A1 | 5.840150188 | 2.834428421 | 0.000507932 | 0.002290556 |
| LOC111095136 | 5.840040821 | -1.399636417 | 0.009020877 | 0.026441639 |
| APBB2 | 5.839677837 | 7.812711714 | 2.09E-20 | 4.01E-18 |
| LOC119867515 | 5.839417277 | 4.560301634 | 2.85E-10 | 7.01E-09 |
| LOC102151152 | 5.837340178 | 3.051414051 | 0.000956359 | 0.00394164 |
| LOC119876957 | 5.832151021 | 5.578809284 | 1.91E-08 | 3.07E-07 |
| KLHL40 | 5.831417919 | 0.963813775 | 0.000418112 | 0.001939294 |
| LOC111098861 | 5.828739969 | 2.424683865 | 2.46E-09 | 4.91E-08 |
| LOC119876979 | 5.828614977 | -0.043061866 | 3.57E-06 | 3.11E-05 |
| ACP7 | 5.82707018 | 1.390618414 | 0.001144326 | 0.004598736 |
| LOC119870970 | 5.825890536 | -0.228264562 | 0.00038692 | 0.00181078 |
| CLIC3 | 5.825730159 | 3.312332235 | 5.44E-05 | 0.000338235 |
| LOC111092192 | 5.825458633 | -1.217428761 | 0.001992928 | 0.0073482 |
| ME1 | 5.824502826 | 5.649994242 | 1.05E-06 | 1.07E-05 |
| LOC119867346 | 5.824092624 | -0.536759969 | 0.00689091 | 0.021040673 |
| LOC102157323 | 5.823610396 | 0.34880906 | 0.000293336 | 0.001428647 |
| LOC119867070 | 5.819072615 | 1.827303234 | 0.000584516 | 0.002577984 |
| CLU | 5.818396875 | 6.610382178 | 2.80E-07 | 3.36E-06 |
| IGDCC4 | 5.817850412 | 5.497598913 | 1.77E-05 | 0.000126542 |
| LOC119865942 | 5.809293584 | 4.272434935 | 3.28E-06 | 2.89E-05 |
| LOC111096642 | 5.805760479 | 3.383174212 | 7.00E-12 | 2.51E-10 |
| CES2 | 5.802185544 | 2.616568766 | 2.86E-08 | 4.45E-07 |
| IGSF5 | 5.798449322 | -0.724373078 | 0.004232466 | 0.013890102 |
| PCP4L1 | 5.792538782 | 2.323277912 | 5.54E-10 | 1.28E-08 |
| FGF5 | 5.787996257 | 1.416848094 | 0.014084965 | 0.038455965 |
| SCARF2 | 5.782409592 | 4.372672266 | 4.11E-08 | 6.13E-07 |
| FGF16 | 5.779597588 | -0.766992307 | 0.000472585 | 0.00214679 |
| SATB2 | 5.778356751 | 4.248145343 | 3.30E-07 | 3.89E-06 |
| SLC25A21 | 5.773536354 | 2.021341374 | 1.49E-11 | 4.88E-10 |
| ARFGEF3 | 5.771928044 | 5.205530609 | 8.48E-10 | 1.90E-08 |
| LOC111097002 | 5.768580771 | 0.418488683 | 3.56E-05 | 0.000233705 |

| | | | | |
|---|---|---|---|---|
| NOVA1 | 5.764858761 | 4.610506872 | 2.10E-07 | 2.60E-06 |
| LOC119866856 | 5.761504591 | -0.334409076 | 7.68E-07 | 8.14E-06 |
| CRYM | 5.759757747 | 2.470128957 | 0.00022798 | 0.001148004 |
| LPAR4 | 5.757925887 | -1.082625622 | 0.002290076 | 0.00826112 |
| LOC119872255 | 5.756132121 | -0.559372266 | 0.003937839 | 0.013113827 |
| HPSE2 | 5.75034092 | 1.036634093 | 0.004300635 | 0.014077581 |
| SDK1 | 5.746672591 | 5.997674 | 8.81E-07 | 9.14E-06 |
| LOC119866190 | 5.743293447 | -0.452996545 | 0.001956843 | 0.007232376 |
| KIAA1217 | 5.739987499 | 6.279290325 | 9.49E-16 | 7.18E-14 |
| CYS1 | 5.739243987 | 5.6455484 | 1.69E-14 | 9.98E-13 |
| LOC111094924 | 5.738943243 | 0.882509066 | 0.00014089 | 0.000762045 |
| HCN1 | 5.735587564 | 4.47231362 | 1.72E-05 | 0.000123456 |
| ARMCX4 | 5.732936178 | 3.721141829 | 3.97E-09 | 7.49E-08 |
| LOC111092958 | 5.732519052 | 2.359669291 | 0.001930266 | 0.007142677 |
| PLIN1 | 5.731422838 | 6.472353207 | 5.42E-05 | 0.000337317 |
| SOWAHC | 5.73113318 | 4.60995183 | 2.30E-11 | 7.19E-10 |
| NOTCH3 | 5.731047103 | 7.727929401 | 9.19E-11 | 2.52E-09 |
| TNFRSF11B | 5.730175798 | 1.800688071 | 2.41E-08 | 3.80E-07 |
| PLCD3 | 5.727480372 | 5.933248292 | 1.82E-17 | 1.81E-15 |
| LOC119875957 | 5.726573589 | 3.161366186 | 2.11E-06 | 1.97E-05 |
| LOC111098304 | 5.723299398 | 0.458918852 | 0.010362032 | 0.029713723 |
| LOC119872481 | 5.721123473 | -0.775840438 | 0.004012182 | 0.013322508 |
| LOC102153260 | 5.720808006 | -1.198591372 | 0.0079778 | 0.023863953 |
| GRIN2D | 5.715674047 | 2.383312572 | 2.73E-06 | 2.47E-05 |
| LOC111096987 | 5.713049942 | -1.303517588 | 0.00547744 | 0.017299994 |
| LOC106557863 | 5.70937064 | -0.223981626 | 0.002513058 | 0.008947842 |
| LOC119877935 | 5.708127387 | 1.443503371 | 2.62E-08 | 4.09E-07 |
| LYPD1 | 5.702745179 | -0.595635885 | 5.62E-05 | 0.000347222 |
| BAMBI | 5.700837843 | 3.909309306 | 0.001537433 | 0.005891227 |
| LOC102155545 | 5.698897921 | 1.728915769 | 0.000924084 | 0.003833783 |
| OBSL1 | 5.697591839 | 6.083201591 | 2.96E-07 | 3.54E-06 |
| ONECUT1 | 5.693843492 | -0.979510715 | 0.011890266 | 0.033302886 |
| CDCP1 | 5.693353795 | 2.638871891 | 0.003530081 | 0.011941229 |
| MS4A13 | 5.693352106 | 0.723426817 | 0.00036668 | 0.001729501 |
| CNTLN | 5.692789182 | 5.555783485 | 4.49E-12 | 1.67E-10 |
| LOC119872614 | 5.691815348 | 3.479191047 | 1.11E-06 | 1.12E-05 |
| LOC119864156 | 5.691439523 | 2.39628727 | 0.001026554 | 0.004181101 |
| CWH43 | 5.69129119 | 1.461988998 | 0.003540438 | 0.011972523 |
| FXYD3 | 5.690761971 | 2.2652062 | 9.77E-07 | 1.00E-05 |

| | | | | |
|---|---|---|---|---|
| LOC111095154 | 5.684125097 | -0.54861331 | 0.00018667 | 0.000967176 |
| SCUBE1 | 5.681177622 | 5.14929465 | 2.88E-07 | 3.45E-06 |
| LOC119871144 | 5.679361169 | 0.11251629 | 0.002201851 | 0.008001462 |
| LOC119868066 | 5.678655662 | 3.926441406 | 2.25E-08 | 3.57E-07 |
| LONRF2 | 5.673099878 | 2.829784027 | 3.01E-05 | 0.000202156 |
| DPF1 | 5.672539255 | 1.676952505 | 9.57E-06 | 7.33E-05 |
| GRIK3 | 5.672356981 | 1.621903528 | 0.00705243 | 0.021470334 |
| LOC119867860 | 5.669803319 | 1.306149743 | 9.89E-07 | 1.01E-05 |
| LIMCH1 | 5.666903838 | 6.506943806 | 3.26E-11 | 9.83E-10 |
| MSX1 | 5.666139346 | 2.460519672 | 1.87E-08 | 3.02E-07 |
| HAS2 | 5.665635155 | 2.881472677 | 3.04E-07 | 3.62E-06 |
| LOC119870901 | 5.66359576 | -0.242838967 | 3.38E-05 | 0.000223161 |
| FERMT1 | 5.656931457 | 1.133232433 | 4.26E-06 | 3.62E-05 |
| GJA8 | 5.656072655 | 1.372931901 | 0.000938927 | 0.003882726 |
| LOC100683963 | 5.655301079 | 2.782529848 | 1.27E-06 | 1.27E-05 |
| EYA2 | 5.652943964 | 3.281214189 | 4.37E-06 | 3.71E-05 |
| EFHD1 | 5.652091449 | 6.130892233 | 0.000714078 | 0.003063381 |
| LMOD1 | 5.650876681 | 6.419396272 | 8.08E-10 | 1.82E-08 |
| LOC119879242 | 5.649535377 | -0.547271394 | 3.25E-05 | 0.000215557 |
| ADAM12 | 5.64732927 | 6.680180638 | 1.35E-09 | 2.87E-08 |
| PACSIN3 | 5.640579368 | 3.913329443 | 2.92E-14 | 1.66E-12 |
| ARHGEF19 | 5.638039257 | 4.368656257 | 9.99E-08 | 1.35E-06 |
| LOC119876645 | 5.636580206 | 1.79756133 | 6.45E-08 | 9.20E-07 |
| LGALS12 | 5.635786011 | 3.134741922 | 0.000144159 | 0.000776626 |
| LOC119872577 | 5.633847434 | -1.065666601 | 0.001103012 | 0.004454178 |
| ABLIM3 | 5.633014625 | 6.353473905 | 3.45E-09 | 6.60E-08 |
| LOC119866292 | 5.630958517 | 6.935988908 | 1.41E-07 | 1.84E-06 |
| CDK14 | 5.629647421 | 6.308780599 | 1.37E-15 | 1.00E-13 |
| PAMR1 | 5.625511618 | 5.034704923 | 5.80E-06 | 4.73E-05 |
| EPHX4 | 5.623667189 | 2.615796943 | 0.002158448 | 0.007863515 |
| LOC111090936 | 5.621029101 | 1.408379641 | 2.04E-06 | 1.92E-05 |
| CACNA1E | 5.61761894 | 2.151671548 | 0.000191585 | 0.000988152 |
| NPR2 | 5.612915272 | 5.179049839 | 2.84E-13 | 1.34E-11 |
| LOC111093294 | 5.611958828 | -0.7134307 | 0.000317267 | 0.001529047 |
| CLEC11A | 5.608357085 | 4.869954852 | 9.89E-06 | 7.54E-05 |
| HGFAC | 5.607608452 | 2.893051971 | 8.42E-09 | 1.48E-07 |
| LOC102156402 | 5.601957781 | 1.579599936 | 3.13E-09 | 6.08E-08 |
| PCDH19 | 5.5985022 | 4.034785735 | 7.69E-09 | 1.36E-07 |
| CCDC160 | 5.59839743 | 0.084178265 | 0.001863036 | 0.006931276 |

| | | | | |
|---|---|---|---|---|
| LOC119878216 | 5.595629779 | 6.417280183 | 3.35E-06 | 2.96E-05 |
| C25H2orf72 | 5.59374729 | -0.841497971 | 0.010695088 | 0.030499294 |
| STRA6 | 5.59194897 | 4.527936334 | 6.34E-05 | 0.000385215 |
| LIMA1 | 5.591621206 | 6.871409385 | 6.21E-17 | 5.73E-15 |
| MUC1 | 5.589629589 | 1.639069495 | 0.001806091 | 0.006752355 |
| ZDHHC1 | 5.585901468 | 4.742644086 | 2.72E-11 | 8.36E-10 |
| LOC111097974 | 5.585365555 | 4.19768984 | 7.29E-08 | 1.02E-06 |
| LOC119877203 | 5.562645817 | 7.03336982 | 6.62E-06 | 5.30E-05 |
| SLC8A2 | 5.556562283 | 2.416009855 | 3.58E-05 | 0.0002348 |
| LOC119875907 | 5.555594552 | 8.269252041 | 7.16E-06 | 5.67E-05 |
| ANGPT4 | 5.546480027 | 6.596070558 | 2.94E-05 | 0.000197792 |
| FKBP7 | 5.544225722 | 2.673564396 | 1.16E-05 | 8.69E-05 |
| TMEM178A | 5.544132782 | 0.506341454 | 0.000754817 | 0.003212083 |
| LOC119872133 | 5.543655568 | 0.630782293 | 0.004535284 | 0.014729987 |
| KCNJ10 | 5.53738911 | 2.689710881 | 0.001837814 | 0.006853223 |
| LOC111094458 | 5.536531601 | 0.213573592 | 3.56E-05 | 0.000233599 |
| FNBP1L | 5.528902455 | 4.867032803 | 1.38E-07 | 1.80E-06 |
| MAPK12 | 5.526782855 | 0.209648312 | 0.000201254 | 0.001033597 |
| EFEMP2 | 5.518408091 | 5.181763247 | 1.19E-11 | 3.96E-10 |
| TNFAIP6 | 5.516982393 | 3.61220019 | 1.80E-05 | 0.000128528 |
| ZNF165 | 5.5161026 | -0.613806179 | 6.79E-07 | 7.28E-06 |
| LGR5 | 5.513820807 | 2.109224509 | 5.81E-06 | 4.73E-05 |
| LOC119877100 | 5.512557919 | -1.017034368 | 0.009964209 | 0.028778558 |
| C2CD2 | 5.509971002 | 4.616977962 | 8.85E-16 | 6.78E-14 |
| LOC484365 | 5.508700973 | 2.201313097 | 1.03E-06 | 1.05E-05 |
| EPPK1 | 5.505372368 | 5.619816659 | 0.000852379 | 0.003569091 |
| LOC111096354 | 5.502441354 | 2.562610416 | 0.000148878 | 0.000796501 |
| LOC119869835 | 5.500927237 | 2.306019594 | 1.07E-05 | 8.08E-05 |
| LOC111091987 | 5.499006322 | 1.082490757 | 2.17E-07 | 2.68E-06 |
| C3 | 5.491211786 | 8.335277951 | 6.64E-09 | 1.19E-07 |
| TMEM182 | 5.489990695 | 0.778947565 | 0.007886678 | 0.023618624 |
| GYG2 | 5.489666264 | 1.771599245 | 1.99E-05 | 0.000140419 |
| LOC111093264 | 5.489277709 | -0.570919373 | 2.54E-05 | 0.000173857 |
| LOC119879267 | 5.485794557 | 11.83848032 | 1.59E-05 | 0.00011466 |
| FHL5 | 5.484898787 | 4.466711129 | 0.000130054 | 0.000713585 |
| RNASE12 | 5.477748187 | -0.687824183 | 0.00081783 | 0.003440379 |
| KCNH2 | 5.477671044 | 3.091827922 | 5.72E-06 | 4.68E-05 |
| DENND2B | 5.474685506 | 5.905079899 | 1.17E-08 | 1.99E-07 |
| TIMD4 | 5.473929252 | 5.009266477 | 5.02E-07 | 5.61E-06 |

| | | | | |
|---|---|---|---|---|
| FST | 5.471234416 | 3.260711903 | 2.93E-08 | 4.54E-07 |
| NAV2 | 5.470724779 | 6.974910435 | 1.06E-10 | 2.86E-09 |
| TNKS1BP1 | 5.469740585 | 8.190951529 | 9.63E-11 | 2.62E-09 |
| LOC111098598 | 5.469694196 | 2.272666424 | 1.24E-08 | 2.08E-07 |
| CILP | 5.46868228 | 5.694122009 | 0.000215551 | 0.001096888 |
| HES2 | 5.464483864 | -0.612949799 | 0.00451089 | 0.014665152 |
| HOXB3 | 5.463595555 | 4.183661926 | 6.49E-11 | 1.83E-09 |
| KLK6 | 5.462007348 | 3.455719084 | 0.007045259 | 0.021451518 |
| STX1B | 5.460845724 | 2.458191697 | 3.28E-07 | 3.87E-06 |
| MEGF6 | 5.457372702 | 4.361913305 | 9.68E-10 | 2.13E-08 |
| MORC4 | 5.457273285 | 5.360683308 | 4.72E-09 | 8.73E-08 |
| NAV3 | 5.456455233 | 6.822864637 | 4.69E-11 | 1.37E-09 |
| SYNPO2 | 5.45336975 | 5.235408922 | 1.04E-08 | 1.79E-07 |
| AP3B2 | 5.449919446 | 4.098727685 | 0.00053661 | 0.002398423 |
| LOC119865671 | 5.44827045 | 2.134496087 | 0.000335457 | 0.001600345 |
| SNAP91 | 5.444312871 | 0.364892877 | 4.68E-07 | 5.29E-06 |
| LOC106557817 | 5.440172311 | 2.916130595 | 3.42E-07 | 4.01E-06 |
| ZNF25 | 5.439856171 | 1.294141715 | 9.05E-09 | 1.57E-07 |
| LOC119868442 | 5.439362356 | -0.946838634 | 0.00052924 | 0.002371841 |
| LOC111091560 | 5.438940924 | 2.39005018 | 2.14E-07 | 2.64E-06 |
| POU3F2 | 5.434597035 | 2.729351224 | 0.0029596 | 0.010301374 |
| NDRG2 | 5.432148973 | 6.317961334 | 5.88E-14 | 3.15E-12 |
| LOC111099012 | 5.430346727 | 1.44558595 | 2.98E-06 | 2.67E-05 |
| LOC102151868 | 5.428439912 | 0.278210049 | 0.001087536 | 0.00439906 |
| PEBP4 | 5.428270657 | 2.171043112 | 0.000219807 | 0.001114356 |
| SEMA3C | 5.425356733 | 6.457304527 | 4.44E-07 | 5.04E-06 |
| PARD3 | 5.420417206 | 6.990893253 | 4.87E-10 | 1.14E-08 |
| DIRAS1 | 5.417942896 | -0.169686887 | 1.16E-05 | 8.71E-05 |
| SASH1 | 5.415631 | 7.092104867 | 1.26E-15 | 9.19E-14 |
| NFIB | 5.411650987 | 7.91219864 | 2.03E-05 | 0.000142768 |
| FAXC | 5.408698473 | 3.232671362 | 4.61E-10 | 1.09E-08 |
| FLT4 | 5.405361671 | 5.92609397 | 7.11E-12 | 2.54E-10 |
| LOC102151706 | 5.403532572 | -0.039003573 | 0.003156722 | 0.010876889 |
| CHAD | 5.402319717 | 1.609319158 | 7.61E-08 | 1.06E-06 |
| TEX13B | 5.401657739 | 0.456397985 | 2.03E-06 | 1.91E-05 |
| LOC119877637 | 5.401253072 | 2.252169683 | 0.000371602 | 0.001749668 |
| CST6 | 5.398139185 | 2.123494953 | 0.000621085 | 0.002718252 |
| SLC16A9 | 5.397001639 | 1.737248887 | 0.002726468 | 0.009600445 |
| LOC100049001 | 5.396424645 | 1.111597298 | 0.012275007 | 0.034216878 |

| | | | | |
|---|---|---|---|---|
| PRDM6 | 5.392811378 | -0.047822189 | 0.000323069 | 0.001551836 |
| EPHB3 | 5.390091307 | 3.829519305 | 2.11E-11 | 6.68E-10 |
| LOC119875531 | 5.387030271 | -0.190022306 | 0.010211644 | 0.029364065 |
| LOC119870567 | 5.386186437 | 3.683071444 | 0.000119638 | 0.000666043 |
| LOC111091473 | 5.37578439 | -0.79708494 | 0.000149634 | 0.000799755 |
| CALCRL | 5.372132941 | 5.673066989 | 9.05E-10 | 2.01E-08 |
| ADAMTS16 | 5.370189029 | 1.732252289 | 0.000240922 | 0.001203664 |
| FAM186A | 5.367839441 | 2.404806942 | 2.85E-06 | 2.56E-05 |
| LOC119874624 | 5.360348165 | -0.316261442 | 2.33E-06 | 2.15E-05 |
| LOC119870595 | 5.358189524 | 6.654104689 | 5.00E-05 | 0.000314394 |
| LOC111092398 | 5.357710435 | 1.010235368 | 0.002684424 | 0.009469322 |
| TGFB1I1 | 5.357108573 | 5.811336096 | 1.32E-14 | 7.90E-13 |
| GABRA3 | 5.356428527 | 0.914202634 | 0.000568149 | 0.002518095 |
| COL15A1 | 5.354304247 | 7.721127803 | 2.12E-09 | 4.32E-08 |
| GLI1 | 5.353414483 | 7.200516391 | 4.58E-07 | 5.19E-06 |
| HOXA4 | 5.353225023 | 1.365084395 | 6.43E-07 | 6.95E-06 |
| FGFR1 | 5.349252829 | 7.076555699 | 4.65E-15 | 3.09E-13 |
| SERTM1 | 5.344742972 | 0.179233858 | 0.000351447 | 0.001667814 |
| RECK | 5.342967358 | 4.495911228 | 6.37E-12 | 2.31E-10 |
| MEIS3 | 5.341920333 | 3.323088839 | 3.85E-09 | 7.28E-08 |
| TTC23 | 5.336448138 | 3.9941024 | 2.64E-12 | 1.04E-10 |
| GIPC2 | 5.333644544 | -0.16365417 | 0.000119534 | 0.000665977 |
| ALK | 5.328753744 | 2.148530767 | 1.15E-06 | 1.16E-05 |
| LOC102155709 | 5.319051991 | 3.004069802 | 6.96E-10 | 1.59E-08 |
| LOC119866187 | 5.318161525 | 2.874124647 | 6.42E-07 | 6.95E-06 |
| LOC119866989 | 5.313827371 | 1.588732294 | 0.000118227 | 0.000660398 |
| PCDH1 | 5.312790692 | 6.10360395 | 1.19E-11 | 3.98E-10 |
| LOC111095843 | 5.311410373 | 1.235919818 | 0.000157702 | 0.000837715 |
| TBX2 | 5.309034371 | 4.930846967 | 3.51E-07 | 4.10E-06 |
| LOC119866551 | 5.302945767 | 1.54506625 | 1.89E-07 | 2.37E-06 |
| LOC111097095 | 5.301711456 | -0.713497389 | 0.012855295 | 0.03560547 |
| LOC119871534 | 5.297665315 | 2.494156109 | 6.64E-07 | 7.14E-06 |
| TPPP3 | 5.297362256 | 4.681296808 | 1.30E-09 | 2.79E-08 |
| HES1 | 5.295186344 | 4.594868008 | 1.01E-08 | 1.74E-07 |
| NPB | 5.293692029 | -0.385954301 | 5.80E-05 | 0.000357076 |
| LOC119878341 | 5.293473224 | 5.736777932 | 0.000709336 | 0.003045452 |
| DNAJB5 | 5.29254216 | 2.330487926 | 5.67E-07 | 6.26E-06 |
| ACSM5 | 5.289884002 | -1.029218949 | 0.003526979 | 0.011934461 |
| LOC111091867 | 5.28809546 | 0.352627348 | 6.42E-07 | 6.95E-06 |

| | | | | |
|---|---|---|---|---|
| LOC106559295 | 5.286108386 | 3.25654408 | 1.75E-06 | 1.68E-05 |
| VWA2 | 5.281349316 | 4.925740885 | 9.87E-08 | 1.34E-06 |
| RAB9B | 5.275350197 | 0.805012482 | 0.010691217 | 0.030492684 |
| CROCC2 | 5.2749874 | 3.619837092 | 3.96E-05 | 0.000256359 |
| METTL24 | 5.27461454 | 2.511843684 | 0.000241788 | 0.001207158 |
| LOC119873580 | 5.274495311 | 2.901861 | 1.18E-06 | 1.19E-05 |
| CALD1 | 5.271405161 | 8.957441859 | 4.65E-06 | 3.91E-05 |
| OMD | 5.270073504 | 5.089773994 | 4.24E-05 | 0.000271684 |
| ACKR2 | 5.269526425 | 4.665364531 | 0.000430157 | 0.001984841 |
| LOC111089996 | 5.263821868 | 0.815496326 | 0.00012438 | 0.000687677 |
| LOC119864345 | 5.258196474 | -0.252565826 | 0.000821522 | 0.0034539 |
| LOC119876292 | 5.258055628 | 4.387462406 | 5.80E-07 | 6.38E-06 |
| DISC1 | 5.257998727 | 4.445369469 | 5.71E-09 | 1.04E-07 |
| RUNX1T1 | 5.248144287 | 5.936560514 | 4.96E-08 | 7.30E-07 |
| HOXA5 | 5.246613178 | 3.042177281 | 5.53E-08 | 8.02E-07 |
| RBPJL | 5.24452489 | 1.630613211 | 4.85E-06 | 4.05E-05 |
| CD276 | 5.241685842 | 6.494496099 | 9.15E-15 | 5.66E-13 |
| DCLK3 | 5.24025895 | 2.179182814 | 0.000585527 | 0.00258192 |
| LOC119876035 | 5.239284952 | 1.312311735 | 7.04E-08 | 9.92E-07 |
| LAMC1 | 5.233217139 | 9.538705531 | 7.82E-15 | 4.91E-13 |
| LOC119878420 | 5.232983585 | 6.628647138 | 1.26E-05 | 9.36E-05 |
| MAP4K3 | 5.231253753 | 5.359727635 | 1.53E-16 | 1.34E-14 |
| TGFB2 | 5.226729967 | 5.869217862 | 2.32E-07 | 2.85E-06 |
| IL13RA2 | 5.225051426 | 1.855965959 | 4.74E-05 | 0.000299808 |
| LOC119874085 | 5.221809605 | 3.909289451 | 8.32E-06 | 6.48E-05 |
| RNF148 | 5.220698221 | 0.428776698 | 5.18E-06 | 4.28E-05 |
| LOC119870193 | 5.218703728 | 4.122958608 | 3.63E-06 | 3.16E-05 |
| LOC111092644 | 5.207526099 | 0.110429643 | 0.007577081 | 0.022836611 |
| ACVR1C | 5.205257324 | 2.517848546 | 0.004554544 | 0.014788108 |
| LOC111092834 | 5.203399135 | 0.205523154 | 2.47E-05 | 0.000170278 |
| SLC24A4 | 5.201599566 | -0.997555679 | 9.14E-06 | 7.04E-05 |
| LOC119879268 | 5.199246797 | 12.41180576 | 6.46E-05 | 0.00039204 |
| BSN | 5.197485655 | 5.644796095 | 3.38E-07 | 3.97E-06 |
| MAPK11 | 5.193904923 | 3.962211362 | 0.000110954 | 0.000626065 |
| LOC111090544 | 5.190714784 | -0.669390215 | 8.10E-06 | 6.34E-05 |
| LOC119870288 | 5.18953016 | 4.907136856 | 2.19E-07 | 2.70E-06 |
| ABLIM2 | 5.188329779 | 4.963337472 | 5.17E-06 | 4.27E-05 |
| LOC106558924 | 5.187930997 | -0.04359689 | 1.01E-05 | 7.69E-05 |
| ZNF300 | 5.184725907 | 1.176878372 | 3.43E-08 | 5.23E-07 |

| | | | | |
|---|---|---|---|---|
| SCG3 | 5.178479034 | 4.294117674 | 8.47E-08 | 1.17E-06 |
| PTGES | 5.178053798 | 2.958123343 | 7.71E-07 | 8.16E-06 |
| DIPK2B | 5.176167956 | 4.489244243 | 3.63E-06 | 3.16E-05 |
| PTPN14 | 5.175467224 | 6.977333047 | 7.66E-13 | 3.35E-11 |
| LOC111093633 | 5.174171451 | -0.282797791 | 0.007520086 | 0.022690289 |
| OGDHL | 5.173094777 | 2.588105387 | 5.26E-05 | 0.000328072 |
| LOC111090631 | 5.171981301 | -0.782657193 | 0.006225301 | 0.019296473 |
| LOC119872477 | 5.170395475 | 5.489826152 | 7.78E-08 | 1.08E-06 |
| HSF4_2 | 5.166900663 | 3.735320536 | 9.86E-07 | 1.01E-05 |
| LOC111095671 | 5.166523085 | 2.624079435 | 1.76E-05 | 0.000125611 |
| LOC111097510 | 5.163718229 | 1.022408568 | 0.000235647 | 0.001182215 |
| SEZ6L2 | 5.163030813 | 4.658950039 | 0.000410151 | 0.001907966 |
| STK33 | 5.159073168 | 3.03157507 | 8.60E-09 | 1.51E-07 |
| LOC106559496 | 5.157597881 | 3.742196828 | 2.14E-06 | 2.00E-05 |
| RTKN | 5.149834344 | 5.299231286 | 5.74E-13 | 2.57E-11 |
| CYP4B1 | 5.147208604 | 0.179778441 | 0.013204678 | 0.036414937 |
| LOC119876215 | 5.14357944 | 2.618363395 | 8.64E-06 | 6.70E-05 |
| ANGPTL2 | 5.142967461 | 7.418437272 | 5.30E-07 | 5.89E-06 |
| LOC111091415 | 5.136962107 | 1.423225853 | 0.001082499 | 0.004381141 |
| KAZALD1 | 5.136595662 | 3.881458114 | 0.000198024 | 0.001018698 |
| TSHZ2 | 5.136054383 | 6.875596891 | 1.12E-07 | 1.49E-06 |
| SMO | 5.134484102 | 4.613222688 | 7.10E-07 | 7.58E-06 |
| LOC111096657 | 5.131204234 | 2.070651523 | 1.00E-06 | 1.03E-05 |
| LOC111093200 | 5.13030709 | -0.020775099 | 7.54E-05 | 0.000449465 |
| COLCA2 | 5.127579221 | 0.635785091 | 0.000150215 | 0.000802272 |
| LOC102155413 | 5.124965051 | 2.51925311 | 0.000130846 | 0.000716855 |
| KCNH1 | 5.124293977 | 2.044043681 | 1.15E-06 | 1.16E-05 |
| LOC119872627 | 5.124191934 | 5.478624925 | 0.000108442 | 0.000613807 |
| TBC1D16 | 5.122245103 | 4.460192327 | 2.17E-06 | 2.01E-05 |
| KCNQ2 | 5.11912406 | 5.977580929 | 0.000647568 | 0.002820485 |
| SYT8 | 5.117786844 | 0.488177704 | 0.003494268 | 0.011851528 |
| ARHGAP29 | 5.112913336 | 6.803093267 | 6.98E-06 | 5.55E-05 |
| MGAT4F | 5.107283736 | 0.124006635 | 5.10E-06 | 4.22E-05 |
| LOC111091159 | 5.104970386 | -1.279485067 | 0.018520136 | 0.047863929 |
| ITGA3 | 5.104146747 | 5.340038427 | 1.58E-09 | 3.29E-08 |
| ERBB2 | 5.103077895 | 5.98922336 | 2.52E-10 | 6.29E-09 |
| PTPRK | 5.102507267 | 4.695239195 | 1.15E-06 | 1.16E-05 |
| CCDC159 | 5.096168968 | 4.304374393 | 0.000157599 | 0.000837372 |
| RADX | 5.095117208 | 3.51E-05 | 0.013923159 | 0.038100584 |

| | | | | |
|---|---|---|---|---|
| RAB27B | 5.09487844 | 1.681388027 | 3.75E-05 | 0.000244524 |
| LTBP2 | 5.093683872 | 7.091214566 | 1.20E-09 | 2.60E-08 |
| TRNASTOP-UCA | 5.092951398 | -0.484935429 | 8.51E-05 | 0.000498226 |
| LOC102152553 | 5.092152528 | -0.218995158 | 2.28E-05 | 0.000158274 |
| LOC100855618 | 5.091457846 | 3.203975342 | 2.90E-09 | 5.68E-08 |
| TIE1 | 5.083557856 | 5.228581022 | 1.20E-06 | 1.20E-05 |
| IGDCC3 | 5.083410773 | 1.754938046 | 5.18E-05 | 0.000323967 |
| PPFIA4 | 5.081552624 | 6.466217845 | 3.85E-08 | 5.79E-07 |
| ACP6 | 5.079009744 | 1.681466176 | 7.83E-07 | 8.25E-06 |
| GPR153 | 5.076786909 | 3.829174054 | 7.83E-07 | 8.25E-06 |
| EPHA5 | 5.076627206 | 1.697064786 | 0.005831698 | 0.01827507 |
| LOC111095248 | 5.074080807 | 2.276548349 | 0.001188028 | 0.004747076 |
| PHLDA3 | 5.072599785 | 6.431495066 | 1.64E-09 | 3.40E-08 |
| LRRC49 | 5.070588789 | 2.978421948 | 2.97E-09 | 5.80E-08 |
| KRT12 | 5.070176874 | 2.239830437 | 0.001393226 | 0.005419129 |
| FGD5 | 5.066249795 | 6.228740335 | 3.78E-09 | 7.17E-08 |
| LOC119864742 | 5.06459856 | -0.50481071 | 0.017739867 | 0.046255551 |
| CDKN1C | 5.056551832 | 4.64525215 | 4.31E-08 | 6.40E-07 |
| CHN1 | 5.053019553 | 4.887330175 | 8.97E-11 | 2.46E-09 |
| LOC119876123 | 5.052305314 | 3.919694775 | 0.00013132 | 0.000718713 |
| LOC102155424 | 5.050121008 | -0.656992634 | 0.002617193 | 0.009262317 |
| LOC119879270 | 5.047830393 | 9.055266567 | 0.003919336 | 0.013062244 |
| LOC111090863 | 5.044252762 | 4.509350275 | 0.00139064 | 0.005411013 |
| LOC102154586 | 5.043507704 | 1.211270784 | 0.000799414 | 0.003372071 |
| ARHGEF17 | 5.041596557 | 6.910530827 | 4.59E-10 | 1.09E-08 |
| LOC119876662 | 5.037745936 | 0.361475813 | 3.37E-05 | 0.000222269 |
| LOC111093520 | 5.030798487 | 1.827615528 | 0.015276129 | 0.041056492 |
| HS3ST1 | 5.023598709 | 1.293707995 | 2.49E-06 | 2.28E-05 |
| LOC119867305 | 5.022697307 | 1.545261667 | 5.08E-08 | 7.45E-07 |
| SPSB1 | 5.021105289 | 6.023075893 | 1.08E-08 | 1.84E-07 |
| GOLGA7B | 5.018014781 | 1.594784763 | 4.19E-07 | 4.79E-06 |
| DKKL1 | 5.012930845 | 0.71790489 | 0.000226056 | 0.001140701 |
| LOC119874830 | 5.012584414 | 2.620255447 | 4.93E-07 | 5.52E-06 |
| SHROOM1 | 5.011599532 | 4.458652811 | 1.76E-11 | 5.67E-10 |
| GLIS3 | 5.011456417 | 4.557041635 | 2.01E-07 | 2.50E-06 |
| SPRED1 | 5.007532101 | 5.6752833 | 4.31E-09 | 8.06E-08 |
| LOC119865042 | 5.00388856 | 4.218880365 | 1.10E-10 | 2.96E-09 |
| CYP21A2 | 4.999622991 | 5.649224828 | 2.64E-07 | 3.19E-06 |
| LOC111090004 | 4.997150395 | 6.047635498 | 3.35E-12 | 1.29E-10 |

| | | | | |
|---|---|---|---|---|
| KRT20 | 4.996153063 | 0.787291787 | 0.000777282 | 0.003295389 |
| RASL11B | 4.995168882 | 3.781984752 | 4.64E-08 | 6.86E-07 |
| OPHN1 | 4.994237663 | 4.523048928 | 1.80E-13 | 8.78E-12 |
| GNG4 | 4.99323543 | 2.277912641 | 7.14E-07 | 7.61E-06 |
| LOC119865387 | 4.992873489 | 1.070019303 | 6.80E-07 | 7.30E-06 |
| CMTM5 | 4.988043689 | 1.955107453 | 0.011174174 | 0.031636561 |
| LOC119866608 | 4.985886813 | 2.083526123 | 3.50E-06 | 3.06E-05 |
| MYORG | 4.978408608 | 2.26357256 | 2.29E-08 | 3.62E-07 |
| CD300LG | 4.97838518 | 3.621531222 | 3.95E-05 | 0.000255618 |
| LOC102155782 | 4.977984402 | 1.514531533 | 0.000254633 | 0.001262851 |
| YAP1 | 4.977858187 | 6.383185944 | 1.00E-14 | 6.14E-13 |
| ABCB11 | 4.97278498 | 3.533375659 | 0.005166768 | 0.016461416 |
| NID2 | 4.972572494 | 7.35312326 | 1.66E-09 | 3.44E-08 |
| IQUB | 4.967298248 | 2.301967946 | 2.10E-05 | 0.000146802 |
| SNCG | 4.966714879 | 3.949022857 | 2.39E-10 | 6.01E-09 |
| FAM171A1 | 4.963089994 | 6.006942121 | 1.23E-07 | 1.63E-06 |
| SGIP1 | 4.962283422 | 4.619720798 | 7.75E-05 | 0.000459276 |
| APC2 | 4.962224837 | 6.229159201 | 3.47E-05 | 0.000228098 |
| INHBA | 4.961765622 | 5.847628559 | 1.54E-07 | 1.98E-06 |
| PCLO | 4.961476249 | 3.566578244 | 0.000237909 | 0.001191912 |
| NEO1 | 4.960036363 | 5.99493237 | 4.56E-11 | 1.33E-09 |
| SLC7A14 | 4.959917717 | 0.986191264 | 0.003241995 | 0.011117303 |
| LOC102155458 | 4.959267934 | 1.051073588 | 0.001142411 | 0.00459189 |
| RASA4B | 4.958072873 | 5.435137855 | 3.25E-12 | 1.26E-10 |
| MIR145 | 4.958059243 | -0.125774976 | 0.018987939 | 0.048839971 |
| MEST | 4.957631116 | 4.808222069 | 0.000685747 | 0.002959418 |
| LOC119878218 | 4.957061265 | 7.278972337 | 6.01E-07 | 6.57E-06 |
| PI3 | 4.952886157 | 5.706219089 | 0.002369877 | 0.008493856 |
| MYH15 | 4.952014874 | 0.694587424 | 0.002005492 | 0.007388243 |
| CHPF | 4.951414732 | 6.481158925 | 1.34E-09 | 2.85E-08 |
| SERPINF1 | 4.949564332 | 5.302134969 | 7.92E-05 | 0.000468301 |
| MRC2 | 4.945327386 | 7.451617765 | 8.51E-09 | 1.50E-07 |
| LOC111094286 | 4.945136213 | 3.374713391 | 0.001298756 | 0.005113157 |
| LOC102152633 | 4.944595903 | -0.57186023 | 0.000113731 | 0.000638737 |
| PALD1 | 4.941267922 | 4.805763149 | 6.46E-08 | 9.21E-07 |
| LOC106559255 | 4.939298447 | 0.616737874 | 0.000236357 | 0.001184682 |
| CHST9 | 4.938667577 | 1.46060272 | 8.63E-06 | 6.69E-05 |
| RHPN2 | 4.938051953 | 3.556891689 | 7.03E-05 | 0.000422076 |
| LOC487387 | 4.936636317 | 4.022747039 | 3.10E-05 | 0.000207079 |

| | | | | |
|---|---|---|---|---|
| LOC102155002 | 4.933058386 | 0.988014851 | 3.60E-06 | 3.13E-05 |
| ITGB5 | 4.928508794 | 7.463414786 | 1.90E-10 | 4.90E-09 |
| LOC119873139 | 4.927769922 | 0.56655515 | 5.18E-05 | 0.000323775 |
| STAC2 | 4.923372997 | 4.061695296 | 7.28E-08 | 1.02E-06 |
| TCEAL4 | 4.922459494 | 6.375490063 | 1.55E-06 | 1.51E-05 |
| LOC119864313 | 4.918355149 | 2.804516137 | 0.000335283 | 0.001599866 |
| CPLX1 | 4.915380767 | 4.752437378 | 8.11E-07 | 8.51E-06 |
| LOC119864844 | 4.914923832 | 3.717484572 | 0.000379322 | 0.001780989 |
| LOC111095290 | 4.906175647 | 1.558018334 | 7.13E-08 | 1.00E-06 |
| PLXND1 | 4.905007754 | 8.125518903 | 4.59E-11 | 1.34E-09 |
| NCR3LG1 | 4.900026098 | 2.681213955 | 4.97E-06 | 4.13E-05 |
| LOC119871970 | 4.899780112 | 2.234044138 | 3.81E-06 | 3.29E-05 |
| ASIC4 | 4.899421726 | 0.122434684 | 0.000216303 | 0.001100197 |
| AADAT | 4.898430259 | 2.23670175 | 0.000124184 | 0.000686767 |
| GLTPD2 | 4.894029734 | 1.41905673 | 0.001379656 | 0.005375994 |
| ADGRL2 | 4.892950415 | 5.167988162 | 3.29E-05 | 0.000217939 |
| IGFBP6 | 4.890206663 | 6.305560181 | 4.27E-09 | 8.01E-08 |
| CHRNA5 | 4.889548006 | 0.434715585 | 0.000105451 | 0.000599536 |
| FKBP10 | 4.8887955 | 5.729531921 | 2.02E-07 | 2.52E-06 |
| PRM3 | 4.882697074 | 0.875801716 | 0.00084346 | 0.003537897 |
| ICAM1 | 4.882526647 | 6.031152612 | 2.66E-16 | 2.20E-14 |
| LOC111098097 | 4.879029401 | 0.531286425 | 0.001576124 | 0.006013969 |
| EPS8L2 | 4.878948395 | 3.483494571 | 1.91E-05 | 0.00013513 |
| RNF165 | 4.878042653 | 1.795003543 | 0.008663773 | 0.025557363 |
| LOC111090065 | 4.877514181 | 3.65245645 | 0.000319569 | 0.001537412 |
| HEY2 | 4.876993973 | 1.416904649 | 0.000328627 | 0.001572955 |
| LOC106559882 | 4.875547506 | 0.948858478 | 2.54E-05 | 0.000173857 |
| LOC102153655 | 4.873291197 | 0.816553734 | 0.000452644 | 0.002070176 |
| PDE6H | 4.872867659 | 0.900192716 | 0.000313205 | 0.001512834 |
| LOC119870760 | 4.870005377 | 1.115089755 | 4.68E-05 | 0.000296599 |
| GALNT5 | 4.868520271 | 3.450385094 | 9.59E-06 | 7.34E-05 |
| LOC111091940 | 4.864366033 | -0.553556307 | 0.000489665 | 0.002217871 |
| FADS2 | 4.863994378 | 6.83800366 | 1.74E-17 | 1.74E-15 |
| P3H4 | 4.857497214 | 4.000206266 | 1.76E-05 | 0.000125861 |
| LOC111095569 | 4.852650853 | 3.13209109 | 4.85E-09 | 8.95E-08 |
| STMN3 | 4.847614376 | 5.393116687 | 1.35E-05 | 9.98E-05 |
| ABCG4 | 4.84636311 | 2.81264682 | 4.88E-05 | 0.000307524 |
| HSD3B7 | 4.845954848 | 4.652024262 | 1.14E-15 | 8.43E-14 |
| MYO7A | 4.841244129 | 4.209920646 | 7.11E-06 | 5.64E-05 |

| | | | | |
|---|---|---|---|---|
| LOC119873408 | 4.840044625 | -1.25590046 | 0.013973196 | 0.038211127 |
| KCNIP3 | 4.839137453 | 2.435362998 | 0.000338674 | 0.001613558 |
| LOC102153053 | 4.838353124 | 2.224644732 | 0.000100659 | 0.000575465 |
| LOC102154915 | 4.837069723 | 0.596283854 | 4.12E-06 | 3.52E-05 |
| LOC119873726 | 4.833332739 | 4.139290474 | 8.84E-05 | 0.000513751 |
| ITGA7 | 4.831741952 | 8.110760319 | 1.90E-06 | 1.80E-05 |
| LOC102151579 | 4.83110606 | 2.14362857 | 1.80E-05 | 0.000128425 |
| ZNF214 | 4.828199409 | 1.93179044 | 2.65E-09 | 5.23E-08 |
| ASIC2 | 4.824928965 | 1.483935316 | 6.75E-05 | 0.000406782 |
| LOC119879489 | 4.823108899 | 5.334846386 | 5.63E-06 | 4.61E-05 |
| LOC119864724 | 4.820801354 | 4.64092439 | 8.74E-06 | 6.76E-05 |
| SLC47A1 | 4.820712137 | 0.334258203 | 0.007375652 | 0.022322652 |
| MAMLD1 | 4.818988838 | 5.915777001 | 4.47E-06 | 3.79E-05 |
| PCDH12 | 4.816071225 | 2.545867926 | 4.29E-05 | 0.000274151 |
| CYP26B1 | 4.813888689 | 4.982083926 | 6.60E-06 | 5.29E-05 |
| TGFBR3 | 4.813711107 | 8.479298932 | 4.00E-17 | 3.75E-15 |
| HOXB4 | 4.813485052 | 1.714824662 | 4.94E-06 | 4.11E-05 |
| TRIP10 | 4.813060699 | 6.708502763 | 4.03E-07 | 4.64E-06 |
| LOC119865606 | 4.801433104 | 2.418488538 | 1.34E-06 | 1.33E-05 |
| LOC111090959 | 4.798335059 | 2.40814097 | 1.70E-08 | 2.78E-07 |
| LMO7 | 4.797177604 | 4.804055995 | 1.09E-05 | 8.26E-05 |
| NKAIN3 | 4.793035668 | 1.363117718 | 0.012364361 | 0.034439379 |
| LOC119877329 | 4.7919355 | 5.17566373 | 1.17E-13 | 5.89E-12 |
| LOC111092741 | 4.791674875 | 1.465249383 | 7.33E-05 | 0.000437887 |
| OTOP3 | 4.790784197 | -1.245884401 | 0.011195919 | 0.031675614 |
| SYDE1 | 4.790340357 | 5.264553494 | 1.27E-10 | 3.38E-09 |
| CFI | 4.789639591 | 4.985018972 | 1.00E-07 | 1.36E-06 |
| LGR4 | 4.788947064 | 6.009885845 | 2.90E-09 | 5.67E-08 |
| NCKAP5 | 4.786210296 | 3.964519059 | 2.62E-06 | 2.39E-05 |
| NMT2 | 4.783518354 | 5.297424907 | 1.81E-09 | 3.71E-08 |
| LOC119875958 | 4.779926851 | 1.058313364 | 3.63E-05 | 0.000237648 |
| TUBB4A | 4.77911343 | 5.233055452 | 1.23E-06 | 1.23E-05 |
| EHD2 | 4.777223226 | 8.292361 | 5.51E-11 | 1.59E-09 |
| LOC119878331 | 4.776657805 | 0.819053905 | 0.000105044 | 0.000597853 |
| LOC119869758 | 4.773854889 | 1.391416882 | 6.65E-05 | 0.000401553 |
| LOC100685214 | 4.773437141 | 4.257082734 | 4.02E-05 | 0.000259525 |
| MAST4 | 4.770274025 | 6.767719965 | 2.16E-11 | 6.82E-10 |
| LOC119864725 | 4.76921365 | 4.205155025 | 7.69E-07 | 8.15E-06 |
| LOC119867623 | 4.766642994 | 0.241534556 | 9.10E-05 | 0.000526927 |

| | | | | |
|---|---|---|---|---|
| PALLD | 4.766526592 | 7.3501899 | 3.45E-15 | 2.32E-13 |
| SDC4 | 4.766152333 | 6.292592852 | 1.03E-11 | 3.52E-10 |
| CUEDC1 | 4.764874425 | 5.13365585 | 4.01E-11 | 1.19E-09 |
| LOC119879485 | 4.762624122 | 6.497818618 | 0.000386037 | 0.001807819 |
| WWC2 | 4.760642609 | 5.943598441 | 4.68E-12 | 1.74E-10 |
| ADCY5 | 4.753534348 | 3.12211721 | 2.27E-07 | 2.80E-06 |
| FAM71E2 | 4.753246968 | 0.662927593 | 0.000315501 | 0.001521551 |
| DENND2A | 4.753026516 | 6.698950344 | 9.84E-13 | 4.20E-11 |
| CDO1 | 4.750947655 | 5.878806354 | 3.51E-08 | 5.33E-07 |
| ZDBF2 | 4.748875358 | 2.563380287 | 0.000691806 | 0.002982004 |
| PDGFA | 4.748553389 | 3.515522025 | 0.00067146 | 0.002909928 |
| LOC607920 | 4.748147662 | 2.082650046 | 3.08E-06 | 2.75E-05 |
| SORBS1 | 4.747023901 | 7.83436632 | 2.64E-10 | 6.54E-09 |
| NREP | 4.746496534 | 4.58564821 | 3.92E-05 | 0.000254231 |
| SEMA5A | 4.745886233 | 7.233603952 | 6.47E-06 | 5.19E-05 |
| PTK7 | 4.745311389 | 4.434863073 | 1.40E-05 | 0.000103022 |
| IDO2 | 4.742614205 | 2.329251505 | 0.000210221 | 0.001074678 |
| APOB | 4.74126114 | 4.426743865 | 1.73E-06 | 1.66E-05 |
| AMOTL1 | 4.740408772 | 6.686995987 | 2.13E-14 | 1.24E-12 |
| CD109 | 4.738904219 | 6.638437691 | 3.27E-09 | 6.32E-08 |
| LOC111092800 | 4.737537843 | 2.752590699 | 7.38E-06 | 5.83E-05 |
| MTARC2 | 4.736188881 | 4.427566449 | 2.70E-10 | 6.67E-09 |
| PDK4 | 4.734223448 | 6.27363872 | 3.81E-06 | 3.29E-05 |
| DOC2B | 4.733686444 | 5.32423983 | 4.85E-09 | 8.95E-08 |
| GXYLT2 | 4.726469292 | 3.779884385 | 5.93E-05 | 0.000364325 |
| LOC102152165 | 4.724792791 | 3.726904082 | 5.02E-06 | 4.17E-05 |
| MMP17 | 4.723255764 | 3.731723471 | 3.13E-06 | 2.79E-05 |
| PRR36 | 4.717504487 | 2.832515647 | 5.49E-05 | 0.000340924 |
| MLIP | 4.716956263 | 4.839055283 | 0.001977744 | 0.007299669 |
| LOC119866579 | 4.71626545 | 3.083899607 | 3.03E-05 | 0.000202828 |
| LOC106559499 | 4.712054346 | -0.014754591 | 0.000971326 | 0.003992328 |
| PERP | 4.711173628 | 3.602317918 | 0.0020747 | 0.007599285 |
| CTXND2 | 4.710441876 | -0.245917234 | 8.80E-05 | 0.000512358 |
| LOC102154995 | 4.707872312 | 1.294400916 | 0.005005428 | 0.016009834 |
| UNC79 | 4.705959825 | 1.867492747 | 0.00254372 | 0.009033269 |
| PLIN4 | 4.703297985 | 7.555086668 | 2.42E-05 | 0.000166703 |
| LOC111099011 | 4.695328799 | 2.677166969 | 7.13E-07 | 7.61E-06 |
| LOC119879136 | 4.694247205 | 0.009939266 | 0.000248754 | 0.001237941 |
| LOC119878340 | 4.6937731 | 6.441930698 | 8.86E-06 | 6.85E-05 |

| | | | | |
|---|---|---|---|---|
| CCBE1 | 4.692059568 | 1.70835631 | 0.001325788 | 0.005198806 |
| INHBB | 4.691825003 | 4.269800344 | 2.49E-06 | 2.28E-05 |
| TCF7L1 | 4.691260638 | 4.514577392 | 1.17E-11 | 3.92E-10 |
| EPB41L2 | 4.689904362 | 8.330409887 | 2.77E-09 | 5.44E-08 |
| SLC1A3 | 4.688913123 | 2.877016955 | 9.48E-05 | 0.000546317 |
| MAPK8IP1 | 4.686508968 | 6.334753689 | 1.20E-05 | 8.94E-05 |
| LGI4 | 4.686012267 | 6.522389294 | 1.22E-08 | 2.04E-07 |
| B4GALNT4 | 4.683351868 | 2.881845935 | 0.000147201 | 0.000790065 |
| XKR5 | 4.678952763 | 1.538559769 | 0.001973258 | 0.007286834 |
| ST8SIA1 | 4.67861951 | 4.474747726 | 0.002286572 | 0.008251227 |
| NLGN2 | 4.67808598 | 4.747571938 | 5.56E-12 | 2.05E-10 |
| CFAP44 | 4.669604406 | 2.643060583 | 8.48E-05 | 0.0004965 |
| GAL3ST1 | 4.66958917 | 0.928569747 | 0.018164775 | 0.047132913 |
| ASNS | 4.668374363 | 3.180094398 | 2.37E-06 | 2.19E-05 |
| LOC119878339 | 4.668043609 | 6.410536119 | 8.43E-06 | 6.56E-05 |
| LOC119869766 | 4.658369078 | 3.245696831 | 0.001700696 | 0.006409177 |
| LOC111096727 | 4.657214138 | -0.103337024 | 0.004523733 | 0.014696876 |
| NKAIN1 | 4.654960091 | 0.982147452 | 2.97E-05 | 0.000199582 |
| LOC102152366 | 4.654885129 | -0.776921665 | 0.000385733 | 0.001807086 |
| GLIS2 | 4.653720825 | 6.756222962 | 1.69E-11 | 5.48E-10 |
| DNER | 4.652207861 | -0.840936217 | 0.003161746 | 0.010886696 |
| NRSN2 | 4.647721004 | 2.271942834 | 9.15E-05 | 0.000529511 |
| SP7 | 4.64625043 | 1.584818516 | 0.007832817 | 0.023473541 |
| LOC119876983 | 4.644494966 | 0.61110929 | 1.94E-05 | 0.0001371 |
| CELF3 | 4.64370924 | 2.574803213 | 0.000119685 | 0.000666137 |
| RIMS3 | 4.642413937 | 3.394626434 | 0.000562646 | 0.002497793 |
| SDR42E2 | 4.640686009 | 1.97838857 | 0.018189802 | 0.047178983 |
| AGAP1 | 4.638610657 | 7.487292032 | 9.85E-10 | 2.17E-08 |
| NINJ2 | 4.636223264 | 2.867507118 | 6.67E-06 | 5.34E-05 |
| MINDY4 | 4.633428613 | 3.708122873 | 1.36E-06 | 1.35E-05 |
| SEPTIN3 | 4.633314145 | 3.210917369 | 0.002528713 | 0.008990287 |
| LOC111092083 | 4.632211797 | 4.051397033 | 0.000361668 | 0.001709581 |
| FAM167B | 4.631807551 | 3.261984761 | 0.00023959 | 0.001199777 |
| LOC111093535 | 4.628523133 | 6.162219354 | 0.006279059 | 0.019440863 |
| LOC102154476 | 4.625633257 | 1.036323772 | 5.93E-07 | 6.50E-06 |
| ADRA2B | 4.625236041 | 2.40976744 | 0.002655665 | 0.009383149 |
| IGSF10 | 4.623566005 | 3.640101964 | 1.68E-05 | 0.000120944 |
| HOOK2 | 4.622160764 | 4.655975908 | 7.62E-05 | 0.000453002 |
| RAI2 | 4.621374521 | 3.105934134 | 0.000137892 | 0.000748613 |

| | | | | |
|---|---|---|---|---|
| ACSBG2 | 4.619076137 | -0.235948616 | 1.13E-05 | 8.53E-05 |
| MYH6 | 4.618968755 | 1.245676566 | 0.01590384 | 0.042406979 |
| CREB3L1 | 4.613687525 | 5.419486178 | 0.000184152 | 0.000956414 |
| C9H9orf50 | 4.610216092 | 0.369151989 | 0.000165167 | 0.000872026 |
| PODXL | 4.608536369 | 7.11057739 | 5.83E-11 | 1.67E-09 |
| IL11RA | 4.607485163 | 3.801175016 | 2.35E-11 | 7.35E-10 |
| ODF3B | 4.606553341 | 5.617463348 | 8.49E-07 | 8.84E-06 |
| LOC111090527 | 4.606397195 | 3.771097399 | 0.000132752 | 0.000725269 |
| DYNC2LI1 | 4.605569053 | 2.521950813 | 9.42E-06 | 7.22E-05 |
| HOXA10 | 4.599106556 | 0.927604402 | 0.009405378 | 0.027398219 |
| IGSF8 | 4.592496802 | 5.713811913 | 4.90E-10 | 1.15E-08 |
| LOC119864307 | 4.591166062 | 3.570111322 | 0.000627269 | 0.002743102 |
| FOXD2 | 4.590310297 | 2.452329544 | 3.45E-06 | 3.03E-05 |
| HSD11B1 | 4.588241447 | 2.010588329 | 0.000976702 | 0.004010621 |
| NPAS1 | 4.587254973 | 3.685195867 | 4.03E-07 | 4.64E-06 |
| ADGRA2 | 4.586582815 | 6.947085566 | 6.95E-09 | 1.25E-07 |
| GSN | 4.586310688 | 10.5649266 | 3.09E-07 | 3.68E-06 |
| LOC102153358 | 4.585267256 | -0.681953283 | 0.001081486 | 0.004377914 |
| SCUBE2 | 4.577281309 | 5.123771899 | 1.36E-07 | 1.78E-06 |
| LOC119876680 | 4.57548264 | 8.324233981 | 1.03E-07 | 1.38E-06 |
| LOC119877341 | 4.57100104 | 3.518795981 | 1.35E-05 | 9.96E-05 |
| CA12 | 4.57054462 | 3.999228664 | 0.000106675 | 0.000605705 |
| ANK2 | 4.570103631 | 6.163313523 | 6.06E-11 | 1.72E-09 |
| ARHGAP21 | 4.569710091 | 8.752009841 | 1.79E-06 | 1.72E-05 |
| ERAS | 4.567282568 | -0.962003359 | 0.000137671 | 0.000747805 |
| KHDRBS3 | 4.566445022 | 4.517385187 | 0.000208817 | 0.00106864 |
| HTR3B | 4.565356076 | 1.093200855 | 4.17E-05 | 0.000267142 |
| MAOB | 4.56385146 | 2.658953058 | 0.00648102 | 0.019974851 |
| LOC119871927 | 4.563761371 | 3.525820212 | 4.57E-07 | 5.18E-06 |
| HOXA1 | 4.560917065 | 2.136688973 | 0.000937848 | 0.003879744 |
| LOC111094898 | 4.560080553 | 0.615280891 | 0.000571257 | 0.002529801 |
| LOC111097610 | 4.556643937 | 1.40534278 | 0.000766068 | 0.003252301 |
| CPED1 | 4.554960985 | 5.445554055 | 7.72E-12 | 2.72E-10 |
| LOC111098999 | 4.554159192 | 0.733423069 | 3.11E-05 | 0.000207571 |
| PLEKHA4 | 4.554055658 | 6.824036496 | 2.24E-06 | 2.08E-05 |
| LOC111091014 | 4.552594284 | 0.317277149 | 9.79E-06 | 7.48E-05 |
| FGFR4 | 4.551462154 | 1.798775966 | 0.000160547 | 0.00085095 |
| TMEM17 | 4.550822671 | 1.690449446 | 2.75E-06 | 2.48E-05 |
| TDRD12 | 4.550368789 | 0.659536545 | 6.90E-06 | 5.50E-05 |

| | | | | |
|---|---|---|---|---|
| LOC111092486 | 4.549256311 | 1.106241548 | 0.000380913 | 0.001787686 |
| ADAMTS3 | 4.549111068 | 1.720470731 | 0.001321527 | 0.005185606 |
| LOC119865971 | 4.546245111 | 2.141353129 | 0.000222823 | 0.001126483 |
| TNNI3K | 4.543378984 | 0.92716996 | 9.43E-05 | 0.000543432 |
| LOC119869452 | 4.542887053 | 4.710957616 | 0.000205327 | 0.001052521 |
| LOC111092415 | 4.541969311 | 1.665750569 | 0.016027681 | 0.042668612 |
| CCDC30 | 4.539483701 | 2.337601906 | 0.000171004 | 0.000897814 |
| LOC111095511 | 4.533897289 | 0.263152472 | 6.38E-06 | 5.13E-05 |
| ARC | 4.533202324 | 2.063757585 | 0.001622311 | 0.006159421 |
| FAM78B | 4.532486651 | 5.278193783 | 3.20E-05 | 0.000212796 |
| HOXC4 | 4.528839054 | 5.672452476 | 9.18E-08 | 1.25E-06 |
| ZNF703 | 4.523618538 | 4.932348303 | 1.74E-06 | 1.67E-05 |
| WASF1 | 4.518255291 | 1.150024422 | 6.60E-05 | 0.000399175 |
| LOC102155202 | 4.518177701 | 1.356890602 | 6.94E-07 | 7.44E-06 |
| KLHL34 | 4.510657573 | 2.698811287 | 6.29E-05 | 0.000383176 |
| COL8A2 | 4.509918948 | 4.9606172 | 1.26E-07 | 1.67E-06 |
| PRAP1 | 4.507797689 | 1.800767643 | 1.08E-05 | 8.15E-05 |
| LOC111096057 | 4.507494466 | 0.726219116 | 0.000418139 | 0.001939294 |
| GABRB1 | 4.502198243 | -0.471533373 | 0.001755918 | 0.006581808 |
| LOC119871041 | 4.501254218 | 2.622880337 | 0.01210341 | 0.03380172 |
| TMEM61 | 4.49976624 | 2.495605044 | 0.000695964 | 0.002997542 |
| NPNT | 4.498670025 | 3.211773922 | 0.001899183 | 0.007036067 |
| SYT1 | 4.495298604 | 3.190335388 | 0.004468735 | 0.01454436 |
| PLK2 | 4.493788536 | 6.33079885 | 2.83E-09 | 5.56E-08 |
| TPBGL | 4.493459625 | 0.925244305 | 0.00147918 | 0.005692165 |
| BMP2 | 4.493022013 | 2.710063551 | 0.000132354 | 0.000723641 |
| RHBDF1 | 4.492772238 | 7.441265727 | 1.35E-08 | 2.24E-07 |
| LOC111090067 | 4.492403988 | 1.206523161 | 0.001685144 | 0.006363847 |
| PCSK1 | 4.489327051 | -0.807127247 | 0.000195672 | 0.001007556 |
| PPL | 4.487675154 | 7.718679862 | 0.000185647 | 0.000962794 |
| TCHH | 4.48585422 | 0.813824062 | 0.001380455 | 0.005378143 |
| KCNE4 | 4.484291544 | 2.668134591 | 0.000138394 | 0.000750794 |
| PLEKHG5 | 4.482681242 | 5.747054977 | 7.27E-09 | 1.30E-07 |
| PKD1 | 4.482273672 | 9.046785247 | 4.23E-11 | 1.24E-09 |
| PEG3 | 4.47836457 | 7.145542165 | 6.56E-12 | 2.37E-10 |
| LOC111094997 | 4.47631777 | 2.253880413 | 0.000121061 | 0.000672917 |
| MYO7B | 4.47388257 | 4.840739327 | 1.19E-09 | 2.59E-08 |
| IL6 | 4.468069797 | 2.446785931 | 0.005198747 | 0.016540288 |
| PALM2AKAP2 | 4.464997878 | 7.650517159 | 1.01E-06 | 1.04E-05 |

| | | | | |
|---|---|---|---|---|
| SPP1 | 4.464482461 | 5.64437391 | 4.91E-05 | 0.000309335 |
| LOC119872412 | 4.463091415 | 7.298419117 | 9.39E-06 | 7.21E-05 |
| PHETA1 | 4.462934973 | 2.555805324 | 9.60E-07 | 9.89E-06 |
| CADM4 | 4.462119609 | 5.981216881 | 3.41E-07 | 4.00E-06 |
| RGN | 4.460965058 | 0.115472068 | 0.002627243 | 0.009293329 |
| ZNF507 | 4.459166414 | 8.438126284 | 0.001209218 | 0.004819305 |
| LOC119868054 | 4.459002725 | 4.396260171 | 0.000162104 | 0.000858151 |
| HEBP2 | 4.458750592 | 1.760643084 | 0.000141668 | 0.000765485 |
| LOC119878345 | 4.457900475 | 4.109612004 | 2.10E-05 | 0.000147043 |
| LOC119870551 | 4.455679069 | 6.766730899 | 3.15E-06 | 2.80E-05 |
| LOC119877781 | 4.453397471 | 2.289660282 | 0.000594275 | 0.002616232 |
| IGF1 | 4.451195604 | 3.813387679 | 0.000845308 | 0.003542904 |
| TMEM176A | 4.450836557 | 3.927976212 | 2.46E-10 | 6.16E-09 |
| BAG3 | 4.448809956 | 5.095257073 | 4.30E-07 | 4.90E-06 |
| LOC119871108 | 4.447532334 | -0.93705576 | 0.000942787 | 0.003894228 |
| ALDH1L2 | 4.447522224 | 4.022658977 | 5.06E-06 | 4.20E-05 |
| TMEM35A | 4.446756725 | 0.975443703 | 0.000340373 | 0.001620229 |
| LOC111096325 | 4.442307063 | 0.105134155 | 0.000123518 | 0.000683958 |
| CATSPERZ | 4.441478332 | 0.856100806 | 0.00817597 | 0.024326989 |
| LOC119879137 | 4.441073655 | 0.499364276 | 0.000232447 | 0.0011678 |
| DST | 4.437896499 | 10.18722488 | 1.18E-11 | 3.95E-10 |
| TNR | 4.435540117 | 3.646647754 | 0.002460049 | 0.008776404 |
| LOC111097360 | 4.434375456 | 1.407598625 | 0.001083591 | 0.004383921 |
| LOC479911 | 4.433801628 | 3.251124459 | 3.52E-06 | 3.08E-05 |
| ZNF286A | 4.433395131 | 2.649823729 | 9.90E-06 | 7.55E-05 |
| GFRA4 | 4.430604406 | 1.684345609 | 0.000461603 | 0.002106173 |
| LOC106557926 | 4.429670045 | 4.673093191 | 3.51E-11 | 1.06E-09 |
| HES5 | 4.42856446 | -0.248589151 | 0.001690915 | 0.006378973 |
| ADRA1D | 4.424335205 | 2.529758312 | 1.16E-07 | 1.54E-06 |
| TSKS | 4.4243083 | 1.069075432 | 0.0048272 | 0.015526848 |
| DLL1 | 4.4230526 | 3.363225884 | 5.37E-05 | 0.000334435 |
| YES1 | 4.422872032 | 4.866695895 | 9.96E-05 | 0.000570252 |
| LIPJ | 4.416916848 | 0.847553646 | 3.20E-06 | 2.84E-05 |
| SLC6A8 | 4.415602498 | 5.210091854 | 5.61E-08 | 8.11E-07 |
| AUTS2 | 4.415331358 | 7.190407559 | 3.84E-09 | 7.27E-08 |
| PLCE1 | 4.413508099 | 6.564706036 | 1.51E-09 | 3.17E-08 |
| ENDOU | 4.40379308 | 5.388095383 | 5.92E-06 | 4.81E-05 |
| SGCG | 4.39838531 | 2.244073549 | 2.78E-05 | 0.000188463 |
| GPC4 | 4.397867199 | 4.228806216 | 0.000271001 | 0.001330932 |

| | | | | |
|---|---|---|---|---|
| LOC119864290 | 4.397533833 | 1.580296995 | 0.000257512 | 0.001273923 |
| LOC119876879 | 4.394068897 | 4.801068433 | 6.69E-07 | 7.19E-06 |
| LOC119864602 | 4.393327458 | 3.643193959 | 2.18E-10 | 5.56E-09 |
| FIGN | 4.391515347 | 3.781468437 | 0.000630018 | 0.00275401 |
| CMTM8 | 4.391382366 | 1.725841151 | 4.66E-05 | 0.000295597 |
| PPARA | 4.390760555 | 2.40771642 | 5.87E-06 | 4.77E-05 |
| WNK2 | 4.389393579 | 4.019461863 | 0.000174187 | 0.000912312 |
| NCKAP5L | 4.385157777 | 6.424178563 | 4.83E-08 | 7.12E-07 |
| KIFC3 | 4.383732504 | 4.94083594 | 7.73E-10 | 1.75E-08 |
| LOC119871207 | 4.381892561 | 3.791856111 | 5.79E-07 | 6.37E-06 |
| MARCHF10 | 4.381380082 | 2.974585858 | 0.001153778 | 0.004629854 |
| LOC106559773 | 4.378691141 | 1.491025716 | 0.000498443 | 0.002252923 |
| LOC119866294 | 4.378496148 | 5.125537506 | 5.55E-07 | 6.13E-06 |
| MAP2 | 4.378084278 | 6.314339903 | 1.29E-06 | 1.28E-05 |
| PPP1R9A | 4.37507985 | 5.888657084 | 1.58E-08 | 2.60E-07 |
| LOC102157048 | 4.37494312 | 0.660378394 | 0.002366464 | 0.008484431 |
| TOPAZ1 | 4.374901984 | 1.793367794 | 0.002759183 | 0.009705599 |
| TULP1 | 4.372883973 | 1.322749049 | 9.02E-05 | 0.000523245 |
| FOLH1B | 4.372444604 | 3.670014865 | 0.000614258 | 0.00269381 |
| FAIM2 | 4.372215362 | 5.36819191 | 3.60E-05 | 0.000235903 |
| JAG1 | 4.370153509 | 7.262750269 | 2.45E-12 | 9.70E-11 |
| NR5A2 | 4.36935354 | 3.071160095 | 0.001405364 | 0.005451916 |
| LOC119867818 | 4.367214744 | 2.763557429 | 0.000395025 | 0.00184515 |
| LOC111094445 | 4.367071667 | 3.127079856 | 6.96E-07 | 7.45E-06 |
| USHBP1 | 4.363993158 | 3.493384613 | 2.28E-05 | 0.000158274 |
| NRIP3 | 4.363603369 | 2.456838247 | 0.00095136 | 0.003925153 |
| LOC111091408 | 4.362360848 | 4.923731762 | 3.31E-06 | 2.92E-05 |
| MIR365-2 | 4.361383906 | 0.077874341 | 0.000138779 | 0.000752501 |
| PRND | 4.361222191 | 1.989797571 | 0.002675198 | 0.009439999 |
| RAPGEF3 | 4.358755014 | 5.896930311 | 1.29E-06 | 1.28E-05 |
| LOC119879490 | 4.350830256 | 0.554891536 | 0.001798529 | 0.006728727 |
| LUZP1 | 4.350562836 | 7.377053037 | 1.09E-08 | 1.87E-07 |
| LOC111091483 | 4.349705076 | 3.713784828 | 2.58E-09 | 5.11E-08 |
| LOC111092832 | 4.346355442 | -0.649812318 | 0.000263332 | 0.001298862 |
| CHGA | 4.344604568 | 5.810107359 | 1.75E-06 | 1.68E-05 |
| EGFL7 | 4.340609284 | 6.243376138 | 2.05E-07 | 2.55E-06 |
| CMYA5 | 4.339755705 | 2.432497766 | 5.61E-06 | 4.59E-05 |
| LOC111091528 | 4.335086896 | 0.597655444 | 0.001187273 | 0.004744931 |
| CABP1 | 4.334879714 | 1.789714501 | 3.66E-06 | 3.17E-05 |

| | | | | |
|---|---|---|---|---|
| DACT3 | 4.332521555 | 3.066036995 | 1.96E-05 | 0.000138294 |
| LOC119871887 | 4.332053818 | 0.516571129 | 0.000116866 | 0.000653802 |
| AMPH | 4.331014431 | 3.793150615 | 4.90E-05 | 0.000308935 |
| DUSP15 | 4.326345359 | 2.205011721 | 0.010343851 | 0.029673366 |
| CCDC9B | 4.326341752 | 5.709380513 | 5.97E-12 | 2.18E-10 |
| LOC489080 | 4.3189442 | 4.202702943 | 0.006283164 | 0.019450795 |
| TGFB3 | 4.31769239 | 6.25024291 | 1.21E-07 | 1.60E-06 |
| TMEM249 | 4.316044712 | 1.320554159 | 0.000213084 | 0.001085861 |
| ADAMTSL5 | 4.315187766 | 6.612570735 | 1.04E-05 | 7.87E-05 |
| LOC106559249 | 4.31495316 | -0.822728198 | 0.001575665 | 0.006013969 |
| LOC119867529 | 4.31233771 | 3.635842078 | 7.94E-06 | 6.23E-05 |
| SH3D21 | 4.311135158 | 5.105497379 | 5.99E-07 | 6.55E-06 |
| LOC119873960 | 4.310952917 | 0.482710747 | 5.40E-06 | 4.45E-05 |
| SLC39A14 | 4.308671701 | 5.402423163 | 4.02E-12 | 1.52E-10 |
| CTTNBP2 | 4.306881954 | 5.712043058 | 6.60E-15 | 4.25E-13 |
| PLAC9 | 4.298300365 | 4.399947335 | 3.42E-06 | 3.00E-05 |
| LOC119866411 | 4.297945535 | 1.038467235 | 0.007798245 | 0.023389338 |
| LOC111096245 | 4.295489966 | 0.782532298 | 9.54E-05 | 0.00054862 |
| GPR176 | 4.294316049 | 2.73008076 | 5.53E-05 | 0.000342629 |
| NECTIN3 | 4.291030475 | 3.456723656 | 3.12E-05 | 0.000207882 |
| HIC1 | 4.2907662 | 6.272033519 | 2.33E-08 | 3.68E-07 |
| ACR | 4.289737184 | -0.376791809 | 0.000815804 | 0.003433188 |
| PTGER3 | 4.287327991 | 3.145730992 | 0.001049941 | 0.004266275 |
| DDX25 | 4.28520466 | 1.516717966 | 3.29E-05 | 0.00021801 |
| VASN | 4.283229924 | 6.163355194 | 1.21E-08 | 2.04E-07 |
| PRDM16 | 4.279081297 | 5.083270975 | 0.003192354 | 0.010973494 |
| ITGA9 | 4.274368585 | 5.934084285 | 2.24E-10 | 5.68E-09 |
| LOC119865418 | 4.270529096 | 1.589113079 | 0.000211102 | 0.001078295 |
| NPAS3 | 4.268739807 | 1.773130471 | 0.000743669 | 0.003172742 |
| LOC119866123 | 4.268534841 | 4.785073767 | 1.43E-06 | 1.40E-05 |
| SERPINB6 | 4.268478925 | 4.939872393 | 7.48E-12 | 2.65E-10 |
| CFAP69 | 4.266633748 | -0.721615326 | 0.004797678 | 0.015446683 |
| CCND1 | 4.266496175 | 5.502104608 | 5.80E-08 | 8.35E-07 |
| LOC119869752 | 4.266298061 | 0.89314278 | 0.005207011 | 0.016559288 |
| DPEP1 | 4.265568481 | 4.959448376 | 0.004453495 | 0.014510109 |
| SHISA2 | 4.264419817 | 4.307730543 | 0.000503071 | 0.002270398 |
| SELENBP1 | 4.263640783 | 3.835626555 | 1.12E-08 | 1.91E-07 |
| DIPK1B | 4.262466648 | 2.49614267 | 0.000114163 | 0.000640823 |
| HCN2 | 4.262227804 | 1.774891781 | 0.000173074 | 0.000907143 |

| | | | | |
|---|---|---|---|---|
| ARMH1 | 4.262216876 | 2.53079539 | 7.45E-06 | 5.88E-05 |
| LOC111094308 | 4.261857984 | -0.664504327 | 0.000160619 | 0.000851122 |
| RILPL1 | 4.261634293 | 5.994976246 | 1.84E-13 | 8.94E-12 |
| LOC106557476 | 4.260556227 | 6.325228141 | 4.99E-06 | 4.15E-05 |
| LOC111091526 | 4.257389421 | 1.957334831 | 0.000767654 | 0.003258396 |
| SYP | 4.257107238 | 1.649277727 | 0.003074636 | 0.010632082 |
| FAT4 | 4.256904019 | 7.748804575 | 1.73E-08 | 2.81E-07 |
| RGS12 | 4.256820092 | 5.341444634 | 1.65E-10 | 4.30E-09 |
| LOC111094516 | 4.253787949 | 0.936185318 | 3.78E-05 | 0.000246056 |
| SH3PXD2B | 4.25122537 | 8.182191075 | 9.14E-10 | 2.03E-08 |
| AXIN2 | 4.251213472 | 6.541022115 | 1.25E-08 | 2.09E-07 |
| MELTF | 4.251014555 | 5.032278137 | 0.000688501 | 0.002968939 |
| ARHGDIG | 4.250801654 | 3.11296727 | 0.000128367 | 0.000706834 |
| TCAF1 | 4.245366403 | 4.157679994 | 2.39E-05 | 0.000165292 |
| HSPB6 | 4.244090919 | 8.103615546 | 1.44E-05 | 0.000105332 |
| TNFAIP8L3 | 4.240828868 | 3.354418592 | 0.000216589 | 0.001101132 |
| EPHA6 | 4.236719031 | 2.229603332 | 0.000112984 | 0.000635201 |
| ARMCX5 | 4.235876978 | 1.028830051 | 9.14E-07 | 9.44E-06 |
| PDE7B | 4.235661934 | 4.910835637 | 1.79E-08 | 2.91E-07 |
| LOC102155151 | 4.23254759 | 0.941107134 | 0.005980448 | 0.018662959 |
| LOC111093426 | 4.231918609 | 1.164184144 | 1.52E-05 | 0.000110786 |
| LOC119867042 | 4.22966266 | 1.17318605 | 0.000307071 | 0.001486397 |
| FBLN2 | 4.226647698 | 8.781873704 | 2.63E-07 | 3.18E-06 |
| ZYG11A | 4.224122 | 1.960825746 | 1.04E-06 | 1.06E-05 |
| LOC119869367 | 4.217405015 | 0.33420382 | 7.57E-05 | 0.000451025 |
| MAFF | 4.209174879 | 4.791816019 | 3.82E-05 | 0.00024855 |
| FBXO44 | 4.207446212 | 2.872694239 | 0.000127415 | 0.000702485 |
| SOX6 | 4.204928896 | 4.182301459 | 0.000701044 | 0.003015481 |
| LOC111090634_1 | 4.203380768 | 1.604624769 | 0.001140748 | 0.004586055 |
| EMILIN3 | 4.202453378 | 4.685520104 | 0.000198327 | 0.001019773 |
| CA2 | 4.201472451 | 2.191874721 | 0.014130082 | 0.038540305 |
| LOC119869994 | 4.200440619 | 1.459961958 | 0.000181725 | 0.000944946 |
| RPPH1 | 4.199542432 | 11.80254274 | 7.63E-05 | 0.00045345 |
| PLEKHG1 | 4.199006322 | 6.328914683 | 3.18E-08 | 4.87E-07 |
| SPATA6 | 4.196636277 | 5.170243642 | 6.36E-06 | 5.12E-05 |
| FAM43A | 4.195772621 | 4.687182573 | 1.36E-08 | 2.26E-07 |
| LOC111094470 | 4.194682946 | 4.545690775 | 4.69E-05 | 0.000296771 |
| LOC111097879 | 4.194219009 | 2.401147803 | 0.00012112 | 0.000672917 |
| LOC119877340 | 4.188343877 | 0.834674733 | 0.001271268 | 0.005024104 |

| | | | | |
|---|---|---|---|---|
| LOC119878217 | 4.185038609 | 1.612576063 | 0.000529617 | 0.002372549 |
| SPRY2 | 4.18409661 | 3.069373966 | 7.30E-05 | 0.000436421 |
| FBLN5 | 4.180373628 | 5.508930551 | 2.83E-06 | 2.55E-05 |
| LOC119876719 | 4.174774094 | 6.104223224 | 2.82E-11 | 8.60E-10 |
| LOC119876692 | 4.17469544 | 4.28966921 | 0.003312811 | 0.01131577 |
| LOC111091857 | 4.17438268 | 0.22519278 | 0.000261245 | 0.001289154 |
| FRMD6 | 4.174194804 | 6.434170488 | 1.05E-11 | 3.56E-10 |
| LOC119865099 | 4.173316729 | 6.192424505 | 3.72E-07 | 4.32E-06 |
| LTBP3 | 4.169356585 | 7.614945239 | 4.87E-14 | 2.67E-12 |
| EVA1B | 4.16799698 | 5.310955294 | 5.92E-06 | 4.81E-05 |
| ENTPD3 | 4.165582388 | 3.864452164 | 0.000151824 | 0.000809865 |
| LOC119864356 | 4.165128552 | 6.300379144 | 7.75E-05 | 0.000459276 |
| AGRN | 4.161933742 | 6.464898876 | 6.95E-08 | 9.83E-07 |
| LOC106558052 | 4.161795216 | 2.315510232 | 2.66E-06 | 2.42E-05 |
| ZNF771 | 4.154991354 | 2.459479248 | 5.47E-06 | 4.49E-05 |
| FAM186B | 4.154438698 | 1.111371594 | 5.24E-05 | 0.000326894 |
| PTPRN | 4.154402079 | 4.845295582 | 0.002642526 | 0.009342815 |
| NMU | 4.152810192 | 0.547822003 | 4.96E-05 | 0.000312318 |
| AOC3_1 | 4.149852547 | 0.752998974 | 0.000928875 | 0.003851448 |
| ADAMTS20 | 4.149131137 | 5.405972606 | 0.003338794 | 0.011393747 |
| ZNF510 | 4.147458665 | 4.457248959 | 9.98E-09 | 1.72E-07 |
| LOC111089962 | 4.145759858 | 2.38111342 | 0.010683885 | 0.030483392 |
| LOC119873678 | 4.145491863 | 2.890120729 | 2.04E-06 | 1.92E-05 |
| LOC611067 | 4.142842351 | 1.936143246 | 1.06E-05 | 8.03E-05 |
| LOC119873948 | 4.141836216 | 1.811279866 | 3.18E-05 | 0.000211662 |
| NRXN3 | 4.141015737 | 2.629648026 | 0.005797257 | 0.018182915 |
| SLC39A4 | 4.137787184 | 1.413020756 | 0.000255551 | 0.001266248 |
| CREG2 | 4.136362047 | 1.993074428 | 0.009666691 | 0.028016517 |
| LOC119871368 | 4.135904505 | 3.507645645 | 0.00294849 | 0.010267132 |
| GADD45G | 4.135462172 | 4.163784245 | 0.003191335 | 0.010973494 |
| SMOC1 | 4.134545388 | 3.83162576 | 0.004409319 | 0.014398549 |
| NMNAT2 | 4.132272681 | 3.141753999 | 1.24E-05 | 9.22E-05 |
| LOC111097618 | 4.129254854 | 1.98444651 | 0.000869007 | 0.003630299 |
| CLSTN2 | 4.127356886 | 3.756015737 | 2.77E-06 | 2.50E-05 |
| LOC119872621 | 4.125249608 | 0.272485774 | 0.000128506 | 0.000707238 |
| LOC119878710 | 4.125062374 | 6.826663278 | 0.000147623 | 0.000791938 |
| CHRNA3 | 4.124637225 | 2.406107877 | 0.002796406 | 0.009819172 |
| SMAD9 | 4.122986528 | 4.209032739 | 0.002809465 | 0.009853836 |
| CACNA1H | 4.120552402 | 5.78830663 | 4.23E-05 | 0.000270685 |

| | | | | |
|---|---|---|---|---|
| ARL9 | 4.117157206 | 0.139972573 | 0.001575915 | 0.006013969 |
| LOC111090252 | 4.116965872 | 0.321769109 | 0.000374755 | 0.001762983 |
| COL7A1 | 4.114506374 | 6.547265833 | 7.91E-06 | 6.21E-05 |
| LOC606974 | 4.114325321 | 1.0468344 | 7.34E-05 | 0.000438415 |
| SLC26A1 | 4.109915642 | 4.479703432 | 8.45E-05 | 0.000495003 |
| GUCY1B1 | 4.104978017 | 4.150703826 | 2.41E-05 | 0.00016629 |
| PYGO1 | 4.104339813 | 5.01440553 | 0.000113215 | 0.000636334 |
| LOC119866288 | 4.10219737 | 3.895352303 | 5.63E-05 | 0.000347946 |
| CEND1 | 4.100136635 | 3.239539949 | 2.83E-05 | 0.000191291 |
| LOC111092742 | 4.099571632 | 3.766564318 | 6.57E-05 | 0.000397917 |
| GPR156 | 4.094385856 | 1.58972392 | 0.000139735 | 0.000757308 |
| LOC119877572 | 4.092651726 | -0.056160104 | 0.001019359 | 0.004161342 |
| NOS3 | 4.091501283 | 5.774738599 | 9.61E-07 | 9.89E-06 |
| ADGRG2 | 4.089191509 | 2.225529089 | 0.003552719 | 0.012004263 |
| LOC611347 | 4.088281334 | 1.138276609 | 5.91E-05 | 0.000363144 |
| CCDC113 | 4.085154364 | 1.891306227 | 0.00033127 | 0.001583154 |
| KIF17 | 4.08283977 | 1.853663465 | 0.004019479 | 0.013342651 |
| TRAM1L1 | 4.082356933 | 2.606068603 | 5.08E-05 | 0.000318486 |
| LOXL2 | 4.080261619 | 5.964465048 | 4.80E-06 | 4.01E-05 |
| CCN6 | 4.077843586 | 2.595151249 | 6.20E-06 | 5.00E-05 |
| ABCC5 | 4.077640973 | 4.968772135 | 2.61E-09 | 5.16E-08 |
| KIF7 | 4.072811418 | 5.50559013 | 2.27E-07 | 2.80E-06 |
| ABCB9 | 4.070796069 | 3.000307661 | 0.00030075 | 0.001460158 |
| RYR3 | 4.069422161 | 4.99287612 | 2.39E-07 | 2.92E-06 |
| CCM2L | 4.067483699 | 3.132985654 | 0.001712941 | 0.00644524 |
| LOC111098576 | 4.066632317 | 3.741112547 | 2.50E-08 | 3.92E-07 |
| CCDC148 | 4.065966149 | 1.035261832 | 1.12E-05 | 8.43E-05 |
| TIMP3 | 4.065869357 | 7.18413005 | 1.69E-05 | 0.00012132 |
| P3H3 | 4.064310994 | 4.706192505 | 4.24E-06 | 3.61E-05 |
| DUOXA1 | 4.06323559 | 0.18802769 | 0.003788775 | 0.012695441 |
| FAAP20 | 4.062074311 | 3.200516839 | 2.08E-06 | 1.95E-05 |
| CEP89 | 4.059940789 | 3.981207766 | 1.74E-05 | 0.000124849 |
| MARVELD1 | 4.057989174 | 1.684056673 | 1.02E-05 | 7.75E-05 |
| ANKRD24 | 4.056072605 | 3.767329336 | 0.000182461 | 0.000948546 |
| EFNA3 | 4.054636457 | 3.028421288 | 8.78E-07 | 9.11E-06 |
| S100A1 | 4.054166148 | 2.912770101 | 0.000945794 | 0.003905162 |
| ANO2 | 4.052307564 | 1.680292966 | 0.000264901 | 0.001305307 |
| LPL | 4.05047705 | 7.108879414 | 2.03E-05 | 0.000142639 |
| LOC119870788 | 4.044577062 | 2.137643937 | 0.00382347 | 0.012787973 |

| | | | | |
|---|---|---|---|---|
| ARHGEF28 | 4.043287527 | 5.058310778 | 2.02E-07 | 2.52E-06 |
| KLRG2 | 4.043038883 | 1.563013116 | 0.006435099 | 0.019866747 |
| DNM1 | 4.038686649 | 6.310647478 | 2.56E-05 | 0.000175261 |
| FGF11 | 4.03710431 | 3.364974478 | 2.31E-07 | 2.83E-06 |
| MIR8897 | 4.036956384 | -0.894705998 | 0.00011578 | 0.000648565 |
| PBX4 | 4.036538813 | 0.839882804 | 2.63E-05 | 0.000179272 |
| VEGFB | 4.033866376 | 6.153203006 | 1.52E-07 | 1.96E-06 |
| HJV | 4.032726409 | 0.188720595 | 0.004217877 | 0.013859013 |
| SUGCT | 4.031229964 | 4.151245149 | 3.26E-05 | 0.000216439 |
| NRXN1 | 4.030957324 | 4.031905464 | 0.00034654 | 0.001647052 |
| LOC102156453 | 4.027104263 | -0.233750236 | 0.000547009 | 0.002436862 |
| LOC111090874 | 4.02538393 | 1.438683886 | 0.001863222 | 0.006931276 |
| LOC119873943 | 4.024855003 | 0.878891924 | 5.70E-05 | 0.000351991 |
| SMAD1 | 4.024185514 | 5.168869605 | 5.28E-12 | 1.96E-10 |
| FGF13 | 4.022933346 | 3.040969079 | 2.68E-05 | 0.000181834 |
| LOC119872739 | 4.021765464 | 5.468698426 | 0.002039551 | 0.007494877 |
| LOC102152723 | 4.021233678 | 2.379338092 | 0.000125415 | 0.000692692 |
| LOC119870883 | 4.020432136 | -0.739750143 | 0.000100111 | 0.000572784 |
| RFTN2 | 4.01978294 | 3.954828708 | 0.000476189 | 0.002161801 |
| LOC100687983 | 4.019692282 | 0.02688283 | 0.000327832 | 0.00156984 |
| COLGALT2 | 4.019289518 | 2.038680484 | 0.000664716 | 0.00288416 |
| SOX4 | 4.016495871 | 4.817064749 | 2.17E-08 | 3.45E-07 |
| C18H11orf80 | 4.014225427 | 0.977626474 | 0.001876927 | 0.006965521 |
| KLHL38 | 4.014159708 | 0.733213263 | 0.001643406 | 0.006226842 |
| LIPC | 4.011753895 | 1.25788499 | 0.004577934 | 0.014855154 |
| LOC119867331 | 4.011061582 | 2.476887597 | 0.004341388 | 0.014195946 |
| ITGA10 | 4.009939535 | 5.645015466 | 6.06E-07 | 6.61E-06 |
| VWDE | 4.008398337 | 1.578915042 | 0.008460826 | 0.025037068 |
| TJP1 | 4.006638397 | 7.524249667 | 2.17E-07 | 2.68E-06 |
| LOC119873474 | 4.006432202 | 6.071177263 | 2.73E-06 | 2.47E-05 |
| LOC102152105 | 3.999082376 | 1.553914693 | 0.002972688 | 0.010331144 |
| ID3 | 3.998104524 | 5.46979188 | 2.41E-11 | 7.50E-10 |
| RIN1 | 3.998023844 | 4.612842039 | 7.70E-08 | 1.07E-06 |
| VKORC1 | 3.997153249 | 3.830523445 | 6.23E-08 | 8.94E-07 |
| LOC119876586 | 3.996234115 | 0.978610766 | 0.001935472 | 0.00716066 |
| LOC111092215 | 3.996059883 | 3.370560456 | 0.000267826 | 0.001317702 |
| DNASE1L3 | 3.994945266 | 2.025659634 | 0.007736058 | 0.023222104 |
| LOC102155774 | 3.99376422 | 2.60027823 | 2.41E-07 | 2.95E-06 |
| PPP4R4 | 3.993666037 | 2.298657423 | 0.000392375 | 0.001834724 |

| | | | | |
|---|---|---|---|---|
| EFR3B | 3.992784406 | 3.850496542 | 1.48E-06 | 1.45E-05 |
| MGAT3 | 3.991821508 | 5.031797985 | 1.87E-05 | 0.000132618 |
| LOC106559704 | 3.991671399 | 0.588634718 | 8.84E-05 | 0.000513698 |
| LOC119873082 | 3.986706537 | 4.380966409 | 0.012786066 | 0.035436368 |
| BCL6B | 3.980422658 | 5.030870099 | 3.18E-09 | 6.16E-08 |
| LOC119867351 | 3.980411703 | 2.480733291 | 0.000346037 | 0.001645021 |
| LOC119866440 | 3.978976577 | 2.903789031 | 1.28E-05 | 9.45E-05 |
| LOC119873316 | 3.97861761 | 2.070522984 | 0.000353855 | 0.001678505 |
| LOC102155777 | 3.976534289 | 2.177837099 | 0.002122582 | 0.007751215 |
| LOC111092057 | 3.975879579 | 1.515234272 | 0.000607693 | 0.002668801 |
| KDELR3 | 3.97570959 | 5.108885614 | 1.99E-07 | 2.48E-06 |
| LOC119878219 | 3.974103457 | 10.69290135 | 4.32E-05 | 0.000276243 |
| LOC119869702 | 3.973784705 | 3.10920522 | 2.40E-05 | 0.00016578 |
| NPHP1 | 3.970077764 | 2.106218668 | 6.22E-05 | 0.000379714 |
| TAGLN | 3.969903178 | 8.270134657 | 4.53E-06 | 3.83E-05 |
| LOC119873452 | 3.969825122 | 9.005969777 | 4.84E-07 | 5.44E-06 |
| CDH13 | 3.96823538 | 5.990469411 | 1.20E-05 | 8.96E-05 |
| SHC2 | 3.967490177 | 3.303059657 | 1.65E-07 | 2.09E-06 |
| CTSV | 3.966713671 | 3.839144146 | 1.40E-05 | 0.000102676 |
| CBS | 3.966296986 | 4.322149841 | 9.34E-05 | 0.000539202 |
| PRL | 3.965789377 | -0.677212339 | 0.004480163 | 0.014574989 |
| LOC119875986 | 3.964845056 | -0.214878058 | 4.09E-06 | 3.50E-05 |
| SUSD2 | 3.96196912 | 4.289367556 | 5.14E-06 | 4.25E-05 |
| NSMF | 3.959685211 | 5.261773268 | 1.39E-09 | 2.95E-08 |
| CACNA2D1 | 3.956447422 | 5.627129404 | 0.002970906 | 0.010327196 |
| UCHL1 | 3.954568945 | 3.399128927 | 0.000230961 | 0.001161587 |
| ANKRD35 | 3.950967483 | 2.513674314 | 5.05E-05 | 0.000316861 |
| ACVR2B | 3.949993383 | 3.677046511 | 2.73E-06 | 2.47E-05 |
| LOC111092897 | 3.948771928 | 0.31616119 | 9.97E-05 | 0.000570928 |
| LOC111092898 | 3.947199047 | 2.519391441 | 0.000118483 | 0.000661483 |
| ATF5 | 3.946450731 | 5.820427388 | 1.38E-05 | 0.000101918 |
| LOC484933 | 3.94631399 | 2.922412926 | 1.33E-06 | 1.32E-05 |
| LOC119866812 | 3.946281518 | 2.124988444 | 0.001362696 | 0.005323309 |
| PTPRD | 3.94182104 | 3.758585803 | 0.00328279 | 0.011234471 |
| LOC119870818 | 3.938405717 | 0.933729472 | 0.001585999 | 0.00604526 |
| KCNN1 | 3.938369278 | 2.524408442 | 5.99E-05 | 0.000367274 |
| RBFOX2 | 3.937877014 | 7.150753529 | 9.51E-10 | 2.10E-08 |
| LRFN4 | 3.934314426 | 3.696998652 | 6.36E-06 | 5.12E-05 |
| INSC | 3.933282461 | 2.538907952 | 0.011264914 | 0.031831094 |

| | | | | |
|---|---|---|---|---|
| NOTCH4 | 3.933094149 | 5.821305001 | 1.54E-09 | 3.22E-08 |
| LOC119877096 | 3.930706624 | 2.677949073 | 0.000234766 | 0.001178068 |
| MAST1 | 3.929832357 | 5.424572668 | 0.000365984 | 0.001726971 |
| FADS3 | 3.924583823 | 7.323354497 | 3.41E-07 | 4.00E-06 |
| LOC479761 | 3.924279067 | 4.262053156 | 7.70E-06 | 6.06E-05 |
| VAV2 | 3.923236058 | 6.658942309 | 1.72E-12 | 6.95E-11 |
| DHRS9 | 3.92074422 | 7.649056518 | 0.003802097 | 0.012730239 |
| UGGT2 | 3.918179071 | 4.434222408 | 3.47E-08 | 5.29E-07 |
| LOC111096233 | 3.917929406 | 2.639277973 | 0.002414169 | 0.008633542 |
| CIART | 3.917609405 | 2.850230078 | 2.08E-05 | 0.000145574 |
| MUC2 | 3.914704738 | 1.434096835 | 0.003250182 | 0.011142231 |
| GGT7 | 3.913544772 | 3.707332275 | 2.06E-07 | 2.56E-06 |
| PLEKHF1 | 3.909082255 | 5.517557495 | 3.21E-06 | 2.84E-05 |
| TMEM59L | 3.907592657 | 3.696271151 | 1.21E-05 | 9.01E-05 |
| C1QL1 | 3.906190076 | 2.208582465 | 6.81E-05 | 0.000409598 |
| OSCP1 | 3.905608684 | 1.910175768 | 2.23E-06 | 2.07E-05 |
| LOC479758 | 3.905364459 | 3.746842254 | 1.31E-06 | 1.30E-05 |
| AOC3_2 | 3.905247836 | 5.630559598 | 0.000684743 | 0.002956849 |
| P2RX3 | 3.904283167 | 0.841773502 | 0.000451378 | 0.002066041 |
| LOC119872925 | 3.902460253 | 0.21516647 | 6.33E-05 | 0.000384957 |
| SERP2 | 3.901466557 | 2.702278347 | 8.68E-07 | 9.02E-06 |
| LOC119869671 | 3.901425162 | 3.66691209 | 0.001079147 | 0.004371655 |
| BEGAIN | 3.899965422 | 4.171485385 | 4.16E-05 | 0.000266742 |
| FAM222A | 3.895928086 | 2.844885582 | 9.89E-05 | 0.000566751 |
| LOC119867079 | 3.89555966 | 1.707446301 | 0.001119818 | 0.004512796 |
| LOC119878308 | 3.894797926 | 3.249299156 | 0.000381686 | 0.001790535 |
| LOC106557987 | 3.893914758 | 5.556553899 | 3.75E-12 | 1.43E-10 |
| LIMS2 | 3.893611778 | 6.458429829 | 5.88E-11 | 1.68E-09 |
| PKIA | 3.892878735 | 4.260482708 | 0.001750753 | 0.00656472 |
| LOC102155115 | 3.892514724 | 3.999167239 | 2.88E-07 | 3.45E-06 |
| PDLIM4 | 3.892071025 | 4.727363697 | 1.26E-09 | 2.70E-08 |
| MAP1LC3A | 3.890658502 | 5.191334088 | 3.19E-08 | 4.90E-07 |
| LOC111098114 | 3.887026208 | -0.361017658 | 0.00248298 | 0.008850925 |
| LOC119878144 | 3.884140984 | 1.211595856 | 0.001841624 | 0.00686389 |
| LOC111098294 | 3.883338146 | 5.976649567 | 3.31E-09 | 6.37E-08 |
| PDE1B | 3.882845619 | 4.937833633 | 0.000936628 | 0.003876177 |
| HFM1 | 3.881645173 | 2.824999285 | 2.28E-05 | 0.000158325 |
| LOC106557839 | 3.88130788 | 1.824694362 | 9.37E-06 | 7.20E-05 |
| LOC106559248 | 3.880531769 | 1.649892628 | 0.003834264 | 0.012820117 |

| | | | | |
|---|---|---|---|---|
| CYBRD1 | 3.878820167 | 7.40889254 | 4.62E-07 | 5.24E-06 |
| LURAP1 | 3.878652068 | 4.730307322 | 5.14E-08 | 7.53E-07 |
| LOC111090703 | 3.874670746 | 0.437303341 | 0.000146533 | 0.00078687 |
| ACTA2 | 3.873285948 | 7.71844041 | 0.007546505 | 0.022760285 |
| LOC119870891 | 3.872898661 | -0.762213914 | 0.002791176 | 0.00980599 |
| BCAS1 | 3.870933561 | 2.329947979 | 0.017884413 | 0.046559626 |
| LOC119871629 | 3.866989746 | 2.3110399 | 6.57E-05 | 0.000397917 |
| SLC6A9 | 3.866413139 | 3.236207956 | 8.14E-06 | 6.37E-05 |
| TCEA2 | 3.86600049 | 5.588060324 | 2.67E-05 | 0.000181313 |
| FAT1 | 3.86588647 | 8.646321677 | 1.65E-09 | 3.42E-08 |
| GRID2IP | 3.865846654 | 0.790350891 | 0.005054872 | 0.016158443 |
| AMN | 3.865085081 | 1.170037173 | 0.001761001 | 0.006598577 |
| LOC119871511 | 3.864416805 | -0.278213745 | 0.000458241 | 0.002092155 |
| METTL7B | 3.864355877 | 2.162670809 | 0.003160216 | 0.010885654 |
| LOC119870882 | 3.863583189 | -0.253796129 | 2.60E-05 | 0.000177537 |
| FAM83H | 3.861858919 | 5.152130547 | 9.67E-05 | 0.000555182 |
| KASH5 | 3.861007024 | 0.909603088 | 0.001879256 | 0.006971775 |
| EFNB2 | 3.860521974 | 3.600416977 | 1.88E-06 | 1.79E-05 |
| CPNE4 | 3.85941528 | 1.155532172 | 0.018775747 | 0.048403325 |
| COL2A1 | 3.856966478 | 2.720371562 | 0.004059565 | 0.013434612 |
| GATA6 | 3.851827762 | 3.086524674 | 0.000312174 | 0.001508863 |
| NTRK2 | 3.850500781 | 7.014950951 | 0.00303524 | 0.010515145 |
| CLIP3 | 3.846962061 | 6.179727232 | 1.73E-07 | 2.18E-06 |
| LOC102152093 | 3.845339982 | 0.938050346 | 0.000225833 | 0.001140067 |
| MYOM1 | 3.84381666 | 5.001856417 | 1.17E-07 | 1.56E-06 |
| LOC102153902 | 3.843277137 | 0.938483171 | 0.000453617 | 0.002074101 |
| SLC16A12 | 3.841116848 | 2.110005865 | 7.86E-05 | 0.000465149 |
| GPR17 | 3.839313916 | 5.23369552 | 2.76E-07 | 3.32E-06 |
| SYNPO2L | 3.837879157 | 1.837522755 | 4.41E-05 | 0.000281396 |
| LRP5 | 3.833585966 | 5.867217341 | 9.97E-09 | 1.72E-07 |
| LOC111097625 | 3.829841184 | -0.349712387 | 0.000537426 | 0.002401079 |
| EPHA4 | 3.82975715 | 3.9525034 | 0.000165939 | 0.000875178 |
| FOXS1 | 3.829695364 | 4.37601086 | 0.001886704 | 0.006994617 |
| SLC12A5 | 3.829141777 | 2.415245091 | 0.000572518 | 0.002534349 |
| COL13A1 | 3.82893345 | 1.679081004 | 0.002896274 | 0.010109671 |
| LHFPL4 | 3.821234825 | 1.901415075 | 0.002191887 | 0.00796926 |
| LOC119873719 | 3.818586791 | 5.510487339 | 9.09E-08 | 1.24E-06 |
| ODF2L | 3.816905099 | 5.228148982 | 8.69E-05 | 0.000506969 |
| FAM227A | 3.813878556 | 0.696931452 | 1.43E-05 | 0.000104781 |

| | | | | |
|---|---|---|---|---|
| C29H8orf37 | 3.811788385 | 3.870046323 | 4.83E-07 | 5.43E-06 |
| LOC111095961 | 3.80927899 | 5.565597981 | 4.88E-07 | 5.47E-06 |
| LOC111098779 | 3.808000976 | 1.033395121 | 0.005529428 | 0.017446395 |
| MIR6516 | 3.806810141 | 5.158511697 | 6.91E-08 | 9.77E-07 |
| MSRB3 | 3.805860901 | 6.400832451 | 1.07E-07 | 1.43E-06 |
| LOC111097177 | 3.805203902 | 3.557274573 | 0.00026331 | 0.001298862 |
| LOC111095920 | 3.804206401 | 0.883656977 | 0.016255618 | 0.043148409 |
| LOC119871703 | 3.801673272 | 0.415830309 | 0.003143714 | 0.010837237 |
| LOC111090634_2 | 3.79987377 | 1.258754424 | 0.002902321 | 0.010119371 |
| KIAA1614 | 3.799069681 | 3.933189947 | 1.56E-07 | 2.00E-06 |
| LOC102154211 | 3.798788527 | 1.201912847 | 0.00134264 | 0.005255302 |
| COL23A1 | 3.797107645 | 1.87667163 | 0.003805268 | 0.012738891 |
| LDLRAD3 | 3.796911297 | 4.795305208 | 5.13E-10 | 1.20E-08 |
| KIFC2 | 3.794610742 | 6.188862196 | 8.88E-06 | 6.87E-05 |
| THNSL2 | 3.789961373 | 3.560737761 | 6.77E-07 | 7.27E-06 |
| LOC106557917 | 3.787171827 | 2.968408139 | 7.06E-05 | 0.000423404 |
| RGL1 | 3.786436882 | 7.229761478 | 6.98E-08 | 9.85E-07 |
| SPC24 | 3.786387939 | 1.176150667 | 0.00076512 | 0.003248912 |
| EGR3 | 3.78554038 | 5.716971275 | 0.000613825 | 0.002692457 |
| SPINK13 | 3.78422279 | 0.790316956 | 0.000956099 | 0.00394164 |
| CDH22 | 3.782468994 | 1.444157734 | 0.001081459 | 0.004377914 |
| RAB40B | 3.776810014 | 1.689622812 | 0.000277227 | 0.001357205 |
| CYP7A1 | 3.7767224 | -0.429223547 | 0.009483898 | 0.027578763 |
| TTBK1 | 3.772902664 | 0.574050491 | 0.00161813 | 0.006146114 |
| THRB | 3.76979595 | 4.337354546 | 8.70E-09 | 1.52E-07 |
| TMEM63C | 3.769781099 | 4.939096472 | 0.000210285 | 0.001074678 |
| CNTN2 | 3.769547968 | 5.706459536 | 0.000207045 | 0.001060573 |
| FAM83G | 3.766722697 | 3.872117386 | 2.63E-05 | 0.000179162 |
| FAM20C | 3.76578192 | 6.517752216 | 1.44E-07 | 1.86E-06 |
| HSPA12B | 3.765517639 | 5.508686591 | 1.65E-08 | 2.70E-07 |
| CASKIN2 | 3.764683975 | 6.377406813 | 1.39E-09 | 2.94E-08 |
| TERT | 3.763238366 | 1.809046617 | 0.003325141 | 0.011352517 |
| RIMS1 | 3.761683332 | 3.109534263 | 0.00016653 | 0.000877937 |
| LOC119872326 | 3.761196282 | 0.962937427 | 0.001468416 | 0.005657774 |
| LOC119865807 | 3.75866026 | 8.276031131 | 3.37E-06 | 2.97E-05 |
| SHC4 | 3.75613446 | 4.48532768 | 3.44E-05 | 0.000226699 |
| TEX49 | 3.753449905 | 1.137811405 | 0.000145772 | 0.000784145 |
| RASSF8 | 3.748396732 | 5.754149169 | 4.36E-07 | 4.97E-06 |
| LMNA | 3.746822562 | 8.779635733 | 2.19E-09 | 4.46E-08 |

| | | | | |
|---|---|---|---|---|
| LOC119864946 | 3.744470889 | 1.103325621 | 6.60E-05 | 0.000399175 |
| VASH2 | 3.743814707 | 4.731125577 | 9.20E-06 | 7.08E-05 |
| PDE1C | 3.741736208 | 3.96452125 | 0.000358935 | 0.001697774 |
| LOC119869707 | 3.740359249 | 2.774134229 | 0.000137005 | 0.000744747 |
| KIAA0895L | 3.738282416 | 3.555060693 | 1.39E-07 | 1.82E-06 |
| BTG3 | 3.738076605 | 3.089272991 | 0.000986633 | 0.004047566 |
| LOC102153410 | 3.733523723 | 2.174199231 | 0.000112304 | 0.000632196 |
| LOC119876681 | 3.73351907 | 9.510027543 | 1.33E-05 | 9.84E-05 |
| LOC119873274 | 3.730524831 | 0.622130686 | 0.000962685 | 0.003963578 |
| LOC102153499 | 3.725417579 | 1.48971892 | 0.003216126 | 0.011036527 |
| LOC119877612 | 3.725101709 | -0.003375036 | 0.00024014 | 0.001201059 |
| MYEF2 | 3.723574211 | 5.54501269 | 5.40E-10 | 1.26E-08 |
| PDIA2 | 3.723142151 | 3.644359024 | 0.000228885 | 0.001151758 |
| BHLHE41 | 3.719343933 | 4.885935284 | 1.55E-07 | 1.98E-06 |
| ARVCF | 3.716511505 | 5.15451661 | 1.95E-08 | 3.14E-07 |
| ANO9 | 3.712040571 | 1.65560451 | 0.002254299 | 0.008153775 |
| CNRIP1 | 3.711656297 | 4.238538694 | 4.02E-06 | 3.44E-05 |
| TRAF3IP1 | 3.711238002 | 4.990827795 | 3.88E-06 | 3.34E-05 |
| VTN | 3.710656565 | 4.301662806 | 0.0003786 | 0.001778371 |
| LOC119877700 | 3.707888702 | 3.75755591 | 0.000112283 | 0.000632196 |
| PBX3 | 3.707803066 | 5.479406424 | 5.80E-06 | 4.73E-05 |
| AQP1 | 3.706870248 | 6.712393812 | 0.000274813 | 0.001346909 |
| LOC102157313 | 3.703872063 | -0.365322152 | 5.02E-05 | 0.000315537 |
| LOC119872246 | 3.703473487 | 1.799019802 | 0.000119503 | 0.000665976 |
| CHI3L1 | 3.702370971 | 3.041363645 | 0.001495189 | 0.005744591 |
| HEPACAM2 | 3.701803795 | -0.786352352 | 0.003328132 | 0.01136094 |
| LOC102154534 | 3.701369784 | 1.341240486 | 0.008061977 | 0.024053857 |
| PAWR | 3.698936111 | 5.728975962 | 1.17E-06 | 1.17E-05 |
| LOC119865793 | 3.69876914 | 3.556605516 | 0.001843016 | 0.006866716 |
| TSPAN15 | 3.697818364 | 3.373542048 | 0.000105734 | 0.000600831 |
| SCRN1 | 3.695142718 | 5.033978459 | 1.53E-06 | 1.49E-05 |
| LOC119871509 | 3.694774915 | 4.489334354 | 1.43E-08 | 2.37E-07 |
| KALRN | 3.693130716 | 5.2461071 | 1.05E-06 | 1.07E-05 |
| LOC111098532 | 3.689443523 | -0.877847516 | 0.013237588 | 0.036477845 |
| PDE9A | 3.686604113 | 3.665873598 | 4.38E-05 | 0.000279218 |
| LOC119868038 | 3.685858818 | 2.534296598 | 0.000162623 | 0.000860269 |
| TUB | 3.680090923 | 4.796849959 | 1.27E-05 | 9.38E-05 |
| SGCD | 3.679105788 | 5.753272444 | 1.48E-06 | 1.45E-05 |
| ADARB2 | 3.67863201 | 0.768421966 | 0.001313624 | 0.005162316 |

| | | | | |
|---|---|---|---|---|
| LOC102154314 | 3.676253996 | 2.556496506 | 0.000249772 | 0.001241871 |
| MFSD3 | 3.676139432 | 2.829789873 | 1.82E-06 | 1.74E-05 |
| MYRIP | 3.675485578 | 4.037933871 | 3.59E-05 | 0.000235071 |
| ITIH3 | 3.673741136 | 3.132247353 | 0.007654236 | 0.023033915 |
| LOC111097903 | 3.673203675 | 2.855282544 | 6.78E-05 | 0.000408019 |
| LOC119877154 | 3.672960922 | 2.343721861 | 4.15E-05 | 0.000266742 |
| LOC119864170 | 3.670401709 | 0.210397021 | 0.006634565 | 0.020369872 |
| CNTNAP2 | 3.670337679 | 1.140634022 | 0.002021083 | 0.007438095 |
| LOC111094851 | 3.670200505 | -0.44218453 | 0.009239546 | 0.026991332 |
| MPZL1 | 3.668232234 | 5.006190022 | 2.38E-06 | 2.20E-05 |
| H1-0 | 3.667737648 | 7.712580607 | 4.45E-05 | 0.000283345 |
| CPAMD8 | 3.666485557 | 5.477939183 | 1.61E-07 | 2.06E-06 |
| SESTD1 | 3.663617878 | 6.299125122 | 2.15E-08 | 3.43E-07 |
| SDC3 | 3.662481634 | 6.897220474 | 1.85E-08 | 3.00E-07 |
| LOC119866762 | 3.655820493 | -0.346127608 | 0.005852366 | 0.018316003 |
| ENPP2 | 3.651930301 | 6.40308673 | 3.78E-05 | 0.0002461 |
| ACTL10 | 3.651728461 | 2.301514832 | 0.000527468 | 0.002365368 |
| FAM25A | 3.651423471 | 3.364765198 | 0.000459523 | 0.002097123 |
| LOC111096307 | 3.649663928 | 5.178285305 | 0.001327853 | 0.005203139 |
| LOC111092771 | 3.647065419 | 1.931374739 | 6.66E-05 | 0.000401861 |
| RARRES1 | 3.644080482 | 5.950208924 | 2.65E-06 | 2.42E-05 |
| CD59 | 3.642113756 | 7.047509911 | 0.000135278 | 0.000737208 |
| LOC119878447 | 3.641938662 | 1.591507972 | 0.004499604 | 0.014633842 |
| LOC106557533 | 3.641817846 | 0.340026616 | 0.005467749 | 0.017271903 |
| EDA | 3.640346978 | 2.874471657 | 0.000317851 | 0.001530412 |
| CCDC68 | 3.637762362 | 2.963680859 | 0.001869164 | 0.006946226 |
| MBOAT2 | 3.636601402 | 3.947906715 | 5.56E-05 | 0.00034413 |
| HECW2 | 3.635991996 | 5.505621992 | 1.83E-07 | 2.30E-06 |
| PKP3 | 3.635492903 | 3.654140149 | 0.00173819 | 0.006524383 |
| CDC42BPG | 3.635321362 | 5.772331971 | 0.000291856 | 0.001422716 |
| C18H11orf24 | 3.635001332 | 4.283527422 | 7.02E-08 | 9.90E-07 |
| DCXR | 3.633576836 | 3.654253791 | 4.16E-07 | 4.76E-06 |
| LOC119869818 | 3.631263098 | 0.739711043 | 0.008093314 | 0.024130746 |
| GLP1R | 3.63025879 | 2.848459457 | 0.001449331 | 0.005596179 |
| PLSCR4 | 3.628951467 | 4.914120637 | 7.04E-06 | 5.59E-05 |
| RASGEF1C | 3.62732472 | 4.874834529 | 0.000194364 | 0.001001296 |
| FRAS1 | 3.626658152 | 5.079896754 | 0.00067738 | 0.002929606 |
| LOC119866119 | 3.623613582 | -0.110202373 | 0.002166063 | 0.007885952 |
| CA11 | 3.623585813 | 4.160872093 | 4.85E-07 | 5.44E-06 |

| | | | | |
|---|---|---|---|---|
| LOC119870751 | 3.622004155 | 0.135219541 | 0.002280686 | 0.008234102 |
| LOC119868486 | 3.618637488 | 0.714149775 | 0.002018073 | 0.007429538 |
| TSNAXIP1 | 3.618023215 | 1.345101646 | 0.000187303 | 0.0009693 |
| LOC102154279 | 3.617983258 | 1.784151797 | 0.001092636 | 0.004416395 |
| LOC102156180 | 3.615632297 | 3.059889268 | 0.000187991 | 0.000972396 |
| LOC102156678 | 3.615572644 | 0.8328845 | 0.000446045 | 0.002044655 |
| GIPC1 | 3.615440331 | 4.57097896 | 1.31E-08 | 2.18E-07 |
| LRP12 | 3.61489458 | 3.876777914 | 6.09E-06 | 4.93E-05 |
| GRPR | 3.613554276 | 1.468005254 | 0.007124492 | 0.021653222 |
| VGF | 3.61321867 | 3.192591401 | 0.003289851 | 0.011253306 |
| LOC111094521 | 3.612343096 | 0.136731615 | 0.006079619 | 0.01891796 |
| LOC119865587 | 3.612056556 | 0.785979147 | 0.000147099 | 0.000789713 |
| ZNF512B | 3.610491299 | 6.56927567 | 1.81E-06 | 1.73E-05 |
| PPFIBP1 | 3.610193456 | 6.313192165 | 2.48E-07 | 3.03E-06 |
| GAS2 | 3.60887124 | 1.728595169 | 0.018182266 | 0.04716508 |
| LOC102157396 | 3.605365728 | 0.502890955 | 0.003815588 | 0.012765551 |
| LOC119869807 | 3.605031238 | 4.142592692 | 0.000137482 | 0.000746969 |
| CDC42BPA | 3.604812606 | 7.552986483 | 4.73E-08 | 6.99E-07 |
| MCAM | 3.602537376 | 7.554133589 | 0.000218876 | 0.001111195 |
| CCDC191 | 3.601007533 | 5.203247961 | 0.000140017 | 0.000758414 |
| REM1 | 3.59770577 | 3.625303761 | 7.80E-05 | 0.0004621 |
| ERG | 3.597375342 | 5.164901267 | 7.74E-10 | 1.75E-08 |
| RASL10B | 3.596723092 | 3.46351825 | 1.89E-05 | 0.000133885 |
| BCL2L2 | 3.595713184 | 2.826419892 | 4.17E-07 | 4.77E-06 |
| SEMA3B | 3.594237388 | 7.658509398 | 2.72E-07 | 3.27E-06 |
| CCN4 | 3.592652062 | 3.022859057 | 0.003138333 | 0.010822132 |
| LOC119867106 | 3.589240402 | 0.367308926 | 0.000133007 | 0.000726477 |
| LOC102156621 | 3.587526158 | -0.343398267 | 0.000465836 | 0.002121468 |
| IDO1 | 3.58687078 | 2.289910051 | 0.016073957 | 0.042750137 |
| ANGEL1 | 3.586130154 | 3.46266909 | 6.17E-06 | 4.98E-05 |
| RBP4 | 3.584696342 | 5.940711317 | 0.000631165 | 0.002757355 |
| PPP1R3G | 3.584244157 | 0.21152166 | 0.009883274 | 0.028582891 |
| CAVIN2 | 3.583947559 | 5.980016967 | 1.12E-05 | 8.41E-05 |
| LOC608913 | 3.583488629 | 0.049492756 | 0.000607872 | 0.002669047 |
| SAXO2 | 3.581883109 | 1.946116043 | 0.000303736 | 0.001473005 |
| SORBS3 | 3.581428955 | 8.517431405 | 2.04E-06 | 1.92E-05 |
| LOC119864243 | 3.581375242 | 3.200490955 | 0.000140405 | 0.000759798 |
| TIGD5 | 3.579362367 | 4.67852668 | 1.99E-09 | 4.07E-08 |
| SATL1 | 3.57808407 | -1.06385373 | 0.006687975 | 0.020513514 |

| | | | | |
|---|---|---|---|---|
| PSD2 | 3.578004634 | 4.686717061 | 5.04E-07 | 5.63E-06 |
| MORN2 | 3.577788098 | -0.559459593 | 0.001266938 | 0.005009732 |
| VLDLR | 3.576252663 | 4.318804327 | 0.000310962 | 0.001503677 |
| CPNE7 | 3.575515204 | 4.936981288 | 0.002560003 | 0.009082166 |
| POGLUT3 | 3.575207041 | 6.458377182 | 5.42E-08 | 7.88E-07 |
| PPP1R26 | 3.57418096 | 5.345615963 | 3.53E-07 | 4.12E-06 |
| SCN4B | 3.574012128 | 3.56308004 | 0.006818807 | 0.020870499 |
| FOXC1 | 3.573172653 | 5.25271937 | 0.000371247 | 0.00174838 |
| LOC119867359 | 3.57253045 | 2.235636881 | 6.15E-05 | 0.000375986 |
| CAMK2N1 | 3.571317139 | 3.236347984 | 5.31E-05 | 0.000331197 |
| C3H4orf48 | 3.570602077 | 2.979631036 | 9.68E-06 | 7.41E-05 |
| PMEPA1 | 3.566682898 | 7.058005344 | 3.19E-06 | 2.82E-05 |
| SLC38A5 | 3.564048097 | 3.155227886 | 3.88E-05 | 0.000251638 |
| NFIC | 3.563745791 | 8.495316954 | 1.08E-06 | 1.10E-05 |
| LOC106559462 | 3.560519727 | 3.450849485 | 0.000116004 | 0.000649653 |
| LOC119877838 | 3.559779053 | 2.316506412 | 0.003517403 | 0.011913216 |
| LOC102156440 | 3.557066773 | 1.280194285 | 0.007078152 | 0.021536542 |
| LOC119877595 | 3.55528203 | 4.95916845 | 7.12E-07 | 7.59E-06 |
| LOC111096956 | 3.554878038 | 1.999247655 | 0.004546083 | 0.014762849 |
| XG | 3.554377726 | 6.751799594 | 1.70E-07 | 2.16E-06 |
| CCL8 | 3.55269201 | 0.366080789 | 0.006064634 | 0.018884887 |
| MRAP2 | 3.550881746 | 0.823219398 | 0.003335594 | 0.011384619 |
| MAP3K7CL | 3.5479033 | 1.66862192 | 0.007566566 | 0.022814439 |
| LOC119873193 | 3.547271768 | 0.009825578 | 0.000626401 | 0.002740283 |
| CHP2 | 3.547061987 | 2.465567296 | 0.0001857 | 0.00096284 |
| TLX2 | 3.544953422 | 2.842539086 | 0.013046286 | 0.03604234 |
| MLF1 | 3.542736692 | 4.177236587 | 2.93E-07 | 3.50E-06 |
| LOC111096026 | 3.542507889 | -0.640742713 | 0.013630212 | 0.037369714 |
| FHL1 | 3.540849767 | 7.237589622 | 0.000355617 | 0.001683918 |
| LOC102156557 | 3.540809635 | 0.264512668 | 0.000809774 | 0.003411124 |
| LOC489640 | 3.538496034 | 2.504771906 | 0.000128902 | 0.000708878 |
| USP54 | 3.536759816 | 5.725709032 | 1.03E-08 | 1.77E-07 |
| ECRG4 | 3.536613239 | 2.10494696 | 0.012103772 | 0.03380172 |
| PBX1 | 3.53603707 | 6.798369987 | 4.01E-06 | 3.44E-05 |
| OGN | 3.535635362 | 4.607110989 | 0.002077182 | 0.00760709 |
| ESR2 | 3.532817 | 2.41039624 | 0.011419736 | 0.032218164 |
| LOC102155783 | 3.532380253 | 0.820604289 | 5.15E-05 | 0.000321985 |
| TSPAN12 | 3.530478018 | 4.060597101 | 3.35E-07 | 3.94E-06 |
| LOC111096205 | 3.530369355 | 0.733587707 | 0.014542706 | 0.039462134 |

| | | | | |
|---|---|---|---|---|
| B9D1 | 3.528727318 | 3.109259491 | 1.22E-05 | 9.08E-05 |
| CDRT1 | 3.527109101 | 2.478071669 | 0.004814685 | 0.015491191 |
| FOXF1 | 3.525951111 | 2.769729374 | 4.16E-05 | 0.000266958 |
| TSPAN6 | 3.524964671 | 3.221562695 | 0.000327201 | 0.001567572 |
| MMP23 | 3.52365285 | 4.213839634 | 0.000128664 | 0.00070793 |
| LOC111090100 | 3.521525232 | 0.957827019 | 0.000419739 | 0.001945053 |
| LOC119870368 | 3.518452621 | 4.941034342 | 2.64E-06 | 2.41E-05 |
| STAB1 | 3.518046044 | 7.61446428 | 2.87E-08 | 4.45E-07 |
| LOC119871051 | 3.516388096 | 2.42195533 | 0.005263525 | 0.016711694 |
| LOC111097489 | 3.515960531 | 2.341202682 | 1.71E-06 | 1.65E-05 |
| LOC119876363 | 3.513755725 | 1.107213982 | 0.000219021 | 0.001111411 |
| NACAD | 3.5133328 | 4.906284028 | 9.97E-05 | 0.000570796 |
| PRSS54 | 3.512756318 | -0.106811954 | 0.009438718 | 0.027469737 |
| PAK4 | 3.510973838 | 5.555988997 | 2.14E-06 | 1.99E-05 |
| LOC119869669 | 3.509266 | 2.985957038 | 0.002110389 | 0.007713063 |
| SLC43A1 | 3.509190006 | 4.198506455 | 2.68E-06 | 2.43E-05 |
| LOC119864619 | 3.508428022 | -0.338521032 | 0.000881777 | 0.003679389 |
| THSD1 | 3.508391908 | 4.079766523 | 2.08E-06 | 1.95E-05 |
| SLC45A3 | 3.508005373 | 2.13513196 | 0.001509725 | 0.005796329 |
| LOC111096182 | 3.506596038 | 1.983670705 | 0.000313812 | 0.001514756 |
| CHTF18 | 3.504030599 | 3.6192261 | 9.89E-06 | 7.54E-05 |
| LOC102156872 | 3.503859822 | 1.238240677 | 0.002535741 | 0.009009363 |
| MTSS2 | 3.502529742 | 6.245988137 | 1.17E-07 | 1.56E-06 |
| SLCO1C1 | 3.501310041 | 1.9165847 | 0.018302496 | 0.047408867 |
| LOC111090873 | 3.500935844 | 1.923451177 | 0.000675873 | 0.002924958 |
| PFN2 | 3.500149866 | 5.009074942 | 1.49E-07 | 1.93E-06 |
| RAB6B | 3.498631524 | 3.002799132 | 3.80E-06 | 3.28E-05 |
| NOTUM | 3.49684097 | 2.943063571 | 2.60E-05 | 0.000177782 |
| ITIH4 | 3.494422642 | 5.244121933 | 0.003930305 | 0.013096786 |
| KCTD19 | 3.492008112 | 2.342349637 | 1.50E-05 | 0.000109127 |
| LOC111094523 | 3.491574012 | 0.826776716 | 0.000240752 | 0.001203092 |
| LOC119870553 | 3.491262766 | 5.65070783 | 0.000320899 | 0.001542776 |
| LOC611908 | 3.488134868 | -0.444583121 | 0.009188836 | 0.026868544 |
| SYNGAP1 | 3.483503336 | 5.535381066 | 2.45E-08 | 3.86E-07 |
| AGAP3 | 3.482687174 | 5.209616783 | 4.22E-08 | 6.27E-07 |
| LOC119869926 | 3.481225549 | 0.3524825 | 0.001757734 | 0.006587475 |
| RARRES2 | 3.480816901 | 5.262869671 | 2.56E-10 | 6.37E-09 |
| SLC1A1 | 3.478726831 | 2.112418891 | 0.004022915 | 0.013347925 |
| FAM171A2 | 3.477523866 | 2.371080965 | 0.000335268 | 0.001599866 |

| | | | | |
|---|---|---|---|---|
| NEK10 | 3.477201921 | 3.290382303 | 3.98E-08 | 5.96E-07 |
| INAVA | 3.476127077 | 2.585294462 | 0.012954003 | 0.035833069 |
| HDAC11 | 3.475719422 | 4.298913932 | 8.16E-08 | 1.13E-06 |
| LOC106558144 | 3.475103592 | -1.317583022 | 0.007667736 | 0.023061734 |
| MISP | 3.473505795 | 0.98952159 | 0.004185417 | 0.013781607 |
| LOC111096252 | 3.470303654 | 0.667725837 | 0.000119602 | 0.000666043 |
| RN7SL1 | 3.466170671 | 14.0433254 | 0.000473165 | 0.002148976 |
| CDNF | 3.466030264 | -0.101507022 | 0.002651129 | 0.00936865 |
| RNF207 | 3.463569479 | 2.56596773 | 0.000921165 | 0.003824716 |
| TNFRSF6B | 3.462336257 | 5.155122708 | 0.00026053 | 0.001285917 |
| DRC3 | 3.461763146 | 1.221589637 | 0.0002232 | 0.001127863 |
| LOC119876103 | 3.461022334 | -0.651672616 | 0.017275063 | 0.045261416 |
| CREB3L3 | 3.459413057 | 2.066927481 | 0.002670299 | 0.009425635 |
| ALOX15B | 3.458773419 | 0.512266874 | 0.008458306 | 0.025033029 |
| LOC111096475 | 3.457001536 | 1.799932638 | 0.000150129 | 0.00080201 |
| MN1 | 3.456133556 | 4.727495292 | 7.55E-05 | 0.000449505 |
| LOC119876467 | 3.4554368 | 0.355777052 | 0.002037113 | 0.007490657 |
| SEC31B | 3.451815983 | 2.167159954 | 0.000107218 | 0.000607513 |
| LOC111097140 | 3.450870255 | 0.28685379 | 0.000175994 | 0.000919875 |
| SPA17 | 3.450014214 | 1.849658586 | 3.54E-06 | 3.09E-05 |
| STXBP1 | 3.449521498 | 6.320139638 | 1.34E-06 | 1.32E-05 |
| ADAM11 | 3.448137987 | 1.534992241 | 0.00119465 | 0.004768194 |
| CC2D2A | 3.447744967 | 5.693540663 | 4.54E-09 | 8.44E-08 |
| LOC102154792 | 3.445838547 | 1.333874927 | 0.000768925 | 0.003263149 |
| FOXD1 | 3.445720901 | 3.041840914 | 0.001969905 | 0.007278171 |
| LOC119869431 | 3.445604641 | 0.43255452 | 0.008068423 | 0.024069778 |
| MEX3D | 3.444818159 | 4.563608929 | 2.80E-06 | 2.53E-05 |
| SYN2 | 3.444790994 | 2.188067537 | 0.000563105 | 0.002498807 |
| CDK15 | 3.443340717 | 1.555897785 | 0.017633041 | 0.046015762 |
| KCNC3 | 3.442806278 | 4.809532336 | 0.000240063 | 0.001201059 |
| LOC111092270 | 3.442751781 | -0.387430403 | 0.002820394 | 0.009887111 |
| AFAP1 | 3.440322922 | 6.903702388 | 3.64E-12 | 1.39E-10 |
| LDLR | 3.440080448 | 5.473566027 | 1.75E-05 | 0.000125371 |
| SDC2 | 3.438341941 | 6.116918605 | 4.76E-06 | 3.99E-05 |
| RASIP1 | 3.437318412 | 5.124061852 | 2.31E-08 | 3.65E-07 |
| LOC119871313 | 3.434352471 | 0.187031257 | 0.012512534 | 0.034789501 |
| DZIP1 | 3.433513053 | 5.496412363 | 1.32E-06 | 1.31E-05 |
| AHDC1 | 3.433023459 | 6.883868054 | 6.22E-09 | 1.12E-07 |
| LOC111091235 | 3.432633151 | 4.436126662 | 1.08E-06 | 1.10E-05 |

| | | | | |
|---|---|---|---|---|
| PROCR | 3.43114031 | 2.504054309 | 0.000896617 | 0.003732873 |
| UNC5B | 3.4300054 | 5.441467786 | 5.17E-05 | 0.000323312 |
| PANK1 | 3.429063149 | 1.384988417 | 0.003125264 | 0.010785643 |
| TJP3 | 3.4282515 | 3.434446807 | 0.007813918 | 0.023429861 |
| CLIC2 | 3.427689642 | 3.20442974 | 6.54E-05 | 0.000396365 |
| LOC119871139 | 3.427485611 | 2.029421946 | 0.000931049 | 0.003859724 |
| LOC476620 | 3.42504654 | 0.13992134 | 0.002501441 | 0.008910869 |
| SLCO2B1 | 3.424427837 | 5.178026293 | 1.89E-06 | 1.79E-05 |
| FZD7 | 3.423461071 | 4.871131987 | 5.45E-05 | 0.000338596 |
| LOC119866386 | 3.422238186 | 2.769812574 | 0.015816106 | 0.04221979 |
| LOC111097662 | 3.418805326 | 1.030418909 | 0.01328796 | 0.036593386 |
| ANO4 | 3.418511159 | 0.840011509 | 0.016059205 | 0.042726633 |
| CRIP2 | 3.418165222 | 5.555558903 | 9.98E-09 | 1.72E-07 |
| S100A13 | 3.417956578 | 3.618694256 | 0.000366315 | 0.001728157 |
| SLC24A5 | 3.417227138 | 3.052668055 | 5.86E-05 | 0.00036073 |
| PTPDC1 | 3.416379116 | 3.925342567 | 8.33E-05 | 0.000488703 |
| PDZD3 | 3.415275433 | 1.804713051 | 7.07E-05 | 0.000423755 |
| LOC102152253 | 3.413790592 | 2.386447113 | 0.001408191 | 0.005460678 |
| LOC119869726 | 3.41246778 | 2.934192313 | 5.85E-05 | 0.000359826 |
| LOXL1 | 3.40995544 | 5.309155246 | 4.03E-07 | 4.64E-06 |
| CD99L2 | 3.407796167 | 4.485515617 | 5.62E-06 | 4.61E-05 |
| LOC111092735 | 3.406515186 | 2.983491888 | 4.08E-07 | 4.68E-06 |
| SERPINE1 | 3.405960543 | 4.409309305 | 0.000122347 | 0.000678986 |
| ASS1 | 3.405918699 | -0.089635386 | 0.019411028 | 0.049786416 |
| LOC111090530 | 3.405556835 | -0.311950195 | 0.000355356 | 0.001683051 |
| GNA11 | 3.405098626 | 4.367180678 | 4.26E-08 | 6.33E-07 |
| CASKIN1 | 3.404244913 | 2.730195348 | 0.000170155 | 0.000894002 |
| LOC102155889 | 3.403596302 | 2.113414573 | 0.000126454 | 0.000697788 |
| EMILIN1 | 3.397643577 | 6.108146963 | 4.56E-06 | 3.85E-05 |
| ACVR1 | 3.396829593 | 5.27464199 | 2.49E-09 | 4.96E-08 |
| MAPRE3 | 3.396770684 | 5.238091047 | 3.17E-06 | 2.82E-05 |
| GPR27 | 3.396723071 | 3.676216369 | 0.008158516 | 0.02428506 |
| TMEM82 | 3.396090552 | 1.09034751 | 0.000547141 | 0.002436947 |
| F8 | 3.394925772 | 3.628642944 | 0.00031152 | 0.001506037 |
| TMPRSS9 | 3.394750638 | 1.919003904 | 0.002149397 | 0.007835809 |
| LOC111093309 | 3.39453756 | 1.741822314 | 0.003525022 | 0.011931565 |
| PTK2 | 3.393761798 | 6.875695855 | 3.76E-08 | 5.67E-07 |
| RBPMS | 3.390972852 | 5.330446763 | 5.94E-07 | 6.50E-06 |
| FGF2 | 3.386168972 | 6.129299687 | 4.14E-05 | 0.000266098 |

| | | | | |
|---|---|---|---|---|
| LOC102153760 | 3.383632483 | 1.284208269 | 0.002289855 | 0.00826112 |
| PAPLN | 3.381727346 | 7.549107104 | 0.000222776 | 0.001126483 |
| NPC1L1 | 3.379599357 | 1.312879396 | 0.001285594 | 0.005072369 |
| LOC119876690 | 3.378595634 | 4.877869218 | 0.004225588 | 0.013878037 |
| LOC119872641 | 3.375554373 | 0.828984609 | 8.39E-05 | 0.000492095 |
| ZNF483 | 3.373191269 | 3.721224316 | 3.45E-05 | 0.000226984 |
| LOC119875648 | 3.372507073 | 1.622216276 | 8.93E-05 | 0.000518211 |
| LOC111098860 | 3.368821769 | 2.474827506 | 0.000597866 | 0.002630442 |
| IFT140 | 3.366650324 | 4.365598111 | 7.74E-08 | 1.08E-06 |
| TULP3 | 3.36537352 | 4.731389342 | 9.73E-08 | 1.32E-06 |
| SSTR1 | 3.36440047 | 0.310557985 | 0.014449797 | 0.039278873 |
| PLAAT3 | 3.361612082 | 5.773535199 | 7.72E-07 | 8.16E-06 |
| TCEAL8 | 3.360035247 | 3.126790694 | 1.68E-05 | 0.000120755 |
| SPTBN1 | 3.357277889 | 11.10458565 | 2.99E-06 | 2.68E-05 |
| LOC111090944 | 3.356685355 | -0.734025884 | 0.011172701 | 0.031636561 |
| NUDT2 | 3.356637111 | 1.331461014 | 2.72E-05 | 0.000184301 |
| LOC119864207 | 3.35539783 | 2.307027159 | 0.000973901 | 0.004000455 |
| LOC119878269 | 3.3537741 | 7.64753415 | 0.001266136 | 0.005007476 |
| LOC474655 | 3.353534382 | -0.075586597 | 0.012644201 | 0.035088062 |
| SLC9B2 | 3.353289516 | 3.731524787 | 0.004276055 | 0.014007699 |
| ADCY3 | 3.351700966 | 5.1598295 | 4.07E-05 | 0.000262408 |
| GRIP1 | 3.351465253 | 3.551732759 | 0.002544161 | 0.009033355 |
| B4GALT2 | 3.350182623 | 4.624989471 | 9.75E-08 | 1.33E-06 |
| RUSC2 | 3.349809201 | 6.629028174 | 1.32E-07 | 1.74E-06 |
| LOC111090957 | 3.349538162 | 3.023019041 | 0.000620121 | 0.002715675 |
| TMEM232 | 3.348709598 | 2.85755033 | 4.21E-06 | 3.59E-05 |
| CTSK | 3.348095378 | 7.312412586 | 2.06E-06 | 1.93E-05 |
| CADM1 | 3.344603602 | 4.702820364 | 2.85E-06 | 2.57E-05 |
| ZNF205 | 3.343244798 | 4.492393278 | 6.38E-05 | 0.00038773 |
| CACNA1B | 3.342879228 | 2.326419586 | 0.001107621 | 0.00446946 |
| LOC102151658 | 3.342315709 | 3.159531772 | 7.63E-05 | 0.00045345 |
| KCTD15 | 3.341645927 | 4.514459022 | 6.59E-05 | 0.000398592 |
| MID2 | 3.34097447 | 5.40361658 | 1.37E-07 | 1.79E-06 |
| LOC119875994 | 3.340366802 | 2.74284437 | 8.87E-05 | 0.000514993 |
| KCTD9 | 3.339347594 | 5.741918965 | 2.92E-06 | 2.62E-05 |
| LOC119866432 | 3.338876592 | 2.49337859 | 5.50E-05 | 0.000340953 |
| LOC119865205 | 3.338062162 | 1.426449511 | 0.004116234 | 0.013588903 |
| LOC119870121 | 3.337815911 | 2.796223278 | 0.00089291 | 0.003720831 |
| LOC102151164 | 3.332429563 | 0.574149599 | 0.001580208 | 0.006027427 |

| | | | | |
|---|---|---|---|---|
| LOC119866898 | 3.330649049 | 4.441672874 | 3.23E-06 | 2.86E-05 |
| ZNF579 | 3.328073065 | 4.016557647 | 1.82E-05 | 0.000129489 |
| HR | 3.325016207 | 4.138409909 | 0.000142907 | 0.000771411 |
| FZD3 | 3.3247756 | 2.850502992 | 0.000956447 | 0.00394164 |
| SAMD4A | 3.324763373 | 5.996421295 | 1.53E-06 | 1.49E-05 |
| TPRN | 3.321364411 | 4.040083159 | 4.20E-06 | 3.58E-05 |
| ABCC4 | 3.320898888 | 3.445631847 | 0.002052277 | 0.007533088 |
| ZNF503 | 3.32075798 | 5.261510671 | 2.45E-06 | 2.25E-05 |
| FER1L6 | 3.316485964 | 1.590272768 | 0.000855834 | 0.003582173 |
| SLC27A1 | 3.312562963 | 6.826346292 | 1.81E-06 | 1.73E-05 |
| TMEM238 | 3.311734723 | 0.201710878 | 0.002508605 | 0.008934921 |
| PAK3 | 3.309618949 | 4.337562049 | 8.40E-05 | 0.000492305 |
| LOC119876067 | 3.308812448 | 1.055743599 | 0.002516593 | 0.008956014 |
| BAIAP2 | 3.306735174 | 7.396169413 | 7.91E-05 | 0.000467663 |
| LOC106559541 | 3.303983224 | 1.947196259 | 5.63E-06 | 4.61E-05 |
| FAM161A | 3.30254312 | 3.578998175 | 4.76E-05 | 0.00030088 |
| RNLS | 3.301099928 | 3.781817398 | 2.28E-07 | 2.81E-06 |
| RASL11A | 3.299892814 | 2.820254754 | 0.000922449 | 0.003829198 |
| DYNLRB2 | 3.297208265 | 0.669469991 | 0.001300865 | 0.005118671 |
| BAALC | 3.295528322 | 1.501521858 | 0.002023178 | 0.007444541 |
| CDC42BPB | 3.294093165 | 7.361284391 | 7.90E-08 | 1.09E-06 |
| SELENOP | 3.293884462 | 9.954144024 | 8.95E-06 | 6.91E-05 |
| LOC111091687 | 3.291929003 | 0.249273435 | 0.019267398 | 0.04948244 |
| SYNDIG1 | 3.291623713 | 1.857235674 | 0.004970232 | 0.015916052 |
| LOC119873493 | 3.28979901 | 0.218520855 | 0.000552979 | 0.002459413 |
| SETBP1 | 3.285291632 | 6.750851389 | 3.67E-08 | 5.55E-07 |
| FKBP14 | 3.284793288 | 3.293878941 | 0.001016238 | 0.00415173 |
| LOC106557811 | 3.283414208 | 2.133970973 | 0.001651875 | 0.006253466 |
| LOC106557956 | 3.2833924 | 1.541567729 | 0.002246842 | 0.008132908 |
| GABBR2 | 3.28287443 | 3.982335522 | 0.000908752 | 0.003778853 |
| RND2 | 3.282611037 | 2.014301528 | 0.002757179 | 0.009700706 |
| ACAP3 | 3.280708087 | 5.793792823 | 2.22E-06 | 2.06E-05 |
| LRRN4CL | 3.27981119 | 3.737333896 | 3.90E-06 | 3.36E-05 |
| FGD6 | 3.275630236 | 5.562889071 | 1.19E-08 | 2.01E-07 |
| BMPR1A | 3.274681453 | 4.255823419 | 8.38E-07 | 8.76E-06 |
| CORIN | 3.271111618 | 4.32385444 | 0.00908247 | 0.026604186 |
| RGS4 | 3.269094277 | 2.075327357 | 0.003698163 | 0.01243604 |
| LOC111098565 | 3.268537806 | 0.234747997 | 0.009646789 | 0.027977366 |
| LOC119871531 | 3.265687742 | 2.308397386 | 4.23E-05 | 0.000270719 |

| | | | | |
|---|---|---|---|---|
| ENOSF1 | 3.264589849 | 1.412981533 | 0.000103583 | 0.000590888 |
| MIR568 | 3.264192324 | 1.898844329 | 0.001974323 | 0.007289522 |
| NR2F6 | 3.264025125 | 3.149049489 | 8.07E-07 | 8.48E-06 |
| LOC102155683 | 3.263025087 | 2.134996985 | 0.013018288 | 0.035974162 |
| LOC102154996 | 3.261087864 | 0.843152372 | 0.011201605 | 0.03168023 |
| ASAP2 | 3.260819476 | 6.194263448 | 1.71E-06 | 1.64E-05 |
| IFT81 | 3.260459384 | 3.127492626 | 2.48E-05 | 0.000170705 |
| CCDC78 | 3.258850275 | 4.618856308 | 1.41E-05 | 0.000103598 |
| LOC119869901 | 3.257791783 | 3.938523686 | 0.00364271 | 0.01227051 |
| IFT43 | 3.256729731 | 3.04015667 | 3.42E-06 | 3.00E-05 |
| BACE1 | 3.255943927 | 6.393425465 | 2.48E-10 | 6.21E-09 |
| C7H1orf105 | 3.255835449 | 0.303561612 | 0.001654648 | 0.006262869 |
| TRPV6 | 3.255799691 | -0.135366424 | 0.00414874 | 0.013677464 |
| MAGI2 | 3.25565271 | 4.099052539 | 0.000378046 | 0.001776293 |
| ADCY6 | 3.25197652 | 5.662528074 | 5.54E-08 | 8.02E-07 |
| DTX1 | 3.251666904 | 4.088388341 | 0.000533879 | 0.002387199 |
| LOC111098972 | 3.247207911 | -0.786131163 | 0.000965198 | 0.003971662 |
| LOC100682937 | 3.2471829 | 1.757521694 | 0.004756304 | 0.015328384 |
| LOC119876510 | 3.246778735 | 0.738087192 | 0.000564396 | 0.002503125 |
| DAPK3 | 3.245650254 | 5.973594466 | 7.15E-06 | 5.66E-05 |
| LOC119873974 | 3.244798026 | 2.010938798 | 0.014304509 | 0.038932767 |
| NPL | 3.243933358 | 4.559788994 | 3.53E-07 | 4.12E-06 |
| GPR150 | 3.242751197 | -0.866360735 | 0.004467082 | 0.014541165 |
| LOC119867936 | 3.242300413 | -0.330498205 | 0.003168185 | 0.010904252 |
| RCN1 | 3.240359381 | 5.898506446 | 4.75E-06 | 3.98E-05 |
| ALX3 | 3.238742288 | 2.34241068 | 0.010640345 | 0.030386631 |
| RECQL4 | 3.237988421 | 3.769645981 | 0.000249659 | 0.001241592 |
| MYO15B | 3.236543773 | 4.25346944 | 0.001197086 | 0.004775345 |
| TRPC4 | 3.236245473 | 1.576501739 | 0.004034678 | 0.013378761 |
| WNT2B | 3.236155613 | 3.052410678 | 1.05E-07 | 1.41E-06 |
| CBARP | 3.234128254 | 3.733291455 | 0.000219954 | 0.001114842 |
| LOC119870959 | 3.234120266 | 0.440045169 | 0.003516157 | 0.011910858 |
| SSC4D | 3.233330075 | -0.427749158 | 0.002333692 | 0.008379408 |
| B3GNT9 | 3.232937483 | 4.366438721 | 2.90E-07 | 3.47E-06 |
| HSD11B1L | 3.232272318 | 3.456282385 | 0.00030571 | 0.001481254 |
| ZDHHC8 | 3.231198865 | 6.706905066 | 7.14E-06 | 5.66E-05 |
| POFUT2 | 3.229527157 | 4.737901437 | 8.69E-08 | 1.20E-06 |
| LOC111092325 | 3.228633649 | -0.248620822 | 0.005987778 | 0.018683144 |
| LOC100686848 | 3.226813548 | 0.31002435 | 0.018248071 | 0.047301812 |

| | | | | |
|---|---|---|---|---|
| LOC119864306 | 3.22639954 | 4.886501182 | 0.001885415 | 0.006991037 |
| CTF1 | 3.223637957 | 0.630703654 | 0.00603008 | 0.018790786 |
| TRIQK | 3.223493198 | 3.435554088 | 4.28E-06 | 3.63E-05 |
| PKP2 | 3.222998319 | 2.732518625 | 1.55E-05 | 0.00011242 |
| KLHL29 | 3.222804502 | 5.380759911 | 0.00035455 | 0.00168039 |
| LOC119873710 | 3.219835 | 3.132686649 | 0.000249484 | 0.001241292 |
| LOC119865625 | 3.219148959 | 1.512869746 | 0.017775199 | 0.046330955 |
| BACE2 | 3.218584293 | 5.333075859 | 7.35E-07 | 7.83E-06 |
| LOC102156662 | 3.216332315 | 1.247582716 | 0.000398026 | 0.001857138 |
| MACROH2A2 | 3.214493223 | 1.943707289 | 0.000655028 | 0.002849505 |
| LOC119875996 | 3.214186735 | 3.655230327 | 4.80E-07 | 5.40E-06 |
| LOC119866957 | 3.21388812 | 0.792181673 | 0.008547054 | 0.025258172 |
| MAG | 3.21353131 | 2.189949274 | 0.007458748 | 0.022533239 |
| MIR8877 | 3.212766496 | -0.097308954 | 0.001193048 | 0.004764498 |
| CCDC183 | 3.211918218 | 1.688246948 | 0.00070029 | 0.003013179 |
| PLXNA3 | 3.208852923 | 5.494464463 | 6.87E-06 | 5.48E-05 |
| LRRC56 | 3.208199338 | 1.975978684 | 0.001241269 | 0.004918102 |
| DOCK9 | 3.205444263 | 8.818387656 | 1.05E-06 | 1.07E-05 |
| GAPDHS | 3.204282725 | 2.245786113 | 0.001727739 | 0.006490775 |
| EPM2A | 3.201293648 | 1.998018938 | 1.85E-05 | 0.000131423 |
| RAB3B | 3.200772908 | 3.608786024 | 7.97E-06 | 6.25E-05 |
| TMEM151B | 3.197961664 | 4.878779505 | 1.57E-07 | 2.01E-06 |
| LOC106558104 | 3.196888267 | 2.625734396 | 0.00191334 | 0.007087305 |
| FHDC1 | 3.196189447 | 2.614586714 | 0.001265882 | 0.005007384 |
| LOC119864718 | 3.195397467 | 1.271219255 | 0.000788232 | 0.003332685 |
| PCSK4 | 3.195255074 | 0.625547771 | 0.010771292 | 0.030688342 |
| LOC119873935 | 3.193397853 | 1.394108138 | 0.000838706 | 0.003521364 |
| LOC611835 | 3.192921056 | 4.518201016 | 3.22E-07 | 3.81E-06 |
| JHY | 3.192901006 | 0.47615389 | 0.002521697 | 0.008971231 |
| LOC102152726 | 3.190792544 | 2.193440156 | 0.000440576 | 0.002024294 |
| HSPA2 | 3.190056645 | 6.243937573 | 0.000181199 | 0.000942661 |
| HTR1F | 3.187060315 | 3.400743145 | 0.005439134 | 0.017206591 |
| CC2D2B | 3.186169201 | 0.705067515 | 0.013909012 | 0.038068448 |
| SMPDL3A | 3.185695392 | 5.792368525 | 0.000581503 | 0.002567307 |
| MYO1E | 3.184983661 | 6.814281254 | 9.51E-09 | 1.65E-07 |
| MGARP | 3.184443755 | 0.922542247 | 0.000158565 | 0.000841681 |
| LOC119867832 | 3.182170871 | 1.125931541 | 0.007475229 | 0.022576732 |
| MARVELD3 | 3.181876573 | -0.684308442 | 0.00928065 | 0.027090346 |
| HES4 | 3.181168119 | 3.103772671 | 0.000331111 | 0.001582897 |

| | | | | |
|---|---|---|---|---|
| LOC119877521 | 3.181067657 | -0.684413783 | 0.011961596 | 0.033463751 |
| LOC102151574 | 3.180186283 | 0.385442714 | 0.000373602 | 0.001757942 |
| ADAMTS7 | 3.179333279 | 4.448063354 | 3.99E-05 | 0.000257585 |
| CBX6 | 3.177056058 | 7.336812938 | 1.21E-07 | 1.60E-06 |
| SLC10A6 | 3.176778675 | 3.415205713 | 1.16E-05 | 8.69E-05 |
| LZTFL1 | 3.174638183 | 3.714139849 | 4.23E-05 | 0.000270719 |
| ARNTL2 | 3.174570738 | 4.724207844 | 0.000143787 | 0.000775198 |
| DEPP1 | 3.174036519 | 3.117741996 | 7.51E-06 | 5.92E-05 |
| LOC102154710 | 3.173031369 | 1.968581718 | 0.000215001 | 0.001094858 |
| S100B | 3.170641352 | 5.254169523 | 0.00172351 | 0.006476009 |
| TMEM25 | 3.169042344 | 2.021920454 | 0.000508807 | 0.002292523 |
| LOC119869520 | 3.168314167 | 2.156968995 | 0.001664735 | 0.00629665 |
| BRSK2 | 3.167516917 | 4.670365716 | 3.46E-06 | 3.03E-05 |
| PRPF40B | 3.167231293 | 5.420595999 | 5.42E-08 | 7.88E-07 |
| WTIP | 3.166897602 | 3.315068292 | 4.94E-05 | 0.000310844 |
| LOC119874147 | 3.166866972 | 2.081825556 | 0.000608772 | 0.002671909 |
| PYROXD2 | 3.164322279 | 5.168954743 | 2.68E-08 | 4.19E-07 |
| LOC119873714 | 3.163210188 | 0.459892549 | 0.001461366 | 0.005637627 |
| PPFIA2 | 3.162001339 | 3.187494147 | 0.00616227 | 0.019132719 |
| LOC102157107 | 3.160048925 | 1.014826815 | 0.001020885 | 0.004165219 |
| LOC119878332 | 3.159831992 | 0.379466457 | 0.011535425 | 0.032494452 |
| LOC119866505 | 3.157699571 | -0.145148345 | 0.011958097 | 0.033460565 |
| LOC119871958 | 3.15630091 | 1.699882373 | 0.006302856 | 0.019503397 |
| SEMA4C | 3.154657325 | 5.349983539 | 1.33E-05 | 9.78E-05 |
| RSPH3 | 3.154597087 | 1.683653432 | 2.71E-05 | 0.000183582 |
| CDC42EP5 | 3.154380755 | 3.684229612 | 2.77E-05 | 0.000187432 |
| LOC119878547 | 3.154212269 | 2.419649193 | 0.003070471 | 0.010620223 |
| PDZD2 | 3.153890734 | 5.673473286 | 1.48E-07 | 1.91E-06 |
| LOC119869611 | 3.152978827 | 3.115682303 | 0.000240695 | 0.001203089 |
| IQCK | 3.152465731 | 3.003425772 | 7.04E-06 | 5.59E-05 |
| LOC119870564 | 3.151469108 | 2.001695614 | 0.004432716 | 0.01445535 |
| LIF | 3.151099883 | 2.674026824 | 0.000657522 | 0.002856952 |
| LRGUK | 3.150266824 | 2.124774328 | 0.000344306 | 0.00163787 |
| VWA1 | 3.147583115 | 6.004390442 | 1.71E-07 | 2.17E-06 |
| LOC119877582 | 3.145824635 | -0.404065937 | 0.00867179 | 0.025571907 |
| DAAM2 | 3.144158975 | 6.22985466 | 2.93E-05 | 0.000196849 |
| CXXC5 | 3.144053393 | 5.520883806 | 6.12E-08 | 8.79E-07 |
| DYNC2I1 | 3.143403084 | 4.141728405 | 8.29E-06 | 6.46E-05 |
| NXPH3 | 3.14193095 | 1.172292943 | 0.001414813 | 0.005481452 |

| | | | | |
|---|---|---|---|---|
| SUMO3 | 3.138297026 | 4.997281115 | 5.26E-08 | 7.68E-07 |
| CLIC4 | 3.137239622 | 8.170281327 | 3.36E-08 | 5.14E-07 |
| CYB5R3 | 3.13234233 | 7.35707932 | 8.54E-06 | 6.64E-05 |
| FSCN2 | 3.130980554 | 0.879967321 | 0.002314795 | 0.008328112 |
| VASH1 | 3.129248525 | 6.143493576 | 4.96E-05 | 0.000312234 |
| LOC119871867 | 3.128283181 | 1.925709791 | 0.016079527 | 0.042759704 |
| TRMT61A | 3.126227388 | 3.641133364 | 1.61E-06 | 1.56E-05 |
| LOC474642 | 3.125634008 | 0.481701781 | 0.003446763 | 0.011707494 |
| DUSP8 | 3.122590541 | 4.386723257 | 0.00223361 | 0.008089746 |
| EFNB1 | 3.121796893 | 5.740823283 | 7.95E-05 | 0.000469805 |
| ARHGAP33 | 3.12109729 | 4.661134871 | 2.06E-05 | 0.000144469 |
| ZNF488 | 3.120653228 | 1.381903824 | 0.009846142 | 0.028490712 |
| LOC119865986 | 3.120610056 | 0.23438189 | 0.018144282 | 0.047094734 |
| TMEM132C | 3.117845009 | 1.59592964 | 0.015304672 | 0.041128104 |
| ITGB4 | 3.115855155 | 8.945546448 | 0.000427543 | 0.001975302 |
| CENPB | 3.112757792 | 6.014767193 | 1.78E-08 | 2.89E-07 |
| LOC119872134 | 3.112694981 | 2.508831576 | 0.001013662 | 0.004141988 |
| LOC119874502_1 | 3.111123426 | -0.246574772 | 0.000729984 | 0.003122348 |
| SYNJ2 | 3.109876351 | 7.011424988 | 2.17E-09 | 4.41E-08 |
| LOC119867912 | 3.107521632 | -0.649287794 | 0.004350068 | 0.014220065 |
| SNX21 | 3.107425509 | 4.044796921 | 2.04E-05 | 0.000143408 |
| LOC119866553 | 3.10686716 | 0.861550553 | 0.000485818 | 0.002201826 |
| LOC102156046 | 3.105373064 | 4.393599465 | 0.000186873 | 0.000967874 |
| SMTNL2 | 3.10479927 | 2.943499699 | 0.005001461 | 0.01600187 |
| WDR90 | 3.104750796 | 6.171179737 | 2.48E-06 | 2.27E-05 |
| KIF1C | 3.101117602 | 7.077221422 | 1.78E-06 | 1.71E-05 |
| SNED1 | 3.100728551 | 7.100925576 | 7.45E-07 | 7.92E-06 |
| PROM1 | 3.100568052 | 2.485412645 | 0.012522535 | 0.034808375 |
| DNAL1 | 3.100513668 | 3.403288912 | 2.11E-07 | 2.62E-06 |
| FKBP9 | 3.09995807 | 6.174596399 | 1.76E-06 | 1.69E-05 |
| SEPTIN10 | 3.099734776 | 5.74858084 | 3.75E-09 | 7.13E-08 |
| OR51F23 | 3.09905552 | -0.558134812 | 0.011395509 | 0.032158187 |
| LOC102153535 | 3.098612617 | 2.626751411 | 4.30E-05 | 0.000275001 |
| LOC106559473 | 3.097566812 | 0.14045536 | 0.010147423 | 0.029226654 |
| SYNE2 | 3.096848596 | 8.493539496 | 2.52E-07 | 3.06E-06 |
| C13H8orf82 | 3.095820025 | 3.387985372 | 0.001990376 | 0.007340039 |
| LOC119863990 | 3.09385894 | 5.761763344 | 9.55E-05 | 0.000549024 |
| KCNAB3 | 3.091867982 | 2.738732779 | 0.003472837 | 0.011788064 |
| SELENOV | 3.090115949 | 0.355602688 | 0.00890483 | 0.026147458 |

| | | | | |
|---|---|---|---|---|
| PKNOX2 | 3.089667481 | 3.227572213 | 0.001026707 | 0.004181101 |
| LOC106559304 | 3.089358282 | 1.955378385 | 0.003621059 | 0.012208963 |
| LOC119871713 | 3.088654229 | 0.68502497 | 0.001927102 | 0.007133404 |
| ECM1 | 3.087277565 | 8.44566678 | 0.000523426 | 0.002349673 |
| PTRH1 | 3.08719457 | 1.644449781 | 0.000591336 | 0.002604354 |
| CSPG4B | 3.086814484 | 5.029259915 | 2.36E-06 | 2.18E-05 |
| SPIRE1 | 3.086702207 | 5.506012819 | 1.04E-05 | 7.89E-05 |
| SHISA4 | 3.086616982 | 5.633553414 | 3.32E-05 | 0.000219265 |
| RAPSN | 3.082270622 | 2.607526706 | 0.001300572 | 0.005118448 |
| NEB | 3.082154281 | 5.775235573 | 0.00011941 | 0.000665803 |
| LOC119872578 | 3.080691723 | 2.23668593 | 0.001878525 | 0.006970255 |
| LOC102157331 | 3.076811856 | 1.109929931 | 0.001554262 | 0.005946254 |
| NFIA | 3.076627869 | 7.766306747 | 1.92E-07 | 2.40E-06 |
| SV2B | 3.075956923 | 3.23672645 | 0.011341139 | 0.032021433 |
| ARTN | 3.071585543 | 1.853952195 | 0.010745731 | 0.030627596 |
| LOC102153634 | 3.071099937 | 5.217890875 | 4.62E-07 | 5.24E-06 |
| B4GALT4 | 3.068973161 | 4.114093665 | 3.35E-05 | 0.000220931 |
| LOC119873992 | 3.068559828 | 4.173031203 | 2.39E-05 | 0.00016516 |
| ABCA4 | 3.065697442 | 2.142602661 | 0.000461983 | 0.00210702 |
| DSCAML1 | 3.06526112 | 2.553331815 | 0.01003766 | 0.028955971 |
| CCDC181 | 3.064921987 | 1.449018417 | 0.000180746 | 0.000940757 |
| LOC119872301 | 3.064539011 | 1.764055188 | 0.004650738 | 0.015057581 |
| MTMR2 | 3.063119399 | 5.136785733 | 5.80E-06 | 4.73E-05 |
| OLFML3 | 3.062618956 | 6.3134073 | 0.000375567 | 0.001766039 |
| MIR8865 | 3.061643612 | 10.11852175 | 9.52E-05 | 0.000547834 |
| LOC100686546 | 3.061501837 | 0.801257891 | 0.001160424 | 0.004653532 |
| GARNL3 | 3.06112962 | 3.57458642 | 0.000218338 | 0.001108725 |
| LOC106559768 | 3.058211877 | 1.612357858 | 0.01141262 | 0.032202279 |
| PABPC5 | 3.057406477 | 2.056023291 | 0.000784802 | 0.00331948 |
| TMEM150C | 3.056583212 | 3.16775701 | 0.000497062 | 0.002247615 |
| PRICKLE1 | 3.056136702 | 5.897522128 | 3.74E-06 | 3.23E-05 |
| CCDC170 | 3.054886032 | 1.701424771 | 0.000282542 | 0.001380729 |
| MSR1 | 3.053874528 | 4.154164988 | 4.36E-05 | 0.0002783 |
| SOX10 | 3.052302774 | 6.157308311 | 0.000799321 | 0.003372071 |
| MDH1B | 3.049544032 | 0.319622159 | 0.001495133 | 0.005744591 |
| TUBB6 | 3.04936511 | 5.876652508 | 4.66E-07 | 5.28E-06 |
| MMP28 | 3.04886508 | 4.67592978 | 0.000241989 | 0.001207885 |
| LOC475605 | 3.048525381 | 2.36861554 | 0.006877578 | 0.021013095 |
| FLYWCH1 | 3.043818052 | 2.190130754 | 0.000149036 | 0.000796952 |

| | | | | |
|---|---|---|---|---|
| TTC28 | 3.042859975 | 7.042504047 | 3.73E-08 | 5.63E-07 |
| LOC119876100 | 3.041760889 | 0.379399857 | 0.000819493 | 0.003446036 |
| LINGO2 | 3.041708616 | 0.740580273 | 0.01165713 | 0.032755735 |
| EPN3 | 3.039783991 | -0.139805805 | 0.007133464 | 0.021677447 |
| MAP3K21 | 3.039486153 | 3.358904626 | 2.12E-06 | 1.98E-05 |
| DENND6B | 3.037822645 | 5.602877076 | 1.06E-05 | 8.01E-05 |
| PLOD1 | 3.035288355 | 6.899284252 | 7.98E-08 | 1.10E-06 |
| LOC119877580 | 3.035033374 | 0.152559874 | 0.003075965 | 0.010632082 |
| LOC119876404 | 3.033100584 | 0.248428184 | 0.005599189 | 0.017635697 |
| ECSCR | 3.031965203 | 4.316647185 | 0.000381679 | 0.001790535 |
| ADGRA3 | 3.031861078 | 4.700832233 | 2.32E-07 | 2.85E-06 |
| LOC119870568 | 3.030506467 | 4.415855932 | 0.002686152 | 0.009470794 |
| SLC25A10 | 3.030188844 | 3.725294272 | 0.000360148 | 0.001703137 |
| FBXL13 | 3.028823749 | 0.667549971 | 0.007212412 | 0.02188667 |
| LOC119872526 | 3.027922908 | 3.109242088 | 0.004031539 | 0.013372444 |
| CORO2B | 3.026694851 | 4.549960617 | 9.99E-06 | 7.61E-05 |
| LOC119869495 | 3.026221235 | 2.004602063 | 0.009402612 | 0.027393844 |
| LOC102157008 | 3.026060511 | -0.666631941 | 0.010827561 | 0.03081707 |
| LOC111095904 | 3.021845775 | 1.114142795 | 0.00419782 | 0.013811954 |
| RBFOX3 | 3.021088925 | 1.107497901 | 0.001690781 | 0.006378973 |
| C5H11orf52 | 3.018523629 | 1.115295462 | 0.001316141 | 0.005171271 |
| SLC34A3 | 3.018324286 | 0.932654073 | 0.009118586 | 0.026695545 |
| LOC119873855 | 3.01561521 | 0.110564124 | 0.006457847 | 0.019911923 |
| LRRC9 | 3.014676038 | 2.24292852 | 2.67E-06 | 2.43E-05 |
| LOC119873489 | 3.014130947 | 2.249180955 | 0.010937487 | 0.031067909 |
| INAFM1 | 3.014125034 | 3.73509476 | 2.60E-06 | 2.37E-05 |
| LOC100688933 | 3.013814501 | 2.402533711 | 6.73E-05 | 0.000405764 |
| PRELID3A | 3.013125943 | 4.477578223 | 1.12E-05 | 8.45E-05 |
| LOC102153215 | 3.01208874 | -0.79215334 | 0.006414124 | 0.019810931 |
| KIF3C | 3.011088583 | 4.895459986 | 1.16E-07 | 1.54E-06 |
| THAP10 | 3.010983344 | 1.596182698 | 0.000918076 | 0.003813615 |
| MYO1C | 3.010257161 | 7.680062068 | 4.39E-09 | 8.19E-08 |
| LOC607080 | 3.009207342 | 4.753688023 | 0.002898834 | 0.010115133 |
| SPTBN2 | 3.009138774 | 3.18192034 | 1.41E-07 | 1.83E-06 |
| LRRN1 | 3.007946972 | 3.22254638 | 0.000839202 | 0.003522764 |
| SOX13 | 3.00606042 | 5.927807958 | 0.000440029 | 0.002022642 |
| PLXNB2 | 3.004099795 | 8.515040124 | 2.04E-06 | 1.91E-05 |
| IFT22 | 3.003278467 | 2.062420287 | 0.000323501 | 0.00155322 |
| TGFBR3L | 3.003245037 | 3.82442925 | 0.003940787 | 0.013121628 |

| | | | | |
|---|---|---|---|---|
| LRRC27 | 3.003209953 | 1.379685914 | 0.009654215 | 0.027995157 |
| RRBP1 | 2.999859013 | 7.896497839 | 1.87E-05 | 0.0001329 |
| LOC119873500 | 2.999449179 | 1.516686538 | 0.012867248 | 0.035634024 |
| LOC102156186 | 2.996285188 | 3.055700674 | 0.000371232 | 0.00174838 |
| LOC119866503 | 2.996283322 | 2.917021857 | 0.012309716 | 0.034304805 |
| LOC119868548 | 2.993970082 | 2.704249415 | 0.001209734 | 0.004819662 |
| THEM6 | 2.993285705 | 2.810079743 | 0.000525555 | 0.002358249 |
| PANX2 | 2.992336263 | 2.874837111 | 0.001545105 | 0.005917487 |
| MYPOP | 2.991920206 | 3.810507302 | 0.000151761 | 0.00080973 |
| SEPTIN8 | 2.991159037 | 6.835276174 | 7.52E-09 | 1.33E-07 |
| NEURL3 | 2.982418207 | -0.40993995 | 0.008061834 | 0.024053857 |
| CBX2 | 2.981907254 | 2.164116314 | 0.010717988 | 0.030560577 |
| LOC111093862 | 2.981426821 | 0.017564037 | 0.014491995 | 0.039359023 |
| LOC100684414 | 2.981236056 | 2.640951521 | 1.93E-06 | 1.83E-05 |
| GAS2L2 | 2.980378065 | 2.53923359 | 0.001281131 | 0.005058217 |
| LOC119876378 | 2.97968236 | 1.61618453 | 0.001360928 | 0.005318318 |
| LOC119877507 | 2.97778601 | 1.606749259 | 0.00650476 | 0.020025237 |
| CBR3 | 2.976774162 | 3.417477926 | 1.41E-05 | 0.000103681 |
| ABCA5 | 2.975786354 | 5.018720751 | 1.12E-07 | 1.49E-06 |
| TMEM8B | 2.975685482 | 3.931900314 | 6.05E-06 | 4.90E-05 |
| NUAK1 | 2.975577989 | 5.731193423 | 6.20E-07 | 6.75E-06 |
| ECHDC2 | 2.97540374 | 2.974558621 | 0.001162589 | 0.004660893 |
| LOC119874173 | 2.972455817 | 0.751119925 | 0.005310825 | 0.016840017 |
| LOC119866956 | 2.972004273 | 4.775267296 | 0.013404159 | 0.036833821 |
| PLXNA1 | 2.97019454 | 7.916173186 | 1.34E-05 | 9.87E-05 |
| VAT1 | 2.970168846 | 7.6347639 | 4.21E-08 | 6.26E-07 |
| LOC111093275 | 2.968715595 | -0.237934182 | 0.001622477 | 0.006159421 |
| LOC111090541 | 2.968707814 | 4.523173356 | 1.80E-05 | 0.00012826 |
| PRDX2 | 2.966877279 | 4.979978001 | 2.90E-08 | 4.49E-07 |
| KIAA1522 | 2.966010913 | 5.436152461 | 1.61E-07 | 2.05E-06 |
| PDGFRL | 2.963944508 | 4.62345383 | 0.001854712 | 0.006905544 |
| SPINK7 | 2.961153201 | 1.254200164 | 0.000278143 | 0.001361071 |
| LOC119873233 | 2.959819509 | 0.146182425 | 0.002894818 | 0.010107336 |
| CSDC2 | 2.955360271 | 2.737790129 | 0.001334655 | 0.005228846 |
| BAHCC1 | 2.953063544 | 6.635053213 | 2.76E-06 | 2.49E-05 |
| ITGAE | 2.951995392 | 5.323206255 | 0.000876675 | 0.003659509 |
| LOC119873514 | 2.951747434 | 6.724628496 | 0.014273749 | 0.038868572 |
| CBX8 | 2.951511115 | 3.053791203 | 0.001557518 | 0.005956605 |
| RETREG1 | 2.95071792 | 4.399884324 | 2.13E-06 | 1.99E-05 |

| | | | | |
|---|---|---|---|---|
| TONSL | 2.950715143 | 4.65530964 | 0.000168654 | 0.000887411 |
| JCAD | 2.949051179 | 6.105818246 | 0.000119302 | 0.000665369 |
| LOC111093268 | 2.948934684 | 7.386841985 | 0.000508897 | 0.002292523 |
| BDKRB2 | 2.946752405 | 2.325099992 | 0.00295214 | 0.010278191 |
| PODNL1 | 2.945519782 | 1.789285503 | 0.000795138 | 0.003356645 |
| CCDC102A | 2.945364834 | 5.55284331 | 2.53E-05 | 0.00017373 |
| LOC119873650 | 2.945025957 | -0.971430968 | 0.002185107 | 0.007945945 |
| LOC111098802 | 2.944644604 | 0.532498321 | 0.011636751 | 0.032706951 |
| LOC111092895 | 2.94417234 | 2.894523197 | 0.001707294 | 0.006427854 |
| KCNN3 | 2.94413397 | 4.605285195 | 1.55E-05 | 0.00011242 |
| WDPCP | 2.94127101 | 2.664364391 | 5.11E-06 | 4.23E-05 |
| RPGRIP1L | 2.940614804 | 3.519847517 | 0.000655681 | 0.00285041 |
| LOC102155268 | 2.939848625 | -0.713055442 | 0.013190029 | 0.036383797 |
| LOC119869745 | 2.939623613 | 3.322852688 | 0.003427434 | 0.011644894 |
| C20H3orf18 | 2.939145078 | 0.716877138 | 0.011271772 | 0.031846318 |
| LOC478825 | 2.937107245 | 0.806598753 | 0.003810686 | 0.012753088 |
| GRM4 | 2.936189332 | 3.745347439 | 0.009668045 | 0.028016517 |
| LOC111098296 | 2.932255669 | -0.026881762 | 0.01537085 | 0.041285466 |
| LOC111097025 | 2.931942942 | -0.278416333 | 0.017238834 | 0.045182883 |
| LOC111098503 | 2.931674407 | -0.102653706 | 0.001705156 | 0.006421519 |
| LOC119867570 | 2.931420652 | 5.132251144 | 0.012505553 | 0.03477455 |
| PTGR2 | 2.931016732 | 3.576184227 | 1.62E-05 | 0.000116816 |
| C24H20orf96 | 2.930980255 | 2.161763253 | 0.000889743 | 0.003708368 |
| LOC119877279 | 2.929969483 | 1.477319455 | 0.014380999 | 0.039121301 |
| PLEKHM2 | 2.929760196 | 8.103226385 | 8.67E-05 | 0.000505813 |
| C27H12orf71 | 2.929399568 | 1.721457768 | 0.0151859 | 0.040859591 |
| SPOCD1 | 2.928516545 | 2.004432838 | 0.001842183 | 0.006864793 |
| TRIM2 | 2.927693907 | 6.698360551 | 1.01E-07 | 1.36E-06 |
| LOC111090945 | 2.925413523 | 1.623530682 | 0.000953169 | 0.003930428 |
| RNF227 | 2.925339621 | 1.507580942 | 0.000676692 | 0.002927335 |
| MMP14 | 2.924639422 | 8.393275268 | 1.89E-06 | 1.80E-05 |
| PIK3R2 | 2.924148801 | 6.393061773 | 6.92E-06 | 5.51E-05 |
| GPSM1 | 2.923894322 | 5.697701466 | 3.91E-07 | 4.52E-06 |
| ARHGEF26 | 2.923845945 | 5.827880879 | 0.000323018 | 0.001551836 |
| KCND1 | 2.921670734 | 4.348396011 | 5.20E-05 | 0.000325072 |
| ODAD1 | 2.920284953 | 2.732365623 | 0.003973144 | 0.013223271 |
| SHANK2 | 2.919896404 | 1.814003638 | 0.010016204 | 0.028897921 |
| FAM133B | 2.919488226 | 0.237140972 | 0.001219521 | 0.004850557 |
| RIC3 | 2.918108641 | 4.805743141 | 0.011814708 | 0.03313408 |

| | | | | |
|---|---|---|---|---|
| TCEAL1 | 2.918024581 | 3.436999962 | 8.34E-05 | 0.000489564 |
| ARHGEF9 | 2.917372533 | 3.052634798 | 3.32E-05 | 0.000219265 |
| FXYD6 | 2.916641741 | 6.217196579 | 0.000454855 | 0.002078447 |
| ZFPM1 | 2.912978733 | 4.884146918 | 0.00018449 | 0.000957712 |
| ZNF572 | 2.912641281 | 1.261187071 | 0.001246254 | 0.004935149 |
| LOC119872600 | 2.91164723 | 1.793941056 | 0.00057094 | 0.002528915 |
| CROCC | 2.908547812 | 5.77201949 | 0.000677487 | 0.002929606 |
| HEY1 | 2.908221084 | 2.896925007 | 0.006825334 | 0.020884579 |
| CLCN4 | 2.907089563 | 3.895445506 | 1.43E-06 | 1.40E-05 |
| JUP | 2.906118793 | 7.089854016 | 2.45E-06 | 2.25E-05 |
| NLGN3 | 2.904271013 | 3.509144754 | 0.000244307 | 0.001218329 |
| ARMC9 | 2.903958045 | 3.848431054 | 1.92E-06 | 1.82E-05 |
| LOC111097476 | 2.903876088 | 2.21504055 | 0.002802521 | 0.00983469 |
| SPATA45 | 2.901911439 | -0.929770102 | 0.013964745 | 0.038197934 |
| LOC119871797 | 2.899173358 | 2.298353492 | 0.019290285 | 0.04953535 |
| MAMDC2 | 2.89820206 | 3.689622381 | 0.001871115 | 0.006948711 |
| LOC119873506 | 2.897537275 | 1.407928439 | 0.004415845 | 0.014415515 |
| LOC119866935 | 2.896988507 | 0.117098256 | 0.013819379 | 0.037854828 |
| MGMT | 2.896082078 | 2.62045194 | 0.001060811 | 0.004306228 |
| MCF2L | 2.893369596 | 4.927634346 | 3.39E-06 | 2.98E-05 |
| LOC102156006 | 2.891551257 | 3.84695668 | 4.74E-06 | 3.97E-05 |
| NPHP4 | 2.891331063 | 4.575791539 | 1.33E-05 | 9.83E-05 |
| SCN1B | 2.890334605 | 3.397442383 | 0.007114897 | 0.021627092 |
| LOC111090394 | 2.888767697 | 3.076107843 | 1.08E-05 | 8.17E-05 |
| LOC102156860 | 2.887628206 | 1.179194194 | 1.00E-04 | 0.000572288 |
| LOC119873361 | 2.887017675 | 0.612231445 | 0.006173263 | 0.019159835 |
| MT2A | 2.885819417 | 5.63161316 | 0.000125216 | 0.000691768 |
| ZNF575 | 2.88568717 | 3.662184575 | 0.000111152 | 0.000626854 |
| LOC119866923 | 2.88553549 | 1.770571415 | 0.003978129 | 0.013233053 |
| FAM221B | 2.885081847 | 1.339085199 | 0.006105648 | 0.018977155 |
| GATA2 | 2.884978761 | 3.150345094 | 0.001553738 | 0.005945297 |
| TCF4 | 2.882191859 | 7.676465442 | 8.63E-08 | 1.19E-06 |
| DCHS1 | 2.880731209 | 6.07806597 | 3.82E-07 | 4.42E-06 |
| CFAP100 | 2.880632902 | 3.825342964 | 0.009855395 | 0.02851368 |
| SPRED2 | 2.88003303 | 5.416730209 | 7.36E-07 | 7.83E-06 |
| RTN4RL1 | 2.878760139 | 1.636425108 | 0.011857055 | 0.033222894 |
| CTH | 2.876909271 | 2.438851388 | 0.001916553 | 0.007097994 |
| PLAG1 | 2.876086727 | 2.171755078 | 0.001186214 | 0.004742449 |
| SLCO5A1 | 2.871782323 | 2.367834908 | 0.017560765 | 0.04584923 |

| | | | | |
|---|---|---|---|---|
| ARL13B | 2.871056451 | 3.618706101 | 3.68E-06 | 3.19E-05 |
| HYAL1 | 2.870404085 | 4.738634349 | 1.52E-06 | 1.49E-05 |
| LOC102152846 | 2.869268518 | 2.370187294 | 0.000161651 | 0.000856174 |
| PRRT4 | 2.86923274 | 1.59060492 | 0.001870624 | 0.006948537 |
| SCTR | 2.868716658 | 1.792671129 | 0.000510096 | 0.00229697 |
| LOC106559908 | 2.867246561 | 2.345099306 | 0.01817484 | 0.047151461 |
| LOC119879327 | 2.866692525 | 9.374809515 | 0.010411051 | 0.029809021 |
| NTHL1 | 2.864863635 | 3.345264898 | 8.30E-05 | 0.000487237 |
| C16H8orf48 | 2.862053957 | 1.507461799 | 0.005433207 | 0.017190351 |
| LOC119872418 | 2.862052485 | 3.067903807 | 0.00192469 | 0.007125694 |
| EPOP | 2.861548292 | 2.820248912 | 0.000177759 | 0.000927659 |
| LOC119864296 | 2.861353857 | 1.018469033 | 0.001020409 | 0.004164062 |
| LOC119867337 | 2.859914724 | 2.784461972 | 0.000752826 | 0.003205501 |
| NRP1 | 2.85833735 | 7.117766861 | 3.06E-07 | 3.64E-06 |
| LOC119872731 | 2.857343426 | 3.775151434 | 0.001191926 | 0.004760896 |
| LOC119872052 | 2.857238127 | 5.013358458 | 0.010753433 | 0.030641489 |
| IFI27L2 | 2.85574564 | 3.732379811 | 6.78E-05 | 0.000408019 |
| ATP7B | 2.855715185 | 3.37983748 | 0.000148809 | 0.000796402 |
| LOC102153635 | 2.855479521 | 0.635022687 | 0.003216292 | 0.011036527 |
| WDR35 | 2.855086433 | 4.041580774 | 1.41E-06 | 1.39E-05 |
| LOC119865927 | 2.854741781 | 0.827590392 | 0.012972043 | 0.035878392 |
| TUFT1 | 2.854512677 | 3.732246838 | 0.000211437 | 0.001079499 |
| MAP7D1 | 2.853433463 | 9.202872485 | 0.000519947 | 0.002335506 |
| RCN3 | 2.850534557 | 6.360267022 | 3.14E-06 | 2.80E-05 |
| LOC119878577 | 2.849682479 | 0.166111969 | 0.00813942 | 0.024241535 |
| SH3BP5 | 2.849134332 | 6.891412906 | 1.60E-07 | 2.04E-06 |
| LOC102154481 | 2.849001493 | 0.887128292 | 0.017424514 | 0.045581351 |
| LOC102151519 | 2.847696633 | 2.446899031 | 9.48E-06 | 7.27E-05 |
| ANKRD29 | 2.846842991 | 4.00822256 | 8.92E-06 | 6.90E-05 |
| RHOD | 2.846783859 | 3.154949893 | 1.35E-05 | 9.97E-05 |
| TNS1 | 2.846339137 | 10.57644235 | 6.53E-05 | 0.000395854 |
| CYP27C1 | 2.846125358 | 1.907306907 | 0.004429877 | 0.014449537 |
| SYNPO | 2.846004576 | 7.301225462 | 5.03E-07 | 5.62E-06 |
| LOC119873600 | 2.845923622 | 3.371592884 | 0.001024118 | 0.00417291 |
| F2RL2 | 2.843811418 | 5.627840006 | 0.001627357 | 0.006174667 |
| OTUD7A | 2.842663461 | 1.734202777 | 0.002551939 | 0.009055037 |
| NR1D1 | 2.841391787 | 6.514530669 | 0.001412523 | 0.005474537 |
| PGF | 2.839515809 | 4.597136507 | 1.66E-06 | 1.60E-05 |
| EMID1 | 2.838946448 | 3.385673484 | 0.00096721 | 0.003977943 |

| | | | | |
|---|---|---|---|---|
| HDGFL2 | 2.838300353 | 5.881597155 | 7.55E-05 | 0.000449598 |
| GPRC5A | 2.836640899 | 3.099639079 | 0.004880924 | 0.015674066 |
| LOC119876689 | 2.83429004 | 1.760028306 | 0.018195156 | 0.047182493 |
| LOC119876016 | 2.834168976 | 2.412383975 | 0.001006238 | 0.00411864 |
| LOC119870120 | 2.833367413 | 2.61206838 | 0.000982678 | 0.004032869 |
| ZSCAN18 | 2.831362036 | 5.214826333 | 8.81E-10 | 1.97E-08 |
| CHRNB2 | 2.829862865 | 2.877388515 | 0.006978362 | 0.02127772 |
| AJM1 | 2.824586568 | 3.185429297 | 0.001464494 | 0.005647684 |
| FAM163B | 2.824183735 | -0.253604493 | 0.013312701 | 0.036652207 |
| FHOD3 | 2.824163702 | 3.591074334 | 0.001272645 | 0.005028631 |
| LOC106559435 | 2.822102026 | 2.370768179 | 6.65E-05 | 0.000401798 |
| SPOCK1 | 2.820446228 | 3.034966783 | 0.007135006 | 0.021679095 |
| LOC106558577 | 2.820086338 | -0.832481907 | 0.007405857 | 0.022398428 |
| PLCXD1 | 2.819622087 | 1.232341667 | 0.003553147 | 0.012004263 |
| LOC111095987 | 2.818855472 | 2.444393413 | 0.00560876 | 0.01766071 |
| HECTD2 | 2.817839405 | 4.701555604 | 3.93E-05 | 0.000254421 |
| FMNL3 | 2.816902064 | 7.101642555 | 1.15E-06 | 1.16E-05 |
| MROH6 | 2.815935077 | 3.873089334 | 0.005420879 | 0.017159783 |
| LOC111094738 | 2.813290008 | 5.29054642 | 8.82E-05 | 0.000512991 |
| LOC119868218 | 2.812448043 | 2.676947889 | 5.30E-05 | 0.000330689 |
| AMIGO1 | 2.811668459 | 4.07412082 | 1.03E-05 | 7.81E-05 |
| ZBTB8A | 2.808610165 | 1.695842354 | 0.00227984 | 0.008232419 |
| LOC119864748 | 2.808604689 | 2.152603743 | 0.000315201 | 0.001520442 |
| LOC119878760 | 2.808314718 | -0.343286291 | 0.005367409 | 0.017009485 |
| LOC119865899 | 2.805153362 | 2.273058047 | 0.000136559 | 0.000743068 |
| CYP39A1 | 2.802402189 | 3.51396805 | 0.00015446 | 0.000822714 |
| CD300LF | 2.79759606 | 2.013679738 | 0.014537334 | 0.039457439 |
| CNKSR1 | 2.797475687 | 4.106375057 | 0.001613339 | 0.006130067 |
| SHB | 2.795716763 | 3.307167612 | 9.23E-05 | 0.000533756 |
| MAF | 2.795493536 | 7.056808527 | 0.001215539 | 0.004839157 |
| TFAP4 | 2.793017211 | 4.529213375 | 2.25E-05 | 0.000156219 |
| NSUN7 | 2.791905437 | 1.457815519 | 0.001986542 | 0.007328397 |
| RAB30 | 2.789777071 | 5.651457054 | 3.83E-05 | 0.000249086 |
| BEAN1 | 2.789557602 | 1.418053307 | 0.002962154 | 0.010304699 |
| LOC119864688 | 2.786824503 | 0.86187482 | 0.003357145 | 0.011445555 |
| CLMN | 2.785432549 | 4.160366037 | 2.02E-05 | 0.00014203 |
| ZNF112 | 2.784514749 | 3.082288447 | 7.76E-07 | 8.19E-06 |
| SLC26A11 | 2.783126305 | 3.753223777 | 1.37E-05 | 0.000100526 |
| PRX | 2.782215214 | 7.781829343 | 0.001049269 | 0.004264967 |

| | | | | |
|---|---|---|---|---|
| NACC2 | 2.781788776 | 5.784602069 | 3.59E-06 | 3.12E-05 |
| CDCA8 | 2.781059487 | 2.349926149 | 0.000588283 | 0.002592979 |
| FAM160A1 | 2.78066479 | 5.458704776 | 2.91E-05 | 0.00019589 |
| PAQR7 | 2.78052877 | 3.87588449 | 0.000180634 | 0.000940397 |
| PXYLP1 | 2.78014622 | 5.133230828 | 7.79E-07 | 8.22E-06 |
| BMP6 | 2.77926812 | 4.948099294 | 0.000227252 | 0.001145402 |
| TEAD2 | 2.778544074 | 5.008053639 | 6.33E-06 | 5.10E-05 |
| C15H1orf122 | 2.777208749 | 3.334943332 | 2.03E-05 | 0.000142421 |
| TWSG1 | 2.777136733 | 6.69348853 | 2.60E-05 | 0.00017786 |
| LOC119864513 | 2.776537652 | 3.065038263 | 0.005083695 | 0.01623621 |
| PLEKHA7 | 2.776066416 | 5.066194947 | 0.01179196 | 0.033083127 |
| LOC100684448 | 2.77573967 | 0.873601832 | 0.01888799 | 0.048634835 |
| LOC111094029 | 2.774542343 | 0.182507111 | 0.009211773 | 0.026924713 |
| PLCB4 | 2.771347875 | 5.539394 | 0.000941551 | 0.003889952 |
| C17H2orf16 | 2.771228774 | 1.271843278 | 0.000636897 | 0.002779594 |
| THY1 | 2.770529933 | 8.603363168 | 2.15E-05 | 0.000150135 |
| TSPAN9 | 2.769495696 | 6.474830597 | 0.001017043 | 0.004154236 |
| LOC100688940 | 2.768897399 | 2.655638024 | 0.000948131 | 0.003914063 |
| RTEL1 | 2.767924751 | 6.858164868 | 0.000367125 | 0.001730845 |
| ARHGEF12 | 2.767652989 | 7.973857249 | 3.28E-06 | 2.89E-05 |
| LOC478001 | 2.764898498 | 2.318454886 | 0.001390112 | 0.005409931 |
| MAGEF1 | 2.764081819 | 3.320217338 | 0.000171951 | 0.000902127 |
| SERF1A | 2.763714532 | 1.811094023 | 0.001518649 | 0.005827492 |
| RENBP | 2.763398386 | 5.368787863 | 7.48E-05 | 0.000446203 |
| LOC111098307 | 2.762924832 | -0.484943272 | 0.018842698 | 0.048538313 |
| LRIG1 | 2.762088828 | 6.171292893 | 8.33E-05 | 0.000489055 |
| CCDC141 | 2.760778756 | 4.280164559 | 0.000114039 | 0.000640302 |
| ASAP3 | 2.760723314 | 6.248537805 | 0.000109023 | 0.000616773 |
| LOC102155565 | 2.759877363 | -0.706220268 | 0.01268628 | 0.03519582 |
| NRBP2 | 2.757159201 | 7.006775547 | 9.28E-06 | 7.13E-05 |
| KLF16 | 2.755982218 | 5.180921089 | 0.001136191 | 0.004570281 |
| MNS1 | 2.755320534 | 2.555561857 | 0.000807071 | 0.003400397 |
| RAB13 | 2.755262963 | 4.31183318 | 2.87E-06 | 2.58E-05 |
| IL17RC | 2.754976548 | 4.152123511 | 7.47E-07 | 7.93E-06 |
| ADGRV1 | 2.75332932 | 2.819158339 | 0.003363001 | 0.011460112 |
| CCDC188 | 2.752135984 | 4.170143979 | 0.00019349 | 0.000997267 |
| LOC119865180 | 2.751071719 | 0.325169469 | 0.000934039 | 0.00386768 |
| NRARP | 2.750258409 | 2.513403208 | 0.000432196 | 0.001992128 |
| LOC102152908 | 2.748201118 | 1.771344378 | 0.003603334 | 0.012156765 |

| | | | | |
|---|---|---|---|---|
| AFAP1L1 | 2.746936498 | 4.295655875 | 4.74E-05 | 0.000299639 |
| KIF5C | 2.745562152 | 5.349530475 | 0.003030041 | 0.010500493 |
| GDPD2 | 2.745091367 | 1.787898009 | 0.001405429 | 0.005451916 |
| SMAD6 | 2.743249128 | 4.228680286 | 5.27E-06 | 4.34E-05 |
| SCAI | 2.741552581 | 6.23545111 | 0.000147918 | 0.000793126 |
| TTC30A | 2.740468295 | 0.945689754 | 0.008336402 | 0.024722884 |
| PAQR4 | 2.739021519 | 4.333977399 | 0.000155392 | 0.000826861 |
| LOC111090550 | 2.737973335 | -0.078603029 | 0.016690146 | 0.04403902 |
| FUOM | 2.737520914 | 1.831924223 | 0.005732865 | 0.018012234 |
| FCGR2B | 2.737199397 | 4.142063192 | 0.002871066 | 0.010041087 |
| GGT5 | 2.73421211 | 4.141458187 | 0.001476403 | 0.005684504 |
| LOC102152002 | 2.733641555 | -0.368625231 | 0.00941176 | 0.027409444 |
| LARGE1 | 2.732569325 | 5.069602376 | 9.12E-06 | 7.03E-05 |
| LOC119877678 | 2.730227065 | -0.858518672 | 0.012939833 | 0.035807584 |
| PLEKHG4 | 2.729954485 | 3.629397098 | 9.16E-05 | 0.00053002 |
| LOC119866410 | 2.728737182 | 1.21894227 | 0.010614676 | 0.030333845 |
| KIT | 2.727719982 | 3.427005864 | 0.000211576 | 0.001079955 |
| PLEKHA6 | 2.726399919 | 5.79387163 | 0.001073471 | 0.00435273 |
| CCDC92 | 2.726366099 | 4.485378526 | 6.25E-05 | 0.000380874 |
| FASN | 2.726260624 | 8.496594116 | 6.24E-05 | 0.000380636 |
| ABTB2 | 2.723834496 | 3.45046039 | 0.000150679 | 0.000804547 |
| LOC111095223 | 2.717459194 | -0.139208435 | 0.016838491 | 0.044352492 |
| LOC102154725 | 2.716147451 | 1.347131297 | 0.006635967 | 0.020371292 |
| TTYH2 | 2.715348961 | 5.024393002 | 0.000176097 | 0.000920095 |
| LOC102151760 | 2.714091954 | 1.279550772 | 0.004205584 | 0.013829105 |
| RIIAD1 | 2.713632385 | 2.133979877 | 0.00650413 | 0.020025237 |
| GTSE1 | 2.712666306 | 2.23249112 | 0.001782898 | 0.006672551 |
| CELSR1 | 2.712408764 | 4.416198554 | 4.71E-05 | 0.000298355 |
| GPT2 | 2.711436736 | 4.071068302 | 0.00012237 | 0.000678986 |
| LOC607011 | 2.709417359 | 4.498331807 | 1.75E-05 | 0.000125285 |
| LOC111095615 | 2.708854131 | 1.252022403 | 0.002970981 | 0.010327196 |
| AK1 | 2.705613972 | 3.713530057 | 0.001211438 | 0.004824605 |
| AK4 | 2.70391218 | 5.668197809 | 0.00011113 | 0.000626854 |
| MIIP | 2.703507678 | 3.338907097 | 2.82E-05 | 0.000190368 |
| ACYP1 | 2.702949515 | 0.911771703 | 0.005079383 | 0.016224828 |
| GIPC3 | 2.702606455 | 1.377127815 | 0.009811937 | 0.028403113 |
| LOC106558496 | 2.70058528 | 2.342647593 | 0.000399594 | 0.001862851 |
| MICALL2 | 2.69786698 | 5.555612383 | 1.18E-05 | 8.84E-05 |
| LOC119877092 | 2.695811996 | 4.454512465 | 2.58E-05 | 0.00017666 |

| | | | | |
|---|---|---|---|---|
| SLC25A34 | 2.695738577 | 3.630010716 | 1.99E-05 | 0.000140254 |
| PRRT2 | 2.694921611 | 5.284022241 | 0.000226234 | 0.001141332 |
| CLTB | 2.694101944 | 5.544651903 | 1.70E-06 | 1.64E-05 |
| LOC119879325 | 2.693506911 | 0.02724243 | 0.004754065 | 0.015323445 |
| ITSN1 | 2.693467109 | 7.408889569 | 0.000179412 | 0.000935611 |
| LOC119872509 | 2.693332272 | 1.828663863 | 0.000699989 | 0.003012482 |
| MERTK | 2.691033741 | 4.842195202 | 0.000493368 | 0.002232311 |
| ADGRF1 | 2.690916095 | 1.77076126 | 0.017336551 | 0.045389597 |
| LOC102151797 | 2.690297009 | 0.373926145 | 0.015624605 | 0.041785792 |
| LOC102157224 | 2.689563274 | 0.70886176 | 0.001252768 | 0.004959132 |
| LOC119866943 | 2.689554584 | 0.576874136 | 0.008841719 | 0.025987447 |
| RNF152 | 2.687424038 | 5.10227573 | 0.001496277 | 0.005747751 |
| LOC111090977 | 2.687411876 | 3.551972921 | 0.000332072 | 0.001586639 |
| LOC119873713 | 2.687090667 | 0.066806413 | 0.018666743 | 0.04817389 |
| P4HA3 | 2.685894252 | 4.81553442 | 8.76E-05 | 0.000510097 |
| KIRREL2 | 2.685060378 | -0.489958514 | 0.014887964 | 0.040227759 |
| LOC102157295 | 2.684694172 | 0.974326215 | 0.001627146 | 0.006174667 |
| LOC119871842 | 2.684221041 | 1.98708882 | 0.000129063 | 0.000709553 |
| LOC119871828 | 2.682042549 | 0.015865824 | 0.006961451 | 0.02123811 |
| FNDC10 | 2.680309517 | 3.442509029 | 5.15E-06 | 4.26E-05 |
| TSNARE1 | 2.680127718 | 4.253351446 | 0.000157407 | 0.00083656 |
| PWWP2B | 2.678786823 | 4.765838704 | 0.000333973 | 0.001594316 |
| GNB3 | 2.678401648 | 3.858273994 | 0.000188526 | 0.000974929 |
| LOC119877086 | 2.678007936 | -0.096837916 | 0.007759797 | 0.023290136 |
| ANKUB1 | 2.67638497 | -0.834377509 | 0.014819363 | 0.04007737 |
| LOC119876062 | 2.676278027 | -0.085239736 | 0.002924331 | 0.010191191 |
| C1QL3 | 2.674771992 | 1.774231295 | 0.003501138 | 0.011871116 |
| BCAR3 | 2.673332376 | 4.344471962 | 6.66E-05 | 0.000401925 |
| PLEC | 2.673244888 | 11.68065251 | 0.000718826 | 0.003079487 |
| GNAI1 | 2.669207507 | 5.118407923 | 7.09E-05 | 0.000424602 |
| TNFRSF13C | 2.668868264 | 2.727364777 | 0.001287053 | 0.005075383 |
| FBXO41 | 2.668171631 | 2.340819033 | 0.004216802 | 0.013857582 |
| IDUA | 2.668116649 | 5.266416064 | 5.09E-05 | 0.000318757 |
| RGS22 | 2.667060199 | 4.124493631 | 6.15E-05 | 0.000375986 |
| PCSK6 | 2.665103986 | 3.737755091 | 0.006018314 | 0.018762216 |
| IFITM5 | 2.663437605 | 1.980646313 | 0.008679556 | 0.02558999 |
| BBS12 | 2.662422099 | 3.350490855 | 0.000139162 | 0.000754394 |
| DAG1 | 2.662264763 | 7.642220929 | 3.25E-06 | 2.87E-05 |
| SPEM3 | 2.662038801 | -0.267372567 | 0.016760695 | 0.044181962 |

| | | | | |
|---|---|---|---|---|
| ZFHX2 | 2.657508777 | 5.966790284 | 0.000178238 | 0.000929711 |
| HAGHL | 2.656832624 | 3.228283486 | 0.00029273 | 0.001426035 |
| SYNGR1 | 2.656301614 | 3.886221857 | 0.000434673 | 0.002002266 |
| ARHGEF37 | 2.653775099 | 4.358965372 | 0.000615208 | 0.002697432 |
| ARL6 | 2.653557303 | 3.80171028 | 2.33E-05 | 0.000161101 |
| TCTE1 | 2.653172821 | 2.309766831 | 0.000490661 | 0.002220987 |
| MEGF8 | 2.652589262 | 6.538237403 | 4.02E-08 | 6.01E-07 |
| COL9A2 | 2.649877884 | 5.751702829 | 0.000333026 | 0.001590496 |
| RPP25 | 2.648812884 | 1.594537875 | 0.006613431 | 0.020310739 |
| MST1 | 2.647908062 | 2.297761073 | 0.006552081 | 0.02014802 |
| DEPTOR | 2.645843313 | 6.948673392 | 0.003033859 | 0.010512044 |
| ETV4 | 2.645498302 | 3.894504409 | 0.004690646 | 0.015152835 |
| CTDSPL | 2.643460355 | 5.606657164 | 3.14E-06 | 2.79E-05 |
| HOOK1 | 2.643196181 | 4.030624811 | 0.000266456 | 0.001311881 |
| PDZD7 | 2.643185102 | 2.947418025 | 0.000212301 | 0.001082635 |
| DUSP23 | 2.642828321 | 4.270500931 | 6.53E-05 | 0.000395854 |
| PLXDC2 | 2.640278986 | 9.252413907 | 0.000556722 | 0.002473519 |
| ENHO | 2.639577758 | 1.927908421 | 0.006536541 | 0.020113488 |
| DLGAP4 | 2.639258701 | 7.222641055 | 2.47E-05 | 0.000170202 |
| LOC102151792 | 2.637132054 | -0.771370881 | 0.008724122 | 0.025700817 |
| PCYT2 | 2.636989074 | 4.779914311 | 0.000101328 | 0.000578981 |
| LOC102155379 | 2.63674471 | 1.992294727 | 4.99E-05 | 0.00031368 |
| SMIM1 | 2.636291745 | 1.088059569 | 0.010842213 | 0.030853898 |
| LOC119876928 | 2.636239418 | 0.092522935 | 0.001847315 | 0.006879184 |
| KCNJ12 | 2.63607144 | 1.409867644 | 0.016706569 | 0.044064923 |
| LOC119866207 | 2.635878804 | 3.723814001 | 1.94E-06 | 1.83E-05 |
| LOC119864722 | 2.634331377 | -0.135021837 | 0.00561355 | 0.017673223 |
| RHOBTB2 | 2.632985208 | 4.497796295 | 9.79E-05 | 0.000561637 |
| ANXA4 | 2.631114226 | 6.028109253 | 8.30E-06 | 6.46E-05 |
| NCDN | 2.628773631 | 4.546258231 | 1.46E-06 | 1.43E-05 |
| SCN2B | 2.628193495 | 2.879795645 | 0.001885124 | 0.006991037 |
| NCOR2 | 2.627717374 | 9.684327458 | 0.000314908 | 0.001519366 |
| LOC106558034 | 2.626387622 | -0.064956392 | 0.001644458 | 0.00622974 |
| LOC119870675 | 2.624962518 | 0.166466188 | 0.013965784 | 0.038197934 |
| RARB | 2.623152959 | 3.999055704 | 0.000136402 | 0.000742435 |
| PLXNB3 | 2.622944191 | 6.565481951 | 0.000193374 | 0.000996906 |
| GPR34 | 2.622385579 | 4.0271228 | 8.81E-05 | 0.000512358 |
| LOC119865621 | 2.620459358 | 2.584248879 | 3.16E-05 | 0.000210781 |
| TCF7L2 | 2.62034309 | 7.767948537 | 0.003160907 | 0.010886118 |

| | | | | |
|---|---|---|---|---|
| MYOT | 2.620329325 | 0.610708838 | 0.008199525 | 0.02438034 |
| TMEM141 | 2.618790923 | 2.48701007 | 0.000142441 | 0.000769281 |
| LOC119874395 | 2.618744748 | 3.444857626 | 0.001017679 | 0.004155352 |
| ARHGAP20 | 2.618544948 | 2.859548096 | 9.17E-05 | 0.00053049 |
| RASGRF2 | 2.617490281 | 6.356184147 | 0.009946541 | 0.028735187 |
| PCGF2 | 2.617269457 | 5.754296543 | 6.66E-06 | 5.33E-05 |
| LOC102153020 | 2.616813623 | -0.736542419 | 0.011173496 | 0.031636561 |
| LOC119868044 | 2.614981961 | 1.950540668 | 0.000928429 | 0.003850335 |
| SGCE | 2.613939217 | 3.508202414 | 0.018814923 | 0.048481255 |
| CHAC1 | 2.61379912 | 1.383418875 | 0.003383764 | 0.011521811 |
| EBI3 | 2.613583882 | 1.965766367 | 0.000764921 | 0.003248705 |
| REEP1 | 2.612241961 | 1.479614863 | 0.006878 | 0.021013095 |
| KANK3 | 2.611983121 | 4.751220143 | 3.15E-05 | 0.000210209 |
| LMNTD2 | 2.61185121 | 3.335172451 | 0.01597988 | 0.042574474 |
| DMTN | 2.610107199 | 4.45940365 | 0.008672243 | 0.025571907 |
| DIP2A | 2.610085812 | 7.529305053 | 4.89E-05 | 0.000308005 |
| OR51E2 | 2.607562145 | -0.083286317 | 0.015574658 | 0.041698526 |
| FZD9 | 2.607163063 | 1.618610805 | 0.013133544 | 0.036241827 |
| VPS37D | 2.606341513 | 0.940575819 | 0.001870414 | 0.006948537 |
| LOC102157379 | 2.605810376 | 0.02193223 | 0.007390326 | 0.022359376 |
| SOCS2 | 2.604470696 | 4.181044903 | 0.000939373 | 0.00388383 |
| CMSS1_2 | 2.603515665 | 3.162342569 | 6.63E-05 | 0.00040087 |
| RTL6 | 2.602484412 | 6.347951129 | 4.06E-07 | 4.66E-06 |
| HHAT | 2.601287735 | 2.822759655 | 0.002357049 | 0.008454871 |
| LOC119870210 | 2.601025482 | 4.895778756 | 1.12E-05 | 8.43E-05 |
| TRPM3 | 2.599327111 | 3.622047556 | 0.001720059 | 0.006466254 |
| JUN | 2.598713176 | 7.710232216 | 0.000121598 | 0.000675397 |
| BCAT2 | 2.598193525 | 4.79677576 | 9.00E-06 | 6.94E-05 |
| SH3RF1 | 2.597962413 | 4.661681837 | 1.54E-05 | 0.000111665 |
| LOC119876980 | 2.59605562 | 0.450369465 | 0.006808387 | 0.020853332 |
| LOC483462 | 2.595680023 | 0.996350037 | 0.002227005 | 0.008071216 |
| LOC102154523 | 2.59405113 | 0.349843854 | 0.008201781 | 0.024383702 |
| DLC1 | 2.591957498 | 7.368872842 | 1.98E-05 | 0.000139968 |
| FOXP4 | 2.589674885 | 5.849643683 | 4.89E-06 | 4.07E-05 |
| LOC119871943 | 2.588902499 | 1.661059189 | 0.01150836 | 0.032434421 |
| LOC119877185 | 2.588661885 | 1.022559616 | 0.00147079 | 0.0056639 |
| SMPD3 | 2.588101974 | 2.373879048 | 0.012207565 | 0.034046398 |
| VANGL2 | 2.584879277 | 2.529079762 | 0.004685062 | 0.015139308 |
| PHLPP1 | 2.584359869 | 5.622493944 | 0.000513411 | 0.002308059 |

| | | | | |
|---|---|---|---|---|
| LOC100685532 | 2.584320587 | 2.761656606 | 0.001989682 | 0.007338729 |
| RIMBP2 | 2.582209043 | 2.806165756 | 0.013396637 | 0.036817817 |
| SYT7 | 2.581663796 | 4.61235059 | 2.57E-05 | 0.000176144 |
| LOC102155087 | 2.577597864 | -0.084048019 | 0.014932159 | 0.040306972 |
| EPHB6 | 2.577419189 | 4.358983545 | 0.000180602 | 0.000940397 |
| LOC119869609 | 2.576777698 | -0.31261989 | 0.004394209 | 0.014355696 |
| PEA15 | 2.57563588 | 7.883032473 | 2.69E-06 | 2.44E-05 |
| SRRM4 | 2.575125006 | 0.784192246 | 0.009557686 | 0.027763533 |
| LOC119863913 | 2.574984325 | 0.66195416 | 0.017639072 | 0.046025959 |
| TMCC3 | 2.574114438 | 5.093646849 | 6.96E-06 | 5.54E-05 |
| PLEKHN1 | 2.573531093 | 4.07980466 | 0.000598159 | 0.002631195 |
| LOC102154060 | 2.57287223 | 2.296680208 | 0.000245691 | 0.001224388 |
| CDK3 | 2.570873927 | 1.477324028 | 0.009901084 | 0.028619123 |
| LOC484356 | 2.569397762 | 4.169379109 | 0.000407413 | 0.001896448 |
| EVI5L | 2.567394023 | 5.210485872 | 2.06E-05 | 0.000144587 |
| FLT1 | 2.5657687 | 6.073716152 | 0.001589223 | 0.006054869 |
| PROX1 | 2.564980752 | 1.376728366 | 0.018298698 | 0.047408867 |
| DMWD | 2.564662628 | 5.030975675 | 5.64E-06 | 4.61E-05 |
| LRFN3 | 2.562952936 | 3.372859881 | 0.000435428 | 0.002005319 |
| PRTFDC1 | 2.56281994 | 2.992809036 | 0.002990884 | 0.010386385 |
| NPRL3 | 2.561632736 | 4.763354093 | 0.0002818 | 0.001377723 |
| PDE8B | 2.561156307 | 2.725794342 | 0.011505819 | 0.032431475 |
| AVIL | 2.559943172 | 4.083086653 | 0.00029261 | 0.001426035 |
| FARP2 | 2.559690656 | 6.269786012 | 0.000110874 | 0.000625772 |
| SLC1A4 | 2.5593955 | 4.48205116 | 3.10E-05 | 0.000207533 |
| LOC119865629 | 2.558188253 | 1.351374866 | 0.012714613 | 0.035260884 |
| ARHGAP23 | 2.558072162 | 6.385773498 | 2.98E-06 | 2.67E-05 |
| LOC119863959 | 2.556904271 | 0.57306252 | 0.018811887 | 0.048479196 |
| LMNB2 | 2.555732267 | 4.408340406 | 0.000237685 | 0.001191062 |
| POGLUT2 | 2.554936839 | 2.75213498 | 0.000430126 | 0.001984841 |
| STARD9 | 2.55367798 | 7.391779706 | 2.50E-06 | 2.29E-05 |
| LOC119865179 | 2.552653716 | 2.830765969 | 0.000190704 | 0.000984784 |
| BAD | 2.552039509 | 1.73217505 | 0.005197754 | 0.016539557 |
| CFAP410 | 2.551168137 | 4.046463163 | 0.000236242 | 0.001184381 |
| PTPRR | 2.548733879 | 2.377691486 | 0.00289975 | 0.010115289 |
| SPRED3 | 2.546042214 | 3.322120026 | 0.000537233 | 0.00240071 |
| LOC111097081 | 2.544031755 | 1.409637306 | 0.003878489 | 0.012948011 |
| ARHGEF33 | 2.543656289 | 1.777337584 | 0.000949467 | 0.003918834 |
| LOC119869194 | 2.543592071 | 0.632785906 | 0.014656858 | 0.039727123 |

| | | | | |
|---|---|---|---|---|
| CCDC152 | 2.542803529 | 3.144605083 | 0.002539072 | 0.00901972 |
| LOC119877091 | 2.541836493 | 3.950171988 | 4.87E-06 | 4.05E-05 |
| LOXL3 | 2.540724079 | 6.007294764 | 4.91E-07 | 5.50E-06 |
| LOC102154667 | 2.539099248 | -0.431086447 | 0.011587468 | 0.03260649 |
| CPQ | 2.534772839 | 3.661458218 | 0.008640514 | 0.025514256 |
| LOC100682872 | 2.534497641 | 1.525464104 | 0.003671982 | 0.012355669 |
| FZD8 | 2.534425844 | 3.831615379 | 0.000884913 | 0.003691054 |
| PTX4 | 2.533581513 | 1.360599429 | 0.004261378 | 0.013970178 |
| GDPD5 | 2.531759853 | 4.549021021 | 1.17E-05 | 8.76E-05 |
| LOC119878089 | 2.531560942 | -0.507372793 | 0.014922417 | 0.040285693 |
| ANAPC11 | 2.530352763 | 5.236335041 | 0.000153607 | 0.000818366 |
| GALM | 2.528281923 | 3.987254153 | 0.000130418 | 0.000715344 |
| MTMR11 | 2.526852973 | 5.552865023 | 4.16E-05 | 0.000266958 |
| TBKBP1 | 2.526476976 | 4.24625251 | 6.97E-05 | 0.000418876 |
| ADARB1 | 2.526452905 | 5.200028255 | 3.78E-05 | 0.000246131 |
| LOC102154979 | 2.526357863 | 1.234738431 | 0.01393603 | 0.038126177 |
| IFT27 | 2.525175084 | 2.706488868 | 0.001701068 | 0.006409409 |
| TMOD2 | 2.524925188 | 4.878702824 | 0.018992539 | 0.048846007 |
| NPTXR | 2.521740638 | 3.458831258 | 0.010366849 | 0.029723604 |
| PARP2 | 2.520036329 | 3.531615021 | 2.42E-05 | 0.000166924 |
| GALR1 | 2.519786625 | 1.848007755 | 0.012307462 | 0.034302935 |
| FBXL8 | 2.519268205 | 3.428830156 | 0.001114802 | 0.004494251 |
| LOC111097651 | 2.517125816 | 2.428255747 | 0.00085796 | 0.003589683 |
| MEI1 | 2.516744062 | 2.764369662 | 0.002582326 | 0.009153868 |
| METTL1 | 2.514197568 | 3.247222008 | 0.000110759 | 0.000625346 |
| SLC52A2 | 2.512827123 | 4.652047193 | 5.15E-05 | 0.000321985 |
| MDK | 2.510651634 | 5.772627663 | 0.000329816 | 0.001577946 |
| CTNND1 | 2.510522873 | 7.968815832 | 1.79E-06 | 1.72E-05 |
| LOC106558258 | 2.508564668 | 4.172399176 | 9.16E-05 | 0.000529898 |
| EVPL | 2.506463819 | 4.263063017 | 0.00079264 | 0.003349362 |
| BNIP3 | 2.505502589 | 5.080900963 | 2.64E-05 | 0.000179439 |
| ANXA5 | 2.505085152 | 6.897455457 | 1.80E-05 | 0.000128355 |
| LOC119864089 | 2.504652532 | 4.582836959 | 0.00022087 | 0.001118179 |
| LOC607201 | 2.504292125 | 2.57743593 | 0.003749577 | 0.012577711 |
| MIXL1 | 2.504261785 | 0.226137393 | 0.014905795 | 0.040255861 |
| GPRIN1 | 2.503700421 | 2.660188587 | 0.000722535 | 0.003092318 |
| LOC119876345 | 2.502781829 | 2.103391612 | 0.011696035 | 0.03284802 |
| CCNF | 2.500672576 | 2.671424098 | 0.002196248 | 0.007983779 |
| LOC100682707 | 2.500175823 | 0.561099332 | 0.003269426 | 0.011195808 |

| | | | | |
|---|---|---|---|---|
| VEGFA | 2.499485468 | 7.092552514 | 0.002765678 | 0.009724293 |
| GPX7 | 2.498106693 | 3.22672009 | 0.011540815 | 0.032500531 |
| PCYOX1L | 2.496624674 | 3.595293811 | 0.000546691 | 0.002435946 |
| LOC111092399 | 2.495763578 | 0.338181656 | 0.001614075 | 0.006131786 |
| LOC119872972 | 2.494468373 | 1.160608061 | 0.001359861 | 0.005315107 |
| BDH2 | 2.494265084 | 2.023645045 | 0.000590498 | 0.00260119 |
| LOC111095184 | 2.494165132 | 1.289910658 | 0.012046362 | 0.033670467 |
| GAMT | 2.494097877 | 3.664329579 | 0.000503148 | 0.002270398 |
| RND3 | 2.493897488 | 6.26483735 | 0.000114336 | 0.000641137 |
| PKMYT1 | 2.492085596 | 2.983612187 | 0.001764599 | 0.006610918 |
| LOC119866695 | 2.49067314 | -0.342919093 | 0.006539216 | 0.020117025 |
| NRP2 | 2.489391007 | 8.187583972 | 1.31E-05 | 9.71E-05 |
| FRRS1 | 2.489264899 | 1.350653005 | 0.018411484 | 0.047651313 |
| NFE2L1 | 2.48890369 | 7.951773778 | 1.02E-07 | 1.38E-06 |
| LOC100856611 | 2.48601781 | 0.119903601 | 0.010464339 | 0.0299358 |
| ARHGAP31 | 2.48111891 | 8.239953413 | 0.000168514 | 0.000887103 |
| MT1E | 2.481006715 | 3.362511758 | 0.016024505 | 0.042665735 |
| L3MBTL1 | 2.480723285 | 4.614335631 | 8.50E-05 | 0.000497381 |
| CYP2W1 | 2.480689717 | 3.197078785 | 1.49E-05 | 0.000108434 |
| ZSWIM9 | 2.479013165 | 2.011107233 | 0.002578002 | 0.009140035 |
| NEURL1 | 2.478983137 | 2.022951807 | 0.00874562 | 0.02574624 |
| PKIG | 2.478124129 | 4.598643781 | 1.54E-05 | 0.00011172 |
| LOC102156785 | 2.477529523 | 1.542011663 | 0.001154272 | 0.004630978 |
| PLCD1 | 2.47610655 | 6.264712066 | 0.000337043 | 0.001606496 |
| SLC4A2 | 2.475326855 | 6.81322568 | 9.76E-05 | 0.000559942 |
| LOC119877175 | 2.475242906 | 1.688285849 | 0.006886634 | 0.02103058 |
| VWA5A | 2.473999841 | 5.134878565 | 0.006263298 | 0.019397606 |
| CERS4 | 2.471988356 | 5.271470834 | 0.000945787 | 0.003905162 |
| NECTIN2 | 2.469148241 | 4.970540593 | 3.97E-06 | 3.41E-05 |
| LOC119878549 | 2.468898266 | 1.337073512 | 0.003360223 | 0.011452448 |
| MRC1 | 2.468721095 | 6.017299847 | 0.013529024 | 0.037115763 |
| CPT1B | 2.468164026 | 5.49302678 | 0.001870748 | 0.006948537 |
| PLCB3 | 2.467881463 | 6.636427928 | 0.000289398 | 0.001411299 |
| MICAL3 | 2.467702797 | 7.202793802 | 0.000124703 | 0.000689284 |
| LOC102157101 | 2.466627248 | 3.969998986 | 9.08E-07 | 9.39E-06 |
| LOC106558962 | 2.466306678 | 2.387993646 | 0.002859301 | 0.010006398 |
| APP | 2.464854299 | 8.598071103 | 1.25E-05 | 9.26E-05 |
| CYB5R2 | 2.464734823 | 3.258697419 | 0.004143924 | 0.013665743 |
| COMP | 2.464627633 | 6.321160896 | 0.017373821 | 0.045465205 |

| | | | | |
|---|---|---|---|---|
| SMKR1 | 2.463762841 | -0.450219865 | 0.0142314 | 0.038762992 |
| MTA3 | 2.463645256 | 4.267436503 | 0.000679994 | 0.00293986 |
| OSBP2 | 2.462649754 | 1.512044008 | 0.003287059 | 0.011245529 |
| DAPK2 | 2.461702326 | 1.589463034 | 0.000668149 | 0.002897319 |
| USP36 | 2.461416009 | 6.60328326 | 0.000198161 | 0.00101916 |
| CNN1 | 2.461309938 | 3.956094585 | 0.016657091 | 0.043971879 |
| LRRC20 | 2.458860323 | 3.510547964 | 1.47E-05 | 0.000107143 |
| ZNF316 | 2.45859437 | 5.861334038 | 1.50E-05 | 0.000109429 |
| TIAM2 | 2.457158385 | 5.606737376 | 2.86E-05 | 0.000193092 |
| TTC39A | 2.456770221 | 6.772232168 | 0.000367 | 0.001730633 |
| LOC119869542 | 2.453607569 | 4.007061906 | 0.006813677 | 0.020866585 |
| LOC119867842 | 2.452923858 | 1.391695066 | 0.002939503 | 0.010239128 |
| LOC102155446 | 2.451295388 | 1.522768634 | 0.003526491 | 0.011934461 |
| ARMH4 | 2.450438421 | 2.866402939 | 0.001702329 | 0.006411989 |
| LOC119876295 | 2.450392838 | 0.009360048 | 0.009896454 | 0.028609556 |
| RDH10 | 2.450201413 | 5.162136748 | 0.00041929 | 0.001943803 |
| LOC119863916 | 2.449814926 | 4.06955906 | 2.08E-06 | 1.95E-05 |
| LOC100856555 | 2.449067246 | -0.368889152 | 0.002830789 | 0.009919433 |
| C11H5orf63 | 2.449009311 | 1.861232806 | 0.007695938 | 0.023127297 |
| YPEL2 | 2.448632601 | 5.674208295 | 2.33E-07 | 2.85E-06 |
| DLG4 | 2.447798187 | 5.603581706 | 0.000135058 | 0.000736194 |
| SRMS | 2.447328748 | 2.929224382 | 0.006770455 | 0.02074454 |
| LOC111092918 | 2.446877422 | 2.673320398 | 0.006835188 | 0.020905876 |
| APBB1 | 2.443600944 | 5.372372787 | 4.38E-07 | 4.99E-06 |
| MIER2 | 2.442490279 | 5.770794467 | 0.000191575 | 0.000988152 |
| WDR88 | 2.441397284 | 1.40463041 | 0.018625829 | 0.048096937 |
| PPP2R3B | 2.441111735 | 4.762804518 | 0.000547664 | 0.002438778 |
| LZTS2 | 2.440872046 | 6.57042476 | 1.13E-05 | 8.47E-05 |
| RASD1 | 2.438638081 | 2.532668063 | 0.002282363 | 0.008238783 |
| LOC111096666 | 2.438029454 | 0.196120887 | 0.015503387 | 0.041564188 |
| VMO1 | 2.437911321 | 1.765501589 | 0.009704088 | 0.028102178 |
| SFT2D3 | 2.437902685 | -0.154384761 | 0.004609595 | 0.014944467 |
| LOC111096392 | 2.435765514 | 2.813736632 | 0.002735287 | 0.009626807 |
| LOC119873060 | 2.435746032 | 3.958197558 | 0.012590431 | 0.034961231 |
| NGEF | 2.435669733 | 2.801113236 | 0.003730442 | 0.012525154 |
| CSF1 | 2.435408647 | 7.183260456 | 0.000275599 | 0.001349845 |
| NPDC1 | 2.434558894 | 5.517308979 | 5.35E-05 | 0.000333148 |
| LOC119864426 | 2.43435794 | 1.291361769 | 0.009978783 | 0.028809133 |
| LPAR1 | 2.433968118 | 6.805595635 | 0.001952557 | 0.007217768 |

| | | | | |
|---|---|---|---|---|
| WDR93 | 2.433765095 | 1.218450998 | 0.004342035 | 0.014195946 |
| SPRY1 | 2.429964777 | 4.932956262 | 1.61E-05 | 0.00011622 |
| FAM210B | 2.429112107 | 4.099350156 | 3.40E-05 | 0.000223993 |
| LOC119878020 | 2.426282236 | 3.517541813 | 0.001424639 | 0.005511639 |
| NOX5 | 2.426118461 | 3.789083124 | 0.018987815 | 0.048839971 |
| LOC119872164 | 2.422674114 | 3.840116251 | 0.000816656 | 0.003436107 |
| LOC102151240 | 2.422483306 | 3.456918607 | 0.000840396 | 0.003526408 |
| LOC111091327 | 2.422247487 | 2.628841419 | 0.000513153 | 0.00230738 |
| MARVELD2 | 2.421864172 | 1.260549882 | 0.015588237 | 0.041714268 |
| LOC102153220 | 2.421021828 | 0.676289665 | 0.014489807 | 0.039358014 |
| LOC478678 | 2.420523736 | 4.162107382 | 0.00420406 | 0.013827853 |
| NECTIN1 | 2.420489812 | 6.171095153 | 8.76E-05 | 0.000510097 |
| DNAAF3 | 2.417592753 | 3.03544718 | 0.010197265 | 0.029332983 |
| LOC106559925 | 2.415465726 | 6.701622773 | 0.000117416 | 0.00065654 |
| CCDC112 | 2.415165317 | 3.781103001 | 0.000156305 | 0.000831516 |
| RIOX2 | 2.412808323 | 4.79414652 | 0.00010128 | 0.000578859 |
| HS1BP3 | 2.412544264 | 5.189285082 | 8.14E-06 | 6.36E-05 |
| LOC119877186 | 2.409812877 | 6.017507838 | 1.46E-05 | 0.000106458 |
| TMEM200B | 2.408524727 | 3.419465355 | 0.000136408 | 0.000742435 |
| MPRIP | 2.407834331 | 9.120008295 | 1.31E-05 | 9.69E-05 |
| LOC119873226 | 2.405349599 | 1.128104385 | 0.006070221 | 0.018897241 |
| PLD2 | 2.401949155 | 5.374473354 | 2.71E-06 | 2.45E-05 |
| WSCD2 | 2.401413356 | 3.701004204 | 0.018286588 | 0.04738465 |
| LOC102152748 | 2.400967184 | -0.033297273 | 0.016268932 | 0.043173326 |
| GPR173 | 2.400744773 | 3.56276163 | 5.36E-06 | 4.41E-05 |
| STYXL1 | 2.399356678 | 2.797518726 | 0.005796831 | 0.018182915 |
| LOC111096751 | 2.399298373 | 1.53172391 | 0.006224501 | 0.019296473 |
| BPNT2 | 2.398422257 | 6.019404948 | 0.000206733 | 0.001059224 |
| CASTOR2 | 2.398169775 | 5.260014192 | 2.61E-06 | 2.38E-05 |
| PTGFRN | 2.397563208 | 5.863902789 | 0.00103788 | 0.004224671 |
| LOC100855995 | 2.397374263 | 0.914492339 | 0.005200387 | 0.016543078 |
| TRIM3 | 2.396085003 | 4.724255753 | 3.95E-06 | 3.40E-05 |
| TRAF3IP2 | 2.395703159 | 5.537264672 | 0.000340483 | 0.001620397 |
| RSAD1 | 2.393802601 | 3.663946537 | 1.78E-05 | 0.000127372 |
| LOC119872883 | 2.393116256 | 1.914132569 | 0.014263678 | 0.03884603 |
| LOC119869693 | 2.392746484 | 6.334329588 | 0.000952963 | 0.003930428 |
| ZBTB12 | 2.39068676 | 3.19113768 | 0.000268206 | 0.001318698 |
| TLCD3A | 2.389970149 | 3.087465391 | 0.000588471 | 0.002592979 |
| C10H12orf56 | 2.38879152 | 3.345949946 | 7.66E-05 | 0.000455088 |

| | | | | |
|---|---|---|---|---|
| DHRS11 | 2.387619745 | 2.70677893 | 0.000211646 | 0.001079961 |
| SVBP | 2.386200619 | 3.734416918 | 0.000175455 | 0.000918066 |
| TEF | 2.384801225 | 5.634610991 | 3.33E-05 | 0.000220175 |
| ADGRB2 | 2.384284567 | 3.220704019 | 0.000762604 | 0.003240132 |
| PC | 2.383058567 | 4.819990739 | 0.00073877 | 0.003153083 |
| GET1 | 2.38293506 | 4.066028581 | 4.63E-05 | 0.00029378 |
| IL34 | 2.382213136 | 3.708310754 | 0.001138878 | 0.004579389 |
| CYTH3 | 2.381541083 | 6.427637171 | 1.15E-06 | 1.16E-05 |
| DHRS4 | 2.380948415 | 4.64475983 | 0.000136826 | 0.000744147 |
| LOC102151464 | 2.37863469 | -0.114876224 | 0.013386243 | 0.036806697 |
| LOC111096218 | 2.378293561 | 1.256149409 | 0.004021395 | 0.013345472 |
| ZNF618 | 2.377973991 | 5.519606675 | 0.000325446 | 0.001560483 |
| TMEM64 | 2.376688955 | 5.437707031 | 0.003262476 | 0.011175538 |
| HYKK | 2.37423348 | 3.056022495 | 4.48E-05 | 0.000285549 |
| HRK | 2.373707305 | 0.315087503 | 0.003128399 | 0.010794731 |
| CCDC187 | 2.373157006 | 1.622595409 | 0.011223049 | 0.031733483 |
| KDR | 2.373106258 | 5.815105709 | 0.003224144 | 0.011061717 |
| EPB41L4A | 2.37284449 | 4.237471168 | 0.000443027 | 0.002033401 |
| TRAIP | 2.371354861 | 2.945357817 | 0.002300772 | 0.008287293 |
| LOC102152627 | 2.370976266 | -0.319193012 | 0.0173118 | 0.045335747 |
| DBF4B | 2.370073948 | 3.074493202 | 0.000688095 | 0.002967778 |
| HINT3 | 2.369045868 | 4.044835912 | 5.59E-05 | 0.000345876 |
| FGFR3 | 2.368419268 | 3.044689124 | 0.01855126 | 0.047932935 |
| TRPC1 | 2.367562648 | 4.231953117 | 6.10E-05 | 0.000373607 |
| LOC111090138 | 2.367206204 | 2.631699078 | 0.014227699 | 0.038757783 |
| LOC106558323 | 2.365404482 | 4.674108345 | 0.00092299 | 0.003830709 |
| LOC111097086 | 2.363685603 | 4.42776 | 0.000164607 | 0.000869279 |
| ENKD1 | 2.362660932 | 1.920564227 | 0.016321922 | 0.043292745 |
| BRF1 | 2.361537649 | 6.019228566 | 0.001153773 | 0.004629854 |
| LOC119865992 | 2.36153496 | 6.356572738 | 0.000896661 | 0.003732873 |
| PLEKHH1 | 2.36123508 | 3.487048475 | 9.35E-05 | 0.000539408 |
| UGDH | 2.359805132 | 4.869423935 | 8.19E-06 | 6.39E-05 |
| SIRT4 | 2.359340407 | 5.371830094 | 0.000632017 | 0.002759965 |
| PEAR1 | 2.359038684 | 6.890517399 | 0.001712156 | 0.006443403 |
| SOGA1 | 2.358916848 | 7.422658271 | 1.99E-06 | 1.87E-05 |
| LOC612587 | 2.358539396 | 0.685373935 | 0.002221377 | 0.008060256 |
| SLC23A2 | 2.358372447 | 6.810006669 | 0.003568807 | 0.012047782 |
| LOC119872142 | 2.355558622 | 1.828868374 | 0.001611059 | 0.006123552 |
| DYNC2I2 | 2.353240311 | 2.824401416 | 0.000162761 | 0.000860789 |

| | | | | |
|---|---|---|---|---|
| TMEM132A | 2.352985513 | 5.962984043 | 5.97E-07 | 6.54E-06 |
| CAND2 | 2.351688509 | 3.344220969 | 0.000528897 | 0.002371284 |
| NDST1 | 2.351260771 | 7.18484243 | 3.11E-05 | 0.000207816 |
| TMEM169 | 2.350008084 | 1.350593599 | 0.003064398 | 0.010602603 |
| IRF2BPL | 2.349215921 | 6.819829682 | 0.000601085 | 0.002642995 |
| IZUMO4 | 2.347856705 | 2.477206959 | 0.001707431 | 0.006427854 |
| HSPBP1 | 2.343793349 | 3.256424384 | 0.000108367 | 0.00061354 |
| C15H1orf109 | 2.342862503 | 2.099745082 | 0.000181607 | 0.000944559 |
| LOC119872514 | 2.342853444 | 2.292770604 | 0.005496537 | 0.017352724 |
| LOC111092428 | 2.340663659 | 2.066844183 | 0.001419398 | 0.005494306 |
| LOC119877301 | 2.340432247 | 0.354166109 | 0.012036846 | 0.03365255 |
| MTA1 | 2.339961082 | 6.456280796 | 0.000835499 | 0.003509256 |
| TRPV3 | 2.339929132 | 1.899199473 | 0.006060715 | 0.018875397 |
| ZNF774 | 2.338937952 | 1.79318807 | 0.000659592 | 0.002864797 |
| DDAH2 | 2.338166601 | 5.417719083 | 0.000877213 | 0.00366105 |
| LOC119866653 | 2.337737526 | 2.055147048 | 0.001373314 | 0.005355926 |
| LOC479969 | 2.337092713 | 1.752863631 | 0.000339511 | 0.00161648 |
| LOC119870708 | 2.336320437 | 1.369868796 | 0.003510829 | 0.01189653 |
| LOC119876300 | 2.336154576 | 1.664274406 | 0.012723464 | 0.035280914 |
| FANCC | 2.335952019 | 4.975001495 | 8.04E-05 | 0.000473959 |
| RUSC1 | 2.335730084 | 5.866227472 | 0.000180455 | 0.00093992 |
| LOC119873717 | 2.335521297 | -0.48480738 | 0.01288282 | 0.035668033 |
| HEG1 | 2.333699084 | 8.299415254 | 8.59E-06 | 6.66E-05 |
| PFKL | 2.33330471 | 6.341657228 | 0.000120578 | 0.000670419 |
| RHBDL2 | 2.333108819 | 1.51298556 | 0.002881343 | 0.010073779 |
| HRAS | 2.331043436 | 3.250222924 | 0.000541513 | 0.002416845 |
| DVL1 | 2.329546177 | 5.88576511 | 0.000523835 | 0.002351019 |
| LOC481187 | 2.327236259 | 4.580731413 | 0.001110756 | 0.004479609 |
| NT5DC3 | 2.325203835 | 3.207396451 | 0.000306811 | 0.001486256 |
| EPAS1 | 2.325185813 | 8.123982192 | 0.000987048 | 0.004048502 |
| DSTN | 2.322317051 | 6.124467072 | 0.002667783 | 0.009422891 |
| LOC119869433 | 2.321810102 | 4.546526204 | 0.000204787 | 0.001050498 |
| NSG1 | 2.319707721 | 4.854332899 | 0.001745616 | 0.006548854 |
| DMD | 2.317917976 | 7.008204114 | 0.000933573 | 0.003866972 |
| C20H19orf44 | 2.317260819 | 5.322442911 | 0.000105877 | 0.000601484 |
| LOC111097267 | 2.316805591 | 2.796463471 | 0.000620772 | 0.002717429 |
| AIF1L | 2.316449798 | 5.669230152 | 0.013660107 | 0.03744694 |
| ACCS | 2.314362011 | 4.602468795 | 3.02E-05 | 0.000202254 |
| LOC100688091 | 2.312384586 | 2.790958128 | 0.00016894 | 0.0008887 |

| | | | | |
|---|---|---|---|---|
| GRK4 | 2.312247253 | 2.329752295 | 0.005646453 | 0.017761333 |
| CSRP2 | 2.311993745 | 3.358238309 | 0.001145431 | 0.004602322 |
| PHEX | 2.310842327 | 3.124190543 | 0.003918461 | 0.013061334 |
| FAM171B | 2.310817197 | 4.184074314 | 0.002560537 | 0.009082574 |
| LOC111097105 | 2.310603491 | 1.050016432 | 0.017159768 | 0.045057385 |
| ZNF853 | 2.310391603 | 4.246821379 | 0.00039467 | 0.001844263 |
| HDLBP | 2.307480571 | 9.032728845 | 0.000118595 | 0.000661768 |
| LOC119873203 | 2.304995037 | 4.367671584 | 0.000582443 | 0.002570935 |
| CRACD | 2.304961399 | 3.720041897 | 0.002613244 | 0.009249853 |
| LOC119866753 | 2.304558919 | 2.462741184 | 0.015050868 | 0.040581919 |
| LOC102156231 | 2.303834537 | 1.614637732 | 0.00405886 | 0.013434612 |
| MAP4 | 2.30278623 | 9.400713226 | 0.000127913 | 0.00070469 |
| LOC102152879 | 2.300728555 | 1.714875945 | 0.013482156 | 0.03702041 |
| LOC111090737 | 2.300486014 | -0.520216097 | 0.017835274 | 0.046459601 |
| ARHGAP39 | 2.299869591 | 5.005104513 | 6.99E-05 | 0.000419469 |
| LOC102153565 | 2.299861592 | 1.90342378 | 0.018810585 | 0.048479196 |
| LOC119878766 | 2.299123633 | 1.319361405 | 0.002258412 | 0.00816568 |
| LOC119873279 | 2.295581476 | 3.725385764 | 0.000110043 | 0.000621767 |
| LATS2 | 2.294253068 | 6.095885038 | 8.22E-07 | 8.61E-06 |
| LOC608573 | 2.292286765 | 1.617231428 | 0.001555817 | 0.005951151 |
| SIMC1 | 2.291772136 | 4.746782909 | 0.000240179 | 0.001201059 |
| ZNF608 | 2.291295319 | 5.848305987 | 0.000122161 | 0.000678178 |
| LOC485387 | 2.290912447 | 4.104235079 | 0.004453525 | 0.014510109 |
| LRRC24 | 2.290262745 | 4.678996226 | 0.000371233 | 0.00174838 |
| PEPD | 2.289163095 | 5.334528948 | 0.000705176 | 0.003029393 |
| EBPL | 2.288905423 | 2.461981267 | 0.000356391 | 0.001686479 |
| LOC102151135 | 2.288760262 | 2.182735404 | 0.000209237 | 0.001070283 |
| TPGS1 | 2.288404141 | 3.731871434 | 0.001236977 | 0.004904685 |
| AURKB | 2.286511742 | 4.897738217 | 0.000710578 | 0.00305018 |
| RNASE10 | 2.285821252 | 1.307397155 | 0.011877227 | 0.033274967 |
| PARD6G | 2.283713061 | 2.935357755 | 0.001711158 | 0.006440764 |
| LOC111095906 | 2.28252838 | 0.928394255 | 0.005990107 | 0.018687721 |
| POLD2 | 2.281126944 | 4.648723011 | 7.24E-05 | 0.000432955 |
| ZNF704 | 2.281001815 | 3.606011508 | 0.00016289 | 0.000861051 |
| RMI2 | 2.280492443 | 1.493835185 | 0.009052019 | 0.026522159 |
| HMG20B | 2.278826952 | 5.680898488 | 0.000772682 | 0.003278452 |
| SLC2A8 | 2.277880206 | 3.405432305 | 9.31E-05 | 0.000537263 |
| LOC119876450 | 2.27696914 | 0.782504554 | 0.007920267 | 0.023709386 |
| DLG5 | 2.27649904 | 6.767058262 | 1.84E-05 | 0.000130663 |

| | | | | |
|---|---|---|---|---|
| MRPS34 | 2.276299488 | 2.147635534 | 0.001188416 | 0.004747748 |
| DLG2 | 2.275695928 | 2.563857563 | 0.00165503 | 0.006263219 |
| ST8SIA5 | 2.275585533 | 1.636746782 | 0.002602723 | 0.009215622 |
| NEXN | 2.274365607 | 3.48086846 | 0.0015971 | 0.00607902 |
| PKDCC | 2.274048755 | 6.074451016 | 0.000121742 | 0.000676021 |
| LOC119868078 | 2.274034193 | 2.834178162 | 0.00397612 | 0.013229114 |
| P4HA2 | 2.273218668 | 4.931881506 | 0.000751506 | 0.003201138 |
| LOC106558651 | 2.272254859 | 2.868379835 | 0.006622595 | 0.020336003 |
| LOC102151970 | 2.271949562 | 0.267530491 | 0.005149148 | 0.016413809 |
| C21H11orf54 | 2.271584009 | 2.987140973 | 0.000731268 | 0.003126089 |
| TSPYL2 | 2.270544551 | 6.180844651 | 0.00618228 | 0.01917958 |
| TROAP | 2.269124968 | 2.539937359 | 0.013246477 | 0.036493059 |
| CHRNE | 2.268595627 | 5.450713659 | 8.17E-05 | 0.000480588 |
| SRC | 2.266154689 | 5.563520785 | 5.39E-07 | 5.97E-06 |
| LOC119866448 | 2.265263918 | 1.568241305 | 0.002273814 | 0.008214763 |
| AVPI1 | 2.264235839 | 3.952355083 | 0.000265 | 0.001305307 |
| MARCHF3 | 2.264044318 | 1.894384642 | 0.016659442 | 0.04397273 |
| PLTP | 2.262605975 | 6.048598809 | 0.000469545 | 0.002136564 |
| TEK | 2.261526402 | 6.169814131 | 0.00060975 | 0.002675124 |
| LOC119878307 | 2.261129914 | 6.832184684 | 0.01086689 | 0.030903831 |
| LOC119865740 | 2.261071036 | 4.826610331 | 0.000176013 | 0.000919875 |
| LOC119876372 | 2.260504371 | 2.087662766 | 0.002204334 | 0.008009142 |
| RAB3IL1 | 2.258625819 | 6.289857525 | 0.001486326 | 0.00571459 |
| LOC111097305 | 2.256351774 | 1.349923033 | 0.012614295 | 0.035009554 |
| LOC111094002 | 2.256090168 | 2.18447926 | 0.002378116 | 0.008520567 |
| MLXIPL | 2.255734929 | 3.256433196 | 0.011799183 | 0.033099105 |
| GLI4 | 2.254886552 | 4.819678933 | 0.000451914 | 0.002067623 |
| NAV1 | 2.253966468 | 9.444479181 | 0.002472597 | 0.008817521 |
| LOC102156973 | 2.252283945 | 1.516131678 | 0.005766454 | 0.018107267 |
| LOC119868281 | 2.251750948 | 1.694087639 | 0.010156947 | 0.029241698 |
| TMBIM1 | 2.250678176 | 6.796993826 | 9.78E-06 | 7.47E-05 |
| LOC111094421 | 2.250168469 | 5.22491942 | 0.000521885 | 0.002343239 |
| EFNA1 | 2.248673377 | 3.202331237 | 0.000267973 | 0.001317855 |
| LOC102153929 | 2.247139545 | 1.011588853 | 0.009378063 | 0.0273407 |
| ADAMTSL4 | 2.245539135 | 7.176747112 | 0.000630872 | 0.002757185 |
| SIK1 | 2.244358884 | 6.489683618 | 0.002167187 | 0.007888719 |
| LOC111092136 | 2.243439646 | 1.719116396 | 0.004951442 | 0.015872297 |
| TNS3 | 2.242513507 | 7.266983566 | 0.000469714 | 0.002136883 |
| CTIF | 2.242446481 | 4.855152036 | 0.000260189 | 0.001284819 |

| | | | | |
|---|---|---|---|---|
| ZNF598 | 2.242095464 | 6.565015318 | 0.001153032 | 0.004628574 |
| RAB23 | 2.240951916 | 5.249981826 | 0.001339758 | 0.005246942 |
| CMTM4 | 2.239558196 | 5.408571705 | 2.16E-05 | 0.000150907 |
| CALU | 2.238336086 | 7.794510386 | 0.000371135 | 0.00174838 |
| RNASEH2A | 2.237771408 | 3.51208635 | 0.000320409 | 0.001540767 |
| PPP1R16A | 2.237109481 | 4.579681333 | 0.001826686 | 0.006822295 |
| LOC119874445 | 2.236483146 | 3.263426644 | 0.00354318 | 0.011979928 |
| IL17RD | 2.235866652 | 5.689408035 | 0.005445514 | 0.017219233 |
| DEXI | 2.235773493 | 5.211904034 | 3.98E-05 | 0.000257509 |
| LOC111095969 | 2.233840131 | 1.444647787 | 0.011824715 | 0.033149274 |
| RPTOR | 2.233213156 | 6.737097065 | 0.00033068 | 0.001581457 |
| LOC106558641 | 2.233168199 | 2.681797341 | 0.003895849 | 0.012993958 |
| THAP4 | 2.232942231 | 4.397739892 | 0.000355869 | 0.00168474 |
| LOC119871843 | 2.230522393 | 0.332866993 | 0.015688556 | 0.04192578 |
| SIPA1L3 | 2.230124612 | 7.393783754 | 0.000322743 | 0.001550958 |
| CLDN23 | 2.229058824 | 1.376491442 | 0.011857107 | 0.033222894 |
| TEC | 2.227399116 | 4.463316334 | 0.000735908 | 0.003142107 |
| ABL1 | 2.227389049 | 7.886433165 | 0.000146216 | 0.00078556 |
| CCDC150 | 2.226784725 | 3.677482582 | 0.002330665 | 0.008373837 |
| SSNA1 | 2.226615617 | 3.017584985 | 0.000644371 | 0.002808819 |
| PTPN21 | 2.226200277 | 7.10536153 | 2.44E-05 | 0.000167856 |
| LOC119866590 | 2.223299818 | 6.365446943 | 0.01181454 | 0.03313408 |
| LOC111096709 | 2.222092144 | 1.028417347 | 0.004738848 | 0.015283489 |
| SULT1A1 | 2.220612541 | 3.757951739 | 0.000195596 | 0.001007404 |
| HRH2 | 2.217298899 | 3.00955695 | 0.002714117 | 0.009561616 |
| LOC119865440 | 2.217038402 | 1.424543607 | 0.010926373 | 0.03104041 |
| LOC111094310 | 2.217020725 | 1.873836403 | 0.00144756 | 0.005591334 |
| KLF5 | 2.216514517 | 5.960126645 | 0.011428005 | 0.032233966 |
| PCNT | 2.21646204 | 6.923288257 | 0.001176315 | 0.004708084 |
| ENO2 | 2.216406273 | 5.0496668 | 7.98E-05 | 0.000471195 |
| NBEAL1 | 2.213661781 | 7.110797784 | 9.10E-05 | 0.000526937 |
| LOC111094751 | 2.213389767 | 2.381968557 | 0.011903478 | 0.033335582 |
| SEPTIN5 | 2.213350683 | 4.809126175 | 0.000837783 | 0.003518169 |
| COPRS | 2.212645083 | 2.879210744 | 0.00259779 | 0.00920417 |
| CDKN2B | 2.211820047 | 3.499886906 | 0.017077696 | 0.044879946 |
| ST14 | 2.210632932 | 0.992106878 | 0.005074183 | 0.016210608 |
| COBLL1 | 2.210286171 | 5.644018526 | 2.88E-05 | 0.000193929 |
| CDCA2 | 2.210252638 | 3.446498939 | 0.004248254 | 0.013933478 |
| GTF2F1 | 2.209249386 | 5.866937919 | 0.002633208 | 0.00931139 |

| | | | | |
|---|---|---|---|---|
| CENPX | 2.207871473 | 3.214055054 | 0.00134413 | 0.005258794 |
| NID1 | 2.207601206 | 7.957097745 | 0.000183278 | 0.000952562 |
| PON2 | 2.207487665 | 4.988963313 | 0.000543975 | 0.002425839 |
| SEMA6C | 2.206533499 | 4.460855535 | 5.96E-05 | 0.000365947 |
| SRPX | 2.20609314 | 4.293819415 | 0.000165832 | 0.000874897 |
| PRRT3 | 2.205712211 | 1.855153389 | 0.015980088 | 0.042574474 |
| LOC119870209 | 2.205648963 | 4.585368121 | 0.000227631 | 0.001146779 |
| SYTL4 | 2.204527096 | 4.003479952 | 0.000894641 | 0.003727329 |
| SCARF1 | 2.204231226 | 4.441459279 | 0.002546094 | 0.009038739 |
| GAB1 | 2.203234922 | 7.473040516 | 0.000472446 | 0.00214661 |
| LOC489638 | 2.202941708 | 0.715184677 | 0.008258263 | 0.024528072 |
| ANKS6 | 2.200594352 | 6.53023469 | 0.004274516 | 0.014004775 |
| MBD3 | 2.200290658 | 5.523261659 | 0.000438728 | 0.002018373 |
| C14H1orf35 | 2.200067472 | 4.889995676 | 0.005783888 | 0.018151491 |
| TTLL7 | 2.199027966 | 3.772383248 | 0.000644594 | 0.002809228 |
| THOP1 | 2.198325725 | 4.041715474 | 0.000230999 | 0.001161587 |
| LOC119870875 | 2.197641539 | -0.0211358 | 0.008096956 | 0.024138284 |
| FBF1 | 2.196439882 | 4.969043973 | 0.002669381 | 0.009425635 |
| LOC119876310 | 2.195591908 | 2.483560483 | 0.000902846 | 0.003755734 |
| CAPN12 | 2.194902291 | 4.185860264 | 0.00345062 | 0.011718159 |
| LOC607624 | 2.194438436 | 2.978040599 | 0.004014701 | 0.013328831 |
| LOC119874406 | 2.193753658 | 2.786379885 | 0.013497673 | 0.037048515 |
| SH3BP4 | 2.191566249 | 5.46880329 | 2.07E-05 | 0.000145005 |
| FAM174C | 2.191565355 | 4.5012805 | 0.001080806 | 0.004377558 |
| TOMM7 | 2.191325944 | 2.012133237 | 0.000410929 | 0.00191117 |
| LOC100856547 | 2.191039431 | 8.631906424 | 0.000416921 | 0.001935717 |
| RTL5 | 2.189765632 | 7.855885282 | 0.000782227 | 0.003311173 |
| FSTL3 | 2.189695648 | 5.797465803 | 0.001232415 | 0.004889279 |
| FN3K | 2.187573583 | 2.985385621 | 0.009121238 | 0.026699703 |
| LOC100684379 | 2.186872516 | 2.285736431 | 0.003393306 | 0.011549557 |
| TRIM66 | 2.186275591 | 2.529346199 | 0.002313093 | 0.008323372 |
| ZFP41 | 2.185387141 | 5.012885607 | 0.000327213 | 0.001567572 |
| LPCAT4 | 2.185195254 | 4.594321383 | 4.35E-05 | 0.000277707 |
| VEGFC | 2.184114702 | 3.198816196 | 0.001562039 | 0.005969684 |
| CABLES1 | 2.184086161 | 3.625440375 | 0.002225556 | 0.008067314 |
| LOC111091695 | 2.182882003 | 2.721231883 | 0.000814837 | 0.003429786 |
| STARD13 | 2.179264436 | 6.999850121 | 5.76E-05 | 0.000355358 |
| SHE | 2.178359554 | 4.9396907 | 0.011817042 | 0.033136337 |
| SYT17 | 2.178324025 | 4.057124496 | 0.001312201 | 0.00515766 |

| | | | | |
|---|---|---|---|---|
| LOC119871211 | 2.17729486 | 0.856068583 | 0.003563962 | 0.012035174 |
| GALNT2 | 2.176634452 | 6.624005261 | 5.21E-06 | 4.30E-05 |
| ZDHHC14 | 2.176225498 | 3.967175445 | 0.000800973 | 0.003377331 |
| LOC119878972 | 2.175816647 | 0.207081866 | 0.006154166 | 0.019114249 |
| LOC480918 | 2.174901749 | -0.600101548 | 0.010288542 | 0.029538161 |
| NKD2 | 2.174542673 | 2.185887804 | 0.004451456 | 0.014507729 |
| TUBA1C | 2.174090913 | 5.428349429 | 0.000701103 | 0.003015481 |
| LOC111092504 | 2.173489394 | 5.403645846 | 2.67E-05 | 0.000181307 |
| ARMC2 | 2.172256574 | 1.872765125 | 0.005097315 | 0.016270119 |
| CHST14 | 2.171533318 | 3.492191932 | 5.09E-05 | 0.000318757 |
| ST18 | 2.170472257 | 1.624352066 | 0.00753127 | 0.022720659 |
| NT5E | 2.169924658 | 4.245200863 | 0.016461619 | 0.043546034 |
| TNFAIP1 | 2.169704451 | 5.756630161 | 1.42E-05 | 0.000104417 |
| KDELR1 | 2.169505745 | 6.979082074 | 0.000471691 | 0.002144077 |
| CEP162 | 2.168797034 | 4.934988002 | 0.00099917 | 0.004093574 |
| CCDC124 | 2.165900439 | 3.709764395 | 0.004921923 | 0.015794023 |
| LOC111093313 | 2.163122879 | 3.652363773 | 0.000980554 | 0.004025676 |
| TLE1 | 2.16257831 | 5.654995953 | 0.001176985 | 0.004709895 |
| SPAG16 | 2.162450891 | 3.914683123 | 0.001372743 | 0.005354834 |
| ABCA2 | 2.1621373 | 7.155242288 | 0.000419708 | 0.001945053 |
| SHMT2 | 2.161888993 | 4.809454169 | 0.000400608 | 0.001867178 |
| LOC119867358 | 2.160600636 | 2.81165617 | 0.004626249 | 0.01498949 |
| XRCC1 | 2.159876752 | 5.170782022 | 9.02E-05 | 0.000523245 |
| NT5DC2 | 2.159767893 | 6.177463998 | 1.06E-05 | 8.06E-05 |
| NUDC | 2.159529134 | 5.737141532 | 0.000655253 | 0.002849505 |
| SGSH | 2.158853771 | 4.711863463 | 0.000162877 | 0.000861051 |
| PPP1R37 | 2.157813928 | 5.950911728 | 0.001312138 | 0.00515766 |
| MTG1 | 2.156788077 | 3.452090347 | 0.000305109 | 0.001479004 |
| PASK | 2.156317049 | 3.90874552 | 0.000458361 | 0.00209226 |
| EEF1AKMT3 | 2.156076984 | 2.62381512 | 0.002866325 | 0.010026122 |
| DTWD1 | 2.155219464 | 4.00342129 | 0.000562904 | 0.002498426 |
| ADAT3 | 2.154663933 | 2.330342615 | 0.00742536 | 0.022448017 |
| BEX4 | 2.152892359 | 2.211306134 | 0.008092836 | 0.024130746 |
| RRAS | 2.151753383 | 5.3109712 | 0.000117752 | 0.000657911 |
| ZNF875 | 2.151565072 | 2.132464054 | 0.008161303 | 0.02429002 |
| SLC24A2 | 2.151174621 | 0.920963649 | 0.009329335 | 0.027213282 |
| CRACR2B | 2.147324483 | 3.764733682 | 0.004745867 | 0.015301575 |
| ENGASE | 2.147115102 | 2.999183828 | 0.00167889 | 0.00634549 |
| RABEP1 | 2.146786967 | 6.420163669 | 0.001241011 | 0.004917982 |

| | | | | |
|---|---|---|---|---|
| SHF | 2.145931039 | 2.619439081 | 0.008622265 | 0.025466116 |
| DISP1 | 2.144838824 | 5.844217318 | 0.00017283 | 0.00090608 |
| ARHGAP10 | 2.14422166 | 6.39864333 | 0.000713711 | 0.003062609 |
| INCA1 | 2.143873012 | 2.827801115 | 0.008267144 | 0.024551086 |
| LOC119876531 | 2.143621184 | 3.039111728 | 0.012033976 | 0.033648867 |
| SRM | 2.143555214 | 4.236161105 | 0.000395945 | 0.001848228 |
| GRAMD2A | 2.143058073 | 0.124184462 | 0.011057474 | 0.03137373 |
| ZNF219 | 2.139955397 | 3.533219833 | 0.004424615 | 0.014437719 |
| FOSL1 | 2.13936212 | 2.797426066 | 0.003620277 | 0.012208225 |
| TMEM67 | 2.139329229 | 3.238247929 | 0.00026415 | 0.001302304 |
| ZNF215 | 2.136203286 | 2.673288951 | 0.001299471 | 0.005115046 |
| PGPEP1L | 2.136089691 | 4.216409041 | 0.000219663 | 0.001113888 |
| MARK4 | 2.134778365 | 5.335606339 | 0.00028121 | 0.001375148 |
| AGBL3 | 2.132349201 | 2.281112163 | 0.006540848 | 0.020119187 |
| LOC106559551 | 2.130357908 | -0.410749589 | 0.015221776 | 0.040935792 |
| CHCHD6 | 2.129441836 | 3.583398134 | 0.00025686 | 0.001271763 |
| WDR62 | 2.128235426 | 3.488544926 | 0.001417381 | 0.005489438 |
| FAM174B | 2.128176396 | 3.161353108 | 0.00778656 | 0.023361098 |
| LOC111096427 | 2.128128684 | 2.098000945 | 0.009438801 | 0.027469737 |
| BSG | 2.12631508 | 7.629256298 | 0.000127884 | 0.00070469 |
| ENG | 2.124953562 | 6.927057093 | 0.000201043 | 0.001033003 |
| COMTD1 | 2.124934901 | 3.010262024 | 0.007660374 | 0.023045987 |
| HADH | 2.124077892 | 5.216974462 | 0.000208021 | 0.00106532 |
| BCL9 | 2.120950446 | 6.033854403 | 0.000148461 | 0.000795448 |
| LOC119877747 | 2.120464726 | 0.833660115 | 0.01715704 | 0.04505568 |
| GLB1L | 2.118082213 | 5.285590309 | 9.65E-07 | 9.93E-06 |
| IFT57 | 2.117577227 | 3.367155304 | 0.003516082 | 0.011910858 |
| KLHL21 | 2.116781548 | 4.676859895 | 0.000619807 | 0.002714851 |
| LOC119865386 | 2.115773197 | 1.154708388 | 0.007329282 | 0.022200913 |
| LOC111093763 | 2.113739478 | 3.010743145 | 0.015069481 | 0.04062705 |
| FAM166B | 2.113689632 | 3.593750854 | 0.000364408 | 0.001719907 |
| ASB9 | 2.111247944 | 1.923229171 | 0.01110559 | 0.03148763 |
| PAGR1 | 2.110804144 | 3.851086208 | 0.00130485 | 0.005132488 |
| CACNB3 | 2.110671887 | 5.435450291 | 2.40E-05 | 0.000166126 |
| ATXN7L2 | 2.109719647 | 5.301472009 | 0.002841826 | 0.009953281 |
| KCNQ4 | 2.109570185 | 3.114292627 | 0.007978508 | 0.023863953 |
| GM2A | 2.106565731 | 4.336487632 | 0.006208557 | 0.019250078 |
| CRPPA | 2.104154853 | 3.85625889 | 0.000296434 | 0.001441787 |
| SCN2A | 2.103820281 | 0.285750482 | 0.011256263 | 0.031810795 |

| | | | | |
|---|---|---|---|---|
| POLRMT | 2.101019568 | 5.265791365 | 0.000889748 | 0.003708368 |
| MCRIP1 | 2.097774018 | 5.389694933 | 0.000164222 | 0.000867672 |
| PDE2A | 2.096708912 | 5.465224164 | 0.000757718 | 0.003220635 |
| GDF11 | 2.095338833 | 4.05449442 | 0.00552433 | 0.017435386 |
| PPP1R13B | 2.095304031 | 5.477229979 | 8.22E-05 | 0.000483231 |
| LOC119873619 | 2.094736419 | 2.794135354 | 0.003545169 | 0.011983104 |
| LOC119872199 | 2.093945686 | 2.106098921 | 0.010263084 | 0.029472879 |
| ANOS1 | 2.09264943 | 4.359647401 | 0.004095765 | 0.013526562 |
| INF2 | 2.092304907 | 8.568832629 | 0.006443563 | 0.019887686 |
| LOC119877843 | 2.091495752 | 1.234141163 | 0.004896132 | 0.015720574 |
| ASPHD1 | 2.091183753 | 1.413106792 | 0.012831124 | 0.035547608 |
| POP7 | 2.08935057 | 2.781382818 | 0.000301882 | 0.001464999 |
| TBXA2R | 2.088455741 | 3.655480774 | 0.012095425 | 0.033790172 |
| PPP1R14B | 2.088034285 | 4.07929906 | 0.00010042 | 0.00057425 |
| MAP10 | 2.087663728 | 1.653915385 | 0.013601283 | 0.037295118 |
| RAB11FIP3 | 2.087607674 | 5.83395054 | 5.88E-05 | 0.000361729 |
| SH3RF3 | 2.08594444 | 4.532524871 | 0.001670036 | 0.006314498 |
| MELK | 2.085748954 | 3.102304992 | 0.012981089 | 0.035894251 |
| RFLNB | 2.085691186 | 4.820274208 | 0.000859794 | 0.00359597 |
| VOPP1 | 2.085582692 | 5.000660083 | 0.014910639 | 0.040263926 |
| TSPYL4 | 2.080548349 | 2.715598898 | 0.006341955 | 0.019613184 |
| CC2D1A | 2.080389359 | 5.8375847 | 0.000916555 | 0.003809106 |
| TANC2 | 2.079688114 | 6.414786505 | 0.000511714 | 0.00230282 |
| LOC111098596 | 2.079601294 | 1.843973334 | 0.002087158 | 0.007638464 |
| HIBADH | 2.078319996 | 4.278967856 | 0.000604769 | 0.002657575 |
| DZANK1 | 2.077157972 | 3.445108564 | 0.000216378 | 0.00110032 |
| ARSK | 2.074039077 | 2.455210462 | 0.004730363 | 0.01526294 |
| SRD5A1 | 2.071206476 | 3.306330012 | 0.001217519 | 0.004845261 |
| LOC111093403 | 2.070930197 | 3.564867659 | 0.005175792 | 0.016483914 |
| LOC119876350 | 2.070880983 | 1.341388721 | 0.009857046 | 0.028514651 |
| FMNL2 | 2.068698153 | 7.759874225 | 0.007934192 | 0.023747791 |
| MTHFR | 2.066990142 | 4.638838759 | 5.40E-05 | 0.000335851 |
| SLX4 | 2.066254172 | 4.850294325 | 0.000235751 | 0.001182463 |
| CACNA2D2 | 2.064414306 | 4.24198459 | 0.003212643 | 0.011029242 |
| E2F1 | 2.064049625 | 3.121683723 | 0.003893304 | 0.012989467 |
| LOC102152872 | 2.063817827 | 0.946740711 | 0.003982147 | 0.013243066 |
| TP53I11 | 2.06229928 | 5.50439037 | 0.001135438 | 0.0045681 |
| TTC26 | 2.061013781 | 2.855825869 | 0.004605305 | 0.01493279 |
| KIFAP3 | 2.059803249 | 5.387658549 | 0.000151623 | 0.000809189 |

| | | | | |
|---|---|---|---|---|
| ZNF70 | 2.059639484 | 4.757596612 | 5.42E-06 | 4.45E-05 |
| HYAL2 | 2.05553266 | 4.298702691 | 0.000531552 | 0.002378757 |
| LOC119865100 | 2.055507357 | 3.728748456 | 0.008736595 | 0.025730157 |
| LOC608750 | 2.055408754 | 1.77335302 | 0.001679112 | 0.00634549 |
| BCR | 2.054941598 | 7.304644147 | 0.001558526 | 0.005959412 |
| PHPT1 | 2.053111655 | 4.32996492 | 0.000119293 | 0.000665369 |
| LOC119865988 | 2.052558809 | 1.643320744 | 0.004568335 | 0.014828444 |
| DRAP1 | 2.051583405 | 5.303281258 | 0.001209752 | 0.004819662 |
| HCN3 | 2.048872902 | 2.504445378 | 0.014529746 | 0.039446723 |
| TOM1L2 | 2.04845656 | 7.114006611 | 6.67E-05 | 0.000402276 |
| SAMD11 | 2.047930116 | 5.111312442 | 0.000180018 | 0.00093832 |
| TMEM88 | 2.047257704 | 3.270911584 | 0.007351511 | 0.022258912 |
| GNAO1 | 2.047149812 | 4.646558953 | 0.006050412 | 0.018848695 |
| LOC111091843 | 2.046374674 | 1.827915016 | 0.002714716 | 0.009562172 |
| SLC22A23 | 2.046265238 | 3.518654122 | 0.00635062 | 0.019634379 |
| PACS2 | 2.043583321 | 7.627999946 | 0.005067432 | 0.016196202 |
| C2 | 2.042547962 | 4.94316981 | 0.00078415 | 0.003317371 |
| ST3GAL3 | 2.041981754 | 3.764977971 | 0.000567597 | 0.002516166 |
| LOC610677 | 2.040296148 | 3.595880263 | 0.002039169 | 0.007494877 |
| PTMS | 2.039908104 | 8.113756475 | 0.000239967 | 0.001201059 |
| IL6ST | 2.037559359 | 8.356410187 | 0.000239901 | 0.001201057 |
| PIR | 2.032218918 | 4.26818981 | 0.016954645 | 0.044605246 |
| LOC119872476 | 2.032076389 | 4.001927978 | 0.002990045 | 0.010386385 |
| NAT14 | 2.031683032 | 2.537624414 | 0.011772416 | 0.033036847 |
| WNK4 | 2.030974226 | 4.90878617 | 0.001197051 | 0.004775345 |
| LOC119876495 | 2.030609527 | 2.859775622 | 0.00267524 | 0.009439999 |
| ST3GAL4 | 2.029846031 | 5.881334766 | 0.001410773 | 0.005468809 |
| PLAT | 2.029338799 | 5.103744342 | 0.010391978 | 0.029775957 |
| TRIP6 | 2.028838111 | 4.850156014 | 8.12E-05 | 0.000478068 |
| PRDM11 | 2.027417801 | 4.609649902 | 4.57E-05 | 0.000290342 |
| MAGED2 | 2.026779793 | 5.521285403 | 4.53E-05 | 0.000288091 |
| CMTM3 | 2.026166652 | 5.816037855 | 8.19E-05 | 0.000481779 |
| BATF2 | 2.02584197 | 4.246408789 | 0.000141903 | 0.000766565 |
| SELENON | 2.025769459 | 7.483518829 | 0.00019626 | 0.001010344 |
| PDE4A | 2.025414942 | 5.622736564 | 0.000544728 | 0.002428694 |
| GTPBP3 | 2.024851266 | 3.97196034 | 0.000703851 | 0.003024899 |
| CRLF1 | 2.024176751 | 4.649494385 | 0.014161567 | 0.038601892 |
| LOC119865347 | 2.024130772 | 2.513690791 | 0.000249888 | 0.001242159 |
| EXOC6B | 2.02386645 | 7.175279743 | 0.001194854 | 0.004768194 |

| | | | | |
|---|---|---|---|---|
| ZNF250 | 2.021835383 | 3.95836835 | 0.000274088 | 0.001344265 |
| CEP112 | 2.021307725 | 5.291241982 | 0.007246166 | 0.021979368 |
| NENF | 2.020004291 | 4.268582325 | 6.80E-05 | 0.000408809 |
| LOC119864957 | 2.019323971 | 1.675766705 | 0.009946467 | 0.028735187 |
| CHST7 | 2.018594933 | 3.356747563 | 0.002019728 | 0.007434369 |
| LOC111097293 | 2.018167088 | 4.443637475 | 0.008950765 | 0.026264546 |
| LAMB2 | 2.016527033 | 8.28119087 | 5.17E-05 | 0.000323576 |
| SOX12 | 2.016283219 | 5.280650226 | 0.001587106 | 0.006048415 |
| SGF29 | 2.01503174 | 3.742205914 | 0.001183439 | 0.004733974 |
| LYPLAL1 | 2.014529168 | 2.449241585 | 0.005837282 | 0.018279356 |
| LOC111096175 | 2.014192147 | 2.672852326 | 0.01016858 | 0.02926353 |
| MRAS | 2.01030344 | 6.407636283 | 0.005945268 | 0.018563866 |
| MYRF | 2.010225228 | 5.771623036 | 0.003780745 | 0.012670493 |
| MED25 | 2.009708918 | 6.304408612 | 0.002224371 | 0.008065717 |
| KIF13A | 2.006966286 | 6.457918535 | 3.51E-05 | 0.000230498 |
| MIF | 2.005889033 | 5.77604494 | 0.000489615 | 0.002217871 |
| FNDC5 | 2.002612048 | 2.700303814 | 0.016037792 | 0.042680139 |
| LOC100687594 | 2.00229168 | 6.742285085 | 0.003273755 | 0.011208861 |
| ZFP36L1 | 2.001357673 | 8.879716992 | 6.34E-05 | 0.000385484 |
| LRRC75A | 2.00113915 | 4.560836363 | 0.001110662 | 0.004479609 |
| IL20RB | 1.999605829 | -0.233625805 | 0.018712253 | 0.048268347 |
| LOC102154540 | 1.999089423 | 2.289312837 | 0.004942114 | 0.015854121 |
| ABCB6 | 1.998779143 | 4.710878892 | 0.000465313 | 0.00211953 |
| SLC18B1 | 1.998175019 | 4.175910056 | 5.11E-05 | 0.000320085 |
| ISOC2 | 1.997927524 | 2.892725303 | 0.001296484 | 0.00510607 |
| PYCR3 | 1.99772099 | 3.74819183 | 0.000275878 | 0.001350902 |
| LOC111096398 | 1.996206682 | 2.426322571 | 0.019197002 | 0.049319182 |
| FKBP2 | 1.995566954 | 3.788398517 | 0.001466645 | 0.00565497 |
| MMAB | 1.995022032 | 2.706003908 | 0.000436994 | 0.002011677 |
| AVEN | 1.994986381 | 3.327276037 | 0.001068913 | 0.004335058 |
| LOC481722 | 1.994890982 | 7.216817993 | 0.001000442 | 0.004097237 |
| HCFC1R1 | 1.993350508 | 5.321135989 | 0.000791637 | 0.003345777 |
| LOC119877541 | 1.990836061 | 5.771298488 | 0.004682571 | 0.015138032 |
| CENPM | 1.987948087 | 1.946467832 | 0.011058179 | 0.03137373 |
| LOC106557691 | 1.987183692 | 0.834348371 | 0.001610191 | 0.006121325 |
| RNF215 | 1.984884255 | 4.189728579 | 9.49E-05 | 0.000546558 |
| TMPRSS5 | 1.982086993 | 1.956039048 | 0.004876354 | 0.01566171 |
| GPHN | 1.9820001 | 4.635779148 | 2.47E-05 | 0.000170202 |
| BRSK1 | 1.981860882 | 4.701441481 | 0.001172964 | 0.004695541 |

| | | | | |
|---|---|---|---|---|
| POLR3D | 1.981321128 | 4.792579943 | 4.32E-05 | 0.000275882 |
| PRNP | 1.981147814 | 6.049966369 | 0.005939481 | 0.018550642 |
| TEP1 | 1.980466914 | 6.802419265 | 0.002799051 | 0.009825695 |
| LOC119878548 | 1.979564166 | 2.58460981 | 0.00320718 | 0.011012231 |
| N4BP3 | 1.979265483 | 3.913609567 | 0.007082899 | 0.021544936 |
| TTC3 | 1.976853351 | 8.418598101 | 0.000184668 | 0.000958408 |
| TJP2 | 1.976426382 | 5.990302474 | 0.00025698 | 0.001271871 |
| LOC102152612 | 1.975896603 | 0.983520519 | 0.01018138 | 0.029296475 |
| IER5L | 1.975084179 | 5.456018918 | 0.004028194 | 0.013363395 |
| OXLD1 | 1.974481298 | 2.371221372 | 0.002386653 | 0.008548326 |
| TYRO3 | 1.97428957 | 3.623210902 | 0.003256863 | 0.011161605 |
| BTBD2 | 1.974209023 | 6.322936912 | 0.000861251 | 0.003601368 |
| NUDT6 | 1.974107687 | 4.223090229 | 0.001090014 | 0.004406635 |
| ASRGL1 | 1.972739602 | 3.925462654 | 0.000284574 | 0.001389093 |
| HDHD5 | 1.97193404 | 3.658038548 | 0.001384602 | 0.005391389 |
| LOC119865725 | 1.97170028 | 1.480259274 | 0.010789344 | 0.030731694 |
| LOC119864272 | 1.971304703 | 1.855885614 | 0.018195509 | 0.047182493 |
| MOSMO | 1.971097831 | 3.401012389 | 0.000378076 | 0.001776293 |
| BTBD3 | 1.970297199 | 6.028278308 | 0.000581196 | 0.002566474 |
| PROSER3 | 1.968818234 | 5.092595433 | 0.002863163 | 0.010018296 |
| LOC119872280 | 1.96581452 | 2.757342184 | 0.001295227 | 0.005102973 |
| SLC9A5 | 1.965783001 | 4.135423383 | 0.000273131 | 0.001340482 |
| LAMA3 | 1.964912827 | 4.945427728 | 0.017903602 | 0.046592793 |
| LOC111095514 | 1.964146989 | 1.497098333 | 0.012450195 | 0.034638393 |
| HASPIN | 1.963545098 | 3.50396228 | 0.012107321 | 0.033805973 |
| LOC611670 | 1.961457983 | 3.611329033 | 0.000279599 | 0.001367581 |
| TP53INP2 | 1.957760657 | 6.006981189 | 0.000320323 | 0.001540693 |
| ZNF358 | 1.957261296 | 5.409889227 | 0.002222787 | 0.00806311 |
| LOC478038 | 1.955513616 | 8.552050921 | 3.68E-05 | 0.000240461 |
| PKN3 | 1.954502727 | 4.79972097 | 0.000254966 | 0.001263635 |
| MIR8867 | 1.953580948 | 1.209181431 | 0.013511762 | 0.037077795 |
| THPO | 1.95299588 | 2.146665359 | 0.007106472 | 0.021607543 |
| HOMER3 | 1.951687561 | 4.666920575 | 0.001404232 | 0.005450202 |
| ACSS3 | 1.950688917 | 5.003769372 | 0.004942966 | 0.015854507 |
| LOC119870761 | 1.950503059 | 2.869583043 | 0.001126303 | 0.004535557 |
| LOC106560149 | 1.949495906 | 0.267636096 | 0.01943177 | 0.049833719 |
| P4HB | 1.948897068 | 8.542473404 | 0.000419183 | 0.001943722 |
| OR10V4 | 1.948542323 | 2.293737019 | 0.008282499 | 0.024587511 |
| ARHGEF40 | 1.946770693 | 6.759908013 | 0.000325772 | 0.001561702 |

| | | | | |
|---|---|---|---|---|
| NCKAP1 | 1.946420838 | 7.279171947 | 0.004076461 | 0.013480187 |
| NUDT7 | 1.945938467 | 2.198977243 | 0.002143552 | 0.007815813 |
| LOC119866908 | 1.94564143 | 1.946419125 | 0.008456653 | 0.025031554 |
| CRIM1 | 1.944588293 | 7.441746461 | 0.001805245 | 0.006752355 |
| SETD7 | 1.94421052 | 7.47061594 | 6.08E-05 | 0.000372249 |
| GNA12 | 1.943903936 | 5.485013197 | 9.58E-06 | 7.33E-05 |
| PCGF1 | 1.943732006 | 4.868292821 | 0.00412149 | 0.01360418 |
| HRH1 | 1.943687554 | 3.209469293 | 0.003502821 | 0.011874962 |
| PLPP1 | 1.941868023 | 6.711017195 | 0.000716626 | 0.003071277 |
| TOMM40 | 1.941342839 | 4.570617251 | 0.000191046 | 0.000986079 |
| TPM1 | 1.940178385 | 7.98946788 | 0.003301239 | 0.011279797 |
| NRIP1 | 1.939981532 | 6.607081491 | 0.001995816 | 0.007357597 |
| SNAPC2 | 1.939450035 | 3.804302179 | 0.00565906 | 0.017798405 |
| PRKRA | 1.938331873 | 3.125322358 | 0.002013062 | 0.007413611 |
| MRTFB | 1.937859864 | 7.465769855 | 2.22E-05 | 0.000154656 |
| LDHD | 1.937771202 | 2.505098902 | 0.008033536 | 0.023995429 |
| VWA5B2 | 1.93658602 | 3.995286278 | 0.000140967 | 0.00076227 |
| DOCK1 | 1.936354955 | 8.065476985 | 0.000713756 | 0.003062609 |
| LOC111094331 | 1.935922052 | 0.971295042 | 0.005221615 | 0.016593558 |
| EFNA4 | 1.935384191 | 2.989135648 | 0.002067429 | 0.007579056 |
| ANKRD13C | 1.934193598 | 6.198295591 | 0.000175169 | 0.000916911 |
| LOC119869720 | 1.933849412 | 4.275558559 | 0.000642934 | 0.002803684 |
| LOC111098591 | 1.932471386 | 4.125162342 | 0.000355981 | 0.001684902 |
| CHST10 | 1.932337261 | 5.883415997 | 0.000429498 | 0.001982645 |
| MB21D2 | 1.931907825 | 2.592783101 | 0.014124168 | 0.038529023 |
| TCHP | 1.931866492 | 4.583561327 | 0.003509431 | 0.011893715 |
| LOC111091842 | 1.929851569 | 2.210795012 | 0.009359532 | 0.027290346 |
| MACROD1 | 1.929251648 | 2.854505078 | 0.005211746 | 0.016569483 |
| CLCN6 | 1.928363067 | 6.054474535 | 0.001006557 | 0.004119169 |
| MUC5B | 1.926103514 | 2.25323842 | 0.011099303 | 0.03147804 |
| SLC29A4 | 1.924412952 | 2.77317922 | 0.006599135 | 0.020275453 |
| LOC119871488 | 1.923740239 | 4.029849685 | 0.001131476 | 0.004553848 |
| ZBTB20 | 1.923084962 | 7.629725949 | 0.000960622 | 0.003956589 |
| LOC119863930 | 1.922315014 | 2.349966049 | 0.00483352 | 0.015544869 |
| LOC606845 | 1.921898978 | 5.050558572 | 0.000378756 | 0.001778715 |
| HOGA1 | 1.921544354 | 2.145411779 | 0.002261407 | 0.00817403 |
| ATP6V0E2 | 1.920926935 | 3.257562101 | 0.001521409 | 0.005837052 |
| MLLT1 | 1.919905011 | 6.829801936 | 0.001765495 | 0.006613131 |
| KIFC1 | 1.919665036 | 4.022086321 | 0.000389271 | 0.001820994 |

| | | | | |
|---|---|---|---|---|
| RNF32 | 1.919516007 | 1.839779908 | 0.015026738 | 0.040526942 |
| SFXN2 | 1.91944633 | 3.999261034 | 0.001603548 | 0.00609928 |
| ICMT | 1.918521363 | 4.183132528 | 0.001231952 | 0.004888337 |
| LOC111093714 | 1.91830871 | 3.47663045 | 0.001358617 | 0.005311201 |
| MTHFD1L | 1.917754647 | 5.681419014 | 0.00388487 | 0.012965321 |
| PEX14 | 1.916960589 | 5.553551485 | 0.002863937 | 0.010019387 |
| CNTROB | 1.916249072 | 5.188726933 | 0.001559111 | 0.005960039 |
| DOK4 | 1.915127822 | 4.66854623 | 0.000655282 | 0.002849505 |
| LRRC36 | 1.914103192 | 3.648808917 | 0.000449215 | 0.002057033 |
| S1PR2 | 1.913722973 | 4.366133316 | 0.000508768 | 0.002292523 |
| ZNF428 | 1.911822191 | 3.612999045 | 0.000232879 | 0.001169684 |
| UBXN2A | 1.911471989 | 6.258837614 | 0.000314729 | 0.001518842 |
| PGGHG | 1.909142516 | 5.102131238 | 0.00102272 | 0.004169564 |
| EPN2 | 1.907749091 | 6.673290436 | 0.001078806 | 0.004371089 |
| SLC12A4 | 1.907282018 | 6.50833125 | 2.29E-05 | 0.000158767 |
| GRIK2 | 1.906392422 | 4.091422549 | 0.012598008 | 0.034969191 |
| LOC119869290 | 1.906166731 | 2.275535744 | 0.005053563 | 0.01615664 |
| C1H9orf3 | 1.90524577 | 7.221273492 | 3.99E-05 | 0.000257531 |
| LOC102157082 | 1.902025558 | 2.381053841 | 0.009131277 | 0.02672187 |
| LYRM9 | 1.901637889 | 1.81101084 | 0.007571015 | 0.022821504 |
| RGS16 | 1.900552199 | 3.368557033 | 0.002258721 | 0.00816568 |
| LIX1L | 1.900546746 | 4.440246605 | 0.000296089 | 0.001440433 |
| ZNF213 | 1.898895005 | 3.221988254 | 0.003717013 | 0.012489741 |
| CACNB2 | 1.898764928 | 3.728090242 | 0.003978525 | 0.013233053 |
| NEK11 | 1.898591052 | 2.073794335 | 0.010012592 | 0.028893047 |
| LOC119873800 | 1.897885877 | 4.227544064 | 0.002329567 | 0.00837292 |
| MKS1 | 1.897745795 | 3.705624134 | 0.000220794 | 0.001118056 |
| ZKSCAN2 | 1.897005466 | 4.494298918 | 0.000247451 | 0.001232027 |
| LOC102157251 | 1.896390131 | 5.385127451 | 0.001643276 | 0.006226842 |
| ZNF775 | 1.896123225 | 2.276809199 | 0.008323319 | 0.024690841 |
| ACYP2 | 1.892968504 | 2.803903132 | 0.010950649 | 0.031101221 |
| LOC119872071 | 1.892444771 | 3.493444158 | 0.003984479 | 0.013246757 |
| SYT11 | 1.89160957 | 5.664479775 | 0.00175516 | 0.006580104 |
| ECE1 | 1.889324697 | 7.395040719 | 6.86E-05 | 0.00041215 |
| ARL6IP4 | 1.889277367 | 3.977806833 | 0.002913339 | 0.01015615 |
| FAM189B | 1.887856823 | 6.913323999 | 0.001320446 | 0.005184425 |
| ZMIZ2 | 1.887655045 | 6.843984242 | 0.00407087 | 0.01346375 |
| BMP4 | 1.886780467 | 2.873977871 | 0.008383515 | 0.024849003 |
| EPB41L5 | 1.885744587 | 2.651601542 | 0.0070237 | 0.021400004 |

| | | | | |
|---|---|---|---|---|
| LOC106558083 | 1.884704085 | 1.139289261 | 0.003732664 | 0.012528733 |
| MICU3 | 1.88421595 | 3.229595541 | 0.001585876 | 0.00604526 |
| LOC119865046 | 1.884211188 | 2.438517925 | 0.012979039 | 0.035893161 |
| GRAMD1C | 1.883212439 | 1.147038322 | 0.019179271 | 0.049285314 |
| SPNS2 | 1.882986976 | 4.424979355 | 0.004236909 | 0.013900472 |
| ZHX3 | 1.881214084 | 7.14337805 | 0.00140206 | 0.005444694 |
| ZNF311 | 1.879990659 | 4.480250988 | 0.000926344 | 0.00384242 |
| KATNAL1 | 1.879823935 | 4.802338165 | 0.006374539 | 0.019702708 |
| TMEM63B | 1.878748646 | 6.148117792 | 0.000685316 | 0.002958146 |
| FBXL6 | 1.876980664 | 3.642150828 | 0.002396615 | 0.008582589 |
| NIBAN2 | 1.87613996 | 7.099391309 | 0.000731291 | 0.003126089 |
| LOC119873220 | 1.875786538 | 5.609122348 | 0.004955058 | 0.01587919 |
| TMEM37 | 1.875152017 | 2.385877347 | 0.014285279 | 0.038895083 |
| DTD2 | 1.874657339 | 2.633251672 | 0.004001564 | 0.013293361 |
| LOC119866556 | 1.873908385 | 1.942960726 | 0.00315271 | 0.010864794 |
| LOC111096251 | 1.873804101 | 5.644440321 | 3.56E-05 | 0.00023396 |
| AGBL4 | 1.873703269 | 3.303832209 | 0.006933257 | 0.02116103 |
| ACKR4 | 1.872961808 | 2.368405234 | 0.017081208 | 0.044883732 |
| BRMS1L | 1.872818683 | 3.895552741 | 0.000216068 | 0.00109926 |
| LOC111092216 | 1.872694127 | 2.757675113 | 0.005070629 | 0.016201644 |
| MOK | 1.872156827 | 3.983498386 | 0.014680333 | 0.03977583 |
| CLDN15 | 1.871095603 | 2.954084023 | 0.015208522 | 0.040905223 |
| CRYBG3 | 1.870766083 | 7.391378642 | 0.001717125 | 0.006457619 |
| SLC22A15 | 1.870621015 | 4.3789031 | 0.002238933 | 0.008106315 |
| TSPAN4 | 1.869367051 | 4.553284989 | 0.000188815 | 0.00097619 |
| NINL | 1.86904285 | 5.243679869 | 0.000472142 | 0.002145676 |
| ODF2 | 1.866450503 | 5.521964201 | 0.001023501 | 0.004171965 |
| ITGAV | 1.86603746 | 8.372897923 | 0.003054693 | 0.010572399 |
| MORC2 | 1.865660751 | 6.271311694 | 0.000258599 | 0.001277258 |
| SLC2A4RG | 1.864569702 | 6.159609177 | 0.002669644 | 0.009425635 |
| PTER | 1.86367594 | 1.909989174 | 0.003895379 | 0.012993958 |
| CMKLR1 | 1.863474235 | 5.56695956 | 0.003393512 | 0.011549557 |
| SLC26A2 | 1.863158654 | 5.546476812 | 0.00092366 | 0.003832756 |
| AK8 | 1.861813797 | 1.370724753 | 0.014160612 | 0.038601892 |
| IQCE | 1.860957995 | 4.923972346 | 0.001810189 | 0.006764177 |
| NOS2 | 1.859175938 | 3.053830463 | 0.00350945 | 0.011893715 |
| PTPN13 | 1.859055116 | 6.413257074 | 0.003790196 | 0.012698242 |
| CTNNB1 | 1.858934099 | 8.441534893 | 0.001169772 | 0.004684491 |
| STK25 | 1.857706497 | 5.959209009 | 0.001235037 | 0.004897886 |

| | | | | |
|---|---|---|---|---|
| TTLL1 | 1.857571026 | 3.667526066 | 0.001400152 | 0.00543934 |
| RPP38 | 1.856905697 | 3.541658037 | 0.000551043 | 0.002452813 |
| DPY19L3 | 1.856350626 | 5.598824696 | 0.000134028 | 0.000731316 |
| DNMT3B | 1.856146489 | 3.509490211 | 0.002368119 | 0.008488958 |
| LOC111091205 | 1.855560593 | 3.789233941 | 0.009410621 | 0.027409444 |
| WIPI1 | 1.851332243 | 5.615074112 | 0.00146747 | 0.005656139 |
| LOC111092837 | 1.849456662 | 4.178959798 | 0.000258341 | 0.001276569 |
| CDC42EP2 | 1.848619838 | 5.665660163 | 0.00046171 | 0.002106218 |
| MINK1 | 1.847721277 | 7.596892126 | 0.002345304 | 0.008418311 |
| CFDP1 | 1.847020076 | 5.106621996 | 0.002699815 | 0.009514323 |
| CCHCR1 | 1.843446175 | 5.76324932 | 0.002265741 | 0.008188327 |
| ZNF404 | 1.842023134 | 2.483795878 | 0.001682396 | 0.006356795 |
| DOP1B | 1.841664109 | 5.772189595 | 0.007379602 | 0.022331487 |
| LOC119872567 | 1.841231349 | 6.408613133 | 0.002127267 | 0.00776558 |
| LOC111093101 | 1.840546535 | 4.24585561 | 0.000971694 | 0.003993085 |
| EXT1 | 1.840408935 | 6.548712675 | 0.000526758 | 0.002362671 |
| ME3 | 1.840124143 | 4.637358186 | 0.002217165 | 0.00804632 |
| ZFHX3 | 1.839514931 | 8.324906759 | 0.001186714 | 0.004743574 |
| RND1 | 1.838394383 | 2.770362323 | 0.015072053 | 0.040628931 |
| TASP1 | 1.837954137 | 4.327182544 | 0.00014844 | 0.000795448 |
| FLNB | 1.837822307 | 8.270623147 | 0.000387415 | 0.001812705 |
| MARCHF9 | 1.83773261 | 2.258019733 | 0.010389737 | 0.029773471 |
| EARS2 | 1.836617716 | 3.246478473 | 0.003379465 | 0.011508979 |
| PJA1 | 1.835926314 | 4.109680378 | 0.001088585 | 0.004402481 |
| LOC102153308 | 1.835511104 | 1.877915928 | 0.010013181 | 0.028893047 |
| DBNDD2 | 1.83494035 | 4.351295183 | 0.000330696 | 0.001581457 |
| XYLB | 1.831614341 | 4.168407441 | 0.001041363 | 0.004236009 |
| CLIP4 | 1.831495032 | 3.498760806 | 0.00386259 | 0.012900895 |
| RALA | 1.830231237 | 5.719953591 | 0.001632587 | 0.006191499 |
| BCAP31 | 1.828823631 | 5.532498952 | 4.13E-05 | 0.000265692 |
| SCRIB | 1.827200019 | 7.245938036 | 0.001595489 | 0.006073958 |
| ATXN10 | 1.82482221 | 5.543657769 | 7.06E-05 | 0.000423404 |
| PIAS3 | 1.824795928 | 5.312928807 | 0.000464467 | 0.002117458 |
| RAB26 | 1.823063791 | 2.861373575 | 0.005686002 | 0.017880545 |
| RHBDL1 | 1.82305588 | 2.596814778 | 0.0027456 | 0.009661535 |
| SLC38A10 | 1.822617079 | 6.793187044 | 0.000682266 | 0.002947919 |
| ARHGAP42 | 1.821410752 | 7.053243292 | 0.001637072 | 0.006206099 |
| ZNF461 | 1.821149134 | 2.905629563 | 0.001832632 | 0.006838609 |
| LOC119865224 | 1.820702345 | 3.986558389 | 0.003722392 | 0.012502 |

| | | | | |
|---|---|---|---|---|
| PELP1 | 1.820439825 | 5.284746282 | 0.00075518 | 0.003212997 |
| PTDSS2 | 1.819417833 | 4.980835796 | 0.001426022 | 0.005516005 |
| ADM5 | 1.819075405 | 3.399305867 | 0.017153462 | 0.045051742 |
| LOC111092530 | 1.814802881 | 2.443154539 | 0.015527737 | 0.041614028 |
| PLIN3 | 1.812816709 | 4.932647605 | 0.004631271 | 0.015003517 |
| BLVRA | 1.811930759 | 4.141133764 | 0.007309907 | 0.022148416 |
| STEAP3 | 1.811296724 | 4.375642548 | 0.005428595 | 0.017180773 |
| PLPP7 | 1.810700564 | 3.057781631 | 0.014097269 | 0.03848471 |
| LOC102152881 | 1.809925293 | 2.16435561 | 0.016428588 | 0.043481513 |
| FAM3A | 1.809634266 | 4.610855367 | 0.001105627 | 0.004463077 |
| RCAN2 | 1.809629411 | 4.696101171 | 0.001341034 | 0.005250989 |
| LOC476816 | 1.808169124 | 4.55673055 | 0.014788704 | 0.04000444 |
| AJUBA | 1.807118906 | 5.30536234 | 3.86E-05 | 0.000250701 |
| FAXDC2 | 1.805403893 | 6.228304394 | 0.000938693 | 0.003882498 |
| CCDC14 | 1.805045673 | 4.51887204 | 0.002010436 | 0.007405197 |
| ARL3 | 1.804565425 | 2.66524042 | 0.009004372 | 0.026403971 |
| BCL7C | 1.804237491 | 3.875387531 | 0.003046874 | 0.010550392 |
| ALG12 | 1.804081002 | 4.3095302 | 0.000584257 | 0.002577368 |
| LOC119873573 | 1.803980486 | 7.033118078 | 0.001837357 | 0.006852699 |
| LOC111097287 | 1.803436332 | 0.089757081 | 0.014108735 | 0.038501463 |
| ERCC8 | 1.803385396 | 2.192151534 | 0.007631553 | 0.02297472 |
| ZNF229 | 1.802002253 | 2.309319527 | 0.003192197 | 0.010973494 |
| SPTAN1 | 1.800944138 | 9.983581287 | 0.003075229 | 0.010632082 |
| IZUMO1 | 1.8007466 | 2.712706206 | 0.007067619 | 0.021510535 |
| PIH1D1 | 1.800390627 | 4.337282204 | 0.00285381 | 0.009992025 |
| MORN4 | 1.798062834 | 2.822540304 | 0.001407955 | 0.005460678 |
| ABHD10 | 1.796048346 | 3.185061228 | 0.001342863 | 0.005255302 |
| LOC111096317 | 1.795701417 | 3.639347123 | 0.001296396 | 0.00510607 |
| IGF1R | 1.795161356 | 7.358905676 | 0.000812728 | 0.003422408 |
| GMPPA | 1.795106767 | 5.953188353 | 0.00031285 | 0.001511458 |
| MAFB | 1.795001484 | 8.667848517 | 0.013570602 | 0.037225118 |
| LOC111094502 | 1.794847979 | 1.657695298 | 0.016336039 | 0.043314584 |
| ITPKC | 1.794402599 | 4.395013056 | 0.000271973 | 0.0013351 |
| LOC102152738 | 1.792386243 | 1.806373623 | 0.003195419 | 0.010980548 |
| LOC119876626 | 1.790819623 | 3.015851555 | 0.004511308 | 0.014665152 |
| LOC111098324 | 1.790716301 | 2.820928334 | 0.013361476 | 0.036758561 |
| SPART | 1.788441738 | 5.546618806 | 0.000129273 | 0.000710198 |
| CCDC51 | 1.785380173 | 3.9006578 | 0.001747866 | 0.006555027 |
| NR1H3 | 1.783438374 | 4.575938498 | 0.000454656 | 0.002078029 |

| | | | | |
|---|---|---|---|---|
| CAMKK1 | 1.782201463 | 4.080468323 | 0.000185374 | 0.00096161 |
| RFNG | 1.78216799 | 5.397242464 | 0.00163418 | 0.006196219 |
| ARHGEF4 | 1.780241519 | 3.946660817 | 0.004314063 | 0.014113009 |
| GATB | 1.77862336 | 4.051060159 | 0.009889337 | 0.028592795 |
| ACTN4 | 1.778376654 | 9.095532781 | 0.000466773 | 0.002125286 |
| BPHL | 1.777587082 | 3.333676749 | 0.001058725 | 0.004298565 |
| KIAA0825 | 1.777307039 | 3.675584314 | 0.000637912 | 0.002783439 |
| DHODH | 1.775935073 | 2.334395607 | 0.002461641 | 0.008780635 |
| LOC119868047 | 1.77515683 | 3.518285945 | 0.003418316 | 0.011619379 |
| LOC119877936 | 1.774639925 | 2.177848221 | 0.010091891 | 0.029089182 |
| PAFAH1B3 | 1.773963442 | 4.524321118 | 0.004456995 | 0.01451705 |
| ANKRD13B | 1.772657284 | 3.085412927 | 0.011995316 | 0.033545094 |
| ARF5 | 1.772656676 | 3.940278599 | 0.001131144 | 0.004553357 |
| RTN4 | 1.772238447 | 7.847797514 | 0.003130169 | 0.010797416 |
| DTX3 | 1.771329029 | 7.138724038 | 0.002082701 | 0.00762344 |
| PDLIM5 | 1.771249127 | 6.824788961 | 0.00163265 | 0.006191499 |
| LBX2 | 1.770746571 | 3.63423037 | 0.016530928 | 0.04369205 |
| GORAB | 1.766614096 | 5.192806904 | 0.000519544 | 0.002334177 |
| SCAMP4 | 1.765147039 | 5.759603654 | 0.000148972 | 0.00079681 |
| TPRG1L | 1.764925123 | 5.544656374 | 0.000234727 | 0.001178068 |
| SERHL2 | 1.760462909 | 3.807859663 | 0.00147776 | 0.00568872 |
| TRIO | 1.760336001 | 8.757044693 | 0.00609166 | 0.018941829 |
| MKNK2 | 1.759966804 | 6.777480204 | 0.005872169 | 0.018366032 |
| C19H2orf76 | 1.759211607 | 2.018394085 | 0.003757704 | 0.012601069 |
| MRPS15 | 1.758451071 | 4.473687774 | 0.003403802 | 0.011579124 |
| GRWD1 | 1.757809477 | 2.82048807 | 0.012579492 | 0.034935331 |
| ADCK5 | 1.757282 | 4.019977031 | 0.000958607 | 0.003949787 |
| ELMO3 | 1.755792558 | 2.89007469 | 0.013374681 | 0.036790135 |
| POLM | 1.753950878 | 3.211573039 | 0.004059874 | 0.013434612 |
| CIAO3 | 1.753677596 | 4.833254387 | 0.001592254 | 0.006062706 |
| MAN2C1 | 1.751443283 | 5.357901809 | 0.001049984 | 0.004266275 |
| RHOB | 1.750677216 | 6.5259675 | 0.001713344 | 0.006445637 |
| LOC102156128 | 1.748475595 | 3.416695812 | 0.004904251 | 0.015739645 |
| LOC111097117 | 1.746521634 | 0.138109486 | 0.011653381 | 0.032749446 |
| SREBF1 | 1.745537906 | 7.010385431 | 0.004229217 | 0.013885746 |
| ZNF35 | 1.742608606 | 5.201388142 | 0.00077948 | 0.003302123 |
| DNAJC18 | 1.742286811 | 5.118448172 | 0.000478037 | 0.002169737 |
| GOLGA2 | 1.741192467 | 6.865019145 | 0.001714628 | 0.006449347 |
| ARFGAP1 | 1.739822501 | 5.774907731 | 0.001026397 | 0.004181101 |

| | | | | |
|---|---|---|---|---|
| PRICKLE4 | 1.739594556 | 4.131531783 | 0.015908974 | 0.04241056 |
| GPT | 1.739049734 | 3.773192851 | 0.012064167 | 0.033711538 |
| RUVBL2 | 1.738628159 | 4.359753204 | 0.000196553 | 0.00101161 |
| ISCA2 | 1.736781721 | 5.76860166 | 0.000186517 | 0.000966613 |
| MZT2B | 1.734399309 | 2.872499307 | 0.008968809 | 0.026313928 |
| PRRT1 | 1.734242831 | 2.323376209 | 0.016558684 | 0.043754739 |
| IRF2BP1 | 1.733430288 | 4.021758843 | 0.001147485 | 0.004608867 |
| DNAJC15 | 1.73333927 | 2.711483606 | 0.008986663 | 0.026359174 |
| HS6ST1 | 1.733109218 | 6.186640955 | 0.009845947 | 0.028490712 |
| GABBR1 | 1.731418034 | 5.99619911 | 0.000525948 | 0.002359526 |
| RRP9 | 1.731186889 | 3.459929697 | 0.004589365 | 0.014887786 |
| CHAF1B | 1.729654463 | 3.733989679 | 0.009327611 | 0.027211916 |
| INTU | 1.727467552 | 3.331790675 | 0.004307554 | 0.014098099 |
| AACS | 1.726659219 | 3.056349795 | 0.002351123 | 0.008435009 |
| LAPTM4A | 1.726201663 | 6.956072247 | 0.004683813 | 0.015139308 |
| DHCR7 | 1.723153599 | 2.986224033 | 0.012050659 | 0.033678134 |
| LOC119871908 | 1.720086356 | 3.298770267 | 0.016414693 | 0.043459032 |
| SNX9 | 1.719565922 | 6.721595984 | 0.002584458 | 0.00915843 |
| RITA1 | 1.718220369 | 4.01666208 | 0.002122616 | 0.007751215 |
| SLC38A7 | 1.717328807 | 4.20943527 | 0.000511481 | 0.00230225 |
| TBC1D19 | 1.717007064 | 3.874858005 | 0.000248396 | 0.001236448 |
| JOSD2 | 1.715805969 | 3.244570058 | 0.002684952 | 0.009469646 |
| C20H19orf25 | 1.71574749 | 4.476728179 | 0.002857099 | 0.010001924 |
| SLC39A1 | 1.715714871 | 5.064832068 | 0.001376095 | 0.005364048 |
| KDM4B | 1.715156279 | 7.198748096 | 0.01646983 | 0.043562439 |
| GAS8 | 1.714739725 | 3.920364017 | 0.01857736 | 0.047988929 |
| C4H1orf198 | 1.714488047 | 6.381744127 | 0.013142156 | 0.036260974 |
| ENDOV | 1.714087543 | 3.431142081 | 0.003197445 | 0.01098577 |
| KLHDC1 | 1.712194462 | 3.097813881 | 0.001697735 | 0.006401358 |
| FAM135A | 1.708714808 | 5.04532104 | 0.001469838 | 0.005661241 |
| MAPK8IP3 | 1.707865987 | 7.272532748 | 0.004448216 | 0.01449935 |
| TNK1 | 1.707102925 | 3.583351635 | 0.01257439 | 0.034925639 |
| TMEM52 | 1.704728719 | 3.000577291 | 0.017180169 | 0.045083642 |
| VPS37B | 1.703621073 | 5.946245194 | 0.008144912 | 0.024254558 |
| TMEM201 | 1.703440682 | 4.73417442 | 0.002548451 | 0.009044143 |
| PRDM5 | 1.703169438 | 4.31989755 | 0.001372568 | 0.005354834 |
| LOC609068 | 1.70300058 | 4.044436078 | 0.002793876 | 0.009813885 |
| NUCKS1 | 1.702909037 | 8.482259639 | 0.000696579 | 0.002999596 |
| TAOK2 | 1.702904061 | 7.143024578 | 0.000918167 | 0.003813615 |

| | | | | |
|---|---|---|---|---|
| CLCN2 | 1.702610893 | 4.377911703 | 0.006051272 | 0.018848695 |
| FAM234B | 1.702416739 | 4.230742899 | 0.011552659 | 0.032529659 |
| PSPH | 1.702095557 | 3.489704955 | 0.003258348 | 0.011164929 |
| ITFG1_2 | 1.701280087 | 4.982559495 | 0.015579195 | 0.041700371 |
| LOC478670 | 1.70004828 | 8.519607789 | 0.001646304 | 0.006235644 |
| STK36 | 1.697809006 | 4.556558671 | 0.000498705 | 0.002253637 |
| SCAF1 | 1.697508188 | 7.54393476 | 0.006347333 | 0.019627014 |
| APLP1 | 1.696797613 | 4.314641472 | 0.009011844 | 0.026422307 |
| INKA1 | 1.695525054 | 3.481659298 | 0.005182173 | 0.016499666 |
| LOC119865055 | 1.694790551 | 3.608595316 | 0.00796967 | 0.023844099 |
| ASL | 1.693586764 | 4.008519409 | 0.001007834 | 0.004122835 |
| UAP1 | 1.692990535 | 5.849978954 | 0.002057389 | 0.007547358 |
| FBXO32 | 1.692629534 | 6.291073796 | 0.003263355 | 0.011176786 |
| FLRT1 | 1.691765966 | 3.65431447 | 0.002498767 | 0.008902809 |
| LOC106558788 | 1.690460831 | 3.84500257 | 0.005526327 | 0.01743915 |
| RYK | 1.689660646 | 5.241967901 | 0.00344694 | 0.011707494 |
| LOC119870886 | 1.688929983 | 3.354035611 | 0.009400334 | 0.027390889 |
| KAT2A | 1.687901648 | 5.787127425 | 0.01172999 | 0.032926314 |
| LOC119864195 | 1.687536143 | 2.656989133 | 0.011569473 | 0.032566187 |
| TMEM184B | 1.68716423 | 6.407846956 | 0.000529086 | 0.00237164 |
| RRP7 | 1.687153152 | 3.994539031 | 0.000794204 | 0.003354663 |
| CRAT | 1.686973296 | 5.223558825 | 0.000575479 | 0.002545898 |
| LOC102151620 | 1.685766902 | 3.035308351 | 0.010246622 | 0.029437307 |
| RACGAP1 | 1.685755719 | 3.595164579 | 0.014748786 | 0.039921375 |
| RAPH1 | 1.682436116 | 7.337265225 | 0.002412171 | 0.008629742 |
| LOC111095870 | 1.680817564 | 1.267844384 | 0.009045111 | 0.026505628 |
| LSS | 1.680758485 | 4.472588665 | 0.01730574 | 0.045330831 |
| RAB7B | 1.680094456 | 4.407179866 | 0.007055589 | 0.021476935 |
| LOC119870992 | 1.67867167 | 2.79481037 | 0.015032359 | 0.040537055 |
| CIPC | 1.677463112 | 5.032068049 | 0.000312473 | 0.001509975 |
| TBC1D24 | 1.677048555 | 4.446068929 | 0.000806605 | 0.003399362 |
| LOC119871840 | 1.676822473 | 1.912771404 | 0.008550779 | 0.025265307 |
| DOK1 | 1.675310846 | 5.239970885 | 0.000405446 | 0.001887695 |
| GOLGA4 | 1.675186602 | 7.540393987 | 0.006450918 | 0.019904719 |
| TNRC18 | 1.674273548 | 9.070808639 | 0.009665357 | 0.028016517 |
| MBNL2 | 1.674028768 | 6.631806522 | 0.00050361 | 0.002272011 |
| F2R | 1.672357299 | 6.443491193 | 0.005264038 | 0.016711694 |
| BMI1 | 1.671142558 | 4.981397492 | 0.000166607 | 0.00087813 |
| STX1A | 1.66952405 | 3.659169918 | 0.011143089 | 0.031577427 |

| | | | | |
|---|---|---|---|---|
| NAA10 | 1.669423561 | 5.459896961 | 0.003741894 | 0.012553881 |
| SLC39A3 | 1.669413291 | 1.907192858 | 0.004642722 | 0.015036234 |
| PHLDA1 | 1.668000331 | 5.50375513 | 0.005151508 | 0.016418918 |
| TBCD | 1.667033912 | 6.622061348 | 0.001403614 | 0.005448779 |
| ZNF613 | 1.665489044 | 2.623797731 | 0.004994783 | 0.015985222 |
| SRCIN1 | 1.665439349 | 4.37771281 | 0.008445723 | 0.025002616 |
| CBY1 | 1.66494441 | 2.458173531 | 0.005967618 | 0.018625601 |
| MXRA7 | 1.663340798 | 6.232238045 | 0.010221878 | 0.029385697 |
| LOC119869363 | 1.660853325 | 5.960852876 | 0.013840974 | 0.0379044 |
| LOC100686484 | 1.659529043 | 4.029155858 | 0.014815601 | 0.040072198 |
| LOC119867565 | 1.659158087 | 4.823950796 | 0.001491069 | 0.005730793 |
| D2HGDH | 1.659088876 | 3.474297114 | 0.002108174 | 0.007708867 |
| HOMER1 | 1.659015911 | 4.702635284 | 0.004367653 | 0.01427109 |
| LOC119877827 | 1.65685231 | 2.459430071 | 0.015938793 | 0.042479275 |
| LOC119867914 | 1.656629254 | 0.472084192 | 0.016257551 | 0.043148409 |
| OTUD7B | 1.655785887 | 5.968424942 | 0.001283388 | 0.005064614 |
| LOC100683760 | 1.655677451 | 6.217729849 | 0.002418612 | 0.008643757 |
| PLBD2 | 1.655091824 | 5.938554448 | 0.000318683 | 0.001533824 |
| LOC119874144 | 1.653213935 | 2.901184476 | 0.004758948 | 0.015334625 |
| CYB561 | 1.652293796 | 4.993630223 | 0.000845212 | 0.003542904 |
| PLA2G4B | 1.651829808 | 4.586729605 | 0.018112712 | 0.047029688 |
| UBTD2 | 1.65172779 | 5.557465352 | 0.002309739 | 0.008312683 |
| CHD9 | 1.650291448 | 7.769928593 | 0.004791273 | 0.0154319 |
| RNF126 | 1.649027813 | 5.618545821 | 0.013315194 | 0.036654414 |
| OPLAH | 1.64863428 | 5.64581646 | 0.000953183 | 0.003930428 |
| RANBP17 | 1.648121865 | 4.054011279 | 0.001768198 | 0.006622112 |
| DBN1 | 1.646962175 | 5.930732277 | 0.010857088 | 0.03088001 |
| TBCA | 1.646613294 | 3.787206289 | 0.002109266 | 0.00771156 |
| NF2 | 1.645901369 | 6.365859517 | 0.000273887 | 0.001343583 |
| AGPAT3 | 1.645144558 | 4.641168246 | 0.002965691 | 0.01031377 |
| PHB | 1.644322232 | 4.53680287 | 0.000289232 | 0.001410876 |
| TSFM | 1.642802051 | 4.438320222 | 0.000631121 | 0.002757355 |
| QTRT1 | 1.642791355 | 3.110991602 | 0.00599515 | 0.018700761 |
| ZFYVE21 | 1.641263134 | 4.094142878 | 0.000398444 | 0.001858689 |
| LOC119870885 | 1.640259859 | 2.366283401 | 0.012552039 | 0.034876967 |
| CAMKK2 | 1.639797549 | 7.270762529 | 0.004793025 | 0.015435251 |
| THEM4 | 1.63947225 | 3.204759822 | 0.004262535 | 0.013970713 |
| PLEKHH3 | 1.638963283 | 4.915654306 | 0.004204566 | 0.013827853 |
| HDAC7 | 1.638779096 | 8.144987959 | 0.001309225 | 0.00514783 |

| | | | | |
|---|---|---|---|---|
| ERF | 1.637257399 | 6.113576704 | 0.00075668 | 0.003217484 |
| DTNBP1 | 1.636857255 | 5.66046489 | 0.015759095 | 0.042103903 |
| DESI2 | 1.634254283 | 4.863050482 | 0.003284467 | 0.011238436 |
| SEPHS1 | 1.633891218 | 5.655727304 | 0.000307062 | 0.001486397 |
| RNFT2 | 1.633265791 | 3.168191032 | 0.012338923 | 0.034372941 |
| DOHH | 1.630933317 | 2.751618675 | 0.008711008 | 0.025668754 |
| MPV17 | 1.628639412 | 2.459732173 | 0.007795141 | 0.023383265 |
| CYB5RL | 1.627593802 | 2.846968404 | 0.002801139 | 0.00983143 |
| CFAP92 | 1.62753864 | 2.719999558 | 0.005560303 | 0.017533601 |
| UCKL1 | 1.627192429 | 6.527048304 | 0.01067042 | 0.030448982 |
| TANGO2 | 1.626106599 | 4.219745366 | 0.005780199 | 0.018142543 |
| B3GNTL1 | 1.625984414 | 5.289337509 | 0.001971262 | 0.007281945 |
| MYO6 | 1.62591401 | 5.915270324 | 0.000555178 | 0.00246818 |
| SETD1A | 1.625010972 | 7.009910274 | 0.01429184 | 0.038903171 |
| EOLA1 | 1.624126723 | 2.657168176 | 0.008809501 | 0.025909663 |
| LPIN3 | 1.622966262 | 5.479874409 | 0.003318457 | 0.011333268 |
| WDR18 | 1.622181054 | 3.400329131 | 0.005167121 | 0.016461416 |
| SPECC1L | 1.622171505 | 6.345764532 | 0.003853522 | 0.012876563 |
| PGBD1 | 1.620997326 | 2.550760793 | 0.006132305 | 0.019051811 |
| EFHC1 | 1.620551624 | 0.851626564 | 0.012394982 | 0.034502497 |
| CKAP4 | 1.620313396 | 5.867343703 | 0.017891085 | 0.0465714 |
| IRAK1BP1 | 1.62024485 | 2.256435697 | 0.014894418 | 0.04023015 |
| CEP164 | 1.620137711 | 5.231411871 | 0.002514158 | 0.008950287 |
| MRRF | 1.616704374 | 3.98267198 | 0.004711112 | 0.015209882 |
| ZCCHC14 | 1.616470388 | 6.723451494 | 0.003819841 | 0.012777808 |
| INSR | 1.615919818 | 5.689248767 | 0.001163484 | 0.004663617 |
| KCTD10 | 1.612965958 | 6.287011136 | 0.00059604 | 0.002622937 |
| AIFM2 | 1.612899133 | 3.831718022 | 0.005838756 | 0.018281327 |
| DUT | 1.612863916 | 5.130031592 | 0.001373517 | 0.005355926 |
| CASTOR1 | 1.612849107 | 4.773802145 | 0.000779024 | 0.003301482 |
| LOC106558983 | 1.612683104 | 2.871784698 | 0.009656263 | 0.02799735 |
| PLOD3 | 1.612121322 | 6.109048577 | 0.001198145 | 0.00477781 |
| ZNF366 | 1.611329448 | 4.577193085 | 0.006483386 | 0.019976459 |
| AIG1 | 1.610720835 | 4.405593791 | 0.001075164 | 0.004357962 |
| TPX2 | 1.609580532 | 4.630844465 | 0.01191277 | 0.033352985 |
| SMIM13 | 1.608962254 | 3.664223732 | 0.010423633 | 0.029835104 |
| PRKCA | 1.608633919 | 7.159975218 | 0.002398541 | 0.008588069 |
| ELAC2 | 1.607368028 | 4.414806061 | 0.006380418 | 0.019715255 |
| NMRAL1 | 1.606814717 | 3.420860997 | 0.004220346 | 0.013865023 |

| | | | | |
|---|---|---|---|---|
| MAN2A2 | 1.606650365 | 6.867763683 | 0.009586342 | 0.0278244 |
| SESN3 | 1.602494581 | 8.212272057 | 0.004958103 | 0.015885446 |
| S100A6 | 1.601001878 | 6.914035206 | 0.001730624 | 0.006500489 |
| CCDC107 | 1.599809722 | 2.079572963 | 0.018700919 | 0.048244853 |
| LMO4 | 1.59968675 | 6.252230732 | 0.002293041 | 0.008267652 |
| KATNB1 | 1.599162473 | 4.790783227 | 0.014079614 | 0.038446197 |
| CDK4 | 1.598797672 | 5.412957655 | 0.002669961 | 0.009425635 |
| LRRTM2 | 1.598406695 | 3.368829707 | 0.01062587 | 0.030361771 |
| TMTC4 | 1.597654711 | 3.274297334 | 0.014031323 | 0.038342226 |
| RAPGEF5 | 1.596545037 | 6.611339609 | 0.008547198 | 0.025258172 |
| CENPP | 1.595626277 | 2.325622177 | 0.017188167 | 0.045093711 |
| ZMYND19 | 1.59416062 | 2.744580194 | 0.005084597 | 0.016236697 |
| CALR | 1.593869407 | 7.9768645 | 0.001324434 | 0.005194435 |
| HEXD | 1.591146237 | 3.859702464 | 0.004768732 | 0.015361582 |
| ADAP2 | 1.590510278 | 5.214000148 | 0.002623573 | 0.009281864 |
| ERC1 | 1.589703301 | 7.032067502 | 0.002768552 | 0.009732817 |
| ADGRF5 | 1.588514552 | 7.113358135 | 0.008655852 | 0.025544428 |
| DYNLL1 | 1.587475829 | 4.67281951 | 0.009192931 | 0.026876891 |
| LOC102156193 | 1.586291928 | 3.708375481 | 0.009487501 | 0.027585538 |
| FRMD4A | 1.585277978 | 6.739399309 | 0.001224771 | 0.004866773 |
| TMEM245 | 1.583407881 | 6.124603628 | 0.003099399 | 0.010699789 |
| ARHGEF11 | 1.580498375 | 7.649634647 | 0.00325956 | 0.011167318 |
| ATP5F1D | 1.579144785 | 5.346447554 | 0.000747593 | 0.00318635 |
| KCTD21 | 1.578141674 | 2.022781224 | 0.009236969 | 0.026990265 |
| CCDC130 | 1.576965742 | 4.307527686 | 0.01287206 | 0.035642794 |
| LOC106558875 | 1.574337715 | 2.969777634 | 0.006507611 | 0.020031167 |
| MARCKS | 1.574087424 | 8.165757519 | 0.010327524 | 0.029634373 |
| NF1 | 1.572529982 | 7.581229832 | 0.0010815 | 0.004377914 |
| FBRSL1 | 1.570356541 | 6.960191549 | 0.016444182 | 0.04351053 |
| TNFAIP8L1 | 1.570306114 | 4.235886575 | 0.002990741 | 0.010386385 |
| IQSEC2 | 1.569909062 | 3.324866844 | 0.004049898 | 0.013416912 |
| KIAA1841 | 1.569766696 | 2.61775215 | 0.014287827 | 0.038897134 |
| ZNF362 | 1.569200405 | 6.597021148 | 0.002572498 | 0.009122013 |
| RAI1 | 1.568026109 | 6.524504296 | 0.001393938 | 0.005420926 |
| CSRP1 | 1.567289693 | 9.004371357 | 0.001151596 | 0.004623665 |
| PIGQ | 1.565384689 | 4.796850276 | 0.000605201 | 0.002658932 |
| PDXK | 1.565343765 | 7.679232137 | 0.004423185 | 0.014437303 |
| MRPL35 | 1.565340786 | 1.801879004 | 0.009490252 | 0.027589837 |
| ABCA1 | 1.564933057 | 8.407798484 | 0.003453613 | 0.011726484 |

| | | | | |
|---|---|---|---|---|
| CARMIL1 | 1.564772685 | 5.534946716 | 0.001975367 | 0.007292135 |
| BCKDHB | 1.563260843 | 3.002308175 | 0.003775646 | 0.01265732 |
| SLC29A1 | 1.562362701 | 6.279511695 | 0.007277938 | 0.022060805 |
| NICN1 | 1.560953271 | 5.485189453 | 0.008949864 | 0.026264546 |
| LOC119865448 | 1.560674573 | 5.168342196 | 0.017803023 | 0.046392318 |
| FN3KRP | 1.560508731 | 4.611200901 | 0.002446324 | 0.008734626 |
| HDDC2 | 1.55875176 | 2.696953509 | 0.012375933 | 0.03446718 |
| TIMP2 | 1.557486686 | 7.033167152 | 0.01415665 | 0.038598301 |
| GRAMD4 | 1.556321816 | 5.517241775 | 0.006208255 | 0.019250078 |
| TRIM62 | 1.555490055 | 3.970313218 | 0.003827945 | 0.012800964 |
| CFAP221 | 1.555454803 | 1.903368785 | 0.017366575 | 0.045451731 |
| EFCAB2 | 1.555061448 | 2.099900961 | 0.010425369 | 0.029836132 |
| CELSR3 | 1.55446387 | 4.849847825 | 0.014032692 | 0.038342226 |
| RAB11FIP5 | 1.554038542 | 5.81221623 | 0.005334745 | 0.016913388 |
| NPAS2 | 1.551657836 | 4.051389874 | 0.004430267 | 0.014449537 |
| LOC119866603 | 1.550963867 | 4.07094679 | 0.018075687 | 0.046944802 |
| MRPL28 | 1.550849927 | 3.879326893 | 0.001815161 | 0.006781586 |
| PPA1 | 1.550447832 | 4.941043988 | 0.00395005 | 0.013148431 |
| PTPRA | 1.547732772 | 6.213846307 | 0.001830586 | 0.006832151 |
| GMDS | 1.547475752 | 4.799461618 | 0.004614169 | 0.014957057 |
| RNH1 | 1.547261755 | 6.48473207 | 0.005570618 | 0.017558859 |
| CACNB1 | 1.545800249 | 3.441904127 | 0.010421649 | 0.029833364 |
| CAPN15 | 1.545287778 | 6.298557572 | 0.009904364 | 0.028624788 |
| RBMS2 | 1.544045414 | 4.395486243 | 0.013417763 | 0.036861855 |
| DNAJB4 | 1.543116986 | 5.061514201 | 0.00295981 | 0.010301374 |
| PPIP5K1 | 1.542008114 | 5.115735799 | 0.00067281 | 0.002914294 |
| LOC111096443 | 1.540587719 | 1.404184728 | 0.016978714 | 0.044657728 |
| LOC119863882 | 1.540478369 | 5.054566587 | 0.000375747 | 0.001766498 |
| ZNF282 | 1.539754177 | 4.56273765 | 0.001622486 | 0.006159421 |
| LOC100686099 | 1.539719866 | 5.068099596 | 0.000456677 | 0.002085892 |
| LOC102155229 | 1.539178428 | 2.102540843 | 0.014074988 | 0.038441567 |
| CTSF | 1.539030416 | 5.722038242 | 0.000634789 | 0.002770952 |
| SHC1 | 1.538929179 | 7.473887004 | 0.001451606 | 0.005602966 |
| CFAP73 | 1.538671208 | 4.937460278 | 0.003650029 | 0.012293251 |
| CTNNA1 | 1.538250631 | 7.709225171 | 0.002081896 | 0.007621781 |
| JRK | 1.537553462 | 5.67756882 | 0.003293179 | 0.011259358 |
| ERICH5 | 1.53556995 | 3.25811128 | 0.005463912 | 0.017262299 |
| IPO13 | 1.535441943 | 4.878432364 | 0.001566347 | 0.005983198 |
| PRKN | 1.534910262 | 3.17024269 | 0.013241207 | 0.036483178 |

| | | | | |
|---|---|---|---|---|
| ICOSLG | 1.534702848 | 4.197167789 | 0.014661845 | 0.039735671 |
| ABL2 | 1.533707546 | 6.375776988 | 0.00352472 | 0.011931565 |
| PURA | 1.533367224 | 6.793900017 | 0.002197889 | 0.007988402 |
| LOC119871214 | 1.533081087 | 3.248737288 | 0.004454262 | 0.014510329 |
| LOC607300 | 1.53289656 | 6.981824474 | 0.006091034 | 0.018941829 |
| LOC479912 | 1.531935544 | 4.725653224 | 0.010208163 | 0.029357952 |
| SMARCD3 | 1.531524253 | 4.340186554 | 0.001168374 | 0.004679759 |
| GSTP1 | 1.531162616 | 4.428261149 | 0.001122113 | 0.004520362 |
| DYNC1LI2 | 1.53091942 | 6.779041333 | 0.000796388 | 0.003361269 |
| YIF1A | 1.530381713 | 3.579571553 | 0.003676789 | 0.012369923 |
| LOC111094535 | 1.529045655 | 2.464039926 | 0.009353534 | 0.027280197 |
| H1-10 | 1.52885539 | 7.141536608 | 0.011026678 | 0.031292548 |
| PAQR6 | 1.52728474 | 5.416309892 | 0.006464801 | 0.019930531 |
| LOC119874443 | 1.526965198 | 3.862105533 | 0.016578784 | 0.043787845 |
| SAMD4B | 1.526207818 | 6.813982693 | 0.002404267 | 0.008605729 |
| LOC102152875 | 1.52311793 | 4.462911295 | 0.006009229 | 0.018736588 |
| AKAP1 | 1.521830087 | 5.726007852 | 0.001057657 | 0.004295838 |
| STN1 | 1.517350302 | 2.157600275 | 0.015841746 | 0.042275981 |
| SVIL | 1.516905937 | 7.759811206 | 0.010386601 | 0.029768421 |
| FAAP100 | 1.516693025 | 5.221291256 | 0.005271166 | 0.016726497 |
| AAAS | 1.516046965 | 4.496177222 | 0.001395186 | 0.005424806 |
| LOC119875931 | 1.515583512 | 2.73217916 | 0.009985723 | 0.028825331 |
| LOC111095534 | 1.514476492 | 4.525902359 | 0.004172033 | 0.013743801 |
| GFPT1 | 1.514304106 | 6.633568744 | 0.002455878 | 0.008764405 |
| HSF1 | 1.513214619 | 6.131519889 | 0.008657591 | 0.025545791 |
| SMARCB1 | 1.51167321 | 5.416351565 | 0.006898459 | 0.021060755 |
| FBXL19 | 1.510662279 | 5.173857663 | 0.009173976 | 0.026835955 |
| HAUS7 | 1.510403006 | 1.219667844 | 0.01873836 | 0.048329938 |
| CBX1 | 1.509485364 | 4.746542687 | 0.000408483 | 0.001900614 |
| NIPSNAP1 | 1.509243705 | 4.040652852 | 0.013207241 | 0.036417372 |
| C1H19orf54 | 1.509150338 | 2.612275897 | 0.005698643 | 0.017915097 |
| ZFYVE28 | 1.507404682 | 3.687613106 | 0.01238263 | 0.034474669 |
| KIAA1671 | 1.507174443 | 6.433397518 | 0.004685036 | 0.015139308 |
| CTBP2 | 1.50708343 | 5.850302811 | 0.002070834 | 0.007588974 |
| APTX | 1.50671376 | 2.543152487 | 0.006881435 | 0.021017665 |
| PANX1 | 1.506365104 | 6.669445232 | 0.015909097 | 0.04241056 |
| IFTAP | 1.504236001 | 4.595103717 | 0.015133502 | 0.040759089 |
| WIZ | 1.503724789 | 5.954457452 | 0.006096119 | 0.018950255 |
| MYBL2 | 1.503528161 | 3.799601905 | 0.014842123 | 0.040118899 |

| | | | | |
|---|---|---|---|---|
| ANKZF1 | 1.503436499 | 5.326062083 | 0.001517317 | 0.005823412 |
| CORO6 | 1.502916315 | 4.702824546 | 0.016961055 | 0.044616696 |
| NR6A1 | 1.502850505 | 3.939483593 | 0.008865443 | 0.02604592 |
| RXYLT1 | 1.502008809 | 3.012034776 | 0.014144043 | 0.038573529 |
| NAA50 | 1.501226675 | 6.378683683 | 0.001282836 | 0.005063358 |
| PRR12 | 1.501107972 | 7.733190905 | 0.010161689 | 0.029251465 |
| LOC106557731 | 1.500297687 | 2.0872304 | 0.015447205 | 0.041449458 |
| TTC38 | 1.499409511 | 4.350297457 | 0.005026395 | 0.016072155 |
| LOC106558650 | 1.499334785 | 5.703801565 | 0.002525906 | 0.00898473 |
| MRTO4 | 1.499251596 | 4.027490865 | 0.008482438 | 0.025090743 |
| TTLL11 | 1.499216061 | 2.655483771 | 0.013078474 | 0.036113627 |
| ZNHIT6 | 1.499084444 | 4.209541402 | 0.002534844 | 0.009009132 |
| LOC119865980 | 1.49897326 | 1.447995064 | 0.010346604 | 0.029677336 |
| IGSF3 | 1.495880862 | 4.758237115 | 0.0099759 | 0.028807542 |
| ZNF782 | 1.495831303 | 3.696031813 | 0.004705851 | 0.01519516 |
| TCF3 | 1.495692966 | 5.888016685 | 0.001402707 | 0.005446233 |
| SCRN2 | 1.495310193 | 2.789264717 | 0.01641413 | 0.043459032 |
| APEX1 | 1.495102834 | 5.693691708 | 0.002629155 | 0.009298577 |
| FAM53A | 1.493508509 | 3.035115706 | 0.012571442 | 0.034921925 |
| PRPF6 | 1.493132909 | 7.073633255 | 0.016134689 | 0.042885346 |
| MAZ | 1.490972899 | 7.282832019 | 0.002156255 | 0.007856844 |
| VEZT | 1.488957965 | 6.122787597 | 0.010642962 | 0.030386631 |
| PRXL2C | 1.48780854 | 3.896173756 | 0.010149684 | 0.029226654 |
| LOC106560188 | 1.48748543 | 2.12307811 | 0.015155308 | 0.040794982 |
| TCFL5 | 1.487406463 | 3.547661878 | 0.017929258 | 0.046648357 |
| SLC5A6 | 1.486966366 | 4.880902126 | 0.002427692 | 0.008672385 |
| FPGS | 1.48329388 | 6.466587005 | 0.002717183 | 0.009569306 |
| RNF217 | 1.482255431 | 5.780143266 | 0.006372198 | 0.01969828 |
| ACBD4 | 1.482188044 | 3.927112271 | 0.008420944 | 0.024942889 |
| LOC119865160 | 1.478767725 | 0.910869181 | 0.018353291 | 0.047517726 |
| LOC479185 | 1.475168764 | 6.174661892 | 0.001692324 | 0.006383177 |
| ZNF239 | 1.475121101 | 3.537770007 | 0.014451854 | 0.039279537 |
| LOC100688768 | 1.474927082 | 2.61165653 | 0.009523762 | 0.027672407 |
| COMMD4 | 1.472813857 | 3.696177382 | 0.007763088 | 0.023296788 |
| ARFRP1 | 1.472617997 | 6.101720869 | 0.002534264 | 0.009008546 |
| TCTN3 | 1.471647269 | 3.294044421 | 0.008442453 | 0.02499635 |
| REEP4 | 1.4697525 | 2.8429471 | 0.015251482 | 0.041000417 |
| VGLL4 | 1.468883909 | 6.861401428 | 0.001526048 | 0.005852774 |
| LOC119874446 | 1.466950712 | 4.092541448 | 0.012210041 | 0.034048923 |

| | | | | |
|---|---|---|---|---|
| KEL | 1.466307559 | 3.665189082 | 0.017038433 | 0.044803927 |
| APLP2 | 1.465905025 | 8.198696861 | 0.004407626 | 0.014395191 |
| JDP2 | 1.464554741 | 5.113526819 | 0.009019433 | 0.02644098 |
| FARP1 | 1.463775022 | 6.157391848 | 0.013007485 | 0.035953479 |
| HGH1 | 1.462741321 | 3.847352482 | 0.00316208 | 0.010886696 |
| DHX37 | 1.46123298 | 6.038447226 | 0.015586137 | 0.0417138 |
| PYM1 | 1.45904889 | 3.823925172 | 0.008158152 | 0.02428506 |
| BOP1 | 1.458833882 | 5.510364545 | 0.003666978 | 0.012340746 |
| PWWP3A | 1.458818974 | 4.021620326 | 0.003912081 | 0.013042074 |
| ZNF550 | 1.456668756 | 1.572604133 | 0.018757935 | 0.048368913 |
| NCL | 1.456037519 | 8.285819547 | 0.016343804 | 0.043320534 |
| CHAF1A | 1.456019571 | 3.970118482 | 0.013585229 | 0.037259828 |
| UBE2E3 | 1.455163711 | 4.343690568 | 0.002889526 | 0.01009262 |
| LOC608162 | 1.454704012 | 5.167812387 | 0.005493709 | 0.017346323 |
| SMYD5 | 1.452596258 | 4.854144185 | 0.000693646 | 0.002988747 |
| SPATS2 | 1.451891019 | 4.979906767 | 0.002090144 | 0.007648097 |
| SYNE4 | 1.451756752 | 2.994206392 | 0.01789543 | 0.046577118 |
| CAMSAP1 | 1.451439307 | 6.580712242 | 0.010061509 | 0.029013183 |
| HDAC4 | 1.451209419 | 6.219383376 | 0.004657103 | 0.015073685 |
| ZCCHC17 | 1.449160589 | 3.745529978 | 0.007196202 | 0.021846657 |
| PCGF3 | 1.44746546 | 6.216554437 | 0.005580969 | 0.01758342 |
| UBIAD1 | 1.447066238 | 4.12495577 | 0.00314207 | 0.010833293 |
| LOC106558999 | 1.446907027 | 2.936814275 | 0.010798471 | 0.030753649 |
| PMVK | 1.44561493 | 4.772199756 | 0.004660331 | 0.015078458 |
| CDC42EP4 | 1.445377385 | 6.126847128 | 0.003321651 | 0.011342387 |
| C5H11orf1 | 1.444972285 | 2.163724948 | 0.017968489 | 0.046721109 |
| MPG | 1.444520706 | 4.114593884 | 0.017226166 | 0.045166065 |
| HOMEZ | 1.444301218 | 3.169966956 | 0.006581041 | 0.020231335 |
| PEBP1 | 1.443905722 | 5.801482561 | 0.006646321 | 0.020394409 |
| ZNF865 | 1.443487277 | 5.47532586 | 0.011202754 | 0.03168023 |
| EHMT2 | 1.442957209 | 6.280638882 | 0.006598229 | 0.020275453 |
| FAM50A | 1.439983463 | 5.302927424 | 0.009838073 | 0.028474968 |
| KLHL17 | 1.439387081 | 4.468718696 | 0.003206862 | 0.011012231 |
| SRBD1 | 1.437738013 | 4.745607787 | 0.001778521 | 0.006658471 |
| PORCN | 1.437574671 | 2.337329696 | 0.012461144 | 0.034659392 |
| EEA1 | 1.437090499 | 7.595747755 | 0.017353008 | 0.045421709 |
| GALT | 1.434952389 | 3.94194157 | 0.004712412 | 0.015211813 |
| AP4M1 | 1.434694186 | 3.791610911 | 0.005094699 | 0.016264165 |
| SNX29 | 1.433684076 | 6.435068153 | 0.001003862 | 0.004109692 |

| | | | | |
|---|---|---|---|---|
| AIDA | 1.432270084 | 6.948987627 | 0.004812676 | 0.015487026 |
| KIF18A | 1.43060887 | 3.583520697 | 0.016168129 | 0.042947893 |
| LOC119874161 | 1.429941166 | 2.714990308 | 0.018508651 | 0.047844056 |
| LOC119870017 | 1.427549726 | 3.371157985 | 0.010814811 | 0.030796136 |
| KDELR2 | 1.426300947 | 6.368000503 | 0.00966745 | 0.028016517 |
| NSRP1 | 1.426151926 | 4.240001723 | 0.00928163 | 0.027090346 |
| PHF13 | 1.425787208 | 3.736200756 | 0.003879745 | 0.012950209 |
| ATP8B2 | 1.423479914 | 7.327346111 | 0.003402 | 0.01157481 |
| LOC607874 | 1.422856465 | 8.692903762 | 0.014892525 | 0.04023015 |
| CIBAR1 | 1.422776748 | 4.012863769 | 0.002212981 | 0.008033827 |
| KLHL3 | 1.421593613 | 5.519085557 | 0.009250167 | 0.027018716 |
| COA8 | 1.420350001 | 3.729494947 | 0.007025378 | 0.021400004 |
| ACBD3 | 1.420309044 | 6.09976904 | 0.002224833 | 0.008066043 |
| MYDGF | 1.420212286 | 4.001830505 | 0.003791223 | 0.012699717 |
| ZNF185 | 1.420011795 | 2.755274235 | 0.019495234 | 0.049990559 |
| RMDN2 | 1.419984125 | 2.919177368 | 0.008698707 | 0.025640338 |
| EPN1 | 1.419121015 | 6.159571835 | 0.00498863 | 0.015970247 |
| DBNDD1 | 1.418577489 | 4.422793537 | 0.004413851 | 0.014411176 |
| GAA | 1.418391569 | 6.429856111 | 0.017150371 | 0.045050147 |
| TTLL12 | 1.418240604 | 6.031236876 | 0.00544338 | 0.017215016 |
| SPAG9 | 1.418113751 | 7.633406116 | 0.002601144 | 0.009211536 |
| LOC119875949 | 1.416790558 | 2.245133717 | 0.013479364 | 0.03702041 |
| AASDH | 1.416119237 | 4.093047496 | 0.004950118 | 0.015870401 |
| EBNA1BP2 | 1.415354978 | 4.058802342 | 0.008037106 | 0.023999476 |
| NEIL2 | 1.415307869 | 2.715434223 | 0.010148476 | 0.029226654 |
| ARL6IP5 | 1.415101346 | 5.683947076 | 0.005883437 | 0.018397307 |
| ZBTB45 | 1.413237989 | 3.160578357 | 0.01511033 | 0.040711857 |
| PHKG1 | 1.411946035 | 3.878451022 | 0.019138918 | 0.049199118 |
| TSC2 | 1.411588268 | 6.956983524 | 0.00749347 | 0.022625519 |
| ZNF438 | 1.411362567 | 4.568253922 | 0.007244769 | 0.021978708 |
| TCTN2 | 1.409634876 | 3.576653172 | 0.017684446 | 0.046127693 |
| PUS7 | 1.40740294 | 4.75164552 | 0.01232936 | 0.034350716 |
| FMN1 | 1.407169148 | 5.145473779 | 0.005608339 | 0.01766071 |
| DAAM1 | 1.407089061 | 5.586086854 | 0.003192223 | 0.010973494 |
| LZTR1 | 1.406871178 | 5.292793492 | 0.004261224 | 0.013970178 |
| B4GALT7 | 1.406188111 | 4.094990286 | 0.006993691 | 0.02132146 |
| MTERF4 | 1.406106221 | 6.176279217 | 0.002181075 | 0.007933441 |
| RBM28 | 1.406020246 | 3.890779161 | 0.018165516 | 0.047132913 |
| TEX264 | 1.405010392 | 4.465843853 | 0.00134424 | 0.005258794 |

| | | | | |
|---|---|---|---|---|
| SEC24D | 1.403945231 | 6.015984158 | 0.006001514 | 0.018717919 |
| IFT46 | 1.403767305 | 4.661764414 | 0.00229969 | 0.008286147 |
| SMIM11A | 1.403087031 | 2.337722661 | 0.015437745 | 0.0414446 |
| LOC102152916 | 1.401140686 | 3.244751069 | 0.012742201 | 0.035323833 |
| RABAC1 | 1.399751673 | 5.394915414 | 0.004196789 | 0.013811954 |
| BTBD6 | 1.399315037 | 4.13903065 | 0.013387496 | 0.036806697 |
| CLN6 | 1.397918415 | 3.917862388 | 0.01482837 | 0.040096727 |
| PBDC1 | 1.397664289 | 4.828996979 | 0.016158191 | 0.042932018 |
| PHF2 | 1.396527333 | 6.525634379 | 0.005940177 | 0.018550642 |
| PEAK1 | 1.393790922 | 7.767305771 | 0.001816887 | 0.006786864 |
| CBR4 | 1.393221716 | 4.807949533 | 0.004049643 | 0.013416912 |
| PHC2 | 1.393183119 | 6.518952844 | 0.001457033 | 0.00562191 |
| LOC100856339 | 1.392660384 | 4.216999274 | 0.0147478 | 0.039921375 |
| GLE1 | 1.391963704 | 4.481316834 | 0.003693471 | 0.01242219 |
| IRGQ | 1.390525949 | 5.132597583 | 0.001982784 | 0.007315777 |
| INSYN2A | 1.389961445 | 3.863945422 | 0.004615433 | 0.014958918 |
| FAM234A | 1.389798565 | 6.719539833 | 0.003715011 | 0.012484949 |
| RCC1 | 1.388813268 | 2.936749177 | 0.016068632 | 0.042741219 |
| KRBA1 | 1.388650952 | 5.184199789 | 0.014458284 | 0.039285386 |
| CHD3 | 1.387623527 | 8.890252974 | 0.016166319 | 0.042947893 |
| CTBP1 | 1.387122109 | 6.547527966 | 0.007868071 | 0.023566154 |
| ACADSB | 1.386841302 | 5.827541931 | 0.001945133 | 0.007191551 |
| RNF214 | 1.38571815 | 5.588137415 | 0.003088746 | 0.010669811 |
| CENPE | 1.385643202 | 4.018943232 | 0.011428311 | 0.032233966 |
| LOC610164 | 1.385519324 | 4.326884717 | 0.015081379 | 0.040644526 |
| CRK | 1.384670964 | 4.206806664 | 0.001498835 | 0.005756558 |
| CLUAP1 | 1.383063284 | 3.436620766 | 0.007909257 | 0.023679697 |
| SPICE1 | 1.382919426 | 4.945976312 | 0.014325459 | 0.038979998 |
| LOC478000 | 1.381718533 | 3.729903959 | 0.005717327 | 0.017969418 |
| EGLN2 | 1.380925677 | 5.717895568 | 0.004961256 | 0.015890362 |
| LOC119876935 | 1.379892352 | 2.734279685 | 0.015644481 | 0.041818307 |
| GEMIN4 | 1.378862458 | 2.967188569 | 0.015112565 | 0.040712818 |
| FCGRT | 1.378586321 | 5.717629388 | 0.017926269 | 0.046646179 |
| ZNF664 | 1.378218978 | 6.669722658 | 0.00420817 | 0.013835506 |
| DUSP18 | 1.377348467 | 4.043482858 | 0.016885637 | 0.044456246 |
| ZFP28 | 1.375863336 | 3.906404106 | 0.00237371 | 0.008506185 |
| POP4 | 1.375736656 | 4.331407195 | 0.004424645 | 0.014437719 |
| SAC3D1 | 1.375583473 | 2.31618236 | 0.018927855 | 0.048720119 |
| MORF4L2 | 1.375163285 | 6.04073182 | 0.002318144 | 0.008338777 |

| | | | | |
|---|---|---|---|---|
| SPINDOC | 1.375012833 | 4.703464856 | 0.00403944 | 0.013392503 |
| CRAMP1 | 1.374810032 | 6.721223658 | 0.010921824 | 0.031031554 |
| ATP5MC1 | 1.374587193 | 3.237452707 | 0.017456982 | 0.045644249 |
| NPM3 | 1.373814069 | 3.818417225 | 0.005767844 | 0.018109008 |
| TIMM13 | 1.373103849 | 2.849089089 | 0.018937691 | 0.048739648 |
| LRCH1 | 1.371982676 | 7.030234916 | 0.017209578 | 0.045128032 |
| SEC23A | 1.371484186 | 5.262147719 | 0.002413399 | 0.008632712 |
| PIDD1 | 1.371482817 | 4.306586399 | 0.002349321 | 0.00842994 |
| NOC4L | 1.370879943 | 4.025841669 | 0.001563977 | 0.005976034 |
| CD151 | 1.37009762 | 6.720592585 | 0.005447849 | 0.017224105 |
| UNC13B | 1.36777315 | 5.338967359 | 0.014863436 | 0.0401715 |
| TPCN1 | 1.367545543 | 6.149349782 | 0.004940316 | 0.015850699 |
| ZNF697 | 1.367246344 | 3.705856799 | 0.015641556 | 0.041815647 |
| ADRM1 | 1.366130073 | 5.771868966 | 0.004089355 | 0.013508398 |
| RABL6 | 1.36551414 | 6.652757647 | 0.015402705 | 0.041365901 |
| FAM161B | 1.363580902 | 3.146282942 | 0.014435363 | 0.039244558 |
| CRELD2 | 1.362489347 | 4.28908484 | 0.00922982 | 0.026973825 |
| HTATSF1 | 1.36080449 | 5.686468889 | 0.00950062 | 0.027616272 |
| TBC1D17 | 1.36077816 | 5.936010627 | 0.011570598 | 0.032566187 |
| SSRP1 | 1.360685889 | 6.221799868 | 0.003291293 | 0.011256401 |
| FYCO1 | 1.359582292 | 7.792517238 | 0.014115146 | 0.038509261 |
| ARL2 | 1.359500875 | 5.658535972 | 0.009638545 | 0.027957197 |
| PRADC1 | 1.359300544 | 4.891738787 | 0.001844806 | 0.006872205 |
| MAP1S | 1.358027645 | 5.542809073 | 0.005349185 | 0.016956691 |
| CHMP2B | 1.357665601 | 4.898709385 | 0.004290959 | 0.014050152 |
| MAN1C1 | 1.355891962 | 4.924808491 | 0.001571808 | 0.006001726 |
| TCF12 | 1.35424449 | 6.858189594 | 0.001597666 | 0.006080107 |
| SLC10A3 | 1.353601214 | 4.222180485 | 0.006644378 | 0.020391334 |
| GOLGA3 | 1.352919781 | 6.760885397 | 0.013348224 | 0.036736007 |
| GPRASP1 | 1.352623752 | 2.341207865 | 0.011169712 | 0.031636328 |
| PRMT1 | 1.35257062 | 4.752817205 | 0.003999086 | 0.013289204 |
| UBAC1 | 1.351816848 | 5.301518215 | 0.005026281 | 0.016072155 |
| GPS1 | 1.351803327 | 5.682734048 | 0.006521541 | 0.020071194 |
| CARS1 | 1.350677823 | 4.968131135 | 0.007831763 | 0.023473541 |
| TMEM104 | 1.347807013 | 5.117403447 | 0.007845814 | 0.023505991 |
| CENPJ | 1.346479152 | 3.498210138 | 0.018419831 | 0.047664169 |
| CALM3 | 1.346114368 | 6.462319418 | 0.003008043 | 0.010434268 |
| SH3GL1 | 1.345461138 | 6.390973092 | 0.018926319 | 0.048720119 |
| BRD3 | 1.345080012 | 6.672452722 | 0.016405612 | 0.043450915 |

| | | | | |
|---|---|---|---|---|
| MYBBP1A | 1.344744933 | 6.594618247 | 0.006044754 | 0.018833806 |
| FAM53C | 1.344488156 | 4.768987809 | 0.004679741 | 0.015131139 |
| STXBP4 | 1.342818034 | 4.975403092 | 0.002727804 | 0.009603589 |
| CLPP | 1.340639168 | 4.349929245 | 0.016029522 | 0.042668612 |
| FAM114A2 | 1.340610941 | 4.960168039 | 0.003524157 | 0.011931565 |
| ARFIP2 | 1.339564121 | 3.456942659 | 0.004972038 | 0.015919484 |
| PCYT1A | 1.338936408 | 6.05981837 | 0.002882468 | 0.010076086 |
| PELI3 | 1.338212483 | 4.32735765 | 0.01294374 | 0.035813824 |
| ASB1 | 1.337654358 | 4.129897098 | 0.007308567 | 0.022147452 |
| RHOT2 | 1.336603046 | 5.802663664 | 0.014101026 | 0.038485561 |
| ENTPD5 | 1.336194233 | 3.879828906 | 0.010821656 | 0.03080929 |
| NAB2 | 1.334408874 | 4.664542084 | 0.009463442 | 0.027534054 |
| PPP2R5B | 1.333281679 | 4.673288272 | 0.007093133 | 0.021570011 |
| POGK | 1.330611182 | 5.31285699 | 0.008003523 | 0.023928868 |
| STK3 | 1.330200689 | 5.085624388 | 0.015440471 | 0.041446784 |
| MAP4K4 | 1.329982473 | 8.875175947 | 0.009679952 | 0.028043523 |
| SLC4A8 | 1.328733988 | 4.27932813 | 0.01244648 | 0.034632501 |
| SEMA4G | 1.327963543 | 3.879131208 | 0.01215238 | 0.033914309 |
| KSR1 | 1.327856608 | 6.12602822 | 0.006755236 | 0.020705167 |
| CLUH | 1.327758595 | 5.956922497 | 0.002837042 | 0.009938129 |
| FOCAD | 1.327078954 | 5.694910386 | 0.003498884 | 0.011865327 |
| CDPF1 | 1.326898792 | 4.100654993 | 0.004625393 | 0.014988957 |
| TMCC2 | 1.325526208 | 4.171713516 | 0.007831491 | 0.023473541 |
| TAMALIN | 1.323891493 | 4.007151603 | 0.017286739 | 0.045286535 |
| TMEM175 | 1.322884591 | 6.057042501 | 0.014508373 | 0.039398566 |
| SSH3 | 1.322857184 | 5.130197854 | 0.010198128 | 0.029332983 |
| PDK2 | 1.321320815 | 5.149546524 | 0.012160574 | 0.033932807 |
| LRRC42 | 1.320587903 | 3.436463827 | 0.005821599 | 0.01825342 |
| LOC111090086 | 1.319677049 | 3.463223135 | 0.009592608 | 0.027838857 |
| PAOX | 1.319012622 | 4.061192089 | 0.008468802 | 0.025057248 |
| VSTM4 | 1.318898801 | 4.391365055 | 0.008841943 | 0.025987447 |
| PDIA6 | 1.318261795 | 6.714280574 | 0.017602326 | 0.045946672 |
| ACACA | 1.317267938 | 7.041750118 | 0.005419155 | 0.017159258 |
| BCKDHA | 1.315017188 | 4.957258124 | 0.002776177 | 0.009754879 |
| IHO1 | 1.314458654 | 2.644535416 | 0.012515594 | 0.034793544 |
| MLH3 | 1.314404733 | 5.050706478 | 0.004084573 | 0.01350289 |
| PDP1 | 1.313979788 | 5.457145717 | 0.010535854 | 0.030128456 |
| FAM114A1 | 1.310995285 | 5.680881839 | 0.014595101 | 0.039584485 |
| ZNF346 | 1.309537282 | 3.410929175 | 0.016337426 | 0.043314584 |

| | | | | |
|---|---|---|---|---|
| DNPEP | 1.306453373 | 3.904543419 | 0.004154052 | 0.013692894 |
| POLE | 1.303491754 | 4.468504249 | 0.015306703 | 0.041128462 |
| ADI1 | 1.303411865 | 4.101271754 | 0.014982425 | 0.040417489 |
| PGM3 | 1.303263343 | 5.198407995 | 0.010910823 | 0.031008429 |
| ZSCAN12 | 1.301848718 | 3.926737895 | 0.007700408 | 0.023137522 |
| NDRG4 | 1.30063822 | 5.984795752 | 0.011598524 | 0.032620662 |
| EEF1D | 1.299135517 | 7.110062718 | 0.009674315 | 0.02803094 |
| XPC | 1.298978592 | 4.913404056 | 0.01483066 | 0.040097916 |
| MGRN1 | 1.298165001 | 6.267256603 | 0.002729067 | 0.009606474 |
| DNMBP | 1.297400162 | 6.064047048 | 0.007520156 | 0.022690289 |
| WDR13 | 1.296498499 | 4.985478406 | 0.009355169 | 0.027281295 |
| LDAH | 1.295149959 | 4.076687264 | 0.011705841 | 0.032871298 |
| CUL7 | 1.292919759 | 6.865628034 | 0.00427266 | 0.014000809 |
| PTPN23 | 1.29284857 | 7.147626635 | 0.019058111 | 0.048997204 |
| CDKL5 | 1.291940438 | 3.96432261 | 0.017465416 | 0.045655285 |
| TMEM135 | 1.28980396 | 4.645979772 | 0.003291795 | 0.011256401 |
| GPAM | 1.288351849 | 6.317842682 | 0.010127076 | 0.029178956 |
| RUSF1 | 1.287028282 | 5.955660965 | 0.001605162 | 0.006104349 |
| C23H3orf33 | 1.284136366 | 3.451923643 | 0.014951411 | 0.040348888 |
| CYHR1 | 1.283565106 | 3.967276599 | 0.006827499 | 0.020888254 |
| ELOF1 | 1.283067975 | 3.823069937 | 0.006820855 | 0.020873821 |
| SLC31A1 | 1.282388403 | 4.645973414 | 0.009585698 | 0.0278244 |
| ZNF846 | 1.281295072 | 3.471550925 | 0.012175789 | 0.033970891 |
| RNF26 | 1.281106965 | 4.014816504 | 0.004131369 | 0.013630561 |
| AP1S1 | 1.275478456 | 4.203112144 | 0.009559195 | 0.027764195 |
| RAD51D | 1.274000916 | 3.716653733 | 0.008994227 | 0.026377791 |
| EXD3 | 1.273731603 | 5.153245878 | 0.012665278 | 0.035142054 |
| ABCF1 | 1.272868765 | 6.15367992 | 0.015594268 | 0.041720106 |
| RAD23B | 1.272176926 | 7.181828846 | 0.005517576 | 0.017416608 |
| NRIP2 | 1.272038492 | 5.02232293 | 0.010691363 | 0.030492684 |
| ALS2 | 1.271682286 | 5.33522706 | 0.008641692 | 0.025514256 |
| SLC25A22 | 1.271553305 | 5.502928532 | 0.017245091 | 0.04518835 |
| HAGH | 1.271142162 | 3.747915141 | 0.007819472 | 0.023443273 |
| STX2 | 1.268587171 | 3.857853331 | 0.013434167 | 0.036902242 |
| KNOP1 | 1.267562374 | 4.503864451 | 0.002047543 | 0.007520147 |
| LOC106559047 | 1.266772338 | 3.772839341 | 0.006071297 | 0.018897488 |
| SULT1C4 | 1.265774927 | 5.297742792 | 0.018952544 | 0.048772084 |
| PAXX | 1.265465381 | 4.62862884 | 0.006206949 | 0.019250078 |
| AKAP7 | 1.265346432 | 4.481306356 | 0.002331841 | 0.008374668 |

| | | | | |
|---|---|---|---|---|
| LOC100685940 | 1.262349432 | 4.375119427 | 0.016088914 | 0.042779417 |
| LOC102151514 | 1.262127702 | 3.665534314 | 0.013394173 | 0.036815715 |
| HSPB1 | 1.261870308 | 6.592347335 | 0.005833813 | 0.018277851 |
| PKP4 | 1.261075798 | 6.148800406 | 0.010656325 | 0.03041677 |
| ESF1 | 1.260902812 | 4.591050153 | 0.005913816 | 0.018478969 |
| STK11IP | 1.259671459 | 4.445604921 | 0.013528165 | 0.037115763 |
| CAPRIN2 | 1.254514939 | 4.669183395 | 0.017123474 | 0.044989338 |
| MTOR | 1.254310013 | 7.236397651 | 0.011082158 | 0.031437644 |
| CIZ1 | 1.253973745 | 5.766696817 | 0.01548872 | 0.041535144 |
| SLC2A11 | 1.25390073 | 4.123753316 | 0.00905572 | 0.026529415 |
| DMXL2 | 1.252417129 | 5.1116192 | 0.01491872 | 0.040280729 |
| CARD6 | 1.251645638 | 5.413688044 | 0.01650376 | 0.043625563 |
| COLGALT1 | 1.250759554 | 6.132741484 | 0.003641712 | 0.012269053 |
| PDCD5 | 1.248170082 | 4.213768675 | 0.009114402 | 0.026686901 |
| ZNF365 | 1.247793319 | 4.571046464 | 0.016830843 | 0.044338744 |
| PMS1 | 1.247678705 | 5.412261529 | 0.001469242 | 0.005659951 |
| WDR97 | 1.2468078 | 5.041816674 | 0.015207977 | 0.040905223 |
| EID1 | 1.245966908 | 6.902177551 | 0.007390881 | 0.022359376 |
| IFT122 | 1.244850418 | 4.968386659 | 0.003378382 | 0.011507099 |
| RING1 | 1.244508567 | 6.080658301 | 0.017573129 | 0.045875984 |
| VARS1 | 1.242459236 | 6.506706595 | 0.008936217 | 0.026228958 |
| SLC41A1 | 1.242353595 | 5.57190657 | 0.00426283 | 0.013970713 |
| ARHGAP35 | 1.242103208 | 7.247299874 | 0.007077774 | 0.021536542 |
| ERCC2 | 1.241031329 | 3.450952259 | 0.015498825 | 0.041557101 |
| ARSA | 1.238349632 | 6.909907601 | 0.006402105 | 0.019776629 |
| PDIA4 | 1.237314722 | 7.317350461 | 0.01460319 | 0.039601468 |
| PCBP4 | 1.237244556 | 5.761957421 | 0.009177053 | 0.026841331 |
| TULP4 | 1.235028312 | 6.235747126 | 0.009141207 | 0.026747317 |
| DDX54 | 1.234824465 | 6.009065059 | 0.011908671 | 0.033345818 |
| EIF2B3 | 1.234162646 | 3.845471726 | 0.01012275 | 0.029170371 |
| HSF2BP | 1.232987308 | 3.526212996 | 0.014027729 | 0.03833833 |
| ACD | 1.230112729 | 3.999036263 | 0.01611438 | 0.042841875 |
| ARHGAP5 | 1.229430325 | 6.881388756 | 0.009468316 | 0.027544538 |
| LOC119875988 | 1.228447517 | 4.47218707 | 0.005750601 | 0.018062722 |
| LOC100856607 | 1.228282116 | 4.167439471 | 0.016488884 | 0.043596874 |
| TXNRD3 | 1.226967694 | 3.473691353 | 0.012892611 | 0.035690582 |
| POLD1 | 1.224178222 | 4.416505768 | 0.010493683 | 0.030015785 |
| ARL10 | 1.223747915 | 4.343404478 | 0.003350804 | 0.01142933 |
| TMEM161A | 1.221768529 | 3.905294582 | 0.012735751 | 0.035310469 |

| | | | | |
|---|---|---|---|---|
| SCP2 | 1.220204428 | 5.835369623 | 0.0147593 | 0.03993487 |
| TIMELESS | 1.219739119 | 4.815947094 | 0.010317296 | 0.029608946 |
| CD99 | 1.219311505 | 8.938441166 | 0.011196745 | 0.031675614 |
| LOC119876729 | 1.2190452 | 3.572099692 | 0.015763894 | 0.042111533 |
| APEH | 1.217185697 | 5.055922776 | 0.004163942 | 0.013719233 |
| TIGD7 | 1.216900637 | 5.693497783 | 0.010921122 | 0.031031554 |
| FAM120C | 1.21668717 | 5.927152244 | 0.002152104 | 0.007843037 |
| CFL2 | 1.215053556 | 5.775822864 | 0.015194662 | 0.040878093 |
| SPDL1 | 1.214225331 | 3.443315558 | 0.019030574 | 0.048937109 |
| MESD | 1.212238743 | 5.267451279 | 0.006750728 | 0.020694275 |
| FAM160B2 | 1.208754781 | 5.707766344 | 0.014178352 | 0.038642783 |
| SLC7A1 | 1.2086844 | 5.045205865 | 0.01207227 | 0.033729831 |
| NLRX1 | 1.207364291 | 4.565901869 | 0.011888496 | 0.033302233 |
| NDOR1 | 1.20636198 | 4.884343182 | 0.009625288 | 0.027929959 |
| CPLANE1 | 1.206148977 | 7.237682885 | 0.013586694 | 0.037259828 |
| LXN | 1.205751131 | 5.224005095 | 0.017940796 | 0.046672774 |
| ATXN7L3B | 1.203194194 | 6.283560191 | 0.010727502 | 0.030583681 |
| GTF2I | 1.20266292 | 7.253184962 | 0.005618612 | 0.017686592 |
| NUCB1 | 1.199792782 | 7.203383013 | 0.007801155 | 0.02339483 |
| TMEM237 | 1.199553188 | 4.867273638 | 0.004459721 | 0.014523747 |
| THRA | 1.194512371 | 5.803042775 | 0.006998646 | 0.021330565 |
| LOC111096351 | 1.194492728 | 4.444236447 | 0.018828141 | 0.048509547 |
| STX6 | 1.194005179 | 5.821113617 | 0.010877768 | 0.030926652 |
| KCTD3 | 1.193140513 | 6.162597261 | 0.010411206 | 0.029809021 |
| ZNF699 | 1.193065263 | 3.911542438 | 0.015476876 | 0.041513654 |
| GNL1 | 1.192617455 | 5.374868155 | 0.008005406 | 0.023931196 |
| ZMIZ1 | 1.191701977 | 8.340905102 | 0.01716879 | 0.045064694 |
| DNAJB2 | 1.190679454 | 6.263108527 | 0.008786277 | 0.025851886 |
| RPUSD1 | 1.189310061 | 4.014869779 | 0.010431691 | 0.029850283 |
| AKT1S1 | 1.185611862 | 6.012262114 | 0.00911106 | 0.026680719 |
| IDS | 1.18280011 | 6.49995472 | 0.012121207 | 0.033840388 |
| ZNF436 | 1.181720104 | 5.426823654 | 0.008025518 | 0.023978088 |
| TUBG1 | 1.179658666 | 3.994562724 | 0.014076144 | 0.038441567 |
| BET1L | 1.177964611 | 5.980564608 | 0.005886956 | 0.018403128 |
| SOD1 | 1.177498709 | 5.348591292 | 0.009717296 | 0.028136668 |
| APPL2 | 1.172269682 | 5.367345279 | 0.004400756 | 0.014374917 |
| ALKBH6 | 1.172167488 | 4.886558934 | 0.016495275 | 0.043608453 |
| TECPR2 | 1.1720475 | 5.038007356 | 0.01633815 | 0.043314584 |
| NUDCD3 | 1.169913824 | 5.949308275 | 0.015546512 | 0.041659193 |

| | | | | |
|---|---|---|---|---|
| CAMSAP2 | 1.169782081 | 6.842466473 | 0.008857966 | 0.026031009 |
| TMEM109 | 1.169161942 | 6.129066096 | 0.009188303 | 0.026868544 |
| CRELD1 | 1.168427387 | 3.709686576 | 0.017233575 | 0.045174562 |
| CDK16 | 1.167249753 | 5.781774809 | 0.019158831 | 0.049238628 |
| FBXO31 | 1.166137325 | 4.926342569 | 0.010236006 | 0.029418507 |
| GNB1L | 1.165585995 | 4.358524622 | 0.012711787 | 0.035257557 |
| PES1 | 1.165080434 | 4.503028564 | 0.012104238 | 0.03380172 |
| LANCL1 | 1.163021953 | 5.01681015 | 0.013377065 | 0.036792021 |
| HMGN1 | 1.162981546 | 6.840577884 | 0.013293887 | 0.036605059 |
| TMEM134 | 1.160274732 | 4.285402288 | 0.014753199 | 0.039923347 |
| LOC102154270 | 1.15615327 | 5.009949722 | 0.018962197 | 0.04879113 |
| CDK5RAP3 | 1.156034903 | 4.653622616 | 0.007228483 | 0.021932368 |
| TPD52L2 | 1.154976457 | 5.2302001 | 0.005622258 | 0.017695497 |
| KIF16B | 1.150422049 | 5.837561534 | 0.016435266 | 0.04349225 |
| APLF | 1.148209419 | 4.936112626 | 0.018688425 | 0.04821836 |
| IKBIP | 1.143650315 | 5.677275224 | 0.008197636 | 0.024378069 |
| CEP41 | 1.140849587 | 4.767239697 | 0.014366627 | 0.039087108 |
| CTDSP2 | 1.138614568 | 8.093550618 | 0.008836182 | 0.025977559 |
| WDR77 | 1.135291852 | 3.965153594 | 0.009885317 | 0.028584985 |
| FTO | 1.135005897 | 5.763570356 | 0.015431188 | 0.041437262 |
| MRPL4 | 1.134353239 | 4.937715494 | 0.010856246 | 0.03088001 |
| SCYL1 | 1.126133544 | 5.390303762 | 0.007589621 | 0.022865707 |
| NDFIP1 | 1.125244038 | 6.299046371 | 0.01151379 | 0.032441286 |
| ZNF517 | 1.125048974 | 5.513383526 | 0.011571643 | 0.032566187 |
| INTS1 | 1.12495808 | 6.30158008 | 0.018401265 | 0.047630554 |
| IMPDH2 | 1.120679921 | 5.841824731 | 0.006500859 | 0.02002176 |
| DIXDC1 | 1.119018473 | 6.208586172 | 0.011433323 | 0.032243904 |
| ZNF268 | 1.118078522 | 4.849015562 | 0.015275821 | 0.041056492 |
| GATD3A | 1.114370847 | 6.131955529 | 0.010193442 | 0.029327289 |
| RNF144A | 1.111655195 | 7.195121704 | 0.009420016 | 0.0274298 |
| UTP4 | 1.110236824 | 4.384416418 | 0.018302244 | 0.047408867 |
| GOLPH3L | 1.109107572 | 4.41575415 | 0.015465296 | 0.041487728 |
| YIPF2 | 1.102172475 | 4.974031365 | 0.010541692 | 0.030141172 |
| PIGG | 1.100187772 | 4.963807027 | 0.017524479 | 0.04577655 |
| CHMP7 | 1.094082917 | 4.508580336 | 0.016324194 | 0.043293471 |
| CKAP5 | 1.09254347 | 6.830987619 | 0.017973681 | 0.046721109 |
| LOC119869347 | 1.092017928 | 5.667730923 | 0.012383394 | 0.034474669 |
| ZNF862 | 1.091778903 | 4.417061557 | 0.017009384 | 0.044732966 |
| DUSP3 | 1.09121655 | 5.69059052 | 0.011145924 | 0.031581331 |

| | | | | |
|---|---|---|---|---|
| NPHP3 | 1.089516467 | 4.642660529 | 0.009550768 | 0.027747156 |
| MRPL54 | 1.088944214 | 4.594441098 | 0.016310786 | 0.043268503 |
| UBQLN2 | 1.088389929 | 5.622690549 | 0.012382611 | 0.034474669 |
| TNPO2 | 1.087932535 | 6.032133873 | 0.010688053 | 0.03049127 |
| EXT2 | 1.084123742 | 5.930066325 | 0.012598138 | 0.034969191 |
| FBXW8 | 1.08389811 | 6.302884135 | 0.011832086 | 0.033165648 |
| FKRP | 1.082952114 | 3.828469664 | 0.017163065 | 0.045060583 |
| COX19 | 1.082122051 | 4.232074268 | 0.014631 | 0.039666959 |
| LMF2 | 1.079989111 | 5.702881582 | 0.017815001 | 0.046417951 |
| IFRD2 | 1.07993739 | 4.807139408 | 0.013996373 | 0.038262283 |
| HYOU1 | 1.079440159 | 6.202816909 | 0.018921939 | 0.048716464 |
| QSOX2 | 1.078082193 | 4.698361286 | 0.017431463 | 0.045594026 |
| ZNF7 | 1.075516699 | 5.99816147 | 0.015980406 | 0.042574474 |
| MEAK7 | 1.074264891 | 3.618010479 | 0.01915482 | 0.049234158 |
| GOT1 | 1.073095581 | 4.063322847 | 0.016187855 | 0.042995023 |
| C1QBP | 1.068854536 | 5.136515766 | 0.017868326 | 0.046534514 |
| MRPL18 | 1.068589614 | 4.29784479 | 0.017309389 | 0.045334912 |
| ACAD8 | 1.0630442 | 4.817214609 | 0.014431541 | 0.039239091 |
| ACP2 | 1.062256466 | 5.162108856 | 0.011535658 | 0.032494452 |
| LOC487150 | 1.059638218 | 4.303523446 | 0.012827897 | 0.035543213 |
| ABHD12 | 1.05864954 | 4.630156358 | 0.017439699 | 0.04560561 |
| WASL | 1.057331692 | 6.083611394 | 0.012571014 | 0.034921925 |
| NOC2L | 1.057142935 | 5.330288503 | 0.010090469 | 0.029088952 |
| USP5 | 1.055802545 | 5.556163223 | 0.016788112 | 0.04424231 |
| RABGAP1 | 1.051695093 | 6.408976484 | 0.017513276 | 0.045758317 |
| JRKL | 1.051295026 | 4.156438769 | 0.015633665 | 0.041804863 |
| SIL1 | 1.050432273 | 4.570005491 | 0.01475087 | 0.039922027 |
| EMC10 | 1.042944587 | 5.38308296 | 0.012946759 | 0.035817605 |
| PHF14 | 1.041884756 | 5.911449582 | 0.017722696 | 0.046216339 |
| POLR1D | 1.036161174 | 5.094722987 | 0.018503692 | 0.047844056 |
| SNRPD2 | 1.034081069 | 5.281516616 | 0.019406868 | 0.049781638 |
| PLSCR3 | 1.033509353 | 5.278506992 | 0.01290125 | 0.035709936 |
| SMAD5 | 0.997423908 | 6.714285544 | 0.018854289 | 0.048559598 |
| TIMM21 | 0.989233661 | 4.256578497 | 0.019316818 | 0.04959173 |
| GOLM2 | 0.987343898 | 5.636226214 | 0.015873328 | 0.042336037 |
| TRIM52 | 0.984805509 | 4.927593758 | 0.01670112 | 0.044055911 |
| KDM1B | -0.867798447 | 5.575019789 | 0.017687818 | 0.046130937 |
| RTF2 | -0.898675122 | 4.540532963 | 0.017661698 | 0.046079448 |
| LOC111092104 | -0.908834693 | 4.778344119 | 0.014307443 | 0.038935862 |

| | | | | |
|---|---|---|---|---|
| FBXL5 | -0.91165099 | 5.850260373 | 0.015843008 | 0.042275981 |
| TSTD2 | -0.915255579 | 5.108094603 | 0.013766208 | 0.037718714 |
| NUP50 | -0.918477759 | 6.275242922 | 0.013109704 | 0.036189864 |
| C7H18orf25 | -0.918751576 | 5.908951341 | 0.017879669 | 0.046552867 |
| SP3 | -0.920287587 | 6.858608636 | 0.018576019 | 0.047988929 |
| PRDM4 | -0.921312542 | 5.038095545 | 0.012935094 | 0.035799042 |
| INPP5K | -0.922222654 | 5.192860905 | 0.011569943 | 0.032566187 |
| LOC111096733 | -0.922383437 | 4.73463313 | 0.016901389 | 0.044486737 |
| TNFRSF1A | -0.929247973 | 6.845265486 | 0.018784274 | 0.04841955 |
| R3HCC1L | -0.930062594 | 4.508705034 | 0.014515657 | 0.039413411 |
| CTR9 | -0.932288141 | 5.679925287 | 0.010232418 | 0.029412094 |
| ZFAND5 | -0.938377936 | 7.334832845 | 0.018966272 | 0.048795822 |
| TMPO | -0.938955126 | 5.779857009 | 0.008781063 | 0.025840051 |
| EML4 | -0.941956233 | 6.603779715 | 0.019239627 | 0.049416974 |
| UBE2L3 | -0.946775099 | 5.476803523 | 0.012462541 | 0.034659392 |
| PDCD10 | -0.94687494 | 4.415842853 | 0.019366328 | 0.049695291 |
| SNAP29 | -0.947786909 | 6.318395447 | 0.016698683 | 0.044054845 |
| TMEM183A | -0.953411344 | 5.375645348 | 0.011988659 | 0.033532112 |
| GAN | -0.954208519 | 5.900917166 | 0.017065198 | 0.044852539 |
| PANK2 | -0.955056543 | 5.414756181 | 0.01620618 | 0.043027874 |
| PRDX5 | -0.955181003 | 5.546176273 | 0.018343586 | 0.047498273 |
| ARPC4 | -0.95531504 | 6.64935478 | 0.017789748 | 0.0463633 |
| VRK2 | -0.956413061 | 4.447969433 | 0.014695225 | 0.039806226 |
| LOC119866139 | -0.957216729 | 4.755552798 | 0.010583647 | 0.030255499 |
| SNRPA1 | -0.957776683 | 4.518413494 | 0.016801123 | 0.044271214 |
| CAPN7 | -0.95887021 | 5.423904744 | 0.011331533 | 0.031998477 |
| CLK1 | -0.960324533 | 6.032093272 | 0.010406014 | 0.029804352 |
| NAPB | -0.961853279 | 4.279492599 | 0.018393334 | 0.04761571 |
| STXBP3 | -0.963187825 | 5.488925784 | 0.01582205 | 0.042230453 |
| ATF4 | -0.964909677 | 7.92211707 | 0.016687946 | 0.04403902 |
| DDX6 | -0.96744631 | 7.350322416 | 0.017204241 | 0.045123793 |
| PSMD6 | -0.96781834 | 4.702534541 | 0.015774054 | 0.042133481 |
| BACH1 | -0.968562654 | 6.879823418 | 0.014624798 | 0.039655104 |
| ABHD6 | -0.969851397 | 3.760694743 | 0.018621982 | 0.048092732 |
| ZNF131 | -0.971700461 | 4.585875182 | 0.007354074 | 0.022263563 |
| ZBTB7B | -0.972062088 | 5.458344867 | 0.019337136 | 0.049638014 |
| ARSB | -0.972987396 | 4.816526113 | 0.018506678 | 0.047844056 |
| TTC13 | -0.97367508 | 5.018044621 | 0.01132054 | 0.031971601 |
| GSS | -0.975598141 | 5.001196958 | 0.011928693 | 0.033391547 |

| | | | | |
|---|---|---|---|---|
| RIPK1 | -0.976768946 | 6.546208522 | 0.007621444 | 0.022951167 |
| LOC102152446 | -0.978784711 | 5.836340538 | 0.016429203 | 0.043481513 |
| PKD1L3 | -0.981207838 | 5.817485936 | 0.013485646 | 0.037024879 |
| UBE2F | -0.981208903 | 4.115158215 | 0.009323178 | 0.027202644 |
| SFT2D1 | -0.983338397 | 4.838949266 | 0.017150776 | 0.045050147 |
| OARD1 | -0.983951844 | 4.831493961 | 0.009385458 | 0.02735858 |
| PNRC1 | -0.986334218 | 7.43746183 | 0.00946223 | 0.027534054 |
| RLF | -0.988201625 | 5.333462017 | 0.008643952 | 0.025516259 |
| SUDS3 | -0.989208512 | 6.247838709 | 0.006591021 | 0.020259139 |
| BTBD9 | -0.992323929 | 5.129316434 | 0.007714846 | 0.023168055 |
| SPTY2D1 | -0.995314997 | 5.744700804 | 0.005570743 | 0.017558859 |
| CSNK1G3 | -0.995649541 | 5.958442005 | 0.011931165 | 0.033391547 |
| RHOF | -0.999119292 | 4.885458772 | 0.010357234 | 0.029703895 |
| FLOT1 | -1.001252824 | 7.28214182 | 0.008009243 | 0.023939365 |
| SH2B3 | -1.002200086 | 6.873767368 | 0.015003506 | 0.040469319 |
| RB1 | -1.003421239 | 5.949795922 | 0.006960436 | 0.021238003 |
| TFAM | -1.005122128 | 4.638327533 | 0.016718452 | 0.044090899 |
| AUH | -1.006402035 | 4.313815395 | 0.007998633 | 0.023920846 |
| C6H16orf72 | -1.007455793 | 6.918006429 | 0.008489935 | 0.025106067 |
| PHTF1 | -1.00748153 | 5.256545446 | 0.015641212 | 0.041815647 |
| FAM13B | -1.009591873 | 6.449675133 | 0.015578408 | 0.041700371 |
| GDI1 | -1.01099057 | 7.315081359 | 0.009633577 | 0.027946529 |
| CRKL | -1.011764709 | 6.070670789 | 0.010966533 | 0.031138171 |
| ZNF143 | -1.014870869 | 4.050792524 | 0.01640856 | 0.043453411 |
| NFATC3 | -1.015386686 | 6.426823725 | 0.010309798 | 0.029591345 |
| RAB11A | -1.019430157 | 5.410201997 | 0.018843785 | 0.048538313 |
| IKZF5 | -1.019819057 | 5.217652495 | 0.005172134 | 0.016474966 |
| MARK2 | -1.020993649 | 6.652957419 | 0.018206332 | 0.047204908 |
| MEMO1 | -1.024280186 | 5.498473214 | 0.009308312 | 0.027162926 |
| DONSON | -1.025067771 | 5.746027697 | 0.015850556 | 0.042280504 |
| CPOX | -1.025523325 | 4.328979705 | 0.015081589 | 0.040644526 |
| DGKA | -1.026675015 | 6.948601606 | 0.016017287 | 0.042656999 |
| ZNF394 | -1.027639835 | 4.151699661 | 0.007081567 | 0.021543907 |
| LOC119863905 | -1.028822196 | 5.964966162 | 0.014393497 | 0.039150385 |
| RAB5A | -1.030914949 | 4.645419509 | 0.014561061 | 0.039502051 |
| CRBN | -1.032303027 | 5.118857039 | 0.006070345 | 0.018897241 |
| RPL18 | -1.034155988 | 7.567441777 | 0.011176134 | 0.031637976 |
| GMEB1 | -1.035361255 | 4.786189255 | 0.004946196 | 0.015861866 |
| ZNF212 | -1.036948272 | 3.981244098 | 0.013177033 | 0.036352575 |

| | | | | |
|---|---|---|---|---|
| ZNF217 | -1.042962927 | 6.641678453 | 0.008892791 | 0.026119184 |
| LOC111096963 | -1.046535169 | 4.611833623 | 0.011254706 | 0.031810795 |
| COX4I1 | -1.049543383 | 5.112042815 | 0.008619054 | 0.025460101 |
| PPP1R15B | -1.050643163 | 6.275715452 | 0.007254133 | 0.021997878 |
| RRAGD | -1.051175043 | 4.401020808 | 0.016422719 | 0.043474971 |
| HPS3 | -1.05128085 | 5.544492783 | 0.00557589 | 0.017572527 |
| TTPAL | -1.051804061 | 6.473644338 | 0.011190394 | 0.031670073 |
| LOC119864282 | -1.053766328 | 5.392154301 | 0.004737635 | 0.015281853 |
| VCPIP1 | -1.054559141 | 6.269477141 | 0.007269569 | 0.022038518 |
| LOC610321 | -1.05533346 | 4.588353107 | 0.017667609 | 0.046089322 |
| NIP7 | -1.056373929 | 4.229597569 | 0.008320042 | 0.024684499 |
| GALNS | -1.060194481 | 4.481078597 | 0.011493195 | 0.032400106 |
| TFEB | -1.060519623 | 5.431994531 | 0.016129864 | 0.042877781 |
| PHF10 | -1.060689788 | 5.508943836 | 0.010403575 | 0.029801306 |
| CNOT11 | -1.061887569 | 4.966280061 | 0.003355141 | 0.011440525 |
| PDSS1 | -1.063027356 | 4.116375759 | 0.010250487 | 0.029444508 |
| LOC119871462 | -1.063627576 | 3.57497038 | 0.01193009 | 0.033391547 |
| PIGV | -1.065834058 | 4.216905196 | 0.007632445 | 0.02297472 |
| KLF6 | -1.06656699 | 8.232329479 | 0.013679707 | 0.037495926 |
| CLINT1 | -1.067718252 | 6.699835612 | 0.015721062 | 0.042007468 |
| RAVER2 | -1.067956888 | 4.149121469 | 0.018497704 | 0.047840183 |
| ITM2B | -1.068195954 | 8.366694597 | 0.014975703 | 0.040404384 |
| LOC119868555 | -1.069293807 | 6.891169892 | 0.006234342 | 0.019318971 |
| ACTR8 | -1.071560728 | 5.334935794 | 0.003183022 | 0.0109501 |
| BCL10 | -1.071636538 | 4.753522798 | 0.006834412 | 0.020905876 |
| TARDBP | -1.071664504 | 6.319677037 | 0.005927619 | 0.018519431 |
| CCP110 | -1.071782586 | 5.173325184 | 0.01110413 | 0.031487612 |
| NDFIP2 | -1.073138133 | 5.463515803 | 0.005350504 | 0.016958392 |
| LOC106557719 | -1.074703871 | 4.128132462 | 0.016061592 | 0.042727738 |
| BIRC2 | -1.07511794 | 5.364913757 | 0.004479414 | 0.014574739 |
| RIPK2 | -1.076516692 | 4.423343205 | 0.011115921 | 0.031507138 |
| UBXN2B | -1.079500254 | 5.794958262 | 0.007691643 | 0.023117596 |
| DLEC1 | -1.079846486 | 6.666901125 | 0.005949097 | 0.018573147 |
| LOC102156733 | -1.080051665 | 5.236075872 | 0.002175015 | 0.007913227 |
| RIT1 | -1.080422441 | 5.415844324 | 0.004820177 | 0.015506561 |
| ATP6V0D1 | -1.080672244 | 5.780777967 | 0.003792643 | 0.012702511 |
| PPT1 | -1.08436514 | 5.999980178 | 0.013017499 | 0.035974162 |
| PLEKHO2 | -1.086946642 | 6.467563136 | 0.004586799 | 0.014881689 |
| ALPK1 | -1.089032989 | 4.919798363 | 0.006770954 | 0.02074454 |

| | | | | |
|---|---|---|---|---|
| TRIM35 | -1.089241486 | 5.366837999 | 0.007595178 | 0.022878428 |
| MTMR6 | -1.090504743 | 6.10733874 | 0.01284665 | 0.035586074 |
| CDH23 | -1.091090281 | 5.31237148 | 0.015800949 | 0.042184524 |
| LOC488190 | -1.091752462 | 3.590589487 | 0.017759608 | 0.046301453 |
| MIR8885 | -1.092307986 | 3.035905067 | 0.018631519 | 0.0481059 |
| PAN3 | -1.092484459 | 7.113791464 | 0.008366264 | 0.024801263 |
| TMPPE | -1.092674271 | 3.903825509 | 0.01383843 | 0.037902223 |
| SNAP23 | -1.094448225 | 4.910939415 | 0.011385088 | 0.032132962 |
| CHD7 | -1.09566576 | 6.764218303 | 0.01842085 | 0.047664169 |
| LOC111091026 | -1.09603985 | 4.99152211 | 0.004210841 | 0.013840092 |
| PDE12 | -1.096521014 | 4.308746165 | 0.007432398 | 0.022466161 |
| SLC25A38 | -1.097693045 | 4.264264453 | 0.011958914 | 0.033460565 |
| CHML | -1.098291048 | 6.098248105 | 0.018500265 | 0.047841099 |
| MFSD9 | -1.100317671 | 3.567505358 | 0.008081909 | 0.024103375 |
| RMND5A | -1.100521545 | 5.928514098 | 0.007683618 | 0.023096678 |
| OXR1 | -1.107334069 | 5.877321831 | 0.017056193 | 0.044839747 |
| BICRAL | -1.107898505 | 6.236002838 | 0.004571052 | 0.014835043 |
| CTPS1 | -1.109489843 | 4.842614033 | 0.008800728 | 0.025887374 |
| LOC119876977 | -1.111295251 | 4.89870398 | 0.008334121 | 0.024719502 |
| ABCB7 | -1.112783687 | 4.923306555 | 0.008400324 | 0.024888614 |
| IER5 | -1.114457044 | 4.955814834 | 0.015784279 | 0.04215548 |
| RAB8A | -1.115668395 | 4.960524838 | 0.003974725 | 0.013226502 |
| TOPORS | -1.115977371 | 5.837593777 | 0.004009573 | 0.013315886 |
| IQGAP1 | -1.116322664 | 9.333429044 | 0.011179318 | 0.031642857 |
| GADD45B | -1.117197988 | 5.264660478 | 0.017852646 | 0.046499266 |
| JMJD6 | -1.117674035 | 4.789127239 | 0.006768067 | 0.020741561 |
| TSC22D3 | -1.117692523 | 7.178439815 | 0.008432142 | 0.024972645 |
| KCNJ13 | -1.118189601 | 2.678982452 | 0.015564431 | 0.041681442 |
| SPSB2 | -1.119082895 | 2.731172005 | 0.016480842 | 0.043586246 |
| ATF7IP | -1.120182751 | 7.763553173 | 0.005264958 | 0.016711694 |
| WDFY2 | -1.120341663 | 6.544305687 | 0.005135124 | 0.01637633 |
| ABHD5 | -1.122417657 | 4.766091546 | 0.011603061 | 0.032620722 |
| AFTPH | -1.122742664 | 5.781682517 | 0.003024428 | 0.010482716 |
| LOC102151385 | -1.123216931 | 4.383291033 | 0.004702059 | 0.015185177 |
| LTA4H | -1.124292948 | 5.487920121 | 0.002331985 | 0.008374668 |
| TMEM53 | -1.124973678 | 2.754692209 | 0.016690652 | 0.04403902 |
| LOC111098391 | -1.126358121 | 4.19080492 | 0.015605549 | 0.041745133 |
| SLC12A6 | -1.128294094 | 6.703482109 | 0.003987888 | 0.013256057 |
| CERT1 | -1.128818597 | 5.8567214 | 0.00359201 | 0.012120449 |

| | | | | |
|---|---|---|---|---|
| ARFIP1 | -1.131058863 | 5.481796647 | 0.001735567 | 0.006515667 |
| SLC49A4 | -1.131187031 | 5.670947311 | 0.004898459 | 0.015725716 |
| NFKB1 | -1.13156027 | 6.15708303 | 0.005628285 | 0.017711894 |
| ZBTB37 | -1.1331164 | 6.229914054 | 0.008301202 | 0.024635348 |
| CLASP1 | -1.134035899 | 7.224272292 | 0.003999824 | 0.013289619 |
| C1GALT1 | -1.134838706 | 5.234934239 | 0.016576847 | 0.043787845 |
| USP15 | -1.137460256 | 5.792935792 | 0.003698935 | 0.012436707 |
| ACSS2 | -1.138481271 | 6.590720124 | 0.010641988 | 0.030386631 |
| ZC3HAV1 | -1.139365061 | 6.827429112 | 0.004437084 | 0.014467418 |
| LOC111098987 | -1.140477569 | 5.902033007 | 0.002418112 | 0.008643757 |
| LOC111097376 | -1.140963267 | 7.230189863 | 0.003945034 | 0.013133753 |
| LOC119871892 | -1.144116537 | 4.095100116 | 0.008282814 | 0.024587511 |
| CSNK1G1 | -1.144887359 | 6.31374881 | 0.003069028 | 0.010616927 |
| LOC119872212 | -1.145481901 | 3.044688408 | 0.016643719 | 0.043947285 |
| TDG | -1.145532697 | 4.836538703 | 0.005868036 | 0.018357327 |
| RC3H1 | -1.146571718 | 7.020770252 | 0.003485415 | 0.011825204 |
| PGLS | -1.148711747 | 4.461904851 | 0.007786673 | 0.023361098 |
| MICU1 | -1.149501167 | 5.238795731 | 0.002885867 | 0.01008309 |
| PSAP | -1.149676531 | 9.741690595 | 0.015558295 | 0.04167531 |
| LIN52 | -1.151176167 | 4.081204915 | 0.006664108 | 0.020446094 |
| PCGF5 | -1.151618279 | 6.98624306 | 0.013909659 | 0.038068448 |
| UPP1 | -1.153183773 | 6.837429764 | 0.01348231 | 0.03702041 |
| SLC41A3 | -1.156830009 | 4.313106425 | 0.009571218 | 0.027791667 |
| PPP2R3C | -1.156915991 | 3.781887867 | 0.006141671 | 0.019078176 |
| BNIP3L | -1.157254435 | 6.795785184 | 0.01723093 | 0.045173092 |
| LOC119867740 | -1.157669953 | 6.470075418 | 0.016344392 | 0.043320534 |
| RBM22 | -1.158870874 | 5.731125126 | 0.001742121 | 0.006536876 |
| LPGAT1 | -1.159327739 | 6.666747135 | 0.011694906 | 0.03284802 |
| REPS1 | -1.160122471 | 5.911129262 | 0.00229155 | 0.008263689 |
| CYREN | -1.160711027 | 5.651371608 | 0.005268978 | 0.016722002 |
| SRSF5 | -1.161034868 | 7.271318128 | 0.006737515 | 0.020656692 |
| EXOSC1 | -1.161381116 | 4.136436178 | 0.011776992 | 0.033045411 |
| MXD3 | -1.161805135 | 4.80462306 | 0.002886368 | 0.010083213 |
| LOC609365 | -1.163250607 | 8.939866683 | 0.007935312 | 0.023747862 |
| TLK1 | -1.163409513 | 6.668143931 | 0.003045122 | 0.010546011 |
| LOC119871953 | -1.164274654 | 3.519281084 | 0.004697508 | 0.015172739 |
| COA6 | -1.164569117 | 3.118702122 | 0.009199016 | 0.026891055 |
| GNE | -1.16473469 | 5.496825037 | 0.003610077 | 0.012177617 |
| DENND3 | -1.165125768 | 6.298001007 | 0.018745659 | 0.048343012 |

| | | | | |
|---|---|---|---|---|
| EML5 | -1.165592922 | 4.888538508 | 0.010524356 | 0.030099547 |
| CEP120 | -1.165683426 | 7.018255765 | 0.003412568 | 0.011606863 |
| FCHSD1 | -1.166965884 | 5.239933708 | 0.006498611 | 0.02001768 |
| LOC119868892 | -1.167476886 | 5.004111383 | 0.004742569 | 0.015293214 |
| MTMR14 | -1.168007664 | 4.884494913 | 0.006537139 | 0.020113488 |
| PRMT9 | -1.168359705 | 4.630736083 | 0.008002273 | 0.02392843 |
| DCUN1D4 | -1.168578259 | 5.852410491 | 0.004850642 | 0.015595307 |
| UHRF1BP1L | -1.168813929 | 6.125292771 | 0.003112991 | 0.010744999 |
| PRELID1 | -1.169927191 | 5.434077223 | 0.002962607 | 0.010304699 |
| RGS19 | -1.170224274 | 5.905836284 | 0.019342262 | 0.049645293 |
| POU2F1 | -1.170876869 | 6.462402269 | 0.003718688 | 0.012493432 |
| MADD | -1.171694419 | 6.077771095 | 0.002997206 | 0.010403337 |
| ERBIN | -1.171772672 | 8.169254519 | 0.016019297 | 0.042657111 |
| LOC119877503 | -1.171834874 | 3.260811052 | 0.006817836 | 0.020870499 |
| RNF138 | -1.17209509 | 4.202676709 | 0.004856299 | 0.015610156 |
| PAPOLG | -1.172714896 | 4.958021699 | 0.005430041 | 0.01718284 |
| LIMK2 | -1.174036532 | 5.750537553 | 0.002527456 | 0.008987294 |
| M6PR | -1.174549237 | 6.683631678 | 0.006954649 | 0.021223331 |
| RNASEH2B | -1.175356152 | 3.968115543 | 0.008522254 | 0.025194764 |
| BCAP29 | -1.175703741 | 4.529319495 | 0.01335444 | 0.036748448 |
| PHTF2 | -1.179440253 | 4.888714665 | 0.014545334 | 0.039464324 |
| ATP2B1 | -1.180818659 | 7.224111849 | 0.009264539 | 0.027052313 |
| FOXO4 | -1.180868255 | 3.980605903 | 0.007589684 | 0.022865707 |
| RAB24 | -1.181919327 | 5.562056034 | 0.001534289 | 0.005880219 |
| PTEN | -1.182163714 | 7.339568772 | 0.005419421 | 0.017159258 |
| EHD1 | -1.184033852 | 6.360825235 | 0.0038475 | 0.012859282 |
| ETV3 | -1.185329989 | 5.999490973 | 0.001935787 | 0.00716066 |
| TAPT1 | -1.187689221 | 5.09872346 | 0.00408679 | 0.013504041 |
| PIGB | -1.188869072 | 4.816489498 | 0.007085259 | 0.02154909 |
| NOL4L | -1.189375969 | 5.931060004 | 0.009422859 | 0.027434393 |
| TRIM14 | -1.189416998 | 5.74932403 | 0.007108895 | 0.021611879 |
| ARMC8 | -1.189466254 | 4.954162649 | 0.002400368 | 0.008593191 |
| DTX3L | -1.189999382 | 5.486356457 | 0.012542758 | 0.034855647 |
| PPM1M | -1.19018077 | 5.050953961 | 0.003991883 | 0.013267304 |
| SRI | -1.191920975 | 6.115400182 | 0.006968659 | 0.021254116 |
| IRF2 | -1.1930305 | 6.314347758 | 0.008349093 | 0.024753746 |
| MON1B | -1.193469801 | 5.796116179 | 0.001881795 | 0.006980001 |
| LOC119872806 | -1.193879401 | 4.641099275 | 0.009913572 | 0.028647578 |
| LOC111096394 | -1.197894555 | 2.742967587 | 0.008909703 | 0.026158223 |

| | | | | |
|---|---|---|---|---|
| ESD | -1.198163077 | 4.307795193 | 0.003889111 | 0.012977476 |
| ATP10A | -1.199863626 | 4.836819289 | 0.010335138 | 0.029652296 |
| ADAP1 | -1.200351111 | 5.043348442 | 0.003364879 | 0.011464711 |
| CPEB2 | -1.201036366 | 6.830503346 | 0.013974133 | 0.038211127 |
| KLF2 | -1.202331359 | 7.247354703 | 0.008535019 | 0.02522906 |
| LOC111097452 | -1.203766355 | 4.766659834 | 0.003558923 | 0.012021904 |
| TBC1D1 | -1.205071674 | 6.580102879 | 0.006605313 | 0.020291559 |
| PPP1CA | -1.205324575 | 5.731840537 | 0.000868753 | 0.003629936 |
| POP5 | -1.207914413 | 2.720941501 | 0.01259289 | 0.03496358 |
| TMBIM4 | -1.210197581 | 4.900373472 | 0.006876079 | 0.021013095 |
| LOC119873249 | -1.212375458 | 4.273766165 | 0.007009537 | 0.021360755 |
| APMAP | -1.212695351 | 5.407611124 | 0.01648681 | 0.043596711 |
| ITPR2 | -1.213188422 | 7.667158879 | 0.019364091 | 0.049695291 |
| RBM38 | -1.213405615 | 5.420225478 | 0.008884673 | 0.026098877 |
| GNG5 | -1.213598311 | 5.41534567 | 0.005176465 | 0.016483914 |
| IRAK4 | -1.213793779 | 4.744001726 | 0.005215476 | 0.01657891 |
| TPM3 | -1.214010883 | 7.911528932 | 0.00148186 | 0.005700452 |
| PPM1L | -1.215256555 | 4.157068537 | 0.0186383 | 0.048117675 |
| LARP1B | -1.21824512 | 4.803018871 | 0.006324243 | 0.019561199 |
| SUSD6 | -1.218641279 | 6.979774796 | 0.004231331 | 0.013888481 |
| PPP2R5A | -1.218773239 | 6.442596518 | 0.007570471 | 0.022821504 |
| LOC119876629 | -1.219352263 | 6.26136904 | 0.010822273 | 0.03080929 |
| YWHAH | -1.219760802 | 6.981198147 | 0.015129723 | 0.040753975 |
| ERAP1 | -1.220296159 | 6.464092147 | 0.007452582 | 0.02252403 |
| PLA1A | -1.220979986 | 4.122376859 | 0.005540969 | 0.017475177 |
| RNF19B | -1.222357975 | 5.15769787 | 0.004252592 | 0.013945594 |
| TRA2B | -1.222518033 | 6.3551231 | 0.001805921 | 0.006752355 |
| PGM2 | -1.222648247 | 5.569827512 | 0.00230898 | 0.00831133 |
| GSKIP | -1.222954242 | 4.676789906 | 0.00194188 | 0.007180747 |
| PFKFB4 | -1.223947349 | 4.308195896 | 0.004087775 | 0.013505237 |
| MOB3A | -1.226836933 | 5.546491763 | 0.007732183 | 0.023213686 |
| CDKN2AIP | -1.227693257 | 4.554120156 | 0.005264376 | 0.016711694 |
| LOC119874198 | -1.229159679 | 7.026630706 | 0.005849884 | 0.018310882 |
| NIPBL | -1.229725154 | 7.918236479 | 0.002448258 | 0.008738651 |
| B3GLCT | -1.231164746 | 5.378975259 | 0.001530214 | 0.005866679 |
| EIF2AK1 | -1.231216586 | 4.20611093 | 0.004516108 | 0.014674303 |
| TLE3 | -1.231394181 | 7.26301488 | 0.002070713 | 0.007588974 |
| PIGBOS1 | -1.234723055 | 2.637937351 | 0.005951243 | 0.01857717 |
| TRIAP1 | -1.236846273 | 4.538205882 | 0.003523853 | 0.011931565 |

| | | | | |
|---|---|---|---|---|
| GRB2 | -1.237499159 | 6.270726837 | 0.002049191 | 0.007524925 |
| PLCXD2 | -1.23756103 | 2.690684699 | 0.01821893 | 0.047231923 |
| LOC102156251 | -1.239553382 | 2.461844461 | 0.016840154 | 0.044352492 |
| UHRF2 | -1.241422637 | 5.597322502 | 0.004732263 | 0.015266798 |
| LOC119869553 | -1.241568615 | 3.408938085 | 0.014692608 | 0.039804111 |
| LOC119864442 | -1.241598332 | 2.328542692 | 0.01724104 | 0.045183201 |
| ARHGEF6 | -1.242473871 | 6.53533557 | 0.01007392 | 0.029045107 |
| H3-3B | -1.242493058 | 8.31244384 | 0.005218997 | 0.01658767 |
| LAMTOR3 | -1.242684077 | 4.47639063 | 0.007043008 | 0.02145069 |
| LOC111095140 | -1.243301824 | 6.187399731 | 0.004055513 | 0.013429356 |
| LOC111093328 | -1.24332571 | 2.450112977 | 0.015845341 | 0.042276999 |
| RTN3 | -1.243965007 | 6.75779935 | 0.008100073 | 0.024144257 |
| LOC119866174 | -1.244430191 | 2.850639003 | 0.010386492 | 0.029768421 |
| ACTR3 | -1.244607968 | 7.633125951 | 0.003529708 | 0.011941229 |
| LOC100682670 | -1.245372899 | 4.276631449 | 0.013855633 | 0.037939753 |
| CAMKMT | -1.245791613 | 2.623336578 | 0.018333368 | 0.047477485 |
| ADSL | -1.2465334 | 3.46586742 | 0.010150372 | 0.029226654 |
| TAF5L | -1.248064535 | 4.361843453 | 0.000757411 | 0.003219963 |
| STK38 | -1.248121426 | 7.466589964 | 0.014156688 | 0.038598301 |
| WNT8A | -1.248855138 | 2.758031421 | 0.013023191 | 0.035983122 |
| MAP3K3 | -1.251417167 | 6.585344562 | 0.000940103 | 0.003885364 |
| LOC102154889 | -1.251755669 | 3.29852558 | 0.006879307 | 0.021014126 |
| ARRDC2 | -1.253177687 | 5.282378915 | 0.011720201 | 0.032903098 |
| MFSD5 | -1.253694184 | 4.192710544 | 0.007536302 | 0.022732675 |
| IL10RB | -1.253997168 | 5.525443294 | 0.001545624 | 0.00591843 |
| PPP4R3A | -1.256323239 | 6.800489652 | 0.002290588 | 0.008261593 |
| GLUL | -1.256526192 | 8.533812005 | 0.004311172 | 0.014107812 |
| ATG3 | -1.258076396 | 6.332662802 | 0.005101434 | 0.016280868 |
| ARL5B | -1.260078159 | 6.025801119 | 0.014943434 | 0.040332383 |
| VPS26B | -1.260359081 | 5.648179431 | 0.00044922 | 0.002057033 |
| RFFL | -1.260410641 | 6.465537056 | 0.0012813 | 0.005058217 |
| MAPRE1 | -1.261541777 | 5.84130091 | 0.000444096 | 0.002037015 |
| ANTXR2 | -1.262912908 | 7.652386378 | 0.009158203 | 0.026793432 |
| LOC485601 | -1.264286482 | 4.78551591 | 0.016816686 | 0.044306835 |
| TACC3 | -1.264743318 | 4.451836404 | 0.002834358 | 0.009930333 |
| LOC611178 | -1.265848793 | 3.935417433 | 0.002127242 | 0.00776558 |
| ATAD2B | -1.266990743 | 6.246085719 | 0.006178231 | 0.019169764 |
| FMO5 | -1.267960327 | 3.061425653 | 0.00579228 | 0.018172567 |
| HK1 | -1.273032505 | 6.468918831 | 0.004079745 | 0.013488986 |

| | | | | |
|---|---|---|---|---|
| GPN2 | -1.274079491 | 4.329362264 | 0.001442829 | 0.005576041 |
| PIAS1 | -1.275070149 | 6.678000517 | 0.000991653 | 0.004065152 |
| SAP30 | -1.27561984 | 3.274855251 | 0.005407274 | 0.017130806 |
| TRIM24 | -1.276551244 | 5.184319361 | 0.002898243 | 0.010114914 |
| FAS | -1.276603363 | 5.023745353 | 0.007835948 | 0.023479677 |
| SEMA7A | -1.276801694 | 6.142249529 | 0.019032478 | 0.048937109 |
| ZNF296 | -1.277372964 | 3.450277099 | 0.008015696 | 0.023955347 |
| LOC111095929 | -1.27753974 | 1.329724942 | 0.011116831 | 0.031507138 |
| KCTD12 | -1.277602107 | 9.187968327 | 0.004466454 | 0.014541165 |
| MAPKAPK3 | -1.278620965 | 5.427914485 | 0.00402156 | 0.013345472 |
| HIVEP1 | -1.279275395 | 6.542213571 | 0.003546151 | 0.011984363 |
| LOC119866853 | -1.279995115 | 1.635736965 | 0.015231373 | 0.040956519 |
| USP3 | -1.282765055 | 6.203101843 | 0.000876426 | 0.003659177 |
| UBE2D3 | -1.282987044 | 7.417411593 | 0.001418728 | 0.005493672 |
| TAF8 | -1.28329427 | 4.552340057 | 0.004065743 | 0.013448846 |
| LOC111097062 | -1.285571775 | 2.416191879 | 0.011602846 | 0.032620722 |
| ST3GAL6 | -1.286707705 | 7.245720038 | 0.013866739 | 0.037965365 |
| GIT2 | -1.288814664 | 7.641065364 | 0.00205226 | 0.007533088 |
| PPP2R5C | -1.289466949 | 7.106433918 | 0.000500142 | 0.002259188 |
| MSN | -1.290060997 | 8.617573932 | 0.00237917 | 0.008522936 |
| ARPC5 | -1.291250509 | 7.41428643 | 0.001439344 | 0.005563564 |
| MAPK6 | -1.291654868 | 5.630645837 | 0.00768354 | 0.023096678 |
| PELI1 | -1.292414264 | 5.900423579 | 0.013078755 | 0.036113627 |
| CAB39 | -1.292555361 | 7.003460044 | 0.001038487 | 0.004225351 |
| MSL2 | -1.296458759 | 6.202926538 | 0.001172106 | 0.004692973 |
| LOC119866146 | -1.296732069 | 3.126839368 | 0.00822847 | 0.024452986 |
| ANKRD49 | -1.298810186 | 3.070142235 | 0.006639891 | 0.02038045 |
| ZNF800 | -1.299634994 | 5.757155132 | 0.002042097 | 0.007502689 |
| LOC102156449 | -1.301455352 | 1.587283618 | 0.01028395 | 0.029528888 |
| STK24 | -1.301811162 | 6.45042959 | 0.00182807 | 0.006826286 |
| LOC119877428 | -1.30254337 | 8.279362143 | 0.017198898 | 0.045116403 |
| PDZD8 | -1.305695168 | 6.831278544 | 0.000447667 | 0.002051659 |
| NIBAN1 | -1.305699642 | 7.298428277 | 0.004860935 | 0.015619133 |
| PRKACB | -1.307845943 | 6.32370597 | 0.006398458 | 0.019768181 |
| LOC119864149 | -1.308050497 | 5.521511161 | 0.002622797 | 0.009280634 |
| PTRH2 | -1.308108481 | 2.964196655 | 0.0060933 | 0.01894421 |
| RSRP1 | -1.308628493 | 5.158594919 | 0.002150817 | 0.007839665 |
| OSBPL11 | -1.309259044 | 6.296728566 | 0.000716249 | 0.003070276 |
| PIP4K2A | -1.311529526 | 6.94774221 | 0.001224946 | 0.004866773 |

| | | | | |
|---|---|---|---|---|
| HDAC1 | -1.312223456 | 2.908085327 | 0.004200392 | 0.01381832 |
| UBALD2 | -1.31251263 | 6.314632182 | 0.002304631 | 0.008298433 |
| LOC106559951 | -1.312602916 | 3.868325487 | 0.004721795 | 0.015237562 |
| MAP3K5 | -1.314039275 | 5.027366797 | 0.003078939 | 0.010639324 |
| C30H15orf65 | -1.314206733 | 0.896801818 | 0.018268644 | 0.047343816 |
| CCNL1 | -1.314326326 | 6.969338002 | 0.00305195 | 0.010564592 |
| TLN1 | -1.316495339 | 9.90767925 | 0.000819423 | 0.003446036 |
| FAM217B | -1.319719297 | 3.280712509 | 0.007024462 | 0.021400004 |
| LOC119877482 | -1.321931826 | 2.51514804 | 0.01402005 | 0.038322177 |
| LOC102152339 | -1.323158053 | 2.027965404 | 0.018428674 | 0.047678722 |
| NCEH1 | -1.32524108 | 5.03155978 | 0.002039625 | 0.007494877 |
| LOC119864182 | -1.325587235 | 2.876022295 | 0.016354688 | 0.043342523 |
| TMC4 | -1.326801186 | 2.231064703 | 0.013764953 | 0.037718714 |
| ZNF710 | -1.326969693 | 4.744096291 | 0.004147193 | 0.013674445 |
| SLC49A3 | -1.328018183 | 3.918669501 | 0.011687315 | 0.032832038 |
| REPS2 | -1.333086238 | 3.025455375 | 0.014477203 | 0.039328707 |
| LOC111098080 | -1.335646414 | 4.213257904 | 0.009265414 | 0.027052313 |
| PAK2 | -1.335704021 | 7.161655052 | 0.002535616 | 0.009009363 |
| UBR2 | -1.337254815 | 7.077819028 | 0.000411241 | 0.001912215 |
| LOC119876321 | -1.337684802 | 2.978332314 | 0.004317412 | 0.014121833 |
| ZBTB11 | -1.338634801 | 5.719941271 | 0.000315777 | 0.001522543 |
| LOC119871820 | -1.339042277 | 2.709195783 | 0.008270868 | 0.024558777 |
| LOC111097597 | -1.339245107 | 2.736772422 | 0.017461216 | 0.045649811 |
| LOC102156144 | -1.340232148 | 4.061797874 | 0.011595269 | 0.032615738 |
| STRADB | -1.340308633 | 4.132182891 | 0.007207579 | 0.021875069 |
| RAC1 | -1.340332876 | 6.671075948 | 0.002688257 | 0.009476674 |
| CYTH1 | -1.340938649 | 7.337648785 | 0.007402212 | 0.022390529 |
| LOC111098213 | -1.345428578 | 2.308014338 | 0.0136928 | 0.037527069 |
| ATP2C1 | -1.34553547 | 6.270367524 | 0.002876005 | 0.010056739 |
| LOC119865644 | -1.346606455 | 5.841473735 | 0.000794749 | 0.003355659 |
| TXNDC16 | -1.349231624 | 4.977963259 | 0.000827616 | 0.003478169 |
| LOC119877473 | -1.349413817 | 3.75127809 | 0.006547603 | 0.020137108 |
| LOC111095448 | -1.349748176 | 1.840875267 | 0.013488493 | 0.037028007 |
| HIPK3 | -1.349755895 | 8.701791726 | 0.000750649 | 0.003198743 |
| LOC119867171 | -1.350096755 | 1.445730444 | 0.009566775 | 0.027782488 |
| MAN2B1 | -1.350827584 | 8.048971908 | 0.001732642 | 0.006506941 |
| TMBIM6 | -1.352763278 | 7.957710065 | 0.006189026 | 0.019197762 |
| SHKBP1 | -1.354329139 | 6.08380019 | 0.002137523 | 0.007797767 |
| LOC111098937 | -1.355031875 | 2.302735425 | 0.018141807 | 0.047093949 |

| | | | | |
|---|---|---|---|---|
| RAP1GDS1 | -1.356274199 | 5.582263365 | 0.004874125 | 0.01565687 |
| BLOC1S6 | -1.358890102 | 4.861042699 | 0.003194131 | 0.010977862 |
| LOC119877492 | -1.360482973 | 1.632244361 | 0.012182891 | 0.033986329 |
| LOC111097442 | -1.362347035 | 4.599324011 | 0.011092717 | 0.031463482 |
| TRAFD1 | -1.363260895 | 7.384995034 | 0.000898127 | 0.003738255 |
| SRGAP3 | -1.365706684 | 5.577456022 | 0.015444779 | 0.041449458 |
| OTULINL | -1.367387029 | 4.97573648 | 0.003651639 | 0.012294854 |
| LOC102153082 | -1.36867085 | 2.469242 | 0.007163187 | 0.021752523 |
| LIG4 | -1.370388536 | 4.200302215 | 0.002936526 | 0.010232047 |
| LOC119865810 | -1.370669019 | 1.6894086 | 0.018650093 | 0.048136652 |
| CYBA | -1.371242325 | 5.288787393 | 0.003561104 | 0.012027397 |
| R3HDM4 | -1.372020756 | 5.354105547 | 0.012039136 | 0.033654611 |
| UBN1 | -1.37260736 | 6.823699849 | 0.000464916 | 0.002118287 |
| MYCBP2 | -1.373234618 | 8.677489711 | 0.001696742 | 0.006398728 |
| LOC111093051 | -1.373605865 | 4.887479435 | 0.003201543 | 0.010996361 |
| ASTE1 | -1.37432082 | 3.855580619 | 0.004803818 | 0.015460818 |
| LOC119864736 | -1.374602956 | 6.141030908 | 0.001256295 | 0.004972183 |
| ARPC2 | -1.376162659 | 7.666047964 | 0.000325269 | 0.001559983 |
| ZFP36 | -1.376895728 | 8.021480789 | 0.016638783 | 0.043939605 |
| BYSL | -1.37723171 | 4.327375127 | 0.000950648 | 0.003922961 |
| MCL1 | -1.377478076 | 8.124027711 | 0.000344967 | 0.001640294 |
| RNF130 | -1.3788956 | 6.247775894 | 0.002645804 | 0.009351355 |
| NIN | -1.379115693 | 7.585183807 | 0.003013917 | 0.010449628 |
| PLEKHM3 | -1.380362278 | 6.115888092 | 0.000343851 | 0.001636066 |
| GREB1L | -1.380407011 | 5.350995784 | 0.005912287 | 0.018476856 |
| LOC119872924 | -1.381228951 | 3.55717225 | 0.007307704 | 0.022147452 |
| ACOX2 | -1.381908956 | 2.141352043 | 0.018131441 | 0.047072677 |
| CAPZA1 | -1.383182866 | 7.19442067 | 0.000532635 | 0.002383111 |
| GPR146 | -1.383237184 | 5.755534269 | 0.001864776 | 0.006934675 |
| LOC119867996 | -1.383724351 | 1.042058801 | 0.015793789 | 0.042170603 |
| CLIC5 | -1.383765846 | 5.269940981 | 0.017486594 | 0.045699619 |
| LOC119877771 | -1.383867131 | 3.579241511 | 0.004471757 | 0.014552011 |
| LOC111097734 | -1.386139103 | 1.629814555 | 0.008127406 | 0.024209082 |
| FBXO33 | -1.386729715 | 4.211256195 | 0.0010177 | 0.004155352 |
| LOC119864621 | -1.388752311 | 2.278567122 | 0.00748538 | 0.022604243 |
| KRTCAP3 | -1.389290611 | 2.209556299 | 0.006285413 | 0.019454978 |
| STAMBPL1 | -1.390708318 | 4.338907727 | 0.005820605 | 0.01825342 |
| ARRDC4 | -1.391366598 | 5.477211841 | 0.005836379 | 0.018279167 |
| BCL2L11 | -1.392611142 | 5.466571258 | 0.000716251 | 0.003070276 |

| | | | | |
|---|---|---|---|---|
| LOC102154256 | -1.392969103 | 0.838952524 | 0.014735621 | 0.039895706 |
| NHSL2 | -1.395299596 | 6.455166647 | 0.001063644 | 0.004316919 |
| CFAP94 | -1.395381086 | 3.976245122 | 0.004599623 | 0.014916597 |
| TIGD4 | -1.395419526 | 3.456293334 | 0.00510884 | 0.016302104 |
| LOC102152829 | -1.395899467 | 5.058150875 | 0.000551277 | 0.002452848 |
| FAM110A | -1.397751364 | 2.750304496 | 0.015552532 | 0.041665021 |
| LOC119868480 | -1.398256283 | 3.712666694 | 0.013121552 | 0.036213345 |
| CTSH | -1.399310297 | 6.548075884 | 0.00123103 | 0.004886466 |
| OAS3 | -1.400449224 | 5.726417298 | 0.00681608 | 0.020870499 |
| CDADC1 | -1.401856255 | 5.303448734 | 0.001356677 | 0.005304574 |
| H2AC8 | -1.402251894 | 2.735030343 | 0.007505804 | 0.022656448 |
| NADK | -1.402494565 | 7.527509137 | 0.000689441 | 0.0029724 |
| LOC111094869 | -1.404565808 | 1.693895342 | 0.015461862 | 0.041483652 |
| PIGA | -1.404926978 | 3.483910567 | 0.001861066 | 0.006926821 |
| SAMHD1 | -1.405464027 | 6.908101947 | 0.005118879 | 0.016331732 |
| AKT3 | -1.406553015 | 6.538257211 | 0.001867683 | 0.006941914 |
| ITGB1BP2 | -1.406748684 | 3.253593799 | 0.005834687 | 0.018277851 |
| ST3GAL5 | -1.406774949 | 5.085076058 | 0.005576961 | 0.017573348 |
| OTULIN | -1.40797242 | 4.902657521 | 0.0002226 | 0.00112601 |
| BRWD3 | -1.409177648 | 5.639409497 | 0.004223889 | 0.013874559 |
| LAT2 | -1.409231238 | 5.954632537 | 0.003911867 | 0.013042074 |
| PKD2L2 | -1.410423958 | 3.369655462 | 0.005410909 | 0.017139816 |
| ELMO1 | -1.411959047 | 7.116534012 | 0.004113882 | 0.013583207 |
| YOD1 | -1.412863605 | 4.932397662 | 0.003423608 | 0.011635542 |
| LOC119879075 | -1.414383928 | 1.053715738 | 0.016569371 | 0.043777641 |
| DENND11 | -1.415441677 | 5.22596728 | 0.000244778 | 0.001220398 |
| RAB11FIP1 | -1.415499968 | 5.695878005 | 0.007461847 | 0.022539458 |
| LOC111094566 | -1.41577741 | 0.058891163 | 0.016727993 | 0.044110693 |
| LRRC8C | -1.4166977 | 6.434703667 | 0.001872957 | 0.006953169 |
| DDA1 | -1.417265455 | 5.988898283 | 0.000577023 | 0.002551953 |
| LOC119876115 | -1.418615798 | 2.116389802 | 0.004328616 | 0.014156344 |
| OAS1 | -1.418641855 | 4.663676527 | 0.008710412 | 0.025668754 |
| LOC111097809 | -1.42135774 | 2.266163752 | 0.006308327 | 0.01951754 |
| LOC102153861 | -1.421990196 | 3.013993061 | 0.001219176 | 0.004850072 |
| PGAP2 | -1.422179464 | 4.534354107 | 0.000382013 | 0.001791683 |
| SLC37A3 | -1.422392729 | 4.291237214 | 0.001484392 | 0.005708166 |
| LOC119867335 | -1.422789954 | 1.148013158 | 0.018101907 | 0.047007265 |
| SGMS1 | -1.423768732 | 6.228149585 | 0.000480601 | 0.002179547 |
| LOC119866156 | -1.425324843 | 6.135143015 | 0.008033285 | 0.023995429 |

| | | | | |
|---|---|---|---|---|
| GRPEL2 | -1.426725098 | 3.984136462 | 0.00571758 | 0.017969418 |
| WDSUB1 | -1.427226165 | 4.165039522 | 0.002014388 | 0.007417233 |
| ATP6V0B | -1.42733094 | 4.682105201 | 0.001865683 | 0.006935669 |
| PPP2R1B | -1.427421937 | 6.142613052 | 0.000858348 | 0.003590615 |
| LOC111093849 | -1.428650831 | 3.220353614 | 0.013978483 | 0.038218198 |
| CBLN3 | -1.430635628 | 1.463804596 | 0.01628433 | 0.043208899 |
| LOC119877696 | -1.430679741 | 0.216062035 | 0.01340686 | 0.036836572 |
| LOC100683076 | -1.431048128 | 0.095601864 | 0.019372174 | 0.049704407 |
| CFLAR | -1.431756052 | 5.957812184 | 0.002712831 | 0.009558637 |
| MANBA | -1.43220787 | 5.16563796 | 0.004445161 | 0.014491574 |
| ADPRH | -1.433220354 | 5.132189005 | 0.000289449 | 0.001411299 |
| PARP12 | -1.433580982 | 5.23493242 | 0.001198397 | 0.004777936 |
| RFC4 | -1.434580239 | 4.094094413 | 0.001324375 | 0.005194435 |
| LOC111092890 | -1.435081752 | 4.445799831 | 0.010397483 | 0.029787793 |
| RBM48 | -1.435618978 | 3.680816219 | 0.000702659 | 0.003020975 |
| LOC102156637 | -1.436381014 | 2.123985609 | 0.01393009 | 0.038114738 |
| FGD4 | -1.436647141 | 5.668081498 | 0.000868358 | 0.003628985 |
| RASA3 | -1.437171796 | 7.662941579 | 0.001897402 | 0.007030672 |
| LOC119873868 | -1.438251754 | 0.148609146 | 0.015165005 | 0.040813505 |
| NAPEPLD | -1.438483924 | 4.251588249 | 0.001351729 | 0.005287137 |
| CD14 | -1.439355671 | 5.059162589 | 0.007200732 | 0.021857349 |
| INTS10 | -1.442628463 | 5.437658589 | 0.000432681 | 0.001993516 |
| SUPT4H1 | -1.443061377 | 3.799059483 | 0.002052524 | 0.007533088 |
| GBE1 | -1.444079169 | 6.246536951 | 0.000917595 | 0.0038127 |
| CAP1 | -1.444538553 | 8.008204098 | 0.000398895 | 0.001860395 |
| RAB8B | -1.4445628 | 7.227234944 | 0.001381802 | 0.005382423 |
| SLC15A4 | -1.445272467 | 5.786208226 | 0.000478941 | 0.002172929 |
| EIF4E3 | -1.446215752 | 5.164506976 | 0.000307114 | 0.001486397 |
| IFNGR1 | -1.447346716 | 6.707305924 | 0.001321704 | 0.005185606 |
| ATP6V0A1 | -1.447441158 | 6.725901491 | 0.00116523 | 0.004669108 |
| PPEF1 | -1.449183426 | 2.955542927 | 0.010239625 | 0.029421103 |
| PTPN1 | -1.449833238 | 6.972520861 | 6.67E-05 | 0.000402276 |
| AGAP2 | -1.450908591 | 5.4364864 | 0.003869873 | 0.012921238 |
| DHX34 | -1.45115521 | 5.543766856 | 0.000546517 | 0.002435672 |
| LOC102152432 | -1.451240248 | 2.175215709 | 0.008476948 | 0.025077926 |
| LOC102154363 | -1.451452727 | -0.054946322 | 0.014702873 | 0.039821966 |
| LNPEP | -1.451808184 | 6.455867589 | 0.00241888 | 0.008643757 |
| ZBTB24 | -1.452111521 | 6.33698854 | 0.000137923 | 0.000748613 |
| ARPC3 | -1.452457943 | 5.532930096 | 0.000123144 | 0.000682231 |

| | | | | |
|---|---|---|---|---|
| LOC119872026 | -1.452623901 | 0.189747451 | 0.011348808 | 0.032038911 |
| CSGALNACT2 | -1.453388686 | 6.432318852 | 0.000298329 | 0.001449706 |
| H2AC13 | -1.454173902 | 1.973417554 | 0.011592786 | 0.032612988 |
| LOC102155367 | -1.456873693 | 3.164582254 | 0.002895139 | 0.010107336 |
| KCNAB1 | -1.458614329 | 4.782859418 | 0.017044223 | 0.044813715 |
| LOC119877440 | -1.459169401 | 1.776634639 | 0.016454979 | 0.043533785 |
| PARP8 | -1.459677001 | 6.845185499 | 0.008511623 | 0.025166767 |
| LOC119876881 | -1.46128757 | 2.457077449 | 0.014837332 | 0.040110952 |
| IFNGR2 | -1.46155688 | 6.67992921 | 0.000888189 | 0.003703294 |
| MORN5 | -1.461600287 | 1.965393955 | 0.015148785 | 0.040788634 |
| NDEL1 | -1.46275644 | 6.159014038 | 0.000129486 | 0.00071119 |
| SH2D3C | -1.463560789 | 5.688182344 | 0.001210237 | 0.004820707 |
| MYOZ3 | -1.465080449 | 3.124820499 | 0.005823344 | 0.018254172 |
| MSL1 | -1.465547479 | 6.370656195 | 0.000452663 | 0.002070176 |
| ARHGAP24 | -1.465908235 | 5.16094905 | 0.007267833 | 0.022036337 |
| SMCHD1 | -1.4661799 | 7.26377322 | 0.005849172 | 0.018310882 |
| DUSP10 | -1.466859932 | 3.214974104 | 0.008191628 | 0.024363546 |
| NCOA3 | -1.467627927 | 7.294664256 | 0.000425092 | 0.001966073 |
| NMNAT3 | -1.468253877 | 2.818516816 | 0.00465952 | 0.015078458 |
| LOC119876447 | -1.468510191 | 3.388150132 | 0.004282707 | 0.01402525 |
| LOC119864213 | -1.46882201 | 4.332445947 | 0.004057788 | 0.013434612 |
| ANXA3 | -1.469303188 | 4.400941198 | 0.011255378 | 0.031810795 |
| LOC119872903 | -1.472396279 | 2.248477146 | 0.016395685 | 0.043435237 |
| IL23A | -1.472420762 | 3.216378207 | 0.00507011 | 0.016201644 |
| LOC111097453 | -1.472513376 | 1.461323412 | 0.018760971 | 0.048370988 |
| RBKS | -1.472993816 | 3.801081225 | 0.004845596 | 0.015581395 |
| FHL3 | -1.473426897 | 5.918828032 | 0.000751877 | 0.003202089 |
| INPP4A | -1.473782848 | 6.451695701 | 9.34E-05 | 0.000539122 |
| TAPBPL | -1.474236217 | 4.864170995 | 0.014535709 | 0.039457439 |
| TMEM184A | -1.475985645 | 1.515859712 | 0.010666831 | 0.030442749 |
| RNF13 | -1.478097443 | 5.821128429 | 0.001008635 | 0.004125333 |
| GPATCH3 | -1.47843394 | 3.930952649 | 0.000313641 | 0.001514266 |
| EDARADD | -1.47862129 | 3.465939817 | 0.005193291 | 0.016527783 |
| C11H9orf72 | -1.478680965 | 5.023935096 | 0.000904479 | 0.003761808 |
| LOC119864712 | -1.478706615 | 3.223997245 | 0.00228636 | 0.008251227 |
| LOC111091261 | -1.478967256 | 4.981623792 | 0.000721161 | 0.003087656 |
| HERC3 | -1.479342842 | 6.807562638 | 0.004124862 | 0.013611164 |
| NCOA1 | -1.479661302 | 6.651259081 | 0.000145888 | 0.000784185 |
| TET1 | -1.479915082 | 5.590670898 | 0.013223889 | 0.036449365 |

| | | | | |
|---|---|---|---|---|
| LOC119876396 | -1.481116968 | 0.720479698 | 0.010628651 | 0.030361771 |
| CASP7 | -1.48132345 | 4.871620994 | 8.78E-05 | 0.000511066 |
| APAF1 | -1.481742135 | 5.979391304 | 0.000886866 | 0.003698491 |
| LOC111096617 | -1.481878233 | 0.584877736 | 0.013285301 | 0.036590714 |
| CNOT6L | -1.482317411 | 6.692353359 | 0.010628328 | 0.030361771 |
| LOC119878095 | -1.482536942 | 1.981622969 | 0.003213499 | 0.011030435 |
| PARVB | -1.482836874 | 6.613795517 | 0.006972544 | 0.021262971 |
| SIAH2 | -1.483592678 | 4.635296808 | 0.002515694 | 0.008954284 |
| LOC102156147 | -1.486256196 | 1.826082982 | 0.004669611 | 0.015102892 |
| SLC12A9 | -1.487741647 | 4.745030077 | 0.000575301 | 0.002545631 |
| PYCARD | -1.489840811 | 3.806228184 | 0.011198198 | 0.031675614 |
| PRDM1 | -1.491056777 | 5.493714075 | 0.003911898 | 0.013042074 |
| LOC119869549 | -1.491586048 | 2.423757929 | 0.005453246 | 0.017233624 |
| LOC111096389 | -1.493742741 | 4.333388719 | 0.001257434 | 0.004975784 |
| TBK1 | -1.493852567 | 6.059120754 | 0.000123625 | 0.000684197 |
| PPP2CA | -1.494535804 | 5.273150271 | 0.000384495 | 0.001802155 |
| LOC111098815 | -1.496841069 | 4.818455442 | 0.001319364 | 0.005182052 |
| LOC119874426 | -1.4986535 | 1.052294671 | 0.013116893 | 0.036205098 |
| LOC119872488 | -1.498889153 | 2.083211246 | 0.004366308 | 0.014268848 |
| PLIN2 | -1.500035006 | 5.43235218 | 0.000184269 | 0.000956795 |
| ATG4D | -1.500623241 | 4.826243072 | 0.000303875 | 0.00147335 |
| PRKCH | -1.501090531 | 5.332598221 | 0.017944276 | 0.046676225 |
| GBP6 | -1.501328797 | 4.141203572 | 0.007625259 | 0.022959466 |
| PLXNA4 | -1.5043667 | 5.691310611 | 0.007725773 | 0.023197657 |
| MSRA | -1.504426802 | 3.718300412 | 0.004197405 | 0.013811954 |
| LOC106559017 | -1.505783812 | 1.013064635 | 0.011590915 | 0.032611955 |
| LOC102154188 | -1.506924749 | 3.214136018 | 0.00696685 | 0.02125159 |
| EVI2A | -1.507185554 | 4.288639286 | 0.002037439 | 0.007490657 |
| LACTB | -1.509903289 | 4.732582366 | 0.000604281 | 0.00265597 |
| MTF1 | -1.50994628 | 5.883024513 | 0.000120021 | 0.000667663 |
| LOC480043 | -1.510268064 | 6.675930153 | 0.000530038 | 0.002373944 |
| ADCY7 | -1.511392024 | 6.37967047 | 0.005229461 | 0.016613618 |
| LOC111096019 | -1.511606287 | 2.00043869 | 0.008834513 | 0.025976176 |
| JAK2 | -1.511716579 | 6.919754257 | 0.000317258 | 0.001529047 |
| LOC119868461 | -1.513235508 | 1.068685626 | 0.015921854 | 0.042439349 |
| MINDY2 | -1.513886523 | 6.317092616 | 0.001113698 | 0.004490639 |
| LOC608697 | -1.514063758 | 5.783341784 | 0.000136634 | 0.000743288 |
| LOC102153750 | -1.514077895 | 0.521688956 | 0.014721617 | 0.039867749 |
| LUZP6 | -1.514764515 | 6.438326492 | 0.003001224 | 0.010415617 |

| | | | | |
|---|---|---|---|---|
| LOC119871773 | -1.515554268 | 4.425700642 | 8.03E-05 | 0.000473488 |
| COL17A1 | -1.515665041 | 3.321540102 | 0.000615813 | 0.002698994 |
| ROMO1 | -1.517211915 | 3.236266726 | 0.002759913 | 0.009705599 |
| LOC119865885 | -1.517506511 | 4.299906363 | 0.001669626 | 0.006314048 |
| LOC111094073 | -1.520842204 | 2.492255026 | 0.017497792 | 0.045723372 |
| LOC111096201 | -1.52202259 | 2.527219155 | 0.016763174 | 0.044181962 |
| SYN1 | -1.522537857 | 3.520173137 | 0.019186128 | 0.049297089 |
| DLA88 | -1.523188901 | 8.857490288 | 0.016042233 | 0.042686719 |
| SLC31A2 | -1.52342241 | 3.583320064 | 0.000631666 | 0.002758989 |
| LOC102154490 | -1.525462516 | 2.947837672 | 0.005191699 | 0.016525142 |
| LOC119874202 | -1.529140305 | 3.731899685 | 0.000447875 | 0.002052176 |
| LOC612235 | -1.529714195 | 1.45462056 | 0.008646809 | 0.025521217 |
| LOC119877334 | -1.529934943 | 1.764203976 | 0.006258577 | 0.019385756 |
| NEDD9 | -1.530212035 | 6.209178368 | 0.001467043 | 0.005655499 |
| ACTB | -1.530443955 | 11.64667169 | 0.000175542 | 0.000918303 |
| LOC111090540 | -1.531228736 | 2.875107071 | 0.001160538 | 0.004653532 |
| CCDC38 | -1.531269021 | 2.476815413 | 0.00117736 | 0.004710526 |
| LOC119877066 | -1.533840346 | 1.264934052 | 0.007705725 | 0.023150288 |
| OSTF1 | -1.534275545 | 5.652287696 | 0.000205543 | 0.001053379 |
| TRIB1 | -1.535056271 | 6.528219058 | 0.006379246 | 0.019714445 |
| METTL9 | -1.535133222 | 5.95700904 | 0.000606761 | 0.002665246 |
| LIN54 | -1.535607045 | 4.423244237 | 0.000462387 | 0.002108418 |
| LOC119873225 | -1.537313876 | 3.967757314 | 0.000728872 | 0.003118207 |
| HIPK1 | -1.538142508 | 7.797145368 | 3.27E-05 | 0.000216886 |
| NUMB | -1.538752087 | 6.698612239 | 5.45E-05 | 0.000338316 |
| ATP2A3 | -1.541123266 | 6.039563257 | 0.008899233 | 0.026134564 |
| MTPN | -1.542656605 | 5.515741303 | 0.000478936 | 0.002172929 |
| CSK | -1.543319112 | 6.839874684 | 0.000783208 | 0.00331403 |
| LOC111093613 | -1.543526733 | 0.439454701 | 0.016864199 | 0.044410424 |
| TANK | -1.544067953 | 5.180495039 | 0.000899706 | 0.003744111 |
| YPEL5 | -1.545571249 | 6.542169794 | 0.000176608 | 0.000922539 |
| IRF1 | -1.547504453 | 6.563686031 | 0.0049528 | 0.015874302 |
| LOC102154412 | -1.547881313 | 0.357582757 | 0.008124105 | 0.024202575 |
| LOC119878065 | -1.552012912 | 6.156608772 | 0.000226884 | 0.001143812 |
| LOC119871449 | -1.552185052 | -0.536078145 | 0.01041177 | 0.029809021 |
| LOC111094480 | -1.553855579 | 2.729204084 | 0.006817907 | 0.020870499 |
| HSPA8 | -1.554159328 | 8.272144331 | 0.000445063 | 0.00204102 |
| RIOK3 | -1.555418356 | 6.052152198 | 7.94E-05 | 0.000469138 |
| LOC119869532 | -1.555622493 | 4.4502873 | 0.005208636 | 0.016562027 |

| | | | | |
|---|---|---|---|---|
| LOC106559997 | -1.556529144 | 3.882180188 | 0.018070389 | 0.046941961 |
| LOC102155278 | -1.557537105 | 5.498953259 | 0.000294313 | 0.001432757 |
| CRACR2A | -1.56051596 | 5.087944775 | 0.004961476 | 0.015890362 |
| CD47 | -1.561483302 | 5.456837188 | 0.000845651 | 0.003543659 |
| RBPJ | -1.563489719 | 7.259663726 | 0.000190155 | 0.000982183 |
| PTBP3 | -1.563944124 | 7.608155464 | 0.000543158 | 0.002423192 |
| LOC111090901 | -1.56464369 | 3.262845447 | 0.001395792 | 0.005426187 |
| LOC111095575 | -1.567272099 | 3.632068897 | 0.001632543 | 0.006191499 |
| PLCB2 | -1.567746885 | 6.326042002 | 0.010749603 | 0.030634601 |
| C9H17orf98 | -1.567903658 | -0.028330135 | 0.011732275 | 0.032928463 |
| LOC119877880 | -1.568135111 | 0.923405368 | 0.005453079 | 0.017233624 |
| LOC111090677 | -1.570238919 | 0.257848784 | 0.011510854 | 0.03243723 |
| PLEKHG6 | -1.570862517 | 3.672240849 | 0.003739257 | 0.012548917 |
| TMEM131L | -1.571637766 | 6.197798873 | 0.002432759 | 0.008687623 |
| GDE1 | -1.572181941 | 4.504327651 | 0.000967275 | 0.003977943 |
| LOC111090828 | -1.572561038 | -0.177297617 | 0.016296283 | 0.043235322 |
| ATP11B | -1.573998863 | 7.374549185 | 0.000399455 | 0.001862604 |
| HERC5 | -1.574910138 | 3.412884246 | 0.008285002 | 0.024590639 |
| LOC119876299 | -1.57550109 | 3.369328481 | 0.006852963 | 0.020948417 |
| TBC1D14 | -1.576118906 | 5.802103031 | 9.36E-05 | 0.000540009 |
| LOC119865898 | -1.57732658 | 5.55671785 | 0.004859823 | 0.015617874 |
| MTMR10 | -1.577888846 | 5.949000364 | 6.21E-05 | 0.000379286 |
| CCDC88B | -1.578102557 | 4.315822887 | 0.01082785 | 0.03081707 |
| OSGIN1 | -1.579387591 | 2.287101899 | 0.003413027 | 0.011606863 |
| LOC119875953 | -1.581083118 | 1.405381931 | 0.006675883 | 0.020479323 |
| NCOA2 | -1.58181519 | 6.990734564 | 7.50E-05 | 0.000447079 |
| LOC102151810 | -1.583431487 | 3.272741082 | 0.00811062 | 0.024169047 |
| LOC119866896 | -1.58355125 | 2.069810103 | 0.002405089 | 0.008607249 |
| LOC111098398 | -1.584405947 | 1.232788061 | 0.018007712 | 0.046796299 |
| FAM117B | -1.584619564 | 4.921771962 | 7.74E-05 | 0.000458998 |
| CCNYL1 | -1.584637566 | 5.324865422 | 0.01077296 | 0.030689059 |
| DOCK10 | -1.585485712 | 7.318863979 | 0.002774577 | 0.009750836 |
| P2RX7 | -1.585987602 | 4.252442489 | 0.010221294 | 0.029385697 |
| CD5 | -1.58636777 | 5.394637301 | 0.003740441 | 0.012550949 |
| LOC111091321 | -1.587435106 | 4.082315902 | 0.000331143 | 0.001582897 |
| LOC119864429 | -1.587643233 | 3.402909978 | 0.001508709 | 0.005793454 |
| ATP1B1 | -1.589168721 | 4.642373204 | 0.000719752 | 0.003082839 |
| LOC111091625 | -1.589175455 | 0.510103056 | 0.01442746 | 0.039232915 |
| LOC119865643 | -1.589771713 | 3.104026885 | 0.003659478 | 0.012319334 |

| | | | | |
|---|---|---|---|---|
| SOAT2 | -1.591152145 | 1.845893453 | 0.00403358 | 0.013377164 |
| LOC102156976 | -1.591258257 | -0.547143765 | 0.008487326 | 0.025101777 |
| ZCCHC2 | -1.592145918 | 6.348416069 | 0.000123573 | 0.000684085 |
| LOC111097881 | -1.59452184 | 3.071895228 | 0.005631125 | 0.01771826 |
| PLEKHF2 | -1.595487773 | 5.304735606 | 0.009427275 | 0.027443563 |
| SLC22A5 | -1.596292003 | 4.526435247 | 0.000746181 | 0.003181828 |
| KLHL28 | -1.596491275 | 5.345258977 | 0.000219334 | 0.001112481 |
| LOC102154094 | -1.597438427 | 1.198135357 | 0.004050981 | 0.013417942 |
| CLCN1 | -1.599638804 | 0.927277995 | 0.010604461 | 0.03030865 |
| RAP1B | -1.600308618 | 7.441001161 | 0.000845133 | 0.003542904 |
| TET2 | -1.600910044 | 7.525493331 | 0.000251796 | 0.001250211 |
| PIK3CB | -1.602757507 | 5.693773041 | 0.000297174 | 0.001444739 |
| TASL | -1.603472683 | 2.93701726 | 0.018442937 | 0.047709927 |
| KRAS | -1.604452694 | 6.141232267 | 0.00316731 | 0.010902972 |
| LOC119864801 | -1.604673946 | -0.087747724 | 0.018510239 | 0.047844056 |
| STING1 | -1.604881637 | 4.523570461 | 0.017440101 | 0.04560561 |
| ARHGAP27 | -1.604895059 | 6.020911875 | 0.001648105 | 0.006241374 |
| LOC111094459 | -1.604978338 | 3.086906437 | 0.012206729 | 0.034046398 |
| TNFRSF1B | -1.605030573 | 7.396656334 | 0.005724968 | 0.01799003 |
| SLC16A7 | -1.605973155 | 5.953117127 | 0.004060471 | 0.013434612 |
| TRAF5 | -1.607080303 | 4.405517292 | 0.014459444 | 0.039285386 |
| LOC119874133 | -1.607096644 | 1.226023845 | 0.010584469 | 0.030255499 |
| NT5C2 | -1.607674204 | 6.417785988 | 2.34E-05 | 0.00016226 |
| LOC119877425 | -1.607701202 | 2.056767332 | 0.008056974 | 0.024053857 |
| SH2B2 | -1.608034867 | 4.319160951 | 0.001689919 | 0.006378973 |
| VDAC3 | -1.60807702 | 4.844062153 | 0.000932808 | 0.003864929 |
| TNFAIP8L2 | -1.609176489 | 3.030654118 | 0.006850286 | 0.020943185 |
| LOC119866855 | -1.61120282 | 3.409668819 | 0.000931874 | 0.003862403 |
| BLOC1S2 | -1.612056477 | 2.807901421 | 0.000733619 | 0.003133417 |
| GADD45A | -1.612496149 | 6.483019099 | 0.006086101 | 0.018931406 |
| CCR1 | -1.613002951 | 4.287174863 | 0.013097454 | 0.036160652 |
| LOC119876868 | -1.615571527 | 8.072931331 | 0.004946724 | 0.015861866 |
| LOC119876637 | -1.616049376 | 4.639189993 | 0.005769218 | 0.018110697 |
| TMEM170A | -1.617280598 | 3.781252859 | 0.000733728 | 0.003133417 |
| SCARB2 | -1.619944472 | 7.022937837 | 0.001297266 | 0.005108221 |
| LOC102154169 | -1.620428269 | 2.225746553 | 0.008153792 | 0.024277665 |
| LOC102157195 | -1.620895784 | 3.975775866 | 0.003545225 | 0.011983104 |
| HERC6 | -1.621766961 | 5.519081974 | 0.002298706 | 0.008283982 |
| LOC111091357 | -1.621855187 | 1.100711779 | 0.009771667 | 0.028290321 |

| | | | | |
|---|---|---|---|---|
| SLA2 | -1.622362305 | 5.103645491 | 0.001023831 | 0.004172526 |
| AKIRIN2 | -1.622761905 | 4.864389272 | 1.68E-05 | 0.000120944 |
| NRN1L | -1.622835549 | 0.863812483 | 0.008698999 | 0.025640338 |
| TXNIP | -1.62579333 | 9.293676859 | 0.000339378 | 0.00161648 |
| PTGER4 | -1.625835247 | 5.072257998 | 0.00755095 | 0.022770521 |
| LOC119872826 | -1.628445299 | 1.185699627 | 0.005253557 | 0.016686922 |
| POLG2 | -1.628487731 | 4.295078272 | 9.54E-05 | 0.000548828 |
| LOC111097359 | -1.628939725 | -0.102547433 | 0.016754106 | 0.044174177 |
| GCNT2 | -1.629200567 | 4.991902902 | 0.000646317 | 0.002815602 |
| LOC611446 | -1.629339113 | 3.289561814 | 0.01065361 | 0.030413026 |
| QPCT | -1.631074895 | 4.006911592 | 0.017174837 | 0.045075109 |
| LOC111097036 | -1.631460677 | 1.002279445 | 0.003651524 | 0.012294854 |
| TRPM5 | -1.631735157 | 1.334530544 | 0.008658671 | 0.025545791 |
| FLVCR2 | -1.63192472 | 4.441601853 | 0.000683925 | 0.002954496 |
| UCP2 | -1.632008875 | 6.900382456 | 0.001037991 | 0.004224671 |
| SGO1 | -1.632190683 | 1.778885597 | 0.014406271 | 0.039180212 |
| LOC611813 | -1.632597017 | 1.127143608 | 0.003801374 | 0.012729788 |
| LOC119870774 | -1.632699886 | 1.380338626 | 0.008862787 | 0.026041646 |
| LOC102154803 | -1.633208603 | -0.167762437 | 0.010969832 | 0.031143459 |
| LOC119872905 | -1.63334421 | 2.331788284 | 0.016656484 | 0.043971879 |
| LOC102156639 | -1.633795391 | 1.098766097 | 0.018049249 | 0.046892997 |
| LOC119876039 | -1.633989555 | 0.953597109 | 0.003809793 | 0.01275207 |
| LOC119876636 | -1.634153474 | 3.602957839 | 0.004647961 | 0.015050838 |
| EXOC6 | -1.636188122 | 5.639375619 | 0.001478565 | 0.005690808 |
| ORAI2 | -1.637269645 | 5.89718322 | 0.002857892 | 0.010003085 |
| LOC111092516 | -1.637427925 | 3.221710033 | 0.006268518 | 0.019411002 |
| LOC100685381 | -1.638978762 | 0.316923643 | 0.006111838 | 0.018993671 |
| CHD1 | -1.640827468 | 6.60675049 | 0.000404705 | 0.001885461 |
| ZSWIM5 | -1.64188936 | 3.847579451 | 0.005443386 | 0.017215016 |
| LOC119866134 | -1.643461062 | 2.811869499 | 0.000643804 | 0.002806912 |
| LOC119864177 | -1.645589532 | 0.814126708 | 0.016008749 | 0.0426395 |
| LOC119866580 | -1.64662304 | 3.340640961 | 0.000746239 | 0.003181828 |
| LOC119873707 | -1.646773959 | -0.185838449 | 0.013261332 | 0.036529341 |
| LOC119866197 | -1.646849635 | 3.57124931 | 0.006175173 | 0.019163018 |
| MYBL1 | -1.647340316 | 3.085621824 | 0.00442748 | 0.014444794 |
| SAMD9L | -1.6478242 | 7.371028072 | 0.009880944 | 0.028579968 |
| LOC111097975 | -1.650629329 | 0.164439282 | 0.013877093 | 0.037988913 |
| RP2 | -1.651982604 | 4.950976235 | 1.40E-05 | 0.000102995 |
| SSBP2 | -1.653275988 | 4.563931434 | 0.015505977 | 0.041565991 |

| | | | | |
|---|---|---|---|---|
| FXYD5 | -1.654457185 | 5.969542779 | 4.84E-05 | 0.000305622 |
| LOC102156726 | -1.654651194 | 2.920312663 | 0.000650956 | 0.002833536 |
| GFOD2 | -1.656439412 | 4.117494329 | 0.000144453 | 0.000778014 |
| BTK | -1.65670554 | 4.480092416 | 0.006480748 | 0.019974851 |
| ACSL4 | -1.658110346 | 6.253110752 | 0.000190994 | 0.000986046 |
| LOC102153833 | -1.659475822 | 0.658330081 | 0.00370171 | 0.012444107 |
| MED10 | -1.65978156 | 4.040237473 | 0.000323598 | 0.001553344 |
| ERICH6 | -1.661226585 | 1.336540805 | 0.006789599 | 0.020798724 |
| AGO4 | -1.662094375 | 6.89906811 | 6.39E-05 | 0.000387881 |
| LOC119864723 | -1.66374234 | 3.468509656 | 0.000650431 | 0.002831819 |
| VTCN1 | -1.664467738 | 0.947125255 | 0.012235628 | 0.034111495 |
| LOC119876089 | -1.666197842 | 3.550968974 | 0.003865613 | 0.012909004 |
| STAT4 | -1.667101111 | 5.352139382 | 0.001400417 | 0.00543934 |
| LRRC19 | -1.668457351 | 2.430022358 | 0.010854598 | 0.03088001 |
| LOC119874183 | -1.668659905 | 4.123481627 | 0.001217971 | 0.004846168 |
| LOC106560032 | -1.669080523 | 6.032932188 | 0.002884736 | 0.010082386 |
| LOC119864941 | -1.671198182 | 3.229138597 | 0.0107309 | 0.030589347 |
| LOC111098019 | -1.67207861 | 2.315958401 | 0.014893033 | 0.04023015 |
| OIP5 | -1.672400509 | 2.733597563 | 0.00100998 | 0.004130056 |
| AGTPBP1 | -1.678717721 | 5.992249687 | 1.27E-05 | 9.37E-05 |
| DAZAP2 | -1.679220636 | 7.433220251 | 4.02E-05 | 0.000259301 |
| ERO1B | -1.679394751 | 5.239825499 | 0.000181075 | 0.000942244 |
| FBXO9 | -1.679682264 | 5.242574064 | 7.31E-06 | 5.78E-05 |
| LOC119871967 | -1.680859376 | -0.556323653 | 0.017314411 | 0.045337106 |
| LOC102156997 | -1.681707078 | 5.804615097 | 0.00038539 | 0.001805958 |
| PTGR1 | -1.682626207 | 5.678071011 | 0.000871473 | 0.003639899 |
| IFNAR2 | -1.68319476 | 4.564921841 | 0.002326766 | 0.008367016 |
| ATP8A1 | -1.683302687 | 6.665570715 | 0.000794579 | 0.003355592 |
| SLC9A9 | -1.683351748 | 5.155108301 | 0.00139882 | 0.005436985 |
| LOC106557567 | -1.684385861 | 3.497587852 | 0.000139924 | 0.000758141 |
| BPI | -1.687904097 | 7.234378074 | 0.004686143 | 0.015140545 |
| AGMAT | -1.689759359 | -0.619663845 | 0.018072428 | 0.046941961 |
| LOC102153616 | -1.690797622 | 5.295576504 | 0.000802335 | 0.003382416 |
| NLRC5 | -1.690900913 | 7.101791249 | 0.010876679 | 0.030926652 |
| PAAF1 | -1.69111843 | 3.584906566 | 0.000274887 | 0.001346967 |
| GNAQ | -1.691474137 | 5.661812339 | 0.001020348 | 0.004164062 |
| IL7 | -1.692485467 | 0.763025257 | 0.015571487 | 0.041695186 |
| PDE4B | -1.693761483 | 5.960558891 | 0.010962581 | 0.031131029 |
| H2BC8 | -1.694011434 | 3.657762386 | 0.000941572 | 0.003889952 |

| | | | | |
|---|---|---|---|---|
| HMGB2 | -1.694681436 | 5.695657995 | 8.52E-06 | 6.62E-05 |
| LOC111096199 | -1.695082046 | 2.175788592 | 0.00616276 | 0.019132719 |
| SGK3 | -1.69744971 | 3.363507893 | 0.000564261 | 0.002503125 |
| LOC119873696 | -1.697555256 | 1.248602287 | 0.012458105 | 0.034655949 |
| BAG4 | -1.69928965 | 5.835323436 | 4.75E-05 | 0.000300457 |
| LOC102155157 | -1.70138389 | 3.826519764 | 0.009698597 | 0.028093784 |
| LMLN2 | -1.703103386 | -0.565566987 | 0.00699579 | 0.02132486 |
| S1PR1 | -1.703390861 | 6.198768652 | 0.008058943 | 0.024053857 |
| LOC111097066 | -1.703506148 | 2.498791798 | 0.016885702 | 0.044456246 |
| LOC119869301 | -1.704702047 | 3.999409258 | 3.19E-05 | 0.000212043 |
| LOC119872277 | -1.706096281 | 4.205848071 | 0.017530328 | 0.045780793 |
| LOC106559329 | -1.707480396 | 1.171547631 | 0.000939977 | 0.003885364 |
| HMCES | -1.707927606 | 4.252667284 | 1.26E-05 | 9.37E-05 |
| TPD52 | -1.709157562 | 5.368810163 | 0.000189032 | 0.000977077 |
| SMIM3 | -1.709923675 | 5.612028253 | 0.000777002 | 0.003294848 |
| CRB3 | -1.71025607 | 5.187122286 | 0.006123702 | 0.019027813 |
| LOC102156091 | -1.710382952 | 2.90387355 | 0.01418077 | 0.038644515 |
| PMEL | -1.710457364 | 4.108557234 | 0.000297916 | 0.001448022 |
| DOK2 | -1.710986092 | 4.958928164 | 0.000164527 | 0.000869068 |
| LOC102154353 | -1.71125419 | -0.301876149 | 0.013221889 | 0.036448487 |
| LOC111091006 | -1.711755139 | 3.677356226 | 0.000395775 | 0.001847831 |
| LOC111097580 | -1.71194802 | 2.507079062 | 0.017982075 | 0.046735281 |
| TRMT9B | -1.71310911 | 2.775661687 | 0.013196578 | 0.036397231 |
| FLT3 | -1.715314732 | 3.997451237 | 0.006453786 | 0.019907899 |
| LOC119868650 | -1.715380065 | 0.01999725 | 0.015487123 | 0.041535144 |
| LOC119873307 | -1.718258635 | -0.023626927 | 0.016892065 | 0.044467598 |
| MOV10 | -1.719031061 | 6.353139971 | 0.000293752 | 0.001430349 |
| MTHFS | -1.719295922 | 4.981253496 | 1.40E-05 | 0.000102963 |
| LOC119870966 | -1.720162316 | 3.656601074 | 0.000603252 | 0.002651986 |
| LOC119876487 | -1.720190229 | 2.198216029 | 0.008104272 | 0.024153451 |
| GRK6 | -1.720601175 | 6.09566552 | 5.75E-05 | 0.000354906 |
| LOC111095909 | -1.720885067 | 1.496869569 | 0.004856701 | 0.015610156 |
| LOC111092940 | -1.721793425 | 2.220297634 | 0.0026771 | 0.009445024 |
| GBP1 | -1.723682485 | 5.934747309 | 0.01131735 | 0.031966756 |
| PTK2B | -1.72534616 | 6.626630557 | 0.000180171 | 0.000938889 |
| SLC25A37 | -1.725748036 | 5.778307976 | 4.30E-05 | 0.000274949 |
| LOC102156658 | -1.72855916 | 5.243648926 | 0.002424112 | 0.008661026 |
| LOC111094575 | -1.730245468 | -0.716224315 | 0.006593756 | 0.020264674 |
| LOC106559812 | -1.730404822 | 1.045040373 | 0.015181487 | 0.040852788 |

| | | | | |
|---|---|---|---|---|
| UNC119 | -1.730822492 | 4.18559785 | 0.000470899 | 0.002140928 |
| SUSD1 | -1.733182938 | 4.761723813 | 0.000439748 | 0.002021779 |
| LOC119876493 | -1.733693107 | 1.9538807 | 0.00408585 | 0.013504041 |
| LOC111091807 | -1.733931201 | 0.74326743 | 0.011192705 | 0.031672475 |
| NCK2 | -1.734245195 | 5.925962505 | 5.55E-05 | 0.000343421 |
| LOC106557683 | -1.734756713 | 0.24172108 | 0.01691499 | 0.044517131 |
| LOC119865808 | -1.734895509 | 0.655260923 | 0.007712072 | 0.023162934 |
| LOC102156211 | -1.734962274 | 3.190437044 | 0.004528207 | 0.014709207 |
| DIAPH1 | -1.735292553 | 7.732928865 | 1.25E-05 | 9.29E-05 |
| ACVR1B | -1.73729881 | 4.873209925 | 4.84E-05 | 0.000305622 |
| LOC100685007 | -1.73779511 | 0.390591064 | 0.009995063 | 0.028848449 |
| ELF4 | -1.739263282 | 5.593800378 | 2.57E-06 | 2.35E-05 |
| LOC111094380 | -1.741231854 | 4.788928039 | 0.000268888 | 0.001321754 |
| LOC102155710 | -1.741539624 | 2.488785161 | 0.000974042 | 0.004000455 |
| LOC100688306 | -1.742670459 | 2.403218625 | 0.001022375 | 0.004168945 |
| LOC119878074 | -1.743515213 | 0.892085833 | 0.008761007 | 0.025788034 |
| RNF39 | -1.743600781 | 2.798909667 | 0.00101035 | 0.004130789 |
| HPX | -1.743766965 | 0.238083362 | 0.00704487 | 0.021451518 |
| TBC1D8 | -1.743792746 | 6.029261529 | 0.000202593 | 0.001039735 |
| LOC111094687 | -1.744190832 | 1.624906735 | 0.00770798 | 0.023153852 |
| LOC119865243 | -1.744506292 | 2.93762192 | 0.008212021 | 0.024410799 |
| PPP4R1 | -1.74537943 | 5.896011007 | 0.000220489 | 0.001117029 |
| LOC119871502 | -1.745677982 | 3.265893314 | 4.06E-05 | 0.000261952 |
| LOC119869793 | -1.746107521 | 2.715378315 | 0.003345052 | 0.011413305 |
| EDEM1 | -1.746239152 | 7.241736303 | 0.002231016 | 0.008084403 |
| LOC111096020 | -1.747046027 | 2.579604226 | 0.016005512 | 0.042636119 |
| LOC111093141 | -1.747790368 | 4.756577408 | 6.33E-05 | 0.000385215 |
| LOC111094419 | -1.748105737 | 2.171440989 | 0.002174515 | 0.007912738 |
| FOXN2 | -1.748571175 | 6.033866496 | 0.00361191 | 0.012181905 |
| ETS1 | -1.749499859 | 8.222620662 | 0.002606874 | 0.009228811 |
| GPCPD1 | -1.750116698 | 8.008535009 | 0.000346928 | 0.001648175 |
| TRNAE-UUC_7 | -1.750289563 | -0.53015876 | 0.006229119 | 0.019305549 |
| TFEC | -1.750388185 | 4.6525262 | 0.00418281 | 0.01377721 |
| MPEG1 | -1.751406672 | 7.658669401 | 0.008060222 | 0.024053857 |
| SLC2A6 | -1.751437056 | 4.193718153 | 2.03E-05 | 0.000142424 |
| KCTD20 | -1.752216633 | 6.516721787 | 2.50E-05 | 0.000172055 |
| SULF2 | -1.754404984 | 6.935985988 | 0.006838374 | 0.020912669 |
| LOC119865246 | -1.755703221 | 3.915056138 | 0.000935801 | 0.003873494 |
| KBTBD7 | -1.756415723 | 3.179634678 | 0.002993228 | 0.010391194 |

| | | | | |
|---|---|---|---|---|
| LOC119868464 | -1.757047168 | -0.416638334 | 0.009573119 | 0.027793464 |
| SLC46A2 | -1.7574117 | 3.723442684 | 0.001998368 | 0.007365754 |
| LOC102153619 | -1.75914745 | 5.725883809 | 0.002300082 | 0.008286182 |
| RABGAP1L | -1.759506595 | 7.254749595 | 3.59E-05 | 0.000235071 |
| LST1 | -1.761782885 | 4.23544725 | 0.001321343 | 0.005185606 |
| GSR | -1.762488898 | 4.907060566 | 0.000282501 | 0.001380729 |
| SOWAHD | -1.762982519 | 1.606105096 | 0.017879176 | 0.046552867 |
| TMSB4X | -1.763736194 | 9.481961029 | 0.000220594 | 0.0011173 |
| LOC607937 | -1.765078872 | 4.993671706 | 0.006164572 | 0.019135602 |
| LOC119868600 | -1.765148448 | -0.895891644 | 0.017974466 | 0.046721109 |
| TNFSF13B | -1.766083062 | 3.252254782 | 0.003281982 | 0.011233481 |
| DRC7 | -1.767339102 | 0.139133622 | 0.014783693 | 0.039995879 |
| GIMAP2 | -1.768454924 | 2.80263952 | 0.016937883 | 0.04457197 |
| LOC119872205 | -1.768793408 | -0.352206775 | 0.011820513 | 0.033141783 |
| ELSPBP1 | -1.768817281 | 0.018376834 | 0.00767794 | 0.023089219 |
| SCIMP | -1.770953412 | 5.046574124 | 0.00110227 | 0.004452012 |
| ARL14EP | -1.773059932 | 5.395606586 | 3.04E-05 | 0.000203658 |
| FOXN1 | -1.773064044 | 0.453241748 | 0.015515498 | 0.04158637 |
| PLEKHG7 | -1.77419398 | 0.969014567 | 0.0075899 | 0.022865707 |
| CCDC115 | -1.774329768 | 5.53925112 | 0.000246485 | 0.00122806 |
| MALT1 | -1.778311084 | 6.325134061 | 0.000132737 | 0.000725269 |
| DENND2D | -1.778958455 | 5.11009814 | 0.00279674 | 0.009819172 |
| LOC111093470 | -1.781052049 | 1.358696769 | 0.001316638 | 0.005172286 |
| OPN3 | -1.78178176 | 4.384519554 | 1.98E-05 | 0.000139619 |
| PRKCD | -1.781805372 | 6.31610311 | 3.35E-05 | 0.000221183 |
| EGLN3 | -1.781908859 | 4.478654171 | 0.003399655 | 0.011568647 |
| ITGB3 | -1.784429738 | 6.804368244 | 0.014878566 | 0.040207376 |
| DENND4A | -1.78499227 | 6.730763344 | 0.000269555 | 0.001324729 |
| LOC119873472 | -1.787445546 | 1.684863702 | 0.002055282 | 0.007540904 |
| SLAMF8 | -1.789260646 | 4.12583788 | 0.01720588 | 0.045123793 |
| ACSL1 | -1.789343109 | 5.774949094 | 0.000141333 | 0.000763867 |
| ABHD2 | -1.792957389 | 7.312040665 | 1.99E-05 | 0.000140419 |
| STAT5B | -1.794257866 | 6.656562049 | 2.37E-05 | 0.000164078 |
| TNFSF13 | -1.794465829 | 3.532337037 | 0.000431322 | 0.001989434 |
| LOC111091234 | -1.794999989 | 3.601330159 | 0.000788549 | 0.003333378 |
| ZBTB18 | -1.795011186 | 5.114074021 | 0.001400192 | 0.00543934 |
| LOC111093108 | -1.795591892 | -0.736046776 | 0.018585299 | 0.048003716 |
| COL10A1 | -1.795702589 | 2.816695767 | 0.000570362 | 0.002526871 |
| CXHXorf38 | -1.795949416 | 3.34731491 | 0.000148352 | 0.000795262 |

| | | | | |
|---|---|---|---|---|
| TEX30 | -1.796038471 | 1.388399376 | 0.003417936 | 0.011619379 |
| LOC106558596 | -1.796146878 | 0.8279641 | 0.002991954 | 0.010388436 |
| CEACAM19 | -1.796220998 | 0.640932407 | 0.011163942 | 0.031624117 |
| LOC111095167 | -1.797260078 | -0.069065505 | 0.012445223 | 0.034632501 |
| DENND1C | -1.797395993 | 5.785127352 | 0.005741942 | 0.018038138 |
| ADIG | -1.798073686 | 1.33982093 | 0.009976902 | 0.028807542 |
| TUT7 | -1.798137516 | 7.315147824 | 2.06E-06 | 1.93E-05 |
| LOC119873566 | -1.799366462 | 2.523109743 | 0.000213254 | 0.001086472 |
| BTG2 | -1.800126497 | 6.472829514 | 0.000169115 | 0.000889186 |
| BEND7 | -1.801449847 | 4.222923101 | 0.013389673 | 0.036808015 |
| MMD | -1.802111161 | 4.815058618 | 0.004505565 | 0.014651028 |
| LOC491028 | -1.802818394 | 2.268919741 | 0.000731252 | 0.003126089 |
| IQGAP2 | -1.80321693 | 6.440081961 | 7.37E-05 | 0.000440137 |
| GTF2B | -1.803872942 | 4.104337775 | 3.41E-05 | 0.00022453 |
| PANK3 | -1.804524713 | 7.70597408 | 8.56E-05 | 0.000500185 |
| LOC119864183 | -1.804662125 | -0.432998415 | 0.012777827 | 0.035418065 |
| FAM131B | -1.80589792 | 2.524343241 | 0.001388491 | 0.005404591 |
| DOCK3 | -1.806137841 | 5.392072571 | 0.001683951 | 0.006361562 |
| PRSS8 | -1.807579057 | 1.971245565 | 0.000392194 | 0.001834271 |
| RINL | -1.80814217 | 3.627156617 | 0.00165133 | 0.006252495 |
| LOC102157222_2 | -1.808650614 | 1.034772895 | 0.003390065 | 0.011541453 |
| EMILIN2 | -1.808686801 | 6.818317143 | 0.000564424 | 0.002503125 |
| SEMA4A | -1.810940276 | 5.571559485 | 0.001938623 | 0.007169926 |
| KIAA0513 | -1.811031051 | 6.187253825 | 1.26E-05 | 9.37E-05 |
| HOPX | -1.811834821 | 3.762303535 | 0.005899294 | 0.01843891 |
| LOC111095592 | -1.812055691 | 3.532827166 | 0.000660777 | 0.002869368 |
| LOC111091477 | -1.812801031 | 3.521113065 | 0.005145104 | 0.016403332 |
| LOC111093364 | -1.815222026 | 1.724189798 | 0.002113178 | 0.007721955 |
| SYN3 | -1.816017443 | 2.793467093 | 5.70E-05 | 0.000351991 |
| TCP11L2 | -1.816333016 | 5.567758053 | 3.94E-05 | 0.000255439 |
| PHF7 | -1.816555599 | 2.683818606 | 0.001435473 | 0.005550579 |
| LOC102154652 | -1.816632095 | 1.686329952 | 0.015150118 | 0.040788634 |
| LOC111090694 | -1.816863938 | 2.153485992 | 0.001264905 | 0.005004434 |
| LOC119879477 | -1.817747001 | 1.893627315 | 0.001064227 | 0.004318479 |
| RALGAPA2 | -1.81893152 | 6.409749333 | 3.89E-06 | 3.35E-05 |
| LOC119865978 | -1.818971623 | 0.630253544 | 0.004006516 | 0.013307773 |
| KAT2B | -1.820261366 | 6.915364054 | 6.46E-05 | 0.00039204 |
| LOC119873403 | -1.82040518 | 0.098952957 | 0.008242962 | 0.024485981 |
| SYCE1L | -1.821112125 | 2.829120602 | 0.000106029 | 0.000602191 |

| | | | | |
|---|---|---|---|---|
| MFAP3L | -1.822042915 | 5.622507569 | 0.000364294 | 0.001719783 |
| LOC119869428 | -1.822096374 | 0.383479902 | 0.004227294 | 0.013881537 |
| CYTH4 | -1.823382035 | 6.592842212 | 0.004194383 | 0.013806934 |
| LOC111091273 | -1.824049024 | 0.222451182 | 0.011485252 | 0.032381926 |
| LOC102156269 | -1.825732179 | 4.314385747 | 3.87E-06 | 3.33E-05 |
| PPM1D | -1.825779164 | 4.777511128 | 4.99E-07 | 5.59E-06 |
| RSU1 | -1.825983891 | 6.845697034 | 0.00022291 | 0.001126661 |
| PARP4 | -1.826272583 | 6.802812913 | 0.000483313 | 0.00219139 |
| LOC111096589 | -1.8265408 | 1.658751667 | 0.012984794 | 0.035899915 |
| PYGM | -1.827210297 | 3.673119314 | 0.000543437 | 0.002423935 |
| LOC119871268 | -1.828201309 | 0.266645516 | 0.007407989 | 0.022399618 |
| SHMT1 | -1.832877316 | 4.981329061 | 0.000240126 | 0.001201059 |
| TBC1D23 | -1.833596267 | 6.317624759 | 1.40E-05 | 0.000102995 |
| LOC119873254 | -1.836167359 | 2.498174982 | 0.000504773 | 0.002276783 |
| GIMAP6 | -1.836993844 | 5.246410291 | 0.004230464 | 0.013887739 |
| LOC119864983 | -1.837505237 | 0.773961485 | 0.00602643 | 0.018784814 |
| NUP210L | -1.838208537 | 3.3432425 | 0.000617497 | 0.002705826 |
| LOC119871933 | -1.83968394 | 1.435466061 | 0.002414428 | 0.008633542 |
| LOC611209 | -1.840911914 | 5.188581396 | 1.15E-06 | 1.16E-05 |
| LOC610465 | -1.841095883 | 1.04640357 | 0.004664332 | 0.015088071 |
| LOC119864942 | -1.842165466 | 1.583808782 | 0.015140214 | 0.040772099 |
| LOC111093278 | -1.842239184 | 1.88392352 | 0.007645187 | 0.02300988 |
| LOC111090721 | -1.84234677 | 4.687647241 | 0.008184996 | 0.024347161 |
| LPXN | -1.842682147 | 4.503993094 | 0.001998936 | 0.007366594 |
| WAS | -1.845484805 | 5.036753657 | 0.001480661 | 0.005696852 |
| LOC102151157 | -1.846612115 | 2.618300603 | 0.000470787 | 0.002140865 |
| LOC119864497 | -1.8466742 | 3.221723678 | 0.002328957 | 0.008372117 |
| LOC111091795 | -1.847488056 | 1.247023176 | 0.013501738 | 0.037054979 |
| LOC119877387 | -1.847527597 | -0.243226015 | 0.010454753 | 0.029912325 |
| LOC111092991 | -1.848848767 | 4.844661452 | 0.000386588 | 0.001809618 |
| LOC102155521 | -1.849700889 | 6.318248131 | 6.25E-05 | 0.000380874 |
| PITPNC1 | -1.851742743 | 6.679646053 | 0.001307575 | 0.005142273 |
| REL | -1.851820159 | 7.126794663 | 7.93E-05 | 0.000468416 |
| CLEC12A | -1.851890304 | 2.889905739 | 0.014040565 | 0.038358901 |
| LOC119872172 | -1.853877284 | 2.448888176 | 0.006086561 | 0.018931406 |
| IL13RA1 | -1.853896468 | 8.181956556 | 0.00123872 | 0.004910697 |
| LOC119869417 | -1.854788837 | 2.833257045 | 0.000533758 | 0.002387154 |
| IL17RA | -1.855281141 | 8.28364729 | 1.86E-05 | 0.000131948 |
| TNFAIP3 | -1.856003561 | 6.648824226 | 7.84E-05 | 0.000464531 |

| | | | | |
|---|---|---|---|---|
| LOC119871259 | -1.856472558 | 0.625341791 | 0.017537904 | 0.045795058 |
| LOC111093244 | -1.857714343 | 2.980562706 | 0.000163782 | 0.000865555 |
| MRPL16_2 | -1.85818136 | 0.674548141 | 0.014953841 | 0.040350422 |
| USP32 | -1.859001899 | 6.697227979 | 8.05E-07 | 8.46E-06 |
| LOC119872819 | -1.859128465 | 0.60697522 | 0.004136341 | 0.013642811 |
| LOC102154431 | -1.861162898 | 2.817220014 | 0.003370947 | 0.011483579 |
| LOC119868031 | -1.861582314 | 0.993169829 | 0.006254298 | 0.019378042 |
| LOC119867317 | -1.862535237 | -1.042013853 | 0.016382858 | 0.043406563 |
| LOC478220 | -1.86401794 | 0.55323204 | 0.015366779 | 0.041279647 |
| NFE2L3 | -1.86418578 | 4.245123543 | 0.000739509 | 0.003155616 |
| MAP3K9 | -1.864289872 | 3.553765342 | 0.003634105 | 0.012247234 |
| KCTD6 | -1.865371962 | 2.993209596 | 0.000194139 | 0.001000377 |
| ECT2L | -1.865603838 | 3.044222094 | 0.000130646 | 0.000716105 |
| MED20 | -1.866415588 | 3.405433944 | 0.000317768 | 0.001530412 |
| ATP8B4 | -1.867416006 | 6.390388003 | 0.003011547 | 0.010444753 |
| LOC119871188 | -1.867456883 | 3.511626715 | 0.000218212 | 0.001108346 |
| LOC119870865 | -1.867609523 | 2.852736764 | 0.000753441 | 0.00320749 |
| LOC111091186 | -1.870753834 | 0.164359352 | 0.015102499 | 0.040695817 |
| ACOD1 | -1.871733468 | 2.777372954 | 0.008119553 | 0.024192338 |
| LOC106559265 | -1.873887341 | 0.9313322 | 0.001540887 | 0.005903419 |
| LOC119878954_2 | -1.874054786 | -0.549379455 | 0.013361507 | 0.036758561 |
| GPR183 | -1.874487364 | 3.732050053 | 0.016372877 | 0.043385423 |
| LOC111090896 | -1.875546045 | 2.762337694 | 0.001542145 | 0.005907195 |
| LOC119867083 | -1.876052341 | 2.069101299 | 0.000483951 | 0.002193824 |
| MAP3K1 | -1.87734468 | 7.346332954 | 0.000309481 | 0.001496847 |
| FTH1 | -1.877926356 | 1.659574075 | 0.003663555 | 0.012331144 |
| LOC102153048 | -1.878043049 | 1.050118972 | 0.00073664 | 0.003144613 |
| ENPP5 | -1.878321767 | 4.595492791 | 0.000362688 | 0.001713656 |
| AP1S2 | -1.878877631 | 6.250845838 | 9.57E-05 | 0.00055 |
| LOC119866204 | -1.879540836 | -0.723939389 | 0.014643192 | 0.039695047 |
| LOC102151885 | -1.88008333 | 0.631035037 | 0.013798478 | 0.037802352 |
| RLBP1 | -1.880761977 | 0.61246955 | 0.002510271 | 0.008939385 |
| LOC111090013 | -1.881153363 | 2.46764706 | 0.001128195 | 0.004542333 |
| DNAH6 | -1.881397688 | 1.930354197 | 0.011989127 | 0.033532112 |
| MIR6529 | -1.881696963 | -0.887244314 | 0.016157479 | 0.042932018 |
| LOC111098018 | -1.885207295 | 5.865435489 | 0.000516064 | 0.002319505 |
| ARHGAP30 | -1.886662482 | 6.202918127 | 0.000133679 | 0.000729778 |
| LOC119868077 | -1.886913607 | 4.385777734 | 0.005299925 | 0.016807912 |
| NOCT | -1.886949757 | 5.083075636 | 0.000201163 | 0.001033372 |

| | | | | |
|---|---|---|---|---|
| LOC119873614 | -1.887196807 | 3.762759893 | 0.00127764 | 0.005047445 |
| LOC102152785 | -1.887300358 | 0.53072913 | 0.001584225 | 0.006040624 |
| LOC111091559 | -1.887961124 | 0.889907556 | 0.001846268 | 0.006876467 |
| DEDD2 | -1.888103813 | 4.662895513 | 1.29E-05 | 9.52E-05 |
| TRANK1 | -1.888339428 | 6.188148837 | 0.002456964 | 0.008766838 |
| LOC111098825 | -1.890768897 | 5.204552147 | 0.002432303 | 0.008687425 |
| DAGLB | -1.891520358 | 5.879641158 | 7.71E-07 | 8.16E-06 |
| TIPARP | -1.891975602 | 6.09036759 | 8.12E-05 | 0.000478068 |
| GPR84 | -1.892589994 | 0.963868979 | 0.001220345 | 0.004852941 |
| LOC111097331 | -1.894054466 | 1.538675536 | 0.000895757 | 0.00373054 |
| LOC119872858 | -1.894616403 | 3.619430114 | 9.30E-05 | 0.000537093 |
| LOC119874472 | -1.896414711 | 3.127782209 | 0.002694194 | 0.009496058 |
| LOC102155525 | -1.897024149 | -0.13188212 | 0.011370616 | 0.032096296 |
| LOC119876688 | -1.89834584 | 0.366704778 | 0.011244204 | 0.031789152 |
| LOC119872180 | -1.899235912 | -0.444579165 | 0.004086486 | 0.013504041 |
| LMOD2 | -1.899410566 | 0.471885369 | 0.008075076 | 0.024086308 |
| MYD88 | -1.900289848 | 5.900854941 | 1.52E-06 | 1.49E-05 |
| CRYBG1 | -1.901821097 | 7.132255506 | 0.000124086 | 0.0006864 |
| LOC111095821 | -1.902372636 | -0.340886743 | 0.012126818 | 0.03385169 |
| LOC111096630 | -1.902376668 | 2.91847813 | 0.004460463 | 0.014523981 |
| LOC100856351 | -1.903975357 | -0.179763389 | 0.00556454 | 0.01754441 |
| NEK5 | -1.904314989 | 0.829693237 | 0.007619372 | 0.022948118 |
| DLA-64 | -1.904641281 | 7.974699538 | 0.001872455 | 0.006952497 |
| LOC111096978 | -1.904858607 | 3.123223386 | 0.00373125 | 0.012525927 |
| LOC119873942 | -1.905190989 | 5.949750575 | 7.87E-06 | 6.18E-05 |
| ACSS1 | -1.908199692 | 6.686777275 | 4.61E-06 | 3.88E-05 |
| INSM2 | -1.908775343 | -0.348773285 | 0.005645177 | 0.017759896 |
| NRROS | -1.908859509 | 4.817732578 | 0.001090018 | 0.004406635 |
| CDC42SE2 | -1.910934918 | 5.580660949 | 0.000160254 | 0.000849811 |
| VAV1 | -1.913300915 | 5.561864541 | 0.000855648 | 0.003582088 |
| CYP4V2 | -1.913394466 | 5.953804989 | 0.001051612 | 0.004272089 |
| LOC111091322 | -1.916496111 | 4.159520038 | 8.14E-06 | 6.37E-05 |
| GDF15 | -1.916779622 | 0.300031804 | 0.011155216 | 0.03160353 |
| LOC111095911 | -1.918061196 | -0.328545597 | 0.006446964 | 0.019895351 |
| LOC106559759 | -1.91903688 | 1.520448043 | 0.000298673 | 0.001451049 |
| TMEM170B | -1.919502592 | 6.378931488 | 6.40E-06 | 5.15E-05 |
| P2RY8 | -1.921432699 | 4.081372427 | 0.008171275 | 0.02431636 |
| LOC102154658 | -1.922484311 | 4.537031808 | 0.000107122 | 0.000607354 |
| SQOR | -1.923086845 | 4.665260862 | 1.21E-05 | 9.04E-05 |

| | | | | |
|---|---|---|---|---|
| LOC111093581 | -1.923160443 | 1.125789355 | 0.002071308 | 0.007589425 |
| LOC102152824 | -1.923371767 | 2.937466264 | 0.008724261 | 0.025700817 |
| IL6R | -1.923784888 | 5.68899931 | 0.0004876 | 0.00220944 |
| LOC119872197 | -1.923964731 | -0.987691735 | 0.015897274 | 0.042394687 |
| LOC111097027 | -1.923969515 | 1.341271899 | 0.008773939 | 0.025822592 |
| TMEM140 | -1.924455026 | 5.579645423 | 1.65E-05 | 0.00011897 |
| MAP4K1 | -1.924458454 | 4.691939969 | 0.002117338 | 0.00773585 |
| LOC111097886 | -1.925875176 | 3.724438365 | 1.46E-05 | 0.000106469 |
| LOC119870831 | -1.926457482 | 0.527903345 | 0.002685445 | 0.009469841 |
| LOC111091460 | -1.927263372 | 0.227807676 | 0.001862446 | 0.006930768 |
| CBL | -1.92917774 | 7.875017092 | 3.54E-06 | 3.09E-05 |
| LOC111094434 | -1.931058356 | 2.350054194 | 0.003636407 | 0.012253087 |
| LOC111094365 | -1.931348742 | 2.302477811 | 0.003778571 | 0.012665168 |
| LOC119873622 | -1.933578685 | 1.467487337 | 0.000467218 | 0.002126868 |
| LOC106559772 | -1.934026233 | 0.756446761 | 0.0076806 | 0.023094013 |
| LOC111095257 | -1.934101769 | 1.040338062 | 0.00296242 | 0.010304699 |
| LOC119864552 | -1.935244275 | 1.091915171 | 0.010040569 | 0.028960509 |
| SOX15 | -1.935973462 | 1.000840519 | 0.008732021 | 0.025720181 |
| LOC111095226 | -1.937778182 | 3.330871965 | 0.000666853 | 0.002892275 |
| CDC42SE1 | -1.938944868 | 5.283189801 | 7.38E-07 | 7.85E-06 |
| PLAGL2 | -1.939164188 | 5.459981087 | 2.97E-06 | 2.66E-05 |
| PSTPIP1 | -1.939515405 | 5.45352986 | 0.002924135 | 0.010191191 |
| LOC100686073 | -1.939574935 | 4.018989442 | 0.000676564 | 0.002927335 |
| ABI3 | -1.939621327 | 4.325456379 | 0.000232274 | 0.001167456 |
| LOC119877401 | -1.939937639 | 2.969669186 | 0.000438519 | 0.002017942 |
| LOC111094212 | -1.941442245 | 1.717944755 | 0.001337185 | 0.005237812 |
| LMNB1 | -1.943095755 | 5.32899665 | 1.42E-06 | 1.40E-05 |
| EFNA2 | -1.943203069 | 2.440457158 | 0.000376902 | 0.001771545 |
| CD37 | -1.94343311 | 5.762082141 | 0.000284861 | 0.001390181 |
| ZAP70 | -1.943511785 | 4.727191291 | 0.014734621 | 0.039895706 |
| LOC111097713 | -1.944126238 | 0.615062711 | 0.001607108 | 0.006110676 |
| LOC111098824 | -1.944159393 | 1.522810882 | 0.007408317 | 0.022399618 |
| MIR1842 | -1.944494017 | 1.246876416 | 0.000965083 | 0.003971662 |
| FRY | -1.945053016 | 7.61601125 | 1.76E-06 | 1.69E-05 |
| LOC102154370 | -1.945600503 | 1.24055107 | 0.003622618 | 0.012212176 |
| LOC119869688 | -1.947230372 | 1.844847295 | 0.003478504 | 0.011803604 |
| LOC102151630 | -1.947504492 | 2.353683062 | 0.004958476 | 0.015885446 |
| LOC111092639 | -1.948549935 | 4.259190471 | 3.54E-06 | 3.09E-05 |
| CD180 | -1.949503221 | 5.205884792 | 0.005453107 | 0.017233624 |

| | | | | |
|---|---|---|---|---|
| SMAP2 | -1.950677733 | 7.322820149 | 1.63E-05 | 0.000117508 |
| AKAP14 | -1.953738603 | -0.618117164 | 0.01214404 | 0.033895399 |
| CMTM6 | -1.954320026 | 5.012592134 | 0.000473713 | 0.002151014 |
| LOC119876647 | -1.955290439 | 1.46462168 | 0.010889304 | 0.03095539 |
| PTPN12 | -1.955506288 | 6.836595879 | 6.00E-05 | 0.000367593 |
| RS1 | -1.955523415 | 2.101921891 | 0.001363378 | 0.005324052 |
| LOC119871938 | -1.95680762 | 1.213644548 | 0.005886997 | 0.018403128 |
| LOC111091665 | -1.957168707 | 2.066367016 | 0.013075837 | 0.036113627 |
| TKT | -1.95786397 | 7.76580455 | 6.61E-06 | 5.29E-05 |
| MCUB | -1.959458449 | 4.381182422 | 0.00939964 | 0.027390889 |
| LOC119873207 | -1.959625593 | 3.520119292 | 0.000107039 | 0.000607134 |
| SLC25A33 | -1.960201943 | 2.97389292 | 0.00025013 | 0.001243059 |
| LOC119864635 | -1.960884309 | 0.356683088 | 0.010296135 | 0.029556044 |
| SERPINB1 | -1.961072367 | 6.312848901 | 2.15E-05 | 0.000149867 |
| LOC111091090 | -1.961777449 | 2.546723109 | 0.002211248 | 0.008030223 |
| LOC111092390 | -1.96198791 | -0.550352302 | 0.003048033 | 0.010552718 |
| GMFG | -1.96276219 | 4.92058126 | 0.000134359 | 0.000732567 |
| TMEM163 | -1.9633822 | 1.669232827 | 0.003476601 | 0.011798992 |
| LOC119864721 | -1.96654959 | 3.689459717 | 3.47E-05 | 0.00022828 |
| ST6GAL1 | -1.967030126 | 6.914096863 | 0.000156849 | 0.000834203 |
| LOC111097550 | -1.968439632 | 0.359835014 | 0.01867007 | 0.048176737 |
| CASR | -1.968546019 | 0.103404979 | 0.003811801 | 0.01275485 |
| APBB1IP | -1.971126879 | 6.194504382 | 4.74E-05 | 0.000299639 |
| IL27RA | -1.972065903 | 5.643047619 | 0.001404856 | 0.005451646 |
| LOC102156314 | -1.972828652 | -0.587552757 | 0.01556135 | 0.041678342 |
| BAZ1A | -1.972971554 | 6.244779724 | 0.000128444 | 0.000707081 |
| E2F3 | -1.973255093 | 4.637268457 | 4.88E-05 | 0.000307419 |
| LOC111094396 | -1.973329736 | 3.482283364 | 0.008018394 | 0.023960106 |
| LOC102152256 | -1.973477174 | 0.114376862 | 0.004096102 | 0.013526562 |
| LOC119871361 | -1.97474638 | -0.522052032 | 0.006919327 | 0.021121489 |
| LOC111090215 | -1.975078677 | 1.151541905 | 0.008741061 | 0.025736316 |
| LOC111094620 | -1.975260812 | 0.178065019 | 0.003841401 | 0.012841999 |
| RNF125 | -1.975953032 | 4.962891467 | 0.000270775 | 0.001330122 |
| LOC111097670 | -1.976726307 | -0.576885212 | 0.014185127 | 0.038651527 |
| CD58 | -1.977669027 | 5.99349548 | 1.76E-05 | 0.000126017 |
| LOC119865612 | -1.978437273 | 0.651469565 | 0.00513807 | 0.016383313 |
| SP100 | -1.979635344 | 6.834752437 | 6.00E-05 | 0.000367817 |
| ESYT3 | -1.981137211 | 4.15689137 | 0.000176877 | 0.0009235 |
| CASP8 | -1.983643553 | 4.320161249 | 0.000714854 | 0.003066105 |

| | | | | |
|---|---|---|---|---|
| PRKAR2B | -1.984064919 | 5.777605661 | 0.00108273 | 0.004381255 |
| LOC119877921 | -1.985212206 | 3.034036833 | 0.002330981 | 0.008373837 |
| LOC119868063 | -1.986047677 | 1.178715308 | 0.003566779 | 0.01204281 |
| CST7 | -1.988070102 | 2.959611568 | 0.001184557 | 0.004737571 |
| LOC111098970 | -1.988173257 | 2.989113008 | 0.002215197 | 0.008040523 |
| FERMT3 | -1.989357244 | 5.989927424 | 0.001109767 | 0.004477287 |
| FAR2 | -1.989678465 | 3.985385553 | 0.004798001 | 0.015446683 |
| LOC119866913 | -1.990450777 | 4.320056676 | 0.000302581 | 0.001467736 |
| LOC119873834 | -1.990605545 | -0.141460059 | 0.006846683 | 0.020935125 |
| CRYBA1 | -1.991114281 | 0.510899363 | 0.012527547 | 0.034817841 |
| LOC119866890 | -1.991996145 | 1.10595396 | 0.005191254 | 0.016525142 |
| LOC111092526 | -1.994380738 | 0.951077143 | 0.004161133 | 0.013714146 |
| MPP7 | -1.996015535 | 2.492159009 | 0.001419014 | 0.0054938 |
| XPO6 | -1.996141787 | 7.893025022 | 3.37E-07 | 3.96E-06 |
| KLHDC9 | -1.998817926 | 0.868354984 | 0.001244033 | 0.004927253 |
| SNRK | -1.99900207 | 7.340989751 | 1.69E-06 | 1.63E-05 |
| DUSP1 | -2.000221131 | 9.083630411 | 0.000107208 | 0.000607513 |
| LOC102153976 | -2.003409297 | 0.841228913 | 0.00331193 | 0.011314544 |
| LOC111091315 | -2.004158968 | 4.215752981 | 3.37E-06 | 2.96E-05 |
| TSPAN32 | -2.005757694 | 1.060085387 | 0.016034382 | 0.042676307 |
| LOC111096372 | -2.00588108 | 1.440337904 | 0.008984573 | 0.026356612 |
| LOC119868088 | -2.007625965 | 0.763542916 | 0.001690564 | 0.006378973 |
| LOC119873137 | -2.008471376 | -0.474350869 | 0.005405428 | 0.017127462 |
| LOC119872572 | -2.008755377 | -0.245517757 | 0.004244355 | 0.013922793 |
| LOC119872726 | -2.009186569 | 0.990266625 | 0.003240381 | 0.011113906 |
| LOC119870476 | -2.009747139 | 0.685484835 | 0.004752268 | 0.015319933 |
| LOC119866269 | -2.010230447 | 0.722028079 | 0.002052804 | 0.007533088 |
| TTC39C | -2.01561048 | 5.447417605 | 0.000119618 | 0.000666043 |
| LOC102157045 | -2.015655313 | 1.451524583 | 0.001735169 | 0.006515302 |
| ALG9 | -2.015916279 | 5.034240969 | 0.000211002 | 0.00107804 |
| TNIK | -2.016040751 | 6.142958473 | 0.00232731 | 0.008367585 |
| KCNQ5 | -2.016906266 | 3.399360391 | 0.019394579 | 0.049756002 |
| H2BC18 | -2.017001174 | 4.380942647 | 5.07E-06 | 4.20E-05 |
| PTGDR | -2.017364836 | 2.393992133 | 0.017352242 | 0.045421709 |
| ANKRD61 | -2.018731695 | 1.138005273 | 0.0076622 | 0.023048281 |
| PDLIM1 | -2.020198418 | 6.87490868 | 0.001194049 | 0.004767616 |
| SUSD5 | -2.020338245 | 2.37999902 | 0.001137539 | 0.004574853 |
| LOC119873132 | -2.022529188 | 1.326109064 | 0.001121635 | 0.004519276 |
| VAV3 | -2.022721683 | 5.977522831 | 0.002233395 | 0.008089746 |

| | | | | |
|---|---|---|---|---|
| MEIOC | -2.023075761 | -0.013348456 | 0.007457244 | 0.022531835 |
| LAX1 | -2.024024307 | 3.218398191 | 0.017966036 | 0.046721109 |
| LOC102156243 | -2.025734587 | 2.91832512 | 0.000672152 | 0.002912347 |
| GNG2 | -2.026444238 | 7.004685575 | 0.000143658 | 0.000774889 |
| LOC111093028 | -2.026743165 | 1.206475526 | 0.014101132 | 0.038485561 |
| LOC119865944 | -2.027224617 | 0.128886417 | 0.008059903 | 0.024053857 |
| AQP7 | -2.028993483 | 1.562136371 | 0.013048169 | 0.036042949 |
| LOC607833 | -2.029073939 | 0.172908022 | 0.00928219 | 0.027090346 |
| LOC102151560 | -2.029176222 | 1.461356402 | 0.006457284 | 0.019911923 |
| LOC119867506 | -2.02967489 | 1.468297154 | 0.007159211 | 0.021743494 |
| PATJ | -2.031980335 | 4.05965361 | 0.002044378 | 0.007509797 |
| DPYD | -2.033170522 | 7.969880753 | 0.000122873 | 0.00068108 |
| SYCP3 | -2.033279143 | 1.49981271 | 0.000625141 | 0.002735451 |
| RASSF3 | -2.03414668 | 6.932145143 | 5.89E-05 | 0.000362169 |
| LOC106558611 | -2.035291938 | 0.804855215 | 0.004642757 | 0.015036234 |
| LOC119863891 | -2.035336085 | 5.071742684 | 0.000227469 | 0.001146229 |
| LOC111095538 | -2.035680217 | 2.154573356 | 8.66E-05 | 0.000505694 |
| LOC609690 | -2.036003407 | 1.642572118 | 0.001591875 | 0.006062706 |
| EIF4EBP1 | -2.036731922 | 4.501838713 | 6.12E-05 | 0.000374472 |
| LOC119873575 | -2.036966754 | 2.322177803 | 0.000256687 | 0.001271293 |
| LOC484867 | -2.037326111 | 2.502526806 | 0.006481947 | 0.019974866 |
| LOC119876459 | -2.037569094 | 2.931684067 | 0.001700311 | 0.006409005 |
| LOC119872893 | -2.038793477 | 0.362435649 | 0.005421171 | 0.017159783 |
| LOC119878070 | -2.040198066 | 1.482343978 | 0.00166414 | 0.006295499 |
| LOC102155256 | -2.040549563 | 2.232255233 | 0.015436299 | 0.0414446 |
| B2M | -2.040659395 | 9.262303413 | 0.001570254 | 0.005996848 |
| CSF2RB | -2.040931696 | 6.861693129 | 0.00098194 | 0.004030604 |
| PARVG | -2.043344362 | 5.569380199 | 0.000411725 | 0.001914051 |
| VASP | -2.043518144 | 6.429876294 | 8.07E-06 | 6.32E-05 |
| INHBC | -2.044389218 | -0.782409942 | 0.017470532 | 0.045663149 |
| RNF144B | -2.044727282 | 5.652361293 | 3.60E-05 | 0.000235648 |
| LGALSL | -2.044895149 | 3.163316867 | 0.000703306 | 0.003023157 |
| SLC38A8 | -2.045065622 | -0.380790794 | 0.016539133 | 0.043708406 |
| LOC100856400 | -2.045562103 | 0.536457023 | 0.006157806 | 0.019122814 |
| MEGF11 | -2.045628666 | 3.59840292 | 5.88E-05 | 0.0003614 |
| CIDEB | -2.046018864 | 1.339789061 | 0.000608894 | 0.002671909 |
| LOC102156722 | -2.046367295 | -0.052984156 | 0.002090722 | 0.007648924 |
| C22H13orf46 | -2.046703668 | 0.868782004 | 0.005454127 | 0.017233896 |
| LOC119865549 | -2.049425196 | 0.216726827 | 0.013219783 | 0.036447316 |

| | | | | |
|---|---|---|---|---|
| LOC119866944 | -2.051920846 | 0.219282075 | 0.001582089 | 0.006033543 |
| INPP4B | -2.052820888 | 6.540656606 | 0.000520482 | 0.002337424 |
| TMEM71 | -2.053061712 | 4.691595397 | 0.000719893 | 0.003082839 |
| LOC111097371 | -2.054548879 | 1.502685944 | 0.005868111 | 0.018357327 |
| LOC111095637 | -2.055034045 | 0.519876183 | 0.012703776 | 0.035239849 |
| LOC119867582 | -2.05583625 | 1.770551236 | 0.01411085 | 0.038502387 |
| GPR65 | -2.056831776 | 4.010614788 | 0.002232553 | 0.008088619 |
| LOC119864211 | -2.057578567 | 1.688372847 | 0.00071854 | 0.003078869 |
| LOC102155052 | -2.058513042 | 0.107291843 | 0.005490264 | 0.017337972 |
| MIR628 | -2.05975249 | -0.901039625 | 0.017393182 | 0.045506461 |
| TAL1 | -2.060311858 | 4.55210079 | 0.00140043 | 0.00543934 |
| IGF2BP2 | -2.061726053 | 4.038332111 | 0.018260407 | 0.047328126 |
| GRAMD1B | -2.062863545 | 6.199141244 | 0.000140079 | 0.000758414 |
| GMIP | -2.063050698 | 6.575414504 | 1.78E-06 | 1.71E-05 |
| ANKRD44 | -2.063316063 | 7.020620224 | 0.001372424 | 0.005354834 |
| CYRIB | -2.064760773 | 5.820558304 | 9.41E-06 | 7.22E-05 |
| LOC119871201 | -2.066256577 | 0.421581766 | 0.01624595 | 0.043128182 |
| LOC119869474 | -2.06753915 | -0.57687046 | 0.015549526 | 0.041662117 |
| KLHL8 | -2.070188533 | 4.557747286 | 0.000123786 | 0.000684915 |
| CACNA1I | -2.070537553 | 2.178142304 | 0.001038548 | 0.004225351 |
| CCDC69 | -2.073495912 | 4.761668107 | 0.00585773 | 0.018330144 |
| LOC119865701 | -2.073920533 | 4.880124038 | 0.002222908 | 0.00806311 |
| H4C11 | -2.074112628 | -0.400463977 | 0.002212707 | 0.008033827 |
| LOC119877479 | -2.074251466 | 5.901440232 | 6.19E-06 | 4.99E-05 |
| LOC111094686 | -2.075587974 | 3.547225911 | 0.006257922 | 0.019385756 |
| LOC111093376 | -2.076087854 | 1.670597875 | 0.000260252 | 0.001284837 |
| TM4SF20 | -2.078184074 | -0.37505716 | 0.013333476 | 0.036700079 |
| ASPRV1 | -2.079865766 | 0.154738329 | 0.002546724 | 0.009039493 |
| LOC612471 | -2.080495716 | 2.689775878 | 0.001427919 | 0.005522359 |
| LOC111093453 | -2.080990837 | 2.612158991 | 8.81E-05 | 0.000512358 |
| ARHGAP26 | -2.082077189 | 7.222143117 | 8.81E-07 | 9.14E-06 |
| LOC102156844 | -2.082923555 | 2.964535252 | 0.000205092 | 0.001051565 |
| LOC119866974 | -2.084523667 | 0.098569595 | 0.006494963 | 0.020009286 |
| SLC7A7 | -2.085753735 | 4.480621355 | 3.89E-06 | 3.35E-05 |
| MYO1G | -2.085850061 | 5.784610543 | 0.002809796 | 0.009853836 |
| KCND3 | -2.086060853 | 1.751219315 | 0.009261097 | 0.027046997 |
| LOC111097030 | -2.086442991 | 2.464831497 | 0.000130619 | 0.000716105 |
| LOC119873867 | -2.086870649 | 4.53500156 | 7.98E-05 | 0.000471195 |
| LOC119867690 | -2.089106153 | 1.763708062 | 0.001559241 | 0.005960039 |

| | | | | |
|---|---|---|---|---|
| GLA | -2.090378236 | 2.89546684 | 0.00017074 | 0.000896644 |
| LOC119871947 | -2.091136079 | 1.104663862 | 0.002901222 | 0.010117166 |
| NLRC3 | -2.09220433 | 6.217156443 | 0.016195898 | 0.043011113 |
| LOC106557930 | -2.092809506 | 3.318376036 | 3.58E-06 | 3.12E-05 |
| LOC119879077 | -2.093773724 | 0.32691769 | 0.001290256 | 0.005085237 |
| NLK | -2.095587261 | 6.285023871 | 3.66E-05 | 0.000239313 |
| LOC111094229 | -2.096785558 | 1.383299631 | 0.003057702 | 0.010581125 |
| LOC111096601 | -2.097018407 | 3.189560541 | 0.001001142 | 0.004099328 |
| LOC119868511 | -2.097300518 | 1.20456623 | 0.000299084 | 0.001452724 |
| CHN2 | -2.097309641 | 4.304140079 | 3.88E-05 | 0.000251775 |
| LOC119869427 | -2.098722719 | -0.244883753 | 0.001550001 | 0.005932047 |
| ASB11 | -2.098956919 | 0.479061858 | 0.000417133 | 0.001936027 |
| LOC119865951 | -2.101770925 | 2.83934254 | 1.82E-05 | 0.000129758 |
| RNF4 | -2.103625312 | 5.046970379 | 0.000317692 | 0.001530412 |
| LOC119872657 | -2.10565578 | 1.928656606 | 0.007455763 | 0.022530501 |
| LOC119873881 | -2.106695668 | 0.986806104 | 0.010165689 | 0.029259094 |
| LOC102152827 | -2.108529326 | 3.845565258 | 0.000138106 | 0.000749417 |
| TRPC5 | -2.109116843 | 0.852512878 | 0.002599002 | 0.009206393 |
| TTC9 | -2.109785162 | 3.211863582 | 0.000756567 | 0.003217484 |
| SLC16A10 | -2.110277495 | 2.493463146 | 0.003931756 | 0.01309817 |
| LOC111097026 | -2.110953326 | 0.551008155 | 0.016579295 | 0.043787845 |
| LOC106558604 | -2.111212639 | 3.803008654 | 5.24E-06 | 4.32E-05 |
| LOC119868082 | -2.112347655 | 2.965755534 | 0.00058066 | 0.002564692 |
| LOC119872886 | -2.112387648 | 1.530711626 | 0.000645696 | 0.002813463 |
| LOC119872926 | -2.112682842 | 0.836897655 | 0.006453577 | 0.019907899 |
| SEPTIN1 | -2.113109219 | 4.027608907 | 0.000493351 | 0.002232311 |
| LOC111096355 | -2.113242833 | 2.872355282 | 0.000210166 | 0.001074678 |
| LOC100855875 | -2.114629846 | 2.549900193 | 0.002475675 | 0.008826337 |
| LOC489516 | -2.117642769 | 2.487451648 | 0.004209356 | 0.013837307 |
| HCRTR1 | -2.11827535 | 1.664787442 | 0.00014007 | 0.000758414 |
| CD101 | -2.119000083 | 3.961253996 | 0.002255911 | 0.008158244 |
| LOC106560145 | -2.120948713 | 0.724961174 | 0.004660665 | 0.015078458 |
| LOC119871177 | -2.121362091 | 0.980047058 | 0.00251708 | 0.008956276 |
| LOC119866865 | -2.121878712 | 2.862743784 | 0.000489953 | 0.002218502 |
| LOC111097342 | -2.123172983 | 4.874634473 | 1.67E-07 | 2.12E-06 |
| LOC111095925 | -2.123305162 | 3.613968241 | 0.000875424 | 0.003655698 |
| GRK2 | -2.125197476 | 6.83147097 | 8.88E-07 | 9.19E-06 |
| LOC111096825 | -2.126556824 | 3.76137497 | 0.000420565 | 0.00194747 |
| NFIL3 | -2.128365907 | 5.28218862 | 2.21E-05 | 0.000154134 |

| | | | | |
|---|---|---|---|---|
| LOC102156001 | -2.128387755 | 2.607517546 | 0.000441651 | 0.002028398 |
| STBD1 | -2.128971668 | 2.505362467 | 0.000732811 | 0.00313105 |
| UNC13D | -2.129295767 | 5.002114719 | 0.000100189 | 0.000573081 |
| BTG1 | -2.130004902 | 7.817971283 | 1.89E-06 | 1.79E-05 |
| F11 | -2.130827191 | 0.823118738 | 0.015249044 | 0.040998949 |
| LOC119867755 | -2.13100334 | -0.275815171 | 0.009478863 | 0.02756782 |
| ANKRD66 | -2.131200695 | 2.70886241 | 0.017971787 | 0.046721109 |
| EPB41 | -2.132899579 | 7.001981185 | 6.87E-06 | 5.48E-05 |
| LOC119876802 | -2.133716991 | -0.313349468 | 0.019238859 | 0.049416974 |
| MYBPC3 | -2.134505633 | 0.60147569 | 0.002136237 | 0.0077957 |
| CYRIA | -2.137118378 | 4.83568061 | 3.61E-06 | 3.14E-05 |
| ARAP2 | -2.138878203 | 6.310127625 | 0.000226503 | 0.001142426 |
| LOC111095589 | -2.13900298 | 2.648363293 | 0.008183617 | 0.0243464 |
| FUT7 | -2.139458479 | 0.872042063 | 0.006079099 | 0.01891796 |
| STXBP5 | -2.139918058 | 5.931927464 | 0.000116478 | 0.000651972 |
| PREX1 | -2.141066376 | 8.248678622 | 4.10E-05 | 0.000263954 |
| LAT | -2.141143578 | 4.073531912 | 0.011112384 | 0.031502774 |
| LOC119871264 | -2.141354259 | 0.72313383 | 0.01515624 | 0.040794982 |
| LOC111091699 | -2.141844263 | -0.227870766 | 0.00513247 | 0.016370276 |
| LOC119870967 | -2.143683338 | 3.188834556 | 0.000655753 | 0.00285041 |
| LOC111096219 | -2.145221932 | -0.104522658 | 0.001165284 | 0.004669108 |
| LOC106557461 | -2.145830192 | 5.347111071 | 0.001115971 | 0.004498128 |
| PTPRE | -2.147085051 | 6.954018008 | 1.21E-05 | 9.04E-05 |
| TRIM34 | -2.150358095 | 5.341849716 | 0.000339468 | 0.00161648 |
| ST8SIA4 | -2.151297542 | 5.859766002 | 3.93E-05 | 0.000254421 |
| LOC111090999 | -2.1529298 | 3.269479272 | 2.00E-05 | 0.000140678 |
| PLCL2 | -2.155703945 | 5.927188854 | 0.000215549 | 0.001096888 |
| MASP2 | -2.157000548 | 4.142450928 | 0.000143837 | 0.000775273 |
| UBASH3B | -2.157599461 | 6.008136718 | 8.38E-05 | 0.000491284 |
| LOC609800 | -2.160066436 | 4.298880454 | 0.00316027 | 0.010885654 |
| LOC102153458 | -2.160208461 | 0.69924933 | 0.010112268 | 0.029144042 |
| LOC111092061 | -2.162233658 | 0.714158985 | 0.017766427 | 0.046313661 |
| DCST2 | -2.162370334 | 4.072713626 | 6.80E-06 | 5.43E-05 |
| LOC119868454 | -2.162374422 | 0.047130039 | 0.001573891 | 0.00600862 |
| LOC100683370 | -2.162495326 | 7.430714324 | 0.001422195 | 0.005503166 |
| NBEAL2 | -2.162861645 | 7.147068498 | 3.18E-06 | 2.82E-05 |
| CREM | -2.166769195 | 3.698304315 | 2.45E-06 | 2.25E-05 |
| GLRX | -2.166849939 | 6.655496505 | 3.83E-05 | 0.000248985 |
| LOC119867926 | -2.167047032 | 2.454339452 | 0.008345255 | 0.024745753 |

| | | | | |
|---|---|---|---|---|
| LOC106557741 | -2.168201448 | 1.727695152 | 0.009962632 | 0.028777838 |
| LOC102154780 | -2.168536313 | 1.387765134 | 0.002139884 | 0.007805067 |
| LOC111091716 | -2.169188884 | 0.689060517 | 0.00049621 | 0.002244232 |
| AP3M2 | -2.17279691 | 2.534001315 | 0.002026869 | 0.007456856 |
| LOC119866169 | -2.173481599 | 1.784604035 | 0.001223409 | 0.00486245 |
| C35H6orf201 | -2.173547114 | 1.566277803 | 0.000508735 | 0.002292523 |
| E2F2 | -2.174408601 | 3.213215967 | 0.000154802 | 0.000824125 |
| PDE7A | -2.176475428 | 4.437470726 | 0.0002112 | 0.001078543 |
| LOC119865372 | -2.1772897 | 3.679956887 | 7.91E-05 | 0.000467727 |
| LOC119876004 | -2.178852171 | 5.068384121 | 0.000499929 | 0.002258696 |
| LOC102152401 | -2.179361886 | 1.140264049 | 0.000779188 | 0.003301531 |
| LOC119873029 | -2.181554655 | 1.821987641 | 9.04E-05 | 0.000524335 |
| LOC119868252 | -2.182158136 | -0.038597381 | 0.006455686 | 0.019910928 |
| LOC119874081 | -2.18345335 | 0.269965976 | 0.005417827 | 0.017159225 |
| IFITM10 | -2.183560046 | 1.836535966 | 0.009389925 | 0.027367919 |
| TRPM2 | -2.187626253 | 5.695816558 | 7.09E-05 | 0.000424448 |
| GRIN2B | -2.19147388 | -0.627343784 | 0.013235904 | 0.036477842 |
| CDC14A | -2.191523858 | 5.295790777 | 1.27E-05 | 9.40E-05 |
| LOC119871676 | -2.192719326 | 0.082452366 | 0.003006594 | 0.010432074 |
| LOC111095784 | -2.195709038 | 3.759493492 | 0.000427903 | 0.001976546 |
| LOC111093566 | -2.196714106 | 4.092770653 | 0.016762888 | 0.044181962 |
| TXLNB | -2.197376295 | 3.70358409 | 0.001527286 | 0.005856487 |
| DKK4 | -2.197687588 | -0.476870523 | 0.008413504 | 0.024924259 |
| NCKAP1L | -2.199787421 | 6.589445253 | 0.00046771 | 0.00212866 |
| ARHGAP25 | -2.201079071 | 6.024897043 | 2.77E-05 | 0.000187432 |
| LOC111095515 | -2.201378975 | 2.186095015 | 0.001374039 | 0.005356998 |
| LIMD2 | -2.203146447 | 5.782071955 | 0.000179492 | 0.000935805 |
| TRIM72 | -2.204061732 | -0.198683697 | 0.00093284 | 0.003864929 |
| INPP5D | -2.205013008 | 7.18253026 | 1.00E-05 | 7.64E-05 |
| ARG2 | -2.206893546 | 4.712856246 | 4.33E-09 | 8.10E-08 |
| LOC119867834 | -2.208345757 | 3.646082264 | 0.000333894 | 0.001594292 |
| LOC119867490 | -2.208552849 | -0.050216526 | 0.017520004 | 0.045770376 |
| LOC119871469 | -2.20900993 | 3.621065439 | 7.58E-05 | 0.000451199 |
| LOC119871782 | -2.210164217 | 2.072129554 | 0.004511938 | 0.014665152 |
| LOC119865119 | -2.210218033 | 0.132687824 | 0.000577647 | 0.002553925 |
| LOC119872486 | -2.213215726 | 5.00074157 | 0.000638035 | 0.002783439 |
| STK17B | -2.215043645 | 7.165011068 | 0.001372516 | 0.005354834 |
| MX2 | -2.215613731 | 5.882068411 | 0.000350569 | 0.001664013 |
| LOC119873979 | -2.216978634 | 3.644025589 | 0.000302518 | 0.001467736 |

| | | | | |
|---|---|---|---|---|
| MIR24-2 | -2.218056249 | 0.141827854 | 0.000850049 | 0.003560711 |
| LOC490629 | -2.218514248 | 1.885627057 | 0.015614632 | 0.041764274 |
| LOC119873479 | -2.219161989 | 0.338380772 | 0.000538929 | 0.002407296 |
| PSMB9 | -2.220610467 | 4.600946044 | 7.58E-05 | 0.000451342 |
| F13A1 | -2.220989456 | 6.699474184 | 0.010253419 | 0.029449027 |
| LOC119871955 | -2.221163993 | 1.556832092 | 0.000692734 | 0.00298541 |
| LOC119873262 | -2.222031124 | 6.334329711 | 0.000190114 | 0.000982183 |
| LOC119867998 | -2.222383548 | 0.081138263 | 0.005533059 | 0.017455313 |
| PRKAG2 | -2.222582481 | 6.19753429 | 1.36E-07 | 1.78E-06 |
| LOC111093774 | -2.222726583 | -0.376104568 | 0.017186046 | 0.045093604 |
| GLIPR2 | -2.223283832 | 6.90899551 | 5.25E-07 | 5.84E-06 |
| LOC106558733 | -2.22390134 | 1.486250306 | 0.000116133 | 0.000650205 |
| FAM78A | -2.224893105 | 5.049680643 | 3.31E-05 | 0.000218821 |
| LOC102155321 | -2.225057459 | 2.398423378 | 2.63E-06 | 2.40E-05 |
| RNF225 | -2.225752158 | 0.569876065 | 0.000541168 | 0.002415807 |
| LOC102154975 | -2.226160857 | 0.317450289 | 0.017613423 | 0.0459701 |
| IL17C | -2.227275611 | -0.641689833 | 0.002491483 | 0.008878313 |
| LOC119866876 | -2.227900757 | 1.202199326 | 0.000300981 | 0.001460955 |
| LOC111095526 | -2.228094256 | 0.84999339 | 0.001377456 | 0.005368389 |
| DTHD1 | -2.228136913 | 2.923613062 | 0.016945371 | 0.044586259 |
| PIGZ | -2.228147049 | 4.86967396 | 6.16E-06 | 4.98E-05 |
| LOC119878408 | -2.228313788 | 1.057206581 | 0.000111998 | 0.000630967 |
| LOC119878087 | -2.22868056 | 3.208288219 | 0.004567525 | 0.014828036 |
| FGD3 | -2.229166562 | 5.25153202 | 0.002165811 | 0.007885952 |
| PTGS1 | -2.229320344 | 4.912249824 | 0.007946055 | 0.023776729 |
| OGFRL1 | -2.22954527 | 7.693428851 | 9.55E-05 | 0.000549002 |
| LOC119865280 | -2.231804414 | 1.400748386 | 0.001656289 | 0.006266892 |
| LOC111097654 | -2.232391437 | 0.27455601 | 0.001203837 | 0.004798741 |
| SSH2 | -2.232633635 | 8.205452686 | 3.28E-07 | 3.87E-06 |
| IL1R2 | -2.234931563 | 1.706831741 | 0.011622572 | 0.032671336 |
| LOC119878870 | -2.234938639 | -1.24108319 | 0.004236856 | 0.013900472 |
| LOC111095729 | -2.235191677 | -0.963906303 | 0.010855254 | 0.03088001 |
| GCNA | -2.235671879 | 2.366101399 | 0.000812769 | 0.003422408 |
| PIK3CG | -2.235975298 | 6.043811519 | 2.66E-05 | 0.000181307 |
| LOC111090251 | -2.236642031 | 3.028565726 | 0.001834542 | 0.006844558 |
| LOC119873498 | -2.236692393 | 0.659290444 | 0.01532443 | 0.041170991 |
| LOC102152592 | -2.237019003 | 0.962451324 | 0.000336761 | 0.001605503 |
| LOC102157137 | -2.23815564 | 4.089657553 | 0.000200296 | 0.001029406 |
| LOC100682812 | -2.239189813 | 0.980265252 | 0.000169083 | 0.000889186 |

| | | | | |
|---|---|---|---|---|
| ADAM19 | -2.239766288 | 7.676459208 | 3.00E-06 | 2.68E-05 |
| RASGRP1 | -2.241184191 | 4.591597936 | 7.02E-05 | 0.000421404 |
| LMBR1L | -2.242487822 | 4.756957238 | 1.96E-05 | 0.000138793 |
| LOC119871977 | -2.242644816 | 0.243185987 | 0.003472196 | 0.011787737 |
| LOC111093322 | -2.242882455 | 2.361687077 | 0.000145568 | 0.000783241 |
| EYS | -2.244212989 | 2.14448879 | 0.000911824 | 0.003790172 |
| C1QTNF12 | -2.245959488 | 0.218689929 | 0.000105107 | 0.000597895 |
| RSAD2 | -2.246505023 | 4.426915827 | 0.014456977 | 0.039285386 |
| CCR10 | -2.247555005 | 0.694557165 | 0.003015738 | 0.010454269 |
| TNFRSF25 | -2.24790698 | 4.873919045 | 1.63E-05 | 0.000117508 |
| BCL6 | -2.248563404 | 6.755759259 | 2.73E-06 | 2.47E-05 |
| GPR3 | -2.248646097 | 0.880043351 | 0.001352956 | 0.005290981 |
| CCDC177 | -2.249615394 | 1.892856328 | 0.000832854 | 0.003499505 |
| BNIPL | -2.249647476 | 2.07829181 | 5.06E-05 | 0.000317458 |
| LOC111096818 | -2.250241992 | -0.579242616 | 0.010904856 | 0.030995534 |
| LOC106559357 | -2.25086188 | 0.185167525 | 0.000661488 | 0.002871308 |
| LOC102155033 | -2.252868288 | 0.250784424 | 0.017168712 | 0.045064694 |
| RASGRP2 | -2.253097019 | 6.107832708 | 0.000145836 | 0.000784185 |
| LOC111091330 | -2.253465136 | 3.520172322 | 0.001342658 | 0.005255302 |
| FLI1 | -2.254263763 | 6.787818936 | 4.51E-06 | 3.82E-05 |
| RAB11FIP4 | -2.254785522 | 5.323320936 | 3.69E-05 | 0.000240783 |
| LOC119878895 | -2.257393978 | -0.709015649 | 0.01885701 | 0.048560834 |
| DIPK1A | -2.257797121 | 4.015484961 | 1.80E-05 | 0.000128355 |
| MBOAT1 | -2.258728172 | 5.595480548 | 1.57E-06 | 1.52E-05 |
| LOC490173 | -2.259038296 | 0.045374327 | 0.004900646 | 0.015730405 |
| LOC119873705 | -2.259072179 | 4.010419793 | 0.000175703 | 0.000918922 |
| LOC111093751 | -2.260099099 | 0.424853784 | 0.001240541 | 0.004917014 |
| LOC119864338 | -2.260540408 | -0.480965747 | 0.00429203 | 0.014051535 |
| TNF | -2.261683153 | 2.004387812 | 0.007334063 | 0.022212289 |
| LONRF3 | -2.262069398 | 6.221871239 | 4.20E-07 | 4.79E-06 |
| STAG3 | -2.262286778 | -0.329081989 | 0.007341914 | 0.022232962 |
| MIR142 | -2.265023444 | 0.54850342 | 0.017393786 | 0.045506461 |
| LOC111091660 | -2.265743588 | 2.088944147 | 0.000252781 | 0.001254815 |
| LOC111097362 | -2.272774453 | 0.189465634 | 0.000185254 | 0.000961218 |
| HNF4A | -2.272886387 | 1.437282237 | 0.00419177 | 0.01380043 |
| PIK3CD | -2.272943543 | 7.257906173 | 2.54E-05 | 0.000173857 |
| LOC106559748 | -2.274920428 | 1.631507175 | 0.001098823 | 0.004439746 |
| ARHGAP9 | -2.277452566 | 5.269894537 | 0.000283253 | 0.001383893 |
| LOC102156776 | -2.277469628 | -0.666470592 | 0.016398629 | 0.043437727 |

| | | | | |
|---|---|---|---|---|
| PHOSPHO1 | -2.277703043 | 2.788767394 | 0.004134753 | 0.013639649 |
| LOC111092186 | -2.278477378 | 0.760101148 | 0.002899239 | 0.010115133 |
| MAL2 | -2.280864879 | 0.160908714 | 0.004122688 | 0.013606062 |
| FASLG | -2.28148635 | 1.502907663 | 0.003094075 | 0.010686517 |
| LOC102152268 | -2.28160381 | 2.332284394 | 0.000112655 | 0.000633683 |
| LOC119873152 | -2.28232418 | -0.901142636 | 0.006320845 | 0.019553478 |
| LOC119872245 | -2.28263682 | 3.418755588 | 0.003300178 | 0.01127795 |
| LOC102156117 | -2.283740679 | 4.643600676 | 0.002330698 | 0.008373837 |
| MIR339-1 | -2.28429664 | -0.373840327 | 0.00067314 | 0.002914294 |
| MFNG | -2.285074844 | 4.624075918 | 5.45E-05 | 0.000338347 |
| SLURP1 | -2.285309002 | 0.343222337 | 0.000583028 | 0.002572991 |
| LOC119873335 | -2.285934466 | 1.657151424 | 0.004721777 | 0.015237562 |
| LFNG | -2.287080948 | 5.86247396 | 0.000322717 | 0.001550958 |
| LOC119878058 | -2.287943583 | 1.310181749 | 0.001421146 | 0.005500089 |
| LOC111098964 | -2.288574365 | -1.009324729 | 0.005537662 | 0.01746729 |
| SPATA32 | -2.290442985 | 3.760044462 | 0.000191926 | 0.000989675 |
| LOC111095229 | -2.291016362 | 1.724214191 | 0.00045686 | 0.002086289 |
| LOC119871826 | -2.292230832 | 0.563429211 | 0.012222401 | 0.034079005 |
| NFKBID | -2.292317461 | 4.191587874 | 5.34E-07 | 5.93E-06 |
| LOC119867922 | -2.293445497 | 3.228809851 | 0.000431373 | 0.001989434 |
| CD300A | -2.293589401 | 3.064681248 | 0.000418001 | 0.001939294 |
| MZB1 | -2.295153625 | 2.127378151 | 0.011715617 | 0.03289449 |
| GRTP1 | -2.296107233 | 2.287770444 | 0.000275051 | 0.001347466 |
| ARRB2 | -2.296860638 | 5.858737088 | 3.84E-07 | 4.44E-06 |
| LOC479821 | -2.297001987 | -0.492175432 | 0.003043881 | 0.010543398 |
| LOC106559562 | -2.297116788 | 4.566546475 | 0.000490009 | 0.002218502 |
| LRRC10 | -2.298944648 | -0.633839403 | 0.008034742 | 0.023995722 |
| PTPN22 | -2.300404623 | 4.235878515 | 0.002062029 | 0.007561819 |
| LOC119866613 | -2.300496468 | 0.554812126 | 0.002295551 | 0.008273985 |
| MIR8891 | -2.300603061 | 0.113922729 | 0.002813885 | 0.009866579 |
| BMX | -2.301207033 | 3.438958388 | 0.006435862 | 0.019866747 |
| LOC119872147 | -2.303407028 | 0.755511091 | 0.005872588 | 0.018366032 |
| RNF183 | -2.303593175 | 0.816612499 | 0.001021734 | 0.004167115 |
| VCPKMT | -2.305010673 | 3.857816279 | 1.37E-06 | 1.35E-05 |
| CCR9 | -2.305521677 | 0.148385046 | 0.010994914 | 0.031210578 |
| C5AR2 | -2.307715009 | 3.499037825 | 0.001841265 | 0.006863732 |
| LOC119865356 | -2.309515561 | 4.416906846 | 2.02E-05 | 0.000142306 |
| SEC11C | -2.309820086 | 5.59223818 | 1.24E-06 | 1.24E-05 |
| LOC111093907 | -2.314330311 | 1.576520993 | 0.000129238 | 0.000710184 |

| | | | | |
|---|---|---|---|---|
| LOC119870389 | -2.31585767 | 2.498442947 | 1.31E-06 | 1.30E-05 |
| LOC102156801 | -2.316965523 | 0.723132857 | 0.000126902 | 0.000700013 |
| LOC119873497 | -2.317449266 | 0.084492241 | 0.018524016 | 0.047868248 |
| LOC111095264 | -2.319426737 | 0.229574169 | 0.001549848 | 0.005932047 |
| H2BC14 | -2.319571557 | -0.446389063 | 0.002110252 | 0.007713063 |
| LOC119865710 | -2.32051225 | 1.306613386 | 0.000140577 | 0.00076054 |
| LOC102153988 | -2.320938082 | 4.679530647 | 0.015677327 | 0.041900938 |
| DENND5A | -2.321931627 | 7.742539218 | 1.14E-07 | 1.53E-06 |
| ITGAL | -2.322825219 | 6.605167139 | 0.000187637 | 0.000970793 |
| CCSER1 | -2.323303676 | 3.325146531 | 0.002136653 | 0.007795907 |
| LOC119871936 | -2.324050955 | 1.824465746 | 4.60E-05 | 0.000292377 |
| POU1F1 | -2.324479145 | 0.228246917 | 0.012324965 | 0.034342885 |
| LOC119864193 | -2.32578295 | 2.703783464 | 0.018148528 | 0.047100114 |
| CD27 | -2.326310868 | 4.327672009 | 0.007247015 | 0.021979368 |
| STK4 | -2.327213574 | 6.40037917 | 2.58E-08 | 4.04E-07 |
| TPST2 | -2.328272231 | 5.378000142 | 8.61E-07 | 8.96E-06 |
| LOC119867958 | -2.329133803 | 0.090894809 | 0.005828485 | 0.018267642 |
| LOC111098236 | -2.329828801 | 2.263703687 | 0.001840121 | 0.006860649 |
| LOC106559158 | -2.330151115 | 1.215799959 | 0.006606352 | 0.020291875 |
| LOC119873791 | -2.332712517 | 0.047852569 | 0.008919222 | 0.026182621 |
| KMO | -2.332992003 | 1.99603191 | 0.003226597 | 0.011068383 |
| LOC119869757 | -2.335834984 | 1.244079136 | 0.009471752 | 0.027550836 |
| RAB32 | -2.336999951 | 5.212594048 | 6.89E-07 | 7.39E-06 |
| LOC111098844 | -2.337069757 | -0.732994071 | 0.004281154 | 0.014022283 |
| PFKFB2 | -2.337300457 | 6.222705016 | 3.28E-09 | 6.33E-08 |
| LITAF | -2.338860285 | 6.062680807 | 4.49E-06 | 3.80E-05 |
| LOC111091679 | -2.33985281 | 0.337867471 | 0.002890716 | 0.010095147 |
| LOC119865480 | -2.3410689 | -0.309966981 | 0.001829368 | 0.006829955 |
| AHSA2P | -2.343951006 | 1.519550387 | 0.000420484 | 0.00194747 |
| MCTP1 | -2.344171264 | 5.558984993 | 1.92E-05 | 0.000136021 |
| RXFP1 | -2.344857419 | 1.028818017 | 0.000426843 | 0.00197291 |
| SNN | -2.34501168 | 5.151842413 | 2.68E-07 | 3.23E-06 |
| ERICH6B | -2.347316131 | 0.796189555 | 0.000571741 | 0.002531429 |
| CRYBG2 | -2.348238712 | 1.79043829 | 0.000530812 | 0.00237643 |
| LOC111094821 | -2.348645431 | -0.352397385 | 0.00252718 | 0.008987294 |
| LOC119871503 | -2.350204114 | 2.047036253 | 7.30E-05 | 0.00043673 |
| DOK3 | -2.35047944 | 2.812604844 | 0.000109389 | 0.000618518 |
| TRNAK-CUU_7 | -2.350778543 | -0.163402455 | 0.001809133 | 0.006761396 |
| RNF43 | -2.351464141 | 3.131867239 | 0.00011066 | 0.000625052 |

| | | | | |
|---|---|---|---|---|
| CRLF3 | -2.351508295 | 5.488396702 | 9.47E-07 | 9.76E-06 |
| LOC119872725 | -2.352044072 | 0.374014564 | 0.001462669 | 0.005641649 |
| NCF1 | -2.352130396 | 4.465474102 | 0.000686157 | 0.002960599 |
| LOC111095500 | -2.353510099 | 0.295009626 | 0.005001457 | 0.01600187 |
| LOC111096129 | -2.353620168 | -1.034349071 | 0.000130848 | 0.000716855 |
| LOC111093657 | -2.354383401 | -0.700833376 | 0.011848545 | 0.033207489 |
| LOC100683019 | -2.354947316 | 0.209061712 | 0.000324281 | 0.00155593 |
| ARHGDIB | -2.355553602 | 6.870948659 | 3.52E-05 | 0.000231053 |
| TENT5C | -2.356051477 | 6.049024292 | 0.000357601 | 0.001691833 |
| DOCK11 | -2.356375955 | 7.296177028 | 4.08E-05 | 0.000262909 |
| AMDHD1 | -2.357213555 | 3.709961963 | 3.09E-07 | 3.68E-06 |
| APOBR | -2.357986109 | 5.201373991 | 0.000649025 | 0.002826263 |
| TCF7 | -2.358251377 | 6.872783672 | 3.41E-05 | 0.000224879 |
| LOC106557734 | -2.358572694 | 4.00581452 | 4.14E-07 | 4.74E-06 |
| LOC119864152 | -2.358622679 | 3.998721718 | 0.001928302 | 0.007136628 |
| ROGDI | -2.359637379 | 3.773186865 | 5.72E-07 | 6.31E-06 |
| LOC119876056 | -2.359730305 | 0.06095721 | 0.010139162 | 0.029209897 |
| IL32 | -2.360414532 | 0.899873959 | 0.001718416 | 0.006461352 |
| LOC119867660 | -2.361136551 | 0.646083768 | 0.000270191 | 0.001327555 |
| FMNL1 | -2.3622823 | 7.375454891 | 6.59E-05 | 0.000398592 |
| LOC486692 | -2.363299134 | 3.249669972 | 0.005160766 | 0.016446007 |
| SKAP1 | -2.366746526 | 3.190591269 | 0.003006929 | 0.010432074 |
| LOC119867194 | -2.36724616 | 0.341988067 | 0.002171131 | 0.00790175 |
| LOC119872732 | -2.367791367 | 1.076089959 | 6.72E-05 | 0.000405305 |
| LPAR5 | -2.369699554 | 4.418910904 | 0.000394383 | 0.001843316 |
| LOC119873934 | -2.37040766 | 0.971678944 | 9.06E-05 | 0.00052529 |
| LOC119867469 | -2.371102933 | 3.925929499 | 1.91E-05 | 0.00013513 |
| LBR | -2.372707765 | 7.143099002 | 1.23E-05 | 9.15E-05 |
| LOC111095173 | -2.37418265 | 2.213349859 | 3.82E-05 | 0.00024808 |
| LOC119864699 | -2.375929876 | 1.748589391 | 0.000107134 | 0.000607354 |
| LOC119870718 | -2.376143484 | 0.495233085 | 0.003085321 | 0.010659677 |
| LOC119877252 | -2.377120872 | 1.794895986 | 0.014194259 | 0.03867155 |
| DUSP5 | -2.377368656 | 4.596482203 | 0.000105169 | 0.000598094 |
| GPR137B | -2.377492707 | 6.598117607 | 1.20E-08 | 2.03E-07 |
| LOC102151249 | -2.378479783 | 0.565373337 | 0.000127294 | 0.000701997 |
| LOC119873642 | -2.378635538 | -0.081485396 | 0.008225828 | 0.024448487 |
| LOC491979 | -2.38161426 | 2.487222662 | 0.008740907 | 0.025736316 |
| FRAT1 | -2.382049576 | 2.341805697 | 1.09E-05 | 8.22E-05 |
| TESC | -2.383296039 | 2.837277196 | 0.000762205 | 0.00323907 |

| | | | | |
|---|---|---|---|---|
| ANKRD22 | -2.386930542 | 2.022253659 | 0.000405274 | 0.001887304 |
| PPIC | -2.390903659 | 5.702755556 | 1.47E-05 | 0.000107229 |
| SLC11A1 | -2.394629527 | 5.818317026 | 8.78E-07 | 9.11E-06 |
| LOC480788 | -2.396212869 | 4.604123777 | 0.011602455 | 0.032620722 |
| GPR152 | -2.396330053 | -1.184501452 | 0.002181303 | 0.007933441 |
| TIFAB | -2.396462187 | 1.086050682 | 0.00512203 | 0.016339381 |
| PGD | -2.397250089 | 7.741666413 | 1.15E-07 | 1.53E-06 |
| LOC111092023 | -2.398317425 | 3.522881448 | 5.79E-07 | 6.37E-06 |
| LOC102156098 | -2.40157786 | 1.744546462 | 0.00023245 | 0.0011678 |
| GAPDH | -2.40160373 | 8.994777612 | 1.35E-07 | 1.77E-06 |
| LOC119872191 | -2.402084016 | 0.190570289 | 0.000779999 | 0.003303679 |
| LOC111097599 | -2.402178279 | -0.369086804 | 0.001101423 | 0.00444942 |
| LOC119876528 | -2.40329297 | 0.218823791 | 0.000338259 | 0.001611935 |
| LOC111091500 | -2.404115347 | 1.788221704 | 0.000685298 | 0.002958146 |
| LOC119873704 | -2.407133201 | 2.040224221 | 0.001131719 | 0.00455398 |
| LOC119864830 | -2.407546928 | -0.426685447 | 0.001546682 | 0.005921434 |
| CD6 | -2.407574284 | 5.546271579 | 0.014540553 | 0.039461232 |
| LOC102155845 | -2.408860107 | 0.674280949 | 0.000264726 | 0.001304847 |
| LCN2 | -2.410957831 | 2.49538561 | 0.018329407 | 0.047472901 |
| LOC119871686 | -2.41124044 | 0.297830707 | 0.001106353 | 0.004465174 |
| HACD4 | -2.411716297 | 4.489709051 | 1.15E-06 | 1.16E-05 |
| LOC119866915 | -2.411876988 | 0.261612966 | 0.000212601 | 0.001083912 |
| ABHD12B | -2.412731399 | 4.393547092 | 1.34E-06 | 1.32E-05 |
| LOC111094184 | -2.412873266 | 5.475467108 | 1.13E-08 | 1.92E-07 |
| LOC119871807 | -2.413564407 | 3.670673954 | 0.000408295 | 0.001900145 |
| LOC100683001 | -2.415216122 | 4.570639985 | 4.65E-07 | 5.27E-06 |
| LOC111095296 | -2.415699516 | -0.48516274 | 0.002206512 | 0.008015711 |
| LOC111097600 | -2.416569901 | 2.303148961 | 7.46E-05 | 0.000445007 |
| MIR8893 | -2.416602475 | 0.206049388 | 0.002447427 | 0.008737124 |
| LOC111091917 | -2.416921005 | 2.131223932 | 0.000348317 | 0.001654411 |
| LOC119867530 | -2.417831235 | 2.234250729 | 0.000143637 | 0.000774889 |
| IL10RA | -2.419249365 | 6.446940079 | 8.22E-07 | 8.61E-06 |
| STK26 | -2.420358655 | 4.527343272 | 0.000673052 | 0.002914294 |
| LOC102153224 | -2.420617898 | 1.714607544 | 0.000186894 | 0.000967874 |
| LOC111095886 | -2.420956176 | 0.943139588 | 0.005755911 | 0.018076778 |
| GIMAP8 | -2.421503762 | 5.086193741 | 6.25E-05 | 0.000380874 |
| LOC480466 | -2.422090598 | 1.278736841 | 0.000687276 | 0.002964837 |
| PYGL | -2.422291655 | 6.377765038 | 9.13E-07 | 9.43E-06 |
| LOC119865641 | -2.423821932 | -0.216829455 | 0.000324217 | 0.00155593 |

| | | | | |
|---|---|---|---|---|
| LOC111092341 | -2.42469027 | -0.278292962 | 0.000857823 | 0.003589683 |
| MKRN1 | -2.42798752 | 5.251930982 | 1.68E-10 | 4.38E-09 |
| LOC119864685 | -2.428426547 | 1.01328119 | 0.000135618 | 0.000738876 |
| RUBCNL | -2.428576019 | 4.438556537 | 0.002247029 | 0.008132908 |
| LOC100688969 | -2.429366366 | -0.067283705 | 7.06E-05 | 0.000423404 |
| LOC119866361 | -2.430017534 | 1.793939802 | 0.017526785 | 0.045777057 |
| LOC119874344 | -2.432398645 | 1.489646393 | 0.001677637 | 0.006342128 |
| CIBAR2 | -2.432418346 | 0.009825158 | 0.00010161 | 0.000580442 |
| CISH | -2.432635905 | 4.446823506 | 1.45E-07 | 1.88E-06 |
| LOC119872542 | -2.433920704 | 2.193300415 | 2.70E-05 | 0.00018314 |
| SLC25A18 | -2.434303154 | 1.247191252 | 0.000934725 | 0.003869779 |
| SLC16A6 | -2.434665163 | 3.770737722 | 0.000361123 | 0.001707377 |
| LOC119873670 | -2.435273487 | -0.202396005 | 0.001445445 | 0.005584159 |
| NRAP | -2.43578624 | 0.348801518 | 0.000937246 | 0.003877995 |
| LOC111093134 | -2.43662349 | 0.391561591 | 0.000135899 | 0.000740219 |
| LOC119877251 | -2.436633514 | 2.020443346 | 0.00288573 | 0.01008309 |
| CD80 | -2.437944582 | 4.075487672 | 3.25E-06 | 2.87E-05 |
| LTB4R2 | -2.437973209 | 1.04119959 | 0.00058613 | 0.002584049 |
| LOC111094295 | -2.438379151 | 0.019843754 | 0.000619595 | 0.002714469 |
| MAP3K15 | -2.438617424 | 3.528896414 | 1.58E-05 | 0.000114412 |
| LOC106557777 | -2.438636352 | 2.932828822 | 0.000296514 | 0.001441856 |
| SIT1 | -2.438836833 | 2.675898034 | 0.000255981 | 0.001268088 |
| CMTM7 | -2.439861273 | 5.713943485 | 1.33E-06 | 1.32E-05 |
| ABCC11 | -2.440121477 | -0.616039184 | 0.005289563 | 0.016779962 |
| LOC119866706 | -2.440905755 | 2.557794714 | 0.001415324 | 0.005482454 |
| SASH3 | -2.442396107 | 6.23527529 | 0.000425613 | 0.001968062 |
| SGK1 | -2.442667489 | 6.777118691 | 2.28E-06 | 2.11E-05 |
| LOC111094684 | -2.446036723 | 0.048133282 | 0.000965882 | 0.003973721 |
| CBY2 | -2.44736009 | 1.120679753 | 0.000225746 | 0.001139931 |
| ITGB7 | -2.4480155 | 6.590739943 | 3.62E-07 | 4.22E-06 |
| LOC119876146 | -2.448257617 | 1.105776651 | 0.001295039 | 0.005102973 |
| LOC119874123 | -2.44856388 | 0.117990914 | 0.001246856 | 0.004936631 |
| LOC102151554 | -2.451154358 | -0.252369334 | 0.003076353 | 0.010632082 |
| LOC111097945 | -2.452645268 | -1.132971601 | 0.003360008 | 0.011452448 |
| CHST15 | -2.454234785 | 5.909392101 | 3.65E-05 | 0.00023858 |
| LOC111095908 | -2.455987557 | 1.657014066 | 0.000232171 | 0.001167209 |
| METTL7A | -2.457980693 | 5.114793622 | 0.00012576 | 0.000694417 |
| LOC119878082 | -2.458992774 | 1.824335854 | 0.000732672 | 0.00313105 |
| LOC119872549 | -2.462534885 | 2.394710517 | 0.001326525 | 0.005200755 |

| | | | | |
|---|---|---|---|---|
| LOC119870504 | -2.466366715 | 4.053480089 | 3.38E-05 | 0.000223161 |
| CYB5R4 | -2.466834169 | 4.377008101 | 1.56E-09 | 3.27E-08 |
| LOC106560115 | -2.467259732 | 2.413759777 | 2.69E-05 | 0.000182755 |
| LOC119864011 | -2.46831074 | 2.722361477 | 0.000554072 | 0.00246377 |
| LOC119874162 | -2.469451702 | 3.306484883 | 1.45E-06 | 1.43E-05 |
| LINGO4 | -2.471812605 | -0.048381846 | 0.007158942 | 0.021743494 |
| SH2D1A | -2.472503242 | 3.004597307 | 0.007909089 | 0.023679697 |
| LOC111091484 | -2.473621104 | -0.803617986 | 0.003571678 | 0.012055597 |
| SKA3 | -2.473868593 | 2.938832796 | 1.45E-05 | 0.000106105 |
| LOC100682736 | -2.477454095 | 0.971748686 | 1.88E-05 | 0.000133475 |
| IL18 | -2.47937181 | 4.668923332 | 0.001533929 | 0.005879881 |
| AFF2 | -2.479388647 | 3.896214109 | 1.25E-05 | 9.27E-05 |
| LCP2 | -2.480068588 | 7.25030862 | 0.000273838 | 0.001343583 |
| SMIM2 | -2.480924635 | 0.810669782 | 0.001372121 | 0.005354834 |
| SKAP2 | -2.484005507 | 7.038568516 | 1.21E-08 | 2.04E-07 |
| LOC119864701 | -2.484672663 | 3.516546953 | 3.51E-07 | 4.10E-06 |
| MILR1 | -2.484876519 | 3.95664186 | 7.88E-05 | 0.000466268 |
| LOC106559460 | -2.485168134 | 1.499936528 | 0.00037222 | 0.0017522 |
| GYG1 | -2.48654124 | 5.571842192 | 3.78E-09 | 7.17E-08 |
| SAMSN1 | -2.486820598 | 4.344372029 | 4.40E-05 | 0.000280739 |
| LOC119870963 | -2.487132149 | 3.016620484 | 9.08E-07 | 9.39E-06 |
| LOC106559676 | -2.487448963 | -0.271791435 | 0.000615612 | 0.002698657 |
| LOC119867380 | -2.488503725 | -0.199833482 | 0.002541444 | 0.009026668 |
| IRAG2 | -2.4885417 | 4.930012786 | 0.000884143 | 0.003688553 |
| LOC111092727 | -2.488806146 | -0.917730388 | 0.017970264 | 0.046721109 |
| LCK | -2.489431638 | 4.925943637 | 0.002345706 | 0.008418362 |
| LOC102152396 | -2.489904003 | 2.677570526 | 0.000845044 | 0.003542904 |
| PTPN6 | -2.49680012 | 6.485979127 | 0.000346695 | 0.001647428 |
| LOC119865505 | -2.497373775 | -0.908600517 | 0.005937566 | 0.018547832 |
| HIGD1C | -2.498550994 | 0.389443274 | 0.000800918 | 0.003377331 |
| LOC111098549 | -2.498608446 | 1.276315112 | 0.01864649 | 0.048133083 |
| ZNF683 | -2.500934116 | 2.230476704 | 0.007502621 | 0.022649995 |
| LOC119865967 | -2.502457068 | 3.823124602 | 1.90E-06 | 1.80E-05 |
| RGS14 | -2.504715041 | 4.755749088 | 0.000497176 | 0.002247661 |
| LOC119865018 | -2.506074195 | 6.425819245 | 0.00093369 | 0.003866972 |
| LOC119869821 | -2.509310617 | 0.307941163 | 0.000988341 | 0.004053039 |
| LOC476396 | -2.50951149 | 2.503474002 | 0.000145165 | 0.000781268 |
| LOC119867195 | -2.512461874 | 3.866789172 | 9.01E-05 | 0.000523037 |
| LOC119864898 | -2.513083665 | 0.087969963 | 0.001281064 | 0.005058217 |

| | | | | |
|---|---|---|---|---|
| KYNU | -2.513496234 | 3.748505997 | 0.002143245 | 0.007815813 |
| LOC119873833 | -2.514722982 | -0.300310418 | 4.14E-05 | 0.000266067 |
| LOC119877474 | -2.516651381 | 2.279026317 | 0.000972269 | 0.003994691 |
| C20H19orf38 | -2.517028989 | 2.610066009 | 9.72E-06 | 7.44E-05 |
| TLR1 | -2.518254216 | 3.910278515 | 0.001147783 | 0.004609208 |
| CAT | -2.518541493 | 6.992704002 | 1.84E-08 | 2.98E-07 |
| LOC111097382 | -2.520440453 | 0.338986466 | 0.001722235 | 0.006472342 |
| TMEM156 | -2.52247247 | 3.461982525 | 0.000354562 | 0.00168039 |
| LOC111093070 | -2.523289617 | 1.779304651 | 0.017064943 | 0.044852539 |
| ASTL | -2.526306834 | -0.867632609 | 0.014053804 | 0.038390231 |
| LOC111093943 | -2.527113448 | 0.463622375 | 0.00224959 | 0.008139459 |
| LOC119876918 | -2.528166654 | 5.136682186 | 0.000133953 | 0.000731092 |
| LOC119873637 | -2.528851281 | -0.193572371 | 9.08E-05 | 0.000526411 |
| LOC119865375 | -2.530215819 | 2.884913986 | 0.011537332 | 0.032494945 |
| LOC119865248 | -2.531548029 | -0.427238109 | 0.000170623 | 0.000896248 |
| LY9 | -2.531681844 | 5.981130112 | 0.000715921 | 0.003070074 |
| LOC111093580 | -2.532091557 | 3.612976465 | 1.69E-05 | 0.000121247 |
| GCNT3 | -2.532316709 | -0.371242307 | 0.001320364 | 0.005184425 |
| GPSM3 | -2.532813186 | 5.180505289 | 2.50E-08 | 3.92E-07 |
| LOC102152682 | -2.533340366 | 0.071351258 | 0.000119848 | 0.000666874 |
| SHCBP1L | -2.533986117 | 0.084191821 | 0.003128893 | 0.010794731 |
| LOC106559238 | -2.53411046 | -0.526660052 | 0.000403074 | 0.001878268 |
| ADGRG3 | -2.534887501 | 2.125207966 | 4.41E-05 | 0.000281368 |
| MAPK14 | -2.536749533 | 5.961466474 | 1.84E-10 | 4.75E-09 |
| RPS6KA1 | -2.538689504 | 5.653695143 | 6.78E-06 | 5.42E-05 |
| PDE6C | -2.540621596 | -0.363754567 | 0.00185728 | 0.006913917 |
| CATIP | -2.542290917 | 0.998361443 | 0.000122864 | 0.00068108 |
| LOC111095917 | -2.542555799 | 1.898667212 | 0.000146266 | 0.000785631 |
| MBOAT7 | -2.542557382 | 6.232607767 | 6.08E-10 | 1.40E-08 |
| LOC111094587 | -2.542618612 | 2.032870018 | 0.000131178 | 0.000718114 |
| LOC119866255 | -2.543728576 | 1.824909743 | 0.000136972 | 0.000744747 |
| LOC119877669 | -2.544102625 | 2.438885466 | 0.015591479 | 0.041717792 |
| LOC111095118 | -2.54435065 | 4.511880086 | 0.000442162 | 0.00203029 |
| LOC111098741 | -2.54812869 | 1.151013286 | 5.77E-05 | 0.00035602 |
| LOC119871793 | -2.548717963 | 2.682668191 | 0.00075149 | 0.003201138 |
| LOC119865164 | -2.54914986 | 3.223324667 | 1.01E-06 | 1.04E-05 |
| JCHAIN | -2.551236922 | 6.27897836 | 0.00025853 | 0.001277208 |
| LOC111095591 | -2.551806873 | 1.949308506 | 8.99E-06 | 6.94E-05 |
| GZMA | -2.551811883 | 2.112914277 | 0.01037783 | 0.029751152 |

| | | | | |
|---|---|---|---|---|
| H2AC15 | -2.553337403 | 1.323063224 | 7.59E-05 | 0.000451524 |
| LOC119878872 | -2.556258431 | 0.690811824 | 0.008393641 | 0.024872215 |
| RESF1 | -2.55699403 | 7.974946298 | 3.93E-05 | 0.000254855 |
| CYSLTR1 | -2.558277279 | 2.979885341 | 0.000681999 | 0.002947349 |
| INSYN2B | -2.559943492 | 4.442964307 | 0.008642096 | 0.025514256 |
| CYFIP2 | -2.562224761 | 6.994720265 | 1.10E-05 | 8.31E-05 |
| LOC102152224 | -2.563195765 | 2.51203628 | 1.07E-05 | 8.08E-05 |
| GPBAR1 | -2.563414943 | 3.406689544 | 2.67E-07 | 3.21E-06 |
| LOC119869376 | -2.564363748 | 2.182824494 | 0.002321533 | 0.008349582 |
| IRF5 | -2.564376068 | 5.938251613 | 4.77E-05 | 0.000301675 |
| LOC119866412 | -2.565502265 | 4.853846092 | 2.61E-06 | 2.38E-05 |
| KLHL35 | -2.566388843 | 2.315479601 | 3.60E-06 | 3.13E-05 |
| ZCWPW1 | -2.566566461 | 3.703923269 | 2.68E-07 | 3.23E-06 |
| FEM1C | -2.568135256 | 6.88281421 | 2.07E-10 | 5.28E-09 |
| SLC18A2 | -2.568697591 | 4.18929209 | 4.04E-06 | 3.46E-05 |
| SAG | -2.568883088 | -0.400529988 | 0.000267357 | 0.001315721 |
| LOC610636 | -2.56927191 | 3.412436166 | 0.005206721 | 0.016559288 |
| LOC111094368 | -2.569275321 | -0.376249334 | 0.009519215 | 0.027662905 |
| FAM83E | -2.569620293 | -0.87499721 | 0.001684921 | 0.006363847 |
| LOC119873481 | -2.570267076 | 0.224823573 | 2.32E-05 | 0.000160559 |
| LOC119867496 | -2.57095909 | 1.249541531 | 0.000701824 | 0.003017981 |
| LOC119868656 | -2.571072073 | -0.560184806 | 3.22E-05 | 0.000213859 |
| LOC100688849 | -2.571143915 | 6.318980719 | 0.001700354 | 0.006409005 |
| LOC102154421 | -2.572254061 | -0.02862495 | 7.60E-05 | 0.000451687 |
| CXHXorf65 | -2.574986666 | 2.17871195 | 0.015446031 | 0.041449458 |
| RCVRN | -2.576633283 | -0.760809652 | 0.009237936 | 0.026990265 |
| FGL2 | -2.576833767 | 8.429837222 | 1.52E-06 | 1.49E-05 |
| LOC102153032 | -2.57793647 | 2.065458211 | 1.27E-05 | 9.37E-05 |
| LOC119866612 | -2.58106087 | 4.682398119 | 1.29E-05 | 9.56E-05 |
| LOC119873355 | -2.581107379 | 3.328082305 | 3.20E-06 | 2.83E-05 |
| STYK1 | -2.581120186 | 0.82775039 | 0.006698249 | 0.020542118 |
| ITGA4 | -2.582912761 | 7.252377923 | 0.000214745 | 0.001093813 |
| TP53TG5 | -2.583756531 | 0.78482908 | 0.000108621 | 0.000614657 |
| CD53 | -2.58389984 | 5.63029754 | 0.002583434 | 0.0091563 |
| BCL2A1 | -2.585768988 | 4.535759437 | 5.40E-05 | 0.000335851 |
| ID2 | -2.586187211 | 5.831566568 | 6.25E-08 | 8.97E-07 |
| LYST | -2.586191267 | 8.445566783 | 3.51E-08 | 5.33E-07 |
| CNST | -2.586221403 | 4.695268298 | 2.88E-06 | 2.59E-05 |
| CCR7 | -2.586322624 | 4.698780666 | 0.010047969 | 0.028977995 |

| | | | | |
|---|---|---|---|---|
| CHPT1 | -2.587625498 | 3.773127554 | 1.53E-05 | 0.000110819 |
| LOC106558210 | -2.590965479 | 2.638202822 | 8.85E-05 | 0.00051439 |
| TNFAIP8 | -2.595241692 | 5.419264216 | 7.09E-06 | 5.63E-05 |
| DOCK2 | -2.595304981 | 7.227621963 | 0.000354694 | 0.001680646 |
| SEC14L5 | -2.596257134 | 2.538972271 | 0.010642289 | 0.030386631 |
| LOC119867707 | -2.596311076 | 0.777510003 | 0.000256899 | 0.001271763 |
| CASP4 | -2.596594382 | 4.395136106 | 7.86E-05 | 0.000465446 |
| ICAM3 | -2.596766539 | 4.776411383 | 1.77E-07 | 2.23E-06 |
| KCNMB3 | -2.597186484 | 0.503868483 | 0.001392595 | 0.005417646 |
| GLIPR1 | -2.598761984 | 4.960713825 | 2.53E-05 | 0.00017352 |
| SMIM22 | -2.599339548 | 2.386965927 | 5.10E-06 | 4.22E-05 |
| LOC119864978 | -2.601546215 | 3.903702175 | 1.15E-05 | 8.62E-05 |
| LOC111090272 | -2.602714728 | -0.907119308 | 0.000993876 | 0.004073426 |
| LOC100682919 | -2.603133666 | 1.449885931 | 0.005790618 | 0.018169981 |
| HEPH | -2.60385684 | -0.046116578 | 0.016199083 | 0.0430143 |
| LOC111098258 | -2.605873703 | 3.603042627 | 0.0011978 | 0.004777315 |
| TAC4 | -2.608549272 | -0.345159788 | 0.00025474 | 0.001263093 |
| PCED1B | -2.610995534 | 3.496001156 | 0.004492873 | 0.014614144 |
| LRRC63 | -2.611155155 | 1.251786126 | 0.003722106 | 0.012502 |
| LOC119868346 | -2.612496565 | -1.09429533 | 0.003730296 | 0.012525154 |
| SVOPL | -2.613053961 | 0.694233491 | 0.001865521 | 0.006935669 |
| RAB39A | -2.613388126 | 3.509644144 | 0.001021495 | 0.004166923 |
| LOC119867921 | -2.615231425 | 1.959063992 | 0.001741639 | 0.006536196 |
| LOC119872734 | -2.616305816 | 1.623794648 | 0.000173406 | 0.000908665 |
| LOC102156729 | -2.617647743 | 6.191000927 | 4.13E-10 | 9.92E-09 |
| NOXO1 | -2.619270471 | 0.159206563 | 0.000178156 | 0.000929507 |
| SGPP2 | -2.62219505 | 2.36492264 | 1.73E-05 | 0.000123639 |
| LOC119873613 | -2.626381033 | 2.736881908 | 0.000103655 | 0.000591034 |
| NCF4 | -2.627048234 | 4.465728891 | 6.71E-06 | 5.36E-05 |
| H2BC10 | -2.628920013 | 1.262596205 | 9.27E-05 | 0.000535799 |
| RILPL2 | -2.630541079 | 5.371132263 | 2.49E-10 | 6.21E-09 |
| LOC119872624 | -2.630996279 | -0.213112672 | 0.005254074 | 0.016686922 |
| ZBTB8B | -2.631733296 | 0.008765204 | 0.000432318 | 0.001992264 |
| METTL21C | -2.63207884 | 0.712368189 | 0.000775493 | 0.003289734 |
| LOC119877063 | -2.632731598 | 0.830053751 | 0.000158235 | 0.000840136 |
| LOC119867979 | -2.63277681 | -0.847455152 | 0.004655146 | 0.0150696 |
| CDH4 | -2.633164829 | -0.679180519 | 0.014668118 | 0.039747703 |
| LOC111090739 | -2.63339843 | 2.916496807 | 5.21E-06 | 4.30E-05 |
| LOC119866259 | -2.634597813 | 0.679980926 | 0.000217001 | 0.001102453 |

| | | | | |
|---|---|---|---|---|
| LOC119872096 | -2.636287987 | -1.031455888 | 0.000577083 | 0.002551953 |
| LOC119874080 | -2.639762653 | 0.105533021 | 0.000529467 | 0.002372365 |
| LOC119871889 | -2.641070092 | -0.197777466 | 0.000305211 | 0.001479167 |
| LOC111090712 | -2.641094302 | 3.000684152 | 1.43E-06 | 1.41E-05 |
| SATB1 | -2.642499278 | 6.558506629 | 0.00148407 | 0.00570794 |
| ALOX15 | -2.643511098 | 2.871369472 | 0.001967582 | 0.007270829 |
| GAB3 | -2.643538621 | 1.923984117 | 1.40E-05 | 0.000102643 |
| LOC111096966 | -2.644851528 | -0.325868141 | 0.002121752 | 0.00775067 |
| PDE4C | -2.646489158 | 2.904959852 | 0.000122764 | 0.000681 |
| HHEX | -2.646728027 | 4.486281784 | 1.73E-06 | 1.66E-05 |
| SPI1 | -2.646936438 | 6.753546117 | 0.000324942 | 0.001558758 |
| LOC119869680 | -2.648935649 | 2.993557935 | 0.000171791 | 0.000901509 |
| LOC119869673 | -2.649007333 | -0.924422432 | 0.008817123 | 0.025928562 |
| LOC100686511 | -2.649260392 | 4.003779035 | 0.000203931 | 0.001046354 |
| LOC119874381 | -2.65062137 | 0.3437435 | 0.000104795 | 0.000596591 |
| LOC102151549 | -2.651335724 | -0.04364546 | 0.003623139 | 0.012212176 |
| LOC119871825 | -2.651596053 | 3.240813096 | 6.55E-05 | 0.000396727 |
| LOC119864503 | -2.651631466 | 4.67691802 | 3.78E-07 | 4.39E-06 |
| CPVL | -2.656898486 | -0.385837294 | 0.001438625 | 0.005561776 |
| OSM | -2.65764617 | 2.006823052 | 1.16E-05 | 8.68E-05 |
| LOC106559087 | -2.658512656 | -0.389057165 | 0.013378879 | 0.036792341 |
| TNFRSF17 | -2.661161961 | -0.209094078 | 0.018031448 | 0.046852366 |
| MIR8884 | -2.662520208 | -0.116471536 | 0.000558402 | 0.002479971 |
| RAC2 | -2.664101775 | 5.979007319 | 0.002960225 | 0.010301374 |
| LOC111094367 | -2.6670697 | 3.274015506 | 2.62E-05 | 0.000178658 |
| TRIM22 | -2.668443397 | 7.276699951 | 0.002820653 | 0.009887111 |
| LOC111095527 | -2.669711918 | 2.384290311 | 5.09E-06 | 4.22E-05 |
| LOC111094048 | -2.670135969 | 2.026716868 | 0.00043954 | 0.002021678 |
| STK10 | -2.671156817 | 7.808046091 | 2.11E-09 | 4.32E-08 |
| LOC106559407 | -2.671478906 | 1.473361661 | 0.001599644 | 0.006086568 |
| LOC119864761 | -2.671598505 | 5.2322959 | 3.47E-08 | 5.28E-07 |
| LOC100686047 | -2.673572338 | 2.541681814 | 0.001413449 | 0.005477145 |
| EFNB3 | -2.675473302 | 2.790571618 | 0.001789468 | 0.006695985 |
| LOC111097691 | -2.677409685 | 1.446537145 | 0.002027578 | 0.007458198 |
| P2RY12 | -2.679825754 | 1.634860784 | 0.005584249 | 0.017591197 |
| GALNT6 | -2.680407659 | 4.789448847 | 4.54E-06 | 3.84E-05 |
| DOCK8 | -2.680953201 | 7.660927012 | 1.32E-05 | 9.74E-05 |
| LOC111092452 | -2.681260839 | 0.432250003 | 0.002249004 | 0.0081387 |
| LOC100855449 | -2.686979593 | 4.140799238 | 0.0001379 | 0.000748613 |

| | | | | |
|---|---|---|---|---|
| LOC111097366 | -2.687425723 | -0.505842177 | 0.006288099 | 0.019460513 |
| LOC111097343 | -2.687508168 | 5.079641105 | 8.98E-06 | 6.94E-05 |
| RASAL3 | -2.687743496 | 5.046813301 | 4.22E-05 | 0.000270685 |
| DUXB | -2.691087118 | 0.481555303 | 6.50E-05 | 0.000394215 |
| LOC111093077 | -2.691473383 | -0.433466124 | 0.007855254 | 0.023531017 |
| LOC611536 | -2.692230605 | 2.782986709 | 4.82E-06 | 4.03E-05 |
| RORC | -2.692790182 | 1.146327221 | 0.001688189 | 0.006374239 |
| FNDC9 | -2.694459808 | 2.840278106 | 0.001012249 | 0.004136992 |
| FBXO40 | -2.695061144 | 2.793153036 | 0.000695063 | 0.002994256 |
| LOC102153166 | -2.696459522 | 1.123100106 | 0.000288932 | 0.001409729 |
| LOC119877116 | -2.698129101 | -1.093955098 | 0.007510655 | 0.022667932 |
| LOC119864175 | -2.698566803 | -1.069153119 | 0.000851634 | 0.003566661 |
| CLEC7A | -2.700811272 | 3.056724415 | 6.91E-06 | 5.51E-05 |
| LOC111096643 | -2.701340383 | -0.604667383 | 0.003855257 | 0.012878389 |
| LOC119866550 | -2.702384871 | 1.081008386 | 0.001327443 | 0.005202474 |
| LOC119873755 | -2.702635323 | 3.131419437 | 0.00010185 | 0.000581658 |
| HCLS1 | -2.703399502 | 6.056611931 | 5.73E-07 | 6.31E-06 |
| LOC119874790 | -2.703642219 | 0.472226448 | 0.00297307 | 0.010331144 |
| GBP5 | -2.70466297 | 3.910475761 | 0.001921934 | 0.007116706 |
| LOC609053 | -2.705573915 | 4.282765397 | 0.003136132 | 0.010816262 |
| LOC111096259 | -2.705737292 | 1.365593176 | 0.00011433 | 0.000641137 |
| LOC111097825 | -2.708068507 | 3.003418224 | 0.010238474 | 0.029421103 |
| TRNAY-GUA_9 | -2.715811973 | 0.101119636 | 0.002080132 | 0.007616609 |
| GIMAP7 | -2.716134769 | 4.370891058 | 5.14E-05 | 0.000321874 |
| LOC119867760 | -2.717546529 | -0.571559143 | 6.59E-05 | 0.000398592 |
| KCNA3 | -2.718363474 | 6.469950043 | 0.00163771 | 0.006207432 |
| LOC119867717 | -2.719164592 | 0.643955048 | 0.003170984 | 0.010912154 |
| LOC102156918 | -2.719283317 | 1.172707383 | 0.000299284 | 0.001453366 |
| KLRK1 | -2.721108649 | 1.559957422 | 0.002238288 | 0.008105333 |
| LOC111092288 | -2.721565859 | 2.557116863 | 0.000148822 | 0.000796402 |
| GIMAP4 | -2.7221849 | 5.039263924 | 4.27E-05 | 0.000273425 |
| LOC119873515 | -2.724405341 | 0.567033237 | 9.32E-06 | 7.17E-05 |
| LOC102152134 | -2.726240722 | 2.010074004 | 4.73E-06 | 3.97E-05 |
| LOC119864708 | -2.726474181 | 0.189405607 | 0.001270036 | 0.005020153 |
| LOC119877543 | -2.727042248 | 2.659242964 | 6.24E-07 | 6.78E-06 |
| LOC119873457 | -2.727211645 | 0.035499786 | 0.00043214 | 0.001992128 |
| LOC119871606 | -2.727446031 | 1.743914337 | 6.39E-06 | 5.14E-05 |
| LOC488248 | -2.728987702 | 6.277772923 | 0.008307577 | 0.024650891 |
| LOC119867945 | -2.729584102 | 1.085935153 | 4.13E-05 | 0.000265747 |

| | | | | |
|---|---|---|---|---|
| NIPAL4 | -2.731059947 | 1.37462588 | 0.000833043 | 0.00349962 |
| PLSCR1 | -2.731604003 | 2.331708206 | 0.000230406 | 0.001159144 |
| H2BC7 | -2.733277903 | 1.773491625 | 2.69E-05 | 0.000182638 |
| LOC119869499 | -2.734063785 | 1.978438811 | 0.000249629 | 0.001241592 |
| LOC111091400 | -2.735665032 | -0.816477465 | 0.003346338 | 0.011415897 |
| LOC119872733 | -2.737008384 | 1.609304331 | 0.000841885 | 0.003531971 |
| LOC119874195 | -2.737090065 | 5.194153835 | 4.80E-07 | 5.40E-06 |
| MAP3K19 | -2.737608136 | 0.843589097 | 0.000155287 | 0.000826509 |
| LOC106557635 | -2.737784215 | 3.237664589 | 0.000159661 | 0.000846874 |
| LOC111095540 | -2.740198564 | 5.655661483 | 3.13E-05 | 0.000208696 |
| MARCHF1 | -2.742144754 | 5.862529043 | 7.54E-07 | 8.00E-06 |
| PRSS57 | -2.74301683 | 2.580173216 | 2.85E-05 | 0.000192192 |
| SEMA4D | -2.743756069 | 7.137528605 | 1.98E-06 | 1.86E-05 |
| LOC119877531 | -2.744404858 | 3.427964911 | 1.83E-06 | 1.74E-05 |
| PTPRJ | -2.746386774 | 7.648547234 | 9.88E-08 | 1.34E-06 |
| PLAC8B | -2.746671539 | 4.769984598 | 0.015847675 | 0.042278023 |
| LOC119870696 | -2.748523441 | 0.046888172 | 0.009093294 | 0.026632292 |
| LOC102154724 | -2.749151785 | 3.238684344 | 7.45E-08 | 1.04E-06 |
| CD52 | -2.749756024 | -0.188764177 | 0.000652073 | 0.002837827 |
| LOC119869506 | -2.750069373 | 2.685044155 | 0.001230995 | 0.004886466 |
| CD28 | -2.750393842 | 2.559826716 | 0.00731646 | 0.022165171 |
| CERKL | -2.750545578 | 3.884234051 | 0.000172675 | 0.000905487 |
| LOC119876150 | -2.752693034 | 4.281430882 | 0.000336387 | 0.001604074 |
| LOC111095986 | -2.753716367 | -0.487230773 | 0.001327232 | 0.005202474 |
| RFESD | -2.754267545 | 1.134151553 | 9.30E-05 | 0.000537093 |
| RASGRP4 | -2.756839963 | 5.722247299 | 4.16E-11 | 1.23E-09 |
| LOC119872728 | -2.757077442 | 1.869281714 | 2.91E-05 | 0.000196049 |
| DPYS | -2.760438742 | 1.379618658 | 0.008388042 | 0.024859024 |
| NUP210 | -2.762093935 | 7.132543664 | 2.03E-06 | 1.91E-05 |
| LOC111097780 | -2.7628594 | 2.703151365 | 0.000756511 | 0.003217484 |
| CMA1 | -2.764590934 | 1.770965536 | 0.009702661 | 0.028101801 |
| LOC119864626 | -2.76621434 | 0.568354054 | 0.002161084 | 0.007871793 |
| NAMPT | -2.766590727 | 7.540084633 | 4.69E-09 | 8.69E-08 |
| LOC119873281 | -2.766877275 | 1.377173062 | 0.012791745 | 0.035447576 |
| LOC111098259 | -2.766931327 | 2.216834009 | 0.004185239 | 0.013781607 |
| LYN | -2.76861714 | 7.047865581 | 7.43E-07 | 7.90E-06 |
| LOC119869713 | -2.77435371 | 2.203005369 | 7.97E-05 | 0.00047063 |
| LOC111090595 | -2.775646935 | 2.639472632 | 2.99E-06 | 2.68E-05 |
| EFHC2 | -2.77640737 | 2.155563085 | 1.71E-05 | 0.00012252 |

| | | | | |
|---|---|---|---|---|
| LOC119871929 | -2.777333806 | 0.742830975 | 1.86E-05 | 0.000132147 |
| LOC102157140 | -2.77835659 | -0.287870325 | 0.001701349 | 0.006409409 |
| FAM241A | -2.778401429 | 5.628974898 | 9.63E-05 | 0.000552941 |
| CD244 | -2.779401952 | 3.284645844 | 0.001213522 | 0.004832017 |
| LOC111094789 | -2.781070532 | -0.065256485 | 0.001444026 | 0.005579672 |
| STK17A | -2.782124299 | 6.333053195 | 2.48E-07 | 3.03E-06 |
| EVI2B | -2.782923912 | 6.433392727 | 1.14E-08 | 1.94E-07 |
| ITGB2 | -2.782934862 | 6.689524065 | 1.72E-05 | 0.000123451 |
| LOC111098949 | -2.783320883 | 0.395116953 | 0.001874787 | 0.00695877 |
| LOC119871510 | -2.783856259 | 0.846629222 | 0.006029564 | 0.018790786 |
| KCNJ2 | -2.784265577 | 3.460093929 | 7.50E-06 | 5.92E-05 |
| LOC119866364 | -2.788079491 | 2.212088817 | 0.000428268 | 0.001977806 |
| LOC119870695 | -2.79267105 | 0.207034561 | 0.007369562 | 0.022307334 |
| LOC119872243 | -2.79377058 | 3.377294623 | 1.83E-06 | 1.75E-05 |
| PGLYRP2 | -2.79628682 | 0.206949335 | 0.000754242 | 0.003210268 |
| LOC119865960 | -2.797115942 | 2.18498375 | 0.001566404 | 0.005983198 |
| FADS1 | -2.803897231 | 4.360348289 | 1.55E-05 | 0.000112474 |
| LY96 | -2.803994299 | 2.89365702 | 2.26E-06 | 2.09E-05 |
| PIK3R5 | -2.80517818 | 6.647899812 | 2.57E-06 | 2.35E-05 |
| FGF9 | -2.808300189 | 1.875179333 | 0.005092948 | 0.016260969 |
| LOC119863944 | -2.81021391 | 0.703741643 | 0.000263465 | 0.00129922 |
| ABRACL | -2.810426395 | 4.380631678 | 9.36E-07 | 9.66E-06 |
| C6H1orf194 | -2.810509471 | 1.658762306 | 2.18E-06 | 2.03E-05 |
| LOC119877356 | -2.813875926 | 2.868275991 | 0.000354262 | 0.001680069 |
| GPR15 | -2.814990004 | 1.70101867 | 4.55E-06 | 3.84E-05 |
| LOC119866916 | -2.820066979 | 1.206305348 | 0.001286139 | 0.005072699 |
| CD3G | -2.820341604 | 4.535906991 | 0.01146294 | 0.032323224 |
| IL2RG | -2.82122741 | 4.793425184 | 0.001514722 | 0.005814483 |
| KCNE3 | -2.821558418 | 4.948262512 | 6.90E-08 | 9.77E-07 |
| SNX10 | -2.822372029 | 5.39435424 | 1.37E-05 | 0.000100691 |
| NME9 | -2.822456037 | 0.545679399 | 0.000134276 | 0.000732348 |
| LOC111091793 | -2.823542715 | 0.582125681 | 0.000626498 | 0.002740283 |
| ZDHHC18 | -2.827180609 | 6.361581074 | 1.07E-10 | 2.89E-09 |
| GPR171 | -2.832794868 | 2.627101968 | 0.002340485 | 0.008402406 |
| LOC102156504 | -2.83497526 | 0.276370943 | 2.58E-05 | 0.0001764 |
| CD40LG | -2.835934098 | 2.975156892 | 0.000123197 | 0.000682351 |
| LOC119866299 | -2.838118624 | 3.844470772 | 2.25E-05 | 0.000156316 |
| LOC119865177 | -2.838454633 | 1.833552448 | 0.001233628 | 0.004893195 |
| IKZF1 | -2.842737538 | 7.014012432 | 1.85E-05 | 0.000131683 |

| | | | | |
|---|---|---|---|---|
| LOC119866110 | -2.845934706 | 2.944524272 | 0.016139028 | 0.04289162 |
| LOC487977 | -2.846069078 | 2.2643414 | 6.76E-06 | 5.40E-05 |
| SIRPB2 | -2.848186449 | 4.533005794 | 0.000363031 | 0.001714903 |
| LOC119866562 | -2.849251846 | 1.101225339 | 0.000796648 | 0.00336171 |
| LOC111094049 | -2.849367922 | 0.268729212 | 0.003013704 | 0.010449628 |
| LOC100684842 | -2.850754215 | 1.081824449 | 4.03E-05 | 0.000260018 |
| CDA | -2.857578992 | 4.08463068 | 9.30E-05 | 0.000537093 |
| RHOG | -2.863722987 | 6.534235019 | 1.54E-11 | 5.01E-10 |
| LOC102153493 | -2.864950635 | 2.048535276 | 0.01578618 | 0.04215548 |
| MYO1F | -2.865229323 | 6.930878915 | 1.16E-07 | 1.54E-06 |
| LOC111095548 | -2.867637178 | 3.012342334 | 1.64E-08 | 2.68E-07 |
| LOC111090235 | -2.867983214 | 0.478498659 | 0.000103557 | 0.000590888 |
| LOC111097365 | -2.869218019 | 0.666923514 | 0.002852678 | 0.009989675 |
| CD69 | -2.870909829 | 3.428139318 | 0.004993337 | 0.015982955 |
| LOC111097616 | -2.873266408 | 0.615338039 | 0.009045154 | 0.026505628 |
| CORO1A | -2.873785493 | 6.948658845 | 5.05E-05 | 0.000317147 |
| LOC119872724 | -2.877845518 | 0.414705456 | 0.000541855 | 0.002417876 |
| LOC102152043 | -2.878021071 | -0.07873838 | 1.66E-05 | 0.00011922 |
| GPAT3 | -2.878257311 | 4.455297189 | 5.25E-06 | 4.33E-05 |
| LOC119871709 | -2.885736509 | -0.108007037 | 0.000385797 | 0.001807086 |
| TXK | -2.887239864 | 3.91263298 | 0.000436299 | 0.002008905 |
| LOC119871107 | -2.888177362 | 2.37189582 | 4.17E-06 | 3.55E-05 |
| LOC119869554 | -2.888571199 | 1.921263789 | 1.04E-06 | 1.06E-05 |
| TNFSF9 | -2.892189681 | 2.822138492 | 1.96E-07 | 2.45E-06 |
| LOC119876361 | -2.89896549 | 1.135407485 | 0.00035508 | 0.001682111 |
| LOC119869906 | -2.903060952 | 0.90488858 | 9.10E-05 | 0.000527116 |
| SOS2 | -2.903792379 | 6.638758211 | 3.96E-12 | 1.49E-10 |
| NT5C3A | -2.904307422 | 4.150223226 | 1.69E-08 | 2.75E-07 |
| EPB42 | -2.905764989 | 2.384139917 | 7.35E-07 | 7.83E-06 |
| TRNAK-CUU_29 | -2.907753582 | 0.479707216 | 0.000455686 | 0.002081807 |
| LOC106558747 | -2.907924111 | -0.252014555 | 1.96E-05 | 0.000138342 |
| ADAM8 | -2.908103869 | 5.293381256 | 3.31E-10 | 8.07E-09 |
| LOC111091314 | -2.910722518 | 0.910439712 | 1.91E-06 | 1.81E-05 |
| PAG1 | -2.913168574 | 7.440483053 | 5.85E-06 | 4.76E-05 |
| SLAMF9 | -2.914432477 | 3.11236129 | 0.000152726 | 0.000814275 |
| LOC119865427 | -2.916599372 | 0.410557161 | 0.000372365 | 0.001752501 |
| LOC119873172 | -2.917324227 | -0.383875741 | 0.004313055 | 0.014111841 |
| LOC111094000 | -2.922649958 | 0.733474074 | 4.47E-05 | 0.000284637 |
| SLC2A9 | -2.927056954 | 2.497527943 | 5.62E-05 | 0.000347473 |

| | | | | |
|---|---|---|---|---|
| CD48 | -2.92722711 | 5.079518695 | 2.88E-05 | 0.000194284 |
| VSIR | -2.928222256 | 8.593818643 | 7.87E-13 | 3.43E-11 |
| LOC106558146 | -2.929390055 | 0.654095967 | 0.000118364 | 0.000660989 |
| SLC22A11 | -2.929672616 | -0.601374691 | 0.001388307 | 0.005404591 |
| LOC119877532 | -2.929833185 | 3.491239329 | 0.000212817 | 0.001084759 |
| LOC111095547 | -2.932031091 | 4.187800114 | 1.12E-05 | 8.42E-05 |
| LOC102154670 | -2.933824645 | 3.3884239 | 3.77E-05 | 0.00024546 |
| LOC119865482 | -2.934609499 | -0.884593101 | 0.000620559 | 0.002717045 |
| RGS7BP | -2.935657299 | 2.462961262 | 2.38E-05 | 0.000164379 |
| TSPAN13 | -2.937809837 | 5.254524302 | 5.30E-05 | 0.000330591 |
| LOC106559661 | -2.938914255 | 0.689138015 | 5.45E-05 | 0.000338316 |
| LOC102156986 | -2.940380648 | 3.465226807 | 1.40E-05 | 0.000102819 |
| PMFBP1 | -2.941437001 | -0.910166305 | 0.003491994 | 0.011845668 |
| LOC119864892 | -2.941965116 | 1.590770685 | 0.000900833 | 0.003748082 |
| LOC106559529 | -2.943820013 | -0.226691324 | 0.000696818 | 0.003000028 |
| LOC119866882 | -2.943832439 | 2.563391489 | 0.000363333 | 0.001715955 |
| SLC24A1 | -2.944155929 | 2.382669347 | 2.42E-05 | 0.000166872 |
| RCSD1 | -2.946972667 | 6.57804542 | 2.09E-08 | 3.33E-07 |
| MGST1 | -2.947607985 | 4.557071045 | 6.13E-06 | 4.96E-05 |
| LOC111098550 | -2.947916879 | 0.596415234 | 2.16E-05 | 0.00015074 |
| LOC102151269 | -2.949564417 | 0.809414045 | 4.28E-06 | 3.64E-05 |
| LOC102151902 | -2.950032602 | 1.811226783 | 0.001805905 | 0.006752355 |
| LOC106559121 | -2.951533613 | 1.369788235 | 0.011007891 | 0.031243324 |
| LCP1 | -2.952314835 | 8.78613585 | 6.04E-05 | 0.000370178 |
| LOC490269 | -2.953612705 | 6.612995812 | 7.39E-05 | 0.000441321 |
| LOC111097055 | -2.955446338 | 1.485747572 | 0.000551649 | 0.002453999 |
| LOC111093692 | -2.95558811 | 3.705908437 | 9.69E-05 | 0.000555917 |
| LOC111097673 | -2.961409382 | 0.437344524 | 0.00342738 | 0.011644894 |
| CLNK | -2.962834107 | 3.146980109 | 4.25E-06 | 3.62E-05 |
| LOC119873213 | -2.965091 | 1.29748674 | 0.00018981 | 0.000980865 |
| LOC119875832 | -2.969075057 | -1.488805098 | 0.007174998 | 0.021785338 |
| LOC102157377 | -2.970300219 | 3.329822059 | 4.31E-07 | 4.92E-06 |
| LOC111093385 | -2.971294332 | 0.778491586 | 0.000166845 | 0.000878976 |
| GALNT3 | -2.972672396 | 3.878446769 | 0.000344938 | 0.001640294 |
| LOC119871546 | -2.973633359 | 2.23608834 | 0.019301019 | 0.049557041 |
| HCAR2 | -2.973818966 | -0.155447705 | 0.005822262 | 0.01825342 |
| ABCA13 | -2.976263221 | 0.416677536 | 1.66E-05 | 0.000119538 |
| CELA1 | -2.977804695 | 0.204014965 | 0.000762927 | 0.003240872 |
| DDX39A | -2.978167804 | 5.882206232 | 3.83E-13 | 1.77E-11 |

| | | | | |
|---|---|---|---|---|
| LOC119866230 | -2.981564113 | 2.952116071 | 2.57E-07 | 3.12E-06 |
| CCR4 | -2.982461327 | 1.395387692 | 0.001046127 | 0.004253255 |
| LOC100683650 | -2.982845479 | 3.749824925 | 1.60E-08 | 2.62E-07 |
| ZP1 | -2.983886448 | 0.665707844 | 0.000778994 | 0.003301482 |
| SPN | -2.983934093 | 5.870167506 | 0.000721808 | 0.003089818 |
| LOC119867692 | -2.984979408 | 2.06655614 | 0.000806668 | 0.003399362 |
| LRRC75B | -2.985100413 | 2.851252151 | 3.24E-07 | 3.83E-06 |
| STPG4 | -2.986364498 | 0.372990149 | 2.95E-06 | 2.64E-05 |
| CEACAM1 | -2.988769403 | 5.538063262 | 4.38E-12 | 1.64E-10 |
| DOK6 | -2.989366697 | 2.533136625 | 0.002643416 | 0.009344437 |
| GNLY | -2.99025041 | 2.112877858 | 0.005292337 | 0.016786304 |
| LOC111091149 | -2.9925312 | 0.067583359 | 0.000658927 | 0.002862481 |
| LOC119876722 | -2.992644676 | -0.290145031 | 3.76E-05 | 0.000244699 |
| LOC119872741 | -2.993152881 | 1.38871025 | 8.25E-06 | 6.44E-05 |
| GNG11 | -2.993182516 | 5.266822858 | 0.000539254 | 0.002408253 |
| SELPLG | -2.994224331 | 5.733776997 | 3.24E-05 | 0.000215149 |
| TMEM154 | -2.995732858 | 7.250237224 | 1.07E-08 | 1.83E-07 |
| LOC119868447 | -2.996802171 | 0.061384864 | 0.000580635 | 0.002564692 |
| CYBB | -2.998478669 | 6.551727493 | 2.04E-05 | 0.000143517 |
| THEMIS2 | -3.001673754 | 5.256276987 | 2.06E-10 | 5.27E-09 |
| LOC119867034 | -3.004136785 | 2.367617285 | 5.80E-08 | 8.35E-07 |
| LOC119871693 | -3.006019532 | -0.553642024 | 0.003702583 | 0.012445111 |
| LOC119874695 | -3.007493835 | 1.423060063 | 0.006423623 | 0.019837442 |
| TREML2 | -3.008754994 | 3.617498988 | 0.005696053 | 0.017909553 |
| AK3 | -3.009900073 | 5.420893573 | 1.24E-11 | 4.12E-10 |
| GK | -3.0104213 | 4.812760679 | 1.36E-10 | 3.62E-09 |
| LAPTM5 | -3.010714999 | 8.351477139 | 1.51E-06 | 1.47E-05 |
| P2RY1 | -3.011657723 | 4.13540882 | 5.24E-05 | 0.000327292 |
| HTR4 | -3.01392673 | 0.248764546 | 0.002804477 | 0.00983996 |
| LOC119867103 | -3.014419404 | -0.193980523 | 0.000131141 | 0.000718093 |
| LOC102151412 | -3.018018832 | 0.357213095 | 0.003762559 | 0.0126154 |
| JAML | -3.018610754 | 4.73277129 | 1.47E-05 | 0.000107503 |
| IRF4 | -3.019826117 | 4.770222445 | 6.45E-06 | 5.18E-05 |
| CCND3 | -3.020750132 | 6.015473644 | 4.72E-10 | 1.11E-08 |
| TNFSF8 | -3.023056639 | 3.888971614 | 1.00E-05 | 7.62E-05 |
| LOC111090221 | -3.024676009 | -0.166146222 | 0.002293412 | 0.008267652 |
| LOC111096416 | -3.030898856 | 1.941266441 | 6.16E-07 | 6.72E-06 |
| LOC119867915 | -3.031029377 | 1.388761633 | 1.81E-05 | 0.000129273 |
| LOC119866145 | -3.031214321 | 0.48269011 | 5.72E-06 | 4.68E-05 |

| | | | | |
|---|---|---|---|---|
| LOC119864705 | -3.031966182 | 1.847395542 | 1.24E-05 | 9.24E-05 |
| PTGER2 | -3.033428147 | 1.612112227 | 6.43E-05 | 0.000390356 |
| ENKUR | -3.034363559 | 0.840326629 | 0.001194515 | 0.004768194 |
| LOC119877309 | -3.03490567 | 4.003678846 | 9.76E-09 | 1.69E-07 |
| DUSP26 | -3.035145051 | 4.859473184 | 3.97E-06 | 3.41E-05 |
| ADGRE5 | -3.035987779 | 7.540109379 | 1.20E-09 | 2.59E-08 |
| LOC119865167 | -3.037400133 | 2.029010071 | 3.09E-05 | 0.00020698 |
| HMGA2 | -3.037596575 | 1.0585515 | 0.00012304 | 0.000681831 |
| LOC119874702 | -3.040048897 | -0.680040379 | 0.005004307 | 0.016008611 |
| PGLYRP1 | -3.040671115 | 1.732798902 | 2.95E-06 | 2.64E-05 |
| LOC111095895 | -3.041426139 | 0.19474019 | 4.13E-05 | 0.000265419 |
| PADI6 | -3.041918729 | 0.235825874 | 0.000130441 | 0.000715344 |
| FCER1G | -3.042496209 | 6.184516749 | 7.06E-05 | 0.000423404 |
| LOC119869604 | -3.042562841 | 0.863786257 | 0.001746512 | 0.006551081 |
| WDR17 | -3.043103391 | 1.234889744 | 0.000660984 | 0.002869693 |
| RAB25 | -3.046250873 | 0.716015488 | 1.42E-05 | 0.000103811 |
| LOC119865498 | -3.04805207 | -0.562633933 | 3.24E-05 | 0.000215149 |
| MPZL3 | -3.051484653 | 2.805368633 | 5.79E-05 | 0.000356726 |
| SLC25A53 | -3.052324974 | 0.203514209 | 0.00086536 | 0.003617155 |
| MIR590 | -3.053786745 | 0.432488179 | 0.000166923 | 0.000879155 |
| CD3E | -3.054218744 | 5.660597007 | 0.001285821 | 0.005072369 |
| CCR2 | -3.055178815 | 5.226831516 | 3.46E-05 | 0.000227911 |
| LOC482936 | -3.056118178 | 1.255790747 | 0.003854656 | 0.012878366 |
| S100P | -3.057821357 | 2.685250656 | 2.11E-07 | 2.61E-06 |
| PRKCB | -3.057989477 | 7.056860934 | 5.57E-05 | 0.00034491 |
| LOC119872742 | -3.062893525 | 0.888019246 | 3.96E-05 | 0.000255966 |
| LOC119865374 | -3.063079272 | -0.051166478 | 0.000335876 | 0.001601988 |
| LOC119876517 | -3.063673455 | 2.139942727 | 0.008437498 | 0.024985091 |
| LOC111090263 | -3.063858265 | 0.19848643 | 1.77E-06 | 1.70E-05 |
| SVIP | -3.064752554 | 5.230300312 | 8.15E-08 | 1.13E-06 |
| IL1RAP | -3.068165204 | 5.228595623 | 3.73E-11 | 1.12E-09 |
| LY86 | -3.071012507 | 3.106842441 | 2.79E-06 | 2.52E-05 |
| TENM1 | -3.073489918 | 4.847301845 | 0.000417166 | 0.001936027 |
| IL2RA | -3.074939601 | 2.824378391 | 4.65E-06 | 3.91E-05 |
| LOC111093720 | -3.077553341 | 1.928555558 | 1.19E-06 | 1.20E-05 |
| LOC111091084 | -3.081737676 | 1.820704159 | 0.000111733 | 0.000629802 |
| LOC119870755 | -3.081768555 | -0.279207233 | 0.000106774 | 0.000605947 |
| LOC111090605 | -3.083253264 | 0.668476935 | 3.45E-06 | 3.02E-05 |
| LOC106559086 | -3.087442816 | -0.988704279 | 0.003199311 | 0.010990436 |

| | | | | |
|---|---|---|---|---|
| LOC102153375 | -3.088187007 | 0.455450146 | 0.000243787 | 0.001216015 |
| QRICH2 | -3.090685831 | 3.215017751 | 1.33E-07 | 1.75E-06 |
| MYO15A | -3.093907245 | 0.491998912 | 0.000117732 | 0.000657911 |
| LOC119877468 | -3.097292965 | 1.891069883 | 0.000159642 | 0.000846874 |
| TLR8 | -3.101758495 | 4.810149024 | 3.36E-06 | 2.96E-05 |
| LOC119871074 | -3.103130875 | 2.698227703 | 0.001220738 | 0.004853615 |
| LOC111093640 | -3.103717223 | 1.814145813 | 8.96E-09 | 1.56E-07 |
| ABTB1 | -3.104952502 | 6.322730027 | 9.73E-13 | 4.17E-11 |
| LOC119874168 | -3.105450768 | -0.460865569 | 0.00025828 | 0.001276557 |
| LOC119864961 | -3.110294324 | 1.794736173 | 1.47E-05 | 0.000107273 |
| LOC102157349 | -3.111765142 | 3.884373991 | 1.83E-07 | 2.30E-06 |
| RAP2A | -3.112562523 | 7.167817509 | 6.36E-09 | 1.15E-07 |
| SPTA1 | -3.112919907 | -0.725190019 | 0.003187454 | 0.010963609 |
| LOC119877171 | -3.119002355 | -0.31417061 | 0.00962674 | 0.027930434 |
| LOC106558739 | -3.119711063 | 3.723601311 | 0.000684667 | 0.002956849 |
| DNAJB13 | -3.121083598 | 4.801019921 | 6.32E-05 | 0.000384656 |
| F2RL3 | -3.132512034 | 1.610817248 | 7.65E-05 | 0.000454563 |
| ALAS2 | -3.139895088 | 2.217542204 | 2.54E-05 | 0.000173857 |
| C17H1orf162 | -3.141001591 | 2.608880629 | 7.19E-06 | 5.69E-05 |
| KCNS1 | -3.141313508 | 0.90573986 | 3.17E-06 | 2.82E-05 |
| FCRL2 | -3.142279756 | -0.06264819 | 0.008666818 | 0.025562866 |
| THEMIS | -3.142961546 | 3.283564691 | 0.001410793 | 0.005468809 |
| LOC100686622 | -3.146719666 | 1.34994037 | 8.15E-05 | 0.000479732 |
| GGH | -3.148708424 | 6.887901701 | 1.27E-08 | 2.12E-07 |
| EMR4 | -3.150258193 | 2.295846464 | 0.000509313 | 0.002293917 |
| NOL4 | -3.151401153 | 2.462813404 | 0.00435326 | 0.014228354 |
| TLR2 | -3.153909867 | 5.252712429 | 5.90E-10 | 1.37E-08 |
| LOC102157333 | -3.154197034 | 0.01822407 | 1.48E-05 | 0.000107775 |
| LOC119876385 | -3.15699815 | 2.073634963 | 0.000112094 | 0.000631345 |
| MMP25 | -3.161142689 | 5.140647265 | 6.32E-05 | 0.000384478 |
| LOC102151946 | -3.162957516 | 0.372725609 | 0.001289572 | 0.005084329 |
| LOC111090070 | -3.165671903 | -1.113171306 | 0.003931929 | 0.01309817 |
| LOC119866165 | -3.168108796 | 1.616644393 | 7.27E-05 | 0.000434718 |
| TUBA4A | -3.172493009 | 4.63962577 | 1.01E-10 | 2.74E-09 |
| LOC111097592 | -3.173357332 | 2.142259136 | 2.23E-06 | 2.07E-05 |
| CFP | -3.177876145 | 4.081396423 | 5.03E-08 | 7.39E-07 |
| OR4S7 | -3.178876107 | 0.584109655 | 2.41E-05 | 0.00016629 |
| DNAH8 | -3.17901902 | 0.502001097 | 0.001230038 | 0.004885216 |
| NFAM1 | -3.183793005 | 5.894020282 | 2.15E-06 | 2.00E-05 |

| | | | | |
|---|---|---|---|---|
| LOC119868426 | -3.183806905 | -0.571529611 | 0.0016236 | 0.006162571 |
| LOC102156807 | -3.188636139 | 0.996383526 | 0.000776254 | 0.003292319 |
| LOC119869117 | -3.189400681 | 0.176443361 | 0.018464641 | 0.047760374 |
| LOC119873853 | -3.195545233 | 1.408011783 | 3.69E-06 | 3.20E-05 |
| SLA | -3.195776456 | 4.553676366 | 2.23E-05 | 0.000155201 |
| ALPK2 | -3.1958946 | 3.463714599 | 0.000246711 | 0.001228905 |
| LEF1 | -3.196027485 | 6.298172555 | 3.54E-06 | 3.09E-05 |
| LOC119877931 | -3.196908226 | 2.7719575 | 3.11E-05 | 0.000207571 |
| LOC111095595 | -3.198420783 | -0.025508048 | 0.001104988 | 0.004461328 |
| LOC111091085 | -3.201131027 | 1.694243461 | 5.80E-05 | 0.000357514 |
| RASSF5 | -3.201858361 | 6.733657428 | 2.61E-07 | 3.16E-06 |
| LOC100687917 | -3.202780168 | 0.384622623 | 0.000384107 | 0.001800723 |
| NOD2 | -3.21043307 | 4.698007312 | 2.53E-09 | 5.03E-08 |
| SYK | -3.21140718 | 6.691990958 | 5.48E-07 | 6.07E-06 |
| RSPH1 | -3.212538334 | 0.692345909 | 0.002946617 | 0.010262259 |
| LOC111097534 | -3.213616065 | -0.816902602 | 0.000223985 | 0.001131302 |
| RAB44 | -3.213954958 | 0.576045889 | 0.000308783 | 0.001494141 |
| CCIN | -3.214353089 | 4.237740537 | 6.19E-07 | 6.75E-06 |
| SEL1L3 | -3.21438108 | 3.199722244 | 2.25E-05 | 0.000156219 |
| KLRD1 | -3.21925731 | 2.365143966 | 0.001303695 | 0.005128876 |
| LOC119876202 | -3.22016631 | 3.806049118 | 0.014587046 | 0.039567591 |
| LOC102154812 | -3.224574836 | 1.306916985 | 0.000862309 | 0.003605098 |
| FYB1 | -3.226181267 | 7.466673672 | 2.11E-06 | 1.97E-05 |
| LOC111093311 | -3.227833829 | 0.597687745 | 0.000277425 | 0.001357867 |
| NCR1 | -3.231147909 | 1.014187117 | 0.001370578 | 0.005351204 |
| HCK | -3.232977437 | 5.231596171 | 7.68E-08 | 1.07E-06 |
| LOC106559831 | -3.233397005 | -0.558723048 | 4.35E-06 | 3.69E-05 |
| LOC102155441 | -3.233779971 | -0.334642036 | 0.006085554 | 0.018931406 |
| LOC111091150 | -3.235747981 | 1.377315402 | 7.80E-05 | 0.000461998 |
| LOC491494 | -3.236921335 | 2.091632882 | 0.017825251 | 0.046439074 |
| LOC111093112 | -3.23722135 | 2.774415309 | 1.93E-06 | 1.83E-05 |
| LOC102155503 | -3.239240799 | 0.978605768 | 0.000964184 | 0.003968994 |
| LOC111093878 | -3.239928783 | -0.397314652 | 0.000103602 | 0.000590888 |
| DIRAS2 | -3.243035153 | 1.753581725 | 0.00406082 | 0.013434612 |
| PLEK | -3.244160086 | 5.902392825 | 4.33E-08 | 6.42E-07 |
| MIR29C-2 | -3.248255778 | -0.784003235 | 0.000364302 | 0.001719783 |
| LOC484306 | -3.248292275 | 5.385544026 | 1.65E-06 | 1.59E-05 |
| NCR3 | -3.249162053 | 1.132053432 | 4.60E-05 | 0.000292182 |
| LOC119871847 | -3.253449638 | 3.548818212 | 8.81E-08 | 1.21E-06 |

| | | | | |
|---|---|---|---|---|
| LOC111095190 | -3.260181761 | 2.759243189 | 6.10E-07 | 6.65E-06 |
| LOC119875827 | -3.263558178 | -0.728152648 | 0.002251574 | 0.008145278 |
| KLB | -3.265059296 | 3.178733068 | 6.63E-07 | 7.14E-06 |
| LOC102152520 | -3.266860607 | 1.471378525 | 3.25E-05 | 0.000215595 |
| HTR2A | -3.270682129 | 0.616900293 | 0.003145236 | 0.01084076 |
| LOC119867696 | -3.272363515 | 2.165592009 | 2.12E-05 | 0.000148419 |
| LOC611083 | -3.274643159 | 1.388981553 | 3.10E-06 | 2.76E-05 |
| IL2RB | -3.275403153 | 5.54674461 | 3.24E-05 | 0.000215149 |
| LOC119877587 | -3.278458906 | 0.618126732 | 0.000814258 | 0.003428015 |
| LOC102153575 | -3.279261219 | 1.619039355 | 5.81E-07 | 6.38E-06 |
| IL12RB2 | -3.286076712 | 4.418714553 | 1.25E-10 | 3.33E-09 |
| LOC119871549 | -3.286143823 | 3.51083143 | 0.009128831 | 0.026718319 |
| LOC119871121 | -3.287800546 | -0.56789521 | 1.22E-05 | 9.07E-05 |
| GABRQ | -3.287839462 | 1.854920208 | 0.004762019 | 0.015342239 |
| ICOS | -3.288309353 | 3.105010458 | 0.001242547 | 0.004922267 |
| TLR4 | -3.29171916 | 5.863399615 | 4.10E-10 | 9.86E-09 |
| DNTT | -3.293027172 | -1.255180663 | 0.002207296 | 0.008017215 |
| LOC102152131 | -3.294521384 | 0.973040402 | 0.000167901 | 0.000884092 |
| LOC119871560 | -3.298838202 | 3.60573703 | 3.26E-09 | 6.30E-08 |
| LOC102153240 | -3.29972015 | -0.924061843 | 0.001490672 | 0.005730284 |
| LTB4R | -3.301312382 | 2.811201217 | 8.02E-09 | 1.42E-07 |
| CCRL2 | -3.3140952 | 0.634595371 | 1.00E-05 | 7.63E-05 |
| CCDC96 | -3.323463478 | 0.812574771 | 1.90E-06 | 1.80E-05 |
| CASS4 | -3.324330686 | 3.776022849 | 8.83E-09 | 1.54E-07 |
| PTGIR | -3.326604671 | 3.372879878 | 7.48E-05 | 0.000446203 |
| SLC10A1 | -3.332677559 | 3.023625036 | 8.93E-08 | 1.23E-06 |
| PLXNC1 | -3.335678916 | 7.296965653 | 3.41E-12 | 1.31E-10 |
| LOC119873784 | -3.336646842 | 2.307419287 | 9.80E-05 | 0.000562203 |
| TAGAP | -3.345611169 | 5.482715229 | 7.09E-09 | 1.27E-07 |
| LOC111093358 | -3.350067536 | 1.106085488 | 1.28E-07 | 1.69E-06 |
| IRF6 | -3.351679276 | 0.733551971 | 6.19E-05 | 0.000378192 |
| IPCEF1 | -3.352987965 | 5.997191111 | 2.06E-09 | 4.20E-08 |
| LOC119873634 | -3.355932936 | 3.951924468 | 1.19E-05 | 8.89E-05 |
| VSNL1 | -3.372686198 | -0.725699507 | 7.70E-05 | 0.000456718 |
| LOC111090900 | -3.376245638 | 2.029559599 | 3.63E-07 | 4.22E-06 |
| HK3 | -3.376507098 | 5.64928973 | 6.57E-14 | 3.47E-12 |
| LOC102157336 | -3.376964569 | 0.430119098 | 3.94E-06 | 3.39E-05 |
| LOC111091865 | -3.387112464 | -0.500764488 | 0.00010293 | 0.000587516 |
| MXD1 | -3.388510925 | 4.319320806 | 2.75E-10 | 6.78E-09 |

| | | | | |
|---|---|---|---|---|
| LOC119877648 | -3.392290133 | 0.836623716 | 3.07E-05 | 0.000205457 |
| LOC607096 | -3.393023948 | 2.679815178 | 3.79E-05 | 0.000246543 |
| LOC119865009 | -3.395279505 | -0.057737635 | 2.55E-05 | 0.000174998 |
| LOC119872743 | -3.40036053 | 1.496493417 | 3.86E-06 | 3.33E-05 |
| NRG1 | -3.402622993 | 4.488535409 | 0.000895347 | 0.003729551 |
| SOD2 | -3.403755633 | 7.999306785 | 1.92E-11 | 6.15E-10 |
| XDH | -3.409860184 | 6.66746789 | 2.52E-09 | 5.01E-08 |
| LOC119867246 | -3.412669864 | 0.134199749 | 2.84E-06 | 2.56E-05 |
| LRRK2 | -3.414890803 | 8.397717242 | 4.66E-08 | 6.88E-07 |
| LOC119871614 | -3.414918764 | -1.301754042 | 0.000564976 | 0.002505059 |
| LOC119872095 | -3.41710495 | 1.594043821 | 1.66E-05 | 0.000119388 |
| RNF175 | -3.417719662 | 0.389122467 | 4.09E-05 | 0.000263132 |
| LOC111097029 | -3.418752987 | 3.337356995 | 1.72E-08 | 2.81E-07 |
| TNFSF14 | -3.427834676 | 0.905876369 | 0.000454668 | 0.002078029 |
| DNAH14 | -3.433016498 | 1.535092937 | 5.08E-05 | 0.000318607 |
| LOC119871392 | -3.437411554 | -0.138699116 | 0.000103938 | 0.000592332 |
| PTPRC | -3.441472125 | 8.690408204 | 0.000111534 | 0.000628843 |
| ANKRD63 | -3.44272579 | 0.005341495 | 3.43E-06 | 3.01E-05 |
| LOC119876595 | -3.444987624 | -0.623410205 | 3.48E-05 | 0.000228937 |
| CNTD1 | -3.449708343 | 1.422325476 | 1.11E-07 | 1.48E-06 |
| TYROBP | -3.45304093 | 5.546057976 | 1.98E-09 | 4.06E-08 |
| LOC111091812 | -3.453596544 | 1.63726897 | 6.24E-05 | 0.000380636 |
| SCML4 | -3.456026824 | 5.164977825 | 0.000183603 | 0.000954022 |
| LOC119873061 | -3.460462177 | 1.288185901 | 4.97E-05 | 0.000312512 |
| ZNF385B | -3.463200157 | 1.020443846 | 0.00222415 | 0.008065717 |
| LOC111092086 | -3.465029684 | 2.1001892 | 0.000143931 | 0.000775588 |
| SUCNR1 | -3.468389872 | 1.509039284 | 2.17E-05 | 0.000151239 |
| LOC119870470 | -3.468567466 | -0.241339863 | 3.15E-07 | 3.74E-06 |
| LOC608379_3 | -3.471067065 | 0.96536241 | 0.003097222 | 0.010693978 |
| LOC111094505 | -3.472622104 | -0.105652872 | 5.39E-07 | 5.97E-06 |
| LOC119872505 | -3.473890665 | -0.507165044 | 2.00E-05 | 0.000140731 |
| LOC111097806 | -3.481346998 | 4.008123146 | 2.42E-07 | 2.96E-06 |
| LOC119871559 | -3.481951465 | 4.139667 | 6.62E-06 | 5.30E-05 |
| LOC111095623 | -3.483679499 | -0.89835261 | 1.67E-06 | 1.62E-05 |
| STX11 | -3.484789317 | 5.900050981 | 9.32E-15 | 5.75E-13 |
| C4H5orf58 | -3.486784687 | 4.580692464 | 7.80E-08 | 1.08E-06 |
| LOC119874697 | -3.487951605 | 0.418768127 | 0.000412931 | 0.001918012 |
| LRATD2 | -3.488896275 | 5.200618909 | 1.30E-10 | 3.47E-09 |
| LOC612422 | -3.490062548 | 9.1255265 | 1.54E-06 | 1.50E-05 |

| | | | | |
|---|---|---|---|---|
| LOC119877466 | -3.490578434 | 3.614906414 | 4.04E-06 | 3.46E-05 |
| SYTL3 | -3.490847381 | 5.202028595 | 1.21E-08 | 2.03E-07 |
| LOC119872107 | -3.498364621 | -0.087848926 | 2.61E-05 | 0.000178225 |
| LOC111096673 | -3.502407774 | 3.112764077 | 6.34E-07 | 6.87E-06 |
| IGSF6 | -3.502800449 | 5.351920323 | 6.88E-12 | 2.47E-10 |
| SPTY2D1OS | -3.502813627 | 1.846173707 | 4.23E-09 | 7.93E-08 |
| LOC119867503 | -3.503793498 | 1.651032524 | 5.66E-06 | 4.62E-05 |
| GGT1 | -3.503972216 | 2.300459708 | 8.20E-06 | 6.40E-05 |
| LOC119870976 | -3.504036478 | 0.03345561 | 0.001576813 | 0.006015537 |
| MIR29B-2 | -3.505790591 | 1.370897539 | 9.00E-09 | 1.57E-07 |
| LOC119872802 | -3.512591494 | 0.535148005 | 6.82E-05 | 0.000409893 |
| LOC611538 | -3.545818568 | 4.343414625 | 1.29E-07 | 1.70E-06 |
| LOC119876970 | -3.546253529 | 3.788259558 | 9.76E-08 | 1.33E-06 |
| LOC119867601 | -3.546566467 | -0.77627169 | 0.000183797 | 0.000954802 |
| COTL1 | -3.549303448 | 8.980723335 | 8.81E-12 | 3.04E-10 |
| PPP1R3B | -3.555155104 | 6.619818357 | 1.51E-10 | 3.96E-09 |
| EPHA1 | -3.557174591 | 1.39716908 | 4.38E-06 | 3.71E-05 |
| LOC119872198 | -3.559112495 | 0.058099904 | 1.87E-05 | 0.000132597 |
| LOC119876480 | -3.573060755 | 0.723543395 | 0.000530491 | 0.002375481 |
| LOC119864636 | -3.579029152 | 0.863463557 | 9.80E-06 | 7.49E-05 |
| BIN2 | -3.591466582 | 7.471721351 | 6.39E-08 | 9.13E-07 |
| LOC119869121 | -3.593025541 | 1.163951632 | 0.005282479 | 0.016759942 |
| LOC102156642 | -3.595417842 | -0.207054671 | 3.62E-05 | 0.000237092 |
| CDH20 | -3.597754143 | -0.424792652 | 0.00031957 | 0.001537412 |
| PTAFR | -3.606443098 | 4.634038247 | 7.73E-12 | 2.72E-10 |
| MARCHF2 | -3.607756609 | 5.051037546 | 9.47E-13 | 4.07E-11 |
| LOC102151894 | -3.618396537 | 3.026227319 | 7.65E-11 | 2.14E-09 |
| LOC119876773 | -3.618632629 | 1.36005155 | 6.65E-09 | 1.19E-07 |
| CD24 | -3.625365805 | 6.471663066 | 9.57E-08 | 1.30E-06 |
| ALDH1A1 | -3.632527452 | 0.587664956 | 0.003531409 | 0.011943856 |
| LOC102152171 | -3.633053285 | 0.61611992 | 4.84E-06 | 4.03E-05 |
| DAPP1 | -3.63361522 | 4.823348208 | 9.84E-08 | 1.33E-06 |
| LOC102151345 | -3.634728115 | 1.348154719 | 1.71E-09 | 3.54E-08 |
| LOC111098723 | -3.637812283 | -0.192502528 | 3.65E-05 | 0.000238649 |
| RAB27A | -3.641272209 | 5.55781838 | 6.66E-08 | 9.47E-07 |
| COL6A5 | -3.641900561 | 5.130873471 | 0.003417669 | 0.011619379 |
| SNX20 | -3.643417427 | 4.105408216 | 4.27E-06 | 3.63E-05 |
| LOC119869744 | -3.647973043 | 0.802835494 | 4.18E-07 | 4.78E-06 |
| LOC485809 | -3.653592109 | -0.710571352 | 4.02E-05 | 0.000259301 |

| | | | | |
|---|---|---|---|---|
| TBXAS1 | -3.654663504 | 5.114477851 | 7.21E-12 | 2.57E-10 |
| LOC119871795 | -3.6585251 | 2.576108886 | 0.000233141 | 0.001170731 |
| CCR3 | -3.661877369 | -0.322124324 | 7.74E-06 | 6.09E-05 |
| AMD1 | -3.662010006 | 6.988347603 | 4.25E-10 | 1.01E-08 |
| LOC111095432 | -3.662615981 | -0.569832775 | 5.54E-05 | 0.000343168 |
| ITK | -3.665283533 | 5.866199348 | 0.000313584 | 0.001514266 |
| LOC111091184 | -3.66597718 | 2.862525521 | 0.00021908 | 0.001111452 |
| LOC119873878 | -3.671121987 | -0.057555895 | 1.19E-06 | 1.19E-05 |
| TBX21 | -3.673493495 | 3.955447616 | 1.49E-05 | 0.000108264 |
| AQP3 | -3.676380366 | 3.300076623 | 0.000532956 | 0.002384057 |
| CAMK4 | -3.677807578 | 5.099697843 | 3.58E-06 | 3.12E-05 |
| LOC111095976 | -3.680412424 | 0.625817511 | 0.00020495 | 0.001051085 |
| LOC111093000 | -3.68198112 | 0.419343379 | 2.97E-08 | 4.60E-07 |
| RBPMS2 | -3.686462662 | 4.618166313 | 8.42E-07 | 8.77E-06 |
| PRKCQ | -3.686471937 | 4.156577274 | 0.000240418 | 0.00120198 |
| LOC119870700 | -3.696638592 | 0.234165084 | 6.99E-05 | 0.000419795 |
| P2RY13 | -3.697688997 | 3.053920271 | 5.61E-07 | 6.19E-06 |
| ENO4 | -3.697694888 | 0.861859478 | 6.42E-06 | 5.16E-05 |
| FCGR1A | -3.699880178 | 4.732315176 | 1.66E-13 | 8.20E-12 |
| ANKRD34C | -3.70119089 | 0.045734659 | 1.20E-05 | 8.98E-05 |
| LOC119870241 | -3.709809866 | -0.379316966 | 1.22E-05 | 9.07E-05 |
| EPCAM | -3.712853257 | 0.157782554 | 0.000228431 | 0.001150009 |
| LOC111090345 | -3.714404779 | 1.462055969 | 6.95E-07 | 7.45E-06 |
| LOC102155837 | -3.714658212 | 1.591924594 | 9.90E-05 | 0.000567158 |
| IL31RA | -3.716430754 | 4.540142385 | 2.72E-05 | 0.000184528 |
| LOC610614 | -3.728943997 | 2.303752055 | 0.000162594 | 0.000860269 |
| LOC119876664 | -3.733160408 | -1.234640416 | 9.94E-06 | 7.58E-05 |
| LOC119871221 | -3.740963924 | 1.873552672 | 5.68E-07 | 6.26E-06 |
| SLAIN1 | -3.750767732 | 3.442758322 | 1.21E-08 | 2.03E-07 |
| LOC119878923 | -3.757264207 | 1.45929593 | 0.001146494 | 0.004605739 |
| ITGAD | -3.76141075 | 0.397400291 | 0.000131583 | 0.000719972 |
| LOC119874343 | -3.762764444 | 1.009926196 | 8.67E-06 | 6.72E-05 |
| LOC102154687 | -3.768963685 | -1.098401515 | 2.67E-06 | 2.42E-05 |
| LOC119873305 | -3.779122754 | 1.456373268 | 2.65E-07 | 3.19E-06 |
| NCF2 | -3.781222185 | 4.965719239 | 6.06E-11 | 1.72E-09 |
| TSHB | -3.784231634 | -0.668229568 | 0.000111847 | 0.00063028 |
| LOC119876799 | -3.787853472 | 0.205319517 | 0.001058275 | 0.004297544 |
| LOC119870764 | -3.792988478 | 2.792044717 | 1.87E-06 | 1.78E-05 |
| LOC119866669 | -3.795642208 | -0.253225227 | 0.00015301 | 0.000815588 |

| | | | | |
|---|---|---|---|---|
| LOC106558644 | -3.801409651 | 0.056659557 | 0.000420201 | 0.001946777 |
| MYT1 | -3.802603676 | -0.00201668 | 8.09E-06 | 6.33E-05 |
| C6H16orf54 | -3.805992102 | 5.380218368 | 9.06E-12 | 3.12E-10 |
| PLA2G7 | -3.806029761 | 6.552309533 | 1.71E-07 | 2.17E-06 |
| MSRB1 | -3.816012043 | 5.417955999 | 1.29E-14 | 7.76E-13 |
| LOC119868093 | -3.816021332 | -0.277981616 | 3.76E-08 | 5.67E-07 |
| DUSP2 | -3.82013613 | 4.843642859 | 1.57E-05 | 0.000113883 |
| SMIM35 | -3.834036383 | 3.325716693 | 3.58E-10 | 8.67E-09 |
| CX3CR1 | -3.835003942 | 3.365627873 | 4.76E-06 | 3.99E-05 |
| LOC119871235 | -3.835724644 | 1.152009192 | 7.76E-10 | 1.75E-08 |
| PTGDR2 | -3.836058899 | 1.264002623 | 3.93E-08 | 5.90E-07 |
| LOC100686271 | -3.836145395 | 3.951034809 | 3.77E-10 | 9.13E-09 |
| LOC119867311 | -3.838965275 | 0.286348208 | 1.62E-05 | 0.00011723 |
| SLC7A11 | -3.842251902 | 3.27314209 | 5.75E-09 | 1.05E-07 |
| LOC119872032 | -3.842896388 | 0.267930672 | 2.93E-05 | 0.000196849 |
| LOC119875991 | -3.846679867 | 1.776189692 | 8.27E-06 | 6.45E-05 |
| LOC119875962 | -3.84759858 | 2.888310797 | 8.78E-12 | 3.04E-10 |
| LOC119869848 | -3.848482226 | 0.147985295 | 7.69E-06 | 6.06E-05 |
| LOC102155550 | -3.849822779 | 0.077000767 | 4.03E-07 | 4.64E-06 |
| MUC4 | -3.857450917 | 2.146964329 | 3.48E-05 | 0.000228938 |
| LOC106558563 | -3.8594113 | 1.291693653 | 2.29E-08 | 3.62E-07 |
| AASS | -3.86130527 | 2.700705001 | 1.63E-08 | 2.67E-07 |
| PILRA | -3.876731332 | 4.465136221 | 7.01E-08 | 9.89E-07 |
| LOC119868972 | -3.878774814 | 3.663815089 | 1.73E-08 | 2.82E-07 |
| CYTIP | -3.880355285 | 6.657702649 | 1.35E-07 | 1.76E-06 |
| MCTP2 | -3.881970946 | 5.300945468 | 1.32E-06 | 1.31E-05 |
| LOC111097254 | -3.885064864 | 1.790708462 | 0.000438541 | 0.002017942 |
| SELP | -3.893001102 | 5.118021082 | 4.21E-06 | 3.59E-05 |
| GATA3 | -3.893598438 | 4.691805114 | 1.34E-07 | 1.76E-06 |
| LOC106558673 | -3.905971296 | 0.833836354 | 4.18E-09 | 7.85E-08 |
| LOC106559612 | -3.909421656 | 3.184899706 | 2.95E-08 | 4.57E-07 |
| ARHGAP15 | -3.909671799 | 6.0934189 | 7.08E-08 | 9.96E-07 |
| EREG | -3.914811059 | 1.826305612 | 0.000445226 | 0.002041335 |
| CDKN2D | -3.91765193 | 4.229528995 | 4.49E-14 | 2.47E-12 |
| LOC102156672 | -3.917889103 | 0.075296024 | 5.91E-06 | 4.81E-05 |
| RIPOR2 | -3.918613163 | 8.193810294 | 4.46E-06 | 3.78E-05 |
| GZMB | -3.920618345 | 3.41954012 | 9.71E-06 | 7.43E-05 |
| TSPAN33 | -3.921877966 | 1.196560124 | 3.92E-05 | 0.000254058 |
| NLRP3 | -3.923431686 | 4.990903361 | 1.29E-11 | 4.26E-10 |

| | | | | |
|---|---|---|---|---|
| LOC111097157 | -3.923880712 | 1.072893565 | 0.000110031 | 0.000621767 |
| SLC9C2 | -3.929153757 | 0.464728594 | 5.05E-07 | 5.63E-06 |
| MPL | -3.937264792 | 1.22925263 | 7.93E-08 | 1.10E-06 |
| TMEM121B | -3.938998219 | -0.585333899 | 4.24E-07 | 4.84E-06 |
| GCNT4 | -3.939240375 | 2.625243241 | 1.74E-07 | 2.20E-06 |
| CLEC1B | -3.942314554 | 4.184762479 | 6.64E-06 | 5.31E-05 |
| BEST4 | -3.942318084 | 2.419213607 | 7.11E-10 | 1.62E-08 |
| LOC102152359 | -3.944901743 | 0.914972948 | 1.10E-05 | 8.28E-05 |
| LOC100855676 | -3.950929517 | 1.290903005 | 0.002967132 | 0.010317126 |
| CCDC83 | -3.958417286 | 1.759349138 | 2.60E-08 | 4.07E-07 |
| BIK | -3.966110926 | 0.738847829 | 6.65E-07 | 7.15E-06 |
| LOC102154496 | -3.969947269 | 1.882006205 | 0.000216867 | 0.001102031 |
| MYBPC1 | -3.974860874 | -0.446207166 | 0.00398383 | 0.013246633 |
| KLHL25 | -3.975812677 | 5.224107465 | 3.57E-18 | 4.23E-16 |
| LOC119873443 | -3.976062053 | -0.204722427 | 1.13E-06 | 1.14E-05 |
| LOC119869103 | -3.978248301 | 3.376442877 | 0.002901123 | 0.010117166 |
| LOC111090795 | -3.980443595 | 1.270913479 | 1.58E-05 | 0.000114048 |
| G0S2 | -3.982254404 | 2.87532596 | 0.000584195 | 0.002577368 |
| LOC119879138 | -3.982387183 | 1.367059541 | 7.90E-07 | 8.31E-06 |
| LOC100688166 | -3.985134295 | 0.64962955 | 1.24E-06 | 1.24E-05 |
| LOC100685620 | -3.988807076 | 1.286168761 | 7.89E-10 | 1.78E-08 |
| IL1RN | -3.996567146 | 5.226559808 | 4.93E-15 | 3.26E-13 |
| LOC111094076 | -4.004372245 | 0.36649839 | 7.02E-07 | 7.51E-06 |
| GPR55 | -4.004669978 | 2.38953812 | 2.23E-05 | 0.000155415 |
| LOC119869317 | -4.042233546 | 5.909503891 | 1.23E-08 | 2.06E-07 |
| LOC102157022 | -4.050850646 | 1.777489567 | 2.91E-05 | 0.000196049 |
| C5AR1 | -4.057084948 | 5.686580618 | 3.30E-12 | 1.28E-10 |
| GRAP2 | -4.057362318 | 5.02971326 | 3.47E-08 | 5.29E-07 |
| CEBPE | -4.069695785 | 0.186863241 | 4.07E-07 | 4.67E-06 |
| LOC111094344 | -4.070637153 | 1.724439925 | 1.99E-08 | 3.19E-07 |
| DAB1 | -4.091977503 | 2.104683413 | 8.68E-06 | 6.72E-05 |
| LOC119876581 | -4.095746485 | 2.05791579 | 5.98E-05 | 0.000366814 |
| KLRB1 | -4.09966061 | 1.046591246 | 5.83E-05 | 0.00035888 |
| CSF2RA | -4.107451456 | 5.040443547 | 4.50E-10 | 1.07E-08 |
| LOC119864489 | -4.130722846 | 4.187801422 | 0.00013182 | 0.000721086 |
| LOC111097892 | -4.156086523 | 4.773728039 | 4.83E-10 | 1.13E-08 |
| LOC119870724 | -4.158684506 | 2.670973123 | 6.47E-10 | 1.49E-08 |
| LOC119877642 | -4.161125259 | 0.327972929 | 1.19E-06 | 1.20E-05 |
| LOC111094990 | -4.161279994 | 1.01287963 | 0.000187193 | 0.000969153 |

| | | | | |
|---|---|---|---|---|
| LOC484309 | -4.164436339 | 5.713140818 | 3.34E-12 | 1.29E-10 |
| ACOT7 | -4.17773104 | 4.038192163 | 5.81E-07 | 6.38E-06 |
| HAL | -4.180362474 | -0.311893287 | 0.000114225 | 0.000640849 |
| GP9 | -4.181192269 | 2.985527105 | 3.29E-13 | 1.54E-11 |
| LOC102151704 | -4.182013224 | 2.539315729 | 1.11E-07 | 1.49E-06 |
| FGR | -4.187519374 | 6.624904415 | 2.17E-10 | 5.52E-09 |
| NLRP12 | -4.190642195 | 5.370100709 | 1.33E-12 | 5.52E-11 |
| GUCY2C | -4.196719063 | 1.661747279 | 2.17E-06 | 2.02E-05 |
| ADGRE1 | -4.206995885 | 5.602890734 | 2.82E-08 | 4.39E-07 |
| LOC111095428 | -4.215024455 | 3.473649282 | 1.94E-16 | 1.66E-14 |
| LOC111091185 | -4.218311481 | 0.820267109 | 4.11E-05 | 0.000264364 |
| LOC119869334 | -4.231420013 | -0.645033273 | 0.002269399 | 0.00820018 |
| MOGAT1 | -4.238429304 | -0.186057578 | 2.22E-06 | 2.06E-05 |
| LOC119877825 | -4.250633352 | 0.035066722 | 8.70E-05 | 0.000507481 |
| XK | -4.267460945 | 3.852707036 | 4.41E-09 | 8.22E-08 |
| MS4A2 | -4.28805454 | 1.556397181 | 0.001321124 | 0.005185606 |
| RGS2 | -4.292336857 | 7.015349936 | 2.26E-12 | 8.99E-11 |
| OXER1 | -4.307244812 | 1.979983397 | 9.37E-10 | 2.07E-08 |
| IL27 | -4.325278429 | 2.116570489 | 2.56E-09 | 5.06E-08 |
| CHRNA2 | -4.338094754 | 1.764889497 | 4.78E-08 | 7.05E-07 |
| LRG1 | -4.339158571 | 2.872197334 | 1.80E-07 | 2.27E-06 |
| IL1B | -4.343744718 | 1.804297967 | 1.45E-06 | 1.42E-05 |
| LAMB3 | -4.346305845 | 3.229720117 | 8.98E-10 | 2.00E-08 |
| MIR103-2 | -4.347759856 | -0.484364589 | 1.90E-06 | 1.80E-05 |
| LOC119866716 | -4.349917347 | -0.717985533 | 3.96E-06 | 3.40E-05 |
| LOC119865429 | -4.350420133 | -0.833956759 | 2.97E-05 | 0.000199306 |
| HAMP | -4.357968143 | -0.270319564 | 6.76E-05 | 0.000407351 |
| ALOX5 | -4.360830293 | 5.480442691 | 6.60E-14 | 3.48E-12 |
| LOC119871838 | -4.362518337 | 3.487500485 | 1.36E-09 | 2.89E-08 |
| MEGF9 | -4.367076592 | 8.163503735 | 6.57E-19 | 8.85E-17 |
| SPINT1 | -4.367205192 | 4.834308551 | 3.12E-15 | 2.12E-13 |
| LDHC | -4.372315105 | -1.568190131 | 2.69E-07 | 3.23E-06 |
| LOC119874694 | -4.372908253 | 1.004914277 | 5.31E-05 | 0.000331079 |
| LOC106558720 | -4.394106888 | 1.987501686 | 2.18E-11 | 6.86E-10 |
| CCNJL | -4.394279015 | 2.829350176 | 6.27E-09 | 1.13E-07 |
| ARMH2 | -4.394469966 | -0.045109095 | 5.30E-06 | 4.36E-05 |
| LOC119867255 | -4.398556287 | -1.230254581 | 2.07E-05 | 0.000145005 |
| LOC119865687 | -4.401121273 | 4.439290142 | 1.22E-17 | 1.29E-15 |
| LOC111093977 | -4.402423613 | 1.693042518 | 5.48E-07 | 6.07E-06 |

| | | | | |
|---|---|---|---|---|
| NLRP1 | -4.403587332 | 6.463884887 | 3.84E-09 | 7.27E-08 |
| BEST1 | -4.407458628 | 0.835432901 | 3.96E-05 | 0.00025592 |
| PADI3 | -4.408644423 | 1.301993868 | 4.84E-05 | 0.000305622 |
| FAM71A | -4.410531854 | -0.863390971 | 0.000148621 | 0.000795913 |
| MUC19 | -4.416765168 | -0.269817678 | 6.86E-08 | 9.73E-07 |
| LOC111094690 | -4.417711363 | 1.192852211 | 4.42E-09 | 8.23E-08 |
| LOC111093949 | -4.423400694 | -0.426485821 | 2.48E-06 | 2.27E-05 |
| LOC119866191 | -4.428656675 | -0.768339055 | 1.62E-06 | 1.57E-05 |
| SORL1 | -4.442818123 | 8.577205052 | 7.19E-10 | 1.63E-08 |
| CD1E | -4.444500199 | 0.984810478 | 6.20E-06 | 5.00E-05 |
| CTLA4 | -4.444915985 | 3.211782432 | 1.97E-05 | 0.000139178 |
| SH3BGRL2 | -4.447668802 | 6.250054419 | 6.35E-11 | 1.80E-09 |
| LOC119866868 | -4.491878836 | 1.60518678 | 2.90E-06 | 2.61E-05 |
| LOC119866406 | -4.492331702 | 0.006467859 | 5.66E-08 | 8.17E-07 |
| LOC119869514 | -4.499556698 | -0.87275407 | 8.52E-07 | 8.86E-06 |
| LOC111091444 | -4.500085725 | 0.80020523 | 8.00E-09 | 1.42E-07 |
| CD1A8 | -4.501383189 | 2.60161782 | 3.52E-07 | 4.11E-06 |
| ISG20 | -4.524922303 | 4.381802515 | 2.82E-11 | 8.60E-10 |
| LOC111093061 | -4.536266419 | -0.35490175 | 1.06E-09 | 2.33E-08 |
| DGAT2 | -4.539511142 | 5.524102237 | 2.69E-13 | 1.28E-11 |
| LOC119864387 | -4.540275512 | -0.31624417 | 0.000151062 | 0.000806396 |
| LOC111093040 | -4.550003135 | -0.390617128 | 6.94E-07 | 7.44E-06 |
| GALNT17 | -4.550174833 | 4.110537132 | 2.50E-06 | 2.29E-05 |
| LOC119870768 | -4.553579147 | 0.705271191 | 2.81E-11 | 8.60E-10 |
| S1PR5 | -4.574915842 | 0.54576619 | 1.45E-09 | 3.05E-08 |
| CAECAM1 | -4.576683503 | 5.171896322 | 9.04E-15 | 5.61E-13 |
| GSDMA | -4.580079307 | 1.368740526 | 9.62E-09 | 1.66E-07 |
| LYZ | -4.583348085 | 8.322634979 | 2.56E-08 | 4.01E-07 |
| MALRD1 | -4.598219718 | 0.858634866 | 1.18E-05 | 8.85E-05 |
| LOC119870904 | -4.602325704 | 0.382404426 | 9.82E-06 | 7.50E-05 |
| LOC119869775 | -4.603338718 | -1.332785035 | 4.10E-06 | 3.50E-05 |
| LOC119868973 | -4.609546864 | 0.703772243 | 3.60E-09 | 6.87E-08 |
| LOC111091256 | -4.617730377 | 0.840882436 | 7.28E-08 | 1.02E-06 |
| LOC119874184 | -4.619893028 | -0.560976522 | 3.61E-07 | 4.21E-06 |
| LOC608848 | -4.620733267 | 4.496887353 | 9.56E-11 | 2.61E-09 |
| SELL | -4.651965225 | 6.802219924 | 6.34E-05 | 0.000385215 |
| LOC119863983 | -4.662335072 | 3.537141746 | 3.23E-10 | 7.89E-09 |
| ITGAM | -4.676775855 | 6.52390096 | 9.79E-11 | 2.66E-09 |
| GP1BA | -4.68675448 | 4.739184724 | 9.98E-11 | 2.71E-09 |

| | | | | |
|---|---|---|---|---|
| SLCO4C1 | -4.689296464 | 2.17282654 | 3.36E-06 | 2.96E-05 |
| LOC111092010 | -4.691636741 | 1.071418038 | 7.44E-06 | 5.88E-05 |
| CD226 | -4.699228735 | 5.725355725 | 1.88E-08 | 3.03E-07 |
| CLEC5A | -4.700784416 | 3.59511574 | 1.22E-11 | 4.07E-10 |
| LOC106558211 | -4.701627112 | 1.65926964 | 1.17E-08 | 1.98E-07 |
| AOAH | -4.709551194 | 6.11403406 | 5.14E-11 | 1.49E-09 |
| LOC100686711 | -4.710171337 | 2.107778913 | 1.25E-09 | 2.68E-08 |
| SRGN | -4.712858653 | 7.489379969 | 3.01E-12 | 1.18E-10 |
| IGF2BP3 | -4.729987131 | 4.890064924 | 6.60E-09 | 1.19E-07 |
| ATP13A4 | -4.734781509 | 0.66696822 | 1.59E-08 | 2.61E-07 |
| C9H17orf64 | -4.740243203 | 3.339825908 | 1.49E-15 | 1.07E-13 |
| LOC102154832 | -4.751703585 | 0.176369926 | 5.30E-07 | 5.89E-06 |
| NRGN | -4.7521237 | 5.284939852 | 4.55E-10 | 1.08E-08 |
| LOC489231 | -4.754162831 | -0.106225347 | 3.66E-06 | 3.17E-05 |
| LOC111094796 | -4.779601109 | -1.161174623 | 1.17E-05 | 8.76E-05 |
| SLC28A3 | -4.785963462 | 2.032711761 | 2.63E-10 | 6.51E-09 |
| OR9A4 | -4.819966181 | 0.445481244 | 3.59E-08 | 5.45E-07 |
| LOC102156518 | -4.82824842 | 0.467859797 | 4.03E-06 | 3.45E-05 |
| FOXI1 | -4.849575109 | 1.764648411 | 6.36E-07 | 6.89E-06 |
| IL1RL2 | -4.854533119 | 0.907767139 | 1.63E-07 | 2.07E-06 |
| PON3 | -4.870157015 | 2.697817098 | 3.32E-10 | 8.08E-09 |
| PMAIP1 | -4.889707006 | 3.354877854 | 1.05E-13 | 5.32E-12 |
| LOC102151859 | -4.89577625 | 1.642233357 | 1.06E-08 | 1.81E-07 |
| LOC119871796 | -4.899082803 | -0.652749151 | 7.85E-07 | 8.27E-06 |
| IL7R | -4.899963226 | 6.603561557 | 6.90E-08 | 9.77E-07 |
| LOC111097357 | -4.902029828 | -0.093751363 | 6.31E-08 | 9.03E-07 |
| CD300E | -4.921435328 | 2.084165172 | 5.80E-09 | 1.06E-07 |
| CIDEA | -4.924917492 | 3.65125191 | 4.65E-06 | 3.91E-05 |
| LOC111095804 | -4.925175726 | 0.785913282 | 1.82E-06 | 1.74E-05 |
| LOC608975 | -4.946111629 | 5.07865619 | 2.49E-13 | 1.19E-11 |
| LOC111092148 | -4.949234388 | 1.825764642 | 7.52E-08 | 1.05E-06 |
| CEACAM28 | -4.958159603 | 5.929987955 | 7.12E-20 | 1.19E-17 |
| LOC102153134 | -4.987085466 | 0.216500452 | 1.37E-09 | 2.92E-08 |
| RAP1GAP2 | -4.990889937 | 4.851544026 | 7.42E-14 | 3.86E-12 |
| ADGRE2 | -5.005580659 | 4.466973195 | 1.02E-13 | 5.20E-12 |
| RGS18 | -5.009142677 | 7.113849976 | 1.15E-10 | 3.08E-09 |
| LOC102154873 | -5.021655219 | 2.159543716 | 1.18E-08 | 2.00E-07 |
| DGKG | -5.032567608 | 5.358403149 | 6.95E-12 | 2.50E-10 |
| MIR371 | -5.036127991 | 0.509805481 | 5.76E-08 | 8.32E-07 |

| | | | | |
|---|---|---|---|---|
| LOC119866900 | -5.050141288 | 1.488056544 | 1.21E-09 | 2.61E-08 |
| LOC106557449 | -5.076409035 | 2.823482835 | 2.12E-06 | 1.98E-05 |
| LOC102156464 | -5.083863711 | 0.798229063 | 6.01E-08 | 8.64E-07 |
| SERPINB10 | -5.097638147 | 5.340221382 | 4.74E-10 | 1.12E-08 |
| LOC106559783 | -5.114397661 | 2.628445885 | 1.38E-10 | 3.66E-09 |
| LOC119865634 | -5.130790761 | -0.15050097 | 3.77E-09 | 7.16E-08 |
| LOC119871464 | -5.145543973 | 0.218342813 | 2.05E-11 | 6.50E-10 |
| LOC102152290 | -5.150076953 | 0.716355874 | 4.19E-09 | 7.86E-08 |
| LOC102153812 | -5.152954885 | 0.99668215 | 1.62E-05 | 0.000116651 |
| PEAK3 | -5.15586721 | 3.007924055 | 2.67E-14 | 1.53E-12 |
| PADI2 | -5.174233062 | 6.769355904 | 4.10E-18 | 4.71E-16 |
| LOC119864904 | -5.178512629 | 1.516376715 | 0.000140265 | 0.00075923 |
| UNC45B | -5.180168855 | -0.633187155 | 2.54E-07 | 3.09E-06 |
| LOC119866113 | -5.184473062 | 2.206347795 | 2.70E-05 | 0.00018314 |
| LOC119877470 | -5.216411182 | 0.697793398 | 3.08E-08 | 4.74E-07 |
| TREM1 | -5.234112393 | 4.006891111 | 1.58E-11 | 5.14E-10 |
| CD1A6 | -5.242421824 | 1.407541507 | 1.49E-07 | 1.93E-06 |
| IL18RAP | -5.253019672 | 4.622278563 | 9.40E-14 | 4.83E-12 |
| LOC484314 | -5.26131732 | 4.126675982 | 3.80E-12 | 1.44E-10 |
| IL18R1 | -5.272820797 | 3.835144153 | 2.13E-10 | 5.44E-09 |
| CXCL8 | -5.280697018 | 7.247662048 | 6.35E-06 | 5.11E-05 |
| MPIG6B | -5.285562581 | 7.223679092 | 2.70E-10 | 6.67E-09 |
| HPSE | -5.306548866 | 6.830536866 | 5.14E-10 | 1.20E-08 |
| TRAT1 | -5.320725454 | 4.400529425 | 6.14E-06 | 4.96E-05 |
| LOC611642 | -5.335767646 | 2.193601 | 3.21E-10 | 7.86E-09 |
| DCSTAMP | -5.364887634 | 4.377584219 | 8.03E-05 | 0.000473654 |
| SLFN14 | -5.390496636 | 2.953779124 | 1.74E-14 | 1.02E-12 |
| KCNJ15 | -5.420654812 | 5.747968806 | 3.34E-10 | 8.12E-09 |
| LOC111091743 | -5.43398659 | 0.163233499 | 1.09E-07 | 1.46E-06 |
| LOC102151528 | -5.443094533 | 3.463387004 | 1.37E-12 | 5.67E-11 |
| EMB | -5.46647096 | 7.819095052 | 2.63E-13 | 1.25E-11 |
| LOC119864138 | -5.467799285 | 0.662363236 | 1.12E-08 | 1.90E-07 |
| FPR2 | -5.490129056 | 2.170947077 | 6.95E-10 | 1.59E-08 |
| LOC102154760 | -5.514068415 | 8.890263518 | 2.81E-05 | 0.000189877 |
| LOC119865562 | -5.547141128 | 0.218704457 | 2.37E-07 | 2.90E-06 |
| LOC119872023 | -5.595537631 | 0.407956607 | 1.04E-10 | 2.81E-09 |
| BCL2L15 | -5.616480931 | 2.309688342 | 3.80E-11 | 1.13E-09 |
| GFI1B | -5.630096783 | 3.554980227 | 4.89E-27 | 2.94E-24 |
| LOC119871267 | -5.701279397 | 0.169789488 | 3.79E-11 | 1.13E-09 |

| | | | | |
|---|---|---|---|---|
| GPR162 | -5.710221079 | 5.353703361 | 2.41E-14 | 1.39E-12 |
| ACOXL | -5.784134977 | 3.292339433 | 9.85E-15 | 6.07E-13 |
| GATM | -5.822065339 | 3.76214981 | 4.71E-09 | 8.72E-08 |
| SH2D1B | -5.862002959 | 3.891835514 | 4.73E-22 | 1.11E-19 |
| LOC111093462 | -5.872908731 | 1.200497516 | 9.16E-09 | 1.59E-07 |
| MEFV | -5.874320599 | 3.652840175 | 2.58E-13 | 1.23E-11 |
| LOC119864622 | -5.882374691 | 0.580521049 | 8.48E-11 | 2.34E-09 |
| SLC26A8 | -5.907843279 | 1.044089354 | 1.45E-09 | 3.06E-08 |
| CD4 | -5.926688373 | 6.933521918 | 8.90E-10 | 1.98E-08 |
| FFAR2 | -5.970746634 | 6.270662209 | 1.63E-10 | 4.24E-09 |
| LOC119879117 | -5.980982953 | 3.30799019 | 3.50E-09 | 6.70E-08 |
| TREM2 | -5.990964582 | 4.510034078 | 1.13E-09 | 2.47E-08 |
| LOC102157398 | -6.01231799 | -0.19065043 | 9.22E-11 | 2.52E-09 |
| PLBD1 | -6.026662994 | 7.086488501 | 5.74E-19 | 7.77E-17 |
| LOC484865 | -6.028230637 | 1.540330629 | 7.75E-06 | 6.09E-05 |
| LOC119877019 | -6.065370121 | 1.261024889 | 1.40E-09 | 2.96E-08 |
| MCEMP1 | -6.120202528 | 3.417715961 | 1.20E-09 | 2.60E-08 |
| LOC111092740 | -6.163063057 | 6.062978135 | 3.16E-12 | 1.24E-10 |
| LOC119865122 | -6.240452463 | -0.174006016 | 1.93E-11 | 6.16E-10 |
| APOA2 | -6.28590692 | -0.165648596 | 1.29E-12 | 5.38E-11 |
| LOC111095218 | -6.302588865 | 0.999493728 | 3.00E-11 | 9.09E-10 |
| MGAM2 | -6.33379461 | 1.153942201 | 6.30E-12 | 2.30E-10 |
| IL22RA2 | -6.356444641 | 2.062758649 | 2.71E-18 | 3.31E-16 |
| ITGA2B | -6.396073107 | 6.155166323 | 2.36E-21 | 5.23E-19 |
| LOC111097149 | -6.416123821 | 0.910776343 | 8.07E-12 | 2.83E-10 |
| LOC119869313 | -6.572600231 | 2.555160268 | 1.57E-15 | 1.12E-13 |
| ADGRE3 | -6.582074532 | 5.540661459 | 3.67E-16 | 2.94E-14 |
| LOC102156687 | -6.589810835 | 0.569004977 | 3.39E-11 | 1.02E-09 |
| LOC111097288 | -6.590240315 | 0.564635078 | 8.08E-12 | 2.83E-10 |
| LOC119870554 | -6.598261772 | -0.753152819 | 5.97E-11 | 1.70E-09 |
| SERPINA1 | -6.720114584 | 7.873584675 | 1.40E-13 | 6.99E-12 |
| IL5RA | -6.768272622 | 1.861723155 | 4.86E-10 | 1.14E-08 |
| LOC106559021 | -6.789193944 | 0.328432216 | 3.61E-14 | 2.01E-12 |
| S100A8 | -6.809096194 | 6.362815076 | 2.29E-18 | 2.84E-16 |
| AQP9 | -6.880534051 | 5.901089987 | 1.09E-13 | 5.55E-12 |
| PROK2 | -6.930129762 | 3.398478982 | 3.16E-05 | 0.000210477 |
| LOC111095315 | -6.968012103 | -0.147314801 | 5.59E-12 | 2.06E-10 |
| LOC119876409 | -7.043712451 | 0.971780381 | 5.71E-14 | 3.09E-12 |
| MIR147 | -7.141009645 | 2.610618891 | 2.83E-14 | 1.62E-12 |

| | | | | |
|---|---|---|---|---|
| LOC111097386 | -7.149503082 | 1.908280043 | 3.50E-20 | 6.32E-18 |
| LOC119867504 | -7.150945445 | 1.503245723 | 2.08E-17 | 2.04E-15 |
| MGAM | -7.237411738 | 4.283027956 | 2.07E-27 | 1.40E-24 |
| LOC102155995 | -7.331066772 | 1.859464477 | 4.51E-19 | 6.26E-17 |
| LOC119870738 | -7.337689851 | -0.427110687 | 1.79E-10 | 4.64E-09 |
| LOC119866593 | -7.364437589 | -0.961294348 | 8.48E-11 | 2.34E-09 |
| CTSE | -7.408052192 | 3.959448175 | 1.99E-12 | 7.99E-11 |
| LOC119866531 | -7.418642304 | 1.026806179 | 1.35E-10 | 3.59E-09 |
| LOC102155370 | -7.556618245 | 0.479017193 | 1.87E-12 | 7.52E-11 |
| LOC119867645 | -7.638342654 | 2.828330765 | 7.09E-13 | 3.13E-11 |
| LOC119877471 | -8.090914485 | 0.835516789 | 2.28E-17 | 2.22E-15 |
| VSTM1 | -8.122049935 | 2.761143787 | 1.30E-17 | 1.36E-15 |
| LTF | -8.255109606 | 5.301820556 | 1.07E-29 | 1.15E-26 |
| LOC111093315 | -8.289596129 | 1.558970458 | 1.94E-18 | 2.42E-16 |
| LOC119879116 | -8.295532911 | -0.1473931 | 1.52E-12 | 6.23E-11 |
| LOC102151990 | -8.329305289 | 2.158412215 | 3.69E-25 | 1.55E-22 |
| LOC483397 | -8.334570658 | 3.322996937 | 2.61E-18 | 3.21E-16 |
| CXCR1 | -8.345000668 | 5.22799142 | 7.64E-15 | 4.82E-13 |
| NFE2 | -8.71566829 | 5.02325963 | 7.97E-41 | 4.32E-37 |
| LY6G6F | -8.989024026 | 3.167817676 | 2.52E-20 | 4.75E-18 |
| LOC119863926 | -9.057091986 | 7.300484296 | 3.60E-18 | 4.24E-16 |
| TUBB1 | -9.15851194 | 6.981286637 | 3.55E-33 | 6.99E-30 |
| MIR223 | -9.213790717 | 2.156112622 | 1.38E-24 | 4.75E-22 |
| LOC119879118 | -9.227262804 | 2.491810007 | 1.77E-11 | 5.69E-10 |
| LOC106557989 | -9.525757512 | 3.287136857 | 3.96E-35 | 1.07E-31 |
| MMP8 | -9.538981817 | 3.88841919 | 9.74E-18 | 1.06E-15 |
| CSF3R | -9.721082156 | 8.403576134 | 1.14E-25 | 5.36E-23 |
| TRIM58 | -9.781774324 | 2.426300676 | 9.15E-16 | 6.96E-14 |
| OLIG1 | -9.803764323 | 0.428885541 | 9.16E-14 | 4.72E-12 |
| LOC111098957 | -9.860849148 | 6.601036302 | 7.03E-39 | 3.05E-35 |
| EGF | -9.946645592 | 4.010646561 | 1.15E-15 | 8.43E-14 |
| PADI4 | -10.01337908 | 6.057055077 | 1.38E-26 | 7.50E-24 |
| S100A12 | -10.01887917 | 6.891850487 | 1.10E-47 | 2.38E-43 |
| S100A9 | -10.11098066 | 6.986637976 | 3.12E-27 | 1.93E-24 |
| LOC119872704 | -10.36626032 | 0.778209138 | 1.01E-17 | 1.09E-15 |
| CXCR2 | -10.38208392 | 4.767173064 | 3.08E-41 | 2.23E-37 |
| PPBP | -10.41879667 | 7.769194014 | 1.06E-11 | 3.60E-10 |
| LOC102156485 | -10.46360167 | 2.474963339 | 9.19E-21 | 1.84E-18 |
| LOC486765 | -10.47190038 | 2.46447487 | 1.34E-18 | 1.71E-16 |

| | | | | |
|---|---|---|---|---|
| GP6 | -12.9349987 | 3.247529706 | 3.62E-22 | 8.61E-20 |
